# Genetic diversity of Collaborative Cross mice enables the establishment of a novel *Chlamydia muridarum* female genital tract infection model

**DOI:** 10.64898/2025.12.22.695995

**Authors:** Toni Darville, Jenna Girardi, Taylor Poston, Camille Campbell, Xuejun Sun, Yajing Hao, Yu Zhang, Chuwen Liu, Bryce Duncan, Fei Zou, Catherine M. O’Connell, Xiaojing Zheng

**Author notes:** **Correspondence:** Xiaojing Zheng, PhD, University of North Carolina at Chapel Hill, 8340B MBRB, CB# 7509, 111 Mason Farm Road, Chapel Hill, NC 27599-7509, Alternate corresponding author: Catherine O’Connell, PhD, University of North Carolina at Chapel Hill, 8340B MBRB, CB# 7509, 111 Mason Farm Road, Chapel Hill, NC 27599-7509, Fei Zou, PhD, University of North Carolina at Chapel Hill, McGavran Greenberg Hall, CB #7420, Chapel Hill, NC 27599. shared first authorship (Toni Darville, Jenna Girardi). shared corresponding authorship (Fei Zou, Catherine M. O’Connell, Xiaojing Zheng).

## Abstract

Chlamydial infection in women displays wide variation in bacterial burden, persistence, and risk of upper genital tract pathology, yet the host genetic factors underlying this heterogeneity remain poorly defined. We evaluated genital tract infection with *Chlamydia muridarum* across 20 Collaborative Cross (CC) strains, a recombinant inbred mouse panel that captures broad genetic diversity with high within-strain reproducibility. CC strains exhibited striking differences in early bacterial burden, time to clearance, and oviduct pathology, including prolonged low-inflammatory infections and burden–pathology discordance that mirror key features of human disease. Heritability analyses demonstrated that host genetics accounted for most of the variation observed in Day 7 bacterial burden and pathological outcomes. Genome-wide scans identified suggestive quantitative trait loci associated with both traits. Genes within the burden-associated locus converged on host pathways implicated in chlamydial intracellular growth, including membrane dynamics and lipid metabolism, ubiquitin signaling, host cell survival mechanisms, and immune regulatory signaling. In contrast, genes within the pathology-associated locus were enriched for pathways regulating inflammatory cell recruitment, inflammasome activation, and tissue remodeling, processes central to genital tract damage following infection. Cervical transcriptional profiling further revealed strain-dependent innate and adaptive immune programs associated with bacterial burden and disease phenotype. Together, these findings establish the CC as a powerful platform for dissecting the genetic architecture of chlamydial immunopathogenesis, and for improving preclinical evaluation of vaccines and therapeutics.

## INTRODUCTION

*Chlamydia trachomatis* (CT) is the most common sexually transmitted bacterial pathogen worldwide. More than 70% of infections in women and 50% in men are asymptomatic (1), contributing to underdiagnosis, prolonged infection, and sustained transmission. In women, untreated infection can ascend to the upper genital tract and, in ∼10% of cases, cause acute pelvic inflammatory disease (PID) characterized by vaginal discharge, lower abdominal or pelvic pain, dyspareunia, and cervical or adnexal tenderness on examination (2). However, most women have no overt symptoms despite ongoing infection and upper tract inflammation, a condition termed subclinical or “silent” PID (3). Both acute and subclinical PID can lead to serious reproductive consequences, including chronic pelvic pain, tubal factor infertility, and ectopic pregnancy (4). Many women with tubal factor infertility do not recall a prior diagnosis of PID or CT infection but show serologic evidence of prior CT exposure (5, 6). These varied clinical presentations and severe morbidities highlight the urgent need to better define the pathogenesis of chlamydial disease and to develop effective interventions—particularly vaccines capable of reducing bacterial burden, accelerating clearance, and preventing transmission and upper genital tract damage.

Excessive inflammation during infection can lead to tubal scarring and post-obstructive dilation of the oviduct, or hydrosalpinx, in women and female mice. However, human *C. trachomatis* strains are rapidly cleared by laboratory mice, limiting their utility for studying the mechanisms that drive oviduct scarring. To overcome this, investigators commonly use *Chlamydia muridarum*, a natural respiratory pathogen of mice (7), as a genital tract infection model via intravaginal inoculation. *C. muridarum* has evolved mechanisms to evade interferon gamma (IFN-γ)-induced GTPase-mediated restriction of its growth (8), enabling a productive infection that typically lasts for ∼28 days (9). Although this model elicits marked inflammation and severe upper genital tract pathology (9), it does not reflect the chronic, low-grade infection that is more typical for humans (10). These limitations emphasize the need for animal models that more accurately capture the heterogeneity and chronicity of human chlamydial infection.

Host genetic variation plays a substantial role in shaping chlamydial disease outcomes, with evidence from both human (11–16) and animal studies (17–19). We previously reported marked strain-dependent differences in the course and severity of *C. muridarum* genital tract infection among three inbred mouse strains, that were associated with distinct immune response kinetics (9). These findings were extended by Chen et al., who examined 11 inbred strains and reported gross hydrosalpinx incidences ranging from 10% to 87% (20). Together, these studies underscore the strong influence of host genetics on infection trajectory and pathology.

Despite this, a persistent challenge in modeling chlamydial genital tract infection is the striking variability in pathologic outcomes, even when infectious burden and clearance kinetics are comparable (9, 20, 21). Substantial variation occurs both across classical inbred strains and within individual strains. For example, Chen et al. reported broad intra-strain dispersion in oviduct dilatation scores, with mean severity ranging from 5.3 ± 3.2 in a highly susceptible strain to 0.6 ± 1.4 in a resistant strain (20). Similar inconsistency is observed across experiments: in our laboratory, ten independent preclinical vaccine studies using genetically identical C57BL/6 mice under standardized conditions yielded gross hydrosalpinx rates ranging from 30% to 100%, with correspondingly wide variation in histologic pathology (22, 23)(unpublished data). This degree of intrastrain and inter-experimental variability limits the precision of genetic mapping and constrains the accuracy of vaccine efficacy assessments.

The Collaborative Cross (CC) genetic reference population offers a powerful strategy to address variability in chlamydial disease modeling. CC strains were generated by interbreeding eight genetically diverse founder strains—five classical inbred strains (A/J, C57BL/6J, 129S1/SvImJ, NOD/ShiLtJ, and NZO/HlLtJ) and three wild-derived strains (CAST/EiJ, PWK/PhJ, and WSB/EiJ)—followed by inbreeding to produce stable recombinant lines (24). This multi-allelic architecture preserves the experimental reproducibility of inbred mice while buffering against the effects of single detrimental variants and capturing a level of genetic diversity that more closely reflects human populations (25). As a result, CC panels are well suited for dissecting polygenic traits and enabling high-resolution mapping of loci that influence infectious disease susceptibility and vaccine responsiveness (26).

To test the utility of this approach, we infected CC mice intravaginally with *C. muridarum* and evaluated 20 strains selected for documented variation in infection susceptibility or immune-mediated disease (27–30). Across the panel, we observed marked strain-dependent differences in bacterial burden, infection duration, and gross and histologic pathology, with low within-strain variability. Notably, several disease-resistant strains exhibited prolonged or chronic infection kinetics, paralleling features of *C. trachomatis* infection in women. Quantitative trait locus (QTL) mapping identified loci associated with bacterial burden and pathological severity, providing insight into host determinants of chlamydial disease, and demonstrating the value of CC strains for capturing the heterogeneity of infection outcomes, and complementing traditional inbred mouse systems used for preclinical vaccine evaluation.

## MATERIALS AND METHODS

### Ethics statement

All animal procedures were approved by the Institutional Animal Care and Use Committee (IACUC) at the University of North Carolina at Chapel Hill (UNC-CH) and all experiments involving *C. muridarum* were conducted in accordance with institutional biosafety guidelines.

### Mouse model of *Chlamydia* infection

Collaborative Cross mice were obtained from the Systems Genetics Core Facility at UNC-CH (https://csbio.unc.edu/CCstatus/index.py). A total of 154 female mice from 20 CC strains (6–9 mice per strain) were evaluated over 25 time points (see **Fig. 1** for experimental workflow). Strains studied were CC001, CC002, CC004, CC005, CC006, CC012, CC013, CC019, CC023, CC024, CC027, CC030, CC031, CC036, CC037, CC041, CC051, CC065, CC068, and CC078. Mice were age-matched and used between 8 and 12 weeks of age. Mice were housed in the pathogen-free animal facility at UNC-CH and provided food and water ad libitum in an environmentally controlled room with a cycle of 12 hours of light and 12 hours of darkness. All mice were administered 2.5 mg of depot medroxyprogesterone acetate (Depo-Provera; Pfizer) subcutaneously to induce anestrus.

**Figure 1.**
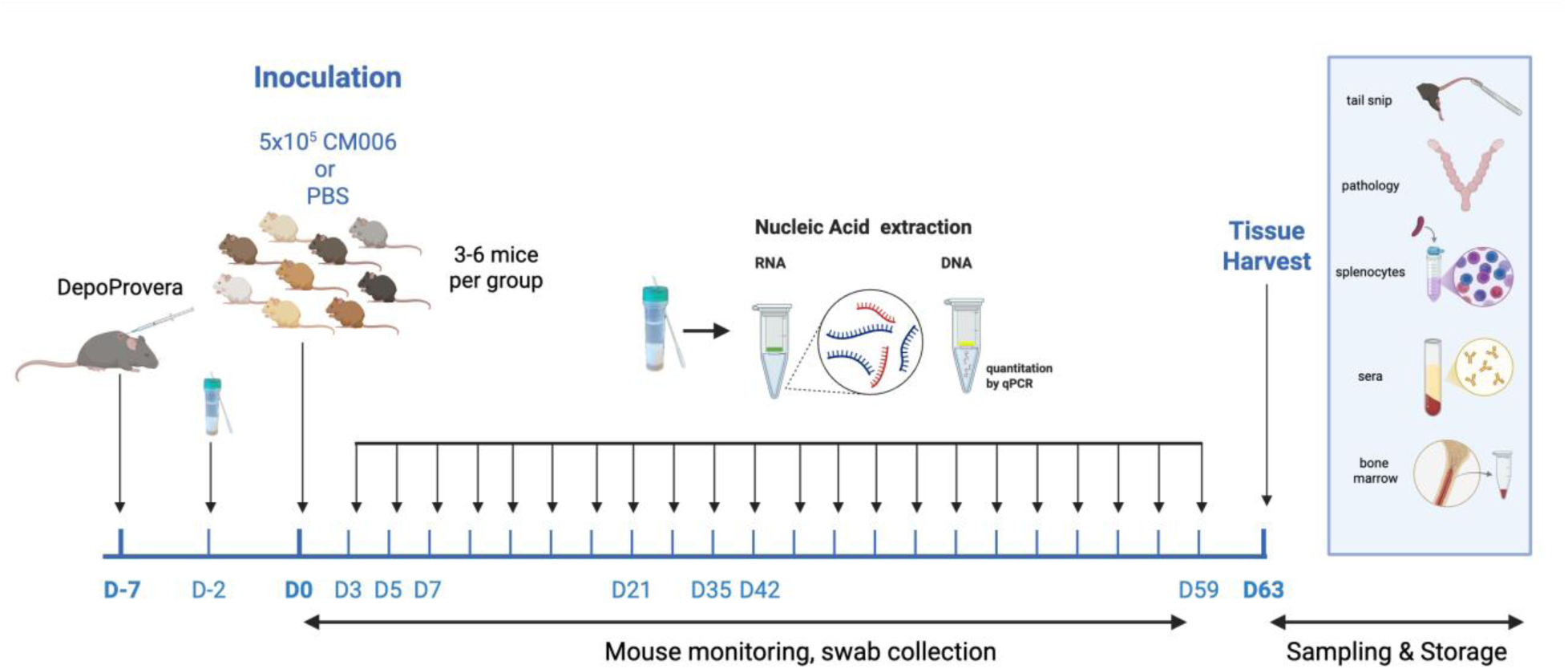
Mouse experimental scheme. Twenty Collaborative Cross (CC) strains were pretreated with Depo-Provera and intravaginally inoculated with either *C. muridarum* strain CM006 or PBS. Cervical swabs were collected throughout infection to monitor bacterial load and clearance. Following resolution of infection, mice were sacrificed for evaluation of genital tract pathology, and serum and immune cells were banked for future analyses. Created in BioRender. O’Connell, C. (2026) https://BioRender.com/t0b0v2x.

Seven days post-progesterone treatment, 79 mice (3–6 per strain) were anesthetized with Nembutal (sodium pentobarbital, 50 mg/ml), diluted 1:10 in sterile PBS, delivered intraperitoneally, in a volume of 10 μl/g mouse weight, and intravaginally inoculated with 5 × 10⁵ inclusion-forming units (IFU) of *C. muridarum* CM006. CM006, a plaque-purified clonal isolate derived from the parental Nigg stock (31), was delivered in 20 μl of sucrose–sodium phosphate–glutamic acid (SPG) buffer containing 250 mM sucrose, 10 mM sodium phosphate, 5 mM l-glutamic acid (pH 7.2). The remaining 75 mice received phosphate-buffered saline (PBS) as mock controls. Cervical swabs were collected pre-infection (day −2) and on post-infection days 2 to 10, 14, 17, 21, 24, 28, 31, 35, 38, 42, 45, 49, 52, 56, 59, and 63. Swabs were stored in 1 mL of DNA/RNA Shield (Zymo Research, Irvine, California, USA) at −80°C for transcriptional response profiling and determination of chlamydial burden. On day 63, mice were euthanized, and reproductive tracts were harvested en bloc for gross and histopathologic analysis.

### Gross and histopathological assessment

In situ examination of the reproductive tract was performed by a trained technician blinded to experimental groups. Gross pathology was scored as follows: 0, normal; 1, enlarged cervix; 2, swollen or discolored uterine horn (left and/or right horn); 3, unilateral hydrosalpinx; and 4, bilateral hydrosalpinx. The genital tract was then collected in its entirety with careful removal of adipose as needed, and mounted on thin cardboard, then fixed in 10% formalin in PBS for 48 h and stored in PBS prior to paraffin embedding. Sections were cut at 4 μm distally, from oviduct to cervix, and stained with hematoxylin and eosin. Oviduct regions of histological samples were evaluated in a masked fashion by a board-certified pathologist, with an assessment of leukocyte infiltration (neutrophils, mononuclear cells, and plasma cells) and dilation reported for each oviduct using a four-tiered semi-quantitative scoring system; 0 = normal or none, 1 = mild, 2 = moderate, 3 = marked, 4 = severe (9).

### Nucleic acid isolation and quantification of bacterial load

DNA and RNA were co-extracted from cervical swabs as previously described (32) using a Quick-DNA/RNA Miniprep Plus Kit (Zymo Research). Chlamydial loads from samples obtained on days −2, 7, 10, 35 and 59 were measured by quantitative PCR using primers (23S_F1 5’ GCTCACGTTCGGAAAGGATAA 3’ and 23S_R1 5’ GTGCTTACACCTCCAACCTATC 3”) that targeted the *C. trachomatis* 23S rRNA loci using the following amplification conditions: 95°C for 15’, 60°C for 45’ for 40 cycles using SsoAdvanced Universal SYBR Green Supermix (BioRad Life Science, Hercules, CA) followed by melt curve analysis. Each specimen was analyzed in a triplicate.

#### Gene expression profiling

Six CC strains (CC005, CC012, CC023, CC030, CC031, CC041) representing a range of phenotypes were selected for targeted transcriptional analysis using probe-based quantitation (33, 34). RNA extracted from swab eluates (3–5 mice per strain) obtained on days −2, 3, 5, 7, and 35 was analyzed using nCounter® Gene Expression Assay (35) (NanoString Technologies, Seattle, WA) at the UNC Lineberger Comprehensive Cancer Center Translational Genomics Laboratory. A custom panel comprised of 51 immune-related mouse genes and 6 internal references (*Gapdh, Hprt, Cltc, Gusb, Pgk1,* and *Tubb5*) was chosen. Probes targeting chlamydial RNAs (23S rRNA, *omcA*, pGP8 anti-sense RNA) were also designed and included in the assay to enable monitoring of chlamydial RNA abundance (**Supplementary Table 1**).

#### Statistical analyses

##### Heritability estimates

Broad-sense heritability (H²) and narrow-sense heritability (h²) were calculated to assess the contribution of genetic variation to phenotypic traits. For each phenotype, genetic variance was determined by subtracting within-strain variance, primarily influenced by environmental factors, from between-strain variance, which includes both genetic and environmental factors. H² accounts for additive, dominance, and epistatic effects (36, 37) whereas h² reflects only additive effects (38). Estimates were derived using the est_herit function in the qtl2 R package (v3.1-0), incorporating a kinship matrix and variance estimated from a mixed-effects model (39).

##### Quantitative trait locus (QTL) mapping

QTL mapping for bacterial burden and pathology scores was performed using CC genotypes obtained from the QTL Archive (https://qtlarchive.org) as implemented in the qtl2 R package (39). Founder haplotype probabilities were calculated via a hidden Markov model (40). A genome-wide (GRCm38, bioproject PRJNA20689) mixed-effects model scan was conducted, and significance thresholds for Logarithm of Odds (LOD) scores were established using 1,000 permutations. Confidence intervals were defined using a Bayesian credible interval estimation implemented in R/qtl2 package. Genes within each QTL interval were annotated using Mouse Genome Informatics (MGI) (41) and the UCSC Genome Browser (42). For both burden- and pathology-associated loci, genes located within 10 Mb upstream and downstream of the peak were examined, and targeted literature searches were performed to identify published evidence that linked these genes with chlamydial growth, replication, or disease pathogenesis.

##### Intraclass correlation and comparative analysis

Within-strain reproducibility of hydrosalpinx (a binary outcome: presence vs. absence) was assessed using intraclass correlation coefficients (ICCs). The ICC is the proportion of total variance in the outcome that is due to differences between strains, with total variance defined as the sum of variance between and within strains. ICCs for the 20 CC strains analyzed in this study and the 11 inbred strains (data from Table 3 of Chen *et al.* (20)) were estimated separately using logistic mixed-effects models in the lme4 R package (43), with a random intercept for strain. To test whether ICCs differed between studies, we applied a hierarchical bootstrap (44) that resampled strains with replacement within each study, followed by resampling mice within strains, and computed the ICC difference. This approach preserves the nested data structure and accounts for unbalanced designs, including differences in the number of strains and the number of replicate mice per strain. The bootstrap distribution of ICC differences was used to construct percentile 95% confidence intervals and assess statistical significance. We also performed a secondary analysis, a meta-analytic comparison of ICCs, as a sensitivity analysis to assess the robustness of our findings to unbalanced sample sizes. A random-effects meta-analysis with inverse-variance weighting was used to account for both between-study heterogeneity and differences in sample sizes across comparisons. Analyses were conducted using the metafor R package (45).

##### Association of cervical cytokine mRNA expression with burden and pathology

NanoString expression data were processed using nSolver v3.0 (46). Quality control filtering excluded flagged or low-expression genes (defined as counts < mean + 3 SD of negative controls). Technical normalization was performed using spike-in control probes, with lane-specific scaling factors derived from their geometric means. Sample-to-sample normalization was then performed using housekeeping genes, applying the same geometric mean approach. Associations between gene expression and 23S RNA load were analyzed using linear mixed-effects models with CC strain as a random effect. Aligned Rank Transform (ART) ANOVA (47, 48) and ARTool R package (49) were used to assess associations with gross pathology scores. To evaluate whether cervical cytokines were associated with pathology independent of bacterial burden, we used a nonparametric factorial ANOVA with align-and-rank transformation. This approach accommodates non-normal distributions and unequal variances while allowing adjustment for bacterial load as a covariate. P-values were obtained from rank-transformed models, and multiple-testing corrections were performed using the Benjamini–Hochberg procedure.

##### Time-series co-expression analysis

Temporal co-expression patterns among host genes were analyzed in the 6 selected CC strains using Lag Penalized Weighted Correlation (LPWC) (50). The optimal number of gene clusters was determined using the Gap Statistic (51). To characterize temporal patterns of cervical host responses, we used the Lag Penalized Weighted Correlation (LPWC) algorithm, which clusters genes based on similarity in expression trajectories while allowing for temporal lags between samples. LPWC was applied to normalized gene-expression data from Days 0, 3, 7, 21, 35, and 42. Cluster stability was assessed using the default penalty and weighting parameters. Gene clusters were interpreted using functional enrichment and manual annotation of immune-related pathways.

## Results

### Chlamydial burden and pathology vary independently across CC strains

We intravaginally inoculated 20 CC strains with *C. muridarum* or sham-infected controls with PBS and monitored them through day 63 for infection outcomes, including chlamydial burden, time to clearance, and genital tract pathology. For analysis, log_10_-transformed cervical chlamydial loads were categorized as: very low (<4), low (4–5), intermediate (5–6), and high (>6). Infection duration was classified as: normal clearance: (no detectable Cm genomic DNA by qPCR on D35, extended (DNA detectable on D35), and prolonged (DNA detected through D59).

All CC strains became infected following the challenge. Bacterial loads detected in the lower genital tract and infection duration varied widely across strains (**Fig. 2, and Supplementary Fig. 1**) but were highly consistent within each strain (**Fig. 3C, D**). One strain, CC004, cleared infection rapidly, with all mice reaching the limit of detection by day 21 (**Fig. 2B**). In contrast, mice from CC013, CC030 and CC019 showed prolonged infection, with detectable chlamydial DNA still present on day 59. Among the remaining strains, five cleared infection by day 35, and ten strains appeared to clear between days 35 and 59 (**Fig. 2C** and **Supplementary Fig. 1**).

**Figure 2.**
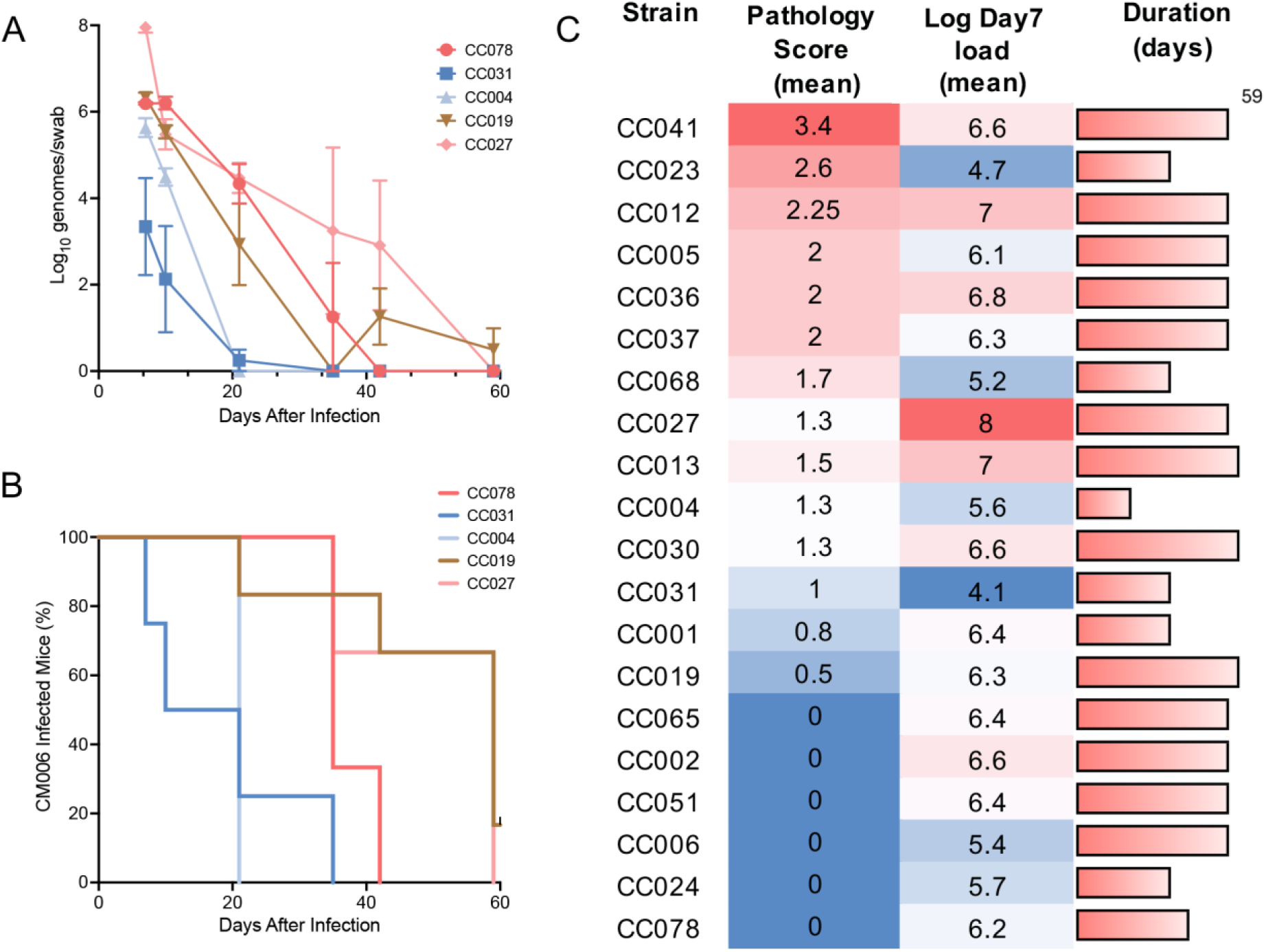
Collaborative Cross (CC) mice exhibit wide variation in infection dynamics following *C. muridarum* challenge. Lower genital tract shedding of strain CM006 was quantified by qPCR targeting the 23S rRNA locus. (A) Mean chlamydial burden ± SEM is shown for a representative subset of CC strains (3–6 mice per group). (B) Time to infection clearance was defined as the first day on which chlamydial genomic DNA fell below the assay limit of detection (<8.3 × 10¹ genomes/swab). (C) Heat map summarizing pathology scores, Day 7 bacterial burden, and infection duration measured through day 59 across all twenty CC strains. Strain sample sizes: CC078 (n=3), CC031 (n=4), CC004 (n=3), CC019 (n=6), CC027 (n=3).

**Figure 3.**
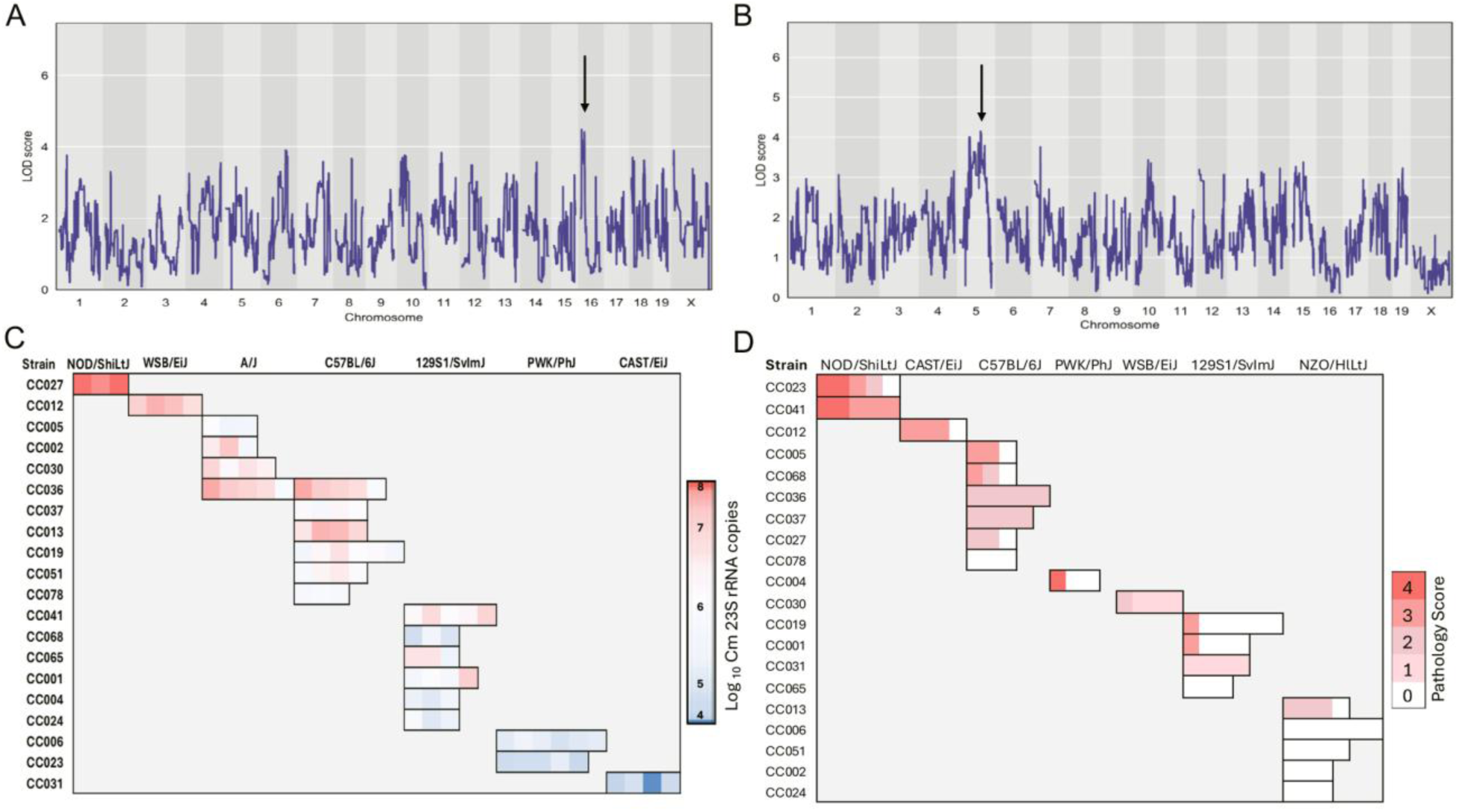
QTL mapping reveals loci on Chromosomes 5 and 16 that influence chlamydial infection outcomes in CC mice. Genome-wide QTL scans (LOD score profiles) identified significant associations between host genetic loci and infection phenotypes. (A) Day 7 chlamydial burden showed its strongest association on Chromosome 16; the corresponding QTL spans a 95% Bayesian credible interval from 6.8 to 21.4 Mb. (B) Day 63 pathology scores showed their strongest association on Chromosome 5; this QTL spans a 95% Bayesian credible interval from 40.6 to 126.0 Mb. Arrow indicates the position of LOD peak. (C–D) Founder haplotype probabilities at the peak markers for these loci are shown for each mouse within each CC strain, illustrating the distribution of inferred founder alleles. Notably, CC036 exhibits an equal (50/50) probability of A/J and C57BL/6J founder alleles at the Chromosome 16 burden locus, so chlamydial burden values are indicated for each.

Across the 20 CC strains the overall hydrosalpinx incidence was 21.5% at tissue harvest. Twelve strains showed no hydrosalpinx; one showed low incidence despite prolonged infection (CC019: 16.7%); another three showed moderate incidence (CC001, CC004, and CC068: 25–33.3%), and four strains displayed high incidence (CC005, CC012, CC023, and CC041: 60-100%). Six CC strains (CC002, CC006, CC024, CC051, CC065, and CC078) showed no gross pathology of the upper or lower genital tract (mean pathology score = 0) despite medium to high Day 7 bacterial loads (**Fig. 2C**). Histopathology supported these findings with none of the oviducts showing a dilatation score > 3. Most mice of CC002, CC006, CC051, and CC078 strains showed no oviduct inflammation, with only occasional mild or moderate mononuclear and plasma cell infiltrates in the oviduct or mesosalpingeal tissues. Strains CC013, CC027, and CC030, had low mean gross pathology scores (1.3- 1.5), no gross hydrosalpinx, and no oviduct dilatation by histology, despite high early bacterial burden and prolonged infection. Despite the absence of gross oviduct dilatation, CC013 mice that were confirmed *C. muridarum* positive by cervical PCR on day 59, exhibited moderate to severe histiocytic inflammation in the bursal and mesosalpingeal tissues, a pathologic change not seen in any PBS-inoculated CC013 controls.

The CC004 strain, despite clearing infection rapidly, developed mild to marked oviduct dilatation; one mouse of three exhibited bilateral hydrosalpinx along with mild inflammatory infiltrates in the mesosalpinx and bursa. CC023 mice had a relatively high mean gross pathology score (2.6), with hydrosalpinx in 5 of 10 oviducts and corresponding histologic dilatation scores of 3-4, despite relatively low bacterial burdens (**Fig. 2C and Supplementary Fig. 1**). None of the PBS-inoculated controls developed hydrosalpinx, although occasional mild uterine hyperemia or hydrometra was observed, likely due to Depo-Provera treatment. Together, these findings demonstrate that in genetically diverse CC mice, neither chlamydial burden nor infection duration reliably predicted pathological outcomes.

### Quantifying genetic influence on chlamydia burden and disease

Heritability describes the proportion of variation in a trait that is explained by genetic differences rather than environmental factors. Narrow-sense heritability reflects the proportion of variance attributable to additive genetic effects, those that sum predictably across alleles. Broad-sense heritability captures all genetic contributions, including additive effects as well as dominance and epistatic (gene–gene interaction) effects. Among the 20 CC strains, heritability of bacterial burden was highest on Day 7, with narrow-sense and broad-sense estimates of 69.3% and 76.4%, respectively (**Table 1**). For gross pathology scores, narrow-sense heritability was 50.1% and broad-sense heritability was 57.5% (**Table 1**). These findings indicate that host genetic factors were major contributors to variation in both bacterial load and pathology following *C. muridarum* infection.

**Table 1.**
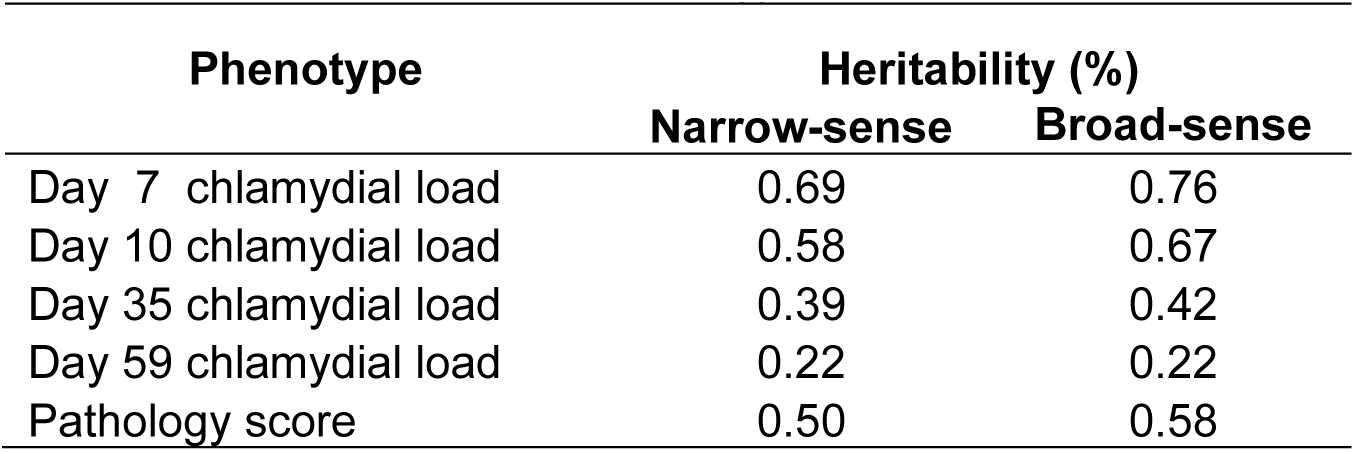
Heritability of chlamydial load and gross pathology.

| Phenotype | Heritability (%) |  |
| --- | --- | --- |
|  | Narrow-sense | Broad-sense |
| Day 7 chlamydial load | 0.69 | 0.76 |
| Day 10 chlamydial load | 0.58 | 0.67 |
| Day 35 chlamydial load | 0.39 | 0.42 |
| Day 59 chlamydial load | 0.22 | 0.22 |
| Pathology score | 0.50 | 0.58 |

### QTL mapping reveals loci associated with chlamydial burden and pathology

We proceeded to examine the influence of genotypes on bacterial load and pathology using a genome-wide scan with CC genotype probabilities obtained from the QTL Archive (https://qtlarchive.org). Although no genome-wide significant QTLs were detected for either trait after permutation correction (p < 0.05), we identified suggestive loci on chromosome 16 (∼7.1 Mb) and 5 (∼98.3 Mb). These peaks yielded LOD scores >4.5 for Day 7 bacterial load (**Fig. 3A**) and 4.15 for pathology (**Fig. 3B**). Such LOD scores correspond to odds >10,000:1 in favor of association compared to chance alone, providing strong suggestive evidence that these regions may contribute to variation in chlamydial infection outcomes.

Genes within the locus associated with chlamydial burden mapped to host pathways previously implicated in chlamydial intracellular development and host control **(Supplementary Table 2)**. Several candidates localized to membrane dynamics and lipid metabolism, processes central to inclusion formation and maintenance. Notably, *Pi4ka*, encoding phosphatidylinositol 4-kinase α, lies within a pathway shown to support inclusion membrane biogenesis through host phosphatidylinositol-4-phosphate supply (52, 53) while *Smpd4*, a sphingomyelinase, maps to sphingolipid metabolic pathways required for inclusion stability and bacterial replication (54). Genes involved in ubiquitin signaling (*Ube2l3, Ube2v2*) were also detected; ubiquitin-dependent targeting of chlamydial inclusions is part of cell-autonomous host defense (55), and *Chlamydia* encodes effectors that modulate this process (56, 57), suggesting that host ubiquitination capacity may influence chlamydial burden.

Additional candidates were linked to host cell stress responses and survival, including *Dnm1l* (*DRP1*), which regulates mitochondrial fission and apoptosis susceptibility, and is inhibited by chlamydial infection (58). Genes involved in DNA damage sensing and repair (*Prkdc, Ercc4, Mcm4, Spidr, Top3b*) were also detected. *Chlamydia* infection induces host DNA damage while altering repair and checkpoint responses (59), making host DNA repair capacity a reasonable modifier of cellular survival and consequently, chlamydial replication. The locus also contained centrosome- and spindle-regulating genes (*Cep20, Mzt2*), processes disrupted during chlamydial infection and associated with enhanced host cell survival and inclusion stability (60). Finally, immune-regulatory genes (*Ciita, Mapk1, Socs1*) were present, implicating variation in interferon-responsive and inflammatory signaling pathways as additional contributors to differences in bacterial burden across Collaborative Cross strains. The identification of *Ciita*, the master regulator of MHC class II expression, is particularly relevant given the established requirement for CD4⁺ T cell–mediated immunity in controlling chlamydial infection. While these associations do not establish causal roles for individual genes, the convergence of multiple candidates within host pathways known to support intracellular growth provides a biologically coherent framework for interpreting genetic effects on chlamydial burden. A full list of genes in this locus is provided in **Supplementary Table 3**.

We next assessed the effects of the eight founder alleles at this locus on Day 7 chlamydial load (**Fig. 3C**). The allelic effects clustered into three groups. Mice carrying the **NOD/ShiLtJ (D)**, **WSB/EiJ (H)**, **A/J (A)**, or **C57BL/6J (B)** alleles showed the highest bacterial loads, with log_10_ values >6. In contrast, mice with the **PWK/PhJ (G)** or **CAST/EiJ (F)** alleles had lower burdens, with log_10_ values of 4–6 and 0–5, respectively. The **129S1/SvImJ (C)** allele produced an intermediate phenotype, with bacterial loads ranging from log_10_ ∼4.5 to 7.

The pathology-associated locus on Chromosome 5 encompasses multiple genes (**Supplementary Table 4**) with established roles in host immune responses and genital tract pathology following chlamydial infection. Included in this locus were genes encoding ELR+ CXC chemokines (*Cxcl1*, *Cxcl2*, *Cxcl3*, *Cxcl5* and *Cxcl15*), key mediators of neutrophil recruitment to sites of injury or infection (61). Neutrophil influx has been consistently linked to tissue damage in murine models of chlamydial genital tract infection (62–64). Also present were genes encoding IFN-inducible chemokines, (*Cxcl9*, *Cxcl10* and *Cxcl11*), that attract CXCR3+ Th1 and cytotoxic T cells to aid pathogen clearance but also implicated in immunopathology (62, 65, 66). *Cxcl13*, which supports ectopic lymphoid follicle formation, contributes to persistent inflammation and fibrosis in chronic infection settings (66, 67). *Bmp3*, part of the BMP/TGF-β superfamily, modulates tissue remodeling and dysregulated BMP signaling has been linked to fallopian tube scarring and infertility in human studies (15, 68). Additional genes in this locus include heparinase (*Hpse*) which influences extracellular matrix remodeling (69); multiple guanylate-binding protein (Gbp) family members involved in inflammasome activation during chlamydial infection (70, 71); and osteopontin (*Spp1*), a multifunctional cytokine linked to inflammatory cell recruitment and tissue remodeling (72, 73). A complete list of genes within this interval is provided in **Supplementary Table 5**.

We next examined the effects of the eight founder alleles at this locus on pathology scores (**Fig. 3D**). The allelic effects grouped into three tiers. Mice carrying the **NOD/ShiLtJ (D)** or **CAST/EiJ (F)** alleles showed the highest pathology scores. In contrast, the **WSB/EiJ (H)**, **129S1/SvImJ (C)**, or **NZO/HlLtJ (E)** alleles were associated with the lowest pathology. The **C57BL/6J (B)** and **PWK/PhJ (G)** alleles produced intermediate pathology scores, with the **C57BL/6J (B)** allele showing the greatest variability both within and across CC strains.

### Within-strain reproducibility of hydrosalpinx rates in CC mice

To quantify within-strain consistency, we calculated the intraclass correlation coefficient (ICC) for hydrosalpinx. The ICC was 0.598 across the 20 CC strains examined here, compared with 0.275 across 11 previously profiled inbred strains (20). To account for unequal sample sizes, we applied both bootstrap and meta-analytic approaches, which yielded a bootstrap P = 0.0003 and a meta-analysis P = 0.007. Together, these findings show that hydrosalpinx outcomes are markedly more consistent within CC strains than within commonly used inbred laboratory strains.

### Relationship between cervical cytokine mRNA expression with bacterial burden and pathology

Human (74–76) and murine (62, 77–79) studies have shown that pro-inflammatory responses contribute substantially to immune-mediated pathology following chlamydial infection. To explore how host genetics shape these responses, we assessed the relationship between expression of 51 cervical cytokine genes and bacterial burden in six CC strains (CC005, CC012, CC023, CC030, CC031, and CC041) spanning a broad range of bacterial loads, infection durations, and gross pathology. Cytokine genes were significantly associated with bacterial burden on days 3, 5, and 7, (listed in **Supplementary Tables 6-8)**. Notably, no cytokine gene displayed a significant negative association with burden at any early time point, and no significant cytokine–burden associations were detected on day 35.

Of the 51 genes measured, 25, 8, and 11 were significantly upregulated in association with increased chlamydial burden on days 3, 5, and 7, respectively. Six genes, *Il17a*, *Il22*, *Il10*, *Il12a*, *Ifng*, and *Ifnb1*, showed significant positive associations with bacterial burden at all three early time points. These included both pro-inflammatory (*Il17a*, *Il22*, *Ifng*, *Ifnb1*) and regulatory/anti-inflammatory (*Il10*, *Il12a*) mediators, with *Il12a* also contributing to Th1 differentiation and the production of IFN-γ and TNF. *Cxcl10,* a chemokine that promotes T cell recruitment, and *Eomes*, a transcription factor critical for T cell differentiation, were significantly associated with bacterial burden on days 3 and 5. Several genes, *Il23*, *Cxcl1*, *Stat3*, *Ccl4*, *Ccl3*, *Tlr2*, *Il6*, *Il1rn*, *Tnf*, *Il15*, *Il18*, and *Gata3*, were uniquely associated with increased bacterial burden on day 3, whereas *Ltb* showed a unique association on Day 7.

We also sought to determine whether cytokine mRNA expression was associated with pathology scores in these 6 CC strains. After adjusting for bacterial burden, *Ccl4, Cxcl9,* and *IL1ra* transcript levels were modestly higher in mice with hydrosalpinx compared to those without (P = 0.039, 0.049, and 0.068, respectively). However, none of these associations were statistically significant after correction for multiple testing.

### Temporal co-expression patterns of cervical host genes in 6 selected CC strains

Because the timing and coordination of immune gene expression are critical determinants of infection outcome, we examined temporal patterns of cervical gene expression using a clustering approach that accommodates differences in activation timing. This analysis resolved two major trajectories: an early-response cluster enriched for genes involved in initiating and recruiting a T-cell response (Fig. 4A), and a later-response cluster dominated by markers of T-cell infiltration (e.g., *CD4, CD3*), differentiation (*Tbx21, Foxp3*), and effector function (*IL-2, IL-21, IL-16*) (Fig. 4B).

**Figure 4.**
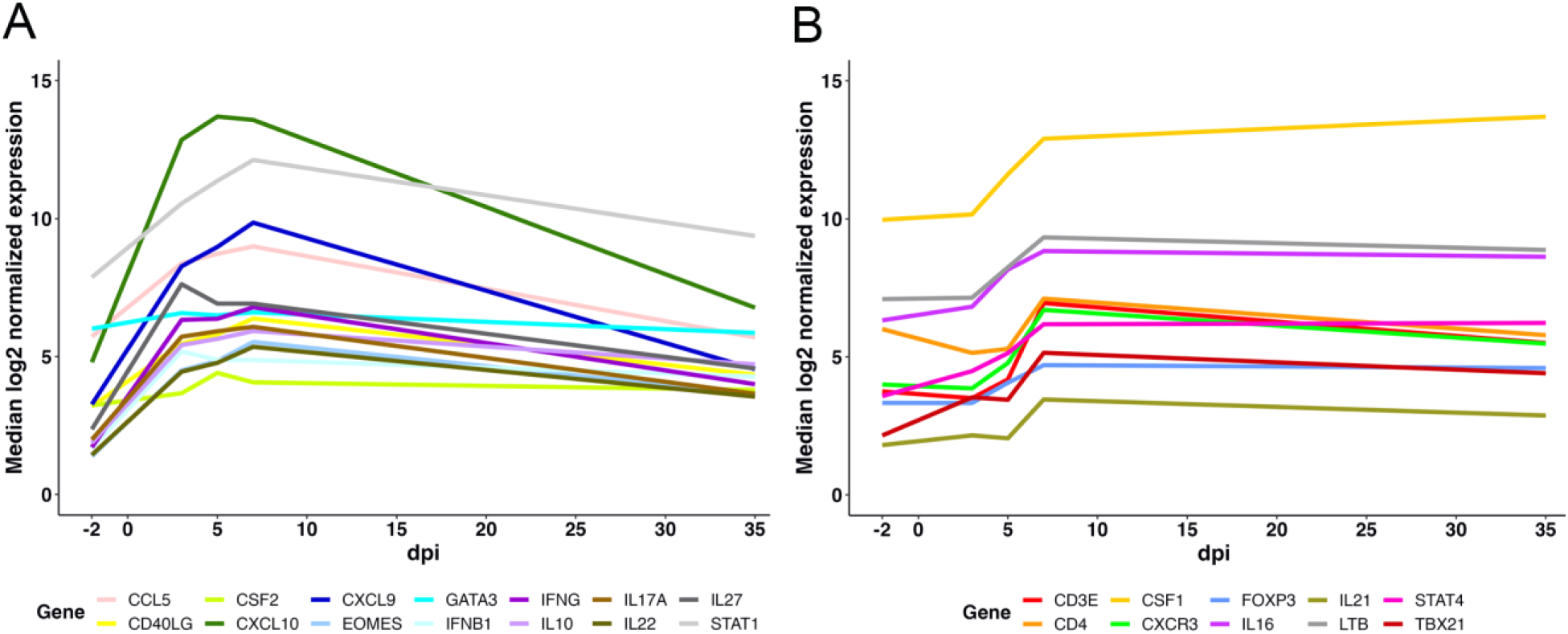
Time series co-expression of cervically-expressed host infection responses for 6 selected CC strains. Lag Penalized Weighted Correlation (LPWC) test used for this analysis allows for temporal offsets and LPWC analysis identified co-expression groups of immune genes upregulated with infection onset (A) and genes showing increased expression levels 5 days post-infection (B). Selected host transcripts (N=50) were quantified in total RNA extracted from mouse swabs (Days −2, 3, 5, 7, 10, 35) using nCounter assay with additional probes used for normalization (host N=6, chlamydiae N=3).

## Discussion

Using 20 CC mouse strains infected with *C. muridarum*, we observed broad, genetically reproducible variation in infection outcomes that parallel the heterogeneity of human chlamydial disease. Several strains exhibited prolonged or near-chronic infection with minimal pathology, resembling persistent, asymptomatic infections in women (3, 10), whereas others developed hydrosalpinx despite low bacterial burden. This burden–pathology discordance, previously noted in select inbred strains (20, 53), was consistently reproduced across the CC panel, enabling systematic genetic dissection of mechanisms that uncouple pathogen load from tissue damage. Together, these findings indicate that host genetic background, rather than bacterial burden alone, is a major determinant of inflammatory regulation and disease severity.

Across all CC strains, hydrosalpinx occurred in ∼20% of mice, comparable to infertility risk estimates following chlamydial PID in women (80, 81), underscoring the translational relevance of the model. Notably, pathology outcomes were far more consistent within CC strains than within classical inbred strains (ICC 0.598 vs. 0.275; p < 0.001). Although inbred strains are often presumed to exhibit phenotypic stability, prior studies demonstrate substantial variability in pathological responses (24, 25, 82), consistent with our observations in C57BL/6 mice and with data reported by Chen et al. (20). Fixed deleterious or threshold-sensitive alleles and limited redundancy in immune regulatory pathways likely render inbred strains vulnerable to stochastic influences. In contrast, CC strains inherit diverse combinations of alleles from eight founders, buffering single-variant effects and producing tighter phenotypic clustering, thereby enhancing reproducibility for genetic mapping and vaccine studies.

Heritability estimates were high for both Day 7 bacterial burden (narrow-sense 69.3%; broad-sense 76.4%) and pathology (narrow-sense 50.1%; broad-sense 57.5%), indicating that host genetics accounted for most observed variation. The modest difference between broad- and narrow-sense estimates suggests that additive effects predominate, with a smaller contribution from non-additive interactions. These findings validate the use of CC strains for detecting genetic loci underlying complex traits such as clearance and immunopathology and indicate that environmental noise is unlikely to obscure genetic signals within strains. Importantly, these results align with human genetic–epidemiologic data, including twin studies showing substantial heritable contributions to immune responses against *C. trachomatis* (83), reinforcing the translational relevance of genetically diverse model systems.

Genome-wide scans identified distinct suggestive QTLs for bacterial burden (Chromosome 16) and pathology (Chromosome 5). Genes within the burden-associated locus converged on host pathways that support chlamydial intracellular growth, including membrane dynamics and lipid metabolism, ubiquitin signaling, host cell survival, DNA damage responses, centrosome regulation, and immune-regulatory signaling, processes that are actively modulated during infection and influence inclusion stability and bacterial replication. Although individual causal genes were not resolved, pathway-level convergence provides a coherent biological framework linking host genetic variation to differences in chlamydial burden.

Genes within the pathology-associated locus converged on host pathways that regulate inflammatory cell recruitment, tissue remodeling, and chronic immune activation, processes central to genital tract damage following chlamydial infection. Prominent representation of chemokine networks governing neutrophil and Th1 cell trafficking, together with genes linked to ectopic lymphoid structure formation, supports a model in which genetically determined variation in immune cell localization and persistence contributes to pathological outcomes. The presence of additional candidates involved in extracellular matrix remodeling, inflammasome activation, and BMP/TGF-β signaling further implicates dysregulated host repair and fibrotic responses as key modifiers of disease severity.

Founder allele analysis further demonstrated dissociation between genetic effects on burden and pathology, reinforcing the concept that pathogen control and tissue damage are partially independent of host processes. Variability associated with certain founder alleles, including C57BL/6J, underscores the limitations of relying on conventional inbred strains and highlights the strength of the CC for uncovering genetic effects otherwise obscured.

Cervical cytokine profiling provided additional insight into immune dynamics. Early expression of inflammatory and regulatory cytokines correlated positively with bacterial burden, suggesting that early inflammation primarily reflects ongoing replication rather than effective control, consistent with observations in women (84). Temporal trajectory analyses revealed a shift from innate to adaptive immune programs, aligning with the known requirement for CD4 Th1–mediated IFN-γ responses for clearance while suggesting that genetic background influences how effectively early innate cues shape protective T-cell responses (16).

Although no cytokine remained significant after multiple-testing correction in this small dataset, CXCL9, CCL4, and IL1RA emerged as the strongest burden-adjusted predictors of pathology. CXCL9, produced by IFN-γ–stimulated immune and epithelial cells, recruits CXCR3⁺ effector T cells to inflamed tissues (62, 85). CCL4, generated by innate immune cells and activated T cells, recruits CCR5⁺ effector T cells, amplifying local inflammatory responses (86–88). In contrast, IL1RA restrains inflammation by blocking IL-1 signaling (89), pointing to a balance between chemokine-driven effector recruitment and inflammatory restraint as a key modifier of tissue damage.

Collectively, these findings demonstrate that CC mice offer a robust and clinically relevant platform for modeling chlamydial disease. Their genetic diversity yields phenotypes that mirror human variability, while fixed genomes ensure high reproducibility. This combination has important implications for vaccine development, particularly for evaluating interventions across genetically distinct hosts that dissociate clearance from pathology. Prioritizing CC strains with extreme but reproducible phenotypes may improve power to detect true vaccine effects and enable mechanistic dissection of protective versus pathogenic immune programs.

Several limitations warrant consideration. *C. muridarum* differs biologically from *C. trachomatis*, and larger cohorts or expansion to CC-F2 or Diversity Outbred populations will increase power for genome-wide significance. Cytokine profiling was limited to six strains, and broader sampling may reveal additional immune architectures. Integration of single-cell transcriptomics, proteomics, and causal network modeling (90) will be essential for pinpointing molecular drivers of protection and pathology.

In summary, this study demonstrates that host genetics profoundly influences chlamydial burden, persistence, and immunopathology and that CC mice recapitulate key features of human disease more comprehensively than conventional inbred strains. By delineating genetically distinct axes of pathogen control and tissue damage, the CC provides a powerful framework for mechanistic discovery and for advancing vaccines and therapeutics in a genetically diverse, clinically relevant context.

## Supporting information

All supplemental figure and tables

## Acknowledgements

We would like to acknowledge the histopathology scoring assessments of Rani Sellers, PHD, DVM, DACVP, UNC Pathology Services Core and the UNC-CH Systems Genetics Core Facility for the maintenance of the Collaborative Cross lines used in these studies. The UNC-CH Systems Genetics Core Facility CC Pilot Program (CC004) provided funding support.

## Bibliography

1. Centers for Disease Control and Prevention. Sexually Transmitted Infections Surveillance, 2024 (Provisional). https://www.cdc.gov/sti-statistics/annual/index.html accessed December 17, 2025.

2. Oakeshott P, Kerry S, Aghaizu A, Atherton H, Hay S, Taylor-Robinson D, Simms I, Hay P. 2010. Randomised controlled trial of screening for *Chlamydia trachomatis* to prevent pelvic inflammatory disease: the POPI (prevention of pelvic infection) trial. BMJ 340:c1642.

3. Wiesenfeld HC, Hillier SL, Meyn LA, Amortegui AJ, Sweet RL. 2012. Subclinical pelvic inflammatory disease and infertility. Obstet Gynecol 120:37–43.

4. Brunham RC, Gottlieb SL, Paavonen J. 2015. Pelvic inflammatory disease. N Engl J Med 372:2039–48.

5. Westrom LV. 1996. Chlamydia and its effect on reproduction. J Br Fer Soc 1:23–30.

6. Tubal infertility: serologic relationship to past chlamydial and gonococcal infection. World Health Organization Task Force on the Prevention and Management of Infertility. Sex Transm Dis 22:71–7.

7. Nigg C. 1942. An unidentified virus which produces pneumonia and systemic infection in mice. Science 95:49–50.

8. Nelson DE, Virok DP, Wood H, Roshick C, Johnson RM, Whitmire WM, Crane DD, Steele-Mortimer O, Kari L, McClarty G, Caldwell HD. 2005. Chlamydial IFN-gamma immune evasion is linked to host infection tropism. Proc Natl Acad Sci USA 102:10658–10663.

9. Darville T, Andrews CW, Jr., Laffoon KK, Shymasani W, Kishen LR, Rank RG. 1997. Mouse strain-dependent variation in the course and outcome of chlamydial genital tract infection is associated with differences in host response. Infect Immun 65:3065–3073.

10. Molano M, Meijer CJ, Weiderpass E, Arslan A, Posso H, Franceschi S, Ronderos M, Munoz N, van den Brule AJ. 2005. The natural course of *Chlamydia trachomatis* infection in asymptomatic Colombian women: a 5-year follow-up study. J Infect Dis 191:907–916.

11. den Hartog JE, Ouburg S, Land JA, Lyons JM, Ito JI, Pena AS, Morre SA. 2006. Do host genetic traits in the bacterial sensing system play a role in the development of *Chlamydia trachomatis*-associated tubal pathology in subfertile women? BMC Infect Dis 6:122.

12. Ouburg S, Spaargaren J, den Hartog JE, Land JA, Fennema JS, Pleijster J, Pena AS, Morre SA. 2005. The CD14 functional gene polymorphism −260 C>T is not involved in either the susceptibility to *Chlamydia trachomatis* infection or the development of tubal pathology. BMC Infect Dis 5:114.

13. Morre SA, Murillo LS, Bruggeman CA, Pena AS. 2003. The role that the functional Asp299Gly polymorphism in the toll-like receptor-4 gene plays in susceptibility to *Chlamydia trachomatis*-associated tubal infertility. J Infect Dis 187:341–2; author reply 342-3.

14. Barr EL, Ouburg S, Igietseme JU, Morre SA, Okwandu E, Eko FO, Ifere G, Belay T, He Q, Lyn D, Nwankwo G, Lillard J, Black CM, Ananaba GA. 2005. Host inflammatory response and development of complications of *Chlamydia trachomatis* genital infection in CCR5-deficient mice and subfertile women with the CCR5delta32 gene deletion. J Microbiol Immunol Infect 38:244–254.

15. Zheng X, Zhong W, O’Connell CM, Liu Y, Haggerty CL, Geisler WM, Anyalechi GE, Kirkcaldy RD, Wiesenfeld HC, Hillier SL, Steinkampf MP, Hammond KR, Fine J, Li Y, Darville T. 2021. Host genetic risk factors for *Chlamydia trachomatis*-related infertility in women. J Infect Dis 224:S64–s71.

16. Zhong W, Kollipara A, Liu Y, Wang Y, O’Connell CM, Poston TB, Yount K, Wiesenfeld HC, Hillier SL, Li Y, Darville T, Zheng X. 2022. Genetic susceptibility loci for *Chlamydia trachomatis* endometrial infection influence expression of genes involved in T cell function, tryptophan metabolism and epithelial integrity. Front Immunol 13:1001255.

17. Ouburg S, Lyons JM, Land JA, den Hartog JE, Fennema JS, de Vries HJ, Bruggeman CA, Ito JI, Pena AS, Lundberg PS, Morre SA. 2009. TLR9 KO mice, haplotypes and CPG indices in *Chlamydia trachomatis* infection. Drugs Today 45 Suppl B:83-93.

18. Lichtenwalner AB, Patton DL, Cosgrove Sweeney YT, Gaur LK, Stamm WE. 1997. Evidence of genetic susceptibility to *Chlamydia trachomatis*-induced pelvic inflammatory disease in the pig-tailed macaque. Infect Immun 65:2250–2253.

19. Tuffrey M, Alexander F, Woods C, Taylor-Robinson D. 1992. Genetic susceptibility to chlamydial salpingitis and subsequent infertility in mice. J Reprod Fertil 95:31–38.

20. Chen J, Zhang H, Zhou Z, Yang Z, Ding Y, Zhou Z, Zhong E, Arulanandam B, Baseman J, Zhong G. 2014. Chlamydial induction of hydrosalpinx in 11 strains of mice reveals multiple host mechanisms for preventing upper genital tract pathology. PLoS One 9:e95076.

21. Ito JI, Jr., Lyons JM, Airo-Brown LP. 1990. Variation in virulence among oculogenital serovars of *Chlamydia trachomatis* in experimental genital tract infection. Infect Immun 58:2021–2023.

22. Poston TB, Girardi J, Kim M, Zwarycz P, Polson AG, Yount KS, Hanlan C, Jaras Salas I, Lammert SM, Arroyo D, Bruno T, Wu M, Rozzelle J, Fairman J, Esser-Kahn AP, Darville T. 2025. Intranasal immunization with CPAF combined with ADU-S100 induces an effector CD4 T cell response and reduces bacterial burden following intravaginal infection with *Chlamydia muridarum*. Vaccine 43:126526.

23. Poston TB, Girardi J, Polson AG, Bhardwaj A, Yount KS, Jaras Salas I, Trim LK, Li Y, O’Connell CM, Leahy D, Harris JM, Beagley KW, Goonetilleke N, Darville T. 2024. Viral-vectored boosting of OmcB- or CPAF-specific T-cell responses fail to enhance protection from *Chlamydia muridarum* in infection-immune mice and elicits a non-protective CD8-dominant response in naive mice. Mucosal Immunol doi:10.1016/j.mucimm.2024.06.012.

24. Noll KE, Ferris MT, Heise MT. 2019. The Collaborative Cross: A Systems Genetics Resource for Studying Host-Pathogen Interactions. Cell Host Microbe 25:484–498.

25. Graham JB, Swarts JL, Mooney M, Choonoo G, Jeng S, Miller DR, Ferris MT, McWeeney S, Lund JM. 2017. Extensive homeostatic T cell phenotypic variation within the Collaborative Cross. Cell Rep 21:2313–2325.

26. Srivastava A, Morgan AP, Najarian ML, Sarsani VK, Sigmon JS, Shorter JR, Kashfeen A, McMullan RC, Williams LH, Giusti-Rodríguez P, Ferris MT, Sullivan P, Hock P, Miller DR, Bell TA, McMillan L, Churchill GA, de Villena FP. 2017. Genomes of the mouse Collaborative Cross. Genetics 206:537–556.

27. Keele GR, Zhang T, Pham DT, Vincent M, Bell TA, Hock P, Shaw GD, Paulo JA, Munger SC, de Villena FP, Ferris MT, Gygi SP, Churchill GA. 2021. Regulation of protein abundance in genetically diverse mouse populations. Cell Genom 1:100003.

28. Nagarajan A, Scoggin K, Gupta J, Aminian M, Adams LG, Kirby M, Threadgill D, Andrews-Polymenis H. 2024. Collaborative Cross mice have diverse phenotypic responses to infection with Methicillin-resistant *Staphylococcus aureus* USA300. PLoS Genet 20:e1011229.

29. Kollmus H, Pilzner C, Leist SR, Heise M, Geffers R, Schughart K. 2018. Of mice and men: the host response to influenza virus infection. Mamm Genome 29:446–470.

30. Pérez Gómez AA, Karmakar M, Carroll RJ, Lawley KS, Amstalden K, Young CR, Threadgill DW, Welsh CJ, Brinkmeyer-Langford C. 2022. Serum cytokines predict neurological damage in genetically diverse mouse models. Cells 11:2044.

31. Poston TB, O’Connell CM, Girardi J, Sullivan JE, Nagarajan UM, Marinov A, Scurlock AM, Darville T. 2018. T Cell-independent gamma interferon and B Cells cooperate to prevent mortality associated with disseminated *Chlamydia muridarum* genital tract infection. Infect Immun 86 e00143–18.

32. O’Connell CM, Brochu H, Girardi J, Harrell E, Jones A, Darville T, Sena AC, Peng X. 2019. Simultaneous profiling of sexually transmitted bacterial pathogens, microbiome, and concordant host response in cervical samples using whole transcriptome sequencing analysis. Microb Cell 6:177–183.

33. Veldman-Jones MH, Brant R, Rooney C, Geh C, Emery H, Harbron CG, Wappett M, Sharpe A, Dymond M, Barrett JC, Harrington EA, Marshall G. 2015. Evaluating robustness and sensitivity of the NanoString Technologies nCounter platform to enable multiplexed gene expression analysis of clinical samples. Cancer Res 75:2587–93.

34. Nielsen T, Wallden B, Schaper C, Ferree S, Liu S, Gao D, Barry G, Dowidar N, Maysuria M, Storhoff J. 2014. Analytical validation of the PAM50-based Prosigna Breast Cancer Prognostic Gene Signature Assay and nCounter Analysis System using formalin-fixed paraffin-embedded breast tumor specimens. BMC Cancer 14:177.

35. Geiss GK, Bumgarner RE, Birditt B, Dahl T, Dowidar N, Dunaway DL, Fell HP, Ferree S, George RD, Grogan T, James JJ, Maysuria M, Mitton JD, Oliveri P, Osborn JL, Peng T, Ratcliffe AL, Webster PJ, Davidson EH, Hood L, Dimitrov K. 2008. Direct multiplexed measurement of gene expression with color-coded probe pairs. Nat Biotechnol 26:317–25.

36. Scoggin K, Gupta J, Lynch R, Nagarajan A, Aminian M, Peterson A, Adams LG, Kirby M, Threadgill DW, Andrews-Polymenis HL. 2022. Elucidating mechanisms of tolerance to *Salmonella* Typhimurium across long-term infections using the Collaborative Cross. mBio 13:e0112022.

37. Scoggin K, Lynch R, Gupta J, Nagarajan A, Sheffield M, Elsaadi A, Bowden C, Aminian M, Peterson A, Adams LG, Kirby M, Threadgill DW, Andrews-Polymenis HL. 2022. Genetic background influences survival of infections with *Salmonella enterica* serovar Typhimurium in the Collaborative Cross. PLoS Genet 18:e1010075.

38. Iraqi FA, Athamni H, Dorman A, Salymah Y, Tomlinson I, Nashif A, Shusterman A, Weiss E, Houri-Haddad Y, Mott R, Soller M. 2014. Heritability and coefficient of genetic variation analyses of phenotypic traits provide strong basis for high-resolution QTL mapping in the Collaborative Cross mouse genetic reference population. Mamm Genome 25:109–19.

39. Broman KW, Gatti DM, Simecek P, Furlotte NA, Prins P, Sen Ś, Yandell BS, Churchill GA. 2019. R/qtl2: Software for mapping quantitative trait loci with high-dimensional data and multiparent populations. Genetics 211:495–502.

40. Hampton BK, Plante KS, Whitmore AC, Linnertz CL, Madden EA, Noll KE, Boyson SP, Parotti B, Xenakis JG, Bell TA, Hock P, Shaw GD, de Villena FP, Ferris MT, Heise MT. 2022. Forward genetic screen of homeostatic antibody levels in the Collaborative Cross identifies MBD1 as a novel regulator of B cell homeostasis. PLoS Genet 18:e1010548.

41. Blake JA, Baldarelli R, Kadin JA, Richardson JE, Smith CL, Bult CJ. 2021. Mouse Genome Database (MGD): Knowledgebase for mouse-human comparative biology. Nucleic Acids Res 49:D981–d987.

42. Perez G, Barber GP, Benet-Pages A, Casper J, Clawson H, Diekhans M, Fischer C, Gonzalez JN, Hinrichs AS, Lee CM, Nassar LR, Raney BJ, Speir ML, van Baren MJ, Vaske CJ, Haussler D, Kent WJ, Haeussler M. 2025. The UCSC Genome Browser database: 2025 update. Nucleic Acids Res 53:D1243–d1249.

43. Bates D, Mächler M, Bolker B, Walker S. 2015. Fitting Linear Mixed-Effects Models Using lme4. Journal of Statistical Software 67:1–48.

44. Efron B, Tibshirani R. 1994. An Introduction to the Bootstrap, Chapman and Hall/CRC, New York.

45. Viechtbauer W. 2010. Conducting Meta-Analyses in R with the metafor Package. J Stat Softw 36:1–48.

46. https://nanostring.com/products/ncounter-analysis-system/ncounter-analysis-solutions/nsolver-data-analysis-support/. Accessed December 2, 2025.

47. Wobbrock JO, Findlater L, Gergle D, Higgins JJ. 2011. The aligned rank transform for nonparametric factorial analyses using only anova procedures, abstr Proceedings of the SIGCHI Conference on Human Factors in Computing Systems, May 7–12, Vancouver, BC, Canada, Association for Computing Machinery.

48. Elkin LA, Kay M, Higgins JJ, Wobbrock JO. 2021. An aligned rank transform procedure for multifactor contrast Tests, abstr The 34th annual ACM symposium on user interface software and technology, October 10–14, Virtual Event, USA, Association for Computing Machinery.

49. Kay M, Elkin L, Higgins JJ, Wobbrock JO. 2025. mjskay/ARTool: ARTool 0.11.2, vv0.11.2. Zenodo, 10.5281/zenodo.15192343.

50. Chandereng T, Gitter A. 2020. Lag penalized weighted correlation for time series clustering. BMC Bioinform 21:21.

51. Tibshirani R, Walther G, Hastie T. 2001. Estimating the number of clusters in a data set via the Gap Statistic. J. R. Stat. Soc., B: Stat. Methodol. 63:411–423.

52. Moorhead AM, Jung JY, Smirnov A, Kaufer S, Scidmore MA. 2010. Multiple host proteins that function in phosphatidylinositol-4-phosphate metabolism are recruited to the chlamydial inclusion. Infect Immun 78:1990–2007.

53. Clayton EL, Minogue S, Waugh MG. 2013. Mammalian phosphatidylinositol 4-kinases as modulators of membrane trafficking and lipid signaling networks. Prog Lipid Res 52:294–304.

54. Robertson DK, Gu L, Rowe RK, Beatty WL. 2009. Inclusion biogenesis and reactivation of persistent *Chlamydia trachomatis* requires host cell sphingolipid biosynthesis. PLOS Pathogens 5:e1000664.

55. Finethy R, Coers J. 2016. Sensing the enemy, containing the threat: cell-autonomous immunity to Chlamydia trachomatis. FEMS Microbiol Rev 40:875–893.

56. Haldar AK, Piro AS, Finethy R, Espenschied ST, Brown HE, Giebel AM, Frickel EM, Nelson DE, Coers J. 2016. *Chlamydia trachomatis* is resistant to inclusion ubiquitination and associated host defense in gamma interferon-primed human epithelial cells. mBio 7: e01417–16.

57. Hausman JM, Kenny S, Iyer S, Babar A, Qiu J, Fu J, Luo ZQ, Das C. 2020. The two deubiquitinating enzymes from *Chlamydia trachomatis* have distinct ubiquitin recognition properties. Biochemistry 59:1604–1617.

58. Yang Y, Lei W, Zhao L, Wen Y, Li Z. 2022. Insights into mitochondrial dynamics in chlamydial infection. Front Cell Infect Microbiol 12:835181.

59. Mi Y, Gurumurthy RK, Zadora PK, Meyer TF, Chumduri C. 2018. *Chlamydia trachomatis* inhibits homologous recombination repair of DNA Breaks by interfering with PP2A Signaling. mBio 9: e01465–18.

60. Johnson KA, Tan M, Sutterlin C. 2009. Centrosome abnormalities during a *Chlamydia trachomatis* infection are caused by dysregulation of the normal duplication pathway. Cell Microbiol 11:1064–73.

61. Rajarathnam K, Schnoor M, Richardson RM, Rajagopal S. 2019. How do chemokines navigate neutrophils to the target site: Dissecting the structural mechanisms and signaling pathways. Cell Signal 54:69–80.

62. Lijek RS, Helble JD, Olive AJ, Seiger KW, Starnbach MN. 2018. Pathology after *Chlamydia trachomatis* infection is driven by nonprotective immune cells that are distinct from protective populations. Proc Natl Acad Sci U S A 115:2216–2221.

63. Scurlock AM, Frazer LC, Andrews CW, Jr., O’Connell CM, Foote IP, Bailey SL, Chandra-Kuntal K, Kolls JK, Darville T. 2011. Interleukin-17 contributes to generation of Th1 immunity and neutrophil recruitment during *Chlamydia muridarum* genital tract infection but is not required for macrophage influx or normal resolution of infection. Infect Immun 79:1349–62.

64. Frazer LC, O’Connell CM, Andrews CW, Jr., Zurenski MA, Darville T. 2011. Enhanced neutrophil longevity and recruitment contribute to the severity of oviduct pathology during *Chlamydia muridarum* infection. Infect Immun 79:4029–41.

65. Olive AJ, Gondek DC, Starnbach MN. 2011. CXCR3 and CCR5 are both required for T cell-mediated protection against *C. trachomatis* infection in the murine genital mucosa. Mucosal Immunol 4:208–16.

66. King M, Poya H, Rao J, Natarajan S, Butch AW, Aziz N, Kok S, Chang MH, Lyons JM, Ault K, Kelly KA. 2009. CXCL13 expression in *Chlamydia trachomatis* infection of the female reproductive tract. Drugs Today (Barc) 45 Suppl B:125–34.

67. Marchesi F, Martin AP, Thirunarayanan N, Devany E, Mayer L, Grisotto MG, Furtado GC, Lira SA. 2009. CXCL13 expression in the gut promotes accumulation of IL-22-producing lymphoid tissue-inducer cells, and formation of isolated lymphoid follicles. Mucosal Immunol 2:486–494.

68. Taylor BD, Zheng X, Darville T, Zhong W, Konganti K, Abiodun-Ojo O, Ness RB, O’Connell CM, Haggerty CL. 2017. Whole-exome sequencing to identify novel biological pathways associated with infertility after pelvic inflammatory disease. Sex Transm Dis 44:35–41.

69. Masola V, Bellin G, Gambaro G, Onisto M. 2018. Heparanase: a multitasking protein involved in extracellular matrix (ECM) remodeling and intracellular events. Cells 7:236.

70. Finethy R, Jorgensen I, Haldar AK, de Zoete MR, Strowig T, Flavell RA, Yamamoto M, Nagarajan UM, Miao EA, Coers J. 2015. Guanylate binding proteins enable rapid activation of canonical and noncanonical inflammasomes in Chlamydia-infected macrophages. Infect Immun 83:4740–9.

71. Haldar AK, Piro AS, Pilla DM, Yamamoto M, Coers J. 2014. The E2-like conjugation enzyme Atg3 promotes binding of IRG and Gbp proteins to Chlamydia- and Toxoplasma-containing vacuoles and host resistance. PLoS One 9:e86684.

72. Wang PH, Liu YF, Tsai HT, Tee YT, Lin LY, Hsieh YH, Yang SF. 2010. Elevated plasma osteopontin level is associated with pelvic inflammatory disease. Reprod Sci17:1052–8.

73. De Filippis A, Buommino E, Domenico MD, Feola A, Brunetti-Pierri R, Rizzo A. 2017. *Chlamydia trachomatis* induces an upregulation of molecular biomarkers podoplanin, Wilms’ tumour gene 1, osteopontin and inflammatory cytokines in human mesothelial cells. Microbiol (Reading) 163:654–663.

74. Kiviat NB, Wolner-Hanssen P, Eschenbach DA, Wasserheit JN, Paavonen JA, Bell TA, Critchlow CW, Stamm WE, Moore DE, Holmes KK. 1990. Endometrial histopathology in patients with culture-proved upper genital tract infection and laparoscopically diagnosed acute salpingitis. Am J Surg Pathol 14:167–75.

75. Zheng X, O’Connell CM, Zhong W, Nagarajan UM, Tripathy M, Lee D, Russell AN, Wiesenfeld H, Hillier S, Darville T. 2018. Discovery of blood transcriptional endotypes in women with pelvic inflammatory disease. J Immunol 200:2941–2956.

76. Zheng X, O’Connell CM, Zhong W, Poston TB, Wiesenfeld HC, Hillier SL, Trent M, Gaydos C, Tseng G, Taylor BD, Darville T. 2018. Gene expression signatures can aid diagnosis of sexually transmitted infection-induced endometritis in women. Front Cell Infect Microbiol 8:307.

77. Dockterman J, Coers J. 2021. Immunopathogenesis of genital Chlamydia infection: insights from mouse models. Pathogens and Disease 79: ftab012.

78. Gyorke CE, Kollipara A, Allen Jt, Zhang Y, Ezzell JA, Darville T, Montgomery SA, Nagarajan UM. 2020. IL-1α is essential for oviduct pathology during genital chlamydial infection in mice. J Immunol 205:3037–3049.

79. Darville T, O’Neill JM, Andrews CW, Jr., Nagarajan UM, Stahl L, Ojcius DM. 2003. Toll-like receptor-2, but not toll-like receptor-4, is essential for development of oviduct pathology in chlamydial genital tract infection. J Immunol 171:6187–6197.

80. Anyalechi GE, Hong J, Danavall DC, Martin DL, Gwyn SE, Horner PJ, Raphael BH, Kirkcaldy RD, Kersh EN, Bernstein KT. 2021. High plasmid gene protein 3 (Pgp3) *Chlamydia trachomatis* seropositivity, pelvic inflammatory disease, and infertility among women, National Health and Nutrition Examination Survey, United States, 2013-2016. Clin Infect Dis 73:1507–1516.

81. Wiesenfeld HC. 2017. Screening for *Chlamydia trachomatis* Infections in Women. N Engl J Med 376:765–773.

82. Tuttle AH, Philip VM, Chesler EJ, Mogil JS. 2018. Comparing phenotypic variation between inbred and outbred mice. Nat Methods 15:994–996.

83. Bailey RL, Natividad-Sancho A, Fowler A, Peeling RW, Mabey DC, Whittle HC, Jepson AP. 2009. Host genetic contribution to the cellular immune response to *Chlamydia trachomatis*: Heritability estimate from a Gambian twin study. Drugs Today 45 Suppl B:45–50.

84. Poston TB, Lee DE, Darville T, Zhong W, Dong L, O’Connell CM, Wiesenfeld HC, Hillier SL, Sempowski GD, Zheng X. 2019. Cervical cytokines associated with *Chlamydia trachomatis* susceptibility and protection. J Infect Dis 220:330–339.

85. Nagarajan UM, Prantner D, Sikes JD, Andrews CW, Jr., Goodwin AM, Nagarajan S, Darville T. 2008. Type I interferon signaling exacerbates *Chlamydia muridarum* genital infection in a murine model. Infect Immun 76:4642–8.

86. Chen R, Ma L, Jiang C, Zhang S. 2022. Expression and potential role of CCL4 in CD8+T cells in NSCLC. Clin Transl Oncol 24:2420–2431.

87. Repeke CE, Ferreira SB, Jr., Claudino M, Silveira EM, de Assis GF, Avila-Campos MJ, Silva JS, Garlet GP. 2010. Evidences of the cooperative role of the chemokines CCL3, CCL4 and CCL5 and its receptors CCR1+ and CCR5+ in RANKL+ cell migration throughout experimental periodontitis in mice. Bone 46:1122–30.

88. Blanpain C, Migeotte I, Lee B, Vakili J, Doranz BJ, Govaerts C, Vassart G, Doms RW, Parmentier M. 1999. CCR5 binds multiple CC-chemokines: MCP-3 acts as a natural antagonist. Blood 94:1899–905.

89. Garlanda C, Di Ceglie I, Jaillon S. 2025. IL-1 family cytokines in inflammation and immunity. Cell Mol Immunol 22:1345–1362.

90. Zhong W, Dong L, Poston TB, Darville T, Spracklen CN, Wu D, Mohlke KL, Li Y, Li Q, Zheng X. 2020. Inferring regulatory networks from mixed observational data using directed acyclic graphs. Front Genet 11:8.

