## Supplementary material for "Genetic diversity of Collaborative Cross mice enables the establishment of a novel *Chlamydia muridarum* female genital tract infection model": All supplemental figure and tables

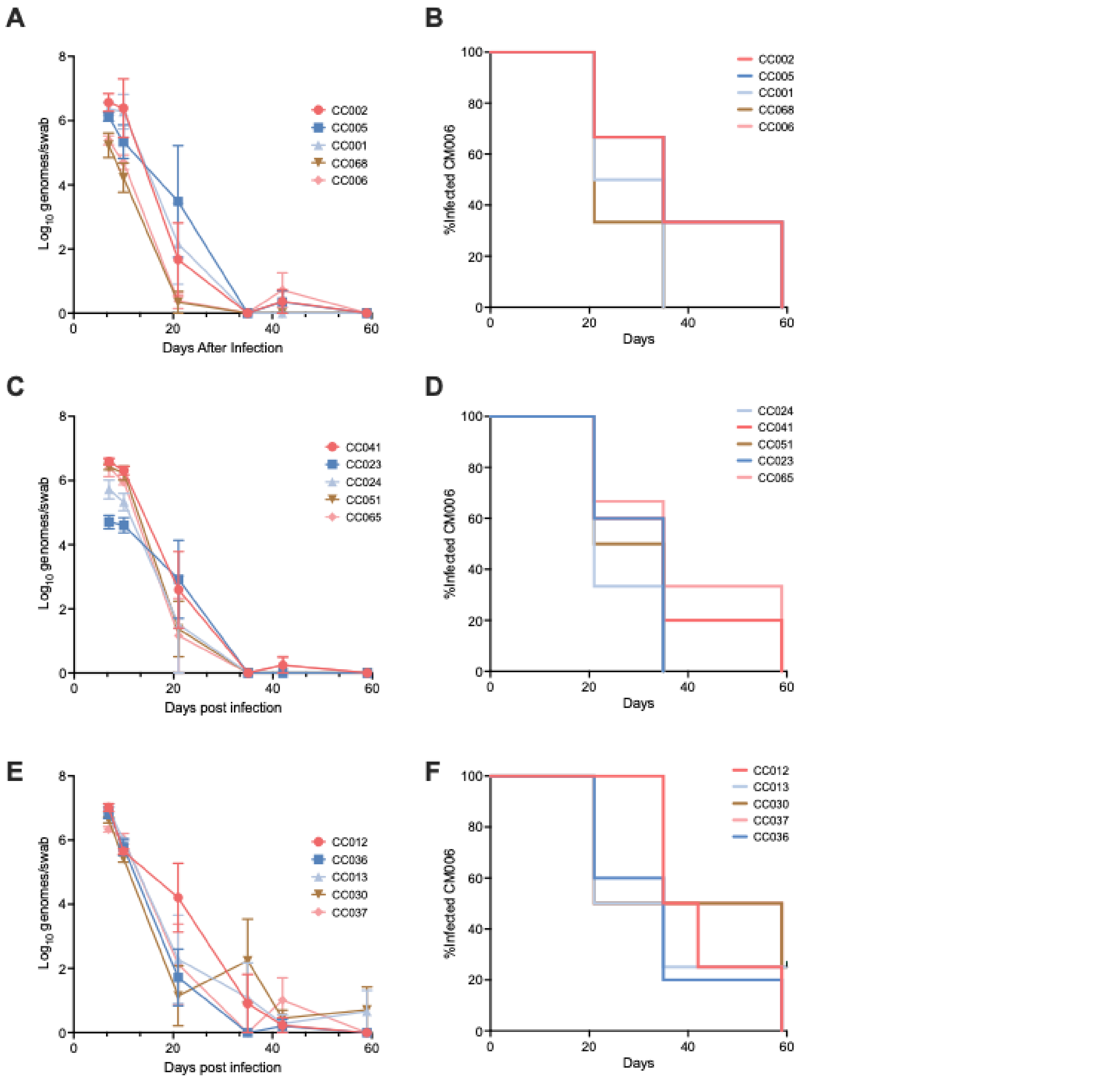

**Supplementary Figure 1. Chlamydial burden and infection clearance across all CC mouse strains inoculated with *C. muridarum* CM006.** Lower genital tract shedding was quantified by qPCR targeting the 23S rRNA locus. Panels A, C, and E show mean bacterial burden ± SEM for each CC strain. Infection clearance (B, D, F) was defined as the first day on which chlamydial genomic DNA fell below the limit of detection ($<8.3 \times 10^1$ genomes/swab). All qPCR reactions were performed in triplicate.

Panels represent groupings of CC strains by cohort size:
(A, B) CC strains with 3–6 mice/group: CC001 (n=4), CC002 (n=3), CC005 (n=3), CC006 (n=6), CC068 (n=3).
(C, D) CC strains with 3–5 mice/group: CC023 (n=5), CC024 (n=3), CC041 (n=5), CC051 (n=4), CC065 (n=3).
(E, F) CC strains with 4–5 mice/group: CC012 (n=4), CC013 (n=4), CC030 (n=4), CC036 (n=5), CC037 (n=4).

**Supplementary Table 1. Host and chlamydial gene probes used for cervical transcriptional profiling in this study**

| Official Symbol | Accession | Alias/Previous Symbol | Target Sequence |
|---|---|---|---|
| ***Selected host targets*** | | | |
| Ccl11 | NM_011330.3 | eotaxin, Scya11 | CAACCTCCTCTCTTGACACTAACCCAGAGCCTAAGAACTGCTTGATTCCTTCTCTTTCCTAAGACGTGCTCTGAGGGAATATCAGCACCAGTCGCCCAAG |
| Ccl19 | NM_011888.2 | exodus-3, Scya19, CKb11 | CTTCTGCCAAGAACAAAGGCAACAGCACCAGAAGGAGCCCTGTGTCTTGAGTAAAGAGATGTGAATCACTCTGGCCCAGGAAACCAAGGACCAGAAGAGA |
| Ccl2 | NM_011333.3 | monocyte chemotactic protein, MCP-1, monocyte chemoattractant protein-1, Sigje, MCAF, SMC-CF, HC11, MCP1, Scya2 | TCTTCAGCACCTTTGAATGTGAAGTTGACCCGTAAATCTGAAGCTAATGCATCCACTACCTTTTCCACAACCACCTCAAGCACTTCTGTAGGAGTGACCA |
| Ccl3 | NM_011337.1 | MIP-1 alpha, MIP1-alpha, MIP1-(a), macrophage inflammatory protein-1alpha, MIP-1alpha, Mip1a, CCL3, G0S19-1, LD78alpha, Scya3 | TCTGTCACCTGCTCAACATCATGAAGGTCTCCACCACTGCCCTTGCTGTTCTTCTCTGTACCATGACACTCTGCAACCAAGTCTTCTCAGCGCCATATGG |
| Ccl4 | NM_013652.1 | MIP-1 beta, MIP-1B, Mip1b, Act-2, AT744.1, Scya4 | TTCTCAGCACCAATGGGCTCTGACCCTCCCACTTCCTGCTGTTTCTCTTACACCTCCCGGCAGCTTCACAGAAGCTTTGTGATGGATTACTATGAGACCA |
| Ccl5 | NM_013653.1 | MuRantes, Scya5, SISd, TCP228, RANTES | CCTCGTGCCCACGTCAAGGAGTATTTCTACACCAGCAGCAAGTGCTCCAATCTTGCAGTCGTGTTTGTCACTCGAAGGAACCGCCAAGTGTGTGCCAACC |
| Cd4 | NM_013488.2 | L3T4, Ly-4 | AAGAGGTGTCCGTACAAAAGTCCACCAAAGACCTCAAGCTCCAGCTGAAGGAAACGCTCCCACTCACCCTCAAGATACCCCAGGTCTCGCTTCAGTTTGC |
| Cd40lg | NM_011616.2 | CD154, gp39, Ly-62, Cd40L, IMD3, HIGM1, T-BAM, Tnfsf5 | AGCAGTGGATCTGAGAGAATCTTACTCAAGGCGGCAAATACCCACAGTTCCTCCCAGCTTTGCGAGCAGCAGTCTGTTCACTTGGGCGGAGTGTTTGAAT |
| Csf1 | NM_001113530.1 | M-CSF, CSF-1, colony-stimulating factor-1, Csfm | TCCAGCTGCTGGAGAAGATCAAGAACTTCTTTAATGAAACAAAGAATCTCCTTGAAAAGGACTGGAACATTTTTACCAAGAACTGCAACAACAGCTTTGC |
| Csf2 | NM_009969.4 | Gm-CSf, Csfgm, MGI-IGM, GMCSF | AAGTCGTCTCTAACGAGTTCTCCTTCAAGAAGCTAACATGTGTGCAGACCCGCCTGAAGATATTCGAGCAGGGTCTACGGGGCAATTTCACCAAACTCAA |

| | | | |
|---|---|---|---|
| Cxcl1 | NM_008176.1 | N51, Mgsa, Fsp, Gro1, Scyb1, KC/GRO-alpha, KC | TGCTAGTAGAAGGGTGTTGTGCGAAAAGAAGTGCAGAGAGATAGAGTTTAGTATTATGTTTTGTATGTATTAGGGTGAGGACATGTGTGGGAGGCTGTGT |
| Cxcl10 | NM_021274.1 | CRG-2, IP-10, Ifi10, mob-1, INP10, C7, gIP-10, Scyb10, IP10 | AGGACGGTCCGCTGCAACTGCATCCATATCGATGACGGGCCAGTGAGAATGAGGGCCATAGGGAAGCTTGAAATCATCCCTGCGAGCCTATCCTGCCCAC |
| Cxcl12 | NM_021704.3 | Sdf1a, Sdf1b, SDF-1, PBSF/SDF-1, Scyb12, Sdf1, TPAR1, TLSF-a, TLSF-b, PBSF | CTGAAAATCCTCAACACTCCAAACTGTGCCCTTCAGATTGTTGCACGGCTGAAGAACAACAACAGACAAGTGTGCATTGACCCGAAATTAAAGTGGATCC |
| Cxcl9 | NM_008599.2 | Mig, CMK, crg-10, Scyb9 | TAGAACTCAGCTCTGCCATGAAGTCCGCTGTTCTTTTCCTCTTGGGCATCATCTTCCTGGAGCAGTGTGGAGTTCGAGGAACCCTAGTGATAAGGAATGC |
| Ifna1 | NM_010502.2 | RP23-360M23.1, Ifa1 | CTGCAAGGCTGTCTGATGCAGCAGGTGGGGGTGCAGGAATTTCCCCTGACCCAGGAAGATGCCCTGCTGGCTGTGAGGAAATACTTCCACAGGATCACTG |
| Ifna2 | NM_010503.2 | RP23-139P14.4, Ifa2 | TAACCTCAGGAACAAGAGGGCCTTGAAGGTCCTGGCACAGATGAGGAGGCTCCCCTTTCTCTCCTGCCTGAAGGACAGGCAGGACTTTGGATTCCCCCTG |
| Ifnb1 | NM_010510.1 | interferon beta 1, fibroblast, Ifb, IFNB, IFN-beta | GATGAACTCCACCAGCAGACAGTGTTTCTGAAGACAGTACTAGAGGAAAAGCAAGAGGAAAGATTGACGTGGGAGATGTCCTCAACTGCTCTCCACTTGA |
| Ifng | NM_008337.1 | IFN-gamma, Ifg | CTAGCTCTGAGACAATGAACGCTACACACTGCATCTTGGCTTTGCAGCTCTTCCTCATGGCTGTTTCTGGCTGTTACTGCCACGGCACAGTCATTGAAAG |
| Il10 | NM_010548.1 | cytokine synthesis inhibitory factor, IL-10 | GGGCCCTTTGCTATGGTGTCCTTTCAATTGCTCTCATCCCTGAGTTCAGAGCTCCTAAGAGAGTTGTGAAGAAACTCATGGGTCTTGGGAAGAGAAACCA |
| Il12a | NM_008351.1 | IL-12p35, p35 | TCATGAAGACATCACACGGGACCAAACCAGCACATTGAAGACCTGTTTACCACTGGAACTACACAAGAACGAGAGTTGCCTGGCTACTAGAGAGACTTCT |
| Il12b | NM_008352.1 | Il-12p40, IL-12 p40, Il-12b, IL-23 subunit p40 | TCGTAGAGAAGACATCTACCGAAGTCCAATGCAAAGGCGGGAATGTCTGCGTGCAAGCTCAGGATCGCTATTACAATTCCTCGTGCAGCAAGTGGGCATG |
| Il13 | NM_008355.2 | Il-13 | AGCTACACAAAGCAACTGTTTCGCCACGGCCCCTTCTAATGAGGAGAGACCATCCCTGGGCATCTCAGCTGTGGACTCATTTTCCTTTCTCACATCAGAC |

| | | | |
|---|---|---|---|
| Il15 | NM_008357.2 | | CTTGCAAACAGCACTCTGTCTTCTAACAAGAATGTAGCAGAATCTGGCTGCAAGGAATGTGAGGAGCTGGAGGAGAAAACCTTCACAGAGTTTTTGCAAA |
| Il17a | NM_010552.3 | Ctla-8, Ctla8, Il17 | ACCTCAAAGTCTTTAACTCCCTTGGCGCAAAAGTGAGCTCCAGAAGGCCCTCAGACTACCTCAACCGTTCCACGTCACCCTGGACTCTCCACCGCAATGA |
| Il18 | NM_008360.1 | Il-18, Igif | ACTTTGGCCGACTTCACTGTACAACCGCAGTAATACGGAATATAAATGACCAAGTTCTCTTCGTTGACAAAAGACAGCCTGTGTTCGAGGATATGACTGA |
| Il1a | NM_010554.4 | Il-1a | ACCTCTGAAACGTCAAAGATGTCCAACTTCACCTTCAAGGAGAGCCGGGTGACAGTATCAGCAACGTCAAGCAACGGGAAGATTCTGAAGAAGAGACGGC |
| Il1b | NM_008361.3 | IL-1B, IL-1beta | GTTGATTCAAGGGGACATTAGGCAGCACTCTCTAGAACAGAACCTAGCTGTCAACGTGTGGGGGATGAATTGGTCATAGCCCGCACTGAGGTCTTTCATT |
| Il1rn | NM_031167.5 | IL-1ra | CAACCAGCTCATTGCTGGGTACTTACAAGGACCAAATATCAAACTAGAAGAAAAGATAGACATGGTGCCTATTGACCTTCATAGTGTGTTCTTGGGCATC |
| Il2 | NM_008366.2 | IL-2 | GCAACTGTGGTGGACTTTCTGAGGAGATGGATAGCCTTCTGTCAAAGCATCATCTCAACAAGCCCTCAATAACTATGTACCTCCTGCTTACAACACATAA |
| Il21 | NM_021782.2 | | ATGGCCTGGGGGATGGTTTTGATCTAAGGAAAAAGGTGTCTGTACCTCACAGTGCCTTTAAAACAAGCAGAGATCCCGTGTACCGCCCTAAGATAGCACA |
| Il22 | NM_016971.1 | mCG_123110, IL-22, IL-22a, ILTIFa, Iltif | AGAAGAATGTCAGAAGGCTGAAGGAGACAGTGAAAAAGCTTGGAGAGAGTGGAGAGATCAAGGCGATTGGGGAACTGGACCTGCTGTTTATGTCTCTGAG |
| Il23a | NM_031252.1 | p19, IL-23, IL-23p19 | CAAGGACAACAGCCAGTTCTGCTTGCAAAGGATCCGCCAAGGTCTGGCTTTTTATAAGCACCTGCTTGACTCTGACATCTTCAAAGGGGAGCCTGCTCTA |
| Il27 | NM_145636.1 | p28, IL-27, Il30, IL-27p28 | GCTATGTCCACAGCTTTGCTGAATCTCGATTGCCAGGAGTGAACCTGGACCTCCTGCCCCTGGGATACCATCTTCCCAATGTTTCCCTGACTTTCCAGGC |
| Il4 | NM_021283.1 | Il-4 | TGCTTGAAGAAGAACTCTAGTGTTCTCATGGAGCTGCAGAGACTCTTTCGGGCTTTTCGATGCCTGGATTCATCGATAAGCTGCACCATGAATGAGTCCA |

| | | | |
|---|---|---|---|
| Il6 | NM_031168.1 | Il-6 | CTCTCTGCAAGAGACTTCCATCCAGTTGCCTTCTTGGGACTGATGCTGGTGACAACCACGGCCTTCCCTACTTCACAAGTCCGGAGAGGAGACTTCACAG |
| Il7 | NM_008371.2 | Il-7, hlb368 | AAACATTCATTGGTGAACCACTGGGGGAGTGGAACTGTCCTGTTTTAGACTGGAGATACTGGAGGGCTCACGGTGATGGATAATGCTCTTGAAAACAAGA |
| Ltb | NM_008518.2 | lymphotoxin beta, Tnfc, LTbeta, Tnfsf3, p33 | ATCAGGGACGTCGGGTTGAGAAGATCATTGGCTCAGGAGCACAGGCTCAGAAAAGACTGGATGACAGCAAACCGTCGTGCATCTTGCCCTCACCCTCTAG |
| Tnf | NM_013693.1 | tumor necrosis factor-alpha, TNF-alpha, TNFalpha, TNF alpha, DIF, Tnfsf1a, Tnfa | TTCCTGAGTTCTGCAAAGGGAGAGTGGTCAGGTTGCCTCTGTCTCAGAATGAGGCTGGATAAGATCTCAGGCCTTCCTACCTTCAGACCTTTCCAGACTC |
| Stat1 | NM_009283.3 | | ACGCTGGGAACAGAACTAATGAGGGGCCTCTCATTGTCACCGAAGAACTTCACTCTCTTAGCTTTGAAACCCAGTTGTGCCAGCCAGGCTTGGTGATTGA |
| Stat3 | NM_213659.2 | Aprf | GGGGTCACTTTCACTTGGGTGGAAAAGGACATCAGTGGCAAGACCCAGATCCAGTCTGTAGAGCCATACACCAAGCAGCAGCTGAACAACATGTCATTTG |
| Ccr5 | NM_009917.5 | CD195, Cmkbr5 | GGAGCAGGGAGAACGAGTCTTTTATCAGGGCCGGGAAATATGCACAAAGAGACTTGAGGCAGGTGCCATGACCCATATGCAAAGGGACGGACACAGGGCC |
| Stat4 | NM_011487.4 | | TAGAGACCAGCTCATTACCTGTCGTGATGATTTCTAATGTCAGCCAACTACCTAATGCATGGGCATCCATCATTTGGTACAATGTATCAACTAACGACTC |
| Tlr2 | NM_011905.2 | Ly105 | GCAGGCGGTCACTGGCAGGAGATGTGTCCGCAATCATAGTTTCTGATGGTGAAGGTTGGACGGCAGTCTCTGCGACCTAGAAGTGGAAAAGATGTCGTTC |
| Cxcr3 | NM_009910.2 | Cxcr3, Cmkar3, Cd183 | GTTGTATGGGGTCTCTGTCTGCTCTTTGCCCTCCCAGATTTCATCTACCTATCAGCCAACTACGATCAGCGCCTCAATGCCACCCATTGCCAGTACAACT |
| Eomes | NM_010136.2 | Tbr2 | ACTGAAAAGGTCGTTCAAGGTGCTGGATTGATTCATTTATGGGAAACGAGAAATGTTCAGAAAAAGCAGGCTATGAAGAACGAGTGCCCGGTGCTATTAA |
| Tbx21 | NM_019507.1 | TBT1, Tblym, T-bet | CACTAAGCAAGGACGGCGAATGTTCCCATTCCTGTCCTTCACCGTGGCTGGGCTGGAGCCCACAAGCCATTACAGGATGTTTGTGGATGTGGTCTTGGTG |

| | | | |
|---|---|---|---|
| Cd3e | NM_007648.4 | T3e, CD3, CD3epsilon | TGAGTACTGTGTGGAGGTGGACCTGACAGCAGTAGCCATAATCATCATTGTTGACATCTGTATCACTCTGGGCTTGCTGATGGTCATTTATTACTGGAGC |
| Il16 | NM_010551.3 | | TGCGAGACAAAGCTGTTGGATGAAAAGGCCAGTAAGCTTTACTCCATCAGCAGCCAGCTATCATCTGCTGTCATGAAATCCCTGCTGTGCCTTCCATCTT |
| Cxcl13 | NM_018866.2 | BLC, BCA-1, ANGIE2, Scyb13 | CTGAGAGGGAATGCTCAAGCTCCGTTGCATACCCAACCCACATCCTTGTTCCTTAAGAAAGGCTATTTGAGAACAGGCATTTAGTGACAACCCACTTCAG |
| Foxp3 | NM_054039.1 | JM2, scurfin | CCAGCTCCCGGCAACTTCTCCTGACTCTGCCTTCAGACGAGACTTGGAAGACAGTCACATCTCAGCAGCTCCTCTGCCGTTATCCAGCCTGCCTCTGACA |
| Gata3 | NM_008091.3 | Gata-3 | CATGCGTGAGGAGTCTCCAAGTGTGCGAAGAGTTCCTCCGACCCCTTCTACTTGCGTTTTTCGCAGGAGCAGTATCATGAAGCCCGAAAGCGACAGATCT |
| ***Chlamydial targets*** | | | |
| plasmid | CT_pGP8_as.1 | CT_pGP8 anti-sense | CACAGCGGTTGCTCGAAGCACGTGCGGGGTTATCTTAAAAGGGATTGCAGCTTGTAGTCCTGCTTGAGAG |
| omcA | CT_444.1 | omcA | CCATGCGCACCTATCCAATGTTCACCTTGTGAATCTAAGAAGAAAGACGTAGACGGTGGTTGCAACTCTTGTAACGGGTATGTCCCAGCTTGCAAACCTT |
| 23S | CT_r02.1 | CT_23SrRNA | AACGGTTAGTAGGCAAATCCGCTAACATAAGATCAGGTCGCGATCAAGGGGAATCTTCGGGGGAACCGATGGTGTGGAGCGAGGCTTTCAAGAAATAATT |
| ***Internal Reference Targets*** | | | |
| Cltc | NM_001003908.1 | 3110065L21Rik, CHC, R74732 | GAGTCTCAGCCAGTGAAAATGTTTGATCGTCATTCTAGCCTCGCAGGATGCCAGATCATCAATTACCGTACAGATGCAAAGCAGAAATGGTTGCTTCTCA |
| Gapdh | NM_001001303.1 | Gapd | AGGTTGTCTCCTGCGACTTCAACAGCAACTCCCACTCTTCCACCTTCGATGCCGGGGCTGGCATTGCTCTCAATGACAACTTTGTCAAGCTCATTTCCTG |

| | | | |
|---|---|---|---|
| Gusb | NM_010368.1 | g, Gut, Gur, asd, adipose storage deficiency, Gus-r, Gus-s, Gus-t, Gus-u, Gus | AATACGTGGTCGGAGAGCTCATCTGGAATTTCGCCGACTTCATGACGAACCAGTCACCGCTGAGAGTAATCGGAAACAAGAAGGGGATCTTCACTCGCCA |
| Hprt | NM_013556.2 | Hprt1 | TGCTGAGGCGGCGAGGGAGAGCGTTGGGCTTACCTCACTGCTTTCCGGAGCGGTAGCACCTCCTCCGCCGGCTTCCTCCTCAGACCGCTTTTTGCCGCGA |
| Pgk1 | NM_008828.2 | Pgk-1 | CCGGCATTCTGCACGCTTCAAAAGCGCACGTCTGCCGCGCTGTTCTCCTCTTCCTCATCTCCGGGCCTTTCGACCTCACGGTGTTGCCAAAATGTCGCTT |
| Tubb5 | NM_011655.4 | AA408537, AI596182, B130022C14Rik, M(beta)5 | ATTGGAAGTGTCTTCCCTGTATTGGTTCTCCTTTCTCGGAGAGATGGGGGTTGGGGGTGCGGCAAGGTCTTGGTCTTGGTCTCTGAACACTCCCAATTCC |

**Supplementary Table 2. Protein-coding genes located within ±10 Mb of the Day 7 bacterial-load peak QTL on chromosome 16 (peak~7.1Mb)**

| Chr | type | start | stop | gene_id | mgi_type | gene_symbol | description |
|---|---|---|---|---|---|---|---|
| 16 | protein_coding | 5124665 | 7229391 | ENSMUSG00000008658 | MGI:1926224 | Rbfox1 | RNA binding protein, fox-1 homolog (C. elegans) 1 [Source:MGI Symbol;Acc:MGI:1926224] |
| 16 | protein_coding | 8227139 | 8243000 | ENSMUSG00000022715 | MGI:1921970 | Tmem114 | transmembrane protein 114 [Source:MGI Symbol;Acc:MGI:1921970] |
| 16 | protein_coding | 8288652 | 8308548 | ENSMUSG00000039345 | MGI:2384301 | Mettl22 | methyltransferase 22, Kin17 lysine [Source:MGI Symbol;Acc:MGI:2384301] |
| 16 | protein_coding | 8331293 | 8439432 | ENSMUSG00000057880 | MGI:2443582 | Abat | 4-aminobutyrate aminotransferase [Source:MGI Symbol;Acc:MGI:2443582] |
| 16 | protein_coding | 8451093 | 8455576 | ENSMUSG00000043140 | MGI:1913940 | Tmem186 | transmembrane protein 186 [Source:MGI Symbol;Acc:MGI:1913940] |
| 16 | protein_coding | 8455538 | 8480331 | ENSMUSG00000022711 | MGI:1859214 | Pmm2 | phosphomannomutase 2 [Source:MGI Symbol;Acc:MGI:1859214] |
| 16 | protein_coding | 8476444 | 8490019 | ENSMUSG00000008393 | MGI:1196368 | Carhsp1 | calcium regulated heat stable protein 1 [Source:MGI Symbol;Acc:MGI:1196368] |
| 16 | protein_coding | 8498634 | 8501775 | ENSMUSG00000107252 | MGI:3643507 | Litafd | LITAF domain containing [Source:MGI Symbol;Acc:MGI:3643507] |
| 16 | protein_coding | 8507459 | 8610172 | ENSMUSG00000022710 | MGI:2182061 | Usp7 | ubiquitin specific peptidase 7 [Source:MGI Symbol;Acc:MGI:2182061] |
| 16 | protein_coding | 8647964 | 8676786 | ENSMUSG00000022507 | MGI:1916303 | Hapstr1 | HUWE1 associated protein modifying stress responses [Source:MGI Symbol;Acc:MGI:1916303] |
| 16 | protein_coding | 9385762 | 9813424 | ENSMUSG00000059003 | MGI:95820 | Grin2a | glutamate receptor, ionotropic, NMDA2A (epsilon 1) [Source:MGI Symbol;Acc:MGI:95820] |
| 16 | protein_coding | 9988090 | 9992775 | ENSMUSG00000039209 | MGI:1915422 | Rpl39l | ribosomal protein L39-like [Source:MGI Symbol;Acc:MGI:1915422] |
| 16 | protein_coding | 10010576 | 10069342 | ENSMUSG00000039200 | MGI:1922579 | Atf7ip2 | activating transcription factor 7 interacting protein 2 [Source:MGI Symbol;Acc:MGI:1922579] |
| 16 | protein_coding | 10099613 | 10131832 | ENSMUSG00000022505 | MGI:1098726 | Emp2 | epithelial membrane protein 2 [Source:MGI Symbol;Acc:MGI:1098726] |
| 16 | protein_coding | 10175812 | 10213354 | ENSMUSG00000039179 | MGI:1917676 | Tekt5 | tektin 5 [Source:MGI Symbol;Acc:MGI:1917676] |

| | | | | | | | |
|---|---|---|---|---|---|---|---|
| 16 | protein_coding | 10229812 | 10242292 | ENSMUSG00000022503 | MGI:1347073 | Nubp1 | nucleotide binding protein 1 [Source:MGI Symbol;Acc:MGI:1347073] |
| 16 | protein_coding | 10238421 | 10265226 | ENSMUSG00000050908 | MGI:3665441 | Tvp23a | trans-golgi network vesicle protein 23A [Source:MGI Symbol;Acc:MGI:3665441] |
| 16 | protein_coding | 10297923 | 10346282 | ENSMUSG00000022504 | MGI:108445 | Ciita | class II transactivator [Source:MGI Symbol;Acc:MGI:108445] |
| 16 | protein_coding | 10320395 | 10360918 | ENSMUSG00000038055 | MGI:1926236 | Dexi | dexamethasone-induced transcript [Source:MGI Symbol;Acc:MGI:1926236] |
| 16 | protein_coding | 10363203 | 10562742 | ENSMUSG00000068663 | MGI:1921624 | Clec16a | C-type lectin domain family 16, member A [Source:MGI Symbol;Acc:MGI:1921624] |
| 16 | protein_coding | 10600104 | 10603400 | ENSMUSG00000038037 | MGI:1354910 | Socs1 | suppressor of cytokine signaling 1 [Source:MGI Symbol;Acc:MGI:1354910] |
| 16 | protein_coding | 10605800 | 10606524 | ENSMUSG00000043050 | MGI:98785 | Tnp2 | transition protein 2 [Source:MGI Symbol;Acc:MGI:98785] |
| 16 | protein_coding | 10608369 | 10608778 | ENSMUSG00000050058 | MGI:106601 | Prm3 | protamine 3 [Source:MGI Symbol;Acc:MGI:106601] |
| 16 | protein_coding | 10609244 | 10613998 | ENSMUSG00000038015 | MGI:97766 | Prm2 | protamine 2 [Source:MGI Symbol;Acc:MGI:97766] |
| 16 | protein_coding | 10614190 | 10623703 | ENSMUSG00000022501 | MGI:97765 | Prm1 | protamine 1 [Source:MGI Symbol;Acc:MGI:97765] |
| 16 | protein_coding | 10652923 | 10710830 | ENSMUSG00000037991 | MGI:2685383 | Rmi2 | RecQ mediated genome instability 2 [Source:MGI Symbol;Acc:MGI:2685383] |
| 16 | protein_coding | 10777139 | 10884021 | ENSMUSG00000022500 | MGI:1929512 | Litaf | LPS-induced TN factor [Source:MGI Symbol;Acc:MGI:1929512] |
| 16 | protein_coding | 10878809 | 10892849 | ENSMUSG00000037972 | MGI:1276549 | Snn | stannin [Source:MGI Symbol;Acc:MGI:1276549] |
| 16 | protein_coding | 10892775 | 10952514 | ENSMUSG00000022498 | MGI:1923620 | Txndc11 | thioredoxin domain containing 11 [Source:MGI Symbol;Acc:MGI:1923620] |
| 16 | protein_coding | 10954456 | 10994257 | ENSMUSG00000037965 | MGI:2445044 | Zc3h7a | zinc finger CCCH type containing 7 A [Source:MGI Symbol;Acc:MGI:2445044] |
| 16 | protein_coding | 11010834 | 11021195 | ENSMUSG00000005846 | MGI:1913659 | Rsl1d1 | ribosomal L1 domain containing 1 [Source:MGI Symbol;Acc:MGI:1913659] |
| 16 | protein_coding | 11037156 | 11072189 | ENSMUSG00000062203 | MGI:1316728 | Gspt1 | G1 to S phase transition 1 [Source:MGI Symbol;Acc:MGI:1316728] |

| | | | | | | | |
|---|---|---|---|---|---|---|---|
| 16 | protein_coding | 11131676 | 11137938 | ENSMUSG00000022496 | MGI:1343050 | Tnfrsf17 | tumor necrosis factor receptor superfamily, member 17 [Source:MGI Symbol;Acc:MGI:1343050] |
| 16 | protein_coding | 11140772 | 11573336 | ENSMUSG00000071669 | MGI:1921728 | Snx29 | sorting nexin 29 [Source:MGI Symbol;Acc:MGI:1921728] |
| 16 | protein_coding | 11621585 | 11727309 | ENSMUSG00000065979 | MGI:2443300 | Cpped1 | calcineurin-like phosphoesterase domain containing 1 [Source:MGI Symbol;Acc:MGI:2443300] |
| 16 | protein_coding | 11801977 | 12088766 | ENSMUSG00000022494 | MGI:1919805 | Shisa9 | shisa family member 9 [Source:MGI Symbol;Acc:MGI:1919805] |
| 16 | protein_coding | 12927548 | 12968481 | ENSMUSG00000022545 | MGI:1354163 | Ercc4 | excision repair cross-complementing rodent repair deficiency, complementation group 4 [Source:MGI Symbol;Acc:MGI:1354163] |
| 16 | protein_coding | 13074345 | 13235393 | ENSMUSG00000009569 | MGI:3050795 | Mrtfb | myocardin related transcription factor B [Source:MGI Symbol;Acc:MGI:3050795] |
| 16 | protein_coding | 13355824 | 13486034 | ENSMUSG00000022685 | MGI:1921358 | Parn | poly(A)-specific ribonuclease (deadenylation nuclease) [Source:MGI Symbol;Acc:MGI:1921358] |
| 16 | protein_coding | 13489722 | 13521476 | ENSMUSG00000022684 | MGI:1914368 | Bfar | bifunctional apoptosis regulator [Source:MGI Symbol;Acc:MGI:1914368] |
| 16 | protein_coding | 13532921 | 13548847 | ENSMUSG00000022683 | MGI:1347522 | Pla2g10 | phospholipase A2, group X [Source:MGI Symbol;Acc:MGI:1347522] |
| 16 | protein_coding | 13572430 | 13576401 | ENSMUSG00000118559 | MGI:3045309 | A930007A09 Rik | RIKEN cDNA A930007A09 gene [Source:MGI Symbol;Acc:MGI:3045309] |
| 16 | protein_coding | 13598572 | 13632703 | ENSMUSG00000022682 | MGI:1925255 | Rrn3 | RRN3 RNA polymerase I transcription factor homolog (yeast) [Source:MGI Symbol;Acc:MGI:1925255] |
| 16 | protein_coding | 13636709 | 13653315 | ENSMUSG00000022681 | MGI:108471 | Ntan1 | N-terminal Asn amidase [Source:MGI Symbol;Acc:MGI:108471] |
| 16 | protein_coding | 13651012 | 13720995 | ENSMUSG00000022680 | MGI:1920909 | Pdxdc1 | pyridoxal-dependent decarboxylase domain containing 1 [Source:MGI Symbol;Acc:MGI:1920909] |
| 16 | protein_coding | 13721025 | 13767483 | ENSMUSG00000022679 | MGI:2135951 | Mpv17l | Mpv17 transgene, kidney disease mutant-like [Source:MGI Symbol;Acc:MGI:2135951] |

| | | | | | | | |
|---|---|---|---|---|---|---|---|
| 16 | protein_coding | 13799563 | 13804752 | ENSMUSG00000065968 | MGI:1921732 | Ifitm7 | interferon induced transmembrane protein 7 [Source:MGI Symbol;Acc:MGI:1921732] |
| 16 | protein_coding | 13804468 | 13919364 | ENSMUSG00000044117 | MGI:1914504 | Bmerb1 | bMERB domain containing 1 [Source:MGI Symbol;Acc:MGI:1914504] |
| 16 | protein_coding | 13927037 | 13981215 | ENSMUSG00000060657 | MGI:2444505 | Marf1 | meiosis regulator and mRNA stability 1 [Source:MGI Symbol;Acc:MGI:2444505] |
| 16 | protein_coding | 13981139 | 14010792 | ENSMUSG00000022678 | MGI:1914453 | Nde1 | nudE neurodevelopment protein 1 [Source:MGI Symbol;Acc:MGI:1914453] |
| 16 | protein_coding | 14012399 | 14109236 | ENSMUSG00000018830 | MGI:102643 | Myh11 | myosin, heavy polypeptide 11, smooth muscle [Source:MGI Symbol;Acc:MGI:102643] |
| 16 | protein_coding | 14117108 | 14135269 | ENSMUSG00000022677 | MGI:1913336 | Cep20 | centrosomal protein 20 [Source:MGI Symbol;Acc:MGI:1913336] |
| 16 | protein_coding | 14179422 | 14293601 | ENSMUSG00000023088 | MGI:102676 | Abcc1 | ATP-binding cassette, sub-family C member 1 [Source:MGI Symbol;Acc:MGI:102676] |
| 16 | protein_coding | 14523716 | 14527249 | ENSMUSG00000022676 | MGI:1096393 | Snai2 | snail family zinc finger 2 [Source:MGI Symbol;Acc:MGI:1096393] |
| 16 | protein_coding | 14724470 | 14799235 | ENSMUSG00000068617 | MGI:1914043 | Clxn | calaxin [Source:MGI Symbol;Acc:MGI:1914043] |
| 16 | protein_coding | 15369941 | 15413637 | ENSMUSG00000022674 | MGI:1917870 | Ube2v2 | ubiquitin-conjugating enzyme E2 variant 2 [Source:MGI Symbol;Acc:MGI:1917870] |
| 16 | protein_coding | 15441761 | 15455264 | ENSMUSG00000022673 | MGI:103199 | Mcm4 | minichromosome maintenance complex component 4 [Source:MGI Symbol;Acc:MGI:103199] |
| 16 | protein_coding | 15455730 | 15660099 | ENSMUSG00000022672 | MGI:104779 | Prkdc | protein kinase, DNA activated, catalytic polypeptide [Source:MGI Symbol;Acc:MGI:104779] |
| 16 | protein_coding | 15666305 | 15681233 | ENSMUSG00000022671 | MGI:1922845 | Mzt2 | mitotic spindle organizing protein 2 [Source:MGI Symbol;Acc:MGI:1922845] |
| 16 | protein_coding | 15705150 | 15708895 | ENSMUSG00000071637 | MGI:103573 | Cebpd | CCAAT/enhancer binding protein delta [Source:MGI Symbol;Acc:MGI:103573] |
| 16 | protein_coding | 15707088 | 15964715 | ENSMUSG00000041974 | MGI:1924834 | Spidr | scaffolding protein involved in DNA repair [Source:MGI Symbol;Acc:MGI:1924834] |
| 16 | protein_coding | 16031182 | 16090576 | ENSMUSG00000041957 | MGI:1914701 | Pkp2 | plakophilin 2 [Source:MGI Symbol;Acc:MGI:1914701] |

| | | | | | | | |
|---|---|---|---|---|---|---|---|
| 16 | protein_coding | 16120829 | 16127504 | ENSMUSG00000022792 | MGI:1917370 | Yars2 | tyrosyl-tRNA synthetase 2 (mitochondrial) [Source:MGI Symbol;Acc:MGI:1917370] |
| 16 | protein_coding | 16130094 | 16176823 | ENSMUSG00000022789 | MGI:1921256 | Dnm1l | dynamin 1-like [Source:MGI Symbol;Acc:MGI:1921256] |
| 16 | protein_coding | 16234781 | 16418413 | ENSMUSG00000022788 | MGI:2183747 | Fgd4 | FYVE, RhoGEF and PH domain containing 4 [Source:MGI Symbol;Acc:MGI:2183747] |
| 16 | protein_coding | 16490092 | 16494269 | ENSMUSG00000048101 | MGI:109316 | Or7a40 | olfactory receptor family 7 subfamily A member 40 [Source:MGI Symbol;Acc:MGI:109316] |
| 16 | protein_coding | 16570880 | 16647320 | ENSMUSG00000022783 | MGI:1354388 | Spag6l | sperm associated antigen 6-like [Source:MGI Symbol;Acc:MGI:1354388] |
| 16 | protein_coding | 16678535 | 16681849 | ENSMUSG00000075370 | MGI:96529 | Igll1 | immunoglobulin lambda-like polypeptide 1 [Source:MGI Symbol;Acc:MGI:96529] |
| 16 | protein_coding | 16686267 | 16688707 | ENSMUSG00000059305 | MGI:98936 | Vpreb1a | V-set pre-B cell surrogate light chain 1A [Source:MGI Symbol;Acc:MGI:98936] |
| 16 | protein_coding | 16688600 | 16710854 | ENSMUSG00000022779 | MGI:1333803 | Top3b | topoisomerase (DNA) III beta [Source:MGI Symbol;Acc:MGI:1333803] |
| 16 | protein_coding | 16714333 | 16745228 | ENSMUSG00000026181 | MGI:1918464 | Ppm1f | protein phosphatase 1F (PP2C domain containing) [Source:MGI Symbol;Acc:MGI:1918464] |
| 16 | protein_coding | 16801246 | 16865317 | ENSMUSG00000063358 | MGI:1346858 | Mapk1 | mitogen-activated protein kinase 1 [Source:MGI Symbol;Acc:MGI:1346858] |
| 16 | protein_coding | 16887560 | 16904909 | ENSMUSG00000022773 | MGI:1913303 | Ypel1 | yippee like 1 [Source:MGI Symbol;Acc:MGI:1913303] |
| 16 | protein_coding | 16904419 | 16929121 | ENSMUSG00000022771 | MGI:2447857 | Ppil2 | peptidylprolyl isomerase (cyclophilin)-like 2 [Source:MGI Symbol;Acc:MGI:2447857] |
| 16 | protein_coding | 16931262 | 16943031 | ENSMUSG00000049916 | MGI:1917708 | 2610318N02 Rik | RIKEN cDNA 2610318N02 gene [Source:MGI Symbol;Acc:MGI:1917708] |
| 16 | protein_coding | 16948002 | 16950247 | ENSMUSG00000022769 | MGI:2149842 | Sdf2l1 | stromal cell-derived factor 2-like 1 [Source:MGI Symbol;Acc:MGI:2149842] |
| 16 | protein_coding | 16956928 | 16965093 | ENSMUSG00000022768 | MGI:1924122 | Ccdc116 | coiled-coil domain containing 116 [Source:MGI Symbol;Acc:MGI:1924122] |
| 16 | protein_coding | 16962485 | 16978565 | ENSMUSG00000041774 | MGI:1916351 | Ydjc | YdjC homolog (bacterial) [Source:MGI Symbol;Acc:MGI:1916351] |
| 16 | protein_coding | 16969877 | 17020513 | ENSMUSG00000038965 | MGI:109240 | Ube2l3 | ubiquitin-conjugating enzyme E2L 3 [Source:MGI Symbol;Acc:MGI:109240] |

| | | | | | | | |
|---|---|---|---|---|---|---|---|
| 16 | protein_coding | 17026467 | 17031846 | ENSMUSG00000071636 | MGI:2685449 | Rimbp3 | RIMS binding protein 3 [Source:MGI Symbol;Acc:MGI:2685449] |
| 16 | protein_coding | 17051436 | 17081294 | ENSMUSG00000050240 | MGI:1929869 | Hic2 | hypermethylated in cancer 2 [Source:MGI Symbol;Acc:MGI:1929869] |
| 16 | protein_coding | 17093941 | 17101093 | ENSMUSG00000055692 | MGI:107238 | Tmem191 | transmembrane protein 191 [Source:MGI Symbol;Acc:MGI:107238] |
| 16 | protein_coding | 17098215 | 17224178 | ENSMUSG00000041720 | MGI:2448506 | Pi4ka | phosphatidylinositol 4-kinase alpha [Source:MGI Symbol;Acc:MGI:2448506] |
| 16 | protein_coding | 17149235 | 17161439 | ENSMUSG00000022766 | MGI:96051 | Serpind1 | serine (or cysteine) peptidase inhibitor, clade D, member 1 [Source:MGI Symbol;Acc:MGI:96051] |

**Supplementary Table 3. All annotated genes within the 95% Bayesian credible interval for the Day 7 bacterial-load QTL on chromosome 16**

| Chr | type | start | stop | gene_id | mgi_type | gene_symbol | description |
|---|---|---|---|---|---|---|---|
| 16 | IG_C_gene | 18880368 | 18880821 | ENSMUSG00000105906 | MGI:99546 | Iglc1 | immunoglobulin lambda constant 1 [Source:MGI Symbol;Acc:MGI:99546] |
| 16 | IG_C_gene | 18883980 | 18884431 | ENSMUSG00000105547 | MGI:99886 | Iglc3 | immunoglobulin lambda constant 3 [Source:MGI Symbol;Acc:MGI:99886] |
| 16 | IG_C_gene | 19013749 | 19014062 | ENSMUSG00000106039 | MGI:3642936 | Iglc4 | immunoglobulin lambda constant 4 [Source:MGI Symbol;Acc:MGI:3642936] |
| 16 | IG_C_gene | 19017146 | 19019101 | ENSMUSG00000076937 | MGI:99547 | Iglc2 | immunoglobulin lambda constant 2 [Source:MGI Symbol;Acc:MGI:99547] |
| 16 | IG_J_gene | 18881975 | 18882012 | ENSMUSG00000106668 | MGI:5009822 | Iglj1 | immunoglobulin lambda joining 1 [Source:MGI Symbol;Acc:MGI:5009822] |
| 16 | IG_J_gene | 18885121 | 18885158 | ENSMUSG00000106405 | MGI:3645901 | Iglj3p | immunoglobulin lambda joining 3 pseudogene [Source:MGI Symbol;Acc:MGI:3645901] |
| 16 | IG_J_gene | 18885791 | 18885828 | ENSMUSG00000105231 | MGI:5293407 | Iglj3 | immunoglobulin lambda joining 3 [Source:MGI Symbol;Acc:MGI:5293407] |
| 16 | IG_J_gene | 19015245 | 19015286 | ENSMUSG00000105066 | MGI:3645510 | Iglj4 | immunoglobulin lambda joining 4 [Source:MGI Symbol;Acc:MGI:3645510] |
| 16 | IG_J_gene | 19018948 | 19018985 | ENSMUSG00000104975 | MGI:5293409 | Iglj2 | immunoglobulin lambda joining 2 [Source:MGI Symbol;Acc:MGI:5293409] |
| 16 | IG_V_gene | 18903767 | 18904250 | ENSMUSG00000076934 | MGI:96530 | Iglv1 | immunoglobulin lambda variable 1 [Source:MGI Symbol;Acc:MGI:96530] |
| 16 | IG_V_gene | 19059957 | 19060440 | ENSMUSG00000076939 | MGI:3645509 | Iglv3 | immunoglobulin lambda variable 3 [Source:MGI Symbol;Acc:MGI:3645509] |
| 16 | IG_V_gene | 19079151 | 19079609 | ENSMUSG00000076940 | MGI:99548 | Iglv2 | immunoglobulin lambda variable 2 [Source:MGI Symbol;Acc:MGI:99548] |
| 16 | lncRNA | 6028260 | 6049376 | ENSMUSG00000143715 | MGI:7805634 | Gm61295 | predicted gene, 61295 [Source:MGI Symbol;Acc:MGI:7805634] |

| | | | | | | | |
|---|---|---|---|---|---|---|---|
| 16 | lncRNA | 6163728 | 6166837 | ENSMUSG00000143829 | MGI:7805644 | Gm61302 | predicted gene, 61302 [Source:MGI Symbol;Acc:MGI:7805644] |
| 16 | lncRNA | 6298226 | 6330952 | ENSMUSG00000143626 | MGI:7805606 | Gm61281 | predicted gene, 61281 [Source:MGI Symbol;Acc:MGI:7805606] |
| 16 | lncRNA | 6582243 | 6597040 | ENSMUSG00000143791 | MGI:7805640 | Gm61298 | predicted gene, 61298 [Source:MGI Symbol;Acc:MGI:7805640] |
| 16 | lncRNA | 6836996 | 6842096 | ENSMUSG00000138950 | MGI:7804973 | Gm60964 | predicted gene, 60964 [Source:MGI Symbol;Acc:MGI:7804973] |
| 16 | lncRNA | 6904486 | 6917736 | ENSMUSG00000123065 | MGI:7802569 | Gm59754 | predicted gene, 59754 [Source:MGI Symbol;Acc:MGI:7802569] |
| 16 | lncRNA | 6977986 | 6979694 | ENSMUSG00000123172 | MGI:7802583 | Gm59761 | predicted gene, 59761 [Source:MGI Symbol;Acc:MGI:7802583] |
| 16 | lncRNA | 7028135 | 7036670 | ENSMUSG00000123107 | MGI:7802573 | Gm59756 | predicted gene, 59756 [Source:MGI Symbol;Acc:MGI:7802573] |
| 16 | lncRNA | 7104704 | 7107338 | ENSMUSG00000123137 | MGI:7802577 | Gm59758 | predicted gene, 59758 [Source:MGI Symbol;Acc:MGI:7802577] |
| 16 | lncRNA | 7237187 | 7241286 | ENSMUSG00000128473 | MGI:7803205 | Gm60076 | predicted gene, 60076 [Source:MGI Symbol;Acc:MGI:7803205] |
| 16 | lncRNA | 7302513 | 7303200 | ENSMUSG00000139398 | MGI:7805035 | Gm60995 | predicted gene, 60995 [Source:MGI Symbol;Acc:MGI:7805035] |
| 16 | lncRNA | 7584296 | 7584981 | ENSMUSG00000125667 | MGI:7802783 | Gm59861 | predicted gene, 59861 [Source:MGI Symbol;Acc:MGI:7802783] |
| 16 | lncRNA | 7585726 | 7588456 | ENSMUSG00000124707 | MGI:5589013 | Gm29854 | predicted gene, 29854 [Source:MGI Symbol;Acc:MGI:5589013] |
| 16 | lncRNA | 7630206 | 7631750 | ENSMUSG00000123267 | MGI:7802603 | Gm59771 | predicted gene, 59771 [Source:MGI Symbol;Acc:MGI:7802603] |
| 16 | lncRNA | 7687966 | 7697723 | ENSMUSG00000116450 | MGI:6155235 | Gm49534 | predicted gene, 49534 [Source:MGI Symbol;Acc:MGI:6155235] |
| 16 | lncRNA | 7765480 | 7769537 | ENSMUSG00000139532 | MGI:7805061 | Gm61008 | predicted gene, 61008 [Source:MGI Symbol;Acc:MGI:7805061] |
| 16 | lncRNA | 7886440 | 7891071 | ENSMUSG00000137024 | MGI:7804593 | Gm60774 | predicted gene, 60774 [Source:MGI Symbol;Acc:MGI:7804593] |
| 16 | lncRNA | 8160162 | 8177767 | ENSMUSG00000126210 | MGI:5589095 | Gm29936 | predicted gene, 29936 [Source:MGI Symbol;Acc:MGI:5589095] |
| 16 | lncRNA | 8275922 | 8289483 | ENSMUSG00000128980 | MGI:5589171 | Gm30012 | predicted gene, 30012 [Source:MGI Symbol;Acc:MGI:5589171] |

| | | | | | | | |
|---|---|---|---|---|---|---|---|
| 16 | lncRNA | 8315029 | 8317226 | ENSMUSG00000143008 | MGI:7805511 | Gm61233 | predicted gene, 61233 [Source:MGI Symbol;Acc:MGI:7805511] |
| 16 | lncRNA | 8495965 | 8500615 | ENSMUSG00000125402 | MGI:7802771 | Gm59855 | predicted gene, 59855 [Source:MGI Symbol;Acc:MGI:7802771] |
| 16 | lncRNA | 8538592 | 8545526 | ENSMUSG00000125362 | MGI:7802763 | Gm59851 | predicted gene, 59851 [Source:MGI Symbol;Acc:MGI:7802763] |
| 16 | lncRNA | 8555352 | 8563430 | ENSMUSG00000125319 | MGI:7802759 | Gm59849 | predicted gene, 59849 [Source:MGI Symbol;Acc:MGI:7802759] |
| 16 | lncRNA | 8588468 | 8588618 | ENSMUSG00000125487 | MGI:7802775 | Gm59857 | predicted gene, 59857 [Source:MGI Symbol;Acc:MGI:7802775] |
| 16 | lncRNA | 8599272 | 8603553 | ENSMUSG00000125445 | MGI:7802773 | Gm59856 | predicted gene, 59856 [Source:MGI Symbol;Acc:MGI:7802773] |
| 16 | lncRNA | 8640009 | 8641685 | ENSMUSG00000122939 | MGI:7802553 | Gm59746 | predicted gene, 59746 [Source:MGI Symbol;Acc:MGI:7802553] |
| 16 | lncRNA | 8700834 | 8702196 | ENSMUSG00000125833 | MGI:7802797 | Gm59868 | predicted gene, 59868 [Source:MGI Symbol;Acc:MGI:7802797] |
| 16 | lncRNA | 8762756 | 8787764 | ENSMUSG00000116525 | MGI:5589530 | Gm30371 | predicted gene, 30371 [Source:MGI Symbol;Acc:MGI:5589530] |
| 16 | lncRNA | 8815371 | 8825701 | ENSMUSG00000136575 | MGI:5589603 | Gm30444 | predicted gene, 30444 [Source:MGI Symbol;Acc:MGI:5589603] |
| 16 | lncRNA | 8820898 | 8822122 | ENSMUSG00000136615 | MGI:7804559 | Gm60757 | predicted gene, 60757 [Source:MGI Symbol;Acc:MGI:7804559] |
| 16 | lncRNA | 8827559 | 8834631 | ENSMUSG00000138549 | MGI:7804869 | Gm60912 | predicted gene, 60912 [Source:MGI Symbol;Acc:MGI:7804869] |
| 16 | lncRNA | 8867836 | 8873284 | ENSMUSG00000116426 | MGI:5589726 | Gm30567 | predicted gene, 30567 [Source:MGI Symbol;Acc:MGI:5589726] |
| 16 | lncRNA | 8897260 | 9038550 | ENSMUSG00000116469 | MGI:6155097 | Gm49448 | predicted gene, 49448 [Source:MGI Symbol;Acc:MGI:6155097] |
| 16 | lncRNA | 8981276 | 8990641 | ENSMUSG00000138255 | MGI:7804833 | Gm60894 | predicted gene, 60894 [Source:MGI Symbol;Acc:MGI:7804833] |
| 16 | lncRNA | 9097790 | 9108447 | ENSMUSG00000133742 | MGI:5624303 | Gm41418 | predicted gene, 41418 [Source:MGI Symbol;Acc:MGI:5624303] |
| 16 | lncRNA | 9114599 | 9116518 | ENSMUSG00000137900 | MGI:7804817 | Gm60886 | predicted gene, 60886 [Source:MGI Symbol;Acc:MGI:7804817] |
| 16 | lncRNA | 9116612 | 9119168 | ENSMUSG00000129757 | MGI:7803371 | Gm60159 | predicted gene, 60159 [Source:MGI Symbol;Acc:MGI:7803371] |

| | | | | | | | |
|---|---|---|---|---|---|---|---|
| 16 | lncRNA | 9212100 | 9213004 | ENSMUSG00000142707 | MGI:7805503 | Gm61229 | predicted gene, 61229 [Source:MGI Symbol;Acc:MGI:7805503] |
| 16 | lncRNA | 9815073 | 9820381 | ENSMUSG00000122586 | MGI:5589827 | Gm30668 | predicted gene, 30668 [Source:MGI Symbol;Acc:MGI:5589827] |
| 16 | lncRNA | 10004157 | 10045388 | ENSMUSG00000115943 | MGI:6155110 | Gm49455 | predicted gene, 49455 [Source:MGI Symbol;Acc:MGI:6155110] |
| 16 | lncRNA | 10141631 | 10169214 | ENSMUSG00000116270 | MGI:2686446 | Gm1600 | predicted gene 1600 [Source:MGI Symbol;Acc:MGI:2686446] |
| 16 | lncRNA | 10167472 | 10171739 | ENSMUSG00000129485 | MGI:5589883 | Gm30724 | predicted gene, 30724 [Source:MGI Symbol;Acc:MGI:5589883] |
| 16 | lncRNA | 10200623 | 10201046 | ENSMUSG00000143676 | MGI:7805632 | Gm61294 | predicted gene, 61294 [Source:MGI Symbol;Acc:MGI:7805632] |
| 16 | lncRNA | 10265251 | 10265526 | ENSMUSG00000138914 | MGI:7804971 | Gm60963 | predicted gene, 60963 [Source:MGI Symbol;Acc:MGI:7804971] |
| 16 | lncRNA | 10267228 | 10269692 | ENSMUSG00000142680 | MGI:7805499 | Gm61227 | predicted gene, 61227 [Source:MGI Symbol;Acc:MGI:7805499] |
| 16 | lncRNA | 10466165 | 10476113 | ENSMUSG00000129964 | MGI:6366384 | Gm52264 | predicted gene, 52264 [Source:MGI Symbol;Acc:MGI:6366384] |
| 16 | lncRNA | 10487112 | 10501456 | ENSMUSG00000087384 | MGI:3783007 | Gm15558 | predicted gene 15558 [Source:MGI Symbol;Acc:MGI:3783007] |
| 16 | lncRNA | 10510191 | 10515959 | ENSMUSG00000130911 | MGI:7803645 | Gm60296 | predicted gene, 60296 [Source:MGI Symbol;Acc:MGI:7803645] |
| 16 | lncRNA | 10566395 | 10582213 | ENSMUSG00000140467 | MGI:7805191 | Gm61073 | predicted gene, 61073 [Source:MGI Symbol;Acc:MGI:7805191] |
| 16 | lncRNA | 10589796 | 10590068 | ENSMUSG00000140592 | MGI:7805209 | Gm61082 | predicted gene, 61082 [Source:MGI Symbol;Acc:MGI:7805209] |
| 16 | lncRNA | 10590215 | 10590598 | ENSMUSG00000140555 | MGI:7805201 | Gm61078 | predicted gene, 61078 [Source:MGI Symbol;Acc:MGI:7805201] |
| 16 | lncRNA | 10602518 | 10630590 | ENSMUSG00000116038 | MGI:5826200 | Gm46563 | predicted gene, 46563 [Source:MGI Symbol;Acc:MGI:5826200] |
| 16 | lncRNA | 10603508 | 10604089 | ENSMUSG00000140525 | MGI:7805199 | Gm61077 | predicted gene, 61077 [Source:MGI Symbol;Acc:MGI:7805199] |
| 16 | lncRNA | 10645765 | 10691449 | ENSMUSG00000096921 | MGI:5477316 | Gm26822 | predicted gene, 26822 [Source:MGI Symbol;Acc:MGI:5477316] |
| 16 | lncRNA | 10729865 | 10777126 | ENSMUSG00000095369 | MGI:5434023 | Gm21859 | predicted gene, 21859 [Source:MGI Symbol;Acc:MGI:5434023] |

| | | | | | | | |
|---|---|---|---|---|---|---|---|
| 16 | lncRNA | 10816946 | 10821356 | ENSMUSG00000125386 | MGI:7802769 | Gm59854 | predicted gene, 59854 [Source:MGI Symbol;Acc:MGI:7802769] |
| 16 | lncRNA | 10849820 | 10854284 | ENSMUSG00000115992 | MGI:3642210 | Gm10832 | predicted gene 10832 [Source:MGI Symbol;Acc:MGI:3642210] |
| 16 | lncRNA | 10864620 | 10868803 | ENSMUSG00000138912 | MGI:7805754 | Gm61360 | predicted gene, 61360 [Source:MGI Symbol;Acc:MGI:7805754] |
| 16 | lncRNA | 10924374 | 10925211 | ENSMUSG00000138960 | MGI:7804977 | Gm60966 | predicted gene, 60966 [Source:MGI Symbol;Acc:MGI:7804977] |
| 16 | lncRNA | 10952801 | 10968472 | ENSMUSG00000141422 | MGI:7805295 | Gm61125 | predicted gene, 61125 [Source:MGI Symbol;Acc:MGI:7805295] |
| 16 | lncRNA | 10994094 | 11010156 | ENSMUSG00000096959 | MGI:1914984 | 4930509G22Rik | RIKEN cDNA 4930509G22 gene [Source:MGI Symbol;Acc:MGI:1914984] |
| 16 | lncRNA | 11020710 | 11082342 | ENSMUSG00000097537 | MGI:1917168 | 2610020C07Rik | RIKEN cDNA 2610020C07 gene [Source:MGI Symbol;Acc:MGI:1917168] |
| 16 | lncRNA | 11021207 | 11033249 | ENSMUSG00000133226 | MGI:7804043 | Gm60496 | predicted gene, 60496 [Source:MGI Symbol;Acc:MGI:7804043] |
| 16 | lncRNA | 11071866 | 11074500 | ENSMUSG00000120347 | MGI:7337108 | Gm56597 | predicted gene, 56597 [Source:MGI Symbol;Acc:MGI:7337108] |
| 16 | lncRNA | 11449930 | 11451039 | ENSMUSG00000127042 | MGI:7802969 | Gm59955 | predicted gene, 59955 [Source:MGI Symbol;Acc:MGI:7802969] |
| 16 | lncRNA | 11506033 | 11509721 | ENSMUSG00000085732 | MGI:3782456 | Gm4279 | predicted gene 4279 [Source:MGI Symbol;Acc:MGI:3782456] |
| 16 | lncRNA | 11596255 | 11599365 | ENSMUSG00000115936 | MGI:1921011 | 4833415N18Rik | RIKEN cDNA 4833415N18 gene [Source:MGI Symbol;Acc:MGI:1921011] |
| 16 | lncRNA | 11719214 | 11748464 | ENSMUSG00000055015 | MGI:3642444 | Gm9961 | predicted gene 9961 [Source:MGI Symbol;Acc:MGI:3642444] |
| 16 | lncRNA | 11796526 | 11803089 | ENSMUSG00000139264 | MGI:5591437 | Gm32278 | predicted gene, 32278 [Source:MGI Symbol;Acc:MGI:5591437] |
| 16 | lncRNA | 12120848 | 12126717 | ENSMUSG00000143211 | MGI:7805533 | Gm61244 | predicted gene, 61244 [Source:MGI Symbol;Acc:MGI:7805533] |
| 16 | lncRNA | 12128412 | 12135411 | ENSMUSG00000143401 | MGI:7805582 | Gm61269 | predicted gene, 61269 [Source:MGI Symbol;Acc:MGI:7805582] |

| | | | | | | | |
|---|---|---|---|---|---|---|---|
| 16 | lncRNA | 12380552 | 12393444 | ENSMUSG00000123409 | MGI:7802611 | Gm59775 | predicted gene, 59775 [Source:MGI Symbol;Acc:MGI:7802611] |
| 16 | lncRNA | 12381586 | 12400760 | ENSMUSG00000123373 | | | novel transcript |
| 16 | lncRNA | 12446948 | 12763828 | ENSMUSG00000116520 | MGI:1926040 | 4930414F18Rik | RIKEN cDNA 4930414F18 gene [Source:MGI Symbol;Acc:MGI:1926040] |
| 16 | lncRNA | 12467885 | 12490637 | ENSMUSG00000140920 | MGI:7805227 | Gm61091 | predicted gene, 61091 [Source:MGI Symbol;Acc:MGI:7805227] |
| 16 | lncRNA | 12522232 | 12530015 | ENSMUSG00000116197 | MGI:5621504 | Gm38619 | predicted gene, 38619 [Source:MGI Symbol;Acc:MGI:5621504] |
| 16 | lncRNA | 12618172 | 12679236 | ENSMUSG00000101565 | MGI:1922743 | 1700003L19Rik | RIKEN cDNA 1700003L19 gene [Source:MGI Symbol;Acc:MGI:1922743] |
| 16 | lncRNA | 12761874 | 12783117 | ENSMUSG00000140400 | MGI:7805165 | Gm61060 | predicted gene, 61060 [Source:MGI Symbol;Acc:MGI:7805165] |
| 16 | lncRNA | 12835898 | 12858295 | ENSMUSG00000139816 | MGI:7805105 | Gm61030 | predicted gene, 61030 [Source:MGI Symbol;Acc:MGI:7805105] |
| 16 | lncRNA | 12893934 | 12903277 | ENSMUSG00000138678 | MGI:7804887 | Gm60921 | predicted gene, 60921 [Source:MGI Symbol;Acc:MGI:7804887] |
| 16 | lncRNA | 12924490 | 12927725 | ENSMUSG00000133609 | MGI:7804069 | Gm60509 | predicted gene, 60509 [Source:MGI Symbol;Acc:MGI:7804069] |
| 16 | lncRNA | 12945386 | 12946851 | ENSMUSG00000133642 | MGI:7804071 | Gm60510 | predicted gene, 60510 [Source:MGI Symbol;Acc:MGI:7804071] |
| 16 | lncRNA | 13007358 | 13007728 | ENSMUSG00000120723 | MGI:7337150 | Gm56618 | predicted gene, 56618 [Source:MGI Symbol;Acc:MGI:7337150] |
| 16 | lncRNA | 13172238 | 13286862 | ENSMUSG00000087526 | MGI:3783180 | Gm15738 | predicted gene 15738 [Source:MGI Symbol;Acc:MGI:3783180] |
| 16 | lncRNA | 13255098 | 13257252 | ENSMUSG00000138767 | MGI:5624306 | Gm41421 | predicted gene, 41421 [Source:MGI Symbol;Acc:MGI:5624306] |
| 16 | lncRNA | 13265280 | 13292725 | ENSMUSG00000109857 | MGI:6435685 | Gm53058 | predicted gene 53058 [Source:MGI Symbol;Acc:MGI:6435685] |
| 16 | lncRNA | 13331995 | 13338971 | ENSMUSG00000099411 | MGI:1917350 | 2310015D24Rik | RIKEN cDNA 2310015D24 gene [Source:MGI Symbol;Acc:MGI:1917350] |
| 16 | lncRNA | 13357121 | 13359618 | ENSMUSG00000129391 | MGI:7803309 | Gm60128 | predicted gene, 60128 [Source:MGI Symbol;Acc:MGI:7803309] |

| | | | | | | | |
|---|---|---|---|---|---|---|---|
| 16 | lncRNA | 13489793 | 13496252 | ENSMUSG00000121810 | MGI:1913848 | Pphln1-ps1 | periphilin 1, pseudogene 1 [Source:MGI Symbol;Acc:MGI:1913848] |
| 16 | lncRNA | 13547660 | 13557951 | ENSMUSG00000086550 | MGI:1923934 | Pla2g10os | phospholipase A2, group X, opposite strand [Source:MGI Symbol;Acc:MGI:1923934] |
| 16 | lncRNA | 13555685 | 13557150 | ENSMUSG00000115907 | MGI:6155072 | Gm49432 | predicted gene, 49432 [Source:MGI Symbol;Acc:MGI:6155072] |
| 16 | lncRNA | 13597524 | 13598492 | ENSMUSG00000133370 | MGI:7804061 | Gm60505 | predicted gene, 60505 [Source:MGI Symbol;Acc:MGI:7804061] |
| 16 | lncRNA | 13735512 | 13747906 | ENSMUSG00000135269 | MGI:7804413 | Gm60684 | predicted gene, 60684 [Source:MGI Symbol;Acc:MGI:7804413] |
| 16 | lncRNA | 13977162 | 13978745 | ENSMUSG00000132903 | MGI:7804023 | Gm60486 | predicted gene, 60486 [Source:MGI Symbol;Acc:MGI:7804023] |
| 16 | lncRNA | 14129705 | 14131878 | ENSMUSG00000085612 | MGI:3801810 | Gm15868 | predicted gene 15868 [Source:MGI Symbol;Acc:MGI:3801810] |
| 16 | lncRNA | 14135026 | 14136102 | ENSMUSG00000134352 | MGI:7804215 | Gm60584 | predicted gene, 60584 [Source:MGI Symbol;Acc:MGI:7804215] |
| 16 | lncRNA | 14282780 | 14285186 | ENSMUSG00000127938 | MGI:7803143 | Gm60045 | predicted gene, 60045 [Source:MGI Symbol;Acc:MGI:7803143] |
| 16 | lncRNA | 14305705 | 14320267 | ENSMUSG00000137348 | MGI:7804727 | Gm60841 | predicted gene, 60841 [Source:MGI Symbol;Acc:MGI:7804727] |
| 16 | lncRNA | 14380181 | 14451972 | ENSMUSG00000075395 | MGI:3584518 | A630010A05Rik | RIKEN cDNA A630010A05 gene [Source:MGI Symbol;Acc:MGI:3584518] |
| 16 | lncRNA | 14522512 | 14523294 | ENSMUSG00000140603 | MGI:7805213 | Gm61084 | predicted gene, 61084 [Source:MGI Symbol;Acc:MGI:7805213] |
| 16 | lncRNA | 14554607 | 14575347 | ENSMUSG00000138555 | MGI:7804871 | Gm60913 | predicted gene, 60913 [Source:MGI Symbol;Acc:MGI:7804871] |
| 16 | lncRNA | 14655394 | 14686705 | ENSMUSG00000129731 | MGI:7803367 | Gm60157 | predicted gene, 60157 [Source:MGI Symbol;Acc:MGI:7803367] |
| 16 | lncRNA | 14710178 | 14810477 | ENSMUSG00000116324 | MGI:5592855 | Gm33696 | predicted gene, 33696 [Source:MGI Symbol;Acc:MGI:5592855] |
| 16 | lncRNA | 14753434 | 14754793 | ENSMUSG00000134967 | MGI:7804365 | Gm60660 | predicted gene, 60660 [Source:MGI Symbol;Acc:MGI:7804365] |
| 16 | lncRNA | 14871190 | 14873987 | ENSMUSG00000122977 | MGI:7802561 | Gm59750 | predicted gene, 59750 [Source:MGI Symbol;Acc:MGI:7802561] |

| | | | | | | | |
|---|---|---|---|---|---|---|---|
| 16 | lncRNA | 14919773 | 14931959 | ENSMUSG00000129831 | MGI:5624308 | Gm41423 | predicted gene, 41423 [Source:MGI Symbol;Acc:MGI:5624308] |
| 16 | lncRNA | 15420808 | 15422665 | ENSMUSG00000132084 | MGI:7803895 | Gm60422 | predicted gene, 60422 [Source:MGI Symbol;Acc:MGI:7803895] |
| 16 | lncRNA | 15422303 | 15422514 | ENSMUSG00000132119 | MGI:7803901 | Gm60425 | predicted gene, 60425 [Source:MGI Symbol;Acc:MGI:7803901] |
| 16 | lncRNA | 15681222 | 15695451 | ENSMUSG00000143526 | MGI:5624310 | Gm41425 | predicted gene, 41425 [Source:MGI Symbol;Acc:MGI:5624310] |
| 16 | lncRNA | 15702802 | 15705084 | ENSMUSG00000120071 | MGI:7337110 | Gm56598 | predicted gene, 56598 [Source:MGI Symbol;Acc:MGI:7337110] |
| 16 | lncRNA | 15967184 | 16037806 | ENSMUSG00000133200 | MGI:7804041 | Gm60495 | predicted gene, 60495 [Source:MGI Symbol;Acc:MGI:7804041] |
| 16 | lncRNA | 16167067 | 16176875 | ENSMUSG00000115923 | MGI:6155216 | Gm49521 | predicted gene, 49521 [Source:MGI Symbol;Acc:MGI:6155216] |
| 16 | lncRNA | 16182848 | 16187925 | ENSMUSG00000126709 | MGI:7805696 | Gm61329 | predicted gene, 61329 [Source:MGI Symbol;Acc:MGI:7805696] |
| 16 | lncRNA | 16502799 | 16536017 | ENSMUSG00000116096 | MGI:1921316 | 4933404G15Rik | RIKEN cDNA 4933404G15 gene [Source:MGI Symbol;Acc:MGI:1921316] |
| 16 | lncRNA | 16536064 | 16549298 | ENSMUSG00000133884 | MGI:5826207 | Gm46570 | predicted gene, 46570 [Source:MGI Symbol;Acc:MGI:5826207] |
| 16 | lncRNA | 16547849 | 16550171 | ENSMUSG00000134003 | MGI:5593499 | Gm34340 | predicted gene, 34340 [Source:MGI Symbol;Acc:MGI:5593499] |
| 16 | lncRNA | 16710097 | 16715212 | ENSMUSG00000133917 | MGI:7804155 | Gm60552 | predicted gene, 60552 [Source:MGI Symbol;Acc:MGI:7804155] |
| 16 | lncRNA | 16727142 | 16730903 | ENSMUSG00000116823 | MGI:6215001 | Gm49595 | predicted gene, 49595 [Source:MGI Symbol;Acc:MGI:6215001] |
| 16 | lncRNA | 16865352 | 16866982 | ENSMUSG00000084792 | MGI:1920642 | 1700056N10Rik | RIKEN cDNA 1700056N10 gene [Source:MGI Symbol;Acc:MGI:1920642] |
| 16 | lncRNA | 16886863 | 16887904 | ENSMUSG00000124450 | MGI:7802675 | Gm59807 | predicted gene, 59807 [Source:MGI Symbol;Acc:MGI:7802675] |
| 16 | lncRNA | 16910679 | 16914214 | ENSMUSG00000116645 | MGI:6214984 | Gm49584 | predicted gene, 49584 [Source:MGI Symbol;Acc:MGI:6214984] |
| 16 | lncRNA | 16950326 | 16962363 | ENSMUSG00000136619 | MGI:7804561 | Gm60758 | predicted gene, 60758 [Source:MGI Symbol;Acc:MGI:7804561] |

| | | | | | | | |
|---|---|---|---|---|---|---|---|
| 16 | lncRNA | 17051465 | 17094582 | ENSMUSG00000116743 | MGI:6214966 | Gm49573 | predicted gene, 49573 [Source:MGI Symbol;Acc:MGI:6214966] |
| 16 | lncRNA | 17269243 | 17269754 | ENSMUSG00000134583 | MGI:7804283 | Gm60618 | predicted gene, 60618 [Source:MGI Symbol;Acc:MGI:7804283] |
| 16 | lncRNA | 17305055 | 17309053 | ENSMUSG00000125877 | MGI:7802811 | Gm59875 | predicted gene, 59875 [Source:MGI Symbol;Acc:MGI:7802811] |
| 16 | lncRNA | 17348623 | 17349393 | ENSMUSG00000116898 | MGI:6215307 | Gm49785 | predicted gene, 49785 [Source:MGI Symbol;Acc:MGI:6215307] |
| 16 | lncRNA | 17372506 | 17372706 | ENSMUSG00000132968 | MGI:7804035 | Gm60492 | predicted gene, 60492 [Source:MGI Symbol;Acc:MGI:7804035] |
| 16 | lncRNA | 17414960 | 17437229 | ENSMUSG00000122877 | MGI:7802545 | Gm59742 | predicted gene, 59742 [Source:MGI Symbol;Acc:MGI:7802545] |
| 16 | lncRNA | 17459885 | 17464310 | ENSMUSG00000122922 | MGI:7802551 | Gm59745 | predicted gene, 59745 [Source:MGI Symbol;Acc:MGI:7802551] |
| 16 | lncRNA | 17637250 | 17640232 | ENSMUSG00000125857 | MGI:7802809 | Gm59874 | predicted gene, 59874 [Source:MGI Symbol;Acc:MGI:7802809] |
| 16 | lncRNA | 17739113 | 17740370 | ENSMUSG00000120128 | MGI:7337122 | Gm56604 | predicted gene, 56604 [Source:MGI Symbol;Acc:MGI:7337122] |
| 16 | lncRNA | 17739423 | 17741632 | ENSMUSG00000123401 | MGI:7802609 | Gm59774 | predicted gene, 59774 [Source:MGI Symbol;Acc:MGI:7802609] |
| 16 | lncRNA | 17829693 | 17834855 | ENSMUSG00000125584 | MGI:7802779 | Gm59859 | predicted gene, 59859 [Source:MGI Symbol;Acc:MGI:7802779] |
| 16 | lncRNA | 17836792 | 17846448 | ENSMUSG00000131237 | MGI:7803737 | Gm60343 | predicted gene, 60343 [Source:MGI Symbol;Acc:MGI:7803737] |
| 16 | lncRNA | 17846509 | 17849732 | ENSMUSG00000116658 | MGI:6214978 | Gm49580 | predicted gene, 49580 [Source:MGI Symbol;Acc:MGI:6214978] |
| 16 | lncRNA | 17907246 | 17911119 | ENSMUSG00000139216 | MGI:7805003 | Gm60979 | predicted gene, 60979 [Source:MGI Symbol;Acc:MGI:7805003] |
| 16 | lncRNA | 17932751 | 17934859 | ENSMUSG00000131794 | MGI:7803837 | Gm60393 | predicted gene, 60393 [Source:MGI Symbol;Acc:MGI:7803837] |
| 16 | lncRNA | 17974398 | 17978508 | ENSMUSG00000127684 | MGI:6722909 | Gm53937 | predicted gene, 53937 [Source:MGI Symbol;Acc:MGI:6722909] |
| 16 | lncRNA | 17987405 | 17992187 | ENSMUSG00000142704 | MGI:5593838 | Gm34679 | predicted gene, 34679 [Source:MGI Symbol;Acc:MGI:5593838] |

| | | | | | | | |
|---|---|---|---|---|---|---|---|
| 16 | lncRNA | 18016530 | 18035679 | ENSMUSG00000116557 | MGI:1914029 | 4933432I09Rik | RIKEN cDNA 4933432I09 gene [Source:MGI Symbol;Acc:MGI:1914029] |
| 16 | lncRNA | 18031117 | 18033910 | ENSMUSG00000116772 | MGI:1925419 | 4930483P17Rik | RIKEN cDNA 4930483P17 gene [Source:MGI Symbol;Acc:MGI:1925419] |
| 16 | lncRNA | 18053109 | 18058284 | ENSMUSG00000131206 | MGI:7803735 | Gm60342 | predicted gene, 60342 [Source:MGI Symbol;Acc:MGI:7803735] |
| 16 | lncRNA | 18107303 | 18118210 | ENSMUSG00000141558 | MGI:7805321 | Gm61138 | predicted gene, 61138 [Source:MGI Symbol;Acc:MGI:7805321] |
| 16 | lncRNA | 18197049 | 18198469 | ENSMUSG00000126711 | MGI:7802891 | Gm59915 | predicted gene, 59915 [Source:MGI Symbol;Acc:MGI:7802891] |
| 16 | lncRNA | 18218314 | 18219589 | ENSMUSG00000126471 | MGI:7802863 | Gm59901 | predicted gene, 59901 [Source:MGI Symbol;Acc:MGI:7802863] |
| 16 | lncRNA | 18235419 | 18241192 | ENSMUSG00000087556 | MGI:3783206 | Gm15764 | predicted gene 15764 [Source:MGI Symbol;Acc:MGI:3783206] |
| 16 | lncRNA | 18317651 | 18320408 | ENSMUSG00000121876 | MGI:3833934 | Rtl10 | retrotransposon Gag like 10 [Source:MGI Symbol;Acc:MGI:3833934] |
| 16 | lncRNA | 18405901 | 18410211 | ENSMUSG00000132162 | | | novel transcript, antisense to Tbx1 |
| 16 | lncRNA | 18409743 | 18420004 | ENSMUSG00000132088 | MGI:7803897 | Gm60423 | predicted gene, 60423 [Source:MGI Symbol;Acc:MGI:7803897] |
| 16 | lncRNA | 18410669 | 18428506 | ENSMUSG00000075335 | MGI:3588254 | 4930588K23Rik | RIKEN cDNA 4930588K23 gene [Source:MGI Symbol;Acc:MGI:3588254] |
| 16 | lncRNA | 18427517 | 18436688 | ENSMUSG00000132123 | MGI:7803903 | Gm60426 | predicted gene, 60426 [Source:MGI Symbol;Acc:MGI:7803903] |
| 16 | lncRNA | 18435825 | 18436786 | ENSMUSG00000132197 | MGI:7803913 | Gm60431 | predicted gene, 60431 [Source:MGI Symbol;Acc:MGI:7803913] |
| 16 | lncRNA | 18500866 | 18573973 | ENSMUSG00000143558 | | | novel transcript |
| 16 | lncRNA | 18513380 | 18530525 | ENSMUSG00000143641 | MGI:7805610 | Gm61283 | predicted gene, 61283 [Source:MGI Symbol;Acc:MGI:7805610] |
| 16 | lncRNA | 18532404 | 18558588 | ENSMUSG00000143599 | MGI:7805600 | Gm61278 | predicted gene, 61278 [Source:MGI Symbol;Acc:MGI:7805600] |
| 16 | lncRNA | 18594613 | 18595436 | ENSMUSG00000129554 | MGI:7803339 | Gm60143 | predicted gene, 60143 [Source:MGI Symbol;Acc:MGI:7803339] |

| | | | | | | | |
|---|---|---|---|---|---|---|---|
| 16 | lncRNA | 18598650 | 18605054 | ENSMUSG00000131881 | MGI:7803867 | Gm60408 | predicted gene, 60408 [Source:MGI Symbol;Acc:MGI:7803867] |
| 16 | lncRNA | 18614470 | 18616329 | ENSMUSG00000131917 | MGI:6366339 | Gm52240 | predicted gene, 52240 [Source:MGI Symbol;Acc:MGI:6366339] |
| 16 | lncRNA | 18695432 | 18695601 | ENSMUSG00000130102 | MGI:7803395 | Gm60171 | predicted gene, 60171 [Source:MGI Symbol;Acc:MGI:7803395] |
| 16 | lncRNA | 18830004 | 18869920 | ENSMUSG00000129794 | MGI:7803377 | Gm60162 | predicted gene, 60162 [Source:MGI Symbol;Acc:MGI:7803377] |
| 16 | lncRNA | 18870082 | 18925657 | ENSMUSG00000106567 | MGI:1917268 | 2010309G21Rik | RIKEN cDNA 2010309G21 gene [Source:MGI Symbol;Acc:MGI:1917268] |
| 16 | lncRNA | 19031405 | 19055691 | ENSMUSG00000104927 | MGI:5663525 | Gm43388 | predicted gene 43388 [Source:MGI Symbol;Acc:MGI:5663525] |
| 16 | lncRNA | 19059925 | 19061438 | ENSMUSG00000120432 | MGI:7337114 | Gm56600 | predicted gene, 56600 [Source:MGI Symbol;Acc:MGI:7337114] |
| 16 | lncRNA | 19249434 | 19264227 | ENSMUSG00000141973 | MGI:7805369 | Gm61162 | predicted gene, 61162 [Source:MGI Symbol;Acc:MGI:7805369] |
| 16 | lncRNA | 19255062 | 19256544 | ENSMUSG00000142056 | MGI:7805415 | Gm61185 | predicted gene, 61185 [Source:MGI Symbol;Acc:MGI:7805415] |
| 16 | lncRNA | 19667792 | 19669093 | ENSMUSG00000139649 | MGI:7805067 | Gm61011 | predicted gene, 61011 [Source:MGI Symbol;Acc:MGI:7805067] |
| 16 | lncRNA | 19702254 | 19703034 | ENSMUSG00000116712 | MGI:6215005 | Gm49598 | predicted gene, 49598 [Source:MGI Symbol;Acc:MGI:6215005] |
| 16 | lncRNA | 19736660 | 19737002 | ENSMUSG00000143559 | MGI:7805598 | Gm61277 | predicted gene, 61277 [Source:MGI Symbol;Acc:MGI:7805598] |
| 16 | lncRNA | 19881953 | 19916124 | ENSMUSG00000121994 | MGI:7802475 | Gm59707 | predicted gene, 59707 [Source:MGI Symbol;Acc:MGI:7802475] |
| 16 | lncRNA | 19994858 | 19995848 | ENSMUSG00000128869 | MGI:7805756 | Gm61361 | predicted gene, 61361 [Source:MGI Symbol;Acc:MGI:7805756] |
| 16 | lncRNA | 19998599 | 20001075 | ENSMUSG00000128829 | MGI:7803257 | Gm60102 | predicted gene, 60102 [Source:MGI Symbol;Acc:MGI:7803257] |
| 16 | lncRNA | 20018424 | 20021080 | ENSMUSG00000134705 | MGI:7804303 | Gm60629 | predicted gene, 60629 [Source:MGI Symbol;Acc:MGI:7804303] |
| 16 | lncRNA | 20059407 | 20060864 | ENSMUSG00000116913 | MGI:6214956 | Gm49566 | predicted gene, 49566 [Source:MGI Symbol;Acc:MGI:6214956] |

| | | | | | | | |
|---|---|---|---|---|---|---|---|
| 16 | lncRNA | 20194384 | 20198690 | ENSMUSG00000134802 | MGI:7804329 | Gm60642 | predicted gene, 60642 [Source:MGI Symbol;Acc:MGI:7804329] |
| 16 | lncRNA | 20263987 | 20270959 | ENSMUSG00000134765 | MGI:7804323 | Gm60639 | predicted gene, 60639 [Source:MGI Symbol;Acc:MGI:7804323] |
| 16 | lncRNA | 20270241 | 20273128 | ENSMUSG00000120468 | MGI:7337120 | Gm56603 | predicted gene, 56603 [Source:MGI Symbol;Acc:MGI:7337120] |
| 16 | lncRNA | 20271000 | 20278095 | ENSMUSG00000134726 | MGI:5624317 | Gm41432 | predicted gene, 41432 [Source:MGI Symbol;Acc:MGI:5624317] |
| 16 | lncRNA | 20292658 | 20303397 | ENSMUSG00000134688 | MGI:7804299 | Gm60627 | predicted gene, 60627 [Source:MGI Symbol;Acc:MGI:7804299] |
| 16 | lncRNA | 20308232 | 20310021 | ENSMUSG00000129936 | MGI:7805758 | Gm61362 | predicted gene, 61362 [Source:MGI Symbol;Acc:MGI:7805758] |
| 16 | lncRNA | 20329883 | 20336516 | ENSMUSG00000086837 | MGI:4439542 | Gm16618 | predicted gene, 16618 [Source:MGI Symbol;Acc:MGI:4439542] |
| 16 | lncRNA | 20337986 | 20339633 | ENSMUSG00000141814 | MGI:7805357 | Gm61156 | predicted gene, 61156 [Source:MGI Symbol;Acc:MGI:7805357] |
| 16 | lncRNA | 20363953 | 20367283 | ENSMUSG00000121849 | | | mitochondrial ribosomal protein S18B pseudogene [Source:NCBI gene (formerly Entrezgene);Acc:100042948] |
| 16 | lncRNA | 20385108 | 20387258 | ENSMUSG00000116604 | MGI:6215237 | Gm49745 | predicted gene, 49745 [Source:MGI Symbol;Acc:MGI:6215237] |
| 16 | lncRNA | 20397869 | 20408808 | ENSMUSG00000134179 | MGI:5594844 | Gm35685 | predicted gene, 35685 [Source:MGI Symbol;Acc:MGI:5594844] |
| 16 | lncRNA | 20408867 | 20409434 | ENSMUSG00000116764 | MGI:6215242 | Gm49748 | predicted gene, 49748 [Source:MGI Symbol;Acc:MGI:6215242] |
| 16 | lncRNA | 20465948 | 20470320 | ENSMUSG00000134212 | MGI:7804197 | Gm60574 | predicted gene, 60574 [Source:MGI Symbol;Acc:MGI:7804197] |
| 16 | lncRNA | 20619042 | 20622135 | ENSMUSG00000116922 | MGI:1921159 | 4833419O12Rik | RIKEN cDNA 4833419O12 gene [Source:MGI Symbol;Acc:MGI:1921159] |
| 16 | lncRNA | 20638540 | 20640264 | ENSMUSG00000143957 | MGI:7805714 | Gm61338 | predicted gene, 61338 [Source:MGI Symbol;Acc:MGI:7805714] |
| 16 | lncRNA | 20647788 | 20732730 | ENSMUSG00000129217 | MGI:7803285 | Gm60116 | predicted gene, 60116 [Source:MGI Symbol;Acc:MGI:7803285] |
| 16 | lncRNA | 20694007 | 20712309 | ENSMUSG00000116850 | MGI:6215232 | Gm49742 | predicted gene, 49742 [Source:MGI Symbol;Acc:MGI:6215232] |

| | | | | | | | |
|---|---|---|---|---|---|---|---|
| 16 | lncRNA | 20747265 | 20749766 | ENSMUSG00000134417 | MGI:7804239 | Gm60596 | predicted gene, 60596 [Source:MGI Symbol;Acc:MGI:7804239] |
| 16 | lncRNA | 20747436 | 20749160 | ENSMUSG00000134456 | | | predicted gene 6551 [Source:NCBI gene (formerly Entrezgene);Acc:625060] |
| 16 | lncRNA | 20880680 | 20882067 | ENSMUSG00000130835 | | | predicted gene 6557 [Source:NCBI gene (formerly Entrezgene);Acc:625123] |
| 16 | lncRNA | 20929759 | 20935770 | ENSMUSG00000141159 | MGI:7805241 | Gm61098 | predicted gene, 61098 [Source:MGI Symbol;Acc:MGI:7805241] |
| 16 | lncRNA | 20931581 | 20938100 | ENSMUSG00000141120 | MGI:7805235 | Gm61095 | predicted gene, 61095 [Source:MGI Symbol;Acc:MGI:7805235] |
| 16 | lncRNA | 21057245 | 21081909 | ENSMUSG00000116843 | MGI:6215246 | Gm49750 | predicted gene, 49750 [Source:MGI Symbol;Acc:MGI:6215246] |
| 16 | lncRNA | 21066244 | 21115622 | ENSMUSG00000096918 | MGI:4439787 | Gm16863 | predicted gene, 16863 [Source:MGI Symbol;Acc:MGI:4439787] |
| 16 | lncRNA | 21083898 | 21090711 | ENSMUSG00000130455 | MGI:7803497 | Gm60222 | predicted gene, 60222 [Source:MGI Symbol;Acc:MGI:7803497] |
| 16 | lncRNA | 21129303 | 21149480 | ENSMUSG00000134551 | MGI:7804275 | Gm60614 | predicted gene, 60614 [Source:MGI Symbol;Acc:MGI:7804275] |
| 16 | lncRNA | 21140249 | 21152139 | ENSMUSG00000121787 | | | MAGE family member F1 [Source:NCBI gene (formerly Entrezgene);Acc:76222] |
| 16 | lncRNA | 21152105 | 21158921 | ENSMUSG00000120420 | MGI:7337138 | Gm56612 | predicted gene, 56612 [Source:MGI Symbol;Acc:MGI:7337138] |
| 16 | lncRNA | 21172798 | 21188491 | ENSMUSG00000122490 | MGI:7802507 | Gm59723 | predicted gene, 59723 [Source:MGI Symbol;Acc:MGI:7802507] |
| 16 | lncRNA | 21177282 | 21214274 | ENSMUSG00000122447 | | | novel transcript |
| 16 | lncRNA | 21210108 | 21217490 | ENSMUSG00000122534 | MGI:5595309 | Gm36150 | predicted gene, 36150 [Source:MGI Symbol;Acc:MGI:5595309] |
| 16 | lncRNA | 21240234 | 21241878 | ENSMUSG00000138453 | MGI:6722913 | Gm53939 | predicted gene, 53939 [Source:MGI Symbol;Acc:MGI:6722913] |
| 16 | lncRNA | 21329875 | 21346379 | ENSMUSG00000138790 | MGI:7804961 | Gm60958 | predicted gene, 60958 [Source:MGI Symbol;Acc:MGI:7804961] |
| 16 | miRNA | 8441027 | 8441134 | ENSMUSG00002075670 | MGI:6846536 | Gm55031 | predicted gene, 55031 [Source:MGI Symbol;Acc:MGI:6846536] |

| | | | | | | | |
|---|---|---|---|---|---|---|---|
| 16 | miRNA | 10091614 | 10091685 | ENSMUSG00002076434 | MGI:6846586 | Gm55056 | predicted gene, 55056 [Source:MGI Symbol;Acc:MGI:6846586] |
| 16 | miRNA | 10783588 | 10783713 | ENSMUSG00002075232 | MGI:6846702 | Gm55114 | predicted gene, 55114 [Source:MGI Symbol;Acc:MGI:6846702] |
| 16 | miRNA | 10961989 | 10962045 | ENSMUSG00002076173 | MGI:6846709 | Gm55118 | predicted gene, 55118 [Source:MGI Symbol;Acc:MGI:6846709] |
| 16 | miRNA | 11072232 | 11072309 | ENSMUSG00000088544 | MGI:3836984 | Mir1945 | microRNA 1945 [Source:MGI Symbol;Acc:MGI:3836984] |
| 16 | miRNA | 13267387 | 13267465 | ENSMUSG00000076972 | MGI:3718458 | Mir193b | microRNA 193b [Source:MGI Symbol;Acc:MGI:3718458] |
| 16 | miRNA | 13271704 | 13271790 | ENSMUSG00000065449 | MGI:3619371 | Mir365-1 | microRNA 365-1 [Source:MGI Symbol;Acc:MGI:3619371] |
| 16 | miRNA | 13288944 | 13289044 | ENSMUSG00000099214 | MGI:5531065 | Mir6365 | microRNA 6365 [Source:MGI Symbol;Acc:MGI:5531065] |
| 16 | miRNA | 13637096 | 13637184 | ENSMUSG00002075895 | MGI:6846797 | Gm55162 | predicted gene, 55162 [Source:MGI Symbol;Acc:MGI:6846797] |
| 16 | miRNA | 13946430 | 13946495 | ENSMUSG00002076089 | MGI:6845248 | Gm54384 | predicted gene, 54384 [Source:MGI Symbol;Acc:MGI:6845248] |
| 16 | miRNA | 13977490 | 13977556 | ENSMUSG00000070074 | MGI:3619421 | Mir484 | microRNA 484 [Source:MGI Symbol;Acc:MGI:3619421] |
| 16 | miRNA | 16941925 | 16942006 | ENSMUSG00000065572 | MGI:3618716 | Mir130b | microRNA 130b [Source:MGI Symbol;Acc:MGI:3618716] |
| 16 | miRNA | 16942264 | 16942360 | ENSMUSG00000076288 | MGI:3691602 | Mir301b | microRNA 301b [Source:MGI Symbol;Acc:MGI:3691602] |
| 16 | miRNA | 17982953 | 17983034 | ENSMUSG00000099293 | MGI:5531102 | Mir6366 | microRNA 6366 [Source:MGI Symbol;Acc:MGI:5531102] |
| 16 | miRNA | 18102103 | 18102181 | ENSMUSG00000080452 | MGI:3836985 | Mir1306 | microRNA 1306 [Source:MGI Symbol;Acc:MGI:3836985] |
| 16 | miRNA | 18102414 | 18102501 | ENSMUSG00000093246 | MGI:5454349 | Mir3618 | microRNA 3618 [Source:MGI Symbol;Acc:MGI:5454349] |
| 16 | miRNA | 18145265 | 18145329 | ENSMUSG00000065464 | MGI:2676849 | Mir185 | microRNA 185 [Source:MGI Symbol;Acc:MGI:2676849] |
| 16 | miRNA | 18179300 | 18179370 | ENSMUSG00002075123 | MGI:6847211 | Gm55370 | predicted gene, 55370 [Source:MGI Symbol;Acc:MGI:6847211] |
| 16 | miRNA | 19138436 | 19138545 | ENSMUSG00002075049 | MGI:6845949 | Gm54736 | predicted gene, 54736 [Source:MGI Symbol;Acc:MGI:6845949] |

| | | | | | | | |
|---|---|---|---|---|---|---|---|
| 16 | miRNA | 19702549 | 19702650 | ENSMUSG00002075497 | MGI:6845973 | Gm54748 | predicted gene, 54748 [Source:MGI Symbol;Acc:MGI:6845973] |
| 16 | miRNA | 20423202 | 20423286 | ENSMUSG00000080669 | MGI:3764925 | Mir1224 | microRNA 1224 [Source:MGI Symbol;Acc:MGI:3764925] |
| 16 | miRNA | 21074763 | 21074816 | ENSMUSG00000098400 | MGI:5530799 | Mir7680 | microRNA 7680 [Source:MGI Symbol;Acc:MGI:5530799] |
| 16 | misc_RNA | 8021839 | 8022112 | ENSMUSG00002075614 | MGI:6846520 | Gm55023 | predicted gene, 55023 [Source:MGI Symbol;Acc:MGI:6846520] |
| 16 | misc_RNA | 8267362 | 8267650 | ENSMUSG00002076758 | MGI:6846522 | Gm55024 | predicted gene, 55024 [Source:MGI Symbol;Acc:MGI:6846522] |
| 16 | misc_RNA | 20264403 | 20264677 | ENSMUSG00002076542 | MGI:6846033 | Gm54778 | predicted gene, 54778 [Source:MGI Symbol;Acc:MGI:6846033] |
| 16 | processed_pseudogene | 7526600 | 7526930 | ENSMUSG00000115989 | MGI:6155234 | Gm49533 | predicted gene, 49533 [Source:MGI Symbol;Acc:MGI:6155234] |
| 16 | processed_pseudogene | 7921238 | 7921652 | ENSMUSG00000116151 | MGI:6155237 | Gm49535 | predicted gene, 49535 [Source:MGI Symbol;Acc:MGI:6155237] |
| 16 | processed_pseudogene | 8441050 | 8441584 | ENSMUSG00000083505 | MGI:3648278 | Gm7541 | predicted gene 7541 [Source:MGI Symbol;Acc:MGI:3648278] |
| 16 | processed_pseudogene | 8443073 | 8443181 | ENSMUSG00000116095 | MGI:6155257 | Gm49549 | predicted gene, 49549 [Source:MGI Symbol;Acc:MGI:6155257] |
| 16 | processed_pseudogene | 8739924 | 8740253 | ENSMUSG00000068674 | MGI:3641838 | Gm10247 | predicted gene 10247 [Source:MGI Symbol;Acc:MGI:3641838] |
| 16 | processed_pseudogene | 8813952 | 8814260 | ENSMUSG00000116352 | MGI:6155096 | Gm49447 | predicted gene, 49447 [Source:MGI Symbol;Acc:MGI:6155096] |
| 16 | processed_pseudogene | 8922667 | 8925634 | ENSMUSG00000116348 | MGI:5010907 | Gm18722 | predicted gene, 18722 [Source:MGI Symbol;Acc:MGI:5010907] |
| 16 | processed_pseudogene | 9862829 | 9863594 | ENSMUSG00000098100 | MGI:3645867 | Gm7572 | predicted gene 7572 [Source:MGI Symbol;Acc:MGI:3645867] |

| | | | | | | | |
|---|---|---|---|---|---|---|---|
| 16 | processed_pseudogene | 9863623 | 9863829 | ENSMUSG00000098170 | MGI:3782063 | Gm3890 | predicted gene 3890 [Source:MGI Symbol;Acc:MGI:3782063] |
| 16 | processed_pseudogene | 10056030 | 10056380 | ENSMUSG00000116187 | MGI:6155112 | Gm49456 | predicted gene, 49456 [Source:MGI Symbol;Acc:MGI:6155112] |
| 16 | processed_pseudogene | 10619245 | 10619577 | ENSMUSG00000071671 | MGI:3642690 | Gm10343 | predicted gene 10343 [Source:MGI Symbol;Acc:MGI:3642690] |
| 16 | processed_pseudogene | 10854538 | 10855195 | ENSMUSG00000115911 | MGI:3782093 | Gm3919 | predicted gene 3919 [Source:MGI Symbol;Acc:MGI:3782093] |
| 16 | processed_pseudogene | 11002997 | 11003963 | ENSMUSG00000080790 | MGI:3643470 | Gm7638 | predicted gene 7638 [Source:MGI Symbol;Acc:MGI:3643470] |
| 16 | processed_pseudogene | 11008118 | 11009129 | ENSMUSG00000081052 | MGI:3644967 | Gm6305 | predicted gene 6305 [Source:MGI Symbol;Acc:MGI:3644967] |
| 16 | processed_pseudogene | 11312685 | 11314170 | ENSMUSG00000082653 | MGI:3801942 | Gm15897 | predicted gene 15897 [Source:MGI Symbol;Acc:MGI:3801942] |
| 16 | processed_pseudogene | 12198994 | 12199366 | ENSMUSG00000116403 | MGI:5011756 | Gm19571 | predicted gene, 19571 [Source:MGI Symbol;Acc:MGI:5011756] |
| 16 | processed_pseudogene | 12226147 | 12227041 | ENSMUSG00000116013 | MGI:2146727 | C87114 | expressed sequence C87114 [Source:MGI Symbol;Acc:MGI:2146727] |
| 16 | processed_pseudogene | 12577896 | 12579230 | ENSMUSG00000091416 | MGI:3646646 | Gm6327 | predicted gene 6327 [Source:MGI Symbol;Acc:MGI:3646646] |
| 16 | processed_pseudogene | 12768645 | 12769327 | ENSMUSG00000116170 | MGI:3648790 | Gm6332 | predicted gene 6332 [Source:MGI Symbol;Acc:MGI:3648790] |
| 16 | processed_pseudogene | 13828626 | 13829580 | ENSMUSG00000082554 | MGI:3802043 | Gm15806 | predicted gene 15806 [Source:MGI Symbol;Acc:MGI:3802043] |

| | | | | | | | |
|---|---|---|---|---|---|---|---|
| 16 | processed_pseudogene | 13855598 | 13856725 | ENSMUSG00000082277 | MGI:3802044 | Gm15807 | predicted gene 15807 [Source:MGI Symbol;Acc:MGI:3802044] |
| 16 | processed_pseudogene | 13885267 | 13885870 | ENSMUSG00000083536 | MGI:3802045 | Gm15808 | predicted gene 15808 [Source:MGI Symbol;Acc:MGI:3802045] |
| 16 | processed_pseudogene | 14176296 | 14176761 | ENSMUSG00000116025 | MGI:6155210 | Gm49517 | predicted gene, 49517 [Source:MGI Symbol;Acc:MGI:6155210] |
| 16 | processed_pseudogene | 14608281 | 14608875 | ENSMUSG00000116400 | MGI:6155231 | Gm49530 | predicted gene, 49530 [Source:MGI Symbol;Acc:MGI:6155231] |
| 16 | processed_pseudogene | 14818186 | 14818353 | ENSMUSG00000116317 | MGI:6155238 | Gm49536 | predicted gene, 49536 [Source:MGI Symbol;Acc:MGI:6155238] |
| 16 | processed_pseudogene | 15034723 | 15035192 | ENSMUSG00000069315 | MGI:3644153 | Gm7731 | predicted gene 7731 [Source:MGI Symbol;Acc:MGI:3644153] |
| 16 | processed_pseudogene | 15036708 | 15039372 | ENSMUSG00000116213 | MGI:3647383 | Gm5404 | predicted gene 5404 [Source:MGI Symbol;Acc:MGI:3647383] |
| 16 | processed_pseudogene | 15244015 | 15245068 | ENSMUSG00000080704 | MGI:3645805 | Gm6031 | predicted gene 6031 [Source:MGI Symbol;Acc:MGI:3645805] |
| 16 | processed_pseudogene | 15269868 | 15270626 | ENSMUSG00000116423 | MGI:3642962 | Gm7085 | predicted gene 7085 [Source:MGI Symbol;Acc:MGI:3642962] |
| 16 | processed_pseudogene | 15405229 | 15405923 | ENSMUSG00000116386 | MGI:6155240 | Gm49538 | predicted gene, 49538 [Source:MGI Symbol;Acc:MGI:6155240] |
| 16 | processed_pseudogene | 15426971 | 15427379 | ENSMUSG00000116140 | MGI:3646851 | Gm7748 | predicted gene 7748 [Source:MGI Symbol;Acc:MGI:3646851] |
| 16 | processed_pseudogene | 15568787 | 15569056 | ENSMUSG00000116118 | MGI:6155202 | Gm49511 | predicted gene, 49511 [Source:MGI Symbol;Acc:MGI:6155202] |

| | | | | | | | |
|---|---|---|---|---|---|---|---|
| 16 | processed_pseudogene | 15689693 | 15690023 | ENSMUSG00000116102 | MGI:5012267 | Gm20082 | predicted gene, 20082 [Source:MGI Symbol;Acc:MGI:5012267] |
| 16 | processed_pseudogene | 15917462 | 15917975 | ENSMUSG00000116385 | MGI:5010519 | Gm18334 | predicted gene, 18334 [Source:MGI Symbol;Acc:MGI:5010519] |
| 16 | processed_pseudogene | 16093467 | 16093811 | ENSMUSG00000115882 | MGI:3648103 | Gm5481 | predicted gene 5481 [Source:MGI Symbol;Acc:MGI:3648103] |
| 16 | processed_pseudogene | 16197076 | 16198072 | ENSMUSG00000116192 | MGI:5011054 | Gm18869 | predicted gene, 18869 [Source:MGI Symbol;Acc:MGI:5011054] |
| 16 | processed_pseudogene | 16200523 | 16200736 | ENSMUSG00000118629 | MGI:6388916 | Gm53024 | predicted gene, 53024 [Source:MGI Symbol;Acc:MGI:6388916] |
| 16 | processed_pseudogene | 16360187 | 16360606 | ENSMUSG00000089861 | MGI:3801880 | Gm16139 | predicted gene 16139 [Source:MGI Symbol;Acc:MGI:3801880] |
| 16 | processed_pseudogene | 16666715 | 16667562 | ENSMUSG00000116729 | MGI:6215076 | Gm49644 | predicted gene, 49644 [Source:MGI Symbol;Acc:MGI:6215076] |
| 16 | processed_pseudogene | 16670179 | 16671721 | ENSMUSG00000116676 | MGI:3647546 | Gm5768 | predicted gene 5768 [Source:MGI Symbol;Acc:MGI:3647546] |
| 16 | processed_pseudogene | 16750725 | 16750959 | ENSMUSG00000116703 | MGI:5826177 | Gm46540 | predicted gene, 46540 [Source:MGI Symbol;Acc:MGI:5826177] |
| 16 | processed_pseudogene | 16786473 | 16786932 | ENSMUSG00000116598 | MGI:3647845 | Gm6438 | predicted gene 6438 [Source:MGI Symbol;Acc:MGI:3647845] |
| 16 | processed_pseudogene | 16899863 | 16900555 | ENSMUSG00000055497 | MGI:3708610 | Gm9974 | predicted gene 9974 [Source:MGI Symbol;Acc:MGI:3708610] |
| 16 | processed_pseudogene | 16955192 | 16955545 | ENSMUSG00000116694 | MGI:3644482 | Gm6440 | predicted gene 6440 [Source:MGI Symbol;Acc:MGI:3644482] |

| | | | | | | | |
|---|---|---|---|---|---|---|---|
| 16 | processed_pseudogene | 17520791 | 17521455 | ENSMUSG00000116561 | MGI:5010520 | Gm18335 | predicted gene, 18335 [Source:MGI Symbol;Acc:MGI:5010520] |
| 16 | processed_pseudogene | 17522207 | 17522964 | ENSMUSG00000116994 | MGI:6215133 | Gm49684 | predicted gene, 49684 [Source:MGI Symbol;Acc:MGI:6215133] |
| 16 | processed_pseudogene | 17529532 | 17529969 | ENSMUSG00000059150 | MGI:3644695 | Rpl26-ps4 | ribosomal protein L26, pseudogene 4 [Source:MGI Symbol;Acc:MGI:3644695] |
| 16 | processed_pseudogene | 17532725 | 17533003 | ENSMUSG00000116696 | MGI:6215132 | Gm49683 | predicted gene, 49683 [Source:MGI Symbol;Acc:MGI:6215132] |
| 16 | processed_pseudogene | 17543240 | 17543549 | ENSMUSG00000116783 | MGI:3645583 | Gm6048 | predicted gene 6048 [Source:MGI Symbol;Acc:MGI:3645583] |
| 16 | processed_pseudogene | 17552033 | 17552550 | ENSMUSG00000116936 | MGI:3646992 | Bod1-ps | biorientation of chromosomes in cell division 1, pseudogene [Source:MGI Symbol;Acc:MGI:3646992] |
| 16 | processed_pseudogene | 17632892 | 17634185 | ENSMUSG00000116777 | MGI:1343185 | Zfp520-ps | zinc finger protein 520, pseudogene [Source:MGI Symbol;Acc:MGI:1343185] |
| 16 | processed_pseudogene | 18500087 | 18500966 | ENSMUSG00000116825 | MGI:3643238 | Rps2-ps7 | ribosomal protein S2, pseudogene 7 [Source:MGI Symbol;Acc:MGI:3643238] |
| 16 | processed_pseudogene | 18756151 | 18756439 | ENSMUSG00000082130 | MGI:3802098 | Gm15799 | predicted gene 15799 [Source:MGI Symbol;Acc:MGI:3802098] |
| 16 | processed_pseudogene | 18763008 | 18763415 | ENSMUSG00000083658 | MGI:3802099 | Gm15798 | predicted gene 15798 [Source:MGI Symbol;Acc:MGI:3802099] |
| 16 | processed_pseudogene | 18836805 | 18837452 | ENSMUSG00000105028 | MGI:5010932 | Gm18747 | predicted gene, 18747 [Source:MGI Symbol;Acc:MGI:5010932] |
| 16 | processed_pseudogene | 18846982 | 18847404 | ENSMUSG00000060811 | MGI:3641695 | Gm10088 | predicted gene 10088 [Source:MGI Symbol;Acc:MGI:3641695] |

| | | | | | | | |
|---|---|---|---|---|---|---|---|
| 16 | processed_pseudogene | 18980177 | 18986796 | ENSMUSG00000106444 | MGI:5010933 | Gm18748 | predicted gene, 18748 [Source:MGI Symbol;Acc:MGI:5010933] |
| 16 | processed_pseudogene | 19450652 | 19451092 | ENSMUSG00000116790 | MGI:6215003 | Gm49596 | predicted gene, 49596 [Source:MGI Symbol;Acc:MGI:6215003] |
| 16 | processed_pseudogene | 19542062 | 19542728 | ENSMUSG00000097945 | MGI:5009985 | Gm17799 | predicted gene, 17799 [Source:MGI Symbol;Acc:MGI:5009985] |
| 16 | processed_pseudogene | 19671205 | 19671517 | ENSMUSG00000116980 | MGI:6215004 | Gm49597 | predicted gene, 49597 [Source:MGI Symbol;Acc:MGI:6215004] |
| 16 | processed_pseudogene | 20384327 | 20384778 | ENSMUSG00000116942 | MGI:6215236 | Gm49744 | predicted gene, 49744 [Source:MGI Symbol;Acc:MGI:6215236] |
| 16 | processed_pseudogene | 20390160 | 20390432 | ENSMUSG00000116538 | MGI:6215239 | Gm49746 | predicted gene, 49746 [Source:MGI Symbol;Acc:MGI:6215239] |
| 16 | processed_pseudogene | 20401859 | 20402174 | ENSMUSG00000116808 | MGI:5012349 | Gm20164 | predicted gene, 20164 [Source:MGI Symbol;Acc:MGI:5012349] |
| 16 | processed_pseudogene | 20482504 | 20483806 | ENSMUSG00000090319 | MGI:3782646 | Gm4462 | predicted gene 4462 [Source:MGI Symbol;Acc:MGI:3782646] |
| 16 | processed_pseudogene | 20615977 | 20616523 | ENSMUSG00000116776 | MGI:5826181 | Gm46544 | predicted gene, 46544 [Source:MGI Symbol;Acc:MGI:5826181] |
| 16 | processed_pseudogene | 20640895 | 20641390 | ENSMUSG00000099764 | MGI:3645627 | Rps10-ps2 | ribosomal protein S10, pseudogene 2 [Source:MGI Symbol;Acc:MGI:3645627] |
| 16 | processed_pseudogene | 20779377 | 20779892 | ENSMUSG00000116732 | MGI:5010954 | Gm18769 | predicted gene, 18769 [Source:MGI Symbol;Acc:MGI:5010954] |
| 16 | processed_pseudogene | 20840078 | 20840592 | ENSMUSG00000116592 | MGI:5010955 | Gm18770 | predicted gene, 18770 [Source:MGI Symbol;Acc:MGI:5010955] |

| | | | | | | | |
|---|---|---|---|---|---|---|---|
| 16 | processed_pseudogene | 20898420 | 20899416 | ENSMUSG00000086936 | MGI:3780104 | Gm9697 | predicted gene 9697 [Source:MGI Symbol;Acc:MGI:3780104] |
| 16 | processed_pseudogene | 20940580 | 20941273 | ENSMUSG00000116731 | MGI:5010956 | Gm18771 | predicted gene, 18771 [Source:MGI Symbol;Acc:MGI:5010956] |
| 16 | protein_coding | 5124665 | 7229391 | ENSMUSG00000008658 | MGI:1926224 | Rbfox1 | RNA binding protein, fox-1 homolog (C. elegans) 1 [Source:MGI Symbol;Acc:MGI:1926224] |
| 16 | protein_coding | 8227139 | 8243000 | ENSMUSG00000022715 | MGI:1921970 | Tmem114 | transmembrane protein 114 [Source:MGI Symbol;Acc:MGI:1921970] |
| 16 | protein_coding | 8288652 | 8308548 | ENSMUSG00000039345 | MGI:2384301 | Mettl22 | methyltransferase 22, Kin17 lysine [Source:MGI Symbol;Acc:MGI:2384301] |
| 16 | protein_coding | 8331293 | 8439432 | ENSMUSG00000057880 | MGI:2443582 | Abat | 4-aminobutyrate aminotransferase [Source:MGI Symbol;Acc:MGI:2443582] |
| 16 | protein_coding | 8451093 | 8455576 | ENSMUSG00000043140 | MGI:1913940 | Tmem186 | transmembrane protein 186 [Source:MGI Symbol;Acc:MGI:1913940] |
| 16 | protein_coding | 8455538 | 8480331 | ENSMUSG00000022711 | MGI:1859214 | Pmm2 | phosphomannomutase 2 [Source:MGI Symbol;Acc:MGI:1859214] |
| 16 | protein_coding | 8476444 | 8490019 | ENSMUSG00000008393 | MGI:1196368 | Carhsp1 | calcium regulated heat stable protein 1 [Source:MGI Symbol;Acc:MGI:1196368] |
| 16 | protein_coding | 8498634 | 8501775 | ENSMUSG00000107252 | MGI:3643507 | Litafd | LITAF domain containing [Source:MGI Symbol;Acc:MGI:3643507] |
| 16 | protein_coding | 8507459 | 8610172 | ENSMUSG00000022710 | MGI:2182061 | Usp7 | ubiquitin specific peptidase 7 [Source:MGI Symbol;Acc:MGI:2182061] |
| 16 | protein_coding | 8647964 | 8676786 | ENSMUSG00000022507 | MGI:1916303 | Hapstr1 | HUWE1 associated protein modifying stress responses [Source:MGI Symbol;Acc:MGI:1916303] |

| | | | | | | | |
|---|---|---|---|---|---|---|---|
| 16 | protein_coding | 9385762 | 9813424 | ENSMUSG00000059003 | MGI:95820 | Grin2a | glutamate receptor, ionotropic, NMDA2A (epsilon 1) [Source:MGI Symbol;Acc:MGI:95820] |
| 16 | protein_coding | 9988090 | 9992775 | ENSMUSG00000039209 | MGI:1915422 | Rpl39l | ribosomal protein L39-like [Source:MGI Symbol;Acc:MGI:1915422] |
| 16 | protein_coding | 10010576 | 10069342 | ENSMUSG00000039200 | MGI:1922579 | Atf7ip2 | activating transcription factor 7 interacting protein 2 [Source:MGI Symbol;Acc:MGI:1922579] |
| 16 | protein_coding | 10099613 | 10131832 | ENSMUSG00000022505 | MGI:1098726 | Emp2 | epithelial membrane protein 2 [Source:MGI Symbol;Acc:MGI:1098726] |
| 16 | protein_coding | 10175812 | 10213354 | ENSMUSG00000039179 | MGI:1917676 | Tekt5 | tektin 5 [Source:MGI Symbol;Acc:MGI:1917676] |
| 16 | protein_coding | 10229812 | 10242292 | ENSMUSG00000022503 | MGI:1347073 | Nubp1 | nucleotide binding protein 1 [Source:MGI Symbol;Acc:MGI:1347073] |
| 16 | protein_coding | 10238421 | 10265226 | ENSMUSG00000050908 | MGI:3665441 | Tvp23a | trans-golgi network vesicle protein 23A [Source:MGI Symbol;Acc:MGI:3665441] |
| 16 | protein_coding | 10297923 | 10346282 | ENSMUSG00000022504 | MGI:108445 | Ciita | class II transactivator [Source:MGI Symbol;Acc:MGI:108445] |
| 16 | protein_coding | 10320395 | 10360918 | ENSMUSG00000038055 | MGI:1926236 | Dexi | dexamethasone-induced transcript [Source:MGI Symbol;Acc:MGI:1926236] |
| 16 | protein_coding | 10363203 | 10562742 | ENSMUSG00000068663 | MGI:1921624 | Clec16a | C-type lectin domain family 16, member A [Source:MGI Symbol;Acc:MGI:1921624] |
| 16 | protein_coding | 10600104 | 10603400 | ENSMUSG00000038037 | MGI:1354910 | Socs1 | suppressor of cytokine signaling 1 [Source:MGI Symbol;Acc:MGI:1354910] |
| 16 | protein_coding | 10605800 | 10606524 | ENSMUSG00000043050 | MGI:98785 | Tnp2 | transition protein 2 [Source:MGI Symbol;Acc:MGI:98785] |
| 16 | protein_coding | 10608369 | 10608778 | ENSMUSG00000050058 | MGI:106601 | Prm3 | protamine 3 [Source:MGI Symbol;Acc:MGI:106601] |
| 16 | protein_coding | 10609244 | 10613998 | ENSMUSG00000038015 | MGI:97766 | Prm2 | protamine 2 [Source:MGI Symbol;Acc:MGI:97766] |

| | | | | | | | |
|---|---|---|---|---|---|---|---|
| 16 | protein_coding | 10614190 | 10623703 | ENSMUSG00000022501 | MGI:97765 | Prm1 | protamine 1 [Source:MGI Symbol;Acc:MGI:97765] |
| 16 | protein_coding | 10652923 | 10710830 | ENSMUSG00000037991 | MGI:2685383 | Rmi2 | RecQ mediated genome instability 2 [Source:MGI Symbol;Acc:MGI:2685383] |
| 16 | protein_coding | 10777139 | 10884021 | ENSMUSG00000022500 | MGI:1929512 | Litaf | LPS-induced TN factor [Source:MGI Symbol;Acc:MGI:1929512] |
| 16 | protein_coding | 10878809 | 10892849 | ENSMUSG00000037972 | MGI:1276549 | Snn | stannin [Source:MGI Symbol;Acc:MGI:1276549] |
| 16 | protein_coding | 10892775 | 10952514 | ENSMUSG00000022498 | MGI:1923620 | Txndc11 | thioredoxin domain containing 11 [Source:MGI Symbol;Acc:MGI:1923620] |
| 16 | protein_coding | 10954456 | 10994257 | ENSMUSG00000037965 | MGI:2445044 | Zc3h7a | zinc finger CCCH type containing 7 A [Source:MGI Symbol;Acc:MGI:2445044] |
| 16 | protein_coding | 11010834 | 11021195 | ENSMUSG00000005846 | MGI:1913659 | Rsl1d1 | ribosomal L1 domain containing 1 [Source:MGI Symbol;Acc:MGI:1913659] |
| 16 | protein_coding | 11037156 | 11072189 | ENSMUSG00000062203 | MGI:1316728 | Gspt1 | G1 to S phase transition 1 [Source:MGI Symbol;Acc:MGI:1316728] |
| 16 | protein_coding | 11131676 | 11137938 | ENSMUSG00000022496 | MGI:1343050 | Tnfrsf17 | tumor necrosis factor receptor superfamily, member 17 [Source:MGI Symbol;Acc:MGI:1343050] |
| 16 | protein_coding | 11140772 | 11573336 | ENSMUSG00000071669 | MGI:1921728 | Snx29 | sorting nexin 29 [Source:MGI Symbol;Acc:MGI:1921728] |
| 16 | protein_coding | 11621585 | 11727309 | ENSMUSG00000065979 | MGI:2443300 | Cpped1 | calcineurin-like phosphoesterase domain containing 1 [Source:MGI Symbol;Acc:MGI:2443300] |
| 16 | protein_coding | 11801977 | 12088766 | ENSMUSG00000022494 | MGI:1919805 | Shisa9 | shisa family member 9 [Source:MGI Symbol;Acc:MGI:1919805] |
| 16 | protein_coding | 12927548 | 12968481 | ENSMUSG00000022545 | MGI:1354163 | Ercc4 | excision repair cross-complementing rodent repair deficiency, complementation group 4 [Source:MGI Symbol;Acc:MGI:1354163] |

| | | | | | | | |
|---|---|---|---|---|---|---|---|
| 16 | protein_coding | 13074345 | 13235393 | ENSMUSG00000009569 | MGI:3050795 | Mrtfb | myocardin related transcription factor B [Source:MGI Symbol;Acc:MGI:3050795] |
| 16 | protein_coding | 13355824 | 13486034 | ENSMUSG00000022685 | MGI:1921358 | Parn | poly(A)-specific ribonuclease (deadenylation nuclease) [Source:MGI Symbol;Acc:MGI:1921358] |
| 16 | protein_coding | 13489722 | 13521476 | ENSMUSG00000022684 | MGI:1914368 | Bfar | bifunctional apoptosis regulator [Source:MGI Symbol;Acc:MGI:1914368] |
| 16 | protein_coding | 13532921 | 13548847 | ENSMUSG00000022683 | MGI:1347522 | Pla2g10 | phospholipase A2, group X [Source:MGI Symbol;Acc:MGI:1347522] |
| 16 | protein_coding | 13572430 | 13576401 | ENSMUSG00000118559 | MGI:3045309 | A930007A09Rik | RIKEN cDNA A930007A09 gene [Source:MGI Symbol;Acc:MGI:3045309] |
| 16 | protein_coding | 13598572 | 13632703 | ENSMUSG00000022682 | MGI:1925255 | Rrn3 | RRN3 RNA polymerase I transcription factor homolog (yeast) [Source:MGI Symbol;Acc:MGI:1925255] |
| 16 | protein_coding | 13636709 | 13653315 | ENSMUSG00000022681 | MGI:108471 | Ntan1 | N-terminal Asn amidase [Source:MGI Symbol;Acc:MGI:108471] |
| 16 | protein_coding | 13651012 | 13720995 | ENSMUSG00000022680 | MGI:1920909 | Pdxdc1 | pyridoxal-dependent decarboxylase domain containing 1 [Source:MGI Symbol;Acc:MGI:1920909] |
| 16 | protein_coding | 13721025 | 13767483 | ENSMUSG00000022679 | MGI:2135951 | Mpv17l | Mpv17 transgene, kidney disease mutant-like [Source:MGI Symbol;Acc:MGI:2135951] |
| 16 | protein_coding | 13799563 | 13804752 | ENSMUSG00000065968 | MGI:1921732 | Ifitm7 | interferon induced transmembrane protein 7 [Source:MGI Symbol;Acc:MGI:1921732] |
| 16 | protein_coding | 13804468 | 13919364 | ENSMUSG00000044117 | MGI:1914504 | Bmerb1 | bMERB domain containing 1 [Source:MGI Symbol;Acc:MGI:1914504] |
| 16 | protein_coding | 13927037 | 13981215 | ENSMUSG00000060657 | MGI:2444505 | Marf1 | meiosis regulator and mRNA stability 1 [Source:MGI Symbol;Acc:MGI:2444505] |

| | | | | | | | |
|---|---|---|---|---|---|---|---|
| 16 | protein_coding | 13981139 | 14010792 | ENSMUSG00000022678 | MGI:1914453 | Nde1 | nudE neurodevelopment protein 1 [Source:MGI Symbol;Acc:MGI:1914453] |
| 16 | protein_coding | 14012399 | 14109236 | ENSMUSG00000018830 | MGI:102643 | Myh11 | myosin, heavy polypeptide 11, smooth muscle [Source:MGI Symbol;Acc:MGI:102643] |
| 16 | protein_coding | 14117108 | 14135269 | ENSMUSG00000022677 | MGI:1913336 | Cep20 | centrosomal protein 20 [Source:MGI Symbol;Acc:MGI:1913336] |
| 16 | protein_coding | 14179422 | 14293601 | ENSMUSG00000023088 | MGI:102676 | Abcc1 | ATP-binding cassette, sub-family C member 1 [Source:MGI Symbol;Acc:MGI:102676] |
| 16 | protein_coding | 14523716 | 14527249 | ENSMUSG00000022676 | MGI:1096393 | Snai2 | snail family zinc finger 2 [Source:MGI Symbol;Acc:MGI:1096393] |
| 16 | protein_coding | 14724470 | 14799235 | ENSMUSG00000068617 | MGI:1914043 | Clxn | calaxin [Source:MGI Symbol;Acc:MGI:1914043] |
| 16 | protein_coding | 15369941 | 15413637 | ENSMUSG00000022674 | MGI:1917870 | Ube2v2 | ubiquitin-conjugating enzyme E2 variant 2 [Source:MGI Symbol;Acc:MGI:1917870] |
| 16 | protein_coding | 15441761 | 15455264 | ENSMUSG00000022673 | MGI:103199 | Mcm4 | minichromosome maintenance complex component 4 [Source:MGI Symbol;Acc:MGI:103199] |
| 16 | protein_coding | 15455730 | 15660099 | ENSMUSG00000022672 | MGI:104779 | Prkdc | protein kinase, DNA activated, catalytic polypeptide [Source:MGI Symbol;Acc:MGI:104779] |
| 16 | protein_coding | 15666305 | 15681233 | ENSMUSG00000022671 | MGI:1922845 | Mzt2 | mitotic spindle organizing protein 2 [Source:MGI Symbol;Acc:MGI:1922845] |
| 16 | protein_coding | 15705150 | 15708895 | ENSMUSG00000071637 | MGI:103573 | Cebpd | CCAAT/enhancer binding protein delta [Source:MGI Symbol;Acc:MGI:103573] |
| 16 | protein_coding | 15707088 | 15964715 | ENSMUSG00000041974 | MGI:1924834 | Spidr | scaffolding protein involved in DNA repair [Source:MGI Symbol;Acc:MGI:1924834] |
| 16 | protein_coding | 16031182 | 16090576 | ENSMUSG00000041957 | MGI:1914701 | Pkp2 | plakophilin 2 [Source:MGI Symbol;Acc:MGI:1914701] |

| | | | | | | | |
|---|---|---|---|---|---|---|---|
| 16 | protein_coding | 16120829 | 16127504 | ENSMUSG00000022792 | MGI:1917370 | Yars2 | tyrosyl-tRNA synthetase 2 (mitochondrial) [Source:MGI Symbol;Acc:MGI:1917370] |
| 16 | protein_coding | 16130094 | 16176823 | ENSMUSG00000022789 | MGI:1921256 | Dnm1l | dynamin 1-like [Source:MGI Symbol;Acc:MGI:1921256] |
| 16 | protein_coding | 16234781 | 16418413 | ENSMUSG00000022788 | MGI:2183747 | Fgd4 | FYVE, RhoGEF and PH domain containing 4 [Source:MGI Symbol;Acc:MGI:2183747] |
| 16 | protein_coding | 16490092 | 16494269 | ENSMUSG00000048101 | MGI:109316 | Or7a40 | olfactory receptor family 7 subfamily A member 40 [Source:MGI Symbol;Acc:MGI:109316] |
| 16 | protein_coding | 16570880 | 16647320 | ENSMUSG00000022783 | MGI:1354388 | Spag6l | sperm associated antigen 6-like [Source:MGI Symbol;Acc:MGI:1354388] |
| 16 | protein_coding | 16678535 | 16681849 | ENSMUSG00000075370 | MGI:96529 | Igll1 | immunoglobulin lambda-like polypeptide 1 [Source:MGI Symbol;Acc:MGI:96529] |
| 16 | protein_coding | 16686267 | 16688707 | ENSMUSG00000059305 | MGI:98936 | Vpreb1a | V-set pre-B cell surrogate light chain 1A [Source:MGI Symbol;Acc:MGI:98936] |
| 16 | protein_coding | 16688600 | 16710854 | ENSMUSG00000022779 | MGI:1333803 | Top3b | topoisomerase (DNA) III beta [Source:MGI Symbol;Acc:MGI:1333803] |
| 16 | protein_coding | 16714333 | 16745228 | ENSMUSG00000026181 | MGI:1918464 | Ppm1f | protein phosphatase 1F (PP2C domain containing) [Source:MGI Symbol;Acc:MGI:1918464] |
| 16 | protein_coding | 16801246 | 16865317 | ENSMUSG00000063358 | MGI:1346858 | Mapk1 | mitogen-activated protein kinase 1 [Source:MGI Symbol;Acc:MGI:1346858] |
| 16 | protein_coding | 16887560 | 16904909 | ENSMUSG00000022773 | MGI:1913303 | Ypel1 | yippee like 1 [Source:MGI Symbol;Acc:MGI:1913303] |
| 16 | protein_coding | 16904419 | 16929121 | ENSMUSG00000022771 | MGI:2447857 | Ppil2 | peptidylprolyl isomerase (cyclophilin)-like 2 [Source:MGI Symbol;Acc:MGI:2447857] |
| 16 | protein_coding | 16931262 | 16943031 | ENSMUSG00000049916 | MGI:1917708 | 2610318N02Rik | RIKEN cDNA 2610318N02 gene [Source:MGI Symbol;Acc:MGI:1917708] |

| | | | | | | | |
|---|---|---|---|---|---|---|---|
| 16 | protein_coding | 16948002 | 16950247 | ENSMUSG00000022769 | MGI:2149842 | Sdf2l1 | stromal cell-derived factor 2-like 1 [Source:MGI Symbol;Acc:MGI:2149842] |
| 16 | protein_coding | 16956928 | 16965093 | ENSMUSG00000022768 | MGI:1924122 | Ccdc116 | coiled-coil domain containing 116 [Source:MGI Symbol;Acc:MGI:1924122] |
| 16 | protein_coding | 16962485 | 16978565 | ENSMUSG00000041774 | MGI:1916351 | Ydjc | YdjC homolog (bacterial) [Source:MGI Symbol;Acc:MGI:1916351] |
| 16 | protein_coding | 16969877 | 17020513 | ENSMUSG00000038965 | MGI:109240 | Ube2l3 | ubiquitin-conjugating enzyme E2L 3 [Source:MGI Symbol;Acc:MGI:109240] |
| 16 | protein_coding | 17026467 | 17031846 | ENSMUSG00000071636 | MGI:2685449 | Rimbp3 | RIMS binding protein 3 [Source:MGI Symbol;Acc:MGI:2685449] |
| 16 | protein_coding | 17051436 | 17081294 | ENSMUSG00000050240 | MGI:1929869 | Hic2 | hypermethylated in cancer 2 [Source:MGI Symbol;Acc:MGI:1929869] |
| 16 | protein_coding | 17093941 | 17101093 | ENSMUSG00000055692 | MGI:107238 | Tmem191 | transmembrane protein 191 [Source:MGI Symbol;Acc:MGI:107238] |
| 16 | protein_coding | 17098215 | 17224178 | ENSMUSG00000041720 | MGI:2448506 | Pi4ka | phosphatidylinositol 4-kinase alpha [Source:MGI Symbol;Acc:MGI:2448506] |
| 16 | protein_coding | 17149235 | 17161439 | ENSMUSG00000022766 | MGI:96051 | Serpind1 | serine (or cysteine) peptidase inhibitor, clade D, member 1 [Source:MGI Symbol;Acc:MGI:96051] |
| 16 | protein_coding | 17223850 | 17248691 | ENSMUSG00000022765 | MGI:1914724 | Snap29 | synaptosomal-associated protein 29 [Source:MGI Symbol;Acc:MGI:1914724] |
| 16 | protein_coding | 17269851 | 17305298 | ENSMUSG00000006134 | MGI:104686 | Crkl | v-crk avian sarcoma virus CT10 oncogene homolog-like [Source:MGI Symbol;Acc:MGI:104686] |
| 16 | protein_coding | 17307475 | 17325349 | ENSMUSG00000022763 | MGI:1919418 | Aifm3 | apoptosis-inducing factor, mitochondrion-associated 3 [Source:MGI Symbol;Acc:MGI:1919418] |
| 16 | protein_coding | 17326552 | 17344197 | ENSMUSG00000022761 | MGI:1914113 | Lztr1 | leucine-zipper-like transcriptional regulator, 1 [Source:MGI Symbol;Acc:MGI:1914113] |

| | | | | | | | |
|---|---|---|---|---|---|---|---|
| 16 | protein_coding | 17345846 | 17349000 | ENSMUSG00000022760 | MGI:1916259 | Thap7 | THAP domain containing 7 [Source:MGI Symbol;Acc:MGI:1916259] |
| 16 | protein_coding | 17362329 | 17379111 | ENSMUSG00000022759 | MGI:1921935 | Lrrc74b | leucine rich repeat containing 74B [Source:MGI Symbol;Acc:MGI:1921935] |
| 16 | protein_coding | 17379749 | 17395664 | ENSMUSG00000022758 | MGI:1337113 | P2rx6 | purinergic receptor P2X, ligand-gated ion channel, 6 [Source:MGI Symbol;Acc:MGI:1337113] |
| 16 | protein_coding | 17389882 | 17401078 | ENSMUSG00000022756 | MGI:2146512 | Slc7a4 | solute carrier family 7 (cationic amino acid transporter, y+ system), member 4 [Source:MGI Symbol;Acc:MGI:2146512] |
| 16 | protein_coding | 17437218 | 17462692 | ENSMUSG00000005899 | MGI:1924876 | Smpd4 | sphingomyelin phosphodiesterase 4 [Source:MGI Symbol;Acc:MGI:1924876] |
| 16 | protein_coding | 17464334 | 17468602 | ENSMUSG00000041617 | MGI:1919565 | Ccdc74a | coiled-coil domain containing 74A [Source:MGI Symbol;Acc:MGI:1919565] |
| 16 | protein_coding | 17469072 | 17550755 | ENSMUSG00000012114 | MGI:2137379 | Med15 | mediator complex subunit 15 [Source:MGI Symbol;Acc:MGI:2137379] |
| 16 | protein_coding | 17577482 | 17611246 | ENSMUSG00000022750 | MGI:1337995 | Klhl22 | kelch-like 22 [Source:MGI Symbol;Acc:MGI:1337995] |
| 16 | protein_coding | 17615146 | 17626157 | ENSMUSG00000012017 | MGI:1858430 | Scarf2 | scavenger receptor class F, member 2 [Source:MGI Symbol;Acc:MGI:1858430] |
| 16 | protein_coding | 17650985 | 17652872 | ENSMUSG00000116652 | MGI:3045365 | Fam246a | family with sequence similarity 246 member A [Source:MGI Symbol;Acc:MGI:3045365] |
| 16 | protein_coding | 17653140 | 17656228 | ENSMUSG00000090236 | MGI:1931324 | Car15 | carbonic anhydrase 15 [Source:MGI Symbol;Acc:MGI:1931324] |
| 16 | protein_coding | 17653141 | 17709535 | ENSMUSG00000092470 | MGI:5141983 | Gm20518 | predicted gene 20518 [Source:MGI Symbol;Acc:MGI:5141983] |
| 16 | protein_coding | 17657346 | 17716426 | ENSMUSG00000003166 | MGI:892866 | Dgcr2 | DiGeorge syndrome critical region gene 2 [Source:MGI Symbol;Acc:MGI:892866] |

| | | | | | | | |
|---|---|---|---|---|---|---|---|
| 16 | protein_coding | 17712087 | 17715786 | ENSMUSG00000041566 | MGI:1347557 | Tssk1 | testis-specific serine kinase 1 [Source:MGI Symbol;Acc:MGI:1347557] |
| 16 | protein_coding | 17716501 | 17717888 | ENSMUSG00000045521 | MGI:1347559 | Tssk2 | testis-specific serine kinase 2 [Source:MGI Symbol;Acc:MGI:1347559] |
| 16 | protein_coding | 17718573 | 17729212 | ENSMUSG00000003527 | MGI:107854 | Ess2 | ess-2 splicing factor [Source:MGI Symbol;Acc:MGI:107854] |
| 16 | protein_coding | 17730971 | 17732923 | ENSMUSG00000022738 | MGI:892006 | Gsc2 | goosecoid homebox 2 [Source:MGI Symbol;Acc:MGI:892006] |
| 16 | protein_coding | 17743087 | 17746083 | ENSMUSG00000003528 | MGI:1345283 | Slc25a1 | solute carrier family 25 (mitochondrial carrier, citrate transporter), member 1 [Source:MGI Symbol;Acc:MGI:1345283] |
| 16 | protein_coding | 17798292 | 17799137 | ENSMUSG00000059280 | MGI:98937 | Vpreb1b | V-set pre-B cell surrogate light chain 1B [Source:MGI Symbol;Acc:MGI:98937] |
| 16 | protein_coding | 17870724 | 17889496 | ENSMUSG00000003531 | MGI:1202877 | Dgcr6 | DiGeorge syndrome critical region gene 6 [Source:MGI Symbol;Acc:MGI:1202877] |
| 16 | protein_coding | 17878221 | 17908067 | ENSMUSG00000003526 | MGI:97770 | Prodh | proline dehydrogenase [Source:MGI Symbol;Acc:MGI:97770] |
| 16 | protein_coding | 17945506 | 17970272 | ENSMUSG00000043811 | MGI:2136886 | Rtn4r | reticulon 4 receptor [Source:MGI Symbol;Acc:MGI:2136886] |
| 16 | protein_coding | 18035743 | 18038212 | ENSMUSG00000090777 | MGI:3647803 | Ccdc188 | coiled-coil domain containing 188 [Source:MGI Symbol;Acc:MGI:3647803] |
| 16 | protein_coding | 18038617 | 18053000 | ENSMUSG00000060166 | MGI:1338012 | Zdhhc8 | zinc finger, DHHC domain containing 8 [Source:MGI Symbol;Acc:MGI:1338012] |
| 16 | protein_coding | 18057648 | 18066596 | ENSMUSG00000005732 | MGI:96269 | Ranbp1 | RAN binding protein 1 [Source:MGI Symbol;Acc:MGI:96269] |
| 16 | protein_coding | 18063079 | 18076185 | ENSMUSG00000116925 | MGI:6215291 | Gm49776 | predicted gene, 49776 [Source:MGI Symbol;Acc:MGI:6215291] |
| 16 | protein_coding | 18066543 | 18072636 | ENSMUSG00000022721 | MGI:96270 | Trmt2a | TRM2 tRNA methyltransferase 2A [Source:MGI Symbol;Acc:MGI:96270] |

| | | | | | | | |
|---|---|---|---|---|---|---|---|
| 16 | protein_coding | 18071812 | 18107110 | ENSMUSG00000022718 | MGI:2151114 | Dgcr8 | DGCR8, microprocessor complex subunit [Source:MGI Symbol;Acc:MGI:2151114] |
| 16 | protein_coding | 18118689 | 18165967 | ENSMUSG00000013539 | MGI:101825 | Tango2 | transport and golgi organization 2 [Source:MGI Symbol;Acc:MGI:101825] |
| 16 | protein_coding | 18166046 | 18225826 | ENSMUSG00000118669 | MGI:109620 | Arvcf | armadillo repeat gene deleted in velocardiofacial syndrome [Source:MGI Symbol;Acc:MGI:109620] |
| 16 | protein_coding | 18225636 | 18245602 | ENSMUSG00000000326 | MGI:88470 | Comt | catechol-O-methyltransferase [Source:MGI Symbol;Acc:MGI:88470] |
| 16 | protein_coding | 18245134 | 18297823 | ENSMUSG00000075704 | MGI:1347023 | Txnrd2 | thioredoxin reductase 2 [Source:MGI Symbol;Acc:MGI:1347023] |
| 16 | protein_coding | 18317463 | 18385429 | ENSMUSG00000000884 | MGI:1338057 | Gnb1l | guanine nucleotide binding protein (G protein), beta polypeptide 1-like [Source:MGI Symbol;Acc:MGI:1338057] |
| 16 | protein_coding | 18399729 | 18409421 | ENSMUSG00000009097 | MGI:98493 | Tbx1 | T-box 1 [Source:MGI Symbol;Acc:MGI:98493] |
| 16 | protein_coding | 18439067 | 18441153 | ENSMUSG00000050761 | MGI:107852 | Gp1bb | glycoprotein Ib, beta polypeptide [Source:MGI Symbol;Acc:MGI:107852] |
| 16 | protein_coding | 18439216 | 18445177 | ENSMUSG00000116594 | MGI:6215011 | Gm49601 | predicted gene, 49601 [Source:MGI Symbol;Acc:MGI:6215011] |
| 16 | protein_coding | 18439252 | 18448704 | ENSMUSG00000072214 | MGI:1195461 | Septin5 | septin 5 [Source:MGI Symbol;Acc:MGI:1195461] |
| 16 | protein_coding | 18595597 | 18597012 | ENSMUSG00000041378 | MGI:1276112 | Cldn5 | claudin 5 [Source:MGI Symbol;Acc:MGI:1276112] |
| 16 | protein_coding | 18599197 | 18630737 | ENSMUSG00000000028 | MGI:1338073 | Cdc45 | cell division cycle 45 [Source:MGI Symbol;Acc:MGI:1338073] |
| 16 | protein_coding | 18630529 | 18654011 | ENSMUSG00000005262 | MGI:109353 | Ufd1 | ubiquitin recognition factor in ER-associated degradation 1 [Source:MGI Symbol;Acc:MGI:109353] |
| 16 | protein_coding | 18655328 | 18658910 | ENSMUSG00000071632 | MGI:1919557 | 2510002D24Rik | RIKEN cDNA 2510002D24 gene [Source:MGI Symbol;Acc:MGI:1919557] |
| 16 | protein_coding | 18655334 | 18774997 | ENSMUSG00000099908 | MGI:5579245 | Gm28539 | predicted gene 28539 [Source:MGI Symbol;Acc:MGI:5579245] |

| | | | | | | | |
|---|---|---|---|---|---|---|---|
| 16 | protein_coding | 18690768 | 18695612 | ENSMUSG00000022706 | MGI:1332635 | Mrpl40 | mitochondrial ribosomal protein L40 [Source:MGI Symbol;Acc:MGI:1332635] |
| 16 | protein_coding | 18695787 | 18789059 | ENSMUSG00000022702 | MGI:99430 | Hira | histone cell cycle regulator [Source:MGI Symbol;Acc:MGI:99430] |
| 16 | protein_coding | 19102854 | 19132814 | ENSMUSG00000050742 | MGI:3029998 | Or2m12 | olfactory receptor family 2 subfamily M member 12 [Source:MGI Symbol;Acc:MGI:3029998] |
| 16 | protein_coding | 19224994 | 19241884 | ENSMUSG00000050158 | MGI:3029999 | Or2m13 | olfactory receptor family 2 subfamily M member 13 [Source:MGI Symbol;Acc:MGI:3029999] |
| 16 | protein_coding | 19302444 | 19308632 | ENSMUSG00000056822 | MGI:3030000 | Or2l13 | olfactory receptor family 2 subfamily L member 13 [Source:MGI Symbol;Acc:MGI:3030000] |
| 16 | protein_coding | 19328308 | 19341016 | ENSMUSG00000045341 | MGI:3030001 | Or2l5 | olfactory receptor family 2 subfamily L member 5 [Source:MGI Symbol;Acc:MGI:3030001] |
| 16 | protein_coding | 19348024 | 19354305 | ENSMUSG00000061361 | MGI:3030002 | Or2l13b | olfactory receptor family 2 subfamily L member 13B [Source:MGI Symbol;Acc:MGI:3030002] |
| 16 | protein_coding | 19383639 | 19390559 | ENSMUSG00000068535 | MGI:3030003 | Or2aj4 | olfactory receptor family 2 subfamily AJ member 4 [Source:MGI Symbol;Acc:MGI:3030003] |
| 16 | protein_coding | 19424446 | 19430632 | ENSMUSG00000062245 | MGI:3030004 | Or2aj5 | olfactory receptor family 2 subfamily AJ member 5 [Source:MGI Symbol;Acc:MGI:3030004] |
| 16 | protein_coding | 19442680 | 19448666 | ENSMUSG00000060480 | MGI:3030005 | Or2aj6 | olfactory receptor family 2 subfamily AJ member 6 [Source:MGI Symbol;Acc:MGI:3030005] |
| 16 | protein_coding | 19472128 | 19525125 | ENSMUSG00000041247 | MGI:2441659 | Lamp3 | lysosomal-associated membrane protein 3 [Source:MGI Symbol;Acc:MGI:2441659] |
| 16 | protein_coding | 19578958 | 19591503 | ENSMUSG00000022686 | MGI:2137302 | B3gnt5 | UDP-GlcNAc:betaGal beta-1,3-N-acetylglucosaminyltransferase 5 [Source:MGI Symbol;Acc:MGI:2137302] |

| | | | | | | | |
|---|---|---|---|---|---|---|---|
| 16 | protein_coding | 19695317 | 19703024 | ENSMUSG00000075330 | MGI:1915412 | A930003A15Rik | RIKEN cDNA A930003A15 gene [Source:MGI Symbol;Acc:MGI:1915412] |
| 16 | protein_coding | 19765246 | 19801787 | ENSMUSG00000043008 | MGI:2686922 | Klhl6 | kelch-like 6 [Source:MGI Symbol;Acc:MGI:2686922] |
| 16 | protein_coding | 19916292 | 19947971 | ENSMUSG00000062901 | MGI:1923035 | Klhl24 | kelch-like 24 [Source:MGI Symbol;Acc:MGI:1923035] |
| 16 | protein_coding | 19959813 | 20051323 | ENSMUSG00000041215 | MGI:2447762 | Yeats2 | YEATS domain containing 2 [Source:MGI Symbol;Acc:MGI:2447762] |
| 16 | protein_coding | 20052053 | 20060108 | ENSMUSG00000041205 | MGI:3607784 | Map6d1 | MAP6 domain containing 1 [Source:MGI Symbol;Acc:MGI:3607784] |
| 16 | protein_coding | 20098568 | 20121137 | ENSMUSG00000033918 | MGI:1277152 | Parl | presenilin associated, rhomboid-like [Source:MGI Symbol;Acc:MGI:1277152] |
| 16 | protein_coding | 20127137 | 20144154 | ENSMUSG00000022818 | MGI:3644957 | Cyp2ab1 | cytochrome P450, family 2, subfamily ab, polypeptide 1 [Source:MGI Symbol;Acc:MGI:3644957] |
| 16 | protein_coding | 20150053 | 20245144 | ENSMUSG00000022822 | MGI:1351644 | Abcc5 | ATP-binding cassette, sub-family C member 5 [Source:MGI Symbol;Acc:MGI:1351644] |
| 16 | protein_coding | 20317567 | 20328073 | ENSMUSG00000003235 | MGI:2446176 | Eif2b5 | eukaryotic translation initiation factor 2B, subunit 5 epsilon [Source:MGI Symbol;Acc:MGI:2446176] |
| 16 | protein_coding | 20335732 | 20352760 | ENSMUSG00000003233 | MGI:108100 | Dvl3 | dishevelled segment polarity protein 3 [Source:MGI Symbol;Acc:MGI:108100] |
| 16 | protein_coding | 20354228 | 20363659 | ENSMUSG00000022841 | MGI:1298405 | Ap2m1 | adaptor-related protein complex 2, mu 1 subunit [Source:MGI Symbol;Acc:MGI:1298405] |
| 16 | protein_coding | 20367327 | 20380129 | ENSMUSG00000003234 | MGI:1351656 | Abcf3 | ATP-binding cassette, sub-family F member 3 [Source:MGI Symbol;Acc:MGI:1351656] |
| 16 | protein_coding | 20408221 | 20424127 | ENSMUSG00000046613 | MGI:2681859 | Vwa5b2 | von Willebrand factor A domain containing 5B2 [Source:MGI Symbol;Acc:MGI:2681859] |

| | | | | | | | |
|---|---|---|---|---|---|---|---|
| 16 | protein_coding | 20424124 | 20430485 | ENSMUSG00000033809 | MGI:1098592 | Alg3 | ALG3 alpha-1,3- mannosyltransferase [Source:MGI Symbol;Acc:MGI:1098592] |
| 16 | protein_coding | 20430343 | 20437761 | ENSMUSG00000115219 | MGI:5903914 | Eef1akmt4 | EEF1A lysine methyltransferase 4 [Source:MGI Symbol;Acc:MGI:5903914] |
| 16 | protein_coding | 20430360 | 20464670 | ENSMUSG00000115293 | MGI:6121518 | Eef1ece2 | Eef1akmt4-endothelin converting enzyme 2 readthrough [Source:MGI Symbol;Acc:MGI:6121518] |
| 16 | protein_coding | 20437965 | 20440037 | ENSMUSG00000051146 | MGI:1920297 | Camk2n2 | calcium/calmodulin-dependent protein kinase II inhibitor 2 [Source:MGI Symbol;Acc:MGI:1920297] |
| 16 | protein_coding | 20448578 | 20465235 | ENSMUSG00000022842 | MGI:1101356 | Ece2 | endothelin converting enzyme 2 [Source:MGI Symbol;Acc:MGI:1101356] |
| 16 | protein_coding | 20470402 | 20482164 | ENSMUSG00000006998 | MGI:1096584 | Psmd2 | proteasome (prosome, macropain) 26S subunit, non-ATPase, 2 [Source:MGI Symbol;Acc:MGI:1096584] |
| 16 | protein_coding | 20487063 | 20511634 | ENSMUSG00000045983 | MGI:2384784 | Eif4g1 | eukaryotic translation initiation factor 4, gamma 1 [Source:MGI Symbol;Acc:MGI:2384784] |
| 16 | protein_coding | 20511991 | 20521798 | ENSMUSG00000050821 | MGI:1925658 | Fam131a | family with sequence similarity 131, member A [Source:MGI Symbol;Acc:MGI:1925658] |
| 16 | protein_coding | 20521714 | 20536496 | ENSMUSG00000022843 | MGI:105061 | Clcn2 | chloride channel, voltage-sensitive 2 [Source:MGI Symbol;Acc:MGI:105061] |
| 16 | protein_coding | 20536415 | 20541017 | ENSMUSG00000021018 | MGI:2384309 | Polr2h | polymerase (RNA) II (DNA directed) polypeptide H [Source:MGI Symbol;Acc:MGI:2384309] |
| 16 | protein_coding | 20543204 | 20553261 | ENSMUSG00000022847 | MGI:101875 | Thpo | thrombopoietin [Source:MGI Symbol;Acc:MGI:101875] |
| 16 | protein_coding | 20551877 | 20561134 | ENSMUSG00000006958 | MGI:1313268 | Chrd | chordin [Source:MGI Symbol;Acc:MGI:1313268] |
| 16 | protein_coding | 20971405 | 20972750 | ENSMUSG00000090356 | MGI:1913811 | Teddm3 | transmembrane epididymal family member 3 [Source:MGI Symbol;Acc:MGI:1913811] |

| | | | | | | | |
|---|---|---|---|---|---|---|---|
| 16 | protein_coding | 21023505 | 21042055 | ENSMUSG00000005958 | MGI:104770 | Ephb3 | Eph receptor B3 [Source:MGI Symbol;Acc:MGI:104770] |
| 16 | protein_coding | 21241868 | 21463430 | ENSMUSG00000033653 | MGI:2146407 | Vps8 | VPS8 CORVET complex subunit [Source:MGI Symbol;Acc:MGI:2146407] |
| 16 | protein_coding | 21467795 | 21513415 | ENSMUSG00000043391 | MGI:1919440 | 2510009E07Rik | RIKEN cDNA 2510009E07 gene [Source:MGI Symbol;Acc:MGI:1919440] |
| 16 | rRNA | 8690943 | 8691035 | ENSMUSG00002075441 | MGI:6846556 | Gm55041 | predicted gene, 55041 [Source:MGI Symbol;Acc:MGI:6846556] |
| 16 | rRNA | 13787212 | 13787321 | ENSMUSG00002076461 | MGI:6845244 | Gm54382 | predicted gene, 54382 [Source:MGI Symbol;Acc:MGI:6845244] |
| 16 | rRNA | 15828764 | 15828873 | ENSMUSG00002076768 | MGI:6845344 | Gm54432 | predicted gene, 54432 [Source:MGI Symbol;Acc:MGI:6845344] |
| 16 | snoRNA | 8220678 | 8220821 | ENSMUSG00000077152 | MGI:5454356 | Gm24579 | predicted gene, 24579 [Source:MGI Symbol;Acc:MGI:5454356] |
| 16 | snoRNA | 8298524 | 8298605 | ENSMUSG00000077557 | MGI:5453483 | Gm23706 | predicted gene, 23706 [Source:MGI Symbol;Acc:MGI:5453483] |
| 16 | snoRNA | 13349292 | 13349415 | ENSMUSG00000077488 | MGI:5455053 | Gm25276 | predicted gene, 25276 [Source:MGI Symbol;Acc:MGI:5455053] |
| 16 | snoRNA | 18058581 | 18058706 | ENSMUSG00000080364 | MGI:5455554 | Gm25777 | predicted gene, 25777 [Source:MGI Symbol;Acc:MGI:5455554] |
| 16 | snoRNA | 19126449 | 19126574 | ENSMUSG00000089036 | MGI:5451935 | Gm22158 | predicted gene, 22158 [Source:MGI Symbol;Acc:MGI:5451935] |
| 16 | snoRNA | 19229908 | 19230039 | ENSMUSG00000077172 | MGI:5455539 | Gm25762 | predicted gene, 25762 [Source:MGI Symbol;Acc:MGI:5455539] |
| 16 | snoRNA | 20502989 | 20503063 | ENSMUSG00000077239 | MGI:3819548 | Snord66 | small nucleolar RNA, C/D box 66 [Source:MGI Symbol;Acc:MGI:3819548] |
| 16 | snRNA | 8154857 | 8154960 | ENSMUSG00000089593 | MGI:5455936 | Gm26159 | predicted gene, 26159 [Source:MGI Symbol;Acc:MGI:5455936] |
| 16 | snRNA | 8584033 | 8584140 | ENSMUSG00000084737 | MGI:5455582 | Gm25805 | predicted gene, 25805 [Source:MGI Symbol;Acc:MGI:5455582] |
| 16 | snRNA | 10055782 | 10055884 | ENSMUSG00000089563 | MGI:5454444 | Gm24667 | predicted gene, 24667 [Source:MGI Symbol;Acc:MGI:5454444] |

| | | | | | | | |
|---|---|---|---|---|---|---|---|
| 16 | snRNA | 10812250 | 10812433 | ENSMUSG00002075097 | MGI:6846704 | Gm55115 | predicted gene, 55115 [Source:MGI Symbol;Acc:MGI:6846704] |
| 16 | snRNA | 10984969 | 10985131 | ENSMUSG00000064565 | MGI:5453922 | Gm24145 | predicted gene, 24145 [Source:MGI Symbol;Acc:MGI:5453922] |
| 16 | snRNA | 14200671 | 14200737 | ENSMUSG00000089106 | MGI:5452893 | Gm23116 | predicted gene, 23116 [Source:MGI Symbol;Acc:MGI:5452893] |
| 16 | snRNA | 14625027 | 14625133 | ENSMUSG00000118928 | MGI:5455324 | Gm25547 | predicted gene, 25547 [Source:MGI Symbol;Acc:MGI:5455324] |
| 16 | snRNA | 16823971 | 16824065 | ENSMUSG00002076351 | MGI:6845440 | Gm54480 | predicted gene, 54480 [Source:MGI Symbol;Acc:MGI:6845440] |
| 16 | snRNA | 17023656 | 17023762 | ENSMUSG00000119426 | MGI:5451906 | Gm22129 | predicted gene, 22129 [Source:MGI Symbol;Acc:MGI:5451906] |
| 16 | snRNA | 17159770 | 17159876 | ENSMUSG00000065331 | MGI:5454704 | Gm24927 | predicted gene, 24927 [Source:MGI Symbol;Acc:MGI:5454704] |
| 16 | snRNA | 17300947 | 17301013 | ENSMUSG00002076579 | MGI:6845504 | Gm54512 | predicted gene, 54512 [Source:MGI Symbol;Acc:MGI:6845504] |
| 16 | snRNA | 18173480 | 18173627 | ENSMUSG00002075609 | MGI:6847209 | Gm55369 | predicted gene, 55369 [Source:MGI Symbol;Acc:MGI:6847209] |
| 16 | snRNA | 20564058 | 20564222 | ENSMUSG00000065372 | MGI:5452518 | Gm22741 | predicted gene, 22741 [Source:MGI Symbol;Acc:MGI:5452518] |
| 16 | TEC | 17739907 | 17741083 | ENSMUSG00000103149 | MGI:1858439 | AA914427 | EST AA914427 [Source:MGI Symbol;Acc:MGI:1858439] |
| 16 | TEC | 18300806 | 18305295 | ENSMUSG00000116842 | MGI:6214991 | Gm49588 | predicted gene, 49588 [Source:MGI Symbol;Acc:MGI:6214991] |
| 16 | TEC | 19050083 | 19053552 | ENSMUSG00000105128 | MGI:5663007 | Gm42870 | predicted gene 42870 [Source:MGI Symbol;Acc:MGI:5663007] |
| 16 | TEC | 19062275 | 19063584 | ENSMUSG00000105160 | MGI:2444617 | A530030E21Rik | RIKEN cDNA A530030E21 gene [Source:MGI Symbol;Acc:MGI:2444617] |
| 16 | transcribed_processed_pseudogene | 10864768 | 10866154 | ENSMUSG00000116274 | MGI:5010909 | Gm18724 | predicted gene, 18724 [Source:MGI Symbol;Acc:MGI:5010909] |

| | | | | | | | |
|---|---|---|---|---|---|---|---|
| 16 | transcribed_processed_pseudogene | 13495005 | 13495835 | ENSMUSG00000079737 | MGI:1913848 | Pphln1-ps1 | periphilin 1, pseudogene 1 [Source:MGI Symbol;Acc:MGI:1913848] |
| 16 | transcribed_processed_pseudogene | 16183048 | 16184021 | ENSMUSG00000098037 | MGI:3645843 | Gm7765 | predicted gene 7765 [Source:MGI Symbol;Acc:MGI:3645843] |
| 16 | transcribed_processed_pseudogene | 16637613 | 16637987 | ENSMUSG00000116648 | MGI:3645866 | Rpl31-ps12 | ribosomal protein L31, pseudogene 1 2 [Source:MGI Symbol;Acc:MGI:3645866] |
| 16 | transcribed_processed_pseudogene | 19995477 | 19995762 | ENSMUSG00000084838 | MGI:3704461 | Gm10241 | predicted pseudogene 10241 [Source:MGI Symbol;Acc:MGI:3704461] |
| 16 | transcribed_processed_pseudogene | 20364668 | 20364950 | ENSMUSG00000080797 | MGI:3642834 | Gm15760 | predicted gene 15760 [Source:MGI Symbol;Acc:MGI:3642834] |
| 16 | transcribed_processed_pseudogene | 20639131 | 20639453 | ENSMUSG00000116721 | MGI:3779946 | Gm9536 | predicted gene 9536 [Source:MGI Symbol;Acc:MGI:3779946] |
| 16 | transcribed_processed_pseudogene | 21151396 | 21151879 | ENSMUSG00000116632 | MGI:1923472 | Magef1 | MAGE family member F1 [Source:MGI Symbol;Acc:MGI:1923472] |

| | | | | | | | |
|---|---|---|---|---|---|---|---|
| 16 | transcribed_unitary_pseudogene | 10679619 | 10756228 | ENSMUSG00000117309 | MGI:6270684 | Gm49960 | predicted gene, 49960 [Source:MGI Symbol;Acc:MGI:6270684] |
| 16 | transcribed_unitary_pseudogene | 18319883 | 18320924 | ENSMUSG00000086965 | MGI:3833934 | Rtl10 | retrotransposon Gag like 10 [Source:MGI Symbol;Acc:MGI:3833934] |
| 16 | transcribed_unprocessed_pseudogene | 20307538 | 20309897 | ENSMUSG00000093381 | MGI:5313062 | Gm20615 | predicted gene 20615 [Source:MGI Symbol;Acc:MGI:5313062] |
| 16 | transcribed_unprocessed_pseudogene | 20747719 | 20748603 | ENSMUSG00000116659 | MGI:3644103 | Gm6551 | predicted gene 6551 [Source:MGI Symbol;Acc:MGI:3644103] |
| 16 | transcribed_unprocessed_pseudogene | 20880963 | 20881844 | ENSMUSG00000116710 | MGI:3779606 | Gm6557 | predicted gene 6557 [Source:MGI Symbol;Acc:MGI:3779606] |
| 16 | unprocessed_pseudogene | 13737291 | 13739263 | ENSMUSG00000089818 | MGI:3802178 | Gm15950 | predicted gene 15950 [Source:MGI Symbol;Acc:MGI:3802178] |
| 16 | unprocessed_pseudogene | 14297762 | 14301610 | ENSMUSG00000116439 | MGI:6155229 | Gm49528 | predicted gene, 49528 [Source:MGI Symbol;Acc:MGI:6155229] |

| | | | | | | | |
|---|---|---|---|---|---|---|---|
| 16 | unprocessed_pseudogene | 14319964 | 14327085 | ENSMUSG00000090808 | MGI:2686604 | Gm1758 | predicted gene 1758 [Source:MGI Symbol;Acc:MGI:2686604] |
| 16 | unprocessed_pseudogene | 16490254 | 16490374 | ENSMUSG00000108454 | MGI:5753508 | Gm44932 | predicted gene 44932 [Source:MGI Symbol;Acc:MGI:5753508] |
| 16 | unprocessed_pseudogene | 20254788 | 20262132 | ENSMUSG00000093649 | MGI:5313061 | Gm20614 | predicted gene 20614 [Source:MGI Symbol;Acc:MGI:5313061] |

**Supplementary Table 4. Protein-coding genes located within ±10 Mb of pathology peak QTL on chromosome 5 (peak~98.3Mb)**

| Chr | type | start | stop | gene_id | mgi_type | gene_symbol | description |
|---|---|---|---|---|---|---|---|
| 5 | protein_coding | 88032888 | 88042033 | ENSMUSG00000009580 | MGI:1916842 | Odam | odontogenic, ameloblast asssociated [Source:MGI Symbol;Acc:MGI:1916842] |
| 5 | protein_coding | 88056638 | 88062030 | ENSMUSG00000105888 | MGI:3512757 | Fdcsp | follicular dendritic cell secreted protein [Source:MGI Symbol;Acc:MGI:3512757] |
| 5 | protein_coding | 88073438 | 88080524 | ENSMUSG00000001622 | MGI:107461 | Csn3 | casein kappa [Source:MGI Symbol;Acc:MGI:107461] |
| 5 | protein_coding | 88117318 | 88120729 | ENSMUSG00000007457 | MGI:1921498 | 2310003L06Rik | RIKEN cDNA 2310003L06 gene [Source:MGI Symbol;Acc:MGI:1921498] |
| 5 | protein_coding | 88117376 | 88156136 | ENSMUSG00000100704 | MGI:5579140 | Gm28434 | predicted gene 28434 [Source:MGI Symbol;Acc:MGI:5579140] |
| 5 | protein_coding | 88127298 | 88129403 | ENSMUSG00000007907 | MGI:1918227 | Cabs1 | calcium binding protein, spermatid specific 1 [Source:MGI Symbol;Acc:MGI:1918227] |
| 5 | protein_coding | 88150408 | 88156393 | ENSMUSG00000029280 | MGI:102763 | Smr3a | submaxillary gland androgen regulated protein 3A [Source:MGI Symbol;Acc:MGI:102763] |
| 5 | protein_coding | 88234415 | 88256912 | ENSMUSG00000029281 | MGI:102762 | Smr2 | submaxillary gland androgen regulated protein 2 [Source:MGI Symbol;Acc:MGI:102762] |
| 5 | protein_coding | 88416778 | 88430694 | ENSMUSG00000090302 | MGI:3646267 | Smr2l | submaxillary gland androgen regulated protein 2 like [Source:MGI Symbol;Acc:MGI:3646267] |
| 5 | protein_coding | 88465171 | 88476673 | ENSMUSG00000064156 | MGI:107496 | Prol1 | proline rich, lacrimal 1 [Source:MGI Symbol;Acc:MGI:107496] |
| 5 | protein_coding | 88523967 | 88533775 | ENSMUSG00000029282 | MGI:1918671 | Amtn | amelotin [Source:MGI Symbol;Acc:MGI:1918671] |
| 5 | protein_coding | 88603850 | 88616390 | ENSMUSG00000029288 | MGI:104655 | Ambn | ameloblastin [Source:MGI Symbol;Acc:MGI:104655] |
| 5 | protein_coding | 88635834 | 88653908 | ENSMUSG00000029286 | MGI:1333772 | Enam | enamelin [Source:MGI Symbol;Acc:MGI:1333772] |
| 5 | protein_coding | 88667668 | 88675750 | ENSMUSG00000067149 | MGI:96493 | Jchain | immunoglobulin joining chain [Source:MGI Symbol;Acc:MGI:96493] |
| 5 | protein_coding | 88702321 | 88703949 | ENSMUSG00000070697 | MGI:1919230 | Utp3 | UTP3 small subunit processome component [Source:MGI Symbol;Acc:MGI:1919230] |

| | | | | | | | |
|---|---|---|---|---|---|---|---|
| 5 | protein_coding | 88712899 | 88799251 | ENSMUSG00000029291 | MGI:106484 | Rufy3 | RUN and FYVE domain containing 3 [Source:MGI Symbol;Acc:MGI:106484] |
| 5 | protein_coding | 88807307 | 88824030 | ENSMUSG00000044221 | MGI:106479 | Grsf1 | G-rich RNA sequence binding factor 1 [Source:MGI Symbol;Acc:MGI:106479] |
| 5 | protein_coding | 88868714 | 88912079 | ENSMUSG00000006262 | MGI:1915723 | Mob1b | MOB kinase activator 1B [Source:MGI Symbol;Acc:MGI:1915723] |
| 5 | protein_coding | 88912855 | 88931140 | ENSMUSG00000029366 | MGI:102726 | Dck | deoxycytidine kinase [Source:MGI Symbol;Acc:MGI:102726] |
| 5 | protein_coding | 89034345 | 89387512 | ENSMUSG00000060961 | MGI:1927555 | Slc4a4 | solute carrier family 4 (anion exchanger), member 4 [Source:MGI Symbol;Acc:MGI:1927555] |
| 5 | protein_coding | 89565381 | 89605757 | ENSMUSG00000035540 | MGI:95669 | Gc | vitamin D binding protein [Source:MGI Symbol;Acc:MGI:95669] |
| 5 | protein_coding | 89675288 | 89731599 | ENSMUSG00000035528 | MGI:1860130 | Npffr2 | neuropeptide FF receptor 2 [Source:MGI Symbol;Acc:MGI:1860130] |
| 5 | protein_coding | 89824946 | 90031193 | ENSMUSG00000043635 | MGI:3045353 | Adamts3 | ADAM metallopeptidase with thrombospondin type 1 motif 3 [Source:MGI Symbol;Acc:MGI:3045353] |
| 5 | protein_coding | 90362583 | 90371860 | ENSMUSG00000035505 | MGI:2448532 | Cox18 | cytochrome c oxidase assembly protein 18 [Source:MGI Symbol;Acc:MGI:2448532] |
| 5 | protein_coding | 90375025 | 90514436 | ENSMUSG00000055204 | MGI:1932101 | Ankrd17 | ankyrin repeat domain 17 [Source:MGI Symbol;Acc:MGI:1932101] |
| 5 | protein_coding | 90608756 | 90624461 | ENSMUSG00000029368 | MGI:87991 | Alb | albumin [Source:MGI Symbol;Acc:MGI:87991] |
| 5 | protein_coding | 90638596 | 90656766 | ENSMUSG00000054932 | MGI:87951 | Afp | alpha fetoprotein [Source:MGI Symbol;Acc:MGI:87951] |
| 5 | protein_coding | 90666791 | 90701402 | ENSMUSG00000029369 | MGI:2429409 | Afm | afamin [Source:MGI Symbol;Acc:MGI:2429409] |
| 5 | protein_coding | 90708966 | 90745730 | ENSMUSG00000070690 | MGI:1923342 | Albfm1 | albumin superfamily member 1 [Source:MGI Symbol;Acc:MGI:1923342] |
| 5 | protein_coding | 90750935 | 90788516 | ENSMUSG00000029370 | MGI:1920496 | Rassf6 | Ras association (RalGDS/AF-6) domain family member 6 [Source:MGI Symbol;Acc:MGI:1920496] |
| 5 | protein_coding | 90907219 | 90909483 | ENSMUSG00000029371 | MGI:1096868 | Cxcl5 | C-X-C motif chemokine ligand 5 [Source:MGI Symbol;Acc:MGI:1096868] |

| | | | | | | | |
|---|---|---|---|---|---|---|---|
| 5 | protein_coding | 90916377 | 90917922 | ENSMUSG00000029372 | MGI:1888712 | Ppbp | pro-platelet basic protein [Source:MGI Symbol;Acc:MGI:1888712] |
| 5 | protein_coding | 90920294 | 90921242 | ENSMUSG00000029373 | MGI:1888711 | Pf4 | platelet factor 4 [Source:MGI Symbol;Acc:MGI:1888711] |
| 5 | protein_coding | 90933962 | 90937459 | ENSMUSG00000029379 | MGI:3037818 | Cxcl3 | C-X-C motif chemokine ligand 3 [Source:MGI Symbol;Acc:MGI:3037818] |
| 5 | protein_coding | 90942393 | 90950926 | ENSMUSG00000029375 | MGI:1339941 | Cxcl15 | C-X-C motif chemokine ligand 15 [Source:MGI Symbol;Acc:MGI:1339941] |
| 5 | protein_coding | 91039100 | 91040974 | ENSMUSG00000029380 | MGI:108068 | Cxcl1 | C-X-C motif chemokine ligand 1 [Source:MGI Symbol;Acc:MGI:108068] |
| 5 | protein_coding | 91051730 | 91053797 | ENSMUSG00000058427 | MGI:1340094 | Cxcl2 | C-X-C motif chemokine ligand 2 [Source:MGI Symbol;Acc:MGI:1340094] |
| 5 | protein_coding | 91078976 | 91169227 | ENSMUSG00000029376 | MGI:1915871 | Mthfd2l | methylenetetrahydrofolate dehydrogenase (NADP+ dependent) 2-like [Source:MGI Symbol;Acc:MGI:1915871] |
| 5 | protein_coding | 91175323 | 91183074 | ENSMUSG00000035020 | MGI:1919170 | Epgn | epithelial mitogen [Source:MGI Symbol;Acc:MGI:1919170] |
| 5 | protein_coding | 91222481 | 91241505 | ENSMUSG00000029377 | MGI:107508 | Ereg | epiregulin [Source:MGI Symbol;Acc:MGI:107508] |
| 5 | protein_coding | 91287458 | 91296291 | ENSMUSG00000029378 | MGI:88068 | Areg | amphiregulin [Source:MGI Symbol;Acc:MGI:88068] |
| 5 | protein_coding | 91505120 | 91550853 | ENSMUSG00000082361 | MGI:99439 | Btc | betacellulin, epidermal growth factor family member [Source:MGI Symbol;Acc:MGI:99439] |
| 5 | protein_coding | 91665474 | 91774753 | ENSMUSG00000034981 | MGI:2443349 | Parm1 | prostate androgen-regulated mucin-like protein 1 [Source:MGI Symbol;Acc:MGI:2443349] |
| 5 | protein_coding | 92096763 | 92110927 | ENSMUSG00000029397 | MGI:1915348 | Rchy1 | ring finger and CHY zinc finger domain containing 1 [Source:MGI Symbol;Acc:MGI:1915348] |
| 5 | protein_coding | 92135334 | 92143179 | ENSMUSG00000096035 | MGI:2685891 | Odaph | odontogenesis associated phosphoprotein [Source:MGI Symbol;Acc:MGI:2685891] |
| 5 | protein_coding | 92153933 | 92191742 | ENSMUSG00000029403 | MGI:1858227 | Cdkl2 | cyclin dependent kinase like 2 [Source:MGI Symbol;Acc:MGI:1858227] |
| 5 | protein_coding | 92200005 | 92231578 | ENSMUSG00000029405 | MGI:2442040 | G3bp2 | G3BP stress granule assembly factor 2 [Source:MGI Symbol;Acc:MGI:2442040] |

| | | | | | | | |
|---|---|---|---|---|---|---|---|
| 5 | protein_coding | 92285797 | 92350657 | ENSMUSG00000029407 | MGI:1929095 | Uso1 | USO1 vesicle docking factor [Source:MGI Symbol;Acc:MGI:1929095] |
| 5 | protein_coding | 92357752 | 92363267 | ENSMUSG00000029409 | MGI:1930915 | U90926 | cDNA sequence U90926 [Source:MGI Symbol;Acc:MGI:1930915] |
| 5 | protein_coding | 92374538 | 92404137 | ENSMUSG00000029410 | MGI:1342304 | Ppef2 | protein phosphatase, EF hand calcium-binding domain 2 [Source:MGI Symbol;Acc:MGI:1342304] |
| 5 | protein_coding | 92405518 | 92426029 | ENSMUSG00000029413 | MGI:1914361 | Naaa | N-acylethanolamine acid amidase [Source:MGI Symbol;Acc:MGI:1914361] |
| 5 | protein_coding | 92431869 | 92458338 | ENSMUSG00000029415 | MGI:2140779 | Sdad1 | SDA1 domain containing 1 [Source:MGI Symbol;Acc:MGI:2140779] |
| 5 | protein_coding | 92469206 | 92475938 | ENSMUSG00000029417 | MGI:1352449 | Cxcl9 | C-X-C motif chemokine ligand 9 [Source:MGI Symbol;Acc:MGI:1352449] |
| 5 | protein_coding | 92479686 | 92562487 | ENSMUSG00000034842 | MGI:1202729 | Art3 | ADP-ribosyltransferase 3 [Source:MGI Symbol;Acc:MGI:1202729] |
| 5 | protein_coding | 92494497 | 92496748 | ENSMUSG00000034855 | MGI:1352450 | Cxcl10 | C-X-C motif chemokine ligand 10 [Source:MGI Symbol;Acc:MGI:1352450] |
| 5 | protein_coding | 92507403 | 92513344 | ENSMUSG00000060183 | MGI:1860203 | Cxcl11 | chemokine (C-X-C motif) ligand 11 [Source:MGI Symbol;Acc:MGI:1860203] |
| 5 | protein_coding | 92563399 | 92583078 | ENSMUSG00000034826 | MGI:1920460 | Nup54 | nucleoporin 54 [Source:MGI Symbol;Acc:MGI:1920460] |
| 5 | protein_coding | 92589173 | 92654692 | ENSMUSG00000029426 | MGI:1196458 | Scarb2 | scavenger receptor class B, member 2 [Source:MGI Symbol;Acc:MGI:1196458] |
| 5 | protein_coding | 92702928 | 92739138 | ENSMUSG00000057068 | MGI:2686227 | Fam47e | family with sequence similarity 47, member E [Source:MGI Symbol;Acc:MGI:2686227] |
| 5 | protein_coding | 92750900 | 92754438 | ENSMUSG00000047963 | MGI:1261768 | Stbd1 | starch binding domain 1 [Source:MGI Symbol;Acc:MGI:1261768] |
| 5 | protein_coding | 92755813 | 92823130 | ENSMUSG00000050050 | MGI:2444555 | Ccdc158 | coiled-coil domain containing 158 [Source:MGI Symbol;Acc:MGI:2444555] |
| 5 | protein_coding | 92831294 | 93113177 | ENSMUSG00000029381 | MGI:1351655 | Shroom3 | shroom family member 3 [Source:MGI Symbol;Acc:MGI:1351655] |
| 5 | protein_coding | 93188982 | 93192881 | ENSMUSG00000045314 | MGI:1925338 | Sowahb | sosondowah ankyrin repeat domain family member B [Source:MGI Symbol;Acc:MGI:1925338] |
| 5 | protein_coding | 93241296 | 93324306 | ENSMUSG00000058013 | MGI:1277214 | Septin11 | septin 11 [Source:MGI Symbol;Acc:MGI:1277214] |

| | | | | | | | |
|---|---|---|---|---|---|---|---|
| 5 | protein_coding | 93329792 | 93354354 | ENSMUSG00000063015 | MGI:1341077 | Ccni | cyclin I [Source:MGI Symbol;Acc:MGI:1341077] |
| 5 | protein_coding | 93354377 | 93361334 | ENSMUSG00000029384 | MGI:1922860 | 2010109A12Rik | RIKEN cDNA 2010109A12 gene [Source:MGI Symbol;Acc:MGI:1922860] |
| 5 | protein_coding | 93415116 | 93424090 | ENSMUSG00000029385 | MGI:1095734 | Ccng2 | cyclin G2 [Source:MGI Symbol;Acc:MGI:1095734] |
| 5 | protein_coding | 93629688 | 93688059 | ENSMUSG00000096044 | MGI:3704103 | Pramel33 | PRAME like 33 [Source:MGI Symbol;Acc:MGI:3704103] |
| 5 | protein_coding | 93783056 | 93819410 | ENSMUSG00000070686 | MGI:2141341 | Pramel34 | PRAME like 34 [Source:MGI Symbol;Acc:MGI:2141341] |
| 5 | protein_coding | 93991623 | 94003966 | ENSMUSG00000107392 | MGI:3704104 | Pramel35 | PRAME like 35 [Source:MGI Symbol;Acc:MGI:3704104] |
| 5 | protein_coding | 94072636 | 94076911 | ENSMUSG00000096230 | MGI:3704105 | Pramel36 | PRAME like 36 [Source:MGI Symbol;Acc:MGI:3704105] |
| 5 | protein_coding | 94214310 | 94218404 | ENSMUSG00000072822 | MGI:2681870 | Pramel37 | PRAME like 37 [Source:MGI Symbol;Acc:MGI:2681870] |
| 5 | protein_coding | 94224244 | 94228794 | ENSMUSG00000072821 | MGI:3647817 | Pramel53 | PRAME like 53 [Source:MGI Symbol;Acc:MGI:3647817] |
| 5 | protein_coding | 94304436 | 94369394 | ENSMUSG00000096259 | MGI:3781282 | Pramel38 | PRAME like 38 [Source:MGI Symbol;Acc:MGI:3781282] |
| 5 | protein_coding | 94460534 | 94465114 | ENSMUSG00000095718 | MGI:3779601 | Pramel40 | PRAME like 40 [Source:MGI Symbol;Acc:MGI:3779601] |
| 5 | protein_coding | 94591865 | 94596410 | ENSMUSG00000074011 | MGI:3779756 | Pramel41 | PRAME like 41 [Source:MGI Symbol;Acc:MGI:3779756] |
| 5 | protein_coding | 94674146 | 94694282 | ENSMUSG00000095074 | MGI:3781318 | Pramel42 | PRAME like 42 [Source:MGI Symbol;Acc:MGI:3781318] |
| 5 | protein_coding | 94759928 | 94764201 | ENSMUSG00000095503 | MGI:3781326 | Pramel43 | PRAME like 43 [Source:MGI Symbol;Acc:MGI:3781326] |
| 5 | protein_coding | 94770074 | 94774605 | ENSMUSG00000094195 | MGI:3779602 | Pramel44 | PRAME like 44 [Source:MGI Symbol;Acc:MGI:3779602] |
| 5 | protein_coding | 95013295 | 95033247 | ENSMUSG00000095954 | MGI:3781362 | Pramel56 | PRAME like 56 [Source:MGI Symbol;Acc:MGI:3781362] |
| 5 | protein_coding | 95254563 | 95259017 | ENSMUSG00000092166 | MGI:3646599 | Pramel45 | PRAME like 45 [Source:MGI Symbol;Acc:MGI:3646599] |
| 5 | protein_coding | 95330483 | 95342825 | ENSMUSG00000095996 | MGI:2141041 | Pramel60 | PRAME like 60 [Source:MGI Symbol;Acc:MGI:2141041] |

| | | | | | | | |
|---|---|---|---|---|---|---|---|
| 5 | protein_coding | 95416352 | 95420627 | ENSMUSG00000096066 | MGI:3704250 | Pramel46 | PRAME like 46 [Source:MGI Symbol;Acc:MGI:3704250] |
| 5 | protein_coding | 95453816 | 95491776 | ENSMUSG00000079424 | MGI:3781437 | Pramel47 | PRAME like 47 [Source:MGI Symbol;Acc:MGI:3781437] |
| 5 | protein_coding | 95604665 | 95633446 | ENSMUSG00000070677 | MGI:1261918 | Pramel48 | PRAME like 48 [Source:MGI Symbol;Acc:MGI:1261918] |
| 5 | protein_coding | 95658617 | 95670261 | ENSMUSG00000079423 | MGI:3781464 | Pramel57 | PRAME like 57 [Source:MGI Symbol;Acc:MGI:3781464] |
| 5 | protein_coding | 95738458 | 95742734 | ENSMUSG00000094043 | MGI:3779772 | Pramel49 | PRAME like 49 [Source:MGI Symbol;Acc:MGI:3779772] |
| 5 | protein_coding | 95879648 | 95884032 | ENSMUSG00000096139 | MGI:3704107 | Pramel50 | PRAME like 50 [Source:MGI Symbol;Acc:MGI:3704107] |
| 5 | protein_coding | 95889873 | 95894414 | ENSMUSG00000072814 | MGI:3648891 | Pramel54 | PRAME like 54 [Source:MGI Symbol;Acc:MGI:3648891] |
| 5 | protein_coding | 95923729 | 95952089 | ENSMUSG00000072813 | MGI:3615333 | Pramel55 | PRAME like 55 [Source:MGI Symbol;Acc:MGI:3615333] |
| 5 | protein_coding | 96104810 | 96108927 | ENSMUSG00000023078 | MGI:1888499 | Cxcl13 | C-X-C motif chemokine ligand 13 [Source:MGI Symbol;Acc:MGI:1888499] |
| 5 | protein_coding | 96218192 | 96312030 | ENSMUSG00000034724 | MGI:2443154 | Cnot6l | CCR4-NOT transcription complex, subunit 6-like [Source:MGI Symbol;Acc:MGI:2443154] |
| 5 | protein_coding | 96357352 | 96414586 | ENSMUSG00000029486 | MGI:2137202 | Mrpl1 | mitochondrial ribosomal protein L1 [Source:MGI Symbol;Acc:MGI:2137202] |
| 5 | protein_coding | 96521814 | 96932587 | ENSMUSG00000034687 | MGI:2385368 | Fras1 | Fraser extracellular matrix complex subunit 1 [Source:MGI Symbol;Acc:MGI:2385368] |
| 5 | protein_coding | 96941198 | 96993825 | ENSMUSG00000029484 | MGI:1201378 | Anxa3 | annexin A3 [Source:MGI Symbol;Acc:MGI:1201378] |
| 5 | protein_coding | 97145548 | 97239726 | ENSMUSG00000034663 | MGI:2155456 | Bmp2k | BMP2 inducible kinase [Source:MGI Symbol;Acc:MGI:2155456] |
| 5 | protein_coding | 97230188 | 97259455 | ENSMUSG00000055725 | MGI:2679683 | Paqr3 | progestin and adipoQ receptor family member III [Source:MGI Symbol;Acc:MGI:2679683] |
| 5 | protein_coding | 97530057 | 97540238 | ENSMUSG00000046000 | MGI:2141314 | Naa11 | N(alpha)-acetyltransferase 11, NatA catalytic subunit [Source:MGI Symbol;Acc:MGI:2141314] |

| | | | | | | | |
|---|---|---|---|---|---|---|---|
| 5 | protein_coding | 97603001 | 97604880 | ENSMUSG00000050553 | MGI:1329027 | Gk2 | glycerol kinase 2 [Source:MGI Symbol;Acc:MGI:1329027] |
| 5 | protein_coding | 98030642 | 98178902 | ENSMUSG00000029338 | MGI:1919164 | Antxr2 | anthrax toxin receptor 2 [Source:MGI Symbol;Acc:MGI:1919164] |
| 5 | protein_coding | 98315057 | 98336850 | ENSMUSG00000035456 | MGI:1924880 | Prdm8 | PR domain containing 8 [Source:MGI Symbol;Acc:MGI:1924880] |
| 5 | protein_coding | 98402043 | 98424889 | ENSMUSG00000029337 | MGI:95519 | Fgf5 | fibroblast growth factor 5 [Source:MGI Symbol;Acc:MGI:95519] |
| 5 | protein_coding | 98477163 | 98949906 | ENSMUSG00000057816 | MGI:1916571 | Cfap299 | cilia and flagella associated protein 299 [Source:MGI Symbol;Acc:MGI:1916571] |
| 5 | protein_coding | 99002274 | 99032255 | ENSMUSG00000029335 | MGI:88179 | Bmp3 | bone morphogenetic protein 3 [Source:MGI Symbol;Acc:MGI:88179] |
| 5 | protein_coding | 99077632 | 99185210 | ENSMUSG00000029334 | MGI:108173 | Prkg2 | protein kinase, cGMP-dependent, type II [Source:MGI Symbol;Acc:MGI:108173] |
| 5 | protein_coding | 99365285 | 99876924 | ENSMUSG00000089809 | MGI:2443755 | Rasgef1b | RasGEF domain family, member 1B [Source:MGI Symbol;Acc:MGI:2443755] |
| 5 | protein_coding | 1.00E+08 | 1.00E+08 | ENSMUSG00000105078 | MGI:5595070 | Vamp9 | vesicle-associated membrane protein 9 [Source:MGI Symbol;Acc:MGI:5595070] |
| 5 | protein_coding | 1.00E+08 | 1.00E+08 | ENSMUSG00000000568 | MGI:101947 | Hnrnpd | heterogeneous nuclear ribonucleoprotein D [Source:MGI Symbol;Acc:MGI:101947] |
| 5 | protein_coding | 1.00E+08 | 1.00E+08 | ENSMUSG00000029328 | MGI:1355299 | Hnrnpdl | heterogeneous nuclear ribonucleoprotein D-like [Source:MGI Symbol;Acc:MGI:1355299] |
| 5 | protein_coding | 1.00E+08 | 1.00E+08 | ENSMUSG00000029326 | MGI:1915120 | Enoph1 | enolase-phosphatase 1 [Source:MGI Symbol;Acc:MGI:1915120] |
| 5 | protein_coding | 1.00E+08 | 1.00E+08 | ENSMUSG00000050640 | MGI:3041258 | Tmem150c | transmembrane protein 150C [Source:MGI Symbol;Acc:MGI:3041258] |
| 5 | protein_coding | 1.01E+08 | 1.01E+08 | ENSMUSG00000035325 | MGI:1916412 | Sec31a | SEC31 homolog A, COPII coat complex component [Source:MGI Symbol;Acc:MGI:1916412] |
| 5 | protein_coding | 1.01E+08 | 1.01E+08 | ENSMUSG00000118665 | MGI:2140902 | Lin54 | lin-54 DREAM MuvB core complex component [Source:MGI Symbol;Acc:MGI:2140902] |
| 5 | protein_coding | 1.01E+08 | 1.01E+08 | ENSMUSG00000035297 | MGI:1349414 | Cops4 | COP9 signalosome subunit 4 [Source:MGI Symbol;Acc:MGI:1349414] |

| | | | | | | | |
|---|---|---|---|---|---|---|---|
| 5 | protein_coding | 1.01E+08 | 1.01E+08 | ENSMUSG00000029322 | MGI:2445289 | Plac8 | placenta-specific 8 [Source:MGI Symbol;Acc:MGI:2445289] |
| 5 | protein_coding | 1.01E+08 | 1.01E+08 | ENSMUSG00000029319 | MGI:1919133 | Coq2 | coenzyme Q2 4-hydroxybenzoate polyprenyltransferase [Source:MGI Symbol;Acc:MGI:1919133] |
| 5 | protein_coding | 1.01E+08 | 1.01E+08 | ENSMUSG00000035273 | MGI:1343124 | Hpse | heparanase [Source:MGI Symbol;Acc:MGI:1343124] |
| 5 | protein_coding | 1.01E+08 | 1.01E+08 | ENSMUSG00000035266 | MGI:2176740 | Helq | helicase, POLQ-like [Source:MGI Symbol;Acc:MGI:2176740] |
| 5 | protein_coding | 1.01E+08 | 1.01E+08 | ENSMUSG00000016833 | MGI:1915985 | Mrps18c | mitochondrial ribosomal protein S18C [Source:MGI Symbol;Acc:MGI:1915985] |
| 5 | protein_coding | 1.01E+08 | 1.01E+08 | ENSMUSG00000035234 | MGI:1917931 | Abraxas1 | BRCA1 A complex subunit [Source:MGI Symbol;Acc:MGI:1917931] |
| 5 | protein_coding | 1.01E+08 | 1.01E+08 | ENSMUSG00000029314 | MGI:3603816 | Gpat3 | glycerol-3-phosphate acyltransferase 3 [Source:MGI Symbol;Acc:MGI:3603816] |
| 5 | protein_coding | 1.02E+08 | 1.02E+08 | ENSMUSG00000035187 | MGI:1206039 | Nkx6-1 | NK6 homeobox 1 [Source:MGI Symbol;Acc:MGI:1206039] |
| 5 | protein_coding | 1.02E+08 | 1.02E+08 | ENSMUSG00000029330 | MGI:1921846 | Cds1 | CDP-diacylglycerol synthase 1 [Source:MGI Symbol;Acc:MGI:1921846] |
| 5 | protein_coding | 1.02E+08 | 1.02E+08 | ENSMUSG00000043940 | MGI:1096875 | Wdfy3 | WD repeat and FYVE domain containing 3 [Source:MGI Symbol;Acc:MGI:1096875] |
| 5 | protein_coding | 1.03E+08 | 1.03E+08 | ENSMUSG00000057315 | MGI:1922647 | Arhgap24 | Rho GTPase activating protein 24 [Source:MGI Symbol;Acc:MGI:1922647] |
| 5 | protein_coding | 1.03E+08 | 1.03E+08 | ENSMUSG00000046709 | MGI:1346863 | Mapk10 | mitogen-activated protein kinase 10 [Source:MGI Symbol;Acc:MGI:1346863] |
| 5 | protein_coding | 1.04E+08 | 1.04E+08 | ENSMUSG00000034573 | MGI:103293 | Ptpn13 | protein tyrosine phosphatase, non-receptor type 13 [Source:MGI Symbol;Acc:MGI:103293] |
| 5 | protein_coding | 1.04E+08 | 1.04E+08 | ENSMUSG00000029321 | MGI:1923000 | Slc10a6 | solute carrier family 10 (sodium/bile acid cotransporter family), member 6 [Source:MGI Symbol;Acc:MGI:1923000] |
| 5 | protein_coding | 1.04E+08 | 1.04E+08 | ENSMUSG00000029320 | MGI:1921468 | 1700016H13Rik | RIKEN cDNA 1700016H13 gene [Source:MGI Symbol;Acc:MGI:1921468] |
| 5 | protein_coding | 1.04E+08 | 1.04E+08 | ENSMUSG00000029313 | MGI:1100819 | Aff1 | AF4/FMR2 family, member 1 [Source:MGI Symbol;Acc:MGI:1100819] |

| | | | | | | | |
|---|---|---|---|---|---|---|---|
| 5 | protein_coding | 1.04E+08 | 1.04E+08 | ENSMUSG00000029312 | MGI:2179430 | Klhl8 | kelch-like 8 [Source:MGI Symbol;Acc:MGI:2179430] |
| 5 | protein_coding | 1.04E+08 | 1.04E+08 | ENSMUSG00000034528 | MGI:2140804 | Hsd17b13 | hydroxysteroid (17-beta) dehydrogenase 13 [Source:MGI Symbol;Acc:MGI:2140804] |
| 5 | protein_coding | 1.04E+08 | 1.04E+08 | ENSMUSG00000029311 | MGI:2149821 | Hsd17b11 | hydroxysteroid (17-beta) dehydrogenase 11 [Source:MGI Symbol;Acc:MGI:2149821] |
| 5 | protein_coding | 1.04E+08 | 1.04E+08 | ENSMUSG00000029310 | MGI:1921417 | Nudt9 | nudix hydrolase 9 [Source:MGI Symbol;Acc:MGI:1921417] |
| 5 | protein_coding | 1.04E+08 | 1.04E+08 | ENSMUSG00000091034 | MGI:4937294 | Scpppq1 | secretory calcium-binding phosphoprotein proline-glutamine rich 1 [Source:MGI Symbol;Acc:MGI:4937294] |
| 5 | protein_coding | 1.04E+08 | 1.04E+08 | ENSMUSG00000029309 | MGI:108110 | Sparcl1 | SPARC-like 1 [Source:MGI Symbol;Acc:MGI:108110] |
| 5 | protein_coding | 1.04E+08 | 1.04E+08 | ENSMUSG00000053268 | MGI:109172 | Dspp | dentin sialophosphoprotein [Source:MGI Symbol;Acc:MGI:109172] |
| 5 | protein_coding | 1.04E+08 | 1.04E+08 | ENSMUSG00000029307 | MGI:94910 | Dmp1 | dentin matrix protein 1 [Source:MGI Symbol;Acc:MGI:94910] |
| 5 | protein_coding | 1.04E+08 | 1.04E+08 | ENSMUSG00000029306 | MGI:96389 | Ibsp | integrin binding sialoprotein [Source:MGI Symbol;Acc:MGI:96389] |
| 5 | protein_coding | 1.04E+08 | 1.04E+08 | ENSMUSG00000053863 | MGI:2137384 | Mepe | matrix extracellular phosphoglycoprotein with ASARM motif (bone) [Source:MGI Symbol;Acc:MGI:2137384] |
| 5 | protein_coding | 1.05E+08 | 1.05E+08 | ENSMUSG00000029304 | MGI:98389 | Spp1 | secreted phosphoprotein 1 [Source:MGI Symbol;Acc:MGI:98389] |
| 5 | protein_coding | 1.05E+08 | 1.05E+08 | ENSMUSG00000034462 | MGI:1099818 | Pkd2 | polycystin 2, transient receptor potential cation channel [Source:MGI Symbol;Acc:MGI:1099818] |
| 5 | protein_coding | 1.05E+08 | 1.05E+08 | ENSMUSG00000097392 | MGI:3040669 | Thoc2l | THO complex subunit 2-like [Source:MGI Symbol;Acc:MGI:3040669] |
| 5 | protein_coding | 1.05E+08 | 1.05E+08 | ENSMUSG00000120166 | MGI:1916873 | Zfp33b | zinc finger protein 33B [Source:MGI Symbol;Acc:MGI:1916873] |
| 5 | protein_coding | 1.05E+08 | 1.05E+08 | ENSMUSG00000072774 | MGI:2441896 | Zfp951 | zinc finger protein 951 [Source:MGI Symbol;Acc:MGI:2441896] |

| | | | | | | | |
|---|---|---|---|---|---|---|---|
| 5 | protein_coding | 1.05E+08 | 1.05E+08 | ENSMUSG00000029299 | MGI:1351624 | Abcg3 | ATP binding cassette subfamily G member 3 [Source:MGI Symbol;Acc:MGI:1351624] |
| 5 | protein_coding | 1.05E+08 | 1.05E+08 | ENSMUSG00000034438 | MGI:1923324 | Gbp8 | guanylate-binding protein 8 [Source:MGI Symbol;Acc:MGI:1923324] |
| 5 | protein_coding | 1.05E+08 | 1.05E+08 | ENSMUSG00000029298 | MGI:3605620 | Gbp9 | guanylate-binding protein 9 [Source:MGI Symbol;Acc:MGI:3605620] |
| 5 | protein_coding | 1.05E+08 | 1.05E+08 | ENSMUSG00000079363 | MGI:97072 | Gbp4 | guanylate binding protein 4 [Source:MGI Symbol;Acc:MGI:97072] |
| 5 | protein_coding | 1.05E+08 | 1.05E+08 | ENSMUSG00000079362 | MGI:5663439 | Gm43302 | predicted gene 43302 [Source:MGI Symbol;Acc:MGI:5663439] |
| 5 | protein_coding | 1.05E+08 | 1.05E+08 | ENSMUSG00000105096 | MGI:4359647 | Gbp10 | guanylate-binding protein 10 [Source:MGI Symbol;Acc:MGI:4359647] |
| 5 | protein_coding | 1.05E+08 | 1.05E+08 | ENSMUSG00000104713 | MGI:2140937 | Gbp6 | guanylate binding protein 6 [Source:MGI Symbol;Acc:MGI:2140937] |
| 5 | protein_coding | 1.05E+08 | 1.05E+08 | ENSMUSG00000092021 | MGI:3646307 | Gbp11 | guanylate binding protein 11 [Source:MGI Symbol;Acc:MGI:3646307] |
| 5 | protein_coding | 1.06E+08 | 1.06E+08 | ENSMUSG00000070639 | MGI:2141353 | Lrrc8b | leucine rich repeat containing 8 family, member B [Source:MGI Symbol;Acc:MGI:2141353] |
| 5 | protein_coding | 1.06E+08 | 1.06E+08 | ENSMUSG00000054720 | MGI:2140839 | Lrrc8c | leucine rich repeat containing 8 family, member C [Source:MGI Symbol;Acc:MGI:2140839] |
| 5 | protein_coding | 1.06E+08 | 1.06E+08 | ENSMUSG00000046079 | MGI:1922368 | Lrrc8d | leucine rich repeat containing 8D [Source:MGI Symbol;Acc:MGI:1922368] |
| 5 | protein_coding | 1.06E+08 | 1.06E+08 | ENSMUSG00000029290 | MGI:1927246 | Zfp326 | zinc finger protein 326 [Source:MGI Symbol;Acc:MGI:1927246] |
| 5 | protein_coding | 1.07E+08 | 1.07E+08 | ENSMUSG00000034384 | MGI:1859314 | Barhl2 | BarH like homeobox 2 [Source:MGI Symbol;Acc:MGI:1859314] |
| 5 | protein_coding | 1.07E+08 | 1.07E+08 | ENSMUSG00000049606 | MGI:1277212 | Zfp644 | zinc finger protein 644 [Source:MGI Symbol;Acc:MGI:1277212] |
| 5 | protein_coding | 1.07E+08 | 1.07E+08 | ENSMUSG00000043410 | MGI:3036246 | Hfm1 | HFM1, ATP-dependent DNA helicase homolog [Source:MGI Symbol;Acc:MGI:3036246] |
| 5 | protein_coding | 1.07E+08 | 1.07E+08 | ENSMUSG00000029283 | MGI:1309511 | Cdc7 | cell division cycle 7 [Source:MGI Symbol;Acc:MGI:1309511] |

| | | | | | | | |
|---|---|---|---|---|---|---|---|
| 5 | protein_coding | 1.07E+08 | 1.07E+08 | ENSMUSG00000029287 | MGI:104637 | Tgfbr3 | transforming growth factor, beta receptor III [Source:MGI Symbol;Acc:MGI:104637] |
| 5 | protein_coding | 1.07E+08 | 1.08E+08 | ENSMUSG00000029279 | MGI:1891374 | Brdt | bromodomain, testis-specific [Source:MGI Symbol;Acc:MGI:1891374] |
| 5 | protein_coding | 1.08E+08 | 1.08E+08 | ENSMUSG00000033805 | MGI:2686228 | Ephx4 | epoxide hydrolase 4 [Source:MGI Symbol;Acc:MGI:2686228] |
| 5 | protein_coding | 1.08E+08 | 1.08E+08 | ENSMUSG00000033794 | MGI:1918152 | Lpcat2b | lysophosphatidylcholine acyltransferase 2B [Source:MGI Symbol;Acc:MGI:1918152] |
| 5 | protein_coding | 1.08E+08 | 1.08E+08 | ENSMUSG00000111375 | MGI:3646208 | Btbd8 | BTB domain containing 8 [Source:MGI Symbol;Acc:MGI:3646208] |
| 5 | protein_coding | 1.08E+08 | 1.08E+08 | ENSMUSG00000106631 | MGI:5662806 | Gm42669 | predicted gene 42669 [Source:MGI Symbol;Acc:MGI:5662806] |
| 5 | protein_coding | 1.08E+08 | 1.08E+08 | ENSMUSG00000089798 | MGI:1923671 | 1700028K03Rik | RIKEN cDNA 1700028K03 gene [Source:MGI Symbol;Acc:MGI:1923671] |
| 5 | protein_coding | 1.08E+08 | 1.08E+08 | ENSMUSG00000029276 | MGI:2141180 | Glmn | glomulin, FKBP associated protein [Source:MGI Symbol;Acc:MGI:2141180] |
| 5 | protein_coding | 1.08E+08 | 1.08E+08 | ENSMUSG00000033773 | MGI:2141142 | Rpap2 | RNA polymerase II associated protein 2 [Source:MGI Symbol;Acc:MGI:2141142] |
| 5 | protein_coding | 1.08E+08 | 1.08E+08 | ENSMUSG00000029275 | MGI:103170 | Gfi1 | growth factor independent 1 transcription repressor [Source:MGI Symbol;Acc:MGI:103170] |
| 5 | protein_coding | 1.08E+08 | 1.08E+08 | ENSMUSG00000011831 | MGI:104736 | Evi5 | ecotropic viral integration site 5 [Source:MGI Symbol;Acc:MGI:104736] |
| 5 | protein_coding | 1.08E+08 | 1.08E+08 | ENSMUSG00000063447 | MGI:1920568 | Ube2d2b | ubiquitin-conjugating enzyme E2D 2B [Source:MGI Symbol;Acc:MGI:1920568] |
| 5 | protein_coding | 1.08E+08 | 1.08E+08 | ENSMUSG00000058558 | MGI:102854 | Rpl5 | ribosomal protein L5 [Source:MGI Symbol;Acc:MGI:102854] |
| 5 | protein_coding | 1.08E+08 | 1.08E+08 | ENSMUSG00000029270 | MGI:1914516 | Dipk1a | divergent protein kinase domain 1A [Source:MGI Symbol;Acc:MGI:1914516] |
| 5 | protein_coding | 1.08E+08 | 1.08E+08 | ENSMUSG00000029267 | MGI:105050 | Mtf2 | metal response element binding transcription factor 2 [Source:MGI Symbol;Acc:MGI:105050] |
| 5 | protein_coding | 1.08E+08 | 1.08E+08 | ENSMUSG00000063406 | MGI:1921586 | Tmed5 | transmembrane p24 trafficking protein 5 [Source:MGI Symbol;Acc:MGI:1921586] |

| | | | | | | | |
|---|---|---|---|---|---|---|---|
| 5 | protein_coding | 1.08E+08 | 1.08E+08 | ENSMUSG00000056531 | MGI:1922974 | Ccdc18 | coiled-coil domain containing 18 [Source:MGI Symbol;Acc:MGI:1922974] |
| 5 | protein_coding | 1.08E+08 | 1.08E+08 | ENSMUSG00000029265 | MGI:1100515 | Dr1 | down-regulator of transcription 1 [Source:MGI Symbol;Acc:MGI:1100515] |
| 5 | protein_coding | 1.08E+08 | 1.08E+08 | ENSMUSG00000029263 | MGI:3576484 | Pigg | phosphatidylinositol glycan anchor biosynthesis, class G [Source:MGI Symbol;Acc:MGI:3576484] |

**Supplementary Table 5. All annotated genes within the 95% Bayesian credible interval for the pathology QTL on chromosome 5**

| Chr | type | start | stop | gene_id | mgi_type | gene_symbol | description |
|---|---|---|---|---|---|---|---|
| 5 | lncRNA | 41161921 | 41172181 | ENSMUSG00000130164 | MGI:7826136 | Gm71684 | predicted gene, 71684 [Source:MGI Symbol;Acc:MGI:7826136] |
| 5 | lncRNA | 41274860 | 41358312 | ENSMUSG00000132413 | MGI:7826738 | Gm71986 | predicted gene, 71986 [Source:MGI Symbol;Acc:MGI:7826738] |
| 5 | lncRNA | 41350377 | 41406253 | ENSMUSG00000132446 | MGI:7826782 | Gm72008 | predicted gene, 72008 [Source:MGI Symbol;Acc:MGI:7826782] |
| 5 | lncRNA | 41765931 | 41771759 | ENSMUSG00000133947 | MGI:5594539 | Gm35380 | predicted gene, 35380 [Source:MGI Symbol;Acc:MGI:5594539] |
| 5 | lncRNA | 41771474 | 41774854 | ENSMUSG00000133992 | MGI:7827302 | Gm72268 | predicted gene, 72268 [Source:MGI Symbol;Acc:MGI:7827302] |
| 5 | lncRNA | 41864464 | 41864958 | ENSMUSG00000134029 | MGI:7827314 | Gm72274 | predicted gene, 72274 [Source:MGI Symbol;Acc:MGI:7827314] |
| 5 | lncRNA | 41865579 | 41893410 | ENSMUSG00000133906 | MGI:7827252 | Gm72243 | predicted gene, 72243 [Source:MGI Symbol;Acc:MGI:7827252] |
| 5 | lncRNA | 42001664 | 42010131 | ENSMUSG00000131938 | MGI:7826668 | Gm71951 | predicted gene, 71951 [Source:MGI Symbol;Acc:MGI:7826668] |
| 5 | lncRNA | 42045334 | 42374141 | ENSMUSG00000067285 | MGI:3648966 | Gm16223 | predicted gene 16223 [Source:MGI Symbol;Acc:MGI:3648966] |
| 5 | lncRNA | 42397049 | 42400594 | ENSMUSG00000135933 | MGI:7827798 | Gm72517 | predicted gene, 72517 [Source:MGI Symbol;Acc:MGI:7827798] |
| 5 | lncRNA | 42590654 | 42658143 | ENSMUSG00000122969 | MGI:7825344 | Gm71283 | predicted gene, 71283 [Source:MGI Symbol;Acc:MGI:7825344] |
| 5 | lncRNA | 42693920 | 42733451 | ENSMUSG00000142458 | MGI:7829582 | Gm73412 | predicted gene, 73412 [Source:MGI Symbol;Acc:MGI:7829582] |
| 5 | lncRNA | 42752178 | 42843407 | ENSMUSG00000135200 | MGI:6723610 | Gm54295 | predicted gene, 54295 [Source:MGI Symbol;Acc:MGI:6723610] |
| 5 | lncRNA | 42929725 | 42949087 | ENSMUSG00000131751 | MGI:7826620 | Gm71927 | predicted gene, 71927 [Source:MGI Symbol;Acc:MGI:7826620] |
| 5 | lncRNA | 43120883 | 43279559 | ENSMUSG00000120713 | MGI:7338483 | Gm57285 | predicted gene, 57285 [Source:MGI Symbol;Acc:MGI:7338483] |
| 5 | lncRNA | 43183885 | 43195679 | ENSMUSG00000107193 | MGI:5663837 | Gm43700 | predicted gene 43700 [Source:MGI Symbol;Acc:MGI:5663837] |
| 5 | lncRNA | 43197080 | 43201404 | ENSMUSG00000106501 | MGI:5662848 | Gm42711 | predicted gene 42711 [Source:MGI Symbol;Acc:MGI:5662848] |

| | | | | | | | |
|---|---|---|---|---|---|---|---|
| 5 | lncRNA | 43206900 | 43217584 | ENSMUSG00000125363 | MGI:7825611 | Gm71419 | predicted gene, 71419 [Source:MGI Symbol;Acc:MGI:7825611] |
| 5 | lncRNA | 43208242 | 43287854 | ENSMUSG00000125280 | MGI:7825603 | Gm71415 | predicted gene, 71415 [Source:MGI Symbol;Acc:MGI:7825603] |
| 5 | lncRNA | 43230866 | 43232064 | ENSMUSG00000125404 | MGI:7825619 | Gm71423 | predicted gene, 71423 [Source:MGI Symbol;Acc:MGI:7825619] |
| 5 | lncRNA | 43306203 | 43392804 | ENSMUSG00000085720 | MGI:3648234 | Gm7854 | predicted gene 7854 [Source:MGI Symbol;Acc:MGI:3648234] |
| 5 | lncRNA | 43388856 | 43390471 | ENSMUSG00000120663 | MGI:7338471 | Gm57279 | predicted gene, 57279 [Source:MGI Symbol;Acc:MGI:7338471] |
| 5 | lncRNA | 43454674 | 43461022 | ENSMUSG00000137961 | MGI:7828504 | Gm72871 | predicted gene, 72871 [Source:MGI Symbol;Acc:MGI:7828504] |
| 5 | lncRNA | 43496505 | 43511399 | ENSMUSG00000137893 | MGI:7828492 | Gm72865 | predicted gene, 72865 [Source:MGI Symbol;Acc:MGI:7828492] |
| 5 | lncRNA | 43508808 | 43514761 | ENSMUSG00000104846 | MGI:5663157 | Gm43020 | predicted gene 43020 [Source:MGI Symbol;Acc:MGI:5663157] |
| 5 | lncRNA | 43533790 | 43536980 | ENSMUSG00000107244 | MGI:5663836 | Gm43699 | predicted gene 43699 [Source:MGI Symbol;Acc:MGI:5663836] |
| 5 | lncRNA | 43579093 | 43580575 | ENSMUSG00000140860 | MGI:7829236 | Gm73239 | predicted gene, 73239 [Source:MGI Symbol;Acc:MGI:7829236] |
| 5 | lncRNA | 43628035 | 43633199 | ENSMUSG00000104691 | MGI:5663158 | Gm43021 | predicted gene 43021 [Source:MGI Symbol;Acc:MGI:5663158] |
| 5 | lncRNA | 43707271 | 43759073 | ENSMUSG00000087290 | MGI:3801874 | Gm15866 | predicted gene 15866 [Source:MGI Symbol;Acc:MGI:3801874] |
| 5 | lncRNA | 43942993 | 43943494 | ENSMUSG00000133185 | MGI:7827010 | Gm72122 | predicted gene, 72122 [Source:MGI Symbol;Acc:MGI:7827010] |
| 5 | lncRNA | 43965047 | 43966085 | ENSMUSG00000104701 | MGI:5662692 | Gm42555 | predicted gene 42555 [Source:MGI Symbol;Acc:MGI:5662692] |
| 5 | lncRNA | 44018654 | 44021803 | ENSMUSG00000133570 | MGI:7827162 | Gm72198 | predicted gene, 72198 [Source:MGI Symbol;Acc:MGI:7827162] |
| 5 | lncRNA | 44028865 | 44029335 | ENSMUSG00000133608 | MGI:7827164 | Gm72199 | predicted gene, 72199 [Source:MGI Symbol;Acc:MGI:7827164] |
| 5 | lncRNA | 44033372 | 44034646 | ENSMUSG00000086479 | MGI:3801812 | Gm16014 | predicted gene 16014 [Source:MGI Symbol;Acc:MGI:3801812] |
| 5 | lncRNA | 44055929 | 44105704 | ENSMUSG00000087281 | MGI:3801811 | Gm16015 | predicted gene 16015 [Source:MGI Symbol;Acc:MGI:3801811] |

| | | | | | | | |
|---|---|---|---|---|---|---|---|
| 5 | lncRNA | 44144508 | 44162382 | ENSMUSG00000139432 | MGI:7828900 | Gm73071 | predicted gene, 73071 [Source:MGI Symbol;Acc:MGI:7828900] |
| 5 | lncRNA | 44179180 | 44182741 | ENSMUSG00000139558 | MGI:7828932 | Gm73087 | predicted gene, 73087 [Source:MGI Symbol;Acc:MGI:7828932] |
| 5 | lncRNA | 44225540 | 44248196 | ENSMUSG00000105264 | MGI:5662853 | Gm42716 | predicted gene 42716 [Source:MGI Symbol;Acc:MGI:5662853] |
| 5 | lncRNA | 44257784 | 44265916 | ENSMUSG00000072962 | MGI:3648398 | Gm16401 | predicted gene 16401 [Source:MGI Symbol;Acc:MGI:3648398] |
| 5 | lncRNA | 44263044 | 44271634 | ENSMUSG00000139480 | MGI:5595059 | Gm35900 | predicted gene, 35900 [Source:MGI Symbol;Acc:MGI:5595059] |
| 5 | lncRNA | 44293962 | 44294263 | ENSMUSG00000105353 | MGI:5662565 | Gm42428 | predicted gene 42428 [Source:MGI Symbol;Acc:MGI:5662565] |
| 5 | lncRNA | 44377025 | 44381345 | ENSMUSG00000105503 | MGI:5663119 | Gm42982 | predicted gene 42982 [Source:MGI Symbol;Acc:MGI:5663119] |
| 5 | lncRNA | 44383929 | 44440444 | ENSMUSG00000104784 | MGI:5663121 | Gm42984 | predicted gene 42984 [Source:MGI Symbol;Acc:MGI:5663121] |
| 5 | lncRNA | 44390266 | 44391130 | ENSMUSG00000136242 | MGI:7827904 | Gm72570 | predicted gene, 72570 [Source:MGI Symbol;Acc:MGI:7827904] |
| 5 | lncRNA | 44417198 | 44419558 | ENSMUSG00000136206 | MGI:7827878 | Gm72557 | predicted gene, 72557 [Source:MGI Symbol;Acc:MGI:7827878] |
| 5 | lncRNA | 44496607 | 44511899 | ENSMUSG00000124218 | MGI:5623181 | Gm40296 | predicted gene, 40296 [Source:MGI Symbol;Acc:MGI:5623181] |
| 5 | lncRNA | 44570134 | 44589024 | ENSMUSG00000104800 | MGI:5595176 | Gm36017 | predicted gene, 36017 [Source:MGI Symbol;Acc:MGI:5595176] |
| 5 | lncRNA | 44589098 | 44593903 | ENSMUSG00000133239 | MGI:5623182 | Gm40297 | predicted gene, 40297 [Source:MGI Symbol;Acc:MGI:5623182] |
| 5 | lncRNA | 44798826 | 44835480 | ENSMUSG00000104894 | MGI:5663644 | Gm43507 | predicted gene 43507 [Source:MGI Symbol;Acc:MGI:5663644] |
| 5 | lncRNA | 44858218 | 44864232 | ENSMUSG00000105224 | MGI:3781542 | Gm3364 | predicted gene 3364 [Source:MGI Symbol;Acc:MGI:3781542] |
| 5 | lncRNA | 45022196 | 45034887 | ENSMUSG00000106656 | MGI:5662887 | Gm42750 | predicted gene 42750 [Source:MGI Symbol;Acc:MGI:5662887] |
| 5 | lncRNA | 45075134 | 45085881 | ENSMUSG00000122500 | MGI:5595235 | Gm36076 | predicted gene, 36076 [Source:MGI Symbol;Acc:MGI:5595235] |
| 5 | lncRNA | 45123525 | 45136147 | ENSMUSG00000062496 | MGI:1923033 | 4930431F12Rik | RIKEN cDNA 4930431F12 gene [Source:MGI Symbol;Acc:MGI:1923033] |

| | | | | | | | |
|---|---|---|---|---|---|---|---|
| 5 | lncRNA | 45166513 | 45581744 | ENSMUSG00000029092 | MGI:1277236 | D5Ertd615e | expressed [Source:MGI Symbol;Acc:MGI:1277236] |
| 5 | lncRNA | 45320381 | 45332525 | ENSMUSG00000139739 | MGI:5826549 | Gm46912 | predicted gene, 46912 [Source:MGI Symbol;Acc:MGI:5826549] |
| 5 | lncRNA | 45577426 | 45605217 | ENSMUSG00000126761 | MGI:7825753 | Gm71490 | predicted gene, 71490 [Source:MGI Symbol;Acc:MGI:7825753] |
| 5 | lncRNA | 45591731 | 45607480 | ENSMUSG00000105255 | MGI:5648986 | Gm42413 | predicted gene, 42413 [Source:MGI Symbol;Acc:MGI:5648986] |
| 5 | lncRNA | 45626025 | 45650599 | ENSMUSG00000125577 | MGI:7825639 | Gm71433 | predicted gene, 71433 [Source:MGI Symbol;Acc:MGI:7825639] |
| 5 | lncRNA | 45796041 | 45802811 | ENSMUSG00000097825 | MGI:2441755 | 9630001P10Rik | RIKEN cDNA 9630001P10 gene [Source:MGI Symbol;Acc:MGI:2441755] |
| 5 | lncRNA | 45815993 | 45820078 | ENSMUSG00000130071 | MGI:7826118 | Gm71675 | predicted gene, 71675 [Source:MGI Symbol;Acc:MGI:7826118] |
| 5 | lncRNA | 46013440 | 46016107 | ENSMUSG00000130050 | MGI:7826114 | Gm71672 | predicted gene, 71672 [Source:MGI Symbol;Acc:MGI:7826114] |
| 5 | lncRNA | 46084153 | 46089779 | ENSMUSG00000106602 | MGI:1925356 | 4930405L22Rik | RIKEN cDNA 4930405L22 gene [Source:MGI Symbol;Acc:MGI:1925356] |
| 5 | lncRNA | 46505008 | 46572771 | ENSMUSG00000140766 | | | predicted pseudogene 7931 [Source:NCBI gene (formerly Entrezgene);Acc:100503799] |
| 5 | lncRNA | 47138572 | 47146867 | ENSMUSG00000124320 | MGI:7825519 | Gm71372 | predicted gene, 71372 [Source:MGI Symbol;Acc:MGI:7825519] |
| 5 | lncRNA | 47344395 | 47376820 | ENSMUSG00000126799 | MGI:7825755 | Gm71491 | predicted gene, 71491 [Source:MGI Symbol;Acc:MGI:7825755] |
| 5 | lncRNA | 47547347 | 47549721 | ENSMUSG00000133558 | MGI:7827158 | Gm72196 | predicted gene, 72196 [Source:MGI Symbol;Acc:MGI:7827158] |
| 5 | lncRNA | 47938737 | 48086939 | ENSMUSG00000128126 | MGI:7825857 | Gm71542 | predicted gene, 71542 [Source:MGI Symbol;Acc:MGI:7825857] |
| 5 | lncRNA | 48145374 | 48163071 | ENSMUSG00000092309 | MGI:1922330 | 4930518C09Rik | RIKEN cDNA 4930518C09 gene [Source:MGI Symbol;Acc:MGI:1922330] |
| 5 | lncRNA | 48163099 | 48165280 | ENSMUSG00000092559 | MGI:5141940 | Gm20475 | predicted gene 20475 [Source:MGI Symbol;Acc:MGI:5141940] |
| 5 | lncRNA | 48748231 | 48757336 | ENSMUSG00000093502 | MGI:5313147 | Gm20700 | predicted gene 20700 [Source:MGI Symbol;Acc:MGI:5313147] |
| 5 | lncRNA | 48956749 | 48973648 | ENSMUSG00000134568 | MGI:7827460 | Gm72347 | predicted gene, 72347 [Source:MGI Symbol;Acc:MGI:7827460] |

| | | | | | | | |
|---|---|---|---|---|---|---|---|
| 5 | lncRNA | 49370218 | 49474233 | ENSMUSG00000141583 | MGI:7829400 | Gm73321 | predicted gene, 73321 [Source:MGI Symbol;Acc:MGI:7829400] |
| 5 | lncRNA | 49474259 | 49532743 | ENSMUSG00000134740 | MGI:7827500 | Gm72367 | predicted gene, 72367 [Source:MGI Symbol;Acc:MGI:7827500] |
| 5 | lncRNA | 49720350 | 49859749 | ENSMUSG00000129598 | MGI:7826001 | Gm71615 | predicted gene, 71615 [Source:MGI Symbol;Acc:MGI:7826001] |
| 5 | lncRNA | 49762811 | 49772775 | ENSMUSG00000129635 | MGI:5623184 | Gm40299 | predicted gene, 40299 [Source:MGI Symbol;Acc:MGI:5623184] |
| 5 | lncRNA | 50217951 | 50230629 | ENSMUSG00000126135 | MGI:5595861 | Gm36702 | predicted gene, 36702 [Source:MGI Symbol;Acc:MGI:5595861] |
| 5 | lncRNA | 50308644 | 50365877 | ENSMUSG00000104870 | MGI:1921215 | 4930448I18Rik | RIKEN cDNA 4930448I18 gene [Source:MGI Symbol;Acc:MGI:1921215] |
| 5 | lncRNA | 50426239 | 50675024 | ENSMUSG00000126829 | | | novel transcript |
| 5 | lncRNA | 50439215 | 50513096 | ENSMUSG00000126863 | MGI:7825759 | Gm71493 | predicted gene, 71493 [Source:MGI Symbol;Acc:MGI:7825759] |
| 5 | lncRNA | 50557102 | 50564135 | ENSMUSG00000126932 | MGI:7825771 | Gm71499 | predicted gene, 71499 [Source:MGI Symbol;Acc:MGI:7825771] |
| 5 | lncRNA | 50606004 | 50701876 | ENSMUSG00000105423 | MGI:5595973 | Gm36814 | predicted gene, 36814 [Source:MGI Symbol;Acc:MGI:5595973] |
| 5 | lncRNA | 50700780 | 50980956 | ENSMUSG00000105320 | MGI:5663153 | Gm43016 | predicted gene 43016 [Source:MGI Symbol;Acc:MGI:5663153] |
| 5 | lncRNA | 50995647 | 50996695 | ENSMUSG00000126009 | MGI:7825675 | Gm71451 | predicted gene, 71451 [Source:MGI Symbol;Acc:MGI:7825675] |
| 5 | lncRNA | 51100591 | 51101634 | ENSMUSG00000122533 | MGI:7825273 | Gm71246 | predicted gene, 71246 [Source:MGI Symbol;Acc:MGI:7825273] |
| 5 | lncRNA | 51124509 | 51202427 | ENSMUSG00000124531 | MGI:7825539 | Gm71383 | predicted gene, 71383 [Source:MGI Symbol;Acc:MGI:7825539] |
| 5 | lncRNA | 51435063 | 51446696 | ENSMUSG00000138939 | MGI:5589083 | Gm29924 | predicted gene, 29924 [Source:MGI Symbol;Acc:MGI:5589083] |
| 5 | lncRNA | 51720599 | 51782721 | ENSMUSG00000129179 | MGI:7825961 | Gm71594 | predicted gene, 71594 [Source:MGI Symbol;Acc:MGI:7825961] |
| 5 | lncRNA | 52088024 | 52091945 | ENSMUSG00000130279 | MGI:7826186 | Gm71709 | predicted gene, 71709 [Source:MGI Symbol;Acc:MGI:7826186] |
| 5 | lncRNA | 52193982 | 52198110 | ENSMUSG00000125968 | MGI:7825671 | Gm71449 | predicted gene, 71449 [Source:MGI Symbol;Acc:MGI:7825671] |

| | | | | | | | |
|---|---|---|---|---|---|---|---|
| 5 | lncRNA | 52269346 | 52270573 | ENSMUSG00000133149 | MGI:7826992 | Gm72113 | predicted gene, 72113 [Source:MGI Symbol;Acc:MGI:7826992] |
| 5 | lncRNA | 52271803 | 52273045 | ENSMUSG00000126882 | MGI:7825763 | Gm71495 | predicted gene, 71495 [Source:MGI Symbol;Acc:MGI:7825763] |
| 5 | lncRNA | 52272010 | 52289297 | ENSMUSG00000126817 | MGI:5623187 | Gm40302 | predicted gene, 40302 [Source:MGI Symbol;Acc:MGI:5623187] |
| 5 | lncRNA | 52273209 | 52275868 | ENSMUSG00000104847 | MGI:5663314 | Gm43177 | predicted gene 43177 [Source:MGI Symbol;Acc:MGI:5663314] |
| 5 | lncRNA | 52285971 | 52286585 | ENSMUSG00000126918 | MGI:7825767 | Gm71497 | predicted gene, 71497 [Source:MGI Symbol;Acc:MGI:7825767] |
| 5 | lncRNA | 52347984 | 52496385 | ENSMUSG00000097145 | MGI:3041212 | 9230114K14Rik | RIKEN cDNA 9230114K14 gene [Source:MGI Symbol;Acc:MGI:3041212] |
| 5 | lncRNA | 52348205 | 52349529 | ENSMUSG00000124543 | MGI:7825541 | Gm71384 | predicted gene, 71384 [Source:MGI Symbol;Acc:MGI:7825541] |
| 5 | lncRNA | 52357326 | 52373776 | ENSMUSG00000106464 | MGI:3041216 | C130083M11Rik | RIKEN cDNA C130083M11 gene [Source:MGI Symbol;Acc:MGI:3041216] |
| 5 | lncRNA | 52499428 | 52499702 | ENSMUSG00000104751 | MGI:5663316 | Gm43179 | predicted gene 43179 [Source:MGI Symbol;Acc:MGI:5663316] |
| 5 | lncRNA | 52512028 | 52513002 | ENSMUSG00000124928 | MGI:7825579 | Gm71403 | predicted gene, 71403 [Source:MGI Symbol;Acc:MGI:7825579] |
| 5 | lncRNA | 52598143 | 52606135 | ENSMUSG00000106380 | MGI:3781696 | Gm3519 | predicted gene 3519 [Source:MGI Symbol;Acc:MGI:3781696] |
| 5 | lncRNA | 52633807 | 52639338 | ENSMUSG00000143439 | MGI:7829810 | Gm73527 | predicted gene, 73527 [Source:MGI Symbol;Acc:MGI:7829810] |
| 5 | lncRNA | 52696173 | 52697421 | ENSMUSG00000104829 | MGI:5663822 | Gm43685 | predicted gene 43685 [Source:MGI Symbol;Acc:MGI:5663822] |
| 5 | lncRNA | 52744234 | 52776414 | ENSMUSG00000052295 | MGI:2443206 | 8030423F21Rik | RIKEN cDNA 8030423F21 gene [Source:MGI Symbol;Acc:MGI:2443206] |
| 5 | lncRNA | 52744861 | 52748627 | ENSMUSG00000122870 | MGI:7825334 | Gm71278 | predicted gene, 71278 [Source:MGI Symbol;Acc:MGI:7825334] |
| 5 | lncRNA | 52864538 | 52870164 | ENSMUSG00000127317 | MGI:7825803 | Gm71515 | predicted gene, 71515 [Source:MGI Symbol;Acc:MGI:7825803] |
| 5 | lncRNA | 52898657 | 52899604 | ENSMUSG00000138208 | MGI:7828538 | Gm72888 | predicted gene, 72888 [Source:MGI Symbol;Acc:MGI:7828538] |
| 5 | lncRNA | 53041447 | 53044211 | ENSMUSG00000125532 | MGI:5589400 | Gm30241 | predicted gene, 30241 [Source:MGI Symbol;Acc:MGI:5589400] |

| | | | | | | | |
|---|---|---|---|---|---|---|---|
| 5 | lncRNA | 53067635 | 53187870 | ENSMUSG00000106538 | MGI:5589460 | Gm30301 | predicted gene, 30301 [Source:MGI Symbol;Acc:MGI:5589460] |
| 5 | lncRNA | 53080694 | 53081256 | ENSMUSG00000122780 | MGI:7825307 | Gm71263 | predicted gene, 71263 [Source:MGI Symbol;Acc:MGI:7825307] |
| 5 | lncRNA | 53130322 | 53157698 | ENSMUSG00000106352 | MGI:1923204 | 5033403H07Rik | RIKEN cDNA 5033403H07 gene [Source:MGI Symbol;Acc:MGI:1923204] |
| 5 | lncRNA | 53205029 | 53206530 | ENSMUSG00000090642 | MGI:4938009 | Gm17182 | predicted gene 17182 [Source:MGI Symbol;Acc:MGI:4938009] |
| 5 | lncRNA | 53369449 | 53395987 | ENSMUSG00000128101 | MGI:5589623 | Gm30464 | predicted gene, 30464 [Source:MGI Symbol;Acc:MGI:5589623] |
| 5 | lncRNA | 53396880 | 53404254 | ENSMUSG00000124368 | MGI:5589678 | Gm30519 | predicted gene, 30519 [Source:MGI Symbol;Acc:MGI:5589678] |
| 5 | lncRNA | 53421512 | 53424670 | ENSMUSG00000142257 | MGI:5623188 | Gm40303 | predicted gene, 40303 [Source:MGI Symbol;Acc:MGI:5623188] |
| 5 | lncRNA | 53439152 | 53440851 | ENSMUSG00000106981 | MGI:5791331 | Gm45495 | predicted gene 45495 [Source:MGI Symbol;Acc:MGI:5791331] |
| 5 | lncRNA | 53517669 | 53529118 | ENSMUSG00000106808 | MGI:5662882 | Gm42745 | predicted gene 42745 [Source:MGI Symbol;Acc:MGI:5662882] |
| 5 | lncRNA | 53556413 | 53561612 | ENSMUSG00000133208 | MGI:7827040 | Gm72137 | predicted gene, 72137 [Source:MGI Symbol;Acc:MGI:7827040] |
| 5 | lncRNA | 53621108 | 53626890 | ENSMUSG00000125364 | MGI:7825613 | Gm71420 | predicted gene, 71420 [Source:MGI Symbol;Acc:MGI:7825613] |
| 5 | lncRNA | 53627024 | 53627431 | ENSMUSG00000125448 | MGI:7825621 | Gm71424 | predicted gene, 71424 [Source:MGI Symbol;Acc:MGI:7825621] |
| 5 | lncRNA | 53676602 | 53680073 | ENSMUSG00000106807 | MGI:3641801 | Gm10441 | predicted gene 10441 [Source:MGI Symbol;Acc:MGI:3641801] |
| 5 | lncRNA | 53703517 | 53704753 | ENSMUSG00000107111 | MGI:5623189 | Gm40304 | predicted gene, 40304 [Source:MGI Symbol;Acc:MGI:5623189] |
| 5 | lncRNA | 53828492 | 53828778 | ENSMUSG00000130376 | MGI:7826198 | Gm71715 | predicted gene, 71715 [Source:MGI Symbol;Acc:MGI:7826198] |
| 5 | lncRNA | 54081392 | 54087398 | ENSMUSG00000132695 | MGI:7826824 | Gm72029 | predicted gene, 72029 [Source:MGI Symbol;Acc:MGI:7826824] |
| 5 | lncRNA | 54134035 | 54153958 | ENSMUSG00000126689 | MGI:5589760 | Gm30601 | predicted gene, 30601 [Source:MGI Symbol;Acc:MGI:5589760] |
| 5 | lncRNA | 54302971 | 54305114 | ENSMUSG00000122231 | MGI:7825241 | Gm71230 | predicted gene, 71230 [Source:MGI Symbol;Acc:MGI:7825241] |

| | | | | | | | |
|---|---|---|---|---|---|---|---|
| 5 | lncRNA | 54424223 | 54671511 | ENSMUSG00000105912 | MGI:3641803 | Gm10440 | predicted gene 10440 [Source:MGI Symbol;Acc:MGI:3641803] |
| 5 | lncRNA | 54521779 | 54580819 | ENSMUSG00000134062 | MGI:7827324 | Gm72279 | predicted gene, 72279 [Source:MGI Symbol;Acc:MGI:7827324] |
| 5 | lncRNA | 54586049 | 54598834 | ENSMUSG00000134107 | MGI:7827366 | Gm72300 | predicted gene, 72300 [Source:MGI Symbol;Acc:MGI:7827366] |
| 5 | lncRNA | 54598112 | 54603067 | ENSMUSG00000134141 | MGI:7827386 | Gm72310 | predicted gene, 72310 [Source:MGI Symbol;Acc:MGI:7827386] |
| 5 | lncRNA | 56169492 | 56177088 | ENSMUSG00000126211 | MGI:7825687 | Gm71457 | predicted gene, 71457 [Source:MGI Symbol;Acc:MGI:7825687] |
| 5 | lncRNA | 57316575 | 57317567 | ENSMUSG00000138900 | MGI:7828752 | Gm72996 | predicted gene, 72996 [Source:MGI Symbol;Acc:MGI:7828752] |
| 5 | lncRNA | 57727421 | 57879729 | ENSMUSG00000097216 | MGI:2441691 | 4932441J04Rik | RIKEN cDNA 4932441J04 gene [Source:MGI Symbol;Acc:MGI:2441691] |
| 5 | lncRNA | 57872321 | 57873985 | ENSMUSG00000140036 | MGI:7829030 | Gm73136 | predicted gene, 73136 [Source:MGI Symbol;Acc:MGI:7829030] |
| 5 | lncRNA | 58205960 | 58217711 | ENSMUSG00000137396 | MGI:7828378 | Gm72808 | predicted gene, 72808 [Source:MGI Symbol;Acc:MGI:7828378] |
| 5 | lncRNA | 58383961 | 58428446 | ENSMUSG00000124009 | MGI:7825475 | Gm71350 | predicted gene, 71350 [Source:MGI Symbol;Acc:MGI:7825475] |
| 5 | lncRNA | 58600579 | 58610346 | ENSMUSG00000105234 | MGI:1925413 | 4930459L07Rik | RIKEN cDNA 4930459L07 gene [Source:MGI Symbol;Acc:MGI:1925413] |
| 5 | lncRNA | 58610360 | 58612683 | ENSMUSG00000100557 | MGI:1919500 | 1700029E06Rik | RIKEN cDNA 1700029E06 gene [Source:MGI Symbol;Acc:MGI:1919500] |
| 5 | lncRNA | 58686983 | 58690013 | ENSMUSG00000136713 | MGI:7828114 | Gm72676 | predicted gene, 72676 [Source:MGI Symbol;Acc:MGI:7828114] |
| 5 | lncRNA | 58735034 | 58847926 | ENSMUSG00000133497 | MGI:7827146 | Gm72190 | predicted gene, 72190 [Source:MGI Symbol;Acc:MGI:7827146] |
| 5 | lncRNA | 58985311 | 59042383 | ENSMUSG00000129218 | MGI:7825963 | Gm71595 | predicted gene, 71595 [Source:MGI Symbol;Acc:MGI:7825963] |
| 5 | lncRNA | 59727045 | 59731585 | ENSMUSG00000106634 | MGI:5663179 | Gm43042 | predicted gene 43042 [Source:MGI Symbol;Acc:MGI:5663179] |
| 5 | lncRNA | 59870661 | 59901815 | ENSMUSG00000138074 | MGI:7828518 | Gm72878 | predicted gene, 72878 [Source:MGI Symbol;Acc:MGI:7828518] |
| 5 | lncRNA | 60106402 | 60113597 | ENSMUSG00000123790 | MGI:7825443 | Gm71333 | predicted gene, 71333 [Source:MGI Symbol;Acc:MGI:7825443] |

| | | | | | | | |
|---|---|---|---|---|---|---|---|
| 5 | lncRNA | 60116730 | 60143384 | ENSMUSG00000106013 | MGI:1918307 | 4933402J10Rik | RIKEN cDNA 4933402J10 gene [Source:MGI Symbol;Acc:MGI:1918307] |
| 5 | lncRNA | 60388621 | 60389202 | ENSMUSG00000129251 | MGI:7825967 | Gm71597 | predicted gene, 71597 [Source:MGI Symbol;Acc:MGI:7825967] |
| 5 | lncRNA | 60468348 | 60471120 | ENSMUSG00000135740 | MGI:7827770 | Gm72503 | predicted gene, 72503 [Source:MGI Symbol;Acc:MGI:7827770] |
| 5 | lncRNA | 60543093 | 60560807 | ENSMUSG00000136235 | MGI:7827898 | Gm72567 | predicted gene, 72567 [Source:MGI Symbol;Acc:MGI:7827898] |
| 5 | lncRNA | 61098281 | 61152246 | ENSMUSG00000105903 | MGI:1924172 | 1700014F14Rik | RIKEN cDNA 1700014F14 gene [Source:MGI Symbol;Acc:MGI:1924172] |
| 5 | lncRNA | 61187951 | 61196673 | ENSMUSG00000133805 | MGI:7827228 | Gm72231 | predicted gene, 72231 [Source:MGI Symbol;Acc:MGI:7827228] |
| 5 | lncRNA | 61520600 | 61531385 | ENSMUSG00000134597 | MGI:7827468 | Gm72351 | predicted gene, 72351 [Source:MGI Symbol;Acc:MGI:7827468] |
| 5 | lncRNA | 61722400 | 61734622 | ENSMUSG00000143250 | MGI:7829752 | Gm73498 | predicted gene, 73498 [Source:MGI Symbol;Acc:MGI:7829752] |
| 5 | lncRNA | 61846082 | 61849804 | ENSMUSG00000124549 | MGI:7825545 | Gm71386 | predicted gene, 71386 [Source:MGI Symbol;Acc:MGI:7825545] |
| 5 | lncRNA | 61921477 | 61966679 | ENSMUSG00000143342 | MGI:5623193 | Gm40308 | predicted gene, 40308 [Source:MGI Symbol;Acc:MGI:5623193] |
| 5 | lncRNA | 61964088 | 61964533 | ENSMUSG00000143380 | MGI:7829796 | Gm73520 | predicted gene, 73520 [Source:MGI Symbol;Acc:MGI:7829796] |
| 5 | lncRNA | 62207719 | 62231006 | ENSMUSG00000138711 | MGI:7828672 | Gm72956 | predicted gene, 72956 [Source:MGI Symbol;Acc:MGI:7828672] |
| 5 | lncRNA | 62456170 | 62456705 | ENSMUSG00000128227 | MGI:7825863 | Gm71545 | predicted gene, 71545 [Source:MGI Symbol;Acc:MGI:7825863] |
| 5 | lncRNA | 62711691 | 62745660 | ENSMUSG00000122017 | MGI:7825185 | Gm71202 | predicted gene, 71202 [Source:MGI Symbol;Acc:MGI:7825185] |
| 5 | lncRNA | 63008684 | 63025265 | ENSMUSG00000124789 | MGI:7825563 | Gm71395 | predicted gene, 71395 [Source:MGI Symbol;Acc:MGI:7825563] |
| 5 | lncRNA | 63718561 | 63781615 | ENSMUSG00000134846 | MGI:5589996 | Gm30837 | predicted gene, 30837 [Source:MGI Symbol;Acc:MGI:5589996] |
| 5 | lncRNA | 63960917 | 63961635 | ENSMUSG00000132514 | MGI:7826796 | Gm72015 | predicted gene, 72015 [Source:MGI Symbol;Acc:MGI:7826796] |
| 5 | lncRNA | 64081244 | 64082313 | ENSMUSG00000087552 | MGI:3802175 | Gm15819 | predicted gene 15819 [Source:MGI Symbol;Acc:MGI:3802175] |

| | | | | | | | |
|---|---|---|---|---|---|---|---|
| 5 | lncRNA | 64187388 | 64200435 | ENSMUSG00000127532 | MGI:7825817 | Gm71522 | predicted gene, 71522 [Source:MGI Symbol;Acc:MGI:7825817] |
| 5 | lncRNA | 64202316 | 64215554 | ENSMUSG00000106219 | MGI:1922007 | 5830416I19Rik | RIKEN cDNA 5830416I19 gene [Source:MGI Symbol;Acc:MGI:1922007] |
| 5 | lncRNA | 64306914 | 64310169 | ENSMUSG00000123942 | MGI:7825463 | Gm71344 | predicted gene, 71344 [Source:MGI Symbol;Acc:MGI:7825463] |
| 5 | lncRNA | 64388000 | 64388359 | ENSMUSG00000132563 | MGI:7826802 | Gm72018 | predicted gene, 72018 [Source:MGI Symbol;Acc:MGI:7826802] |
| 5 | lncRNA | 64480644 | 64521876 | ENSMUSG00000142621 | MGI:7829632 | Gm73438 | predicted gene, 73438 [Source:MGI Symbol;Acc:MGI:7829632] |
| 5 | lncRNA | 64500293 | 64501895 | ENSMUSG00000142836 | MGI:7829676 | Gm73460 | predicted gene, 73460 [Source:MGI Symbol;Acc:MGI:7829676] |
| 5 | lncRNA | 64517664 | 64665195 | ENSMUSG00000105730 | MGI:5663975 | Gm43838 | predicted gene 43838 [Source:MGI Symbol;Acc:MGI:5663975] |
| 5 | lncRNA | 64531911 | 64535171 | ENSMUSG00000142793 | MGI:7829666 | Gm73455 | predicted gene, 73455 [Source:MGI Symbol;Acc:MGI:7829666] |
| 5 | lncRNA | 64612029 | 64630076 | ENSMUSG00000100911 | MGI:1917249 | 1700027F09Rik | RIKEN cDNA 1700027F09 gene [Source:MGI Symbol;Acc:MGI:1917249] |
| 5 | lncRNA | 64635541 | 64645120 | ENSMUSG00000142709 | MGI:7829662 | Gm73453 | predicted gene, 73453 [Source:MGI Symbol;Acc:MGI:7829662] |
| 5 | lncRNA | 64673246 | 64673829 | ENSMUSG00000129223 | MGI:7825965 | Gm71596 | predicted gene, 71596 [Source:MGI Symbol;Acc:MGI:7825965] |
| 5 | lncRNA | 64692554 | 64695172 | ENSMUSG00000123933 | MGI:5590582 | Gm31423 | predicted gene, 31423 [Source:MGI Symbol;Acc:MGI:5590582] |
| 5 | lncRNA | 64751135 | 64768095 | ENSMUSG00000105402 | MGI:3781892 | Gm3716 | predicted gene 3716 [Source:MGI Symbol;Acc:MGI:3781892] |
| 5 | lncRNA | 64752661 | 64763260 | ENSMUSG00000124498 | MGI:7825535 | Gm71381 | predicted gene, 71381 [Source:MGI Symbol;Acc:MGI:7825535] |
| 5 | lncRNA | 64790847 | 64961791 | ENSMUSG00000097640 | MGI:5012218 | Gm20033 | predicted gene, 20033 [Source:MGI Symbol;Acc:MGI:5012218] |
| 5 | lncRNA | 64792704 | 64794731 | ENSMUSG00000105183 | MGI:5662701 | Gm42564 | predicted gene 42564 [Source:MGI Symbol;Acc:MGI:5662701] |
| 5 | lncRNA | 64793183 | 64794959 | ENSMUSG00000104928 | MGI:5662702 | Gm42565 | predicted gene 42565 [Source:MGI Symbol;Acc:MGI:5662702] |
| 5 | lncRNA | 64845044 | 64853943 | ENSMUSG00000136537 | MGI:7828016 | Gm72627 | predicted gene, 72627 [Source:MGI Symbol;Acc:MGI:7828016] |

| | | | | | | | |
|---|---|---|---|---|---|---|---|
| 5 | lncRNA | 64850760 | 64852105 | ENSMUSG00000136771 | MGI:7828162 | Gm72700 | predicted gene, 72700 [Source:MGI Symbol;Acc:MGI:7828162] |
| 5 | lncRNA | 64852804 | 64855931 | ENSMUSG00000097668 | MGI:5477255 | Gm26761 | predicted gene, 26761 [Source:MGI Symbol;Acc:MGI:5477255] |
| 5 | lncRNA | 64872245 | 64880507 | ENSMUSG00000136576 | MGI:7828018 | Gm72628 | predicted gene, 72628 [Source:MGI Symbol;Acc:MGI:7828018] |
| 5 | lncRNA | 64898790 | 64905510 | ENSMUSG00000136618 | MGI:7828020 | Gm72629 | predicted gene, 72629 [Source:MGI Symbol;Acc:MGI:7828020] |
| 5 | lncRNA | 64905617 | 64912317 | ENSMUSG00000136650 | MGI:7828080 | Gm72659 | predicted gene, 72659 [Source:MGI Symbol;Acc:MGI:7828080] |
| 5 | lncRNA | 64955290 | 64959712 | ENSMUSG00000136685 | MGI:7828098 | Gm72668 | predicted gene, 72668 [Source:MGI Symbol;Acc:MGI:7828098] |
| 5 | lncRNA | 65006058 | 65022788 | ENSMUSG00000129045 | MGI:7825951 | Gm71589 | predicted gene, 71589 [Source:MGI Symbol;Acc:MGI:7825951] |
| 5 | lncRNA | 65036877 | 65046210 | ENSMUSG00000126366 | MGI:5591048 | Gm31889 | predicted gene, 31889 [Source:MGI Symbol;Acc:MGI:5591048] |
| 5 | lncRNA | 65055091 | 65063183 | ENSMUSG00000123189 | MGI:7830556 | Gm73906 | predicted gene, 73906 [Source:MGI Symbol;Acc:MGI:7830556] |
| 5 | lncRNA | 65056480 | 65060908 | ENSMUSG00000123224 | MGI:7825377 | Gm71300 | predicted gene, 71300 [Source:MGI Symbol;Acc:MGI:7825377] |
| 5 | lncRNA | 65091094 | 65093321 | ENSMUSG00000140599 | MGI:7829202 | Gm73222 | predicted gene, 73222 [Source:MGI Symbol;Acc:MGI:7829202] |
| 5 | lncRNA | 65117478 | 65126843 | ENSMUSG00000139765 | MGI:7828954 | Gm73098 | predicted gene, 73098 [Source:MGI Symbol;Acc:MGI:7828954] |
| 5 | lncRNA | 65205039 | 65208797 | ENSMUSG00000141992 | MGI:7829484 | Gm73363 | predicted gene, 73363 [Source:MGI Symbol;Acc:MGI:7829484] |
| 5 | lncRNA | 65215935 | 65229709 | ENSMUSG00000125845 | MGI:7825653 | Gm71440 | predicted gene, 71440 [Source:MGI Symbol;Acc:MGI:7825653] |
| 5 | lncRNA | 65260538 | 65264934 | ENSMUSG00000141465 | MGI:7829390 | Gm73316 | predicted gene, 73316 [Source:MGI Symbol;Acc:MGI:7829390] |
| 5 | lncRNA | 65493030 | 65503925 | ENSMUSG00000127609 | MGI:7825827 | Gm71527 | predicted gene, 71527 [Source:MGI Symbol;Acc:MGI:7825827] |
| 5 | lncRNA | 65745059 | 65748783 | ENSMUSG00000107102 | MGI:5662863 | Gm42726 | predicted gene 42726 [Source:MGI Symbol;Acc:MGI:5662863] |
| 5 | lncRNA | 65917839 | 65921732 | ENSMUSG00000106978 | MGI:1921484 | N4bp2os | NEDD4 binding protein 2, opposite strand [Source:MGI Symbol;Acc:MGI:1921484] |

| | | | | | | | |
|---|---|---|---|---|---|---|---|
| 5 | lncRNA | 65922212 | 65922830 | ENSMUSG00000137008 | MGI:7828262 | Gm72750 | predicted gene, 72750 [Source:MGI Symbol;Acc:MGI:7828262] |
| 5 | lncRNA | 65960105 | 66015649 | ENSMUSG00000135056 | MGI:3781698 | Gm3521 | predicted gene 3521 [Source:MGI Symbol;Acc:MGI:3781698] |
| 5 | lncRNA | 66008368 | 66008774 | ENSMUSG00000135096 | MGI:7827602 | Gm72419 | predicted gene, 72419 [Source:MGI Symbol;Acc:MGI:7827602] |
| 5 | lncRNA | 66024584 | 66028133 | ENSMUSG00000137580 | MGI:7828448 | Gm72843 | predicted gene, 72843 [Source:MGI Symbol;Acc:MGI:7828448] |
| 5 | lncRNA | 66041281 | 66070097 | ENSMUSG00000120171 | MGI:5591633 | Gm32474 | predicted gene, 32474 [Source:MGI Symbol;Acc:MGI:5591633] |
| 5 | lncRNA | 66060763 | 66073081 | ENSMUSG00000120641 | MGI:7338467 | Gm57277 | predicted gene, 57277 [Source:MGI Symbol;Acc:MGI:7338467] |
| 5 | lncRNA | 66072341 | 66077025 | ENSMUSG00000137541 | MGI:5591715 | Gm32556 | predicted gene, 32556 [Source:MGI Symbol;Acc:MGI:5591715] |
| 5 | lncRNA | 66086181 | 66086780 | ENSMUSG00000120547 | MGI:7338447 | Gm57267 | predicted gene, 57267 [Source:MGI Symbol;Acc:MGI:7338447] |
| 5 | lncRNA | 66101760 | 66113567 | ENSMUSG00000122150 | MGI:7825215 | Gm71217 | predicted gene, 71217 [Source:MGI Symbol;Acc:MGI:7825215] |
| 5 | lncRNA | 66108660 | 66109554 | ENSMUSG00000122193 | MGI:7825231 | Gm71225 | predicted gene, 71225 [Source:MGI Symbol;Acc:MGI:7825231] |
| 5 | lncRNA | 66136701 | 66161628 | ENSMUSG00000054598 | MGI:3041166 | 9130230L23Rik | RIKEN cDNA 9130230L23 gene [Source:MGI Symbol;Acc:MGI:3041166] |
| 5 | lncRNA | 66226398 | 66227339 | ENSMUSG00000132593 | MGI:7826806 | Gm72020 | predicted gene, 72020 [Source:MGI Symbol;Acc:MGI:7826806] |
| 5 | lncRNA | 66234224 | 66235217 | ENSMUSG00000140414 | MGI:7829146 | Gm73194 | predicted gene, 73194 [Source:MGI Symbol;Acc:MGI:7829146] |
| 5 | lncRNA | 66308366 | 66345219 | ENSMUSG00000097092 | MGI:5477219 | Gm26725 | predicted gene, 26725 [Source:MGI Symbol;Acc:MGI:5477219] |
| 5 | lncRNA | 66323950 | 66348679 | ENSMUSG00000107014 | MGI:1920889 | 1700126H18Rik | RIKEN cDNA 1700126H18 gene [Source:MGI Symbol;Acc:MGI:1920889] |
| 5 | lncRNA | 66332380 | 66333301 | ENSMUSG00000133276 | MGI:7827094 | Gm72164 | predicted gene, 72164 [Source:MGI Symbol;Acc:MGI:7827094] |
| 5 | lncRNA | 66371551 | 66371888 | ENSMUSG00000142467 | MGI:7829588 | Gm73415 | predicted gene, 73415 [Source:MGI Symbol;Acc:MGI:7829588] |
| 5 | lncRNA | 66371969 | 66373272 | ENSMUSG00000141032 | MGI:7829314 | Gm73278 | predicted gene, 73278 [Source:MGI Symbol;Acc:MGI:7829314] |

| | | | | | | | |
|---|---|---|---|---|---|---|---|
| 5 | lncRNA | 66480691 | 66492259 | ENSMUSG00000143772 | MGI:7829966 | Gm73605 | predicted gene, 73605 [Source:MGI Symbol;Acc:MGI:7829966] |
| 5 | lncRNA | 66499835 | 66502882 | ENSMUSG00000143812 | MGI:7829990 | Gm73617 | predicted gene, 73617 [Source:MGI Symbol;Acc:MGI:7829990] |
| 5 | lncRNA | 66563177 | 66564984 | ENSMUSG00000143854 | MGI:7830012 | Gm73628 | predicted gene, 73628 [Source:MGI Symbol;Acc:MGI:7830012] |
| 5 | lncRNA | 66783832 | 66833841 | ENSMUSG00000087601 | MGI:5446771 | Uchl1os | [illegible] opposite strand [Source:MGI Symbol;Acc:MGI:5446771] |
| 5 | lncRNA | 66844671 | 66847320 | ENSMUSG00000143516 | MGI:7829870 | Gm73557 | predicted gene, 73557 [Source:MGI Symbol;Acc:MGI:7829870] |
| 5 | lncRNA | 66874038 | 66877305 | ENSMUSG00000140006 | MGI:1918405 | 4933424C09Rik | RIKEN cDNA 4933424C09 gene [Source:MGI Symbol;Acc:MGI:1918405] |
| 5 | lncRNA | 67001309 | 67002854 | ENSMUSG00000124373 | MGI:7825527 | Gm71377 | predicted gene, 71377 [Source:MGI Symbol;Acc:MGI:7825527] |
| 5 | lncRNA | 67044882 | 67051495 | ENSMUSG00000124297 | MGI:7825513 | Gm71369 | predicted gene, 71369 [Source:MGI Symbol;Acc:MGI:7825513] |
| 5 | lncRNA | 67063518 | 67067487 | ENSMUSG00000124335 | MGI:7825523 | Gm71374 | predicted gene, 71374 [Source:MGI Symbol;Acc:MGI:7825523] |
| 5 | lncRNA | 67142434 | 67142984 | ENSMUSG00000085839 | MGI:3801730 | Gm15949 | predicted gene 15949 [Source:MGI Symbol;Acc:MGI:3801730] |
| 5 | lncRNA | 67146159 | 67148550 | ENSMUSG00000086964 | MGI:3801731 | Gm15948 | predicted gene 15948 [Source:MGI Symbol;Acc:MGI:3801731] |
| 5 | lncRNA | 67256018 | 67287479 | ENSMUSG00000122852 | | | predicted gene, 33167 [Source:NCBI gene (formerly Entrezgene);Acc:102635966] |
| 5 | lncRNA | 67378208 | 67389306 | ENSMUSG00000120155 | MGI:5592452 | Gm33293 | predicted gene, 33293 [Source:MGI Symbol;Acc:MGI:5592452] |
| 5 | lncRNA | 67401114 | 67418605 | ENSMUSG00000107319 | MGI:5662807 | Gm42670 | predicted gene 42670 [Source:MGI Symbol;Acc:MGI:5662807] |
| 5 | lncRNA | 67517660 | 67523998 | ENSMUSG00000107266 | MGI:5663835 | Gm43698 | predicted gene 43698 [Source:MGI Symbol;Acc:MGI:5663835] |
| 5 | lncRNA | 67585021 | 67595333 | ENSMUSG00000097840 | MGI:5477250 | Gm26756 | predicted gene, 26756 [Source:MGI Symbol;Acc:MGI:5477250] |
| 5 | lncRNA | 67601535 | 67602073 | ENSMUSG00000135689 | MGI:7827762 | Gm72499 | predicted gene, 72499 [Source:MGI Symbol;Acc:MGI:7827762] |
| 5 | lncRNA | 67601952 | 67614849 | ENSMUSG00000135654 | | | novel transcript |

| | | | | | | | |
|---|---|---|---|---|---|---|---|
| 5 | lncRNA | 67621240 | 67685733 | ENSMUSG00000107385 | MGI:2444116 | C330024D21Rik | RIKEN cDNA C330024D21 gene [Source:MGI Symbol;Acc:MGI:2444116] |
| 5 | lncRNA | 67663123 | 67699286 | ENSMUSG00000126656 | MGI:7825739 | Gm71483 | predicted gene, 71483 [Source:MGI Symbol;Acc:MGI:7825739] |
| 5 | lncRNA | 67764028 | 67765858 | ENSMUSG00000139038 | MGI:7828790 | Gm73015 | predicted gene, 73015 [Source:MGI Symbol;Acc:MGI:7828790] |
| 5 | lncRNA | 67799744 | 67804119 | ENSMUSG00000085363 | MGI:3705145 | Gm15478 | predicted gene 15478 [Source:MGI Symbol;Acc:MGI:3705145] |
| 5 | lncRNA | 67989308 | 67991617 | ENSMUSG00000138756 | MGI:7828690 | Gm72965 | predicted gene, 72965 [Source:MGI Symbol;Acc:MGI:7828690] |
| 5 | lncRNA | 68004408 | 68134426 | ENSMUSG00000087261 | MGI:3705209 | Gm15477 | predicted gene 15477 [Source:MGI Symbol;Acc:MGI:3705209] |
| 5 | lncRNA | 68090947 | 68100493 | ENSMUSG00000106880 | MGI:1921115 | 4930425K10Rik | RIKEN cDNA 4930425K10 gene [Source:MGI Symbol;Acc:MGI:1921115] |
| 5 | lncRNA | 68833488 | 68862977 | ENSMUSG00000125658 | MGI:7825641 | Gm71434 | predicted gene, 71434 [Source:MGI Symbol;Acc:MGI:7825641] |
| 5 | lncRNA | 69163318 | 69165913 | ENSMUSG00000128313 | MGI:7825879 | Gm71553 | predicted gene, 71553 [Source:MGI Symbol;Acc:MGI:7825879] |
| 5 | lncRNA | 69234412 | 69351194 | ENSMUSG00000129031 | MGI:7825949 | Gm71588 | predicted gene, 71588 [Source:MGI Symbol;Acc:MGI:7825949] |
| 5 | lncRNA | 69870331 | 69879091 | ENSMUSG00000122868 | MGI:7825332 | Gm71277 | predicted gene, 71277 [Source:MGI Symbol;Acc:MGI:7825332] |
| 5 | lncRNA | 69983472 | 69984189 | ENSMUSG00000128577 | MGI:7825921 | Gm71574 | predicted gene, 71574 [Source:MGI Symbol;Acc:MGI:7825921] |
| 5 | lncRNA | 70773071 | 70813819 | ENSMUSG00000141687 | MGI:7829420 | Gm73331 | predicted gene, 73331 [Source:MGI Symbol;Acc:MGI:7829420] |
| 5 | lncRNA | 71253257 | 71272467 | ENSMUSG00000144038 | MGI:7830100 | Gm73674 | predicted gene, 73674 [Source:MGI Symbol;Acc:MGI:7830100] |
| 5 | lncRNA | 71837607 | 71858841 | ENSMUSG00000105489 | MGI:5621447 | Gm38562 | predicted gene, 38562 [Source:MGI Symbol;Acc:MGI:5621447] |
| 5 | lncRNA | 72180652 | 72188828 | ENSMUSG00000139447 | MGI:7828904 | Gm73073 | predicted gene, 73073 [Source:MGI Symbol;Acc:MGI:7828904] |
| 5 | lncRNA | 72188978 | 72189764 | ENSMUSG00000139489 | MGI:7828910 | Gm73076 | predicted gene, 73076 [Source:MGI Symbol;Acc:MGI:7828910] |
| 5 | lncRNA | 72402768 | 72419443 | ENSMUSG00000142102 | MGI:7829496 | Gm73369 | predicted gene, 73369 [Source:MGI Symbol;Acc:MGI:7829496] |

| | | | | | | | |
|---|---|---|---|---|---|---|---|
| 5 | lncRNA | 72412097 | 72422093 | ENSMUSG00000142140 | MGI:7829520 | Gm73381 | predicted gene, 73381 [Source:MGI Symbol;Acc:MGI:7829520] |
| 5 | lncRNA | 72473671 | 72525391 | ENSMUSG00000093400 | MGI:5313094 | Gm20647 | predicted gene 20647 [Source:MGI Symbol;Acc:MGI:5313094] |
| 5 | lncRNA | 72653543 | 72654447 | ENSMUSG00000093771 | MGI:3034643 | AU023070 | expressed sequence AU023070 [Source:MGI Symbol;Acc:MGI:3034643] |
| 5 | lncRNA | 72660812 | 72663757 | ENSMUSG00000141448 | MGI:7829388 | Gm73315 | predicted gene, 73315 [Source:MGI Symbol;Acc:MGI:7829388] |
| 5 | lncRNA | 72792930 | 72793556 | ENSMUSG00000143901 | MGI:7830026 | Gm73636 | predicted gene, 73636 [Source:MGI Symbol;Acc:MGI:7830026] |
| 5 | lncRNA | 72801750 | 72804483 | ENSMUSG00000106136 | MGI:1921334 | 4933408A14Rik | RIKEN cDNA 4933408A14 gene [Source:MGI Symbol;Acc:MGI:1921334] |
| 5 | lncRNA | 72817429 | 72836966 | ENSMUSG00000135140 | MGI:7827610 | Gm72423 | predicted gene, 72423 [Source:MGI Symbol;Acc:MGI:7827610] |
| 5 | lncRNA | 72837080 | 72849347 | ENSMUSG00000139316 | MGI:7828836 | Gm73039 | predicted gene, 73039 [Source:MGI Symbol;Acc:MGI:7828836] |
| 5 | lncRNA | 73064588 | 73066196 | ENSMUSG00000141163 | MGI:7829348 | Gm73295 | predicted gene, 73295 [Source:MGI Symbol;Acc:MGI:7829348] |
| 5 | lncRNA | 73070894 | 73071564 | ENSMUSG00000132452 | MGI:7826788 | Gm72011 | predicted gene, 72011 [Source:MGI Symbol;Acc:MGI:7826788] |
| 5 | lncRNA | 73171243 | 73190770 | ENSMUSG00000142109 | | | novel transcript, antisense to Fryl |
| 5 | lncRNA | 73314344 | 73315233 | ENSMUSG00000086376 | MGI:1920735 | 1700071G01Rik | RIKEN cDNA 1700071G01 gene [Source:MGI Symbol;Acc:MGI:1920735] |
| 5 | lncRNA | 73367499 | 73369986 | ENSMUSG00000090561 | MGI:4938034 | Gm17207 | predicted gene 17207 [Source:MGI Symbol;Acc:MGI:4938034] |
| 5 | lncRNA | 73388934 | 73392244 | ENSMUSG00000136166 | MGI:7827846 | Gm72541 | predicted gene, 72541 [Source:MGI Symbol;Acc:MGI:7827846] |
| 5 | lncRNA | 73413556 | 73436481 | ENSMUSG00000107087 | MGI:5593570 | Gm34411 | predicted gene, 34411 [Source:MGI Symbol;Acc:MGI:5593570] |
| 5 | lncRNA | 73425716 | 73449924 | ENSMUSG00000100175 | MGI:1919526 | 1700025M24Rik | RIKEN cDNA 1700025M24 gene [Source:MGI Symbol;Acc:MGI:1919526] |
| 5 | lncRNA | 73439298 | 73441107 | ENSMUSG00000136196 | MGI:7827874 | Gm72555 | predicted gene, 72555 [Source:MGI Symbol;Acc:MGI:7827874] |
| 5 | lncRNA | 73441541 | 73445986 | ENSMUSG00000136125 | MGI:5623201 | Gm40316 | predicted gene, 40316 [Source:MGI Symbol;Acc:MGI:5623201] |

| | | | | | | | |
|---|---|---|---|---|---|---|---|
| 5 | lncRNA | 73506014 | 73525734 | ENSMUSG00000125783 | MGI:5593655 | Gm34496 | predicted gene, 34496 [Source:MGI Symbol;Acc:MGI:5593655] |
| 5 | lncRNA | 73535307 | 73550115 | ENSMUSG00000121792 | MGI:7828490 | Gm72864 | predicted gene, 72864 [Source:MGI Symbol;Acc:MGI:7828490] |
| 5 | lncRNA | 73633700 | 73638944 | ENSMUSG00000137355 | MGI:7830260 | Gm73755 | predicted gene, 73755 [Source:MGI Symbol;Acc:MGI:7830260] |
| 5 | lncRNA | 73731812 | 73734265 | ENSMUSG00000139088 | MGI:7828794 | Gm73017 | predicted gene, 73017 [Source:MGI Symbol;Acc:MGI:7828794] |
| 5 | lncRNA | 73806349 | 73809172 | ENSMUSG00000137084 | MGI:5594061 | Gm34902 | predicted gene, 34902 [Source:MGI Symbol;Acc:MGI:5594061] |
| 5 | lncRNA | 73955125 | 73962043 | ENSMUSG00000126365 | MGI:5594119 | Gm34960 | predicted gene, 34960 [Source:MGI Symbol;Acc:MGI:5594119] |
| 5 | lncRNA | 74060149 | 74072798 | ENSMUSG00000129782 | MGI:7826029 | Gm71629 | predicted gene, 71629 [Source:MGI Symbol;Acc:MGI:7826029] |
| 5 | lncRNA | 74070190 | 74077770 | ENSMUSG00000107027 | MGI:1916652 | 1700019F05Rik | RIKEN cDNA 1700019F05 gene [Source:MGI Symbol;Acc:MGI:1916652] |
| 5 | lncRNA | 74090336 | 74112514 | ENSMUSG00000143520 | MGI:7829874 | Gm73559 | predicted gene, 73559 [Source:MGI Symbol;Acc:MGI:7829874] |
| 5 | lncRNA | 74102628 | 74106682 | ENSMUSG00000106968 | MGI:2141303 | C78283 | expressed sequence C78283 [Source:MGI Symbol;Acc:MGI:2141303] |
| 5 | lncRNA | 74174805 | 74200657 | ENSMUSG00000121985 | | | novel transcript |
| 5 | lncRNA | 74174859 | 74190341 | ENSMUSG00000087250 | MGI:1921549 | Usp46os1 | ubiquitin specific peptidase 46, opposite strand 1 [Source:MGI Symbol;Acc:MGI:1921549] |
| 5 | lncRNA | 74192929 | 74200664 | ENSMUSG00000086234 | MGI:1924246 | Usp46os2 | ubiquitin specific peptidase 46, opposite strand 2 [Source:MGI Symbol;Acc:MGI:1924246] |
| 5 | lncRNA | 74241653 | 74254064 | ENSMUSG00000128868 | MGI:7825935 | Gm71581 | predicted gene, 71581 [Source:MGI Symbol;Acc:MGI:7825935] |
| 5 | lncRNA | 74253357 | 74284805 | ENSMUSG00000106943 | MGI:1917286 | Dancr | differentiation antagonizing non-protein coding RNA [Source:MGI Symbol;Acc:MGI:1917286] |
| 5 | lncRNA | 74272040 | 74279270 | ENSMUSG00000128908 | MGI:7825939 | Gm71583 | predicted gene, 71583 [Source:MGI Symbol;Acc:MGI:7825939] |
| 5 | lncRNA | 74303305 | 74319238 | ENSMUSG00000122053 | MGI:7825203 | Gm71211 | predicted gene, 71211 [Source:MGI Symbol;Acc:MGI:7825203] |
| 5 | lncRNA | 74322104 | 74325657 | ENSMUSG00000143368 | MGI:5591161 | Gm32002 | predicted gene, 32002 [Source:MGI Symbol;Acc:MGI:5591161] |

| | | | | | | | |
|---|---|---|---|---|---|---|---|
| 5 | lncRNA | 74322227 | 74322947 | ENSMUSG00000143408 | MGI:7829806 | Gm73525 | predicted gene, 73525 [Source:MGI Symbol;Acc:MGI:7829806] |
| 5 | lncRNA | 74334851 | 74348935 | ENSMUSG00000139163 | MGI:7828812 | Gm73027 | predicted gene, 73027 [Source:MGI Symbol;Acc:MGI:7828812] |
| 5 | lncRNA | 74477205 | 74520529 | ENSMUSG00000136303 | MGI:5591223 | Gm32064 | predicted gene, 32064 [Source:MGI Symbol;Acc:MGI:5591223] |
| 5 | lncRNA | 74565860 | 74595439 | ENSMUSG00000136344 | MGI:7827936 | Gm72587 | predicted gene, 72587 [Source:MGI Symbol;Acc:MGI:7827936] |
| 5 | lncRNA | 74808344 | 74819091 | ENSMUSG00000121106 | MGI:5591463 | Gm32304 | predicted gene, 32304 [Source:MGI Symbol;Acc:MGI:5591463] |
| 5 | lncRNA | 74830845 | 74889528 | ENSMUSG00000085657 | MGI:3802050 | Gm15984 | predicted gene 15984 [Source:MGI Symbol;Acc:MGI:3802050] |
| 5 | lncRNA | 74891092 | 74895116 | ENSMUSG00000085497 | MGI:3801786 | Gm15985 | predicted gene 15985 [Source:MGI Symbol;Acc:MGI:3801786] |
| 5 | lncRNA | 74910543 | 74915099 | ENSMUSG00000106987 | MGI:5662712 | Gm42575 | predicted gene 42575 [Source:MGI Symbol;Acc:MGI:5662712] |
| 5 | lncRNA | 74913900 | 74938534 | ENSMUSG00000139893 | MGI:7828968 | Gm73105 | predicted gene, 73105 [Source:MGI Symbol;Acc:MGI:7828968] |
| 5 | lncRNA | 74949995 | 74957654 | ENSMUSG00000131829 | MGI:7826636 | Gm71935 | predicted gene, 71935 [Source:MGI Symbol;Acc:MGI:7826636] |
| 5 | lncRNA | 75101786 | 75109368 | ENSMUSG00000141831 | MGI:6367169 | Gm52779 | predicted gene, 52779 [Source:MGI Symbol;Acc:MGI:6367169] |
| 5 | lncRNA | 75109820 | 75140100 | ENSMUSG00000072874 | MGI:3644088 | Gm6116 | predicted gene 6116 [Source:MGI Symbol;Acc:MGI:3644088] |
| 5 | lncRNA | 75228008 | 75232822 | ENSMUSG00000137041 | MGI:7828266 | Gm72752 | predicted gene, 72752 [Source:MGI Symbol;Acc:MGI:7828266] |
| 5 | lncRNA | 75230570 | 75236830 | ENSMUSG00000137002 | MGI:7828252 | Gm72745 | predicted gene, 72745 [Source:MGI Symbol;Acc:MGI:7828252] |
| 5 | lncRNA | 75270966 | 75307794 | ENSMUSG00000107192 | MGI:5011768 | Gm19583 | predicted gene, 19583 [Source:MGI Symbol;Acc:MGI:5011768] |
| 5 | lncRNA | 75313716 | 75314314 | ENSMUSG00000106760 | MGI:5662939 | Gm42802 | predicted gene 42802 [Source:MGI Symbol;Acc:MGI:5662939] |
| 5 | lncRNA | 75376805 | 75377675 | ENSMUSG00000142591 | MGI:7829630 | Gm73437 | predicted gene, 73437 [Source:MGI Symbol;Acc:MGI:7829630] |
| 5 | lncRNA | 75420845 | 75424125 | ENSMUSG00000127495 | MGI:7825815 | Gm71521 | predicted gene, 71521 [Source:MGI Symbol;Acc:MGI:7825815] |

| | | | | | | | |
|---|---|---|---|---|---|---|---|
| 5 | lncRNA | 75484797 | 75527661 | ENSMUSG00000127213 | MGI:7830290 | Gm73770 | predicted gene, 73770 [Source:MGI Symbol;Acc:MGI:7830290] |
| 5 | lncRNA | 75527165 | 75528751 | ENSMUSG00000127253 | MGI:7825801 | Gm71514 | predicted gene, 71514 [Source:MGI Symbol;Acc:MGI:7825801] |
| 5 | lncRNA | 75538146 | 75548435 | ENSMUSG00000123059 | MGI:7825352 | Gm71287 | predicted gene, 71287 [Source:MGI Symbol;Acc:MGI:7825352] |
| 5 | lncRNA | 75538275 | 75560066 | ENSMUSG00000107386 | MGI:5662937 | Gm42800 | predicted gene 42800 [Source:MGI Symbol;Acc:MGI:5662937] |
| 5 | lncRNA | 75539613 | 75539827 | ENSMUSG00000123097 | MGI:7825354 | Gm71288 | predicted gene, 71288 [Source:MGI Symbol;Acc:MGI:7825354] |
| 5 | lncRNA | 75671908 | 75674187 | ENSMUSG00000138324 | MGI:7828558 | Gm72899 | predicted gene, 72899 [Source:MGI Symbol;Acc:MGI:7828558] |
| 5 | lncRNA | 75817395 | 75835697 | ENSMUSG00000138764 | MGI:7828692 | Gm72966 | predicted gene, 72966 [Source:MGI Symbol;Acc:MGI:7828692] |
| 5 | lncRNA | 75835827 | 75860719 | ENSMUSG00000129581 | MGI:5591717 | Gm32558 | predicted gene, 32558 [Source:MGI Symbol;Acc:MGI:5591717] |
| 5 | lncRNA | 75888098 | 75909603 | ENSMUSG00000106754 | MGI:5663238 | Gm43101 | predicted gene 43101 [Source:MGI Symbol;Acc:MGI:5663238] |
| 5 | lncRNA | 75914216 | 75919546 | ENSMUSG00000106726 | MGI:5663237 | Gm43100 | predicted gene 43100 [Source:MGI Symbol;Acc:MGI:5663237] |
| 5 | lncRNA | 75914261 | 75914931 | ENSMUSG00000143297 | MGI:7829760 | Gm73502 | predicted gene, 73502 [Source:MGI Symbol;Acc:MGI:7829760] |
| 5 | lncRNA | 75993652 | 75996547 | ENSMUSG00000107061 | MGI:5011775 | Gm19590 | predicted gene, 19590 [Source:MGI Symbol;Acc:MGI:5011775] |
| 5 | lncRNA | 75996212 | 76001817 | ENSMUSG00000136464 | MGI:7827970 | Gm72604 | predicted gene, 72604 [Source:MGI Symbol;Acc:MGI:7827970] |
| 5 | lncRNA | 76007093 | 76009434 | ENSMUSG00000137410 | MGI:7828386 | Gm72812 | predicted gene, 72812 [Source:MGI Symbol;Acc:MGI:7828386] |
| 5 | lncRNA | 76009898 | 76014165 | ENSMUSG00000100936 | MGI:5579571 | Gm28865 | predicted gene 28865 [Source:MGI Symbol;Acc:MGI:5579571] |
| 5 | lncRNA | 76012661 | 76041839 | ENSMUSG00000139505 | MGI:7828920 | Gm73081 | predicted gene, 73081 [Source:MGI Symbol;Acc:MGI:7828920] |
| 5 | lncRNA | 76083539 | 76092833 | ENSMUSG00000134112 | MGI:7827374 | Gm72304 | predicted gene, 72304 [Source:MGI Symbol;Acc:MGI:7827374] |
| 5 | lncRNA | 76138510 | 76178133 | ENSMUSG00000120170 | MGI:5591886 | Gm32727 | predicted gene, 32727 [Source:MGI Symbol;Acc:MGI:5591886] |

| | | | | | | | |
|---|---|---|---|---|---|---|---|
| 5 | lncRNA | 76160044 | 76168012 | ENSMUSG00000126916 | MGI:7825765 | Gm71496 | predicted gene, 71496 [Source:MGI Symbol;Acc:MGI:7825765] |
| 5 | lncRNA | 76203710 | 76219434 | ENSMUSG00000106756 | MGI:5591939 | Gm32780 | predicted gene, 32780 [Source:MGI Symbol;Acc:MGI:5591939] |
| 5 | lncRNA | 76222715 | 76225128 | ENSMUSG00000143677 | MGI:7829952 | Gm73598 | predicted gene, 73598 [Source:MGI Symbol;Acc:MGI:7829952] |
| 5 | lncRNA | 76306112 | 76331994 | ENSMUSG00000137209 | MGI:5593097 | Gm33938 | predicted gene, 33938 [Source:MGI Symbol;Acc:MGI:5593097] |
| 5 | lncRNA | 76452723 | 76453374 | ENSMUSG00000098449 | MGI:3648350 | Gm7467 | predicted gene 7467 [Source:MGI Symbol;Acc:MGI:3648350] |
| 5 | lncRNA | 76479085 | 76489732 | ENSMUSG00000087037 | MGI:1921871 | 4930432L08Rik | RIKEN cDNA 4930432L08 gene [Source:MGI Symbol;Acc:MGI:1921871] |
| 5 | lncRNA | 76605488 | 76626206 | ENSMUSG00000143679 | MGI:7829954 | Gm73599 | predicted gene, 73599 [Source:MGI Symbol;Acc:MGI:7829954] |
| 5 | lncRNA | 76741593 | 76768288 | ENSMUSG00000124702 | MGI:7825559 | Gm71393 | predicted gene, 71393 [Source:MGI Symbol;Acc:MGI:7825559] |
| 5 | lncRNA | 76823250 | 76825017 | ENSMUSG00000132122 | MGI:7826714 | Gm71974 | predicted gene, 71974 [Source:MGI Symbol;Acc:MGI:7826714] |
| 5 | lncRNA | 76955148 | 76957770 | ENSMUSG00000132052 | MGI:7826694 | Gm71964 | predicted gene, 71964 [Source:MGI Symbol;Acc:MGI:7826694] |
| 5 | lncRNA | 76964249 | 76978788 | ENSMUSG00000132013 | MGI:6367246 | Gm52818 | predicted gene, 52818 [Source:MGI Symbol;Acc:MGI:6367246] |
| 5 | lncRNA | 76978887 | 76981092 | ENSMUSG00000132086 | MGI:7826708 | Gm71971 | predicted gene, 71971 [Source:MGI Symbol;Acc:MGI:7826708] |
| 5 | lncRNA | 77022258 | 77022921 | ENSMUSG00000137719 | MGI:7828472 | Gm72855 | predicted gene, 72855 [Source:MGI Symbol;Acc:MGI:7828472] |
| 5 | lncRNA | 77053362 | 77063549 | ENSMUSG00000137605 | MGI:7828454 | Gm72846 | predicted gene, 72846 [Source:MGI Symbol;Acc:MGI:7828454] |
| 5 | lncRNA | 77076215 | 77095605 | ENSMUSG00000121495 | MGI:7823087 | Gm70141 | predicted gene, 70141 [Source:MGI Symbol;Acc:MGI:7823087] |
| 5 | lncRNA | 77108327 | 77151020 | ENSMUSG00000137527 | MGI:6723603 | Gm54291 | predicted gene, 54291 [Source:MGI Symbol;Acc:MGI:6723603] |
| 5 | lncRNA | 77149301 | 77151561 | ENSMUSG00000137642 | | | novel transcript |
| 5 | lncRNA | 77150660 | 77166723 | ENSMUSG00000086632 | MGI:3696786 | 1700112J05Rik | RIKEN cDNA 1700112J05 gene [Source:MGI Symbol;Acc:MGI:3696786] |

| | | | | | | | |
|---|---|---|---|---|---|---|---|
| 5 | lncRNA | 77168164 | 77168929 | ENSMUSG00000137680 | MGI:7828468 | Gm72853 | predicted gene, 72853 [Source:MGI Symbol;Acc:MGI:7828468] |
| 5 | lncRNA | 77206097 | 77206721 | ENSMUSG00000137760 | MGI:7828478 | Gm72858 | predicted gene, 72858 [Source:MGI Symbol;Acc:MGI:7828478] |
| 5 | lncRNA | 77215347 | 77219129 | ENSMUSG00000127703 | MGI:7825835 | Gm71531 | predicted gene, 71531 [Source:MGI Symbol;Acc:MGI:7825835] |
| 5 | lncRNA | 77240550 | 77250160 | ENSMUSG00000087541 | MGI:3801917 | Hopxos | HOP homeobox, opposite strand [Source:MGI Symbol;Acc:MGI:3801917] |
| 5 | lncRNA | 77263428 | 77270195 | ENSMUSG00000085858 | MGI:3801916 | Gm15831 | predicted gene 15831 [Source:MGI Symbol;Acc:MGI:3801916] |
| 5 | lncRNA | 77271261 | 77278216 | ENSMUSG00000137325 | MGI:7828356 | Gm72797 | predicted gene, 72797 [Source:MGI Symbol;Acc:MGI:7828356] |
| 5 | lncRNA | 77274414 | 77287851 | ENSMUSG00000120897 | MGI:7338513 | Gm57300 | predicted gene, 57300 [Source:MGI Symbol;Acc:MGI:7338513] |
| 5 | lncRNA | 77283791 | 77288173 | ENSMUSG00000106783 | MGI:5624990 | Chaer1 | [illegible] regulator 1 [Source:MGI Symbol;Acc:MGI:5624990] |
| 5 | lncRNA | 77293927 | 77307047 | ENSMUSG00000130552 | MGI:5593757 | Gm34598 | predicted gene, 34598 [Source:MGI Symbol;Acc:MGI:5593757] |
| 5 | lncRNA | 77415158 | 77416490 | ENSMUSG00000131889 | MGI:7826644 | Gm71939 | predicted gene, 71939 [Source:MGI Symbol;Acc:MGI:7826644] |
| 5 | lncRNA | 77505738 | 77522641 | ENSMUSG00000141011 | MGI:7829276 | Gm73259 | predicted gene, 73259 [Source:MGI Symbol;Acc:MGI:7829276] |
| 5 | lncRNA | 77534435 | 77541539 | ENSMUSG00000106828 | MGI:5593807 | Gm34648 | predicted gene, 34648 [Source:MGI Symbol;Acc:MGI:5593807] |
| 5 | lncRNA | 77581834 | 77621066 | ENSMUSG00000107232 | MGI:1916620 | 1700017L05Rik | RIKEN cDNA 1700017L05 gene [Source:MGI Symbol;Acc:MGI:1916620] |
| 5 | lncRNA | 77610344 | 77613256 | ENSMUSG00000143522 | MGI:7829878 | Gm73561 | predicted gene, 73561 [Source:MGI Symbol;Acc:MGI:7829878] |
| 5 | lncRNA | 77630302 | 77642268 | ENSMUSG00000106985 | MGI:5662898 | Gm42761 | predicted gene 42761 [Source:MGI Symbol;Acc:MGI:5662898] |
| 5 | lncRNA | 77659041 | 77659595 | ENSMUSG00000121772 | MGI:7824207 | Gm70707 | predicted gene, 70707 [Source:MGI Symbol;Acc:MGI:7824207] |
| 5 | lncRNA | 77659055 | 77694753 | ENSMUSG00000123203 | MGI:7825375 | Gm71299 | predicted gene, 71299 [Source:MGI Symbol;Acc:MGI:7825375] |
| 5 | lncRNA | 77661046 | 77681112 | ENSMUSG00000123236 | MGI:7825379 | Gm71301 | predicted gene, 71301 [Source:MGI Symbol;Acc:MGI:7825379] |

| | | | | | | | |
|---|---|---|---|---|---|---|---|
| 5 | lncRNA | 77725236 | 77728017 | ENSMUSG00000129668 | MGI:7826011 | Gm71620 | predicted gene, 71620 [Source:MGI Symbol;Acc:MGI:7826011] |
| 5 | lncRNA | 77756653 | 77792554 | ENSMUSG00000133181 | MGI:7827006 | Gm72120 | predicted gene, 72120 [Source:MGI Symbol;Acc:MGI:7827006] |
| 5 | lncRNA | 77787663 | 77822531 | ENSMUSG00000133218 | MGI:7827046 | Gm72140 | predicted gene, 72140 [Source:MGI Symbol;Acc:MGI:7827046] |
| 5 | lncRNA | 77838851 | 77840079 | ENSMUSG00000123568 | MGI:7825419 | Gm71321 | predicted gene, 71321 [Source:MGI Symbol;Acc:MGI:7825419] |
| 5 | lncRNA | 77891693 | 77909996 | ENSMUSG00000138944 | MGI:7828760 | Gm73000 | predicted gene, 73000 [Source:MGI Symbol;Acc:MGI:7828760] |
| 5 | lncRNA | 77986533 | 78096901 | ENSMUSG00000127046 | | | novel transcript |
| 5 | lncRNA | 78987656 | 79002547 | ENSMUSG00000124166 | MGI:7825493 | Gm71359 | predicted gene, 71359 [Source:MGI Symbol;Acc:MGI:7825493] |
| 5 | lncRNA | 79002682 | 79048131 | ENSMUSG00000134542 | MGI:7827458 | Gm72346 | predicted gene, 72346 [Source:MGI Symbol;Acc:MGI:7827458] |
| 5 | lncRNA | 79298740 | 79299452 | ENSMUSG00000129544 | MGI:7825989 | Gm71609 | predicted gene, 71609 [Source:MGI Symbol;Acc:MGI:7825989] |
| 5 | lncRNA | 79785905 | 79786318 | ENSMUSG00000127094 | MGI:7825789 | Gm71508 | predicted gene, 71508 [Source:MGI Symbol;Acc:MGI:7825789] |
| 5 | lncRNA | 80797068 | 80800099 | ENSMUSG00000137734 | MGI:7828474 | Gm72856 | predicted gene, 72856 [Source:MGI Symbol;Acc:MGI:7828474] |
| 5 | lncRNA | 80925626 | 81002961 | ENSMUSG00000128907 | MGI:7825937 | Gm71582 | predicted gene, 71582 [Source:MGI Symbol;Acc:MGI:7825937] |
| 5 | lncRNA | 81171512 | 81176796 | ENSMUSG00000124141 | MGI:7825489 | Gm71357 | predicted gene, 71357 [Source:MGI Symbol;Acc:MGI:7825489] |
| 5 | lncRNA | 81701680 | 81703966 | ENSMUSG00000121076 | MGI:7338533 | Gm57310 | predicted gene, 57310 [Source:MGI Symbol;Acc:MGI:7338533] |
| 5 | lncRNA | 81939752 | 81946273 | ENSMUSG00000087431 | MGI:3801819 | Gm16054 | predicted gene 16054 [Source:MGI Symbol;Acc:MGI:3801819] |
| 5 | lncRNA | 82057851 | 82074134 | ENSMUSG00000132904 | MGI:7826882 | Gm72058 | predicted gene, 72058 [Source:MGI Symbol;Acc:MGI:7826882] |
| 5 | lncRNA | 82606825 | 82611673 | ENSMUSG00000136333 | MGI:7827932 | Gm72585 | predicted gene, 72585 [Source:MGI Symbol;Acc:MGI:7827932] |
| 5 | lncRNA | 82787205 | 82800131 | ENSMUSG00000135173 | MGI:7827642 | Gm72439 | predicted gene, 72439 [Source:MGI Symbol;Acc:MGI:7827642] |

| | | | | | | | |
|---|---|---|---|---|---|---|---|
| 5 | lncRNA | 82818662 | 82967715 | ENSMUSG00000123351 | MGI:7825395 | Gm71309 | predicted gene, 71309 [Source:MGI Symbol;Acc:MGI:7825395] |
| 5 | lncRNA | 83148035 | 83160095 | ENSMUSG00000128479 | MGI:7825903 | Gm71565 | predicted gene, 71565 [Source:MGI Symbol;Acc:MGI:7825903] |
| 5 | lncRNA | 83159825 | 83164649 | ENSMUSG00000128511 | MGI:7825907 | Gm71567 | predicted gene, 71567 [Source:MGI Symbol;Acc:MGI:7825907] |
| 5 | lncRNA | 83920313 | 83961863 | ENSMUSG00000122804 | MGI:7825313 | Gm71266 | predicted gene, 71266 [Source:MGI Symbol;Acc:MGI:7825313] |
| 5 | lncRNA | 83958251 | 83961799 | ENSMUSG00000122846 | MGI:7825327 | Gm71273 | predicted gene, 71273 [Source:MGI Symbol;Acc:MGI:7825327] |
| 5 | lncRNA | 83996968 | 84037431 | ENSMUSG00000137987 | MGI:7828508 | Gm72873 | predicted gene, 72873 [Source:MGI Symbol;Acc:MGI:7828508] |
| 5 | lncRNA | 84037198 | 84042514 | ENSMUSG00000138029 | MGI:7828516 | Gm72877 | predicted gene, 72877 [Source:MGI Symbol;Acc:MGI:7828516] |
| 5 | lncRNA | 84271075 | 84272260 | ENSMUSG00000138269 | MGI:7830314 | Gm73784 | predicted gene, 73784 [Source:MGI Symbol;Acc:MGI:7830314] |
| 5 | lncRNA | 84565315 | 84614936 | ENSMUSG00000132128 | MGI:7826718 | Gm71976 | predicted gene, 71976 [Source:MGI Symbol;Acc:MGI:7826718] |
| 5 | lncRNA | 84576664 | 84580479 | ENSMUSG00000106598 | MGI:5663669 | Gm43532 | predicted gene 43532 [Source:MGI Symbol;Acc:MGI:5663669] |
| 5 | lncRNA | 84860434 | 84861906 | ENSMUSG00000124137 | MGI:7825487 | Gm71356 | predicted gene, 71356 [Source:MGI Symbol;Acc:MGI:7825487] |
| 5 | lncRNA | 85848190 | 85848619 | ENSMUSG00000142369 | MGI:7829558 | Gm73400 | predicted gene, 73400 [Source:MGI Symbol;Acc:MGI:7829558] |
| 5 | lncRNA | 85853867 | 85859466 | ENSMUSG00000133322 | MGI:5594410 | Gm35251 | predicted gene, 35251 [Source:MGI Symbol;Acc:MGI:5594410] |
| 5 | lncRNA | 85868633 | 85870610 | ENSMUSG00000104878 | MGI:5663704 | Gm43567 | predicted gene 43567 [Source:MGI Symbol;Acc:MGI:5663704] |
| 5 | lncRNA | 86053129 | 86057573 | ENSMUSG00000138698 | MGI:7828670 | Gm72955 | predicted gene, 72955 [Source:MGI Symbol;Acc:MGI:7828670] |
| 5 | lncRNA | 86213524 | 86216687 | ENSMUSG00000130557 | MGI:7826228 | Gm71730 | predicted gene, 71730 [Source:MGI Symbol;Acc:MGI:7826228] |
| 5 | lncRNA | 86214671 | 86215754 | ENSMUSG00000121702 | MGI:7824203 | Gm70705 | predicted gene, 70705 [Source:MGI Symbol;Acc:MGI:7824203] |
| 5 | lncRNA | 86320884 | 86321767 | ENSMUSG00000132971 | MGI:7826904 | Gm72069 | predicted gene, 72069 [Source:MGI Symbol;Acc:MGI:7826904] |

| | | | | | | | |
|---|---|---|---|---|---|---|---|
| 5 | lncRNA | 86831709 | 86834499 | ENSMUSG00000106507 | MGI:5663193 | Gm43056 | predicted gene 43056 [Source:MGI Symbol;Acc:MGI:5663193] |
| 5 | lncRNA | 86876758 | 86907595 | ENSMUSG00000089851 | MGI:4414999 | Gm16579 | predicted gene 16579 [Source:MGI Symbol;Acc:MGI:4414999] |
| 5 | lncRNA | 86876765 | 86964965 | ENSMUSG00000105704 | MGI:5663192 | Gm43055 | predicted gene 43055 [Source:MGI Symbol;Acc:MGI:5663192] |
| 5 | lncRNA | 86914994 | 86930892 | ENSMUSG00000104937 | MGI:5663194 | Gm43057 | predicted gene 43057 [Source:MGI Symbol;Acc:MGI:5663194] |
| 5 | lncRNA | 86951343 | 86952166 | ENSMUSG00000124176 | MGI:7825497 | Gm71361 | predicted gene, 71361 [Source:MGI Symbol;Acc:MGI:7825497] |
| 5 | lncRNA | 87051725 | 87053341 | ENSMUSG00000130987 | MGI:7826348 | Gm71791 | predicted gene, 71791 [Source:MGI Symbol;Acc:MGI:7826348] |
| 5 | lncRNA | 87093304 | 87124242 | ENSMUSG00000131191 | MGI:7826430 | Gm71832 | predicted gene, 71832 [Source:MGI Symbol;Acc:MGI:7826430] |
| 5 | lncRNA | 87108370 | 87109945 | ENSMUSG00000131214 | MGI:7830368 | Gm73811 | predicted gene, 73811 [Source:MGI Symbol;Acc:MGI:7830368] |
| 5 | lncRNA | 87144907 | 87148517 | ENSMUSG00000136916 | MGI:7828234 | Gm72736 | predicted gene, 72736 [Source:MGI Symbol;Acc:MGI:7828234] |
| 5 | lncRNA | 87166312 | 87190056 | ENSMUSG00000122385 | MGI:7825259 | Gm71239 | predicted gene, 71239 [Source:MGI Symbol;Acc:MGI:7825259] |
| 5 | lncRNA | 87520452 | 87527815 | ENSMUSG00000122459 | MGI:7830372 | Gm73813 | predicted gene, 73813 [Source:MGI Symbol;Acc:MGI:7830372] |
| 5 | lncRNA | 87657298 | 87664390 | ENSMUSG00000141278 | MGI:7829378 | Gm73310 | predicted gene, 73310 [Source:MGI Symbol;Acc:MGI:7829378] |
| 5 | lncRNA | 87903460 | 87998719 | ENSMUSG00000104839 | MGI:5662931 | Gm42794 | predicted gene 42794 [Source:MGI Symbol;Acc:MGI:5662931] |
| 5 | lncRNA | 87998218 | 88000872 | ENSMUSG00000121715 | | | DAZ interacting protein 1 pseudogene [Source:NCBI gene (formerly Entrezgene);Acc:654494] |
| 5 | lncRNA | 88055713 | 88127236 | ENSMUSG00000099605 | MGI:1920721 | 1700066N21Rik | RIKEN cDNA 1700066N21 gene [Source:MGI Symbol;Acc:MGI:1920721] |
| 5 | lncRNA | 88111344 | 88114211 | ENSMUSG00000055961 | MGI:2670982 | BC051076 | cDNA sequence BC051076 [Source:MGI Symbol;Acc:MGI:2670982] |
| 5 | lncRNA | 88310430 | 88315615 | ENSMUSG00000139380 | MGI:7828856 | Gm73049 | predicted gene, 73049 [Source:MGI Symbol;Acc:MGI:7828856] |

| | | | | | | | |
|---|---|---|---|---|---|---|---|
| 5 | lncRNA | 88344830 | 88359055 | ENSMUSG00000139858 | | | novel transcript |
| 5 | lncRNA | 88439212 | 88599170 | ENSMUSG00000126970 | MGI:1920562 | 1700041A01Rik | RIKEN cDNA 1700041A01 gene [Source:MGI Symbol;Acc:MGI:1920562] |
| 5 | lncRNA | 88692315 | 88712827 | ENSMUSG00000120316 | MGI:7338457 | Gm57272 | predicted gene, 57272 [Source:MGI Symbol;Acc:MGI:7338457] |
| 5 | lncRNA | 88701671 | 88702091 | ENSMUSG00000134393 | MGI:7827442 | Gm72338 | predicted gene, 72338 [Source:MGI Symbol;Acc:MGI:7827442] |
| 5 | lncRNA | 88728109 | 88732185 | ENSMUSG00000134225 | MGI:7827422 | Gm72328 | predicted gene, 72328 [Source:MGI Symbol;Acc:MGI:7827422] |
| 5 | lncRNA | 88728253 | 88728997 | ENSMUSG00000134311 | MGI:7827436 | Gm72335 | predicted gene, 72335 [Source:MGI Symbol;Acc:MGI:7827436] |
| 5 | lncRNA | 88729359 | 88729963 | ENSMUSG00000134350 | MGI:7827438 | Gm72336 | predicted gene, 72336 [Source:MGI Symbol;Acc:MGI:7827438] |
| 5 | lncRNA | 88824082 | 88832390 | ENSMUSG00000129340 | MGI:7825973 | Gm71600 | predicted gene, 71600 [Source:MGI Symbol;Acc:MGI:7825973] |
| 5 | lncRNA | 88857948 | 88868692 | ENSMUSG00000121172 | MGI:7338561 | Gm57324 | predicted gene, 57324 [Source:MGI Symbol;Acc:MGI:7338561] |
| 5 | lncRNA | 88888201 | 88891403 | ENSMUSG00000132815 | MGI:7826846 | Gm72040 | predicted gene, 72040 [Source:MGI Symbol;Acc:MGI:7826846] |
| 5 | lncRNA | 88941693 | 88961351 | ENSMUSG00000105640 | MGI:5663049 | Gm42912 | predicted gene 42912 [Source:MGI Symbol;Acc:MGI:5663049] |
| 5 | lncRNA | 89351748 | 89353489 | ENSMUSG00000126258 | MGI:7825697 | Gm71462 | predicted gene, 71462 [Source:MGI Symbol;Acc:MGI:7825697] |
| 5 | lncRNA | 89356080 | 89356956 | ENSMUSG00000121166 | MGI:7338557 | Gm57322 | predicted gene, 57322 [Source:MGI Symbol;Acc:MGI:7338557] |
| 5 | lncRNA | 89370386 | 89371742 | ENSMUSG00000130398 | MGI:7826210 | Gm71721 | predicted gene, 71721 [Source:MGI Symbol;Acc:MGI:7826210] |
| 5 | lncRNA | 89393196 | 89410563 | ENSMUSG00000143172 | MGI:7829738 | Gm73491 | predicted gene, 73491 [Source:MGI Symbol;Acc:MGI:7829738] |
| 5 | lncRNA | 89425606 | 89434146 | ENSMUSG00000142798 | MGI:7829670 | Gm73457 | predicted gene, 73457 [Source:MGI Symbol;Acc:MGI:7829670] |
| 5 | lncRNA | 89503065 | 89589568 | ENSMUSG00000137094 | MGI:7828294 | Gm72766 | predicted gene, 72766 [Source:MGI Symbol;Acc:MGI:7828294] |
| 5 | lncRNA | 89537350 | 89540646 | ENSMUSG00000137136 | MGI:7828298 | Gm72768 | predicted gene, 72768 [Source:MGI Symbol;Acc:MGI:7828298] |

| | | | | | | | |
|---|---|---|---|---|---|---|---|
| 5 | lncRNA | 89540266 | 89541409 | ENSMUSG00000137177 | MGI:7828300 | Gm72769 | predicted gene, 72769 [Source:MGI Symbol;Acc:MGI:7828300] |
| 5 | lncRNA | 89611209 | 89647450 | ENSMUSG00000126175 | MGI:7825685 | Gm71456 | predicted gene, 71456 [Source:MGI Symbol;Acc:MGI:7825685] |
| 5 | lncRNA | 89832053 | 89837636 | ENSMUSG00000126217 | MGI:7825689 | Gm71458 | predicted gene, 71458 [Source:MGI Symbol;Acc:MGI:7825689] |
| 5 | lncRNA | 90099232 | 90125740 | ENSMUSG00000130148 | MGI:7826134 | Gm71683 | predicted gene, 71683 [Source:MGI Symbol;Acc:MGI:7826134] |
| 5 | lncRNA | 90267684 | 90298205 | ENSMUSG00000135627 | MGI:7827746 | Gm72491 | predicted gene, 72491 [Source:MGI Symbol;Acc:MGI:7827746] |
| 5 | lncRNA | 90341182 | 90343029 | ENSMUSG00000140834 | MGI:7829230 | Gm73236 | predicted gene, 73236 [Source:MGI Symbol;Acc:MGI:7829230] |
| 5 | lncRNA | 90344681 | 90348377 | ENSMUSG00000140416 | MGI:7829148 | Gm73195 | predicted gene, 73195 [Source:MGI Symbol;Acc:MGI:7829148] |
| 5 | lncRNA | 90394656 | 90436607 | ENSMUSG00000130758 | MGI:7826284 | Gm71758 | predicted gene, 71758 [Source:MGI Symbol;Acc:MGI:7826284] |
| 5 | lncRNA | 90514479 | 90516352 | ENSMUSG00000054945 | MGI:3641866 | Gm9958 | predicted gene 9958 [Source:MGI Symbol;Acc:MGI:3641866] |
| 5 | lncRNA | 90562326 | 90587746 | ENSMUSG00000123171 | MGI:5595158 | Gm35999 | predicted gene, 35999 [Source:MGI Symbol;Acc:MGI:5595158] |
| 5 | lncRNA | 90646939 | 90663790 | ENSMUSG00000106788 | MGI:5663586 | Gm43449 | predicted gene 43449 [Source:MGI Symbol;Acc:MGI:5663586] |
| 5 | lncRNA | 90701058 | 90736936 | ENSMUSG00000106863 | MGI:5624994 | Gm42109 | predicted gene, 42109 [Source:MGI Symbol;Acc:MGI:5624994] |
| 5 | lncRNA | 90750130 | 90754368 | ENSMUSG00000139329 | MGI:7828840 | Gm73041 | predicted gene, 73041 [Source:MGI Symbol;Acc:MGI:7828840] |
| 5 | lncRNA | 90796599 | 90828142 | ENSMUSG00000130379 | MGI:7826200 | Gm71716 | predicted gene, 71716 [Source:MGI Symbol;Acc:MGI:7826200] |
| 5 | lncRNA | 90860787 | 90876723 | ENSMUSG00000141359 | MGI:5595243 | Gm36084 | predicted gene, 36084 [Source:MGI Symbol;Acc:MGI:5595243] |
| 5 | lncRNA | 90906282 | 90910220 | ENSMUSG00000135043 | MGI:7827592 | Gm72413 | predicted gene, 72413 [Source:MGI Symbol;Acc:MGI:7827592] |
| 5 | lncRNA | 90926762 | 90933050 | ENSMUSG00000136230 | MGI:7827894 | Gm72565 | predicted gene, 72565 [Source:MGI Symbol;Acc:MGI:7827894] |
| 5 | lncRNA | 90980187 | 90983810 | ENSMUSG00000107044 | MGI:5663223 | Gm43086 | predicted gene 43086 [Source:MGI Symbol;Acc:MGI:5663223] |

| | | | | | | | |
|---|---|---|---|---|---|---|---|
| 5 | lncRNA | 91007891 | 91036327 | ENSMUSG00000142520 | MGI:7829610 | Gm73427 | predicted gene, 73427 [Source:MGI Symbol;Acc:MGI:7829610] |
| 5 | lncRNA | 91047056 | 91051063 | ENSMUSG00000140717 | MGI:7829222 | Gm73232 | predicted gene, 73232 [Source:MGI Symbol;Acc:MGI:7829222] |
| 5 | lncRNA | 91070591 | 91072764 | ENSMUSG00000107218 | MGI:5662667 | Gm42530 | predicted gene 42530 [Source:MGI Symbol;Acc:MGI:5662667] |
| 5 | lncRNA | 91158799 | 91162369 | ENSMUSG00000131708 | MGI:6367248 | Gm52819 | predicted gene, 52819 [Source:MGI Symbol;Acc:MGI:6367248] |
| 5 | lncRNA | 91198968 | 91283840 | ENSMUSG00000107350 | MGI:5011795 | Gm19610 | predicted gene, 19610 [Source:MGI Symbol;Acc:MGI:5011795] |
| 5 | lncRNA | 91399459 | 91455916 | ENSMUSG00000107198 | MGI:5011804 | Gm19619 | predicted gene, 19619 [Source:MGI Symbol;Acc:MGI:5011804] |
| 5 | lncRNA | 91572101 | 91597783 | ENSMUSG00000133134 | MGI:5624995 | Gm42110 | predicted gene, 42110 [Source:MGI Symbol;Acc:MGI:5624995] |
| 5 | lncRNA | 91596864 | 91604421 | ENSMUSG00000133170 | MGI:7826996 | Gm72115 | predicted gene, 72115 [Source:MGI Symbol;Acc:MGI:7826996] |
| 5 | lncRNA | 91838925 | 91844544 | ENSMUSG00000136387 | MGI:7827950 | Gm72594 | predicted gene, 72594 [Source:MGI Symbol;Acc:MGI:7827950] |
| 5 | lncRNA | 91856334 | 91889188 | ENSMUSG00000129304 | MGI:5595381 | Gm36222 | predicted gene, 36222 [Source:MGI Symbol;Acc:MGI:5595381] |
| 5 | lncRNA | 91888385 | 91915778 | ENSMUSG00000099468 | MGI:5578977 | Gm28271 | predicted gene 28271 [Source:MGI Symbol;Acc:MGI:5578977] |
| 5 | lncRNA | 91974673 | 91992960 | ENSMUSG00000136162 | MGI:7827844 | Gm72540 | predicted gene, 72540 [Source:MGI Symbol;Acc:MGI:7827844] |
| 5 | lncRNA | 91979238 | 92000481 | ENSMUSG00000136120 | MGI:5595462 | Gm36303 | predicted gene, 36303 [Source:MGI Symbol;Acc:MGI:5595462] |
| 5 | lncRNA | 92009494 | 92092978 | ENSMUSG00000143999 | MGI:5595516 | Gm36357 | predicted gene, 36357 [Source:MGI Symbol;Acc:MGI:5595516] |
| 5 | lncRNA | 92059257 | 92061279 | ENSMUSG00000107301 | MGI:5663825 | Gm43688 | predicted gene 43688 [Source:MGI Symbol;Acc:MGI:5663825] |
| 5 | lncRNA | 92110248 | 92170798 | ENSMUSG00000121865 | | | THAP domain containing 6 [Source:NCBI gene (formerly Entrezgene);Acc:381650] |
| 5 | lncRNA | 92116719 | 92127032 | ENSMUSG00000129673 | MGI:7826013 | Gm71621 | predicted gene, 71621 [Source:MGI Symbol;Acc:MGI:7826013] |
| 5 | lncRNA | 92190660 | 92201237 | ENSMUSG00000129634 | MGI:7826007 | Gm71618 | predicted gene, 71618 [Source:MGI Symbol;Acc:MGI:7826007] |

| | | | | | | | |
|---|---|---|---|---|---|---|---|
| 5 | lncRNA | 92198199 | 92198445 | ENSMUSG00000129945 | MGI:7826102 | Gm71666 | predicted gene, 71666 [Source:MGI Symbol;Acc:MGI:7826102] |
| 5 | lncRNA | 92230778 | 92233700 | ENSMUSG00000129828 | MGI:7826064 | Gm71647 | predicted gene, 71647 [Source:MGI Symbol;Acc:MGI:7826064] |
| 5 | lncRNA | 92232457 | 92233892 | ENSMUSG00000129902 | MGI:7826082 | Gm71656 | predicted gene, 71656 [Source:MGI Symbol;Acc:MGI:7826082] |
| 5 | lncRNA | 92273407 | 92285584 | ENSMUSG00000129597 | MGI:7825999 | Gm71614 | predicted gene, 71614 [Source:MGI Symbol;Acc:MGI:7825999] |
| 5 | lncRNA | 92274877 | 92277972 | ENSMUSG00000120423 | MGI:2444469 | D430040L24Rik | RIKEN cDNA D430040L24 gene [Source:MGI Symbol;Acc:MGI:2444469] |
| 5 | lncRNA | 92341678 | 92345320 | ENSMUSG00000129758 | MGI:7826025 | Gm71627 | predicted gene, 71627 [Source:MGI Symbol;Acc:MGI:7826025] |
| 5 | lncRNA | 92358337 | 92360533 | ENSMUSG00000129868 | MGI:5624998 | Gm42113 | predicted gene, 42113 [Source:MGI Symbol;Acc:MGI:5624998] |
| 5 | lncRNA | 92364403 | 92372316 | ENSMUSG00000129715 | MGI:5624997 | Gm42112 | predicted gene, 42112 [Source:MGI Symbol;Acc:MGI:5624997] |
| 5 | lncRNA | 92437911 | 92450619 | ENSMUSG00000106698 | MGI:5663736 | Gm43599 | predicted gene 43599 [Source:MGI Symbol;Acc:MGI:5663736] |
| 5 | lncRNA | 92582183 | 92583916 | ENSMUSG00000131613 | MGI:5624999 | Gm42114 | predicted gene, 42114 [Source:MGI Symbol;Acc:MGI:5624999] |
| 5 | lncRNA | 92653679 | 92660306 | ENSMUSG00000130185 | MGI:7826142 | Gm71687 | predicted gene, 71687 [Source:MGI Symbol;Acc:MGI:7826142] |
| 5 | lncRNA | 92690902 | 92699275 | ENSMUSG00000138080 | MGI:7828520 | Gm72879 | predicted gene, 72879 [Source:MGI Symbol;Acc:MGI:7828520] |
| 5 | lncRNA | 92822888 | 92824425 | ENSMUSG00000139854 | MGI:7828962 | Gm73102 | predicted gene, 73102 [Source:MGI Symbol;Acc:MGI:7828962] |
| 5 | lncRNA | 92830033 | 92831611 | ENSMUSG00000141749 | MGI:7829442 | Gm73342 | predicted gene, 73342 [Source:MGI Symbol;Acc:MGI:7829442] |
| 5 | lncRNA | 92830518 | 92831151 | ENSMUSG00000141780 | MGI:7829448 | Gm73345 | predicted gene, 73345 [Source:MGI Symbol;Acc:MGI:7829448] |
| 5 | lncRNA | 92884064 | 92885301 | ENSMUSG00000092266 | MGI:5141965 | Gm20500 | predicted gene 20500 [Source:MGI Symbol;Acc:MGI:5141965] |
| 5 | lncRNA | 93065634 | 93073037 | ENSMUSG00000142400 | MGI:7829566 | Gm73404 | predicted gene, 73404 [Source:MGI Symbol;Acc:MGI:7829566] |
| 5 | lncRNA | 93151472 | 93167612 | ENSMUSG00000140081 | MGI:5595923 | Gm36764 | predicted gene, 36764 [Source:MGI Symbol;Acc:MGI:5595923] |

| | | | | | | | |
|---|---|---|---|---|---|---|---|
| 5 | lncRNA | 93153559 | 93162449 | ENSMUSG00000140123 | MGI:7829050 | Gm73146 | predicted gene, 73146 [Source:MGI Symbol;Acc:MGI:7829050] |
| 5 | lncRNA | 93174253 | 93176495 | ENSMUSG00000140404 | MGI:7829138 | Gm73190 | predicted gene, 73190 [Source:MGI Symbol;Acc:MGI:7829138] |
| 5 | lncRNA | 93185357 | 93186588 | ENSMUSG00000128457 | MGI:7825899 | Gm71563 | predicted gene, 71563 [Source:MGI Symbol;Acc:MGI:7825899] |
| 5 | lncRNA | 93193166 | 93199034 | ENSMUSG00000133837 | MGI:7827236 | Gm72235 | predicted gene, 72235 [Source:MGI Symbol;Acc:MGI:7827236] |
| 5 | lncRNA | 93225033 | 93241274 | ENSMUSG00000129086 | MGI:7825953 | Gm71590 | predicted gene, 71590 [Source:MGI Symbol;Acc:MGI:7825953] |
| 5 | lncRNA | 93304425 | 93309887 | ENSMUSG00000122927 | MGI:7825338 | Gm71280 | predicted gene, 71280 [Source:MGI Symbol;Acc:MGI:7825338] |
| 5 | lncRNA | 93353325 | 93353458 | ENSMUSG00000122968 | MGI:7825342 | Gm71282 | predicted gene, 71282 [Source:MGI Symbol;Acc:MGI:7825342] |
| 5 | lncRNA | 93365844 | 93388691 | ENSMUSG00000125536 | MGI:7825633 | Gm71430 | predicted gene, 71430 [Source:MGI Symbol;Acc:MGI:7825633] |
| 5 | lncRNA | 93452242 | 93454353 | ENSMUSG00000134089 | MGI:7827340 | Gm72287 | predicted gene, 72287 [Source:MGI Symbol;Acc:MGI:7827340] |
| 5 | lncRNA | 93714055 | 93723875 | ENSMUSG00000125231 | MGI:7825599 | Gm71413 | predicted gene, 71413 [Source:MGI Symbol;Acc:MGI:7825599] |
| 5 | lncRNA | 94168402 | 94212535 | ENSMUSG00000126234 | MGI:7825695 | Gm71461 | predicted gene, 71461 [Source:MGI Symbol;Acc:MGI:7825695] |
| 5 | lncRNA | 94424246 | 94449998 | ENSMUSG00000130196 | MGI:7826144 | Gm71688 | predicted gene, 71688 [Source:MGI Symbol;Acc:MGI:7826144] |
| 5 | lncRNA | 94450328 | 94454655 | ENSMUSG00000121888 | | | PRAME like 39, pseudogene [Source:NCBI gene (formerly Entrezgene);Acc:331195] |
| 5 | lncRNA | 94487608 | 94532938 | ENSMUSG00000121889 | | | PRAME like 52, pseudogene [Source:NCBI gene (formerly Entrezgene);Acc:100554] |
| 5 | lncRNA | 94553273 | 94562789 | ENSMUSG00000125893 | MGI:7825663 | Gm71445 | predicted gene, 71445 [Source:MGI Symbol;Acc:MGI:7825663] |
| 5 | lncRNA | 94734291 | 94759548 | ENSMUSG00000129583 | MGI:7825997 | Gm71613 | predicted gene, 71613 [Source:MGI Symbol;Acc:MGI:7825997] |
| 5 | lncRNA | 94914344 | 94916550 | ENSMUSG00000139846 | MGI:7828960 | Gm73101 | predicted gene, 73101 [Source:MGI Symbol;Acc:MGI:7828960] |
| 5 | lncRNA | 95073209 | 95098491 | ENSMUSG00000125730 | MGI:5625003 | Gm42118 | predicted gene, 42118 [Source:MGI Symbol;Acc:MGI:5625003] |

| | | | | | | | |
|---|---|---|---|---|---|---|---|
| 5 | lncRNA | 95136457 | 95171649 | ENSMUSG00000121887 | MGI:7824219 | Gm70713 | predicted gene, 70713 [Source:MGI Symbol;Acc:MGI:7824219] |
| 5 | lncRNA | 95390289 | 95431161 | ENSMUSG00000138988 | MGI:7830506 | Gm73881 | predicted gene, 73881 [Source:MGI Symbol;Acc:MGI:7830506] |
| 5 | lncRNA | 95751675 | 95753276 | ENSMUSG00000140862 | MGI:7830514 | Gm73885 | predicted gene, 73885 [Source:MGI Symbol;Acc:MGI:7830514] |
| 5 | lncRNA | 95834235 | 95874723 | ENSMUSG00000127000 | MGI:7825775 | Gm71501 | predicted gene, 71501 [Source:MGI Symbol;Acc:MGI:7825775] |
| 5 | lncRNA | 96310583 | 96310919 | ENSMUSG00000135924 | MGI:7827792 | Gm72514 | predicted gene, 72514 [Source:MGI Symbol;Acc:MGI:7827792] |
| 5 | lncRNA | 96351064 | 96357927 | ENSMUSG00000133365 | MGI:7827136 | Gm72185 | predicted gene, 72185 [Source:MGI Symbol;Acc:MGI:7827136] |
| 5 | lncRNA | 96512180 | 96521816 | ENSMUSG00000135575 | MGI:7827738 | Gm72487 | predicted gene, 72487 [Source:MGI Symbol;Acc:MGI:7827738] |
| 5 | lncRNA | 96525726 | 96560691 | ENSMUSG00000124901 | MGI:7825573 | Gm71400 | predicted gene, 71400 [Source:MGI Symbol;Acc:MGI:7825573] |
| 5 | lncRNA | 96604248 | 96605611 | ENSMUSG00000106263 | MGI:5592209 | Gm33050 | predicted gene, 33050 [Source:MGI Symbol;Acc:MGI:5592209] |
| 5 | lncRNA | 96881997 | 96887570 | ENSMUSG00000106526 | MGI:5662741 | Gm42604 | predicted gene 42604 [Source:MGI Symbol;Acc:MGI:5662741] |
| 5 | lncRNA | 96963839 | 96966838 | ENSMUSG00000134078 | MGI:7827334 | Gm72284 | predicted gene, 72284 [Source:MGI Symbol;Acc:MGI:7827334] |
| 5 | lncRNA | 96988999 | 96989775 | ENSMUSG00000134123 | MGI:7827376 | Gm72305 | predicted gene, 72305 [Source:MGI Symbol;Acc:MGI:7827376] |
| 5 | lncRNA | 96997301 | 97000805 | ENSMUSG00000104955 | MGI:1916674 | 1700016F12Rik | RIKEN cDNA 1700016F12 gene [Source:MGI Symbol;Acc:MGI:1916674] |
| 5 | lncRNA | 97004518 | 97005073 | ENSMUSG00000123827 | MGI:7825449 | Gm71336 | predicted gene, 71336 [Source:MGI Symbol;Acc:MGI:7825449] |
| 5 | lncRNA | 97016638 | 97028257 | ENSMUSG00000133145 | MGI:7826990 | Gm72112 | predicted gene, 72112 [Source:MGI Symbol;Acc:MGI:7826990] |
| 5 | lncRNA | 97018970 | 97106952 | ENSMUSG00000106357 | MGI:3648644 | Gm8013 | predicted gene 8013 [Source:MGI Symbol;Acc:MGI:3648644] |
| 5 | lncRNA | 97045768 | 97089184 | ENSMUSG00000105283 | MGI:5592529 | Gm33370 | predicted gene, 33370 [Source:MGI Symbol;Acc:MGI:5592529] |
| 5 | lncRNA | 97104075 | 97131508 | ENSMUSG00000133056 | MGI:7826962 | Gm72098 | predicted gene, 72098 [Source:MGI Symbol;Acc:MGI:7826962] |

| | | | | | | | |
|---|---|---|---|---|---|---|---|
| 5 | lncRNA | 97111030 | 97111176 | ENSMUSG00000133184 | MGI:7827008 | Gm72121 | predicted gene, 72121 [Source:MGI Symbol;Acc:MGI:7827008] |
| 5 | lncRNA | 97144065 | 97145223 | ENSMUSG00000105006 | MGI:3779894 | Gm9484 | predicted gene 9484 [Source:MGI Symbol;Acc:MGI:3779894] |
| 5 | lncRNA | 97258944 | 97357574 | ENSMUSG00000104736 | MGI:5592768 | Gm33609 | predicted gene, 33609 [Source:MGI Symbol;Acc:MGI:5592768] |
| 5 | lncRNA | 97356287 | 97371487 | ENSMUSG00000133503 | MGI:6723653 | Gm54317 | predicted gene, 54317 [Source:MGI Symbol;Acc:MGI:6723653] |
| 5 | lncRNA | 97411967 | 97412790 | ENSMUSG00000131920 | MGI:7826656 | Gm71945 | predicted gene, 71945 [Source:MGI Symbol;Acc:MGI:7826656] |
| 5 | lncRNA | 97418130 | 97419368 | ENSMUSG00000131486 | MGI:7826538 | Gm71886 | predicted gene, 71886 [Source:MGI Symbol;Acc:MGI:7826538] |
| 5 | lncRNA | 97445838 | 97475349 | ENSMUSG00000135690 | MGI:7827764 | Gm72500 | predicted gene, 72500 [Source:MGI Symbol;Acc:MGI:7827764] |
| 5 | lncRNA | 97449018 | 97497427 | ENSMUSG00000104717 | MGI:5593128 | Gm33969 | predicted gene, 33969 [Source:MGI Symbol;Acc:MGI:5593128] |
| 5 | lncRNA | 97503355 | 97511173 | ENSMUSG00000141986 | MGI:7829476 | Gm73359 | predicted gene, 73359 [Source:MGI Symbol;Acc:MGI:7829476] |
| 5 | lncRNA | 97540251 | 97790042 | ENSMUSG00000097516 | MGI:1923064 | 4930467D21Rik | RIKEN cDNA 4930467D21 gene [Source:MGI Symbol;Acc:MGI:1923064] |
| 5 | lncRNA | 97546428 | 97547658 | ENSMUSG00000125869 | MGI:7825659 | Gm71443 | predicted gene, 71443 [Source:MGI Symbol;Acc:MGI:7825659] |
| 5 | lncRNA | 97685798 | 97702355 | ENSMUSG00000125831 | MGI:7825651 | Gm71439 | predicted gene, 71439 [Source:MGI Symbol;Acc:MGI:7825651] |
| 5 | lncRNA | 97848655 | 97980411 | ENSMUSG00000120552 | MGI:6723655 | Gm54318 | predicted gene, 54318 [Source:MGI Symbol;Acc:MGI:6723655] |
| 5 | lncRNA | 97969825 | 97987221 | ENSMUSG00000137242 | MGI:5593256 | Gm34097 | predicted gene, 34097 [Source:MGI Symbol;Acc:MGI:5593256] |
| 5 | lncRNA | 98119363 | 98127015 | ENSMUSG00000127655 | MGI:7825831 | Gm71529 | predicted gene, 71529 [Source:MGI Symbol;Acc:MGI:7825831] |
| 5 | lncRNA | 98164242 | 98175481 | ENSMUSG00000127610 | MGI:5593385 | Gm34226 | predicted gene, 34226 [Source:MGI Symbol;Acc:MGI:5593385] |
| 5 | lncRNA | 98226134 | 98228326 | ENSMUSG00000143266 | MGI:7829754 | Gm73499 | predicted gene, 73499 [Source:MGI Symbol;Acc:MGI:7829754] |
| 5 | lncRNA | 98282899 | 98286917 | ENSMUSG00000130319 | MGI:5625006 | Gm42121 | predicted gene, 42121 [Source:MGI Symbol;Acc:MGI:5625006] |

| | | | | | | | |
|---|---|---|---|---|---|---|---|
| 5 | lncRNA | 98330980 | 98333843 | ENSMUSG00000045238 | MGI:3696858 | A730035I17Rik | RIKEN cDNA A730035I17 gene [Source:MGI Symbol;Acc:MGI:3696858] |
| 5 | lncRNA | 98473766 | 98477169 | ENSMUSG00000136326 | MGI:7827928 | Gm72583 | predicted gene, 72583 [Source:MGI Symbol;Acc:MGI:7827928] |
| 5 | lncRNA | 98627618 | 98661123 | ENSMUSG00000136178 | MGI:5593634 | Gm34475 | predicted gene, 34475 [Source:MGI Symbol;Acc:MGI:5593634] |
| 5 | lncRNA | 98686346 | 98690045 | ENSMUSG00000136289 | MGI:7827920 | Gm72579 | predicted gene, 72579 [Source:MGI Symbol;Acc:MGI:7827920] |
| 5 | lncRNA | 98701480 | 98714873 | ENSMUSG00000067101 | MGI:1922750 | 1700010H22Rik | RIKEN cDNA 1700010H22 gene [Source:MGI Symbol;Acc:MGI:1922750] |
| 5 | lncRNA | 98714907 | 98720731 | ENSMUSG00000136251 | MGI:5625007 | Gm42122 | predicted gene, 42122 [Source:MGI Symbol;Acc:MGI:5625007] |
| 5 | lncRNA | 98981527 | 99002388 | ENSMUSG00000123838 | MGI:5593722 | Gm34563 | predicted gene, 34563 [Source:MGI Symbol;Acc:MGI:5593722] |
| 5 | lncRNA | 99026039 | 99039362 | ENSMUSG00000122946 | MGI:5625008 | Gm42123 | predicted gene, 42123 [Source:MGI Symbol;Acc:MGI:5625008] |
| 5 | lncRNA | 99185150 | 99190769 | ENSMUSG00000137970 | MGI:7828506 | Gm72872 | predicted gene, 72872 [Source:MGI Symbol;Acc:MGI:7828506] |
| 5 | lncRNA | 99186922 | 99319280 | ENSMUSG00000105579 | MGI:5663388 | Gm43251 | predicted gene 43251 [Source:MGI Symbol;Acc:MGI:5663388] |
| 5 | lncRNA | 99368625 | 99369114 | ENSMUSG00000125182 | MGI:7825591 | Gm71409 | predicted gene, 71409 [Source:MGI Symbol;Acc:MGI:7825591] |
| 5 | lncRNA | 99421954 | 99427143 | ENSMUSG00000105686 | MGI:5594553 | Gm35394 | predicted gene, 35394 [Source:MGI Symbol;Acc:MGI:5594553] |
| 5 | lncRNA | 99507200 | 99507903 | ENSMUSG00000129574 | MGI:7825993 | Gm71611 | predicted gene, 71611 [Source:MGI Symbol;Acc:MGI:7825993] |
| 5 | lncRNA | 99639936 | 99650712 | ENSMUSG00000140090 | MGI:7829036 | Gm73139 | predicted gene, 73139 [Source:MGI Symbol;Acc:MGI:7829036] |
| 5 | lncRNA | 99656208 | 99658940 | ENSMUSG00000104753 | MGI:5594331 | Gm35172 | predicted gene, 35172 [Source:MGI Symbol;Acc:MGI:5594331] |
| 5 | lncRNA | 99657729 | 99658626 | ENSMUSG00000090006 | MGI:3801791 | Gm16227 | predicted gene 16227 [Source:MGI Symbol;Acc:MGI:3801791] |
| 5 | lncRNA | 99765244 | 99774928 | ENSMUSG00000090146 | MGI:1921068 | 4930405H06Rik | RIKEN cDNA 4930405H06 gene [Source:MGI Symbol;Acc:MGI:1921068] |
| 5 | lncRNA | 99830241 | 99863867 | ENSMUSG00000089743 | MGI:3801792 | Gm16226 | predicted gene 16226 [Source:MGI Symbol;Acc:MGI:3801792] |

| | | | | | | | |
|---|---|---|---|---|---|---|---|
| 5 | lncRNA | 99862493 | 99869647 | ENSMUSG00000140204 | MGI:7829064 | Gm73153 | predicted gene, 73153 [Source:MGI Symbol;Acc:MGI:7829064] |
| 5 | lncRNA | 99878688 | 99902070 | ENSMUSG00000105218 | MGI:5621298 | Gm38413 | predicted gene, 38413 [Source:MGI Symbol;Acc:MGI:5621298] |
| 5 | lncRNA | 1.00E+08 | 1.00E+08 | ENSMUSG00000132178 | MGI:7826730 | Gm71982 | predicted gene, 71982 [Source:MGI Symbol;Acc:MGI:7826730] |
| 5 | lncRNA | 1.00E+08 | 1.00E+08 | ENSMUSG00000141515 | MGI:7829396 | Gm73319 | predicted gene, 73319 [Source:MGI Symbol;Acc:MGI:7829396] |
| 5 | lncRNA | 1.00E+08 | 1.00E+08 | ENSMUSG00000091623 | MGI:4937919 | Gm17092 | predicted gene 17092 [Source:MGI Symbol;Acc:MGI:4937919] |
| 5 | lncRNA | 1.00E+08 | 1.00E+08 | ENSMUSG00000126730 | MGI:7825749 | Gm71488 | predicted gene, 71488 [Source:MGI Symbol;Acc:MGI:7825749] |
| 5 | lncRNA | 1.00E+08 | 1.00E+08 | ENSMUSG00000140370 | MGI:7829108 | Gm73175 | predicted gene, 73175 [Source:MGI Symbol;Acc:MGI:7829108] |
| 5 | lncRNA | 1.00E+08 | 1.00E+08 | ENSMUSG00000086259 | MGI:1923253 | Tmem150cos | transmembrane protein 150C, opposite strand [Source:MGI Symbol;Acc:MGI:1923253] |
| 5 | lncRNA | 1.00E+08 | 1.00E+08 | ENSMUSG00000136233 | MGI:7827896 | Gm72566 | predicted gene, 72566 [Source:MGI Symbol;Acc:MGI:7827896] |
| 5 | lncRNA | 1.00E+08 | 1.00E+08 | ENSMUSG00000136314 | MGI:7827924 | Gm72581 | predicted gene, 72581 [Source:MGI Symbol;Acc:MGI:7827924] |
| 5 | lncRNA | 1.00E+08 | 1.00E+08 | ENSMUSG00000100157 | MGI:1916821 | 2310034O05Rik | RIKEN cDNA 2310034O05 gene [Source:MGI Symbol;Acc:MGI:1916821] |
| 5 | lncRNA | 1.00E+08 | 1.00E+08 | ENSMUSG00000144029 | MGI:7830096 | Gm73672 | predicted gene, 73672 [Source:MGI Symbol;Acc:MGI:7830096] |
| 5 | lncRNA | 1.00E+08 | 1.00E+08 | ENSMUSG00000121065 | MGI:7338529 | Gm57308 | predicted gene, 57308 [Source:MGI Symbol;Acc:MGI:7338529] |
| 5 | lncRNA | 1.00E+08 | 1.00E+08 | ENSMUSG00000130915 | | | novel transcript |
| 5 | lncRNA | 1.00E+08 | 1.00E+08 | ENSMUSG00000131020 | MGI:7826350 | Gm71792 | predicted gene, 71792 [Source:MGI Symbol;Acc:MGI:7826350] |
| 5 | lncRNA | 1.00E+08 | 1.00E+08 | ENSMUSG00000131094 | MGI:5625013 | Gm42128 | predicted gene, 42128 [Source:MGI Symbol;Acc:MGI:5625013] |
| 5 | lncRNA | 1.00E+08 | 1.00E+08 | ENSMUSG00000130982 | MGI:7826346 | Gm71790 | predicted gene, 71790 [Source:MGI Symbol;Acc:MGI:7826346] |
| 5 | lncRNA | 1.00E+08 | 1.00E+08 | ENSMUSG00000130947 | MGI:5825550 | Gm45913 | predicted gene, 45913 [Source:MGI Symbol;Acc:MGI:5825550] |

| | | | | | | | |
|---|---|---|---|---|---|---|---|
| 5 | lncRNA | 1.00E+08 | 1.00E+08 | ENSMUSG00000131052 | MGI:7826364 | Gm71799 | predicted gene, 71799 [Source:MGI Symbol;Acc:MGI:7826364] |
| 5 | lncRNA | 1.01E+08 | 1.01E+08 | ENSMUSG00000138356 | MGI:7828568 | Gm72904 | predicted gene, 72904 [Source:MGI Symbol;Acc:MGI:7828568] |
| 5 | lncRNA | 1.01E+08 | 1.01E+08 | ENSMUSG00000138317 | MGI:7828556 | Gm72898 | predicted gene, 72898 [Source:MGI Symbol;Acc:MGI:7828556] |
| 5 | lncRNA | 1.01E+08 | 1.01E+08 | ENSMUSG00000097772 | MGI:1918676 | 5430416N02Rik | RIKEN cDNA 5430416N02 gene [Source:MGI Symbol;Acc:MGI:1918676] |
| 5 | lncRNA | 1.01E+08 | 1.01E+08 | ENSMUSG00000131639 | MGI:7826588 | Gm71911 | predicted gene, 71911 [Source:MGI Symbol;Acc:MGI:7826588] |
| 5 | lncRNA | 1.01E+08 | 1.01E+08 | ENSMUSG00000131182 | MGI:7826424 | Gm71829 | predicted gene, 71829 [Source:MGI Symbol;Acc:MGI:7826424] |
| 5 | lncRNA | 1.01E+08 | 1.01E+08 | ENSMUSG00000131120 | MGI:7826404 | Gm71819 | predicted gene, 71819 [Source:MGI Symbol;Acc:MGI:7826404] |
| 5 | lncRNA | 1.01E+08 | 1.01E+08 | ENSMUSG00000131155 | MGI:7830460 | Gm73858 | predicted gene, 73858 [Source:MGI Symbol;Acc:MGI:7830460] |
| 5 | lncRNA | 1.01E+08 | 1.01E+08 | ENSMUSG00000085269 | MGI:3783219 | Gm15777 | predicted gene 15777 [Source:MGI Symbol;Acc:MGI:3783219] |
| 5 | lncRNA | 1.01E+08 | 1.01E+08 | ENSMUSG00000141768 | MGI:7829444 | Gm73343 | predicted gene, 73343 [Source:MGI Symbol;Acc:MGI:7829444] |
| 5 | lncRNA | 1.01E+08 | 1.01E+08 | ENSMUSG00000141699 | MGI:5595778 | Gm36619 | predicted gene, 36619 [Source:MGI Symbol;Acc:MGI:5595778] |
| 5 | lncRNA | 1.01E+08 | 1.01E+08 | ENSMUSG00000105755 | MGI:5663651 | Gm43514 | predicted gene 43514 [Source:MGI Symbol;Acc:MGI:5663651] |
| 5 | lncRNA | 1.01E+08 | 1.01E+08 | ENSMUSG00000087611 | MGI:1923052 | 4930458D05Rik | RIKEN cDNA 4930458D05 gene [Source:MGI Symbol;Acc:MGI:1923052] |
| 5 | lncRNA | 1.01E+08 | 1.01E+08 | ENSMUSG00000141832 | MGI:7829460 | Gm73351 | predicted gene, 73351 [Source:MGI Symbol;Acc:MGI:7829460] |
| 5 | lncRNA | 1.01E+08 | 1.01E+08 | ENSMUSG00000106255 | MGI:5595952 | Gm36793 | predicted gene, 36793 [Source:MGI Symbol;Acc:MGI:5595952] |
| 5 | lncRNA | 1.01E+08 | 1.01E+08 | ENSMUSG00000122123 | MGI:5595899 | Gm36740 | predicted gene, 36740 [Source:MGI Symbol;Acc:MGI:5595899] |
| 5 | lncRNA | 1.01E+08 | 1.01E+08 | ENSMUSG00000126729 | MGI:7825747 | Gm71487 | predicted gene, 71487 [Source:MGI Symbol;Acc:MGI:7825747] |
| 5 | lncRNA | 1.01E+08 | 1.01E+08 | ENSMUSG00000128974 | MGI:7825943 | Gm71585 | predicted gene, 71585 [Source:MGI Symbol;Acc:MGI:7825943] |

| | | | | | | | |
|---|---|---|---|---|---|---|---|
| 5 | lncRNA | 1.01E+08 | 1.01E+08 | ENSMUSG00000136829 | MGI:7828182 | Gm72710 | predicted gene, 72710 [Source:MGI Symbol;Acc:MGI:7828182] |
| 5 | lncRNA | 1.01E+08 | 1.01E+08 | ENSMUSG00000124318 | MGI:7825517 | Gm71371 | predicted gene, 71371 [Source:MGI Symbol;Acc:MGI:7825517] |
| 5 | lncRNA | 1.01E+08 | 1.01E+08 | ENSMUSG00000129567 | MGI:7825991 | Gm71610 | predicted gene, 71610 [Source:MGI Symbol;Acc:MGI:7825991] |
| 5 | lncRNA | 1.01E+08 | 1.01E+08 | ENSMUSG00000142499 | MGI:5596010 | Gm36851 | predicted gene, 36851 [Source:MGI Symbol;Acc:MGI:5596010] |
| 5 | lncRNA | 1.02E+08 | 1.02E+08 | ENSMUSG00000143141 | MGI:5625016 | Gm42131 | predicted gene, 42131 [Source:MGI Symbol;Acc:MGI:5625016] |
| 5 | lncRNA | 1.02E+08 | 1.02E+08 | ENSMUSG00000133793 | MGI:5589141 | Gm29982 | predicted gene, 29982 [Source:MGI Symbol;Acc:MGI:5589141] |
| 5 | lncRNA | 1.02E+08 | 1.02E+08 | ENSMUSG00000133836 | MGI:7827232 | Gm72233 | predicted gene, 72233 [Source:MGI Symbol;Acc:MGI:7827232] |
| 5 | lncRNA | 1.02E+08 | 1.02E+08 | ENSMUSG00000092503 | MGI:5141949 | Gm20484 | predicted gene 20484 [Source:MGI Symbol;Acc:MGI:5141949] |
| 5 | lncRNA | 1.02E+08 | 1.02E+08 | ENSMUSG00000139393 | MGI:7828862 | Gm73052 | predicted gene, 73052 [Source:MGI Symbol;Acc:MGI:7828862] |
| 5 | lncRNA | 1.02E+08 | 1.02E+08 | ENSMUSG00000105097 | MGI:5625018 | Gm42133 | predicted gene, 42133 [Source:MGI Symbol;Acc:MGI:5625018] |
| 5 | lncRNA | 1.02E+08 | 1.02E+08 | ENSMUSG00000106164 | MGI:1924611 | 9430085M18Rik | RIKEN cDNA 9430085M18 gene [Source:MGI Symbol;Acc:MGI:1924611] |
| 5 | lncRNA | 1.02E+08 | 1.02E+08 | ENSMUSG00000122648 | MGI:5625019 | Gm42134 | predicted gene, 42134 [Source:MGI Symbol;Acc:MGI:5625019] |
| 5 | lncRNA | 1.02E+08 | 1.02E+08 | ENSMUSG00000092354 | MGI:5142013 | Gm20548 | predicted gene 20548 [Source:MGI Symbol;Acc:MGI:5142013] |
| 5 | lncRNA | 1.02E+08 | 1.02E+08 | ENSMUSG00000131745 | MGI:7826618 | Gm71926 | predicted gene, 71926 [Source:MGI Symbol;Acc:MGI:7826618] |
| 5 | lncRNA | 1.02E+08 | 1.02E+08 | ENSMUSG00000123489 | MGI:7825409 | Gm71316 | predicted gene, 71316 [Source:MGI Symbol;Acc:MGI:7825409] |
| 5 | lncRNA | 1.02E+08 | 1.02E+08 | ENSMUSG00000132130 | MGI:7826722 | Gm71978 | predicted gene, 71978 [Source:MGI Symbol;Acc:MGI:7826722] |
| 5 | lncRNA | 1.02E+08 | 1.02E+08 | ENSMUSG00000128755 | MGI:5625020 | Gm42135 | predicted gene, 42135 [Source:MGI Symbol;Acc:MGI:5625020] |
| 5 | lncRNA | 1.02E+08 | 1.02E+08 | ENSMUSG00000106492 | MGI:5588866 | Gm29707 | predicted gene, 29707 [Source:MGI Symbol;Acc:MGI:5588866] |

| | | | | | | | |
|---|---|---|---|---|---|---|---|
| 5 | lncRNA | 1.02E+08 | 1.02E+08 | ENSMUSG00000126276 | MGI:7825705 | Gm71466 | predicted gene, 71466 [Source:MGI Symbol;Acc:MGI:7825705] |
| 5 | lncRNA | 1.02E+08 | 1.02E+08 | ENSMUSG00000138981 | MGI:5625021 | Gm42136 | predicted gene, 42136 [Source:MGI Symbol;Acc:MGI:5625021] |
| 5 | lncRNA | 1.02E+08 | 1.02E+08 | ENSMUSG00000133687 | MGI:5826584 | Gm46947 | predicted gene, 46947 [Source:MGI Symbol;Acc:MGI:5826584] |
| 5 | lncRNA | 1.02E+08 | 1.02E+08 | ENSMUSG00000139331 | MGI:7828844 | Gm73043 | predicted gene, 73043 [Source:MGI Symbol;Acc:MGI:7828844] |
| 5 | lncRNA | 1.02E+08 | 1.02E+08 | ENSMUSG00000137432 | MGI:7828396 | Gm72817 | predicted gene, 72817 [Source:MGI Symbol;Acc:MGI:7828396] |
| 5 | lncRNA | 1.02E+08 | 1.02E+08 | ENSMUSG00000132860 | MGI:7826870 | Gm72052 | predicted gene, 72052 [Source:MGI Symbol;Acc:MGI:7826870] |
| 5 | lncRNA | 1.03E+08 | 1.03E+08 | ENSMUSG00000123014 | MGI:7825346 | Gm71284 | predicted gene, 71284 [Source:MGI Symbol;Acc:MGI:7825346] |
| 5 | lncRNA | 1.03E+08 | 1.03E+08 | ENSMUSG00000123054 | MGI:7825350 | Gm71286 | predicted gene, 71286 [Source:MGI Symbol;Acc:MGI:7825350] |
| 5 | lncRNA | 1.03E+08 | 1.03E+08 | ENSMUSG00000106586 | MGI:1922755 | 1700013M08Rik | RIKEN cDNA 1700013M08 gene [Source:MGI Symbol;Acc:MGI:1922755] |
| 5 | lncRNA | 1.03E+08 | 1.03E+08 | ENSMUSG00000143028 | MGI:5589555 | Gm30396 | predicted gene, 30396 [Source:MGI Symbol;Acc:MGI:5589555] |
| 5 | lncRNA | 1.03E+08 | 1.03E+08 | ENSMUSG00000143109 | MGI:7829732 | Gm73488 | predicted gene, 73488 [Source:MGI Symbol;Acc:MGI:7829732] |
| 5 | lncRNA | 1.03E+08 | 1.03E+08 | ENSMUSG00000143146 | MGI:7829736 | Gm73490 | predicted gene, 73490 [Source:MGI Symbol;Acc:MGI:7829736] |
| 5 | lncRNA | 1.03E+08 | 1.03E+08 | ENSMUSG00000122328 | MGI:7825251 | Gm71235 | predicted gene, 71235 [Source:MGI Symbol;Acc:MGI:7825251] |
| 5 | lncRNA | 1.03E+08 | 1.04E+08 | ENSMUSG00000104956 | MGI:1921874 | 4930429D17Rik | RIKEN cDNA 4930429D17 gene [Source:MGI Symbol;Acc:MGI:1921874] |
| 5 | lncRNA | 1.03E+08 | 1.03E+08 | ENSMUSG00000105426 | MGI:1922769 | 1700021F02Rik | RIKEN cDNA 1700021F02 gene [Source:MGI Symbol;Acc:MGI:1922769] |
| 5 | lncRNA | 1.04E+08 | 1.04E+08 | ENSMUSG00000104684 | MGI:2441699 | 5430427N15Rik | RIKEN cDNA 5430427N15 gene [Source:MGI Symbol;Acc:MGI:2441699] |
| 5 | lncRNA | 1.04E+08 | 1.04E+08 | ENSMUSG00000140589 | MGI:5589602 | Gm30443 | predicted gene, 30443 [Source:MGI Symbol;Acc:MGI:5589602] |
| 5 | lncRNA | 1.04E+08 | 1.04E+08 | ENSMUSG00000141609 | MGI:7829404 | Gm73323 | predicted gene, 73323 [Source:MGI Symbol;Acc:MGI:7829404] |

| | | | | | | | |
|---|---|---|---|---|---|---|---|
| 5 | lncRNA | 1.04E+08 | 1.04E+08 | ENSMUSG00000104580 | MGI:5662757 | Gm42620 | predicted gene 42620 [Source:MGI Symbol;Acc:MGI:5662757] |
| 5 | lncRNA | 1.04E+08 | 1.04E+08 | ENSMUSG00000136783 | MGI:5625023 | Gm42138 | predicted gene, 42138 [Source:MGI Symbol;Acc:MGI:5625023] |
| 5 | lncRNA | 1.04E+08 | 1.04E+08 | ENSMUSG00000122606 | MGI:7825279 | Gm71249 | predicted gene, 71249 [Source:MGI Symbol;Acc:MGI:7825279] |
| 5 | lncRNA | 1.04E+08 | 1.04E+08 | ENSMUSG00000142430 | MGI:7829570 | Gm73406 | predicted gene, 73406 [Source:MGI Symbol;Acc:MGI:7829570] |
| 5 | lncRNA | 1.04E+08 | 1.04E+08 | ENSMUSG00000142388 | MGI:7829564 | Gm73403 | predicted gene, 73403 [Source:MGI Symbol;Acc:MGI:7829564] |
| 5 | lncRNA | 1.04E+08 | 1.04E+08 | ENSMUSG00000130284 | MGI:7826190 | Gm71711 | predicted gene, 71711 [Source:MGI Symbol;Acc:MGI:7826190] |
| 5 | lncRNA | 1.04E+08 | 1.04E+08 | ENSMUSG00000140976 | MGI:6367177 | Gm52783 | predicted gene, 52783 [Source:MGI Symbol;Acc:MGI:6367177] |
| 5 | lncRNA | 1.04E+08 | 1.04E+08 | ENSMUSG00000139941 | MGI:7828976 | Gm73109 | predicted gene, 73109 [Source:MGI Symbol;Acc:MGI:7828976] |
| 5 | lncRNA | 1.04E+08 | 1.04E+08 | ENSMUSG00000097877 | MGI:5477197 | Gm26703 | predicted gene, 26703 [Source:MGI Symbol;Acc:MGI:5477197] |
| 5 | lncRNA | 1.04E+08 | 1.04E+08 | ENSMUSG00000105615 | MGI:5590207 | Gm31048 | predicted gene, 31048 [Source:MGI Symbol;Acc:MGI:5590207] |
| 5 | lncRNA | 1.04E+08 | 1.04E+08 | ENSMUSG00000130229 | MGI:7826166 | Gm71699 | predicted gene, 71699 [Source:MGI Symbol;Acc:MGI:7826166] |
| 5 | lncRNA | 1.04E+08 | 1.05E+08 | ENSMUSG00000128790 | MGI:7825931 | Gm71579 | predicted gene, 71579 [Source:MGI Symbol;Acc:MGI:7825931] |
| 5 | lncRNA | 1.05E+08 | 1.05E+08 | ENSMUSG00000130295 | MGI:7826192 | Gm71712 | predicted gene, 71712 [Source:MGI Symbol;Acc:MGI:7826192] |
| 5 | lncRNA | 1.05E+08 | 1.05E+08 | ENSMUSG00000124296 | MGI:7825511 | Gm71368 | predicted gene, 71368 [Source:MGI Symbol;Acc:MGI:7825511] |
| 5 | lncRNA | 1.05E+08 | 1.05E+08 | ENSMUSG00000124334 | MGI:7825521 | Gm71373 | predicted gene, 71373 [Source:MGI Symbol;Acc:MGI:7825521] |
| 5 | lncRNA | 1.05E+08 | 1.05E+08 | ENSMUSG00000122026 | MGI:7825199 | Gm71209 | predicted gene, 71209 [Source:MGI Symbol;Acc:MGI:7825199] |
| 5 | lncRNA | 1.05E+08 | 1.05E+08 | ENSMUSG00000140668 | MGI:7829220 | Gm73231 | predicted gene, 73231 [Source:MGI Symbol;Acc:MGI:7829220] |
| 5 | lncRNA | 1.05E+08 | 1.05E+08 | ENSMUSG00000134138 | MGI:5590414 | Gm31255 | predicted gene, 31255 [Source:MGI Symbol;Acc:MGI:5590414] |

| | | | | | | | |
|---|---|---|---|---|---|---|---|
| 5 | lncRNA | 1.05E+08 | 1.05E+08 | ENSMUSG00000134259 | MGI:7827428 | Gm72331 | predicted gene, 72331 [Source:MGI Symbol;Acc:MGI:7827428] |
| 5 | lncRNA | 1.05E+08 | 1.05E+08 | ENSMUSG00000139462 | MGI:7828906 | Gm73074 | predicted gene, 73074 [Source:MGI Symbol;Acc:MGI:7828906] |
| 5 | lncRNA | 1.05E+08 | 1.05E+08 | ENSMUSG00000134102 | MGI:7827352 | Gm72293 | predicted gene, 72293 [Source:MGI Symbol;Acc:MGI:7827352] |
| 5 | lncRNA | 1.05E+08 | 1.05E+08 | ENSMUSG00000132192 | MGI:7826734 | Gm71984 | predicted gene, 71984 [Source:MGI Symbol;Acc:MGI:7826734] |
| 5 | lncRNA | 1.05E+08 | 1.05E+08 | ENSMUSG00000119978 | MGI:5590944 | Gm31785 | predicted gene, 31785 [Source:MGI Symbol;Acc:MGI:5590944] |
| 5 | lncRNA | 1.06E+08 | 1.06E+08 | ENSMUSG00000134940 | MGI:7827556 | Gm72395 | predicted gene, 72395 [Source:MGI Symbol;Acc:MGI:7827556] |
| 5 | lncRNA | 1.06E+08 | 1.06E+08 | ENSMUSG00000106438 | MGI:5591210 | Gm32051 | predicted gene, 32051 [Source:MGI Symbol;Acc:MGI:5591210] |
| 5 | lncRNA | 1.06E+08 | 1.06E+08 | ENSMUSG00000135055 | MGI:7827596 | Gm72415 | predicted gene, 72415 [Source:MGI Symbol;Acc:MGI:7827596] |
| 5 | lncRNA | 1.06E+08 | 1.06E+08 | ENSMUSG00000135020 | MGI:1923077 | 4930542N06Rik | RIKEN cDNA 4930542N06 gene [Source:MGI Symbol;Acc:MGI:1923077] |
| 5 | lncRNA | 1.06E+08 | 1.06E+08 | ENSMUSG00000126067 | | | novel transcript |
| 5 | lncRNA | 1.06E+08 | 1.06E+08 | ENSMUSG00000126110 | MGI:7825683 | Gm71455 | predicted gene, 71455 [Source:MGI Symbol;Acc:MGI:7825683] |
| 5 | lncRNA | 1.06E+08 | 1.06E+08 | ENSMUSG00000133270 | MGI:7827092 | Gm72163 | predicted gene, 72163 [Source:MGI Symbol;Acc:MGI:7827092] |
| 5 | lncRNA | 1.06E+08 | 1.06E+08 | ENSMUSG00000133188 | MGI:7827012 | Gm72123 | predicted gene, 72123 [Source:MGI Symbol;Acc:MGI:7827012] |
| 5 | lncRNA | 1.06E+08 | 1.06E+08 | ENSMUSG00000133309 | MGI:7827098 | Gm72166 | predicted gene, 72166 [Source:MGI Symbol;Acc:MGI:7827098] |
| 5 | lncRNA | 1.06E+08 | 1.06E+08 | ENSMUSG00000133225 | MGI:7827076 | Gm72155 | predicted gene, 72155 [Source:MGI Symbol;Acc:MGI:7827076] |
| 5 | lncRNA | 1.06E+08 | 1.06E+08 | ENSMUSG00000085771 | MGI:2141339 | C230066G23Rik | RIKEN cDNA C230066G23 gene [Source:MGI Symbol;Acc:MGI:2141339] |
| 5 | lncRNA | 1.06E+08 | 1.06E+08 | ENSMUSG00000143828 | MGI:7831190 | Gm74226 | predicted gene, 74226 [Source:MGI Symbol;Acc:MGI:7831190] |
| 5 | lncRNA | 1.06E+08 | 1.06E+08 | ENSMUSG00000143792 | MGI:7829970 | Gm73607 | predicted gene, 73607 [Source:MGI Symbol;Acc:MGI:7829970] |

| | | | | | | | |
|---|---|---|---|---|---|---|---|
| 5 | lncRNA | 1.06E+08 | 1.06E+08 | ENSMUSG00000131937 | MGI:5591389 | Gm32230 | predicted gene, 32230 [Source:MGI Symbol;Acc:MGI:5591389] |
| 5 | lncRNA | 1.06E+08 | 1.06E+08 | ENSMUSG00000120245 | MGI:7338425 | Gm57256 | predicted gene, 57256 [Source:MGI Symbol;Acc:MGI:7338425] |
| 5 | lncRNA | 1.06E+08 | 1.06E+08 | ENSMUSG00000126152 | MGI:7831156 | Gm74209 | predicted gene, 74209 [Source:MGI Symbol;Acc:MGI:7831156] |
| 5 | lncRNA | 1.06E+08 | 1.06E+08 | ENSMUSG00000121168 | MGI:7338559 | Gm57323 | predicted gene, 57323 [Source:MGI Symbol;Acc:MGI:7338559] |
| 5 | lncRNA | 1.06E+08 | 1.06E+08 | ENSMUSG00000126262 | MGI:7825699 | Gm71463 | predicted gene, 71463 [Source:MGI Symbol;Acc:MGI:7825699] |
| 5 | lncRNA | 1.06E+08 | 1.06E+08 | ENSMUSG00000084878 | MGI:3802116 | Lrrc8dos | leucine rich repeat containing 8D, opposite strand [Source:MGI Symbol;Acc:MGI:3802116] |
| 5 | lncRNA | 1.06E+08 | 1.06E+08 | ENSMUSG00000126228 | MGI:7825691 | Gm71459 | predicted gene, 71459 [Source:MGI Symbol;Acc:MGI:7825691] |
| 5 | lncRNA | 1.06E+08 | 1.06E+08 | ENSMUSG00000123641 | MGI:7825425 | Gm71324 | predicted gene, 71324 [Source:MGI Symbol;Acc:MGI:7825425] |
| 5 | lncRNA | 1.06E+08 | 1.06E+08 | ENSMUSG00000123599 | MGI:7825421 | Gm71322 | predicted gene, 71322 [Source:MGI Symbol;Acc:MGI:7825421] |
| 5 | lncRNA | 1.06E+08 | 1.06E+08 | ENSMUSG00000123655 | MGI:7825427 | Gm71325 | predicted gene, 71325 [Source:MGI Symbol;Acc:MGI:7825427] |
| 5 | lncRNA | 1.06E+08 | 1.06E+08 | ENSMUSG00000106483 | MGI:5591895 | Gm32736 | predicted gene, 32736 [Source:MGI Symbol;Acc:MGI:5591895] |
| 5 | lncRNA | 1.06E+08 | 1.06E+08 | ENSMUSG00000126268 | MGI:7825703 | Gm71465 | predicted gene, 71465 [Source:MGI Symbol;Acc:MGI:7825703] |
| 5 | lncRNA | 1.06E+08 | 1.07E+08 | ENSMUSG00000106594 | MGI:1921031 | 4930432H08Rik | RIKEN cDNA 4930432H08 gene [Source:MGI Symbol;Acc:MGI:1921031] |
| 5 | lncRNA | 1.06E+08 | 1.06E+08 | ENSMUSG00000135775 | MGI:7827772 | Gm72504 | predicted gene, 72504 [Source:MGI Symbol;Acc:MGI:7827772] |
| 5 | lncRNA | 1.06E+08 | 1.06E+08 | ENSMUSG00000105508 | MGI:5592080 | Gm32921 | predicted gene, 32921 [Source:MGI Symbol;Acc:MGI:5592080] |
| 5 | lncRNA | 1.06E+08 | 1.07E+08 | ENSMUSG00000105531 | MGI:2685681 | 4930458A03Rik | RIKEN cDNA 4930458A03 gene [Source:MGI Symbol;Acc:MGI:2685681] |
| 5 | lncRNA | 1.07E+08 | 1.07E+08 | ENSMUSG00000128962 | MGI:7825941 | Gm71584 | predicted gene, 71584 [Source:MGI Symbol;Acc:MGI:7825941] |
| 5 | lncRNA | 1.07E+08 | 1.07E+08 | ENSMUSG00000129002 | MGI:7825947 | Gm71587 | predicted gene, 71587 [Source:MGI Symbol;Acc:MGI:7825947] |

| | | | | | | | |
|---|---|---|---|---|---|---|---|
| 5 | lncRNA | 1.07E+08 | 1.07E+08 | ENSMUSG00000120527 | MGI:5592310 | Gm33151 | predicted gene, 33151 [Source:MGI Symbol;Acc:MGI:5592310] |
| 5 | lncRNA | 1.07E+08 | 1.07E+08 | ENSMUSG00000143832 | MGI:7830010 | Gm73627 | predicted gene, 73627 [Source:MGI Symbol;Acc:MGI:7830010] |
| 5 | lncRNA | 1.07E+08 | 1.07E+08 | ENSMUSG00000097018 | MGI:5477366 | Gm26872 | predicted gene, 26872 [Source:MGI Symbol;Acc:MGI:5477366] |
| 5 | lncRNA | 1.07E+08 | 1.07E+08 | ENSMUSG00000130270 | MGI:7826176 | Gm71704 | predicted gene, 71704 [Source:MGI Symbol;Acc:MGI:7826176] |
| 5 | lncRNA | 1.07E+08 | 1.07E+08 | ENSMUSG00000099061 | MGI:5547786 | Gm28050 | predicted gene, 28050 [Source:MGI Symbol;Acc:MGI:5547786] |
| 5 | lncRNA | 1.07E+08 | 1.07E+08 | ENSMUSG00000131939 | | | novel transcript |
| 5 | lncRNA | 1.07E+08 | 1.07E+08 | ENSMUSG00000098302 | MGI:5547775 | Gm28039 | predicted gene, 28039 [Source:MGI Symbol;Acc:MGI:5547775] |
| 5 | lncRNA | 1.07E+08 | 1.07E+08 | ENSMUSG00000101237 | MGI:5580170 | Gm29464 | predicted gene 29464 [Source:MGI Symbol;Acc:MGI:5580170] |
| 5 | lncRNA | 1.07E+08 | 1.07E+08 | ENSMUSG00000131617 | MGI:7826584 | Gm71909 | predicted gene, 71909 [Source:MGI Symbol;Acc:MGI:7826584] |
| 5 | lncRNA | 1.07E+08 | 1.07E+08 | ENSMUSG00000126214 | MGI:5592486 | Gm33327 | predicted gene, 33327 [Source:MGI Symbol;Acc:MGI:5592486] |
| 5 | lncRNA | 1.07E+08 | 1.07E+08 | ENSMUSG00000140743 | MGI:7829226 | Gm73234 | predicted gene, 73234 [Source:MGI Symbol;Acc:MGI:7829226] |
| 5 | lncRNA | 1.07E+08 | 1.07E+08 | ENSMUSG00000136976 | MGI:7828246 | Gm72742 | predicted gene, 72742 [Source:MGI Symbol;Acc:MGI:7828246] |
| 5 | lncRNA | 1.07E+08 | 1.07E+08 | ENSMUSG00000124898 | MGI:7825571 | Gm71399 | predicted gene, 71399 [Source:MGI Symbol;Acc:MGI:7825571] |
| 5 | lncRNA | 1.07E+08 | 1.07E+08 | ENSMUSG00000134103 | MGI:5592530 | Gm33371 | predicted gene, 33371 [Source:MGI Symbol;Acc:MGI:5592530] |
| 5 | lncRNA | 1.07E+08 | 1.07E+08 | ENSMUSG00000106554 | MGI:5592633 | Gm33474 | predicted gene, 33474 [Source:MGI Symbol;Acc:MGI:5592633] |
| 5 | lncRNA | 1.07E+08 | 1.07E+08 | ENSMUSG00000123782 | MGI:5625031 | Gm42146 | predicted gene, 42146 [Source:MGI Symbol;Acc:MGI:5625031] |
| 5 | lncRNA | 1.07E+08 | 1.07E+08 | ENSMUSG00000138859 | MGI:7828740 | Gm72990 | predicted gene, 72990 [Source:MGI Symbol;Acc:MGI:7828740] |
| 5 | lncRNA | 1.07E+08 | 1.07E+08 | ENSMUSG00000132945 | MGI:7826894 | Gm72064 | predicted gene, 72064 [Source:MGI Symbol;Acc:MGI:7826894] |

| | | | | | | | |
|---|---|---|---|---|---|---|---|
| 5 | lncRNA | 1.08E+08 | 1.08E+08 | ENSMUSG00000125872 | MGI:7825661 | Gm71444 | predicted gene, 71444 [Source:MGI Symbol;Acc:MGI:7825661] |
| 5 | lncRNA | 1.08E+08 | 1.08E+08 | ENSMUSG00000090558 | MGI:4938029 | Gm17202 | predicted gene 17202 [Source:MGI Symbol;Acc:MGI:4938029] |
| 5 | lncRNA | 1.08E+08 | 1.08E+08 | ENSMUSG00000125909 | MGI:7825665 | Gm71446 | predicted gene, 71446 [Source:MGI Symbol;Acc:MGI:7825665] |
| 5 | lncRNA | 1.08E+08 | 1.08E+08 | ENSMUSG00000132608 | MGI:7826808 | Gm72021 | predicted gene, 72021 [Source:MGI Symbol;Acc:MGI:7826808] |
| 5 | lncRNA | 1.08E+08 | 1.08E+08 | ENSMUSG00000134553 | MGI:5593044 | Gm33885 | predicted gene, 33885 [Source:MGI Symbol;Acc:MGI:5593044] |
| 5 | lncRNA | 1.08E+08 | 1.08E+08 | ENSMUSG00000085900 | MGI:1925693 | A930041C12Rik | RIKEN cDNA A930041C12 gene [Source:MGI Symbol;Acc:MGI:1925693] |
| 5 | lncRNA | 1.08E+08 | 1.08E+08 | ENSMUSG00000133374 | MGI:7827138 | Gm72186 | predicted gene, 72186 [Source:MGI Symbol;Acc:MGI:7827138] |
| 5 | lncRNA | 1.08E+08 | 1.08E+08 | ENSMUSG00000106333 | MGI:1923034 | 4930428O21Rik | RIKEN cDNA 4930428O21 gene [Source:MGI Symbol;Acc:MGI:1923034] |
| 5 | lncRNA | 1.08E+08 | 1.08E+08 | ENSMUSG00000090030 | MGI:3041214 | A430072P03Rik | RIKEN cDNA A430072P03 gene [Source:MGI Symbol;Acc:MGI:3041214] |
| 5 | lncRNA | 1.08E+08 | 1.08E+08 | ENSMUSG00000123337 | MGI:7825391 | Gm71307 | predicted gene, 71307 [Source:MGI Symbol;Acc:MGI:7825391] |
| 5 | lncRNA | 1.08E+08 | 1.08E+08 | ENSMUSG00000137433 | MGI:7828406 | Gm72822 | predicted gene, 72822 [Source:MGI Symbol;Acc:MGI:7828406] |
| 5 | lncRNA | 1.08E+08 | 1.08E+08 | ENSMUSG00000137507 | MGI:7828436 | Gm72837 | predicted gene, 72837 [Source:MGI Symbol;Acc:MGI:7828436] |
| 5 | lncRNA | 1.08E+08 | 1.08E+08 | ENSMUSG00000137388 | MGI:7830274 | Gm73762 | predicted gene, 73762 [Source:MGI Symbol;Acc:MGI:7830274] |
| 5 | lncRNA | 1.08E+08 | 1.08E+08 | ENSMUSG00000141797 | MGI:7829452 | Gm73347 | predicted gene, 73347 [Source:MGI Symbol;Acc:MGI:7829452] |
| 5 | lncRNA | 1.08E+08 | 1.08E+08 | ENSMUSG00000123424 | MGI:7830606 | Gm73931 | predicted gene, 73931 [Source:MGI Symbol;Acc:MGI:7830606] |
| 5 | lncRNA | 1.09E+08 | 1.09E+08 | ENSMUSG00000104904 | MGI:1924810 | 9330198I05Rik | RIKEN cDNA 9330198I05 gene [Source:MGI Symbol;Acc:MGI:1924810] |
| 5 | lncRNA | 1.09E+08 | 1.09E+08 | ENSMUSG00000106304 | MGI:5662655 | Gm42518 | predicted gene 42518 [Source:MGI Symbol;Acc:MGI:5662655] |
| 5 | lncRNA | 1.09E+08 | 1.09E+08 | ENSMUSG00000072769 | MGI:3642823 | Gm10419 | predicted gene 10419 [Source:MGI Symbol;Acc:MGI:3642823] |

| | | | | | | | |
|---|---|---|---|---|---|---|---|
| 5 | lncRNA | 1.09E+08 | 1.09E+08 | ENSMUSG00000120483 | MGI:7338441 | Gm57264 | predicted gene, 57264 [Source:MGI Symbol;Acc:MGI:7338441] |
| 5 | lncRNA | 1.09E+08 | 1.09E+08 | ENSMUSG00000121853 | MGI:7830664 | Gm73960 | predicted gene, 73960 [Source:MGI Symbol;Acc:MGI:7830664] |
| 5 | lncRNA | 1.09E+08 | 1.09E+08 | ENSMUSG00000128538 | MGI:7830734 | Gm73995 | predicted gene, 73995 [Source:MGI Symbol;Acc:MGI:7830734] |
| 5 | lncRNA | 1.09E+08 | 1.09E+08 | ENSMUSG00000142352 | MGI:7830736 | Gm73996 | predicted gene, 73996 [Source:MGI Symbol;Acc:MGI:7830736] |
| 5 | lncRNA | 1.09E+08 | 1.09E+08 | ENSMUSG00000133269 | MGI:7830740 | Gm73998 | predicted gene, 73998 [Source:MGI Symbol;Acc:MGI:7830740] |
| 5 | lncRNA | 1.09E+08 | 1.09E+08 | ENSMUSG00000133224 | MGI:7827074 | Gm72154 | predicted gene, 72154 [Source:MGI Symbol;Acc:MGI:7827074] |
| 5 | lncRNA | 1.09E+08 | 1.09E+08 | ENSMUSG00000138567 | MGI:5625036 | Gm42151 | predicted gene, 42151 [Source:MGI Symbol;Acc:MGI:5625036] |
| 5 | lncRNA | 1.09E+08 | 1.09E+08 | ENSMUSG00000101762 | MGI:1920545 | 1700047L14Rik | RIKEN cDNA 1700047L14 gene [Source:MGI Symbol;Acc:MGI:1920545] |
| 5 | lncRNA | 1.09E+08 | 1.09E+08 | ENSMUSG00000093479 | MGI:5313076 | Gm20629 | predicted gene 20629 [Source:MGI Symbol;Acc:MGI:5313076] |
| 5 | lncRNA | 1.09E+08 | 1.09E+08 | ENSMUSG00000129648 | MGI:7826009 | Gm71619 | predicted gene, 71619 [Source:MGI Symbol;Acc:MGI:7826009] |
| 5 | lncRNA | 1.10E+08 | 1.10E+08 | ENSMUSG00000124061 | MGI:6364910 | Gm50598 | predicted gene, 50598 [Source:MGI Symbol;Acc:MGI:6364910] |
| 5 | lncRNA | 1.10E+08 | 1.10E+08 | ENSMUSG00000134481 | MGI:7827446 | Gm72340 | predicted gene, 72340 [Source:MGI Symbol;Acc:MGI:7827446] |
| 5 | lncRNA | 1.10E+08 | 1.10E+08 | ENSMUSG00000136256 | MGI:7827908 | Gm72573 | predicted gene, 72573 [Source:MGI Symbol;Acc:MGI:7827908] |
| 5 | lncRNA | 1.10E+08 | 1.10E+08 | ENSMUSG00000136293 | MGI:7830534 | Gm73895 | predicted gene, 73895 [Source:MGI Symbol;Acc:MGI:7830534] |
| 5 | lncRNA | 1.10E+08 | 1.10E+08 | ENSMUSG00000121595 | | | RIKEN cDNA 4930522L14 gene [Source:NCBI gene (formerly Entrezgene);Acc:100041734] |
| 5 | lncRNA | 1.10E+08 | 1.10E+08 | ENSMUSG00000120227 | MGI:7338503 | Gm57295 | predicted gene, 57295 [Source:MGI Symbol;Acc:MGI:7338503] |
| 5 | lncRNA | 1.10E+08 | 1.10E+08 | ENSMUSG00000133173 | MGI:5594248 | Gm35089 | predicted gene, 35089 [Source:MGI Symbol;Acc:MGI:5594248] |
| 5 | lncRNA | 1.10E+08 | 1.10E+08 | ENSMUSG00000097140 | MGI:5477273 | Gm26779 | predicted gene, 26779 [Source:MGI Symbol;Acc:MGI:5477273] |

| | | | | | | | |
|---|---|---|---|---|---|---|---|
| 5 | lncRNA | 1.10E+08 | 1.10E+08 | ENSMUSG00000133215 | MGI:5477302 | Gm26808 | predicted gene, 26808 [Source:MGI Symbol;Acc:MGI:5477302] |
| 5 | lncRNA | 1.10E+08 | 1.10E+08 | ENSMUSG00000097863 | MGI:1922653 | 1010001B22Rik | RIKEN cDNA 1010001B22 gene [Source:MGI Symbol;Acc:MGI:1922653] |
| 5 | lncRNA | 1.10E+08 | 1.10E+08 | ENSMUSG00000123077 | MGI:6723685 | Gm54334 | predicted gene, 54334 [Source:MGI Symbol;Acc:MGI:6723685] |
| 5 | lncRNA | 1.10E+08 | 1.10E+08 | ENSMUSG00000086247 | MGI:3783229 | Gm15787 | predicted gene 15787 [Source:MGI Symbol;Acc:MGI:3783229] |
| 5 | lncRNA | 1.10E+08 | 1.10E+08 | ENSMUSG00000104720 | MGI:5663275 | Gm43138 | predicted gene 43138 [Source:MGI Symbol;Acc:MGI:5663275] |
| 5 | lncRNA | 1.10E+08 | 1.10E+08 | ENSMUSG00000104970 | MGI:3704253 | 5930411N13Rik | RIKEN cDNA 5930411N13 gene [Source:MGI Symbol;Acc:MGI:3704253] |
| 5 | lncRNA | 1.10E+08 | 1.10E+08 | ENSMUSG00000127072 | MGI:7825785 | Gm71506 | predicted gene, 71506 [Source:MGI Symbol;Acc:MGI:7825785] |
| 5 | lncRNA | 1.10E+08 | 1.10E+08 | ENSMUSG00000087439 | MGI:3783230 | Gm15788 | predicted gene 15788 [Source:MGI Symbol;Acc:MGI:3783230] |
| 5 | lncRNA | 1.10E+08 | 1.10E+08 | ENSMUSG00000085606 | MGI:3783234 | Gm15792 | predicted gene 15792 [Source:MGI Symbol;Acc:MGI:3783234] |
| 5 | lncRNA | 1.10E+08 | 1.10E+08 | ENSMUSG00000132550 | MGI:7826798 | Gm72016 | predicted gene, 72016 [Source:MGI Symbol;Acc:MGI:7826798] |
| 5 | lncRNA | 1.11E+08 | 1.11E+08 | ENSMUSG00000105482 | MGI:5662733 | Gm42596 | predicted gene 42596 [Source:MGI Symbol;Acc:MGI:5662733] |
| 5 | lncRNA | 1.11E+08 | 1.11E+08 | ENSMUSG00000048215 | MGI:2445162 | A630023P12Rik | RIKEN cDNA A630023P12 gene [Source:MGI Symbol;Acc:MGI:2445162] |
| 5 | lncRNA | 1.11E+08 | 1.11E+08 | ENSMUSG00000138464 | MGI:7828604 | Gm72922 | predicted gene, 72922 [Source:MGI Symbol;Acc:MGI:7828604] |
| 5 | lncRNA | 1.11E+08 | 1.11E+08 | ENSMUSG00000134164 | MGI:7827390 | Gm72312 | predicted gene, 72312 [Source:MGI Symbol;Acc:MGI:7827390] |
| 5 | lncRNA | 1.11E+08 | 1.11E+08 | ENSMUSG00000105161 | MGI:5662732 | Gm42595 | predicted gene 42595 [Source:MGI Symbol;Acc:MGI:5662732] |
| 5 | lncRNA | 1.11E+08 | 1.11E+08 | ENSMUSG00000086401 | MGI:3783008 | Gm15559 | predicted gene 15559 [Source:MGI Symbol;Acc:MGI:3783008] |
| 5 | lncRNA | 1.11E+08 | 1.11E+08 | ENSMUSG00000128201 | MGI:5477009 | Gm26515 | predicted gene, 26515 [Source:MGI Symbol;Acc:MGI:5477009] |
| 5 | lncRNA | 1.11E+08 | 1.11E+08 | ENSMUSG00000132579 | MGI:7826804 | Gm72019 | predicted gene, 72019 [Source:MGI Symbol;Acc:MGI:7826804] |

| | | | | | | | |
|---|---|---|---|---|---|---|---|
| 5 | lncRNA | 1.11E+08 | 1.11E+08 | ENSMUSG00000131984 | MGI:7826684 | Gm71959 | predicted gene, 71959 [Source:MGI Symbol;Acc:MGI:7826684] |
| 5 | lncRNA | 1.11E+08 | 1.11E+08 | ENSMUSG00000133651 | MGI:7827166 | Gm72200 | predicted gene, 72200 [Source:MGI Symbol;Acc:MGI:7827166] |
| 5 | lncRNA | 1.11E+08 | 1.11E+08 | ENSMUSG00000140868 | MGI:5594787 | Gm35628 | predicted gene, 35628 [Source:MGI Symbol;Acc:MGI:5594787] |
| 5 | lncRNA | 1.11E+08 | 1.11E+08 | ENSMUSG00000141983 | MGI:7829474 | Gm73358 | predicted gene, 73358 [Source:MGI Symbol;Acc:MGI:7829474] |
| 5 | lncRNA | 1.12E+08 | 1.12E+08 | ENSMUSG00000104559 | MGI:5663255 | Gm43118 | predicted gene 43118 [Source:MGI Symbol;Acc:MGI:5663255] |
| 5 | lncRNA | 1.12E+08 | 1.12E+08 | ENSMUSG00000120465 | MGI:7338437 | Gm57262 | predicted gene, 57262 [Source:MGI Symbol;Acc:MGI:7338437] |
| 5 | lncRNA | 1.12E+08 | 1.12E+08 | ENSMUSG00000132416 | MGI:7826740 | Gm71987 | predicted gene, 71987 [Source:MGI Symbol;Acc:MGI:7826740] |
| 5 | lncRNA | 1.12E+08 | 1.12E+08 | ENSMUSG00000141147 | MGI:5625040 | Gm42155 | predicted gene, 42155 [Source:MGI Symbol;Acc:MGI:5625040] |
| 5 | lncRNA | 1.12E+08 | 1.12E+08 | ENSMUSG00000123711 | MGI:5625039 | Gm42154 | predicted gene, 42154 [Source:MGI Symbol;Acc:MGI:5625039] |
| 5 | lncRNA | 1.12E+08 | 1.12E+08 | ENSMUSG00000144040 | MGI:7830108 | Gm73679 | predicted gene, 73679 [Source:MGI Symbol;Acc:MGI:7830108] |
| 5 | lncRNA | 1.12E+08 | 1.12E+08 | ENSMUSG00000120047 | MGI:5595308 | Cirpil | cardiac ischemia reperfusion associated p53 interacting lncRNA [Source:MGI Symbol;Acc:MGI:5595308] |
| 5 | lncRNA | 1.12E+08 | 1.12E+08 | ENSMUSG00000134002 | MGI:5595382 | Gm36223 | predicted gene, 36223 [Source:MGI Symbol;Acc:MGI:5595382] |
| 5 | lncRNA | 1.12E+08 | 1.12E+08 | ENSMUSG00000141595 | MGI:7829402 | Gm73322 | predicted gene, 73322 [Source:MGI Symbol;Acc:MGI:7829402] |
| 5 | lncRNA | 1.12E+08 | 1.12E+08 | ENSMUSG00000141634 | MGI:7829414 | Gm73328 | predicted gene, 73328 [Source:MGI Symbol;Acc:MGI:7829414] |
| 5 | lncRNA | 1.12E+08 | 1.12E+08 | ENSMUSG00000134952 | MGI:5595433 | Gm36274 | predicted gene, 36274 [Source:MGI Symbol;Acc:MGI:5595433] |
| 5 | lncRNA | 1.12E+08 | 1.12E+08 | ENSMUSG00000052848 | MGI:3041189 | C130026L21Rik | RIKEN cDNA C130026L21 gene [Source:MGI Symbol;Acc:MGI:3041189] |
| 5 | lncRNA | 1.12E+08 | 1.12E+08 | ENSMUSG00000104833 | MGI:5662626 | Gm42489 | predicted gene 42489 [Source:MGI Symbol;Acc:MGI:5662626] |

| | | | | | | | |
|---|---|---|---|---|---|---|---|
| 5 | lncRNA | 1.12E+08 | 1.12E+08 | ENSMUSG00000125147 | MGI:7825585 | Gm71406 | predicted gene, 71406 [Source:MGI Symbol;Acc:MGI:7825585] |
| 5 | lncRNA | 1.12E+08 | 1.12E+08 | ENSMUSG00000122221 | MGI:5595545 | Gm36386 | predicted gene, 36386 [Source:MGI Symbol;Acc:MGI:5595545] |
| 5 | lncRNA | 1.12E+08 | 1.12E+08 | ENSMUSG00000122261 | MGI:5595593 | Gm36434 | predicted gene, 36434 [Source:MGI Symbol;Acc:MGI:5595593] |
| 5 | lncRNA | 1.12E+08 | 1.12E+08 | ENSMUSG00000122182 | MGI:7825227 | Gm71223 | predicted gene, 71223 [Source:MGI Symbol;Acc:MGI:7825227] |
| 5 | lncRNA | 1.12E+08 | 1.12E+08 | ENSMUSG00000123321 | MGI:7825387 | Gm71305 | predicted gene, 71305 [Source:MGI Symbol;Acc:MGI:7825387] |
| 5 | lncRNA | 1.12E+08 | 1.12E+08 | ENSMUSG00000123283 | MGI:7825383 | Gm71303 | predicted gene, 71303 [Source:MGI Symbol;Acc:MGI:7825383] |
| 5 | lncRNA | 1.12E+08 | 1.12E+08 | ENSMUSG00000123154 | MGI:7825360 | Gm71291 | predicted gene, 71291 [Source:MGI Symbol;Acc:MGI:7825360] |
| 5 | lncRNA | 1.12E+08 | 1.12E+08 | ENSMUSG00000142550 | MGI:7829624 | Gm73434 | predicted gene, 73434 [Source:MGI Symbol;Acc:MGI:7829624] |
| 5 | lncRNA | 1.12E+08 | 1.12E+08 | ENSMUSG00000128486 | MGI:5595640 | Gm36481 | predicted gene, 36481 [Source:MGI Symbol;Acc:MGI:5595640] |
| 5 | lncRNA | 1.12E+08 | 1.12E+08 | ENSMUSG00000085919 | MGI:3801872 | Gm16019 | predicted gene 16019 [Source:MGI Symbol;Acc:MGI:3801872] |
| 5 | lncRNA | 1.12E+08 | 1.12E+08 | ENSMUSG00000085271 | MGI:2685527 | E130006D01Rik | RIKEN cDNA E130006D01 gene [Source:MGI Symbol;Acc:MGI:2685527] |
| 5 | lncRNA | 1.12E+08 | 1.12E+08 | ENSMUSG00000105376 | MGI:5595694 | Gm36535 | predicted gene, 36535 [Source:MGI Symbol;Acc:MGI:5595694] |
| 5 | lncRNA | 1.12E+08 | 1.12E+08 | ENSMUSG00000131805 | MGI:7826634 | Gm71934 | predicted gene, 71934 [Source:MGI Symbol;Acc:MGI:7826634] |
| 5 | lncRNA | 1.12E+08 | 1.12E+08 | ENSMUSG00000141916 | MGI:7829468 | Gm73355 | predicted gene, 73355 [Source:MGI Symbol;Acc:MGI:7829468] |
| 5 | lncRNA | 1.12E+08 | 1.12E+08 | ENSMUSG00000128353 | MGI:5589094 | Gm29935 | predicted gene, 29935 [Source:MGI Symbol;Acc:MGI:5589094] |
| 5 | lncRNA | 1.12E+08 | 1.12E+08 | ENSMUSG00000106060 | MGI:5662625 | Gm42488 | predicted gene 42488 [Source:MGI Symbol;Acc:MGI:5662625] |
| 5 | lncRNA | 1.12E+08 | 1.12E+08 | ENSMUSG00000128427 | MGI:7825889 | Gm71558 | predicted gene, 71558 [Source:MGI Symbol;Acc:MGI:7825889] |
| 5 | lncRNA | 1.12E+08 | 1.12E+08 | ENSMUSG00000128317 | MGI:5596039 | Gm36880 | predicted gene, 36880 [Source:MGI Symbol;Acc:MGI:5596039] |

| | | | | | | | |
|---|---|---|---|---|---|---|---|
| 5 | lncRNA | 1.12E+08 | 1.12E+08 | ENSMUSG00000128392 | MGI:5625041 | Gm42156 | predicted gene, 42156 [Source:MGI Symbol;Acc:MGI:5625041] |
| 5 | lncRNA | 1.12E+08 | 1.12E+08 | ENSMUSG00000128275 | MGI:7825869 | Gm71548 | predicted gene, 71548 [Source:MGI Symbol;Acc:MGI:7825869] |
| 5 | lncRNA | 1.12E+08 | 1.12E+08 | ENSMUSG00000128237 | MGI:7825865 | Gm71546 | predicted gene, 71546 [Source:MGI Symbol;Acc:MGI:7825865] |
| 5 | lncRNA | 1.12E+08 | 1.12E+08 | ENSMUSG00000131045 | MGI:5589191 | Gm30032 | predicted gene, 30032 [Source:MGI Symbol;Acc:MGI:5589191] |
| 5 | lncRNA | 1.12E+08 | 1.12E+08 | ENSMUSG00000131083 | MGI:7826374 | Gm71804 | predicted gene, 71804 [Source:MGI Symbol;Acc:MGI:7826374] |
| 5 | lncRNA | 1.12E+08 | 1.12E+08 | ENSMUSG00000139603 | MGI:7828942 | Gm73092 | predicted gene, 73092 [Source:MGI Symbol;Acc:MGI:7828942] |
| 5 | lncRNA | 1.12E+08 | 1.12E+08 | ENSMUSG00000104538 | MGI:1922890 | 1700016B01Rik | RIKEN cDNA 1700016B01 gene [Source:MGI Symbol;Acc:MGI:1922890] |
| 5 | lncRNA | 1.12E+08 | 1.12E+08 | ENSMUSG00000139184 | MGI:7828820 | Gm73031 | predicted gene, 73031 [Source:MGI Symbol;Acc:MGI:7828820] |
| 5 | lncRNA | 1.12E+08 | 1.12E+08 | ENSMUSG00000133363 | MGI:7827134 | Gm72184 | predicted gene, 72184 [Source:MGI Symbol;Acc:MGI:7827134] |
| 5 | lncRNA | 1.12E+08 | 1.12E+08 | ENSMUSG00000101683 | MGI:1922819 | 1700028D13Rik | RIKEN cDNA 1700028D13 gene [Source:MGI Symbol;Acc:MGI:1922819] |
| 5 | lncRNA | 1.12E+08 | 1.12E+08 | ENSMUSG00000097767 | MGI:2444886 | Miat | myocardial infarction associated transcript (non-protein coding) [Source:MGI Symbol;Acc:MGI:2444886] |
| 5 | lncRNA | 1.12E+08 | 1.12E+08 | ENSMUSG00000097986 | MGI:5504068 | Gm26953 | predicted gene, 26953 [Source:MGI Symbol;Acc:MGI:5504068] |
| 5 | lncRNA | 1.12E+08 | 1.12E+08 | ENSMUSG00000133381 | MGI:7827140 | Gm72187 | predicted gene, 72187 [Source:MGI Symbol;Acc:MGI:7827140] |
| 5 | lncRNA | 1.12E+08 | 1.12E+08 | ENSMUSG00000138023 | MGI:5826578 | Gm46941 | predicted gene, 46941 [Source:MGI Symbol;Acc:MGI:5826578] |
| 5 | lncRNA | 1.13E+08 | 1.13E+08 | ENSMUSG00000093637 | MGI:5313083 | Gm20636 | predicted gene 20636 [Source:MGI Symbol;Acc:MGI:5313083] |
| 5 | lncRNA | 1.13E+08 | 1.13E+08 | ENSMUSG00000130751 | MGI:7826280 | Gm71756 | predicted gene, 71756 [Source:MGI Symbol;Acc:MGI:7826280] |
| 5 | lncRNA | 1.13E+08 | 1.13E+08 | ENSMUSG00000124762 | MGI:7825561 | Gm71394 | predicted gene, 71394 [Source:MGI Symbol;Acc:MGI:7825561] |

| | | | | | | | |
|---|---|---|---|---|---|---|---|
| 5 | lncRNA | 1.13E+08 | 1.13E+08 | ENSMUSG00000135548 | MGI:7827732 | Gm72484 | predicted gene, 72484 [Source:MGI Symbol;Acc:MGI:7827732] |
| 5 | lncRNA | 1.13E+08 | 1.13E+08 | ENSMUSG00000124512 | MGI:7825537 | Gm71382 | predicted gene, 71382 [Source:MGI Symbol;Acc:MGI:7825537] |
| 5 | lncRNA | 1.13E+08 | 1.13E+08 | ENSMUSG00000124546 | MGI:7825543 | Gm71385 | predicted gene, 71385 [Source:MGI Symbol;Acc:MGI:7825543] |
| 5 | lncRNA | 1.13E+08 | 1.13E+08 | ENSMUSG00000097962 | MGI:1914575 | 1700034G24Rik | RIKEN cDNA 1700034G24 gene [Source:MGI Symbol;Acc:MGI:1914575] |
| 5 | lncRNA | 1.13E+08 | 1.13E+08 | ENSMUSG00000098027 | MGI:5504119 | Gm27004 | predicted gene, 27004 [Source:MGI Symbol;Acc:MGI:5504119] |
| 5 | lncRNA | 1.13E+08 | 1.13E+08 | ENSMUSG00000122359 | MGI:7825255 | Gm71237 | predicted gene, 71237 [Source:MGI Symbol;Acc:MGI:7825255] |
| 5 | lncRNA | 1.13E+08 | 1.13E+08 | ENSMUSG00000104807 | MGI:5663002 | Gm42865 | predicted gene 42865 [Source:MGI Symbol;Acc:MGI:5663002] |
| 5 | lncRNA | 1.13E+08 | 1.13E+08 | ENSMUSG00000127191 | MGI:7825797 | Gm71512 | predicted gene, 71512 [Source:MGI Symbol;Acc:MGI:7825797] |
| 5 | lncRNA | 1.13E+08 | 1.13E+08 | ENSMUSG00000098230 | MGI:1914610 | 1700095B10Rik | RIKEN cDNA 1700095B10 gene [Source:MGI Symbol;Acc:MGI:1914610] |
| 5 | lncRNA | 1.13E+08 | 1.13E+08 | ENSMUSG00000129505 | MGI:7825985 | Gm71606 | predicted gene, 71606 [Source:MGI Symbol;Acc:MGI:7825985] |
| 5 | lncRNA | 1.13E+08 | 1.13E+08 | ENSMUSG00000134035 | MGI:7827316 | Gm72275 | predicted gene, 72275 [Source:MGI Symbol;Acc:MGI:7827316] |
| 5 | lncRNA | 1.13E+08 | 1.13E+08 | ENSMUSG00000135925 | MGI:7827794 | Gm72515 | predicted gene, 72515 [Source:MGI Symbol;Acc:MGI:7827794] |
| 5 | lncRNA | 1.13E+08 | 1.13E+08 | ENSMUSG00000136353 | MGI:7827946 | Gm72592 | predicted gene, 72592 [Source:MGI Symbol;Acc:MGI:7827946] |
| 5 | lncRNA | 1.13E+08 | 1.13E+08 | ENSMUSG00000137004 | MGI:7828260 | Gm72749 | predicted gene, 72749 [Source:MGI Symbol;Acc:MGI:7828260] |
| 5 | lncRNA | 1.13E+08 | 1.13E+08 | ENSMUSG00000141774 | MGI:1926036 | 4833439F03Rik | RIKEN cDNA 4833439F03 gene [Source:MGI Symbol;Acc:MGI:1926036] |
| 5 | lncRNA | 1.13E+08 | 1.13E+08 | ENSMUSG00000106659 | MGI:5625046 | Gm42161 | predicted gene, 42161 [Source:MGI Symbol;Acc:MGI:5625046] |
| 5 | lncRNA | 1.13E+08 | 1.13E+08 | ENSMUSG00000127988 | MGI:7825843 | Gm71535 | predicted gene, 71535 [Source:MGI Symbol;Acc:MGI:7825843] |
| 5 | lncRNA | 1.13E+08 | 1.13E+08 | ENSMUSG00000138601 | MGI:7828646 | Gm72943 | predicted gene, 72943 [Source:MGI Symbol;Acc:MGI:7828646] |

| | | | | | | | |
|---|---|---|---|---|---|---|---|
| 5 | lncRNA | 1.13E+08 | 1.13E+08 | ENSMUSG00000141062 | MGI:7829336 | Gm73289 | predicted gene, 73289 [Source:MGI Symbol;Acc:MGI:7829336] |
| 5 | lncRNA | 1.13E+08 | 1.13E+08 | ENSMUSG00000104705 | MGI:1921047 | 4930405N21Rik | RIKEN cDNA 4930405N21 gene [Source:MGI Symbol;Acc:MGI:1921047] |
| 5 | lncRNA | 1.13E+08 | 1.13E+08 | ENSMUSG00000105174 | MGI:5663490 | Gm43353 | predicted gene 43353 [Source:MGI Symbol;Acc:MGI:5663490] |
| 5 | lncRNA | 1.13E+08 | 1.13E+08 | ENSMUSG00000143806 | MGI:7829986 | Gm73615 | predicted gene, 73615 [Source:MGI Symbol;Acc:MGI:7829986] |
| 5 | lncRNA | 1.13E+08 | 1.14E+08 | ENSMUSG00000126830 | MGI:1921478 | 1700016C19Rik | RIKEN cDNA 1700016C19 gene [Source:MGI Symbol;Acc:MGI:1921478] |
| 5 | lncRNA | 1.14E+08 | 1.14E+08 | ENSMUSG00000126864 | MGI:7825761 | Gm71494 | predicted gene, 71494 [Source:MGI Symbol;Acc:MGI:7825761] |
| 5 | lncRNA | 1.14E+08 | 1.14E+08 | ENSMUSG00000053873 | MGI:1891321 | Aym1 | activator of yeast meiotic promoters 1 [Source:MGI Symbol;Acc:MGI:1891321] |
| 5 | lncRNA | 1.14E+08 | 1.14E+08 | ENSMUSG00000133155 | MGI:7826994 | Gm72114 | predicted gene, 72114 [Source:MGI Symbol;Acc:MGI:7826994] |
| 5 | lncRNA | 1.14E+08 | 1.14E+08 | ENSMUSG00000143804 | MGI:7829980 | Gm73612 | predicted gene, 73612 [Source:MGI Symbol;Acc:MGI:7829980] |
| 5 | lncRNA | 1.14E+08 | 1.14E+08 | ENSMUSG00000140157 | MGI:7829054 | Gm73148 | predicted gene, 73148 [Source:MGI Symbol;Acc:MGI:7829054] |
| 5 | lncRNA | 1.14E+08 | 1.14E+08 | ENSMUSG00000139134 | MGI:7828798 | Gm73019 | predicted gene, 73019 [Source:MGI Symbol;Acc:MGI:7828798] |
| 5 | lncRNA | 1.14E+08 | 1.14E+08 | ENSMUSG00000131097 | MGI:5589971 | Gm30812 | predicted gene, 30812 [Source:MGI Symbol;Acc:MGI:5589971] |
| 5 | lncRNA | 1.14E+08 | 1.14E+08 | ENSMUSG00000131138 | MGI:7826418 | Gm71826 | predicted gene, 71826 [Source:MGI Symbol;Acc:MGI:7826418] |
| 5 | lncRNA | 1.14E+08 | 1.14E+08 | ENSMUSG00000120761 | MGI:7338541 | Gm57314 | predicted gene, 57314 [Source:MGI Symbol;Acc:MGI:7338541] |
| 5 | lncRNA | 1.14E+08 | 1.14E+08 | ENSMUSG00000126419 | MGI:7825721 | Gm71474 | predicted gene, 71474 [Source:MGI Symbol;Acc:MGI:7825721] |
| 5 | lncRNA | 1.14E+08 | 1.14E+08 | ENSMUSG00000126378 | MGI:7825719 | Gm71473 | predicted gene, 71473 [Source:MGI Symbol;Acc:MGI:7825719] |
| 5 | lncRNA | 1.14E+08 | 1.14E+08 | ENSMUSG00000126337 | MGI:7825715 | Gm71471 | predicted gene, 71471 [Source:MGI Symbol;Acc:MGI:7825715] |
| 5 | lncRNA | 1.14E+08 | 1.14E+08 | ENSMUSG00000129427 | MGI:7825977 | Gm71602 | predicted gene, 71602 [Source:MGI Symbol;Acc:MGI:7825977] |

| | | | | | | | |
|---|---|---|---|---|---|---|---|
| 5 | lncRNA | 1.14E+08 | 1.14E+08 | ENSMUSG00000085713 | MGI:3783178 | Gm15736 | predicted gene 15736 [Source:MGI Symbol;Acc:MGI:3783178] |
| 5 | lncRNA | 1.14E+08 | 1.14E+08 | ENSMUSG00000134876 | MGI:7827550 | Gm72392 | predicted gene, 72392 [Source:MGI Symbol;Acc:MGI:7827550] |
| 5 | lncRNA | 1.14E+08 | 1.14E+08 | ENSMUSG00000125163 | MGI:7825589 | Gm71408 | predicted gene, 71408 [Source:MGI Symbol;Acc:MGI:7825589] |
| 5 | lncRNA | 1.14E+08 | 1.14E+08 | ENSMUSG00000125195 | MGI:7825595 | Gm71411 | predicted gene, 71411 [Source:MGI Symbol;Acc:MGI:7825595] |
| 5 | lncRNA | 1.14E+08 | 1.14E+08 | ENSMUSG00000042184 | MGI:1920726 | 1700069L16Rik | RIKEN cDNA 1700069L16 gene [Source:MGI Symbol;Acc:MGI:1920726] |
| 5 | lncRNA | 1.14E+08 | 1.14E+08 | ENSMUSG00000133974 | MGI:7827298 | Gm72266 | predicted gene, 72266 [Source:MGI Symbol;Acc:MGI:7827298] |
| 5 | lncRNA | 1.14E+08 | 1.14E+08 | ENSMUSG00000135715 | MGI:5624889 | Gm42004 | predicted gene, 42004 [Source:MGI Symbol;Acc:MGI:5624889] |
| 5 | lncRNA | 1.14E+08 | 1.14E+08 | ENSMUSG00000130821 | MGI:7826300 | Gm71766 | predicted gene, 71766 [Source:MGI Symbol;Acc:MGI:7826300] |
| 5 | lncRNA | 1.14E+08 | 1.14E+08 | ENSMUSG00000090458 | MGI:4937949 | Gm17122 | predicted gene 17122 [Source:MGI Symbol;Acc:MGI:4937949] |
| 5 | lncRNA | 1.14E+08 | 1.14E+08 | ENSMUSG00000123600 | MGI:7825423 | Gm71323 | predicted gene, 71323 [Source:MGI Symbol;Acc:MGI:7825423] |
| 5 | lncRNA | 1.14E+08 | 1.14E+08 | ENSMUSG00000105664 | MGI:5662934 | Gm42797 | predicted gene 42797 [Source:MGI Symbol;Acc:MGI:5662934] |
| 5 | lncRNA | 1.14E+08 | 1.14E+08 | ENSMUSG00000123190 | MGI:7825366 | Gm71294 | predicted gene, 71294 [Source:MGI Symbol;Acc:MGI:7825366] |
| 5 | lncRNA | 1.14E+08 | 1.14E+08 | ENSMUSG00000140843 | MGI:7829232 | Gm73237 | predicted gene, 73237 [Source:MGI Symbol;Acc:MGI:7829232] |
| 5 | lncRNA | 1.14E+08 | 1.14E+08 | ENSMUSG00000140808 | MGI:7829228 | Gm73235 | predicted gene, 73235 [Source:MGI Symbol;Acc:MGI:7829228] |
| 5 | lncRNA | 1.14E+08 | 1.14E+08 | ENSMUSG00000127137 | MGI:7825793 | Gm71510 | predicted gene, 71510 [Source:MGI Symbol;Acc:MGI:7825793] |
| 5 | lncRNA | 1.15E+08 | 1.15E+08 | ENSMUSG00000122018 | MGI:5593881 | Gm34722 | predicted gene, 34722 [Source:MGI Symbol;Acc:MGI:5593881] |
| 5 | lncRNA | 1.15E+08 | 1.15E+08 | ENSMUSG00000143223 | MGI:7829748 | Gm73496 | predicted gene, 73496 [Source:MGI Symbol;Acc:MGI:7829748] |
| 5 | lncRNA | 1.15E+08 | 1.15E+08 | ENSMUSG00000137482 | MGI:7828430 | Gm72834 | predicted gene, 72834 [Source:MGI Symbol;Acc:MGI:7828430] |

| | | | | | | | |
|---|---|---|---|---|---|---|---|
| 5 | lncRNA | 1.15E+08 | 1.15E+08 | ENSMUSG00000140498 | MGI:7829156 | Gm73199 | predicted gene, 73199 [Source:MGI Symbol;Acc:MGI:7829156] |
| 5 | lncRNA | 1.15E+08 | 1.15E+08 | ENSMUSG00000086302 | MGI:3650747 | Gm13790 | predicted gene 13790 [Source:MGI Symbol;Acc:MGI:3650747] |
| 5 | lncRNA | 1.15E+08 | 1.15E+08 | ENSMUSG00000127419 | MGI:7825809 | Gm71518 | predicted gene, 71518 [Source:MGI Symbol;Acc:MGI:7825809] |
| 5 | lncRNA | 1.15E+08 | 1.15E+08 | ENSMUSG00000127459 | MGI:7825811 | Gm71519 | predicted gene, 71519 [Source:MGI Symbol;Acc:MGI:7825811] |
| 5 | lncRNA | 1.15E+08 | 1.15E+08 | ENSMUSG00000140177 | MGI:7829060 | Gm73151 | predicted gene, 73151 [Source:MGI Symbol;Acc:MGI:7829060] |
| 5 | lncRNA | 1.15E+08 | 1.15E+08 | ENSMUSG00000122765 | MGI:5594003 | Gm34844 | predicted gene, 34844 [Source:MGI Symbol;Acc:MGI:5594003] |
| 5 | lncRNA | 1.15E+08 | 1.15E+08 | ENSMUSG00000127492 | MGI:7825813 | Gm71520 | predicted gene, 71520 [Source:MGI Symbol;Acc:MGI:7825813] |
| 5 | lncRNA | 1.15E+08 | 1.15E+08 | ENSMUSG00000141988 | MGI:7829480 | Gm73361 | predicted gene, 73361 [Source:MGI Symbol;Acc:MGI:7829480] |
| 5 | lncRNA | 1.15E+08 | 1.15E+08 | ENSMUSG00000139305 | MGI:5594090 | Gm34931 | predicted gene, 34931 [Source:MGI Symbol;Acc:MGI:5594090] |
| 5 | lncRNA | 1.15E+08 | 1.15E+08 | ENSMUSG00000139348 | MGI:7828852 | Gm73047 | predicted gene, 73047 [Source:MGI Symbol;Acc:MGI:7828852] |
| 5 | lncRNA | 1.15E+08 | 1.15E+08 | ENSMUSG00000107340 | MGI:5662926 | Gm42789 | predicted gene 42789 [Source:MGI Symbol;Acc:MGI:5662926] |
| 5 | lncRNA | 1.15E+08 | 1.15E+08 | ENSMUSG00000092183 | MGI:1914892 | 4930515G01Rik | RIKEN cDNA 4930515G01 gene [Source:MGI Symbol;Acc:MGI:1914892] |
| 5 | lncRNA | 1.15E+08 | 1.15E+08 | ENSMUSG00000133025 | MGI:7826956 | Gm72095 | predicted gene, 72095 [Source:MGI Symbol;Acc:MGI:7826956] |
| 5 | lncRNA | 1.15E+08 | 1.15E+08 | ENSMUSG00000133116 | MGI:7826968 | Gm72101 | predicted gene, 72101 [Source:MGI Symbol;Acc:MGI:7826968] |
| 5 | lncRNA | 1.15E+08 | 1.15E+08 | ENSMUSG00000132938 | MGI:7826888 | Gm72061 | predicted gene, 72061 [Source:MGI Symbol;Acc:MGI:7826888] |
| 5 | lncRNA | 1.15E+08 | 1.15E+08 | ENSMUSG00000132979 | MGI:7826908 | Gm72071 | predicted gene, 72071 [Source:MGI Symbol;Acc:MGI:7826908] |
| 5 | lncRNA | 1.15E+08 | 1.15E+08 | ENSMUSG00000131648 | MGI:7826602 | Gm71918 | predicted gene, 71918 [Source:MGI Symbol;Acc:MGI:7826602] |
| 5 | lncRNA | 1.15E+08 | 1.15E+08 | ENSMUSG00000107108 | MGI:3641854 | Gm9936 | predicted gene 9936 [Source:MGI Symbol;Acc:MGI:3641854] |

| | | | | | | | |
|---|---|---|---|---|---|---|---|
| 5 | lncRNA | 1.15E+08 | 1.15E+08 | ENSMUSG00000135230 | MGI:7827654 | Gm72445 | predicted gene, 72445 [Source:MGI Symbol;Acc:MGI:7827654] |
| 5 | lncRNA | 1.15E+08 | 1.15E+08 | ENSMUSG00000087477 | MGI:3650351 | Gm13822 | predicted gene 13822 [Source:MGI Symbol;Acc:MGI:3650351] |
| 5 | lncRNA | 1.15E+08 | 1.15E+08 | ENSMUSG00000122036 | MGI:7825201 | Gm71210 | predicted gene, 71210 [Source:MGI Symbol;Acc:MGI:7825201] |
| 5 | lncRNA | 1.15E+08 | 1.15E+08 | ENSMUSG00000138573 | MGI:7828632 | Gm72936 | predicted gene, 72936 [Source:MGI Symbol;Acc:MGI:7828632] |
| 5 | lncRNA | 1.15E+08 | 1.15E+08 | ENSMUSG00000129685 | MGI:7826015 | Gm71622 | predicted gene, 71622 [Source:MGI Symbol;Acc:MGI:7826015] |
| 5 | lncRNA | 1.15E+08 | 1.15E+08 | ENSMUSG00000086054 | MGI:3652225 | Hnf1aos1 | HNF1 homeobox A, opposite strand 1 [Source:MGI Symbol;Acc:MGI:3652225] |
| 5 | lncRNA | 1.15E+08 | 1.15E+08 | ENSMUSG00000086140 | MGI:3652220 | Hnf1aos2 | HNF1 homeobox A, opposite strand 2 [Source:MGI Symbol;Acc:MGI:3652220] |
| 5 | lncRNA | 1.15E+08 | 1.15E+08 | ENSMUSG00000129727 | MGI:7826021 | Gm71625 | predicted gene, 71625 [Source:MGI Symbol;Acc:MGI:7826021] |
| 5 | lncRNA | 1.15E+08 | 1.15E+08 | ENSMUSG00000121818 | | | ribosomal protein L37, retrotransposed [Source:NCBI gene (formerly Entrezgene);Acc:100502825] |
| 5 | lncRNA | 1.15E+08 | 1.15E+08 | ENSMUSG00000136454 | MGI:7827964 | Gm72601 | predicted gene, 72601 [Source:MGI Symbol;Acc:MGI:7827964] |
| 5 | lncRNA | 1.15E+08 | 1.15E+08 | ENSMUSG00000120554 | MGI:7338449 | Gm57268 | predicted gene, 57268 [Source:MGI Symbol;Acc:MGI:7338449] |
| 5 | lncRNA | 1.15E+08 | 1.15E+08 | ENSMUSG00000128459 | MGI:7825901 | Gm71564 | predicted gene, 71564 [Source:MGI Symbol;Acc:MGI:7825901] |
| 5 | lncRNA | 1.15E+08 | 1.15E+08 | ENSMUSG00000086368 | MGI:3651141 | Gm13830 | predicted gene 13830 [Source:MGI Symbol;Acc:MGI:3651141] |
| 5 | lncRNA | 1.15E+08 | 1.15E+08 | ENSMUSG00000121115 | MGI:7338553 | Gm57320 | predicted gene, 57320 [Source:MGI Symbol;Acc:MGI:7338553] |
| 5 | lncRNA | 1.15E+08 | 1.15E+08 | ENSMUSG00000107288 | MGI:1925175 | A430102K17Rik | RIKEN cDNA A430102K17 gene [Source:MGI Symbol;Acc:MGI:1925175] |
| 5 | lncRNA | 1.15E+08 | 1.16E+08 | ENSMUSG00000086352 | MGI:1921042 | 4930401G09Rik | RIKEN cDNA 4930401G09 gene [Source:MGI Symbol;Acc:MGI:1921042] |
| 5 | lncRNA | 1.15E+08 | 1.16E+08 | ENSMUSG00000087292 | MGI:3650708 | Gm13832 | predicted gene 13832 [Source:MGI Symbol;Acc:MGI:3650708] |

| | | | | | | | |
|---|---|---|---|---|---|---|---|
| 5 | lncRNA | 1.16E+08 | 1.16E+08 | ENSMUSG00000124350 | | | [Source:NCBI gene (formerly Entrezgene);Acc:102639568] |
| 5 | lncRNA | 1.16E+08 | 1.16E+08 | ENSMUSG00000107353 | MGI:1925404 | 4930430O22Rik | RIKEN cDNA 4930430O22 gene [Source:MGI Symbol;Acc:MGI:1925404] |
| 5 | lncRNA | 1.16E+08 | 1.16E+08 | ENSMUSG00000127019 | MGI:7825779 | Gm71503 | predicted gene, 71503 [Source:MGI Symbol;Acc:MGI:7825779] |
| 5 | lncRNA | 1.16E+08 | 1.16E+08 | ENSMUSG00000120846 | MGI:7338509 | Gm57298 | predicted gene, 57298 [Source:MGI Symbol;Acc:MGI:7338509] |
| 5 | lncRNA | 1.16E+08 | 1.16E+08 | ENSMUSG00000121040 | MGI:7338523 | Gm57305 | predicted gene, 57305 [Source:MGI Symbol;Acc:MGI:7338523] |
| 5 | lncRNA | 1.16E+08 | 1.16E+08 | ENSMUSG00000132950 | MGI:7826896 | Gm72065 | predicted gene, 72065 [Source:MGI Symbol;Acc:MGI:7826896] |
| 5 | lncRNA | 1.16E+08 | 1.16E+08 | ENSMUSG00000126088 | MGI:7830410 | Gm73833 | predicted gene, 73833 [Source:MGI Symbol;Acc:MGI:7830410] |
| 5 | lncRNA | 1.16E+08 | 1.16E+08 | ENSMUSG00000130738 | MGI:7826278 | Gm71755 | predicted gene, 71755 [Source:MGI Symbol;Acc:MGI:7826278] |
| 5 | lncRNA | 1.16E+08 | 1.16E+08 | ENSMUSG00000139020 | MGI:7828780 | Gm73010 | predicted gene, 73010 [Source:MGI Symbol;Acc:MGI:7828780] |
| 5 | lncRNA | 1.16E+08 | 1.16E+08 | ENSMUSG00000086849 | MGI:3649479 | Gm13840 | predicted gene 13840 [Source:MGI Symbol;Acc:MGI:3649479] |
| 5 | lncRNA | 1.16E+08 | 1.16E+08 | ENSMUSG00000140383 | MGI:7829134 | Gm73188 | predicted gene, 73188 [Source:MGI Symbol;Acc:MGI:7829134] |
| 5 | lncRNA | 1.16E+08 | 1.16E+08 | ENSMUSG00000107121 | MGI:1913373 | 1110006O24Rik | RIKEN cDNA 1110006O24 gene [Source:MGI Symbol;Acc:MGI:1913373] |
| 5 | lncRNA | 1.16E+08 | 1.16E+08 | ENSMUSG00000135244 | MGI:7827656 | Gm72446 | predicted gene, 72446 [Source:MGI Symbol;Acc:MGI:7827656] |
| 5 | lncRNA | 1.16E+08 | 1.16E+08 | ENSMUSG00000125952 | MGI:7825669 | Gm71448 | predicted gene, 71448 [Source:MGI Symbol;Acc:MGI:7825669] |
| 5 | lncRNA | 1.16E+08 | 1.16E+08 | ENSMUSG00000125719 | MGI:7825645 | Gm71436 | predicted gene, 71436 [Source:MGI Symbol;Acc:MGI:7825645] |
| 5 | lncRNA | 1.16E+08 | 1.16E+08 | ENSMUSG00000134680 | MGI:5624893 | Gm42008 | predicted gene, 42008 [Source:MGI Symbol;Acc:MGI:5624893] |
| 5 | lncRNA | 1.16E+08 | 1.16E+08 | ENSMUSG00000120953 | MGI:5595344 | Gm36185 | predicted gene, 36185 [Source:MGI Symbol;Acc:MGI:5595344] |
| 5 | lncRNA | 1.16E+08 | 1.16E+08 | ENSMUSG00000085376 | MGI:3705279 | Gm14508 | predicted gene 14508 [Source:MGI Symbol;Acc:MGI:3705279] |

| | | | | | | | |
|---|---|---|---|---|---|---|---|
| 5 | lncRNA | 1.16E+08 | 1.16E+08 | ENSMUSG00000084992 | MGI:3650892 | Gm13842 | predicted gene 13842 [Source:MGI Symbol;Acc:MGI:3650892] |
| 5 | lncRNA | 1.16E+08 | 1.16E+08 | ENSMUSG00000131869 | MGI:7826640 | Gm71937 | predicted gene, 71937 [Source:MGI Symbol;Acc:MGI:7826640] |
| 5 | lncRNA | 1.16E+08 | 1.16E+08 | ENSMUSG00000127595 | MGI:7825825 | Gm71526 | predicted gene, 71526 [Source:MGI Symbol;Acc:MGI:7825825] |
| 5 | lncRNA | 1.16E+08 | 1.16E+08 | ENSMUSG00000086655 | MGI:2444781 | C330018A13Rik | RIKEN cDNA C330018A13 gene [Source:MGI Symbol;Acc:MGI:2444781] |
| 5 | lncRNA | 1.16E+08 | 1.16E+08 | ENSMUSG00000086122 | MGI:3705240 | Gm14507 | predicted gene 14507 [Source:MGI Symbol;Acc:MGI:3705240] |
| 5 | lncRNA | 1.16E+08 | 1.16E+08 | ENSMUSG00000086055 | MGI:3650972 | Gm13837 | predicted gene 13837 [Source:MGI Symbol;Acc:MGI:3650972] |
| 5 | lncRNA | 1.16E+08 | 1.16E+08 | ENSMUSG00000131570 | MGI:7826574 | Gm71904 | predicted gene, 71904 [Source:MGI Symbol;Acc:MGI:7826574] |
| 5 | lncRNA | 1.16E+08 | 1.16E+08 | ENSMUSG00000087124 | MGI:3650513 | Gm13839 | predicted gene 13839 [Source:MGI Symbol;Acc:MGI:3650513] |
| 5 | lncRNA | 1.16E+08 | 1.16E+08 | ENSMUSG00000084921 | MGI:3649377 | Gm13838 | predicted gene 13838 [Source:MGI Symbol;Acc:MGI:3649377] |
| 5 | lncRNA | 1.16E+08 | 1.16E+08 | ENSMUSG00000086715 | MGI:1925096 | B230112J18Rik | RIKEN cDNA B230112J18 gene [Source:MGI Symbol;Acc:MGI:1925096] |
| 5 | lncRNA | 1.17E+08 | 1.17E+08 | ENSMUSG00000126742 | MGI:7825751 | Gm71489 | predicted gene, 71489 [Source:MGI Symbol;Acc:MGI:7825751] |
| 5 | lncRNA | 1.17E+08 | 1.17E+08 | ENSMUSG00000086945 | MGI:1922553 | 4930562A09Rik | RIKEN cDNA 4930562A09 gene [Source:MGI Symbol;Acc:MGI:1922553] |
| 5 | lncRNA | 1.17E+08 | 1.17E+08 | ENSMUSG00000133941 | MGI:7827262 | Gm72248 | predicted gene, 72248 [Source:MGI Symbol;Acc:MGI:7827262] |
| 5 | lncRNA | 1.17E+08 | 1.17E+08 | ENSMUSG00000134017 | MGI:5624894 | Gm42009 | predicted gene, 42009 [Source:MGI Symbol;Acc:MGI:5624894] |
| 5 | lncRNA | 1.17E+08 | 1.17E+08 | ENSMUSG00000134063 | MGI:7827330 | Gm72282 | predicted gene, 72282 [Source:MGI Symbol;Acc:MGI:7827330] |
| 5 | lncRNA | 1.17E+08 | 1.17E+08 | ENSMUSG00000086219 | MGI:1924048 | Srrm4os | serine/arginine repetitive matrix 4, opposite strand [Source:MGI Symbol;Acc:MGI:1924048] |
| 5 | lncRNA | 1.17E+08 | 1.17E+08 | ENSMUSG00000134106 | MGI:7827362 | Gm72298 | predicted gene, 72298 [Source:MGI Symbol;Acc:MGI:7827362] |
| 5 | lncRNA | 1.17E+08 | 1.17E+08 | ENSMUSG00000106914 | MGI:5662991 | Gm42854 | predicted gene 42854 [Source:MGI Symbol;Acc:MGI:5662991] |

| | | | | | | | |
|---|---|---|---|---|---|---|---|
| 5 | lncRNA | 1.17E+08 | 1.17E+08 | ENSMUSG00000132062 | MGI:7826700 | Gm71967 | predicted gene, 71967 [Source:MGI Symbol;Acc:MGI:7826700] |
| 5 | lncRNA | 1.17E+08 | 1.17E+08 | ENSMUSG00000132023 | MGI:7826688 | Gm71961 | predicted gene, 71961 [Source:MGI Symbol;Acc:MGI:7826688] |
| 5 | lncRNA | 1.17E+08 | 1.17E+08 | ENSMUSG00000106909 | MGI:5663259 | Gm43122 | predicted gene 43122 [Source:MGI Symbol;Acc:MGI:5663259] |
| 5 | lncRNA | 1.17E+08 | 1.17E+08 | ENSMUSG00000128674 | MGI:7825923 | Gm71575 | predicted gene, 71575 [Source:MGI Symbol;Acc:MGI:7825923] |
| 5 | lncRNA | 1.17E+08 | 1.17E+08 | ENSMUSG00000128640 | MGI:5589002 | Gm29843 | predicted gene, 29843 [Source:MGI Symbol;Acc:MGI:5589002] |
| 5 | lncRNA | 1.17E+08 | 1.17E+08 | ENSMUSG00000107237 | MGI:5589085 | Gm29926 | predicted gene, 29926 [Source:MGI Symbol;Acc:MGI:5589085] |
| 5 | lncRNA | 1.17E+08 | 1.17E+08 | ENSMUSG00000128718 | MGI:7825929 | Gm71578 | predicted gene, 71578 [Source:MGI Symbol;Acc:MGI:7825929] |
| 5 | lncRNA | 1.17E+08 | 1.17E+08 | ENSMUSG00000128565 | MGI:7825917 | Gm71572 | predicted gene, 71572 [Source:MGI Symbol;Acc:MGI:7825917] |
| 5 | lncRNA | 1.17E+08 | 1.17E+08 | ENSMUSG00000133049 | MGI:7826960 | Gm72097 | predicted gene, 72097 [Source:MGI Symbol;Acc:MGI:7826960] |
| 5 | lncRNA | 1.17E+08 | 1.17E+08 | ENSMUSG00000130385 | MGI:7826208 | Gm71720 | predicted gene, 71720 [Source:MGI Symbol;Acc:MGI:7826208] |
| 5 | lncRNA | 1.17E+08 | 1.17E+08 | ENSMUSG00000131824 | MGI:5624895 | Gm42010 | predicted gene, 42010 [Source:MGI Symbol;Acc:MGI:5624895] |
| 5 | lncRNA | 1.17E+08 | 1.17E+08 | ENSMUSG00000128116 | MGI:7825855 | Gm71541 | predicted gene, 71541 [Source:MGI Symbol;Acc:MGI:7825855] |
| 5 | lncRNA | 1.17E+08 | 1.17E+08 | ENSMUSG00000139383 | MGI:7828860 | Gm73051 | predicted gene, 73051 [Source:MGI Symbol;Acc:MGI:7828860] |
| 5 | lncRNA | 1.17E+08 | 1.17E+08 | ENSMUSG00000129200 | MGI:5589131 | Gm29972 | predicted gene, 29972 [Source:MGI Symbol;Acc:MGI:5589131] |
| 5 | lncRNA | 1.17E+08 | 1.17E+08 | ENSMUSG00000129242 | MGI:5589187 | Gm30028 | predicted gene, 30028 [Source:MGI Symbol;Acc:MGI:5589187] |
| 5 | lncRNA | 1.17E+08 | 1.17E+08 | ENSMUSG00000129278 | MGI:7825969 | Gm71598 | predicted gene, 71598 [Source:MGI Symbol;Acc:MGI:7825969] |
| 5 | lncRNA | 1.17E+08 | 1.17E+08 | ENSMUSG00000086047 | MGI:1925671 | 9530046B11Rik | RIKEN cDNA 9530046B11 gene [Source:MGI Symbol;Acc:MGI:1925671] |
| 5 | lncRNA | 1.17E+08 | 1.17E+08 | ENSMUSG00000120368 | MGI:7338439 | Gm57263 | predicted gene, 57263 [Source:MGI Symbol;Acc:MGI:7338439] |

| | | | | | | | |
|---|---|---|---|---|---|---|---|
| 5 | lncRNA | 1.17E+08 | 1.17E+08 | ENSMUSG00000136822 | MGI:7828180 | Gm72709 | predicted gene, 72709 [Source:MGI Symbol;Acc:MGI:7828180] |
| 5 | lncRNA | 1.17E+08 | 1.17E+08 | ENSMUSG00000126795 | MGI:6367266 | Gm52828 | predicted gene, 52828 [Source:MGI Symbol;Acc:MGI:6367266] |
| 5 | lncRNA | 1.17E+08 | 1.17E+08 | ENSMUSG00000126840 | MGI:7825757 | Gm71492 | predicted gene, 71492 [Source:MGI Symbol;Acc:MGI:7825757] |
| 5 | lncRNA | 1.17E+08 | 1.17E+08 | ENSMUSG00000097163 | MGI:3039587 | BC051077 | cDNA sequence BC051077 [Source:MGI Symbol;Acc:MGI:3039587] |
| 5 | lncRNA | 1.17E+08 | 1.17E+08 | ENSMUSG00000072688 | MGI:3704255 | Gm10399 | predicted gene 10399 [Source:MGI Symbol;Acc:MGI:3704255] |
| 5 | lncRNA | 1.17E+08 | 1.17E+08 | ENSMUSG00000122690 | MGI:5624897 | Gm42012 | predicted gene, 42012 [Source:MGI Symbol;Acc:MGI:5624897] |
| 5 | lncRNA | 1.18E+08 | 1.18E+08 | ENSMUSG00000086075 | MGI:3783171 | Gm15728 | predicted gene 15728 [Source:MGI Symbol;Acc:MGI:3783171] |
| 5 | lncRNA | 1.18E+08 | 1.18E+08 | ENSMUSG00000120827 | MGI:6367268 | Gm52829 | predicted gene, 52829 [Source:MGI Symbol;Acc:MGI:6367268] |
| 5 | lncRNA | 1.18E+08 | 1.18E+08 | ENSMUSG00000128985 | MGI:7825945 | Gm71586 | predicted gene, 71586 [Source:MGI Symbol;Acc:MGI:7825945] |
| 5 | lncRNA | 1.18E+08 | 1.18E+08 | ENSMUSG00000138250 | MGI:7828548 | Gm72894 | predicted gene, 72894 [Source:MGI Symbol;Acc:MGI:7828548] |
| 5 | lncRNA | 1.18E+08 | 1.18E+08 | ENSMUSG00000126651 | MGI:7825737 | Gm71482 | predicted gene, 71482 [Source:MGI Symbol;Acc:MGI:7825737] |
| 5 | lncRNA | 1.18E+08 | 1.18E+08 | ENSMUSG00000123983 | MGI:7825465 | Gm71345 | predicted gene, 71345 [Source:MGI Symbol;Acc:MGI:7825465] |
| 5 | lncRNA | 1.18E+08 | 1.18E+08 | ENSMUSG00000128859 | MGI:7831172 | Gm74217 | predicted gene, 74217 [Source:MGI Symbol;Acc:MGI:7831172] |
| 5 | lncRNA | 1.18E+08 | 1.18E+08 | ENSMUSG00000137379 | MGI:5589519 | Gm30360 | predicted gene, 30360 [Source:MGI Symbol;Acc:MGI:5589519] |
| 5 | lncRNA | 1.18E+08 | 1.18E+08 | ENSMUSG00000136798 | MGI:7828174 | Gm72706 | predicted gene, 72706 [Source:MGI Symbol;Acc:MGI:7828174] |
| 5 | lncRNA | 1.18E+08 | 1.18E+08 | ENSMUSG00000098072 | MGI:5504110 | Gm26995 | predicted gene, 26995 [Source:MGI Symbol;Acc:MGI:5504110] |
| 5 | lncRNA | 1.18E+08 | 1.18E+08 | ENSMUSG00000029360 | MGI:3704256 | Gm9754 | predicted gene 9754 [Source:MGI Symbol;Acc:MGI:3704256] |
| 5 | lncRNA | 1.18E+08 | 1.18E+08 | ENSMUSG00000139919 | MGI:7828972 | Gm73107 | predicted gene, 73107 [Source:MGI Symbol;Acc:MGI:7828972] |

| | | | | | | | |
|---|---|---|---|---|---|---|---|
| 5 | lncRNA | 1.18E+08 | 1.18E+08 | ENSMUSG00000141016 | MGI:7829290 | Gm73266 | predicted gene, 73266 [Source:MGI Symbol;Acc:MGI:7829290] |
| 5 | lncRNA | 1.18E+08 | 1.18E+08 | ENSMUSG00000143393 | MGI:7829804 | Gm73524 | predicted gene, 73524 [Source:MGI Symbol;Acc:MGI:7829804] |
| 5 | lncRNA | 1.18E+08 | 1.18E+08 | ENSMUSG00000141505 | MGI:7829394 | Gm73318 | predicted gene, 73318 [Source:MGI Symbol;Acc:MGI:7829394] |
| 5 | lncRNA | 1.18E+08 | 1.18E+08 | ENSMUSG00000140965 | MGI:7829266 | Gm73254 | predicted gene, 73254 [Source:MGI Symbol;Acc:MGI:7829266] |
| 5 | lncRNA | 1.18E+08 | 1.18E+08 | ENSMUSG00000122569 | MGI:5826579 | Gm46942 | predicted gene, 46942 [Source:MGI Symbol;Acc:MGI:5826579] |
| 5 | lncRNA | 1.18E+08 | 1.19E+08 | ENSMUSG00000128713 | MGI:7825925 | Gm71576 | predicted gene, 71576 [Source:MGI Symbol;Acc:MGI:7825925] |
| 5 | lncRNA | 1.19E+08 | 1.19E+08 | ENSMUSG00000141024 | MGI:5589763 | Gm30604 | predicted gene, 30604 [Source:MGI Symbol;Acc:MGI:5589763] |
| 5 | lncRNA | 1.19E+08 | 1.19E+08 | ENSMUSG00000106812 | MGI:5579269 | Gm28563 | predicted gene 28563 [Source:MGI Symbol;Acc:MGI:5579269] |
| 5 | lncRNA | 1.19E+08 | 1.19E+08 | ENSMUSG00000138714 | MGI:7828674 | Gm72957 | predicted gene, 72957 [Source:MGI Symbol;Acc:MGI:7828674] |
| 5 | lncRNA | 1.19E+08 | 1.19E+08 | ENSMUSG00000127080 | MGI:7825787 | Gm71507 | predicted gene, 71507 [Source:MGI Symbol;Acc:MGI:7825787] |
| 5 | lncRNA | 1.19E+08 | 1.19E+08 | ENSMUSG00000126021 | MGI:5589905 | Gm30746 | predicted gene, 30746 [Source:MGI Symbol;Acc:MGI:5589905] |
| 5 | lncRNA | 1.19E+08 | 1.19E+08 | ENSMUSG00000130380 | MGI:5826558 | Gm46921 | predicted gene, 46921 [Source:MGI Symbol;Acc:MGI:5826558] |
| 5 | lncRNA | 1.19E+08 | 1.19E+08 | ENSMUSG00000121735 | MGI:7826194 | Gm71713 | predicted gene, 71713 [Source:MGI Symbol;Acc:MGI:7826194] |
| 5 | lncRNA | 1.19E+08 | 1.19E+08 | ENSMUSG00000128021 | MGI:7825847 | Gm71537 | predicted gene, 71537 [Source:MGI Symbol;Acc:MGI:7825847] |
| 5 | lncRNA | 1.19E+08 | 1.19E+08 | ENSMUSG00000120779 | MGI:7338499 | Gm57293 | predicted gene, 57293 [Source:MGI Symbol;Acc:MGI:7338499] |
| 5 | lncRNA | 1.19E+08 | 1.19E+08 | ENSMUSG00000130938 | MGI:7826312 | Gm71773 | predicted gene, 71773 [Source:MGI Symbol;Acc:MGI:7826312] |
| 5 | lncRNA | 1.19E+08 | 1.19E+08 | ENSMUSG00000139964 | MGI:7828978 | Gm73110 | predicted gene, 73110 [Source:MGI Symbol;Acc:MGI:7828978] |
| 5 | lncRNA | 1.19E+08 | 1.19E+08 | ENSMUSG00000134410 | MGI:7827444 | Gm72339 | predicted gene, 72339 [Source:MGI Symbol;Acc:MGI:7827444] |

| | | | | | | | |
|---|---|---|---|---|---|---|---|
| 5 | lncRNA | 1.19E+08 | 1.19E+08 | ENSMUSG00000135281 | MGI:7827668 | Gm72452 | predicted gene, 72452 [Source:MGI Symbol;Acc:MGI:7827668] |
| 5 | lncRNA | 1.19E+08 | 1.19E+08 | ENSMUSG00000126319 | MGI:7825713 | Gm71470 | predicted gene, 71470 [Source:MGI Symbol;Acc:MGI:7825713] |
| 5 | lncRNA | 1.19E+08 | 1.19E+08 | ENSMUSG00000123304 | MGI:5624900 | Gm42015 | predicted gene, 42015 [Source:MGI Symbol;Acc:MGI:5624900] |
| 5 | lncRNA | 1.19E+08 | 1.19E+08 | ENSMUSG00000132475 | MGI:7826790 | Gm72012 | predicted gene, 72012 [Source:MGI Symbol;Acc:MGI:7826790] |
| 5 | lncRNA | 1.19E+08 | 1.19E+08 | ENSMUSG00000132438 | MGI:7826778 | Gm72006 | predicted gene, 72006 [Source:MGI Symbol;Acc:MGI:7826778] |
| 5 | lncRNA | 1.19E+08 | 1.19E+08 | ENSMUSG00000106708 | MGI:5663919 | Gm43782 | predicted gene 43782 [Source:MGI Symbol;Acc:MGI:5663919] |
| 5 | lncRNA | 1.19E+08 | 1.19E+08 | ENSMUSG00000106745 | MGI:5663479 | Gm43342 | predicted gene 43342 [Source:MGI Symbol;Acc:MGI:5663479] |
| 5 | lncRNA | 1.19E+08 | 1.19E+08 | ENSMUSG00000107144 | MGI:1921877 | 4930413E15Rik | RIKEN cDNA 4930413E15 gene [Source:MGI Symbol;Acc:MGI:1921877] |
| 5 | lncRNA | 1.19E+08 | 1.19E+08 | ENSMUSG00000142875 | MGI:5590054 | Gm30895 | predicted gene, 30895 [Source:MGI Symbol;Acc:MGI:5590054] |
| 5 | lncRNA | 1.19E+08 | 1.19E+08 | ENSMUSG00000142923 | MGI:5590193 | Gm31034 | predicted gene, 31034 [Source:MGI Symbol;Acc:MGI:5590193] |
| 5 | lncRNA | 1.19E+08 | 1.19E+08 | ENSMUSG00000143941 | MGI:7830054 | Gm73650 | predicted gene, 73650 [Source:MGI Symbol;Acc:MGI:7830054] |
| 5 | lncRNA | 1.19E+08 | 1.19E+08 | ENSMUSG00000100573 | MGI:1920727 | 1700081H04Rik | RIKEN cDNA 1700081H04 gene [Source:MGI Symbol;Acc:MGI:1920727] |
| 5 | lncRNA | 1.19E+08 | 1.19E+08 | ENSMUSG00000140902 | MGI:7829242 | Gm73242 | predicted gene, 73242 [Source:MGI Symbol;Acc:MGI:7829242] |
| 5 | lncRNA | 1.19E+08 | 1.19E+08 | ENSMUSG00000140937 | MGI:5590288 | Gm31129 | predicted gene, 31129 [Source:MGI Symbol;Acc:MGI:5590288] |
| 5 | lncRNA | 1.19E+08 | 1.19E+08 | ENSMUSG00000107334 | MGI:3647618 | Gm7538 | predicted gene 7538 [Source:MGI Symbol;Acc:MGI:3647618] |
| 5 | lncRNA | 1.19E+08 | 1.19E+08 | ENSMUSG00000137673 | MGI:7828464 | Gm72851 | predicted gene, 72851 [Source:MGI Symbol;Acc:MGI:7828464] |
| 5 | lncRNA | 1.19E+08 | 1.19E+08 | ENSMUSG00000137711 | MGI:7828470 | Gm72854 | predicted gene, 72854 [Source:MGI Symbol;Acc:MGI:7828470] |
| 5 | lncRNA | 1.19E+08 | 1.19E+08 | ENSMUSG00000140558 | MGI:7829190 | Gm73216 | predicted gene, 73216 [Source:MGI Symbol;Acc:MGI:7829190] |

| | | | | | | | |
|---|---|---|---|---|---|---|---|
| 5 | lncRNA | 1.19E+08 | 1.19E+08 | ENSMUSG00000133340 | MGI:5590349 | Gm31190 | predicted gene, 31190 [Source:MGI Symbol;Acc:MGI:5590349] |
| 5 | lncRNA | 1.19E+08 | 1.19E+08 | ENSMUSG00000141471 | MGI:7829392 | Gm73317 | predicted gene, 73317 [Source:MGI Symbol;Acc:MGI:7829392] |
| 5 | lncRNA | 1.19E+08 | 1.19E+08 | ENSMUSG00000120005 | MGI:7338421 | Gm57254 | predicted gene, 57254 [Source:MGI Symbol;Acc:MGI:7338421] |
| 5 | lncRNA | 1.19E+08 | 1.19E+08 | ENSMUSG00000141549 | MGI:5624901 | Gm42016 | predicted gene, 42016 [Source:MGI Symbol;Acc:MGI:5624901] |
| 5 | lncRNA | 1.20E+08 | 1.20E+08 | ENSMUSG00000143531 | MGI:7829882 | Gm73563 | predicted gene, 73563 [Source:MGI Symbol;Acc:MGI:7829882] |
| 5 | lncRNA | 1.20E+08 | 1.20E+08 | ENSMUSG00000139730 | MGI:7828944 | Gm73093 | predicted gene, 73093 [Source:MGI Symbol;Acc:MGI:7828944] |
| 5 | lncRNA | 1.20E+08 | 1.20E+08 | ENSMUSG00000106938 | MGI:5590473 | Gm31314 | predicted gene, 31314 [Source:MGI Symbol;Acc:MGI:5590473] |
| 5 | lncRNA | 1.20E+08 | 1.20E+08 | ENSMUSG00000087516 | MGI:3780472 | Tbx3os1 | T-box 3, opposite strand 1 [Source:MGI Symbol;Acc:MGI:3780472] |
| 5 | lncRNA | 1.20E+08 | 1.20E+08 | ENSMUSG00000085708 | MGI:3801955 | Gm16063 | predicted gene 16063 [Source:MGI Symbol;Acc:MGI:3801955] |
| 5 | lncRNA | 1.20E+08 | 1.20E+08 | ENSMUSG00000140110 | MGI:7829040 | Gm73141 | predicted gene, 73141 [Source:MGI Symbol;Acc:MGI:7829040] |
| 5 | lncRNA | 1.20E+08 | 1.20E+08 | ENSMUSG00000086847 | MGI:3801954 | Tbx3os2 | T-box 3, opposite strand 2 [Source:MGI Symbol;Acc:MGI:3801954] |
| 5 | lncRNA | 1.20E+08 | 1.20E+08 | ENSMUSG00000127058 | MGI:7825783 | Gm71505 | predicted gene, 71505 [Source:MGI Symbol;Acc:MGI:7825783] |
| 5 | lncRNA | 1.20E+08 | 1.20E+08 | ENSMUSG00000120759 | MGI:7338493 | Gm57290 | predicted gene, 57290 [Source:MGI Symbol;Acc:MGI:7338493] |
| 5 | lncRNA | 1.20E+08 | 1.20E+08 | ENSMUSG00000106795 | MGI:5663187 | Gm43050 | predicted gene 43050 [Source:MGI Symbol;Acc:MGI:5663187] |
| 5 | lncRNA | 1.20E+08 | 1.20E+08 | ENSMUSG00000132877 | MGI:7826876 | Gm72055 | predicted gene, 72055 [Source:MGI Symbol;Acc:MGI:7826876] |
| 5 | lncRNA | 1.20E+08 | 1.20E+08 | ENSMUSG00000107084 | MGI:5663406 | Gm43269 | predicted gene 43269 [Source:MGI Symbol;Acc:MGI:5663406] |
| 5 | lncRNA | 1.20E+08 | 1.20E+08 | ENSMUSG00000122811 | MGI:5590932 | Gm31773 | predicted gene, 31773 [Source:MGI Symbol;Acc:MGI:5590932] |
| 5 | lncRNA | 1.20E+08 | 1.20E+08 | ENSMUSG00000107063 | MGI:5621439 | Gm38554 | predicted gene, 38554 [Source:MGI Symbol;Acc:MGI:5621439] |

| | | | | | | | |
|---|---|---|---|---|---|---|---|
| 5 | lncRNA | 1.20E+08 | 1.20E+08 | ENSMUSG00000125535 | MGI:7825631 | Gm71429 | predicted gene, 71429 [Source:MGI Symbol;Acc:MGI:7825631] |
| 5 | lncRNA | 1.20E+08 | 1.20E+08 | ENSMUSG00000132866 | MGI:7826872 | Gm72053 | predicted gene, 72053 [Source:MGI Symbol;Acc:MGI:7826872] |
| 5 | lncRNA | 1.20E+08 | 1.20E+08 | ENSMUSG00000097368 | MGI:3708725 | Gm10390 | predicted gene 10390 [Source:MGI Symbol;Acc:MGI:3708725] |
| 5 | lncRNA | 1.20E+08 | 1.20E+08 | ENSMUSG00000122430 | MGI:7825265 | Gm71242 | predicted gene, 71242 [Source:MGI Symbol;Acc:MGI:7825265] |
| 5 | lncRNA | 1.20E+08 | 1.20E+08 | ENSMUSG00000122511 | MGI:7825271 | Gm71245 | predicted gene, 71245 [Source:MGI Symbol;Acc:MGI:7825271] |
| 5 | lncRNA | 1.20E+08 | 1.20E+08 | ENSMUSG00000135011 | MGI:7827568 | Gm72401 | predicted gene, 72401 [Source:MGI Symbol;Acc:MGI:7827568] |
| 5 | lncRNA | 1.20E+08 | 1.20E+08 | ENSMUSG00000140328 | MGI:7829102 | Gm73172 | predicted gene, 73172 [Source:MGI Symbol;Acc:MGI:7829102] |
| 5 | lncRNA | 1.20E+08 | 1.20E+08 | ENSMUSG00000136990 | MGI:7828250 | Gm72744 | predicted gene, 72744 [Source:MGI Symbol;Acc:MGI:7828250] |
| 5 | lncRNA | 1.20E+08 | 1.21E+08 | ENSMUSG00000107141 | MGI:5662791 | Gm42654 | predicted gene 42654 [Source:MGI Symbol;Acc:MGI:5662791] |
| 5 | lncRNA | 1.21E+08 | 1.21E+08 | ENSMUSG00000098649 | MGI:5521042 | Lhx5as1 | LIM homeobox 5, antisense 1 [Source:MGI Symbol;Acc:MGI:5521042] |
| 5 | lncRNA | 1.21E+08 | 1.21E+08 | ENSMUSG00000121101 | MGI:7338535 | Gm57311 | predicted gene, 57311 [Source:MGI Symbol;Acc:MGI:7338535] |
| 5 | lncRNA | 1.21E+08 | 1.21E+08 | ENSMUSG00000086165 | MGI:3783131 | Gm15690 | predicted gene 15690 [Source:MGI Symbol;Acc:MGI:3783131] |
| 5 | lncRNA | 1.21E+08 | 1.21E+08 | ENSMUSG00000142125 | MGI:7829516 | Gm73379 | predicted gene, 73379 [Source:MGI Symbol;Acc:MGI:7829516] |
| 5 | lncRNA | 1.21E+08 | 1.21E+08 | ENSMUSG00000107191 | MGI:5663716 | Gm43579 | predicted gene 43579 [Source:MGI Symbol;Acc:MGI:5663716] |
| 5 | lncRNA | 1.21E+08 | 1.21E+08 | ENSMUSG00000134485 | MGI:7827452 | Gm72343 | predicted gene, 72343 [Source:MGI Symbol;Acc:MGI:7827452] |
| 5 | lncRNA | 1.21E+08 | 1.21E+08 | ENSMUSG00000089830 | MGI:3783191 | Gm15749 | predicted gene 15749 [Source:MGI Symbol;Acc:MGI:3783191] |
| 5 | lncRNA | 1.21E+08 | 1.21E+08 | ENSMUSG00000128151 | MGI:7825859 | Gm71543 | predicted gene, 71543 [Source:MGI Symbol;Acc:MGI:7825859] |
| 5 | lncRNA | 1.21E+08 | 1.21E+08 | ENSMUSG00000142087 | MGI:7829494 | Gm73368 | predicted gene, 73368 [Source:MGI Symbol;Acc:MGI:7829494] |

| | | | | | | | |
|---|---|---|---|---|---|---|---|
| 5 | lncRNA | 1.21E+08 | 1.21E+08 | ENSMUSG00000107179 | MGI:5663206 | Gm43069 | predicted gene 43069 [Source:MGI Symbol;Acc:MGI:5663206] |
| 5 | lncRNA | 1.21E+08 | 1.21E+08 | ENSMUSG00000141431 | MGI:5591344 | Gm32185 | predicted gene, 32185 [Source:MGI Symbol;Acc:MGI:5591344] |
| 5 | lncRNA | 1.21E+08 | 1.21E+08 | ENSMUSG00000128029 | MGI:7825849 | Gm71538 | predicted gene, 71538 [Source:MGI Symbol;Acc:MGI:7825849] |
| 5 | lncRNA | 1.21E+08 | 1.21E+08 | ENSMUSG00000143877 | MGI:7830018 | Gm73631 | predicted gene, 73631 [Source:MGI Symbol;Acc:MGI:7830018] |
| 5 | lncRNA | 1.21E+08 | 1.21E+08 | ENSMUSG00000126312 | MGI:7825711 | Gm71469 | predicted gene, 71469 [Source:MGI Symbol;Acc:MGI:7825711] |
| 5 | lncRNA | 1.21E+08 | 1.21E+08 | ENSMUSG00000126344 | MGI:7825717 | Gm71472 | predicted gene, 71472 [Source:MGI Symbol;Acc:MGI:7825717] |
| 5 | lncRNA | 1.21E+08 | 1.21E+08 | ENSMUSG00000142544 | MGI:7829620 | Gm73432 | predicted gene, 73432 [Source:MGI Symbol;Acc:MGI:7829620] |
| 5 | lncRNA | 1.21E+08 | 1.21E+08 | ENSMUSG00000120570 | MGI:7338555 | Gm57321 | predicted gene, 57321 [Source:MGI Symbol;Acc:MGI:7338555] |
| 5 | lncRNA | 1.22E+08 | 1.22E+08 | ENSMUSG00000104985 | MGI:5663054 | Gm42917 | predicted gene 42917 [Source:MGI Symbol;Acc:MGI:5663054] |
| 5 | lncRNA | 1.22E+08 | 1.22E+08 | ENSMUSG00000106209 | MGI:5663055 | Gm42918 | predicted gene 42918 [Source:MGI Symbol;Acc:MGI:5663055] |
| 5 | lncRNA | 1.22E+08 | 1.22E+08 | ENSMUSG00000131095 | MGI:7826388 | Gm71811 | predicted gene, 71811 [Source:MGI Symbol;Acc:MGI:7826388] |
| 5 | lncRNA | 1.22E+08 | 1.22E+08 | ENSMUSG00000126057 | MGI:7825677 | Gm71452 | predicted gene, 71452 [Source:MGI Symbol;Acc:MGI:7825677] |
| 5 | lncRNA | 1.22E+08 | 1.22E+08 | ENSMUSG00000120376 | MGI:7338527 | Gm57307 | predicted gene, 57307 [Source:MGI Symbol;Acc:MGI:7338527] |
| 5 | lncRNA | 1.22E+08 | 1.22E+08 | ENSMUSG00000130280 | MGI:7826188 | Gm71710 | predicted gene, 71710 [Source:MGI Symbol;Acc:MGI:7826188] |
| 5 | lncRNA | 1.22E+08 | 1.22E+08 | ENSMUSG00000130243 | MGI:7826168 | Gm71700 | predicted gene, 71700 [Source:MGI Symbol;Acc:MGI:7826168] |
| 5 | lncRNA | 1.22E+08 | 1.22E+08 | ENSMUSG00000120053 | MGI:1925708 | 1700064N11Rik | RIKEN cDNA 1700064N11 gene [Source:MGI Symbol;Acc:MGI:1925708] |
| 5 | lncRNA | 1.22E+08 | 1.22E+08 | ENSMUSG00000089729 | MGI:4414972 | Gm16552 | predicted gene 16552 [Source:MGI Symbol;Acc:MGI:4414972] |
| 5 | lncRNA | 1.22E+08 | 1.22E+08 | ENSMUSG00000122020 | MGI:7825187 | Gm71203 | predicted gene, 71203 [Source:MGI Symbol;Acc:MGI:7825187] |

| | | | | | | | |
|---|---|---|---|---|---|---|---|
| 5 | lncRNA | 1.22E+08 | 1.22E+08 | ENSMUSG00000104872 | MGI:1913571 | 1700008B11Rik | RIKEN cDNA 1700008B11 gene [Source:MGI Symbol;Acc:MGI:1913571] |
| 5 | lncRNA | 1.22E+08 | 1.22E+08 | ENSMUSG00000125798 | MGI:7825647 | Gm71437 | predicted gene, 71437 [Source:MGI Symbol;Acc:MGI:7825647] |
| 5 | lncRNA | 1.22E+08 | 1.22E+08 | ENSMUSG00000140369 | MGI:7829106 | Gm73174 | predicted gene, 73174 [Source:MGI Symbol;Acc:MGI:7829106] |
| 5 | lncRNA | 1.22E+08 | 1.22E+08 | ENSMUSG00000140332 | MGI:5591581 | Gm32422 | predicted gene, 32422 [Source:MGI Symbol;Acc:MGI:5591581] |
| 5 | lncRNA | 1.22E+08 | 1.22E+08 | ENSMUSG00000131264 | MGI:7826460 | Gm71847 | predicted gene, 71847 [Source:MGI Symbol;Acc:MGI:7826460] |
| 5 | lncRNA | 1.22E+08 | 1.22E+08 | ENSMUSG00000125571 | MGI:7825637 | Gm71432 | predicted gene, 71432 [Source:MGI Symbol;Acc:MGI:7825637] |
| 5 | lncRNA | 1.22E+08 | 1.22E+08 | ENSMUSG00000105795 | MGI:3782144 | Gm3970 | predicted gene 3970 [Source:MGI Symbol;Acc:MGI:3782144] |
| 5 | lncRNA | 1.22E+08 | 1.22E+08 | ENSMUSG00000087386 | MGI:3783081 | Gm15637 | predicted gene 15637 [Source:MGI Symbol;Acc:MGI:3783081] |
| 5 | lncRNA | 1.22E+08 | 1.22E+08 | ENSMUSG00000138489 | MGI:6367183 | Gm52786 | predicted gene, 52786 [Source:MGI Symbol;Acc:MGI:6367183] |
| 5 | lncRNA | 1.22E+08 | 1.22E+08 | ENSMUSG00000123198 | MGI:7825373 | Gm71298 | predicted gene, 71298 [Source:MGI Symbol;Acc:MGI:7825373] |
| 5 | lncRNA | 1.22E+08 | 1.22E+08 | ENSMUSG00000120313 | MGI:7338479 | Gm57283 | predicted gene, 57283 [Source:MGI Symbol;Acc:MGI:7338479] |
| 5 | lncRNA | 1.22E+08 | 1.22E+08 | ENSMUSG00000123162 | MGI:7825362 | Gm71292 | predicted gene, 71292 [Source:MGI Symbol;Acc:MGI:7825362] |
| 5 | lncRNA | 1.22E+08 | 1.22E+08 | ENSMUSG00000123828 | MGI:7825451 | Gm71338 | predicted gene, 71338 [Source:MGI Symbol;Acc:MGI:7825451] |
| 5 | lncRNA | 1.22E+08 | 1.22E+08 | ENSMUSG00000123867 | MGI:7825453 | Gm71339 | predicted gene, 71339 [Source:MGI Symbol;Acc:MGI:7825453] |
| 5 | lncRNA | 1.22E+08 | 1.22E+08 | ENSMUSG00000143197 | MGI:7829746 | Gm73495 | predicted gene, 73495 [Source:MGI Symbol;Acc:MGI:7829746] |
| 5 | lncRNA | 1.22E+08 | 1.22E+08 | ENSMUSG00000134979 | MGI:7827560 | Gm72397 | predicted gene, 72397 [Source:MGI Symbol;Acc:MGI:7827560] |
| 5 | lncRNA | 1.22E+08 | 1.22E+08 | ENSMUSG00000125496 | MGI:1920843 | 1700112N08Rik | RIKEN cDNA 1700112N08 gene [Source:MGI Symbol;Acc:MGI:1920843] |
| 5 | lncRNA | 1.22E+08 | 1.22E+08 | ENSMUSG00000125538 | MGI:7825635 | Gm71431 | predicted gene, 71431 [Source:MGI Symbol;Acc:MGI:7825635] |

| | | | | | | | |
|---|---|---|---|---|---|---|---|
| 5 | lncRNA | 1.22E+08 | 1.22E+08 | ENSMUSG00000134697 | MGI:7827492 | Gm72363 | predicted gene, 72363 [Source:MGI Symbol;Acc:MGI:7827492] |
| 5 | lncRNA | 1.23E+08 | 1.23E+08 | ENSMUSG00000122024 | MGI:7825189 | Gm71204 | predicted gene, 71204 [Source:MGI Symbol;Acc:MGI:7825189] |
| 5 | lncRNA | 1.23E+08 | 1.23E+08 | ENSMUSG00000120495 | MGI:7338443 | Gm57265 | predicted gene, 57265 [Source:MGI Symbol;Acc:MGI:7338443] |
| 5 | lncRNA | 1.23E+08 | 1.23E+08 | ENSMUSG00000105895 | MGI:5662966 | Gm42829 | predicted gene 42829 [Source:MGI Symbol;Acc:MGI:5662966] |
| 5 | lncRNA | 1.23E+08 | 1.23E+08 | ENSMUSG00000133548 | MGI:7827156 | Gm72195 | predicted gene, 72195 [Source:MGI Symbol;Acc:MGI:7827156] |
| 5 | lncRNA | 1.23E+08 | 1.23E+08 | ENSMUSG00000141355 | MGI:7829382 | Gm73312 | predicted gene, 73312 [Source:MGI Symbol;Acc:MGI:7829382] |
| 5 | lncRNA | 1.23E+08 | 1.23E+08 | ENSMUSG00000120573 | MGI:7338495 | Gm57291 | predicted gene, 57291 [Source:MGI Symbol;Acc:MGI:7338495] |
| 5 | lncRNA | 1.23E+08 | 1.23E+08 | ENSMUSG00000130488 | MGI:1920518 | 1700040P09Rik | RIKEN cDNA 1700040P09 gene [Source:MGI Symbol;Acc:MGI:1920518] |
| 5 | lncRNA | 1.23E+08 | 1.23E+08 | ENSMUSG00000105430 | MGI:5663438 | Gm43301 | predicted gene 43301 [Source:MGI Symbol;Acc:MGI:5663438] |
| 5 | lncRNA | 1.23E+08 | 1.23E+08 | ENSMUSG00000130609 | MGI:7826250 | Gm71741 | predicted gene, 71741 [Source:MGI Symbol;Acc:MGI:7826250] |
| 5 | lncRNA | 1.23E+08 | 1.23E+08 | ENSMUSG00000132189 | MGI:7826732 | Gm71983 | predicted gene, 71983 [Source:MGI Symbol;Acc:MGI:7826732] |
| 5 | lncRNA | 1.23E+08 | 1.23E+08 | ENSMUSG00000141260 | MGI:5592147 | Gm32988 | predicted gene, 32988 [Source:MGI Symbol;Acc:MGI:5592147] |
| 5 | lncRNA | 1.23E+08 | 1.23E+08 | ENSMUSG00000129930 | MGI:7826088 | Gm71659 | predicted gene, 71659 [Source:MGI Symbol;Acc:MGI:7826088] |
| 5 | lncRNA | 1.23E+08 | 1.23E+08 | ENSMUSG00000127177 | MGI:7825795 | Gm71511 | predicted gene, 71511 [Source:MGI Symbol;Acc:MGI:7825795] |
| 5 | lncRNA | 1.23E+08 | 1.23E+08 | ENSMUSG00000122493 | MGI:7825269 | Gm71244 | predicted gene, 71244 [Source:MGI Symbol;Acc:MGI:7825269] |
| 5 | lncRNA | 1.23E+08 | 1.23E+08 | ENSMUSG00000140346 | MGI:7829104 | Gm73173 | predicted gene, 73173 [Source:MGI Symbol;Acc:MGI:7829104] |
| 5 | lncRNA | 1.23E+08 | 1.23E+08 | ENSMUSG00000136699 | MGI:7828106 | Gm72672 | predicted gene, 72672 [Source:MGI Symbol;Acc:MGI:7828106] |
| 5 | lncRNA | 1.23E+08 | 1.23E+08 | ENSMUSG00000120987 | MGI:5592203 | Gm33044 | predicted gene, 33044 [Source:MGI Symbol;Acc:MGI:5592203] |

| | | | | | | | |
|---|---|---|---|---|---|---|---|
| 5 | lncRNA | 1.23E+08 | 1.23E+08 | ENSMUSG00000142508 | MGI:5592277 | Gm33118 | predicted gene, 33118 [Source:MGI Symbol;Acc:MGI:5592277] |
| 5 | lncRNA | 1.23E+08 | 1.23E+08 | ENSMUSG00000056735 | MGI:1924414 | A930024E05Rik | RIKEN cDNA A930024E05 gene [Source:MGI Symbol;Acc:MGI:1924414] |
| 5 | lncRNA | 1.23E+08 | 1.23E+08 | ENSMUSG00000105311 | MGI:5753150 | Gm44574 | predicted gene 44574 [Source:MGI Symbol;Acc:MGI:5753150] |
| 5 | lncRNA | 1.23E+08 | 1.23E+08 | ENSMUSG00000142552 | MGI:7829626 | Gm73435 | predicted gene, 73435 [Source:MGI Symbol;Acc:MGI:7829626] |
| 5 | lncRNA | 1.23E+08 | 1.23E+08 | ENSMUSG00000122704 | MGI:7825301 | Gm71260 | predicted gene, 71260 [Source:MGI Symbol;Acc:MGI:7825301] |
| 5 | lncRNA | 1.23E+08 | 1.23E+08 | ENSMUSG00000106555 | MGI:3780646 | Gm2479 | predicted gene 2479 [Source:MGI Symbol;Acc:MGI:3780646] |
| 5 | lncRNA | 1.23E+08 | 1.23E+08 | ENSMUSG00000131184 | MGI:5624908 | Gm42023 | predicted gene, 42023 [Source:MGI Symbol;Acc:MGI:5624908] |
| 5 | lncRNA | 1.23E+08 | 1.23E+08 | ENSMUSG00000131154 | MGI:7826420 | Gm71827 | predicted gene, 71827 [Source:MGI Symbol;Acc:MGI:7826420] |
| 5 | lncRNA | 1.23E+08 | 1.23E+08 | ENSMUSG00000141270 | MGI:7829376 | Gm73309 | predicted gene, 73309 [Source:MGI Symbol;Acc:MGI:7829376] |
| 5 | lncRNA | 1.23E+08 | 1.23E+08 | ENSMUSG00000090086 | MGI:2140910 | AI480526 | expressed sequence AI480526 [Source:MGI Symbol;Acc:MGI:2140910] |
| 5 | lncRNA | 1.23E+08 | 1.23E+08 | ENSMUSG00000104156 | MGI:5611330 | Gm38102 | predicted gene, 38102 [Source:MGI Symbol;Acc:MGI:5611330] |
| 5 | lncRNA | 1.23E+08 | 1.23E+08 | ENSMUSG00000104606 | MGI:5663546 | Gm43409 | predicted gene 43409 [Source:MGI Symbol;Acc:MGI:5663546] |
| 5 | lncRNA | 1.23E+08 | 1.23E+08 | ENSMUSG00000087063 | MGI:3801821 | Gm15857 | predicted gene 15857 [Source:MGI Symbol;Acc:MGI:3801821] |
| 5 | lncRNA | 1.23E+08 | 1.23E+08 | ENSMUSG00000087336 | MGI:3801778 | Gm15860 | predicted gene 15860 [Source:MGI Symbol;Acc:MGI:3801778] |
| 5 | lncRNA | 1.24E+08 | 1.24E+08 | ENSMUSG00000104830 | MGI:1923368 | 5830487J09Rik | RIKEN cDNA 5830487J09 gene [Source:MGI Symbol;Acc:MGI:1923368] |
| 5 | lncRNA | 1.24E+08 | 1.24E+08 | ENSMUSG00000086700 | MGI:3783189 | Gm15747 | predicted gene 15747 [Source:MGI Symbol;Acc:MGI:3783189] |
| 5 | lncRNA | 1.24E+08 | 1.24E+08 | ENSMUSG00000106636 | MGI:5663950 | Gm43813 | predicted gene 43813 [Source:MGI Symbol;Acc:MGI:5663950] |
| 5 | lncRNA | 1.24E+08 | 1.24E+08 | ENSMUSG00000086753 | MGI:3783193 | Gm15751 | predicted gene 15751 [Source:MGI Symbol;Acc:MGI:3783193] |

| | | | | | | | |
|---|---|---|---|---|---|---|---|
| 5 | lncRNA | 1.24E+08 | 1.24E+08 | ENSMUSG00000135480 | MGI:7827704 | Gm72470 | predicted gene, 72470 [Source:MGI Symbol;Acc:MGI:7827704] |
| 5 | lncRNA | 1.24E+08 | 1.24E+08 | ENSMUSG00000125294 | MGI:7825607 | Gm71417 | predicted gene, 71417 [Source:MGI Symbol;Acc:MGI:7825607] |
| 5 | lncRNA | 1.24E+08 | 1.24E+08 | ENSMUSG00000142870 | MGI:7829678 | Gm73461 | predicted gene, 73461 [Source:MGI Symbol;Acc:MGI:7829678] |
| 5 | lncRNA | 1.24E+08 | 1.24E+08 | ENSMUSG00000140043 | MGI:7829032 | Gm73137 | predicted gene, 73137 [Source:MGI Symbol;Acc:MGI:7829032] |
| 5 | lncRNA | 1.24E+08 | 1.24E+08 | ENSMUSG00000142808 | MGI:7829672 | Gm73458 | predicted gene, 73458 [Source:MGI Symbol;Acc:MGI:7829672] |
| 5 | lncRNA | 1.24E+08 | 1.24E+08 | ENSMUSG00000106261 | MGI:5593245 | Gm34086 | predicted gene, 34086 [Source:MGI Symbol;Acc:MGI:5593245] |
| 5 | lncRNA | 1.24E+08 | 1.24E+08 | ENSMUSG00000128052 | MGI:7825851 | Gm71539 | predicted gene, 71539 [Source:MGI Symbol;Acc:MGI:7825851] |
| 5 | lncRNA | 1.24E+08 | 1.24E+08 | ENSMUSG00000104818 | MGI:5663798 | Gm43661 | predicted gene 43661 [Source:MGI Symbol;Acc:MGI:5663798] |
| 5 | lncRNA | 1.24E+08 | 1.24E+08 | ENSMUSG00000126658 | MGI:7825741 | Gm71484 | predicted gene, 71484 [Source:MGI Symbol;Acc:MGI:7825741] |
| 5 | lncRNA | 1.24E+08 | 1.24E+08 | ENSMUSG00000128265 | MGI:7825867 | Gm71547 | predicted gene, 71547 [Source:MGI Symbol;Acc:MGI:7825867] |
| 5 | lncRNA | 1.24E+08 | 1.24E+08 | ENSMUSG00000085433 | MGI:3802119 | Gm16001 | predicted gene 16001 [Source:MGI Symbol;Acc:MGI:3802119] |
| 5 | lncRNA | 1.24E+08 | 1.24E+08 | ENSMUSG00000132124 | MGI:7826716 | Gm71975 | predicted gene, 71975 [Source:MGI Symbol;Acc:MGI:7826716] |
| 5 | lncRNA | 1.24E+08 | 1.24E+08 | ENSMUSG00000089842 | MGI:3840147 | Pitpnm2os2 | phosphatidylinositol transfer protein, membrane-associated 2, opposite strand 2 [Source:MGI Symbol;Acc:MGI:3840147] |
| 5 | lncRNA | 1.24E+08 | 1.24E+08 | ENSMUSG00000133564 | MGI:7827160 | Gm72197 | predicted gene, 72197 [Source:MGI Symbol;Acc:MGI:7827160] |
| 5 | lncRNA | 1.24E+08 | 1.24E+08 | ENSMUSG00000090220 | MGI:1923177 | Pitpnm2os1 | phosphatidylinositol transfer protein, membrane-associated 2, opposite strand 1 [Source:MGI Symbol;Acc:MGI:1923177] |
| 5 | lncRNA | 1.24E+08 | 1.24E+08 | ENSMUSG00000105759 | MGI:5662562 | Gm42425 | predicted gene 42425 [Source:MGI Symbol;Acc:MGI:5662562] |
| 5 | lncRNA | 1.25E+08 | 1.25E+08 | ENSMUSG00000126538 | MGI:7831160 | Gm74211 | predicted gene, 74211 [Source:MGI Symbol;Acc:MGI:7831160] |

| | | | | | | | |
|---|---|---|---|---|---|---|---|
| 5 | lncRNA | 1.25E+08 | 1.25E+08 | ENSMUSG00000136903 | MGI:7828228 | Gm72733 | predicted gene, 72733 [Source:MGI Symbol;Acc:MGI:7828228] |
| 5 | lncRNA | 1.25E+08 | 1.25E+08 | ENSMUSG00000144115 | MGI:7830126 | Gm73688 | predicted gene, 73688 [Source:MGI Symbol;Acc:MGI:7830126] |
| 5 | lncRNA | 1.25E+08 | 1.25E+08 | ENSMUSG00000144077 | MGI:7830124 | Gm73687 | predicted gene, 73687 [Source:MGI Symbol;Acc:MGI:7830124] |
| 5 | lncRNA | 1.25E+08 | 1.25E+08 | ENSMUSG00000144148 | MGI:6367189 | Gm52789 | predicted gene, 52789 [Source:MGI Symbol;Acc:MGI:6367189] |
| 5 | lncRNA | 1.25E+08 | 1.25E+08 | ENSMUSG00000144043 | MGI:7830116 | Gm73683 | predicted gene, 73683 [Source:MGI Symbol;Acc:MGI:7830116] |
| 5 | lncRNA | 1.25E+08 | 1.25E+08 | ENSMUSG00000144003 | MGI:7830082 | Gm73664 | predicted gene, 73664 [Source:MGI Symbol;Acc:MGI:7830082] |
| 5 | lncRNA | 1.25E+08 | 1.25E+08 | ENSMUSG00000106082 | MGI:3605803 | 4930404A12Rik | RIKEN cDNA 4930404A12 gene [Source:MGI Symbol;Acc:MGI:3605803] |
| 5 | lncRNA | 1.25E+08 | 1.25E+08 | ENSMUSG00000105583 | MGI:5663914 | Gm43777 | predicted gene 43777 [Source:MGI Symbol;Acc:MGI:5663914] |
| 5 | lncRNA | 1.25E+08 | 1.25E+08 | ENSMUSG00000125796 | MGI:5623206 | Gm40321 | predicted gene, 40321 [Source:MGI Symbol;Acc:MGI:5623206] |
| 5 | lncRNA | 1.25E+08 | 1.25E+08 | ENSMUSG00000126937 | MGI:5591688 | Gm32529 | predicted gene, 32529 [Source:MGI Symbol;Acc:MGI:5591688] |
| 5 | lncRNA | 1.25E+08 | 1.25E+08 | ENSMUSG00000105604 | MGI:5662975 | Gm42838 | predicted gene 42838 [Source:MGI Symbol;Acc:MGI:5662975] |
| 5 | lncRNA | 1.25E+08 | 1.25E+08 | ENSMUSG00000106663 | MGI:5662976 | Gm42839 | predicted gene 42839 [Source:MGI Symbol;Acc:MGI:5662976] |
| 5 | lncRNA | 1.25E+08 | 1.25E+08 | ENSMUSG00000138261 | MGI:7828552 | Gm72896 | predicted gene, 72896 [Source:MGI Symbol;Acc:MGI:7828552] |
| 5 | lncRNA | 1.25E+08 | 1.25E+08 | ENSMUSG00000105039 | MGI:5591744 | Gm32585 | predicted gene, 32585 [Source:MGI Symbol;Acc:MGI:5591744] |
| 5 | lncRNA | 1.25E+08 | 1.25E+08 | ENSMUSG00000131642 | MGI:7826598 | Gm71916 | predicted gene, 71916 [Source:MGI Symbol;Acc:MGI:7826598] |
| 5 | lncRNA | 1.25E+08 | 1.25E+08 | ENSMUSG00000131568 | MGI:7826572 | Gm71903 | predicted gene, 71903 [Source:MGI Symbol;Acc:MGI:7826572] |
| 5 | lncRNA | 1.25E+08 | 1.25E+08 | ENSMUSG00000131271 | MGI:5591860 | Gm32701 | predicted gene, 32701 [Source:MGI Symbol;Acc:MGI:5591860] |
| 5 | lncRNA | 1.25E+08 | 1.25E+08 | ENSMUSG00000131428 | MGI:7831178 | Gm74220 | predicted gene, 74220 [Source:MGI Symbol;Acc:MGI:7831178] |

| | | | | | | | |
|---|---|---|---|---|---|---|---|
| 5 | lncRNA | 1.25E+08 | 1.25E+08 | ENSMUSG00000131347 | MGI:7826504 | Gm71869 | predicted gene, 71869 [Source:MGI Symbol;Acc:MGI:7826504] |
| 5 | lncRNA | 1.25E+08 | 1.25E+08 | ENSMUSG00000131305 | MGI:7826496 | Gm71865 | predicted gene, 71865 [Source:MGI Symbol;Acc:MGI:7826496] |
| 5 | lncRNA | 1.25E+08 | 1.25E+08 | ENSMUSG00000131385 | MGI:7826506 | Gm71870 | predicted gene, 71870 [Source:MGI Symbol;Acc:MGI:7826506] |
| 5 | lncRNA | 1.25E+08 | 1.25E+08 | ENSMUSG00000104932 | MGI:5623208 | Gm40323 | predicted gene, 40323 [Source:MGI Symbol;Acc:MGI:5623208] |
| 5 | lncRNA | 1.25E+08 | 1.25E+08 | ENSMUSG00000131467 | MGI:7826510 | Gm71872 | predicted gene, 71872 [Source:MGI Symbol;Acc:MGI:7826510] |
| 5 | lncRNA | 1.25E+08 | 1.25E+08 | ENSMUSG00000072612 | MGI:3647829 | Gm10382 | predicted gene 10382 [Source:MGI Symbol;Acc:MGI:3647829] |
| 5 | lncRNA | 1.25E+08 | 1.25E+08 | ENSMUSG00000131682 | MGI:7826606 | Gm71920 | predicted gene, 71920 [Source:MGI Symbol;Acc:MGI:7826606] |
| 5 | lncRNA | 1.25E+08 | 1.25E+08 | ENSMUSG00000120357 | MGI:7338455 | Gm57271 | predicted gene, 57271 [Source:MGI Symbol;Acc:MGI:7338455] |
| 5 | lncRNA | 1.25E+08 | 1.25E+08 | ENSMUSG00000120435 | MGI:7338463 | Gm57275 | predicted gene, 57275 [Source:MGI Symbol;Acc:MGI:7338463] |
| 5 | lncRNA | 1.25E+08 | 1.25E+08 | ENSMUSG00000136838 | MGI:7828190 | Gm72714 | predicted gene, 72714 [Source:MGI Symbol;Acc:MGI:7828190] |
| 5 | lncRNA | 1.25E+08 | 1.25E+08 | ENSMUSG00000132160 | MGI:7826724 | Gm71979 | predicted gene, 71979 [Source:MGI Symbol;Acc:MGI:7826724] |
| 5 | lncRNA | 1.26E+08 | 1.26E+08 | ENSMUSG00000130692 | MGI:7826260 | Gm71746 | predicted gene, 71746 [Source:MGI Symbol;Acc:MGI:7826260] |
| 5 | lncRNA | 1.26E+08 | 1.26E+08 | ENSMUSG00000130651 | MGI:7830302 | Gm73778 | predicted gene, 73778 [Source:MGI Symbol;Acc:MGI:7830302] |
| 5 | lncRNA | 1.26E+08 | 1.26E+08 | ENSMUSG00000130725 | MGI:7826262 | Gm71747 | predicted gene, 71747 [Source:MGI Symbol;Acc:MGI:7826262] |
| 5 | lncRNA | 1.26E+08 | 1.26E+08 | ENSMUSG00000122318 | MGI:5623210 | Gm40325 | predicted gene, 40325 [Source:MGI Symbol;Acc:MGI:5623210] |
| 5 | lncRNA | 1.26E+08 | 1.26E+08 | ENSMUSG00000133223 | MGI:7827072 | Gm72153 | predicted gene, 72153 [Source:MGI Symbol;Acc:MGI:7827072] |
| 5 | lncRNA | 1.26E+08 | 1.26E+08 | ENSMUSG00000133268 | MGI:7827090 | Gm72162 | predicted gene, 72162 [Source:MGI Symbol;Acc:MGI:7827090] |
| 5 | lncRNA | 1.26E+08 | 1.26E+08 | ENSMUSG00000131744 | MGI:7826616 | Gm71925 | predicted gene, 71925 [Source:MGI Symbol;Acc:MGI:7826616] |

| | | | | | | | |
|---|---|---|---|---|---|---|---|
| 5 | lncRNA | 1.26E+08 | 1.26E+08 | ENSMUSG00000129713 | MGI:7826019 | Gm71624 | predicted gene, 71624 [Source:MGI Symbol;Acc:MGI:7826019] |
| 5 | lncRNA | 1.26E+08 | 1.26E+08 | ENSMUSG00000129756 | MGI:5623212 | Gm40327 | predicted gene, 40327 [Source:MGI Symbol;Acc:MGI:5623212] |
| 5 | lncRNA | 1.26E+08 | 1.26E+08 | ENSMUSG00000129792 | MGI:7826031 | Gm71630 | predicted gene, 71630 [Source:MGI Symbol;Acc:MGI:7826031] |
| 5 | lncRNA | 1.26E+08 | 1.26E+08 | ENSMUSG00000129826 | MGI:7830304 | Gm73779 | predicted gene, 73779 [Source:MGI Symbol;Acc:MGI:7830304] |
| 5 | lncRNA | 1.26E+08 | 1.26E+08 | ENSMUSG00000142010 | MGI:5623213 | Gm40328 | predicted gene, 40328 [Source:MGI Symbol;Acc:MGI:5623213] |
| 5 | lncRNA | 1.26E+08 | 1.26E+08 | ENSMUSG00000131107 | MGI:7826394 | Gm71814 | predicted gene, 71814 [Source:MGI Symbol;Acc:MGI:7826394] |
| 5 | miRNA | 42048685 | 42048766 | ENSMUSG00000098264 | MGI:5530900 | Mir6414 | microRNA 6414 [Source:MGI Symbol;Acc:MGI:5530900] |
| 5 | miRNA | 48381284 | 48381393 | ENSMUSG00000065603 | MGI:3618751 | Mir218-1 | microRNA 218-1 [Source:MGI Symbol;Acc:MGI:3618751] |
| 5 | miRNA | 50410104 | 50410210 | ENSMUSG00000099124 | MGI:5530705 | Mir8117 | microRNA 8117 [Source:MGI Symbol;Acc:MGI:5530705] |
| 5 | miRNA | 51380930 | 51381016 | ENSMUSG00002076511 | MGI:6848422 | Gm55981 | predicted gene, 55981 [Source:MGI Symbol;Acc:MGI:6848422] |
| 5 | miRNA | 52293244 | 52293359 | ENSMUSG00000098726 | MGI:5530716 | Mir6417 | microRNA 6417 [Source:MGI Symbol;Acc:MGI:5530716] |
| 5 | miRNA | 64258773 | 64258850 | ENSMUSG00002076449 | MGI:6848597 | Gm56069 | predicted gene, 56069 [Source:MGI Symbol;Acc:MGI:6848597] |
| 5 | miRNA | 64264750 | 64264845 | ENSMUSG00002076429 | MGI:6848599 | Gm56070 | predicted gene, 56070 [Source:MGI Symbol;Acc:MGI:6848599] |
| 5 | miRNA | 64804976 | 64805092 | ENSMUSG00002074850 | MGI:6848635 | Gm56088 | predicted gene, 56088 [Source:MGI Symbol;Acc:MGI:6848635] |
| 5 | miRNA | 65127661 | 65127738 | ENSMUSG00000077042 | MGI:3718549 | Mir574 | microRNA 574 [Source:MGI Symbol;Acc:MGI:3718549] |
| 5 | miRNA | 66146388 | 66146461 | ENSMUSG00002076702 | MGI:6848728 | Gm56135 | predicted gene, 56135 [Source:MGI Symbol;Acc:MGI:6848728] |
| 5 | miRNA | 67211022 | 67211135 | ENSMUSG00002075048 | MGI:6848788 | Gm56165 | predicted gene, 56165 [Source:MGI Symbol;Acc:MGI:6848788] |
| 5 | miRNA | 67789168 | 67789262 | ENSMUSG00002076306 | MGI:6846111 | Gm54817 | predicted gene, 54817 [Source:MGI Symbol;Acc:MGI:6846111] |

| | | | | | | | |
|---|---|---|---|---|---|---|---|
| 5 | miRNA | 69498126 | 69498231 | ENSMUSG00002076624 | MGI:6846134 | Gm54829 | predicted gene, 54829 [Source:MGI Symbol;Acc:MGI:6846134] |
| 5 | miRNA | 75337492 | 75337565 | ENSMUSG00000099060 | MGI:5531242 | Mir7025 | microRNA 7025 [Source:MGI Symbol;Acc:MGI:5531242] |
| 5 | miRNA | 76070487 | 76070587 | ENSMUSG00002075712 | MGI:6847257 | Gm55393 | predicted gene, 55393 [Source:MGI Symbol;Acc:MGI:6847257] |
| 5 | miRNA | 77420604 | 77420685 | ENSMUSG00000093061 | MGI:4950415 | Mir5098 | microRNA 5098 [Source:MGI Symbol;Acc:MGI:4950415] |
| 5 | miRNA | 81883774 | 81883862 | ENSMUSG00002074859 | MGI:6848072 | Gm55803 | predicted gene, 55803 [Source:MGI Symbol;Acc:MGI:6848072] |
| 5 | miRNA | 82946791 | 82946912 | ENSMUSG00000080586 | MGI:3783361 | Mir1187 | microRNA 1187 [Source:MGI Symbol;Acc:MGI:3783361] |
| 5 | miRNA | 83612641 | 83612723 | ENSMUSG00002076941 | MGI:6848080 | Gm55807 | predicted gene, 55807 [Source:MGI Symbol;Acc:MGI:6848080] |
| 5 | miRNA | 87743761 | 87743852 | ENSMUSG00000106074 | MGI:4950408 | Mir3969 | microRNA 3969 [Source:MGI Symbol;Acc:MGI:4950408] |
| 5 | miRNA | 90944045 | 90944114 | ENSMUSG00002076065 | MGI:6848412 | Gm55976 | predicted gene, 55976 [Source:MGI Symbol;Acc:MGI:6848412] |
| 5 | miRNA | 92606416 | 92606515 | ENSMUSG00000099160 | MGI:5531096 | Mir6415 | microRNA 6415 [Source:MGI Symbol;Acc:MGI:5531096] |
| 5 | miRNA | 92936310 | 92936421 | ENSMUSG00000089268 | MGI:3837206 | Mir1961 | microRNA 1961 [Source:MGI Symbol;Acc:MGI:3837206] |
| 5 | miRNA | 98623415 | 98623523 | ENSMUSG00000105814 | MGI:3629657 | Mir703 | microRNA 703 [Source:MGI Symbol;Acc:MGI:3629657] |
| 5 | miRNA | 1.01E+08 | 1.01E+08 | ENSMUSG00002076564 | MGI:6849020 | Gm56281 | predicted gene, 56281 [Source:MGI Symbol;Acc:MGI:6849020] |
| 5 | miRNA | 1.04E+08 | 1.04E+08 | ENSMUSG00000093690 | MGI:5453643 | Mir5619 | microRNA 5619 [Source:MGI Symbol;Acc:MGI:5453643] |
| 5 | miRNA | 1.04E+08 | 1.04E+08 | ENSMUSG00002076521 | MGI:6846437 | Gm54981 | predicted gene, 54981 [Source:MGI Symbol;Acc:MGI:6846437] |
| 5 | miRNA | 1.07E+08 | 1.07E+08 | ENSMUSG00002075799 | MGI:6846624 | Gm55075 | predicted gene, 55075 [Source:MGI Symbol;Acc:MGI:6846624] |
| 5 | miRNA | 1.11E+08 | 1.11E+08 | ENSMUSG00000099284 | MGI:5531370 | Mir7026 | microRNA 7026 [Source:MGI Symbol;Acc:MGI:5531370] |
| 5 | miRNA | 1.11E+08 | 1.11E+08 | ENSMUSG00000076163 | MGI:3629658 | Mir701 | microRNA 701 [Source:MGI Symbol;Acc:MGI:3629658] |

| | | | | | | | |
|---|---|---|---|---|---|---|---|
| 5 | miRNA | 1.13E+08 | 1.13E+08 | ENSMUSG00000105778 | MGI:5562765 | Mir7229 | microRNA 7229 [Source:MGI Symbol;Acc:MGI:5562765] |
| 5 | miRNA | 1.13E+08 | 1.13E+08 | ENSMUSG00000104840 | MGI:5562766 | Mir7230 | microRNA 7230 [Source:MGI Symbol;Acc:MGI:5562766] |
| 5 | miRNA | 1.15E+08 | 1.15E+08 | ENSMUSG00000098731 | MGI:5531213 | Mir7027 | microRNA 7027 [Source:MGI Symbol;Acc:MGI:5531213] |
| 5 | miRNA | 1.15E+08 | 1.15E+08 | ENSMUSG00000098656 | MGI:5531056 | Mir7028 | microRNA 7028 [Source:MGI Symbol;Acc:MGI:5531056] |
| 5 | miRNA | 1.15E+08 | 1.15E+08 | ENSMUSG00000098343 | MGI:5531232 | Mir6240 | microRNA 6240 [Source:MGI Symbol;Acc:MGI:5531232] |
| 5 | miRNA | 1.16E+08 | 1.16E+08 | ENSMUSG00000099047 | MGI:5530675 | Mir7029 | microRNA 7029 [Source:MGI Symbol;Acc:MGI:5530675] |
| 5 | miRNA | 1.16E+08 | 1.16E+08 | ENSMUSG00000099254 | MGI:5530649 | Mir7030 | microRNA 7030 [Source:MGI Symbol;Acc:MGI:5530649] |
| 5 | miRNA | 1.19E+08 | 1.19E+08 | ENSMUSG00002075849 | MGI:6848605 | Gm56073 | predicted gene, 56073 [Source:MGI Symbol;Acc:MGI:6848605] |
| 5 | miRNA | 1.20E+08 | 1.20E+08 | ENSMUSG00002075229 | MGI:6848665 | Gm56103 | predicted gene, 56103 [Source:MGI Symbol;Acc:MGI:6848665] |
| 5 | miRNA | 1.21E+08 | 1.21E+08 | ENSMUSG00002076672 | MGI:6848705 | Gm56123 | predicted gene, 56123 [Source:MGI Symbol;Acc:MGI:6848705] |
| 5 | miRNA | 1.22E+08 | 1.22E+08 | ENSMUSG00000099063 | MGI:5531241 | Mir7031 | microRNA 7031 [Source:MGI Symbol;Acc:MGI:5531241] |
| 5 | miRNA | 1.23E+08 | 1.23E+08 | ENSMUSG00000099324 | MGI:5531394 | Mir8115 | microRNA 8115 [Source:MGI Symbol;Acc:MGI:5531394] |
| 5 | miRNA | 1.24E+08 | 1.24E+08 | ENSMUSG00000098951 | MGI:5530729 | Mir7647 | microRNA 7647 [Source:MGI Symbol;Acc:MGI:5530729] |
| 5 | miRNA | 1.24E+08 | 1.24E+08 | ENSMUSG00000098454 | MGI:5530691 | Mir7032 | microRNA 7032 [Source:MGI Symbol;Acc:MGI:5530691] |
| 5 | miRNA | 1.25E+08 | 1.25E+08 | ENSMUSG00002076658 | MGI:6848996 | Gm56269 | predicted gene, 56269 [Source:MGI Symbol;Acc:MGI:6848996] |
| 5 | miRNA | 1.26E+08 | 1.26E+08 | ENSMUSG00002076951 | MGI:6848100 | Gm55817 | predicted gene, 55817 [Source:MGI Symbol;Acc:MGI:6848100] |
| 5 | misc_RNA | 66483083 | 66483405 | ENSMUSG00002076874 | MGI:6848760 | Gm56151 | predicted gene, 56151 [Source:MGI Symbol;Acc:MGI:6848760] |
| 5 | misc_RNA | 79786662 | 79786956 | ENSMUSG00002075735 | MGI:6847501 | Gm55516 | predicted gene, 55516 [Source:MGI Symbol;Acc:MGI:6847501] |

| | | | | | | | |
|---|---|---|---|---|---|---|---|
| 5 | misc_RNA | 89829775 | 89830098 | ENSMUSG00002075998 | MGI:6848366 | Gm55953 | predicted gene, 55953 [Source:MGI Symbol;Acc:MGI:6848366] |
| 5 | misc_RNA | 92130564 | 92130839 | ENSMUSG00002075701 | MGI:6848420 | Gm55980 | predicted gene, 55980 [Source:MGI Symbol;Acc:MGI:6848420] |
| 5 | misc_RNA | 1.01E+08 | 1.01E+08 | ENSMUSG00002076301 | MGI:6849012 | Gm56277 | predicted gene, 56277 [Source:MGI Symbol;Acc:MGI:6849012] |
| 5 | misc_RNA | 1.01E+08 | 1.01E+08 | ENSMUSG00002076403 | MGI:6849014 | Gm56278 | predicted gene, 56278 [Source:MGI Symbol;Acc:MGI:6849014] |
| 5 | misc_RNA | 1.01E+08 | 1.01E+08 | ENSMUSG00002075475 | MGI:6849016 | Gm56279 | predicted gene, 56279 [Source:MGI Symbol;Acc:MGI:6849016] |
| 5 | misc_RNA | 1.01E+08 | 1.01E+08 | ENSMUSG00002075511 | MGI:6849018 | Gm56280 | predicted gene, 56280 [Source:MGI Symbol;Acc:MGI:6849018] |
| 5 | misc_RNA | 1.04E+08 | 1.04E+08 | ENSMUSG00002075219 | MGI:6846311 | Gm54918 | predicted gene, 54918 [Source:MGI Symbol;Acc:MGI:6846311] |
| 5 | misc_RNA | 1.10E+08 | 1.10E+08 | ENSMUSG00002075066 | MGI:6847414 | Gm55472 | predicted gene, 55472 [Source:MGI Symbol;Acc:MGI:6847414] |
| 5 | misc_RNA | 1.11E+08 | 1.11E+08 | ENSMUSG00002076336 | MGI:6847805 | Gm55669 | predicted gene, 55669 [Source:MGI Symbol;Acc:MGI:6847805] |
| 5 | misc_RNA | 1.11E+08 | 1.11E+08 | ENSMUSG00002075664 | MGI:6847807 | Gm55670 | predicted gene, 55670 [Source:MGI Symbol;Acc:MGI:6847807] |
| 5 | misc_RNA | 1.12E+08 | 1.12E+08 | ENSMUSG00002074930 | MGI:6847823 | Gm55678 | predicted gene, 55678 [Source:MGI Symbol;Acc:MGI:6847823] |
| 5 | misc_RNA | 1.12E+08 | 1.12E+08 | ENSMUSG00002076466 | MGI:6847825 | Gm55679 | predicted gene, 55679 [Source:MGI Symbol;Acc:MGI:6847825] |
| 5 | misc_RNA | 1.12E+08 | 1.12E+08 | ENSMUSG00002075414 | MGI:6847827 | Gm55680 | predicted gene, 55680 [Source:MGI Symbol;Acc:MGI:6847827] |
| 5 | misc_RNA | 1.12E+08 | 1.12E+08 | ENSMUSG00002075709 | MGI:6847837 | Gm55685 | predicted gene, 55685 [Source:MGI Symbol;Acc:MGI:6847837] |
| 5 | misc_RNA | 1.18E+08 | 1.18E+08 | ENSMUSG00002075291 | MGI:6848601 | Gm56071 | predicted gene, 56071 [Source:MGI Symbol;Acc:MGI:6848601] |
| 5 | misc_RNA | 1.24E+08 | 1.24E+08 | ENSMUSG00002075826 | MGI:6847579 | Gm55555 | predicted gene, 55555 [Source:MGI Symbol;Acc:MGI:6847579] |
| 5 | processed_pseudogene | 41499466 | 41500274 | ENSMUSG00000106895 | MGI:3645925 | Gm4754 | predicted gene 4754 [Source:MGI Symbol;Acc:MGI:3645925] |

| | | | | | | | |
|---|---|---|---|---|---|---|---|
| 5 | processed_pseudogene | 41578919 | 41579387 | ENSMUSG00000107135 | MGI:5663943 | Gm43806 | predicted gene 43806 [Source:MGI Symbol;Acc:MGI:5663943] |
| 5 | processed_pseudogene | 42010653 | 42011544 | ENSMUSG00000107259 | MGI:3645526 | Gm5298 | predicted gene 5298 [Source:MGI Symbol;Acc:MGI:3645526] |
| 5 | processed_pseudogene | 42091740 | 42093015 | ENSMUSG00000106884 | MGI:5012341 | Gm20156 | predicted gene, 20156 [Source:MGI Symbol;Acc:MGI:5012341] |
| 5 | processed_pseudogene | 42657324 | 42657757 | ENSMUSG00000106813 | MGI:3644423 | Gm7181 | predicted gene 7181 [Source:MGI Symbol;Acc:MGI:3644423] |
| 5 | processed_pseudogene | 43120747 | 43121952 | ENSMUSG00000106622 | MGI:3648967 | Gm5554 | predicted gene 5554 [Source:MGI Symbol;Acc:MGI:3648967] |
| 5 | processed_pseudogene | 44074095 | 44075856 | ENSMUSG00000093859 | MGI:3647603 | Gm7882 | predicted gene 7882 [Source:MGI Symbol;Acc:MGI:3647603] |
| 5 | processed_pseudogene | 44512203 | 44514096 | ENSMUSG00000106394 | MGI:5662835 | Gm42698 | predicted gene 42698 [Source:MGI Symbol;Acc:MGI:5662835] |
| 5 | processed_pseudogene | 44563299 | 44563935 | ENSMUSG00000104923 | MGI:3645504 | Gm5865 | predicted gene 5865 [Source:MGI Symbol;Acc:MGI:3645504] |
| 5 | processed_pseudogene | 47791844 | 47792648 | ENSMUSG00000106402 | MGI:5010009 | Gm17824 | predicted gene, 17824 [Source:MGI Symbol;Acc:MGI:5010009] |
| 5 | processed_pseudogene | 47857187 | 47857634 | ENSMUSG00000105616 | MGI:3781188 | Gm3010 | predicted gene 3010 [Source:MGI Symbol;Acc:MGI:3781188] |
| 5 | processed_pseudogene | 48132347 | 48132677 | ENSMUSG00000105994 | MGI:5663564 | Gm43427 | predicted gene 43427 [Source:MGI Symbol;Acc:MGI:5663564] |
| 5 | processed_pseudogene | 48823247 | 48824611 | ENSMUSG00000093748 | MGI:5011216 | Gm19031 | predicted gene, 19031 [Source:MGI Symbol;Acc:MGI:5011216] |

| | | | | | | | |
|---|---|---|---|---|---|---|---|
| 5 | processed_pseudogene | 49581184 | 49581496 | ENSMUSG00000093490 | MGI:5012117 | Gm19932 | predicted gene, 19932 [Source:MGI Symbol;Acc:MGI:5012117] |
| 5 | processed_pseudogene | 49709500 | 49710610 | ENSMUSG00000083477 | MGI:3648970 | Gm5555 | predicted pseudogene 5555 [Source:MGI Symbol;Acc:MGI:3648970] |
| 5 | processed_pseudogene | 50057348 | 50059644 | ENSMUSG00000105540 | MGI:3649096 | Gm7988 | predicted gene 7988 [Source:MGI Symbol;Acc:MGI:3649096] |
| 5 | processed_pseudogene | 51078684 | 51079248 | ENSMUSG00000105273 | MGI:5663152 | Gm43015 | predicted gene 43015 [Source:MGI Symbol;Acc:MGI:5663152] |
| 5 | processed_pseudogene | 51209723 | 51210203 | ENSMUSG00000106267 | MGI:5623204 | Gm40319 | predicted gene, 40319 [Source:MGI Symbol;Acc:MGI:5623204] |
| 5 | processed_pseudogene | 52739416 | 52740388 | ENSMUSG00000095078 | MGI:3645339 | Gm5866 | predicted gene 5866 [Source:MGI Symbol;Acc:MGI:3645339] |
| 5 | processed_pseudogene | 53074961 | 53076551 | ENSMUSG00000105054 | MGI:5663801 | Gm43664 | predicted gene 43664 [Source:MGI Symbol;Acc:MGI:5663801] |
| 5 | processed_pseudogene | 54251345 | 54252422 | ENSMUSG00000107105 | MGI:3801941 | Gm15820 | predicted gene 15820 [Source:MGI Symbol;Acc:MGI:3801941] |
| 5 | processed_pseudogene | 54727247 | 54735354 | ENSMUSG00000106529 | MGI:5010636 | Gm18451 | predicted gene, 18451 [Source:MGI Symbol;Acc:MGI:5010636] |
| 5 | processed_pseudogene | 54809310 | 54809463 | ENSMUSG00000105143 | MGI:3645115 | Gm8069 | predicted pseudogene 8069 [Source:MGI Symbol;Acc:MGI:3645115] |
| 5 | processed_pseudogene | 55153962 | 55154790 | ENSMUSG00000106113 | MGI:5663916 | Gm43779 | predicted gene 43779 [Source:MGI Symbol;Acc:MGI:5663916] |
| 5 | processed_pseudogene | 55740987 | 55741692 | ENSMUSG00000105384 | MGI:5663533 | Gm43396 | predicted gene 43396 [Source:MGI Symbol;Acc:MGI:5663533] |

| | | | | | | | |
|---|---|---|---|---|---|---|---|
| 5 | processed_pseudogene | 55741836 | 55742142 | ENSMUSG00000105551 | MGI:5663336 | Gm43199 | predicted gene 43199 [Source:MGI Symbol;Acc:MGI:5663336] |
| 5 | processed_pseudogene | 55781532 | 55782933 | ENSMUSG00000083902 | MGI:3801944 | Tent2-ps1 | terminal nucleotidyltransferase 2, pseudogene 1 [Source:MGI Symbol;Acc:MGI:3801944] |
| 5 | processed_pseudogene | 55879959 | 55880763 | ENSMUSG00000106335 | MGI:5012408 | Gm20223 | predicted gene, 20223 [Source:MGI Symbol;Acc:MGI:5012408] |
| 5 | processed_pseudogene | 56314680 | 56314992 | ENSMUSG00000105685 | MGI:5663452 | Gm43315 | predicted gene 43315 [Source:MGI Symbol;Acc:MGI:5663452] |
| 5 | processed_pseudogene | 57386608 | 57387957 | ENSMUSG00000104739 | MGI:3781461 | Gm3283 | predicted gene 3283 [Source:MGI Symbol;Acc:MGI:3781461] |
| 5 | processed_pseudogene | 57418614 | 57419349 | ENSMUSG00000104560 | MGI:3648794 | Gm8115 | predicted gene 8115 [Source:MGI Symbol;Acc:MGI:3648794] |
| 5 | processed_pseudogene | 57513177 | 57514620 | ENSMUSG00000105291 | MGI:5011086 | Gm18901 | predicted gene, 18901 [Source:MGI Symbol;Acc:MGI:5011086] |
| 5 | processed_pseudogene | 57617502 | 57618132 | ENSMUSG00000104892 | MGI:3644221 | Gm6615 | predicted gene 6615 [Source:MGI Symbol;Acc:MGI:3644221] |
| 5 | processed_pseudogene | 57765911 | 57766409 | ENSMUSG00000051116 | MGI:3645325 | Gm8121 | predicted pseudogene 8121 [Source:MGI Symbol;Acc:MGI:3645325] |
| 5 | processed_pseudogene | 58306965 | 58307186 | ENSMUSG00000105525 | MGI:5663113 | Gm42976 | predicted gene 42976 [Source:MGI Symbol;Acc:MGI:5663113] |
| 5 | processed_pseudogene | 59157643 | 59158336 | ENSMUSG00000106509 | MGI:5663531 | Gm43394 | predicted gene 43394 [Source:MGI Symbol;Acc:MGI:5663531] |
| 5 | processed_pseudogene | 59210942 | 59211895 | ENSMUSG00000094696 | MGI:3644251 | Gm6632 | predicted pseudogene 6632 [Source:MGI Symbol;Acc:MGI:3644251] |

| | | | | | | | |
|---|---|---|---|---|---|---|---|
| 5 | processed_pseudogene | 59669308 | 59669686 | ENSMUSG00000105038 | MGI:5663181 | Gm43044 | predicted gene 43044 [Source:MGI Symbol;Acc:MGI:5663181] |
| 5 | processed_pseudogene | 59935255 | 59935666 | ENSMUSG00000104889 | MGI:5663180 | Gm43043 | predicted gene 43043 [Source:MGI Symbol;Acc:MGI:5663180] |
| 5 | processed_pseudogene | 60182239 | 60182672 | ENSMUSG00000106250 | MGI:5663529 | Gm43392 | predicted gene 43392 [Source:MGI Symbol;Acc:MGI:5663529] |
| 5 | processed_pseudogene | 60183033 | 60183226 | ENSMUSG00000106170 | MGI:5663530 | Gm43393 | predicted gene 43393 [Source:MGI Symbol;Acc:MGI:5663530] |
| 5 | processed_pseudogene | 60333259 | 60333892 | ENSMUSG00000106042 | MGI:5662998 | Gm42861 | predicted gene 42861 [Source:MGI Symbol;Acc:MGI:5662998] |
| 5 | processed_pseudogene | 60388826 | 60389555 | ENSMUSG00000087034 | MGI:1333788 | Cbfa2t2-ps1 | CBFA2/RUNX1 translocation partner 2, pseudogene 1 [Source:MGI Symbol;Acc:MGI:1333788] |
| 5 | processed_pseudogene | 60735973 | 60736393 | ENSMUSG00000106548 | MGI:5663527 | Gm43390 | predicted gene 43390 [Source:MGI Symbol;Acc:MGI:5663527] |
| 5 | processed_pseudogene | 60831660 | 60833451 | ENSMUSG00000105486 | MGI:3781550 | Gm3372 | predicted gene 3372 [Source:MGI Symbol;Acc:MGI:3781550] |
| 5 | processed_pseudogene | 60979002 | 60979770 | ENSMUSG00000104775 | MGI:3643331 | Gm8182 | predicted gene 8182 [Source:MGI Symbol;Acc:MGI:3643331] |
| 5 | processed_pseudogene | 61495239 | 61496076 | ENSMUSG00000106370 | MGI:5663519 | Gm43382 | predicted gene 43382 [Source:MGI Symbol;Acc:MGI:5663519] |
| 5 | processed_pseudogene | 61759913 | 61760381 | ENSMUSG00000105675 | MGI:5663520 | Gm43383 | predicted gene 43383 [Source:MGI Symbol;Acc:MGI:5663520] |
| 5 | processed_pseudogene | 61874373 | 61875009 | ENSMUSG00000105573 | MGI:5663521 | Gm43384 | predicted gene 43384 [Source:MGI Symbol;Acc:MGI:5663521] |

| | | | | | | | |
|---|---|---|---|---|---|---|---|
| 5 | processed_pseudogene | 62161563 | 62162178 | ENSMUSG00000106530 | MGI:5662567 | Gm42430 | predicted gene 42430 [Source:MGI Symbol;Acc:MGI:5662567] |
| 5 | processed_pseudogene | 63851397 | 63851856 | ENSMUSG00000090138 | MGI:3781863 | Gm3687 | predicted gene 3687 [Source:MGI Symbol;Acc:MGI:3781863] |
| 5 | processed_pseudogene | 64147731 | 64148214 | ENSMUSG00000106076 | MGI:5663977 | Gm43840 | predicted gene 43840 [Source:MGI Symbol;Acc:MGI:5663977] |
| 5 | processed_pseudogene | 64347376 | 64348330 | ENSMUSG00000083435 | MGI:3801976 | Gm15824 | predicted gene 15824 [Source:MGI Symbol;Acc:MGI:3801976] |
| 5 | processed_pseudogene | 64409899 | 64410568 | ENSMUSG00000081143 | MGI:3801858 | Gm15823 | predicted gene 15823 [Source:MGI Symbol;Acc:MGI:3801858] |
| 5 | processed_pseudogene | 64477599 | 64478097 | ENSMUSG00000083100 | MGI:3648725 | Gm6044 | predicted gene 6044 [Source:MGI Symbol;Acc:MGI:3648725] |
| 5 | processed_pseudogene | 66355255 | 66358669 | ENSMUSG00000084108 | MGI:3783236 | Gm15794 | predicted gene 15794 [Source:MGI Symbol;Acc:MGI:3783236] |
| 5 | processed_pseudogene | 66514261 | 66514528 | ENSMUSG00000106910 | MGI:3826595 | Gm16273 | predicted gene 16273 [Source:MGI Symbol;Acc:MGI:3826595] |
| 5 | processed_pseudogene | 67047709 | 67048219 | ENSMUSG00000082420 | MGI:3643656 | Gm6517 | predicted gene 6517 [Source:MGI Symbol;Acc:MGI:3643656] |
| 5 | processed_pseudogene | 68145870 | 68146350 | ENSMUSG00000107257 | MGI:5663165 | Gm43028 | predicted gene 43028 [Source:MGI Symbol;Acc:MGI:5663165] |
| 5 | processed_pseudogene | 68434815 | 68435139 | ENSMUSG00000107289 | MGI:5663163 | Gm43026 | predicted gene 43026 [Source:MGI Symbol;Acc:MGI:5663163] |
| 5 | processed_pseudogene | 69618495 | 69619086 | ENSMUSG00000043484 | MGI:3644922 | Gm5867 | predicted gene 5867 [Source:MGI Symbol;Acc:MGI:3644922] |

| | | | | | | | |
|---|---|---|---|---|---|---|---|
| 5 | processed_pseudogene | 70088868 | 70089151 | ENSMUSG00000106284 | MGI:5663276 | Gm43139 | predicted gene 43139 [Source:MGI Symbol;Acc:MGI:5663276] |
| 5 | processed_pseudogene | 70710951 | 70711564 | ENSMUSG00000106613 | MGI:5662789 | Gm42652 | predicted gene 42652 [Source:MGI Symbol;Acc:MGI:5662789] |
| 5 | processed_pseudogene | 72190061 | 72190831 | ENSMUSG00000080736 | MGI:3783062 | Gm15617 | predicted gene 15617 [Source:MGI Symbol;Acc:MGI:3783062] |
| 5 | processed_pseudogene | 72204548 | 72204922 | ENSMUSG00000082705 | MGI:3783061 | Gm15616 | predicted gene 15616 [Source:MGI Symbol;Acc:MGI:3783061] |
| 5 | processed_pseudogene | 72329160 | 72329631 | ENSMUSG00000105016 | MGI:5662706 | Gm42569 | predicted gene 42569 [Source:MGI Symbol;Acc:MGI:5662706] |
| 5 | processed_pseudogene | 72566787 | 72567300 | ENSMUSG00000093644 | MGI:5011745 | Gm19560 | predicted gene, 19560 [Source:MGI Symbol;Acc:MGI:5011745] |
| 5 | processed_pseudogene | 72729517 | 72729775 | ENSMUSG00000105865 | MGI:5593303 | Gm34144 | predicted gene, 34144 [Source:MGI Symbol;Acc:MGI:5593303] |
| 5 | processed_pseudogene | 72883405 | 72883714 | ENSMUSG00000105089 | MGI:5663854 | Gm43717 | predicted gene 43717 [Source:MGI Symbol;Acc:MGI:5663854] |
| 5 | processed_pseudogene | 73554142 | 73555156 | ENSMUSG00000106958 | MGI:5595119 | Gm35960 | predicted gene, 35960 [Source:MGI Symbol;Acc:MGI:5595119] |
| 5 | processed_pseudogene | 75043996 | 75044896 | ENSMUSG00000106806 | MGI:5010091 | Gm17906 | predicted gene, 17906 [Source:MGI Symbol;Acc:MGI:5010091] |
| 5 | processed_pseudogene | 75209140 | 75210697 | ENSMUSG00000107162 | MGI:5010778 | Gm18593 | predicted gene, 18593 [Source:MGI Symbol;Acc:MGI:5010778] |
| 5 | processed_pseudogene | 75269296 | 75269584 | ENSMUSG00000106900 | MGI:3780209 | Gm2040 | predicted gene 2040 [Source:MGI Symbol;Acc:MGI:3780209] |

| | | | | | | | |
|---|---|---|---|---|---|---|---|
| 5 | processed_pseudogene | 75407104 | 75407431 | ENSMUSG00000106704 | MGI:5662936 | Gm42799 | predicted gene 42799 [Source:MGI Symbol;Acc:MGI:5662936] |
| 5 | processed_pseudogene | 76221072 | 76221423 | ENSMUSG00000107020 | MGI:3645584 | Gm6051 | predicted gene 6051 [Source:MGI Symbol;Acc:MGI:3645584] |
| 5 | processed_pseudogene | 76380459 | 76380917 | ENSMUSG00000106831 | MGI:894296 | Ube2n-ps1 | ubiquitin-conjugating enzyme E2N, pseudogene 1 [Source:MGI Symbol;Acc:MGI:894296] |
| 5 | processed_pseudogene | 76558201 | 76558352 | ENSMUSG00000107160 | MGI:5662803 | Gm42666 | predicted gene 42666 [Source:MGI Symbol;Acc:MGI:5662803] |
| 5 | processed_pseudogene | 76598548 | 76599621 | ENSMUSG00000106982 | MGI:5662802 | Gm42665 | predicted gene 42665 [Source:MGI Symbol;Acc:MGI:5662802] |
| 5 | processed_pseudogene | 76889920 | 76890360 | ENSMUSG00000083874 | MGI:3643474 | Gm7494 | predicted gene 7494 [Source:MGI Symbol;Acc:MGI:3643474] |
| 5 | processed_pseudogene | 77375225 | 77375797 | ENSMUSG00000107034 | MGI:5011245 | Gm19060 | predicted gene, 19060 [Source:MGI Symbol;Acc:MGI:5011245] |
| 5 | processed_pseudogene | 77466502 | 77466784 | ENSMUSG00000107238 | MGI:5662895 | Gm42758 | predicted gene 42758 [Source:MGI Symbol;Acc:MGI:5662895] |
| 5 | processed_pseudogene | 78056971 | 78057307 | ENSMUSG00000106960 | MGI:5662810 | Gm42673 | predicted gene 42673 [Source:MGI Symbol;Acc:MGI:5662810] |
| 5 | processed_pseudogene | 78621732 | 78622582 | ENSMUSG00000044058 | MGI:2384077 | Mthfr-ps1 | 5,10-methylenetetrahydrofolate reductase, pseudogene 1 [Source:MGI Symbol;Acc:MGI:2384077] |
| 5 | processed_pseudogene | 78688375 | 78688738 | ENSMUSG00000107309 | MGI:5663369 | Gm43232 | predicted gene 43232 [Source:MGI Symbol;Acc:MGI:5663369] |
| 5 | processed_pseudogene | 79588044 | 79588061 | ENSMUSG00000107308 | MGI:5663371 | Gm43234 | predicted gene 43234 [Source:MGI Symbol;Acc:MGI:5663371] |

| | | | | | | | |
|---|---|---|---|---|---|---|---|
| 5 | processed_pseudogene | 79655248 | 79656358 | ENSMUSG00000107183 | MGI:5010637 | Gm18452 | predicted gene, 18452 [Source:MGI Symbol;Acc:MGI:5010637] |
| 5 | processed_pseudogene | 81048597 | 81049470 | ENSMUSG00000083829 | MGI:3780369 | Gm2199 | predicted gene 2199 [Source:MGI Symbol;Acc:MGI:3780369] |
| 5 | processed_pseudogene | 82484698 | 82485496 | ENSMUSG00000106926 | MGI:3644615 | Rpl7-ps7 | ribosomal protein L7, pseudogene 7 [Source:MGI Symbol;Acc:MGI:3644615] |
| 5 | processed_pseudogene | 83188485 | 83188851 | ENSMUSG00000106606 | MGI:5663354 | Gm43217 | predicted gene 43217 [Source:MGI Symbol;Acc:MGI:5663354] |
| 5 | processed_pseudogene | 83447432 | 83448299 | ENSMUSG00000081822 | MGI:3783070 | Gm15626 | predicted gene 15626 [Source:MGI Symbol;Acc:MGI:3783070] |
| 5 | processed_pseudogene | 84766565 | 84767678 | ENSMUSG00000105658 | MGI:5663668 | Gm43531 | predicted gene 43531 [Source:MGI Symbol;Acc:MGI:5663668] |
| 5 | processed_pseudogene | 85014038 | 85014704 | ENSMUSG00000106389 | MGI:5434361 | Gm21006 | predicted gene, 21006 [Source:MGI Symbol;Acc:MGI:5434361] |
| 5 | processed_pseudogene | 86153266 | 86153476 | ENSMUSG00000106225 | MGI:5663701 | Gm43564 | predicted gene 43564 [Source:MGI Symbol;Acc:MGI:5663701] |
| 5 | processed_pseudogene | 87110235 | 87110423 | ENSMUSG00000118539 | MGI:6388938 | Gm53044 | predicted gene, 53044 [Source:MGI Symbol;Acc:MGI:6388938] |
| 5 | processed_pseudogene | 87145018 | 87146274 | ENSMUSG00000104802 | MGI:3648808 | Gm5869 | predicted gene 5869 [Source:MGI Symbol;Acc:MGI:3648808] |
| 5 | processed_pseudogene | 87162019 | 87163078 | ENSMUSG00000105315 | MGI:5010820 | Gm18635 | predicted gene, 18635 [Source:MGI Symbol;Acc:MGI:5010820] |
| 5 | processed_pseudogene | 87262393 | 87262468 | ENSMUSG00000106502 | MGI:5662932 | Gm42795 | predicted gene 42795 [Source:MGI Symbol;Acc:MGI:5662932] |

| | | | | | | | |
|---|---|---|---|---|---|---|---|
| 5 | processed_pseudogene | 87339481 | 87341032 | ENSMUSG00000106330 | MGI:5663506 | Gm43369 | predicted gene 43369 [Source:MGI Symbol;Acc:MGI:5663506] |
| 5 | processed_pseudogene | 87517662 | 87518342 | ENSMUSG00000104664 | MGI:5594729 | Idi1-ps2 | isopentenyl-diphosphate delta isomerase, pseudogene 2 [Source:MGI Symbol;Acc:MGI:5594729] |
| 5 | processed_pseudogene | 87748930 | 87749338 | ENSMUSG00000106217 | MGI:5663505 | Gm43368 | predicted gene 43368 [Source:MGI Symbol;Acc:MGI:5663505] |
| 5 | processed_pseudogene | 87751561 | 87753475 | ENSMUSG00000105246 | MGI:5434398 | Gm21043 | predicted gene, 21043 [Source:MGI Symbol;Acc:MGI:5434398] |
| 5 | processed_pseudogene | 87884965 | 87885069 | ENSMUSG00000105796 | MGI:5662982 | Gm42845 | predicted gene 42845 [Source:MGI Symbol;Acc:MGI:5662982] |
| 5 | processed_pseudogene | 88210375 | 88212232 | ENSMUSG00000105905 | MGI:3646978 | Gm7709 | predicted gene 7709 [Source:MGI Symbol;Acc:MGI:3646978] |
| 5 | processed_pseudogene | 88218805 | 88219183 | ENSMUSG00000104640 | MGI:5662760 | Gm42623 | predicted gene 42623 [Source:MGI Symbol;Acc:MGI:5662760] |
| 5 | processed_pseudogene | 88292563 | 88292917 | ENSMUSG00000105807 | MGI:5662759 | Gm42622 | predicted gene 42622 [Source:MGI Symbol;Acc:MGI:5662759] |
| 5 | processed_pseudogene | 88379209 | 88381099 | ENSMUSG00000106593 | MGI:3779657 | Gm7048 | predicted gene 7048 [Source:MGI Symbol;Acc:MGI:3779657] |
| 5 | processed_pseudogene | 88571074 | 88571413 | ENSMUSG00000106633 | MGI:5663568 | Gm43431 | predicted gene 43431 [Source:MGI Symbol;Acc:MGI:5663568] |
| 5 | processed_pseudogene | 88575672 | 88576217 | ENSMUSG00000105013 | MGI:3644419 | Gm7721 | predicted gene 7721 [Source:MGI Symbol;Acc:MGI:3644419] |
| 5 | processed_pseudogene | 88692895 | 88693015 | ENSMUSG00000106322 | MGI:5663569 | Gm43432 | predicted gene 43432 [Source:MGI Symbol;Acc:MGI:5663569] |

| | | | | | | | |
|---|---|---|---|---|---|---|---|
| 5 | processed_pseudogene | 88887106 | 88887588 | ENSMUSG00000095524 | MGI:3783124 | Gm15682 | predicted gene 15682 [Source:MGI Symbol;Acc:MGI:3783124] |
| 5 | processed_pseudogene | 88951453 | 88953082 | ENSMUSG00000106305 | MGI:5663048 | Gm42911 | predicted gene 42911 [Source:MGI Symbol;Acc:MGI:5663048] |
| 5 | processed_pseudogene | 88985869 | 88987258 | ENSMUSG00000082389 | MGI:2663191 | Mkrn1-ps1 | makorin, ring finger protein 1, pseudogene 1 [Source:MGI Symbol;Acc:MGI:2663191] |
| 5 | processed_pseudogene | 89647011 | 89647836 | ENSMUSG00000106534 | MGI:3646411 | Gm6366 | predicted gene 6366 [Source:MGI Symbol;Acc:MGI:3646411] |
| 5 | processed_pseudogene | 90280088 | 90280534 | ENSMUSG00000043618 | MGI:3643585 | Eif5al3-ps | eukaryotic translation initiation factor 5A-like 3, pseudogene [Source:MGI Symbol;Acc:MGI:3643585] |
| 5 | processed_pseudogene | 90706708 | 90707278 | ENSMUSG00000106705 | MGI:3780770 | Gm2602 | predicted gene 2602 [Source:MGI Symbol;Acc:MGI:3780770] |
| 5 | processed_pseudogene | 90898384 | 90898697 | ENSMUSG00000107072 | MGI:5663227 | Gm43090 | predicted gene 43090 [Source:MGI Symbol;Acc:MGI:5663227] |
| 5 | processed_pseudogene | 91885564 | 91885938 | ENSMUSG00000107261 | MGI:3643840 | Trmt112-ps1 | tRNA methyltransferase 11-2, pseudogene 1 [Source:MGI Symbol;Acc:MGI:3643840] |
| 5 | processed_pseudogene | 91936457 | 91937340 | ENSMUSG00000107337 | MGI:5663733 | Gm43596 | predicted gene 43596 [Source:MGI Symbol;Acc:MGI:5663733] |
| 5 | processed_pseudogene | 91954438 | 91954932 | ENSMUSG00000096726 | MGI:3644616 | Gm5558 | predicted gene 5558 [Source:MGI Symbol;Acc:MGI:3644616] |
| 5 | processed_pseudogene | 92288153 | 92288788 | ENSMUSG00000084111 | MGI:3783151 | Gm15710 | predicted gene 15710 [Source:MGI Symbol;Acc:MGI:3783151] |
| 5 | processed_pseudogene | 92511763 | 92513370 | ENSMUSG00000082068 | MGI:3780842 | Gm2673 | predicted gene 2673 [Source:MGI Symbol;Acc:MGI:3780842] |

| | | | | | | | |
|---|---|---|---|---|---|---|---|
| 5 | processed_pseudogene | 93151690 | 93152263 | ENSMUSG00000106930 | MGI:3648082 | Gm6450 | predicted gene 6450 [Source:MGI Symbol;Acc:MGI:3648082] |
| 5 | processed_pseudogene | 93343657 | 93344038 | ENSMUSG00000083700 | MGI:3781164 | Gm2986 | predicted gene 2986 [Source:MGI Symbol;Acc:MGI:3781164] |
| 5 | processed_pseudogene | 93538461 | 93540950 | ENSMUSG00000053208 | MGI:3648817 | Kat2b-ps | K(lysine) acetyltransferase 2B, pseudogene [Source:MGI Symbol;Acc:MGI:3648817] |
| 5 | processed_pseudogene | 93854333 | 93854591 | ENSMUSG00000106954 | MGI:5663363 | Gm43226 | predicted gene 43226 [Source:MGI Symbol;Acc:MGI:5663363] |
| 5 | processed_pseudogene | 96009573 | 96010574 | ENSMUSG00000096617 | MGI:3779498 | Gm5559 | predicted gene 5559 [Source:MGI Symbol;Acc:MGI:3779498] |
| 5 | processed_pseudogene | 96122099 | 96123482 | ENSMUSG00000107092 | MGI:3647399 | Gm7993 | predicted gene 7993 [Source:MGI Symbol;Acc:MGI:3647399] |
| 5 | processed_pseudogene | 96321799 | 96322209 | ENSMUSG00000090468 | MGI:3781012 | Gm2840 | predicted gene 2840 [Source:MGI Symbol;Acc:MGI:3781012] |
| 5 | processed_pseudogene | 97412687 | 97413578 | ENSMUSG00000067161 | MGI:3647980 | Gm5560 | predicted pseudogene 5560 [Source:MGI Symbol;Acc:MGI:3647980] |
| 5 | processed_pseudogene | 98204652 | 98204962 | ENSMUSG00000105117 | MGI:5662897 | Gm42760 | predicted gene 42760 [Source:MGI Symbol;Acc:MGI:5662897] |
| 5 | processed_pseudogene | 98223162 | 98223777 | ENSMUSG00000105441 | MGI:3647081 | Gm8041 | predicted gene 8041 [Source:MGI Symbol;Acc:MGI:3647081] |
| 5 | processed_pseudogene | 98301040 | 98302035 | ENSMUSG00000098073 | MGI:3648505 | Gm8048 | predicted gene 8048 [Source:MGI Symbol;Acc:MGI:3648505] |
| 5 | processed_pseudogene | 98601606 | 98601986 | ENSMUSG00000106273 | MGI:5663904 | Gm43767 | predicted gene 43767 [Source:MGI Symbol;Acc:MGI:5663904] |

| | | | | | | | |
|---|---|---|---|---|---|---|---|
| 5 | processed_pseudogene | 99169662 | 99170081 | ENSMUSG00000089659 | MGI:3781646 | Gm3470 | predicted gene 3470 [Source:MGI Symbol;Acc:MGI:3781646] |
| 5 | processed_pseudogene | 99758484 | 99759292 | ENSMUSG00000089627 | MGI:3801790 | Gm16228 | predicted gene 16228 [Source:MGI Symbol;Acc:MGI:3801790] |
| 5 | processed_pseudogene | 99987392 | 99987849 | ENSMUSG00000104731 | MGI:5663386 | Gm43249 | predicted gene 43249 [Source:MGI Symbol;Acc:MGI:5663386] |
| 5 | processed_pseudogene | 1.01E+08 | 1.01E+08 | ENSMUSG00000105181 | MGI:5011805 | Gm19620 | predicted gene, 19620 [Source:MGI Symbol;Acc:MGI:5011805] |
| 5 | processed_pseudogene | 1.01E+08 | 1.01E+08 | ENSMUSG00000105965 | MGI:109499 | Cycs-ps2 | cytochrome c, pseudogene 2 [Source:MGI Symbol;Acc:MGI:109499] |
| 5 | processed_pseudogene | 1.02E+08 | 1.02E+08 | ENSMUSG00000106610 | MGI:5663240 | Gm43103 | predicted gene 43103 [Source:MGI Symbol;Acc:MGI:5663240] |
| 5 | processed_pseudogene | 1.02E+08 | 1.02E+08 | ENSMUSG00000105088 | MGI:5010163 | Gm17978 | predicted gene, 17978 [Source:MGI Symbol;Acc:MGI:5010163] |
| 5 | processed_pseudogene | 1.02E+08 | 1.02E+08 | ENSMUSG00000106189 | MGI:5663070 | Gm42933 | predicted gene 42933 [Source:MGI Symbol;Acc:MGI:5663070] |
| 5 | processed_pseudogene | 1.03E+08 | 1.03E+08 | ENSMUSG00000106295 | MGI:5663069 | Gm42932 | predicted gene 42932 [Source:MGI Symbol;Acc:MGI:5663069] |
| 5 | processed_pseudogene | 1.03E+08 | 1.03E+08 | ENSMUSG00000105253 | MGI:5662659 | Gm42522 | predicted gene 42522 [Source:MGI Symbol;Acc:MGI:5662659] |
| 5 | processed_pseudogene | 1.03E+08 | 1.03E+08 | ENSMUSG00000104688 | MGI:5010887 | Gm18702 | predicted gene, 18702 [Source:MGI Symbol;Acc:MGI:5010887] |
| 5 | processed_pseudogene | 1.04E+08 | 1.04E+08 | ENSMUSG00000084126 | MGI:3802079 | Gm15844 | predicted gene 15844 [Source:MGI Symbol;Acc:MGI:3802079] |

| | | | | | | | |
|---|---|---|---|---|---|---|---|
| 5 | processed_pseudogene | 1.04E+08 | 1.04E+08 | ENSMUSG00000082527 | MGI:3643346 | Gm8200 | predicted gene 8200 [Source:MGI Symbol;Acc:MGI:3643346] |
| 5 | processed_pseudogene | 1.05E+08 | 1.05E+08 | ENSMUSG00000048334 | MGI:3644429 | Gm8258 | predicted gene 8258 [Source:MGI Symbol;Acc:MGI:3644429] |
| 5 | processed_pseudogene | 1.05E+08 | 1.05E+08 | ENSMUSG00000104899 | MGI:5663992 | Gm43855 | predicted gene 43855 [Source:MGI Symbol;Acc:MGI:5663992] |
| 5 | processed_pseudogene | 1.05E+08 | 1.05E+08 | ENSMUSG00000106392 | MGI:3644057 | Gm5870 | predicted pseudogene 5870 [Source:MGI Symbol;Acc:MGI:3644057] |
| 5 | processed_pseudogene | 1.06E+08 | 1.06E+08 | ENSMUSG00000067058 | MGI:3642245 | Rps15a-ps5 | ribosomal protein S15A, pseudogene 5 [Source:MGI Symbol;Acc:MGI:3642245] |
| 5 | processed_pseudogene | 1.06E+08 | 1.06E+08 | ENSMUSG00000094708 | MGI:3708724 | Gm10359 | predicted gene 10359 [Source:MGI Symbol;Acc:MGI:3708724] |
| 5 | processed_pseudogene | 1.06E+08 | 1.06E+08 | ENSMUSG00000105074 | MGI:5663955 | Gm43818 | predicted gene 43818 [Source:MGI Symbol;Acc:MGI:5663955] |
| 5 | processed_pseudogene | 1.06E+08 | 1.06E+08 | ENSMUSG00000105129 | MGI:5011751 | Gm19566 | predicted gene, 19566 [Source:MGI Symbol;Acc:MGI:5011751] |
| 5 | processed_pseudogene | 1.06E+08 | 1.06E+08 | ENSMUSG00000105600 | MGI:3644525 | Gm5987 | predicted gene 5987 [Source:MGI Symbol;Acc:MGI:3644525] |
| 5 | processed_pseudogene | 1.06E+08 | 1.06E+08 | ENSMUSG00000106669 | MGI:5621318 | Gm38433 | predicted gene, 38433 [Source:MGI Symbol;Acc:MGI:5621318] |
| 5 | processed_pseudogene | 1.07E+08 | 1.07E+08 | ENSMUSG00000104604 | MGI:5663052 | Gm42915 | predicted gene 42915 [Source:MGI Symbol;Acc:MGI:5663052] |
| 5 | processed_pseudogene | 1.07E+08 | 1.07E+08 | ENSMUSG00000106142 | MGI:5010122 | Gm17937 | predicted gene, 17937 [Source:MGI Symbol;Acc:MGI:5010122] |

| | | | | | | | |
|---|---|---|---|---|---|---|---|
| 5 | processed_pseudogene | 1.07E+08 | 1.07E+08 | ENSMUSG00000106301 | MGI:3647827 | Gm8365 | predicted gene 8365 [Source:MGI Symbol;Acc:MGI:3647827] |
| 5 | processed_pseudogene | 1.07E+08 | 1.07E+08 | ENSMUSG00000084218 | MGI:3644272 | Gm8145 | predicted gene 8145 [Source:MGI Symbol;Acc:MGI:3644272] |
| 5 | processed_pseudogene | 1.08E+08 | 1.08E+08 | ENSMUSG00000081798 | MGI:3644504 | Gm8152 | predicted gene 8152 [Source:MGI Symbol;Acc:MGI:3644504] |
| 5 | processed_pseudogene | 1.08E+08 | 1.08E+08 | ENSMUSG00000089628 | MGI:3779431 | Gm9727 | predicted gene 9727 [Source:MGI Symbol;Acc:MGI:3779431] |
| 5 | processed_pseudogene | 1.08E+08 | 1.08E+08 | ENSMUSG00000083609 | MGI:3801785 | Gm16007 | predicted gene 16007 [Source:MGI Symbol;Acc:MGI:3801785] |
| 5 | processed_pseudogene | 1.08E+08 | 1.08E+08 | ENSMUSG00000096920 | MGI:3780053 | Rps19-ps8 | ribosomal protein S19, pseudogene 8 [Source:MGI Symbol;Acc:MGI:3780053] |
| 5 | processed_pseudogene | 1.08E+08 | 1.08E+08 | ENSMUSG00000105554 | MGI:3648119 | Gm7118 | predicted gene 7118 [Source:MGI Symbol;Acc:MGI:3648119] |
| 5 | processed_pseudogene | 1.08E+08 | 1.08E+08 | ENSMUSG00000105188 | MGI:3646277 | Gm8190 | predicted gene 8190 [Source:MGI Symbol;Acc:MGI:3646277] |
| 5 | processed_pseudogene | 1.08E+08 | 1.08E+08 | ENSMUSG00000105591 | MGI:5663220 | Gm43083 | predicted gene 43083 [Source:MGI Symbol;Acc:MGI:5663220] |
| 5 | processed_pseudogene | 1.09E+08 | 1.09E+08 | ENSMUSG00000060730 | MGI:3642342 | Gm10086 | predicted pseudogene 10086 [Source:MGI Symbol;Acc:MGI:3642342] |
| 5 | processed_pseudogene | 1.09E+08 | 1.09E+08 | ENSMUSG00000093756 | MGI:5010779 | Gm18594 | predicted gene, 18594 [Source:MGI Symbol;Acc:MGI:5010779] |
| 5 | processed_pseudogene | 1.09E+08 | 1.09E+08 | ENSMUSG00000105808 | MGI:5662764 | Gm42627 | predicted gene 42627 [Source:MGI Symbol;Acc:MGI:5662764] |

| | | | | | | | |
|---|---|---|---|---|---|---|---|
| 5 | processed_pseudogene | 1.09E+08 | 1.09E+08 | ENSMUSG00000105193 | MGI:5011156 | Gm18971 | predicted gene, 18971 [Source:MGI Symbol;Acc:MGI:5011156] |
| 5 | processed_pseudogene | 1.09E+08 | 1.09E+08 | ENSMUSG00000093725 | MGI:5010780 | Gm18595 | predicted gene, 18595 [Source:MGI Symbol;Acc:MGI:5010780] |
| 5 | processed_pseudogene | 1.09E+08 | 1.09E+08 | ENSMUSG00000093545 | MGI:3644058 | Gm5871 | predicted gene 5871 [Source:MGI Symbol;Acc:MGI:3644058] |
| 5 | processed_pseudogene | 1.09E+08 | 1.09E+08 | ENSMUSG00000093543 | MGI:3647680 | Gm8473 | predicted gene 8473 [Source:MGI Symbol;Acc:MGI:3647680] |
| 5 | processed_pseudogene | 1.10E+08 | 1.10E+08 | ENSMUSG00000105303 | MGI:5010740 | Gm18555 | predicted gene, 18555 [Source:MGI Symbol;Acc:MGI:5010740] |
| 5 | processed_pseudogene | 1.10E+08 | 1.10E+08 | ENSMUSG00000093688 | MGI:3644909 | Gm8488 | predicted gene 8488 [Source:MGI Symbol;Acc:MGI:3644909] |
| 5 | processed_pseudogene | 1.10E+08 | 1.10E+08 | ENSMUSG00000094409 | MGI:3761467 | Vmn2r-ps23 | vomeronasal 2, receptor, pseudogene 23 [Source:MGI Symbol;Acc:MGI:3761467] |
| 5 | processed_pseudogene | 1.10E+08 | 1.10E+08 | ENSMUSG00000049477 | MGI:3643536 | Gm8508 | predicted gene 8508 [Source:MGI Symbol;Acc:MGI:3643536] |
| 5 | processed_pseudogene | 1.10E+08 | 1.10E+08 | ENSMUSG00000081865 | MGI:3705726 | Gm15484 | predicted gene 15484 [Source:MGI Symbol;Acc:MGI:3705726] |
| 5 | processed_pseudogene | 1.10E+08 | 1.10E+08 | ENSMUSG00000081256 | MGI:3783233 | Gm15791 | predicted gene 15791 [Source:MGI Symbol;Acc:MGI:3783233] |
| 5 | processed_pseudogene | 1.11E+08 | 1.11E+08 | ENSMUSG00000105603 | MGI:5592155 | Gm32996 | predicted gene, 32996 [Source:MGI Symbol;Acc:MGI:5592155] |
| 5 | processed_pseudogene | 1.11E+08 | 1.11E+08 | ENSMUSG00000104602 | MGI:5662916 | Gm42779 | predicted gene 42779 [Source:MGI Symbol;Acc:MGI:5662916] |

| | | | | | | | |
|---|---|---|---|---|---|---|---|
| 5 | processed_pseudogene | 1.11E+08 | 1.11E+08 | ENSMUSG00000081104 | MGI:3801755 | Gm15989 | predicted gene 15989 [Source:MGI Symbol;Acc:MGI:3801755] |
| 5 | processed_pseudogene | 1.11E+08 | 1.11E+08 | ENSMUSG00000083771 | MGI:3801756 | Gm15988 | predicted gene 15988 [Source:MGI Symbol;Acc:MGI:3801756] |
| 5 | processed_pseudogene | 1.13E+08 | 1.13E+08 | ENSMUSG00000105406 | MGI:5663384 | Gm43247 | predicted gene 43247 [Source:MGI Symbol;Acc:MGI:5663384] |
| 5 | processed_pseudogene | 1.14E+08 | 1.14E+08 | ENSMUSG00000105031 | MGI:3781688 | Gm3511 | predicted gene 3511 [Source:MGI Symbol;Acc:MGI:3781688] |
| 5 | processed_pseudogene | 1.14E+08 | 1.14E+08 | ENSMUSG00000104972 | MGI:5012116 | Gm19931 | predicted gene, 19931 [Source:MGI Symbol;Acc:MGI:5012116] |
| 5 | processed_pseudogene | 1.14E+08 | 1.14E+08 | ENSMUSG00000106484 | MGI:5010018 | Gm17833 | predicted gene, 17833 [Source:MGI Symbol;Acc:MGI:5010018] |
| 5 | processed_pseudogene | 1.14E+08 | 1.14E+08 | ENSMUSG00000084243 | MGI:3649548 | Gm13784 | predicted gene 13784 [Source:MGI Symbol;Acc:MGI:3649548] |
| 5 | processed_pseudogene | 1.15E+08 | 1.15E+08 | ENSMUSG00000082052 | MGI:3650308 | Ywhaq-ps2 | tyrosine 3-monooxygenase/tryptophan 5-monooxygenase activation protein theta, pseudogene 2 [Source:MGI Symbol;Acc:MGI:3650308] |
| 5 | processed_pseudogene | 1.15E+08 | 1.15E+08 | ENSMUSG00000080707 | MGI:3651997 | Gm13823 | predicted gene 13823 [Source:MGI Symbol;Acc:MGI:3651997] |
| 5 | processed_pseudogene | 1.15E+08 | 1.15E+08 | ENSMUSG00000086167 | MGI:3651518 | Gm13827 | predicted gene 13827 [Source:MGI Symbol;Acc:MGI:3651518] |
| 5 | processed_pseudogene | 1.15E+08 | 1.15E+08 | ENSMUSG00000083182 | MGI:3651756 | Gm13828 | predicted gene 13828 [Source:MGI Symbol;Acc:MGI:3651756] |

| | | | | | | | |
|---|---|---|---|---|---|---|---|
| 5 | processed_pseudogene | 1.15E+08 | 1.15E+08 | ENSMUSG00000083875 | MGI:3651085 | Gm13831 | predicted gene 13831 [Source:MGI Symbol;Acc:MGI:3651085] |
| 5 | processed_pseudogene | 1.15E+08 | 1.15E+08 | ENSMUSG00000082610 | MGI:3651367 | Gm13829 | predicted gene 13829 [Source:MGI Symbol;Acc:MGI:3651367] |
| 5 | processed_pseudogene | 1.16E+08 | 1.16E+08 | ENSMUSG00000083833 | MGI:3650890 | Gm13841 | predicted gene 13841 [Source:MGI Symbol;Acc:MGI:3650890] |
| 5 | processed_pseudogene | 1.17E+08 | 1.17E+08 | ENSMUSG00000107369 | MGI:3648223 | Gstm2-ps1 | glutathione S-transferase mu 2 (muscle), pseudogene 1 [Source:MGI Symbol;Acc:MGI:3648223] |
| 5 | processed_pseudogene | 1.17E+08 | 1.17E+08 | ENSMUSG00000107327 | MGI:3648572 | Gm7478 | predicted gene 7478 [Source:MGI Symbol;Acc:MGI:3648572] |
| 5 | processed_pseudogene | 1.17E+08 | 1.17E+08 | ENSMUSG00000106829 | MGI:3781959 | Gm3786 | predicted gene 3786 [Source:MGI Symbol;Acc:MGI:3781959] |
| 5 | processed_pseudogene | 1.17E+08 | 1.17E+08 | ENSMUSG00000081274 | MGI:3783170 | Gm15727 | predicted gene 15727 [Source:MGI Symbol;Acc:MGI:3783170] |
| 5 | processed_pseudogene | 1.21E+08 | 1.21E+08 | ENSMUSG00000107004 | MGI:5662794 | Gm42657 | predicted gene 42657 [Source:MGI Symbol;Acc:MGI:5662794] |
| 5 | processed_pseudogene | 1.22E+08 | 1.22E+08 | ENSMUSG00000082760 | MGI:3782995 | Gm15546 | predicted gene 15546 [Source:MGI Symbol;Acc:MGI:3782995] |
| 5 | processed_pseudogene | 1.22E+08 | 1.22E+08 | ENSMUSG00000082265 | MGI:3782996 | Gm15547 | predicted gene 15547 [Source:MGI Symbol;Acc:MGI:3782996] |
| 5 | processed_pseudogene | 1.22E+08 | 1.22E+08 | ENSMUSG00000082645 | MGI:3705452 | Gm15481 | predicted gene 15481 [Source:MGI Symbol;Acc:MGI:3705452] |
| 5 | processed_pseudogene | 1.22E+08 | 1.22E+08 | ENSMUSG00000083909 | MGI:3801843 | Gm15842 | predicted gene 15842 [Source:MGI Symbol;Acc:MGI:3801843] |

| | | | | | | | |
|---|---|---|---|---|---|---|---|
| 5 | processed_pseudogene | 1.23E+08 | 1.23E+08 | ENSMUSG00000079906 | MGI:3801824 | Gm15846 | predicted gene 15846 [Source:MGI Symbol;Acc:MGI:3801824] |
| 5 | processed_pseudogene | 1.23E+08 | 1.23E+08 | ENSMUSG00000059352 | MGI:3645905 | Gm10064 | predicted gene 10064 [Source:MGI Symbol;Acc:MGI:3645905] |
| 5 | processed_pseudogene | 1.23E+08 | 1.23E+08 | ENSMUSG00000080836 | MGI:3648226 | Gm6444 | predicted gene 6444 [Source:MGI Symbol;Acc:MGI:3648226] |
| 5 | processed_pseudogene | 1.23E+08 | 1.23E+08 | ENSMUSG00000094664 | MGI:3704258 | Rpl35a-ps6 | ribosomal protein L35A, pseudogene 6 [Source:MGI Symbol;Acc:MGI:3704258] |
| 5 | processed_pseudogene | 1.24E+08 | 1.24E+08 | ENSMUSG00000082532 | MGI:3783190 | Rpl31-ps6 | ribosomal protein L31, pseudogene 6 [Source:MGI Symbol;Acc:MGI:3783190] |
| 5 | processed_pseudogene | 1.25E+08 | 1.25E+08 | ENSMUSG00000082402 | MGI:3783066 | Gm15621 | predicted gene 15621 [Source:MGI Symbol;Acc:MGI:3783066] |
| 5 | protein_coding | 41782319 | 41865500 | ENSMUSG00000029128 | MGI:1917285 | Rab28 | RAB28, member RAS oncogene family [Source:MGI Symbol;Acc:MGI:1917285] |
| 5 | protein_coding | 41918826 | 41921844 | ENSMUSG00000049691 | MGI:108015 | Nkx3-2 | NK3 homeobox 2 [Source:MGI Symbol;Acc:MGI:108015] |
| 5 | protein_coding | 41944881 | 42001658 | ENSMUSG00000061755 | MGI:2444804 | Bod1l | biorientation of chromosomes in cell division 1-like [Source:MGI Symbol;Acc:MGI:2444804] |
| 5 | protein_coding | 43390513 | 43447067 | ENSMUSG00000039782 | MGI:2442640 | Cpeb2 | [illegible] protein 2 [Source:MGI Symbol;Acc:MGI:2442640] |
| 5 | protein_coding | 43672881 | 43776145 | ENSMUSG00000061535 | MGI:1925911 | C1qtnf7 | C1q and tumor necrosis factor related protein 7 [Source:MGI Symbol;Acc:MGI:1925911] |
| 5 | protein_coding | 43819688 | 43898314 | ENSMUSG00000039765 | MGI:1924487 | Cc2d2a | coiled-coil and C2 domain containing 2A [Source:MGI Symbol;Acc:MGI:1924487] |
| 5 | protein_coding | 43901957 | 43978980 | ENSMUSG00000039753 | MGI:2152883 | Fbxl5 | F-box and leucine-rich repeat protein 5 [Source:MGI Symbol;Acc:MGI:2152883] |
| 5 | protein_coding | 43976227 | 44001328 | ENSMUSG00000029082 | MGI:105370 | Bst1 | bone marrow stromal cell antigen 1 [Source:MGI Symbol;Acc:MGI:105370] |
| 5 | protein_coding | 44025895 | 44069717 | ENSMUSG00000029084 | MGI:107474 | Cd38 | CD38 antigen [Source:MGI Symbol;Acc:MGI:107474] |

| | | | | | | | |
|---|---|---|---|---|---|---|---|
| 5 | protein_coding | 44070486 | 44071259 | ENSMUSG00000033036 | MGI:3645078 | Ppihl | peptidyl prolyl isomerase H like [Source:MGI Symbol;Acc:MGI:3645078] |
| 5 | protein_coding | 44136200 | 44139121 | ENSMUSG00000048373 | MGI:1096350 | Fgfbp1 | fibroblast growth factor binding protein 1 [Source:MGI Symbol;Acc:MGI:1096350] |
| 5 | protein_coding | 44150962 | 44259374 | ENSMUSG00000029086 | MGI:1100886 | Prom1 | prominin 1 [Source:MGI Symbol;Acc:MGI:1100886] |
| 5 | protein_coding | 44332496 | 44383968 | ENSMUSG00000046985 | MGI:2683537 | Tapt1 | transformation 1 [Source:MGI Symbol;Acc:MGI:2683537] |
| 5 | protein_coding | 44629474 | 44957022 | ENSMUSG00000039706 | MGI:894670 | Ldb2 | LIM domain binding 2 [Source:MGI Symbol;Acc:MGI:894670] |
| 5 | protein_coding | 45591363 | 45607578 | ENSMUSG00000015806 | MGI:97836 | Qdpr | quinoid dihydropteridine reductase [Source:MGI Symbol;Acc:MGI:97836] |
| 5 | protein_coding | 45611093 | 45621491 | ENSMUSG00000049530 | MGI:3646230 | Clrn2 | clarin 2 [Source:MGI Symbol;Acc:MGI:3646230] |
| 5 | protein_coding | 45650716 | 45670033 | ENSMUSG00000039682 | MGI:1914238 | Lap3 | leucine aminopeptidase 3 [Source:MGI Symbol;Acc:MGI:1914238] |
| 5 | protein_coding | 45677571 | 45686618 | ENSMUSG00000015804 | MGI:1914249 | Med28 | mediator complex subunit 28 [Source:MGI Symbol;Acc:MGI:1914249] |
| 5 | protein_coding | 45687047 | 45796956 | ENSMUSG00000015879 | MGI:2442958 | Fam184b | family with sequence similarity 184, member B [Source:MGI Symbol;Acc:MGI:2442958] |
| 5 | protein_coding | 45827261 | 45857888 | ENSMUSG00000015880 | MGI:1930197 | Ncapg | non-SMC condensin I complex, subunit G [Source:MGI Symbol;Acc:MGI:1930197] |
| 5 | protein_coding | 45854523 | 46014957 | ENSMUSG00000015882 | MGI:2651932 | Lcorl | ligand dependent nuclear receptor corepressor-like [Source:MGI Symbol;Acc:MGI:2651932] |
| 5 | protein_coding | 48140480 | 48465075 | ENSMUSG00000031558 | MGI:1315205 | Slit2 | slit guidance ligand 2 [Source:MGI Symbol;Acc:MGI:1315205] |
| 5 | protein_coding | 48485294 | 48571636 | ENSMUSG00000029089 | MGI:1917842 | 5730480H06Rik | RIKEN cDNA 5730480H06 gene [Source:MGI Symbol;Acc:MGI:1917842] |
| 5 | protein_coding | 48546844 | 49682249 | ENSMUSG00000029088 | MGI:1933131 | Kcnip4 | Kv channel interacting protein 4 [Source:MGI Symbol;Acc:MGI:1933131] |
| 5 | protein_coding | 50117298 | 50216348 | ENSMUSG00000029090 | MGI:1917943 | Adgra3 | adhesion G protein-coupled receptor A3 [Source:MGI Symbol;Acc:MGI:1917943] |
| 5 | protein_coding | 51611592 | 51725068 | ENSMUSG00000029167 | MGI:1342774 | Ppargc1a | peroxisome proliferative activated receptor, gamma, coactivator 1 alpha [Source:MGI Symbol;Acc:MGI:1342774] |

| | | | | | | | |
|---|---|---|---|---|---|---|---|
| 5 | protein_coding | 52307545 | 52347856 | ENSMUSG00000029169 | MGI:1099786 | Dhx15 | DEAH-box helicase 15 [Source:MGI Symbol;Acc:MGI:1099786] |
| 5 | protein_coding | 52521133 | 52528760 | ENSMUSG00000072941 | MGI:103181 | Sod3 | superoxide dismutase 3, extracellular [Source:MGI Symbol;Acc:MGI:103181] |
| 5 | protein_coding | 52531993 | 52628863 | ENSMUSG00000045790 | MGI:2685293 | Ccdc149 | coiled-coil domain containing 149 [Source:MGI Symbol;Acc:MGI:2685293] |
| 5 | protein_coding | 52690859 | 52723804 | ENSMUSG00000039252 | MGI:2180196 | Lgi2 | leucine-rich repeat LGI family, member 2 [Source:MGI Symbol;Acc:MGI:2180196] |
| 5 | protein_coding | 52797429 | 52827050 | ENSMUSG00000029173 | MGI:1098791 | Sepsecs | Sep (O-phosphoserine) tRNA:Sec (selenocysteine) tRNA synthase [Source:MGI Symbol;Acc:MGI:1098791] |
| 5 | protein_coding | 52898916 | 52926682 | ENSMUSG00000029186 | MGI:1914323 | Pi4k2b | phosphatidylinositol 4-kinase type 2 beta [Source:MGI Symbol;Acc:MGI:1914323] |
| 5 | protein_coding | 52932751 | 52982007 | ENSMUSG00000029179 | MGI:1926046 | Zcchc4 | zinc finger, CCHC domain containing 4 [Source:MGI Symbol;Acc:MGI:1926046] |
| 5 | protein_coding | 52991354 | 53025139 | ENSMUSG00000029176 | MGI:1098673 | Anapc4 | anaphase promoting complex subunit 4 [Source:MGI Symbol;Acc:MGI:1098673] |
| 5 | protein_coding | 53195423 | 53229006 | ENSMUSG00000029188 | MGI:1342284 | Slc34a2 | [illegible] member 2 [Source:MGI Symbol;Acc:MGI:1342284] |
| 5 | protein_coding | 53264425 | 53371269 | ENSMUSG00000029189 | MGI:1916941 | Sel1l3 | sel-1 suppressor of lin-12-like 3 (C. elegans) [Source:MGI Symbol;Acc:MGI:1916941] |
| 5 | protein_coding | 53424425 | 53435882 | ENSMUSG00000061461 | MGI:1913528 | Smim20 | small integral membrane protein 20 [Source:MGI Symbol;Acc:MGI:1913528] |
| 5 | protein_coding | 53623494 | 53814704 | ENSMUSG00000039191 | MGI:96522 | Rbpj | recombination signal binding protein for immunoglobulin kappa J region [Source:MGI Symbol;Acc:MGI:96522] |
| 5 | protein_coding | 53855118 | 53865047 | ENSMUSG00000029193 | MGI:99478 | Cckar | cholecystokinin A receptor [Source:MGI Symbol;Acc:MGI:99478] |
| 5 | protein_coding | 53966948 | 54061307 | ENSMUSG00000039178 | MGI:1914499 | Tbc1d19 | TBC1 domain family, member 19 [Source:MGI Symbol;Acc:MGI:1914499] |
| 5 | protein_coding | 54155841 | 54278399 | ENSMUSG00000039156 | MGI:2151156 | Stim2 | stromal interaction molecule 2 [Source:MGI Symbol;Acc:MGI:2151156] |
| 5 | protein_coding | 57875309 | 58290572 | ENSMUSG00000029108 | MGI:1860487 | Pcdh7 | protocadherin 7 [Source:MGI Symbol;Acc:MGI:1860487] |
| 5 | protein_coding | 61966159 | 61968506 | ENSMUSG00000089992 | MGI:105977 | G6pd2 | glucose-6-phosphate dehydrogenase 2 [Source:MGI Symbol;Acc:MGI:105977] |

| | | | | | | | |
|---|---|---|---|---|---|---|---|
| 5 | protein_coding | 62759788 | 62923502 | ENSMUSG00000037999 | MGI:2684416 | Arap2 | ArfGAP with RhoGAP domain, ankyrin repeat and PH domain 2 [Source:MGI Symbol;Acc:MGI:2684416] |
| 5 | protein_coding | 62969017 | 63045651 | ENSMUSG00000090326 | MGI:4937018 | Dthd1 | death domain containing 1 [Source:MGI Symbol;Acc:MGI:4937018] |
| 5 | protein_coding | 63806445 | 63967889 | ENSMUSG00000090061 | MGI:1920464 | Nwd2 | NACHT and WD repeat domain containing 2 [Source:MGI Symbol;Acc:MGI:1920464] |
| 5 | protein_coding | 63969706 | 64056968 | ENSMUSG00000060512 | MGI:1923511 | Pgcka1 | PDCD10 and GCKIII kinases associated 1 [Source:MGI Symbol;Acc:MGI:1923511] |
| 5 | protein_coding | 64066240 | 64126240 | ENSMUSG00000047881 | MGI:2140767 | Rell1 | RELT-like 1 [Source:MGI Symbol;Acc:MGI:2140767] |
| 5 | protein_coding | 64250293 | 64285694 | ENSMUSG00000029171 | MGI:97564 | Pgm2 | phosphoglucomutase 2 [Source:MGI Symbol;Acc:MGI:97564] |
| 5 | protein_coding | 64313648 | 64508829 | ENSMUSG00000029174 | MGI:1889508 | Tbc1d1 | TBC1 domain family, member 1 [Source:MGI Symbol;Acc:MGI:1889508] |
| 5 | protein_coding | 64960731 | 64990244 | ENSMUSG00000029178 | MGI:1342773 | Klf3 | Kruppel-like transcription factor 3 (basic) [Source:MGI Symbol;Acc:MGI:1342773] |
| 5 | protein_coding | 65082022 | 65090906 | ENSMUSG00000044827 | MGI:1341295 | Tlr1 | toll-like receptor 1 [Source:MGI Symbol;Acc:MGI:1341295] |
| 5 | protein_coding | 65109374 | 65117440 | ENSMUSG00000051498 | MGI:1341296 | Tlr6 | toll-like receptor 6 [Source:MGI Symbol;Acc:MGI:1341296] |
| 5 | protein_coding | 65127414 | 65199229 | ENSMUSG00000029185 | MGI:1915553 | Fam114a1 | family with sequence similarity 114, member A1 [Source:MGI Symbol;Acc:MGI:1915553] |
| 5 | protein_coding | 65211689 | 65249475 | ENSMUSG00000037913 | MGI:2685292 | Tmem156 | transmembrane protein 156 [Source:MGI Symbol;Acc:MGI:2685292] |
| 5 | protein_coding | 65264882 | 65325531 | ENSMUSG00000054920 | MGI:1919028 | Klhl5 | kelch-like 5 [Source:MGI Symbol;Acc:MGI:1919028] |
| 5 | protein_coding | 65357039 | 65417758 | ENSMUSG00000037890 | MGI:2443231 | Wdr19 | WD repeat domain 19 [Source:MGI Symbol;Acc:MGI:2443231] |
| 5 | protein_coding | 65419193 | 65493013 | ENSMUSG00000029191 | MGI:97891 | Rfc1 | replication factor C (activator 1) 1 [Source:MGI Symbol;Acc:MGI:97891] |
| 5 | protein_coding | 65505657 | 65541350 | ENSMUSG00000029195 | MGI:1932466 | Klb | klotho beta [Source:MGI Symbol;Acc:MGI:1932466] |
| 5 | protein_coding | 65545707 | 65548787 | ENSMUSG00000047215 | MGI:1298373 | Rpl9 | ribosomal protein L9 [Source:MGI Symbol;Acc:MGI:1298373] |

| | | | | | | | |
|---|---|---|---|---|---|---|---|
| 5 | protein_coding | 65548840 | 65568036 | ENSMUSG00000029199 | MGI:1934604 | Lias | lipoic acid synthetase [Source:MGI Symbol;Acc:MGI:1934604] |
| 5 | protein_coding | 65570564 | 65593292 | ENSMUSG00000029201 | MGI:1306785 | Ugdh | UDP-glucose dehydrogenase [Source:MGI Symbol;Acc:MGI:1306785] |
| 5 | protein_coding | 65584770 | 65650176 | ENSMUSG00000105835 | MGI:5663689 | Gm43552 | predicted gene 43552 [Source:MGI Symbol;Acc:MGI:5663689] |
| 5 | protein_coding | 65604187 | 65694527 | ENSMUSG00000037822 | MGI:1915802 | Smim14 | small integral membrane protein 14 [Source:MGI Symbol;Acc:MGI:1915802] |
| 5 | protein_coding | 65694576 | 65756331 | ENSMUSG00000029203 | MGI:1858216 | Ube2k | ubiquitin-conjugating enzyme E2K [Source:MGI Symbol;Acc:MGI:1858216] |
| 5 | protein_coding | 65763064 | 65855616 | ENSMUSG00000029202 | MGI:1918771 | Pds5a | PDS5 cohesin associated factor A [Source:MGI Symbol;Acc:MGI:1918771] |
| 5 | protein_coding | 65920864 | 65987451 | ENSMUSG00000037795 | MGI:2684414 | N4bp2 | NEDD4 binding protein 2 [Source:MGI Symbol;Acc:MGI:2684414] |
| 5 | protein_coding | 66018556 | 66054043 | ENSMUSG00000029204 | MGI:1921984 | Rhoh | ras homolog family member H [Source:MGI Symbol;Acc:MGI:1921984] |
| 5 | protein_coding | 66092264 | 66134669 | ENSMUSG00000029205 | MGI:1202403 | Chrna9 | [illegible] polypeptide 9 [Source:MGI Symbol;Acc:MGI:1202403] |
| 5 | protein_coding | 66173892 | 66330461 | ENSMUSG00000070780 | MGI:2384294 | Rbm47 | RNA binding motif protein 47 [Source:MGI Symbol;Acc:MGI:2384294] |
| 5 | protein_coding | 66417240 | 66455369 | ENSMUSG00000029206 | MGI:1918168 | Nsun7 | NOL1/NOP2/Sun domain family, member 7 [Source:MGI Symbol;Acc:MGI:1918168] |
| 5 | protein_coding | 66456046 | 66776127 | ENSMUSG00000029207 | MGI:108405 | Apbb2 | [illegible] member 2 [Source:MGI Symbol;Acc:MGI:108405] |
| 5 | protein_coding | 66700842 | 66700919 | ENSMUSG00000121580 | MGI:7820357 | Gm68765 | predicted gene, 68765 [Source:MGI Symbol;Acc:MGI:7820357] |
| 5 | protein_coding | 66833434 | 66844577 | ENSMUSG00000029223 | MGI:103149 | Uchl1 | ubiquitin carboxy-terminal hydrolase L1 [Source:MGI Symbol;Acc:MGI:103149] |
| 5 | protein_coding | 66903170 | 67214501 | ENSMUSG00000037736 | MGI:1924819 | Limch1 | LIM and calponin homology domains 1 [Source:MGI Symbol;Acc:MGI:1924819] |
| 5 | protein_coding | 67251742 | 67256644 | ENSMUSG00000012520 | MGI:1100882 | Phox2b | paired-like homeobox 2b [Source:MGI Symbol;Acc:MGI:1100882] |
| 5 | protein_coding | 67417908 | 67448804 | ENSMUSG00000037720 | MGI:1915128 | Tmem33 | transmembrane protein 33 [Source:MGI Symbol;Acc:MGI:1915128] |
| 5 | protein_coding | 67464298 | 67515786 | ENSMUSG00000029221 | MGI:1923690 | Slc30a9 | [illegible] member 9 [Source:MGI Symbol;Acc:MGI:1923690] |

| | | | | | | | |
|---|---|---|---|---|---|---|---|
| 5 | protein_coding | 67549490 | 67585653 | ENSMUSG00000092060 | MGI:3648414 | Bend4 | BEN domain containing 4 [Source:MGI Symbol;Acc:MGI:3648414] |
| 5 | protein_coding | 67765225 | 67780895 | ENSMUSG00000050010 | MGI:3041225 | Shisa3 | shisa family member 3 [Source:MGI Symbol;Acc:MGI:3041225] |
| 5 | protein_coding | 67775483 | 68004777 | ENSMUSG00000037685 | MGI:1330848 | Atp8a1 | ATPase phospholipid transporting 8A1 [Source:MGI Symbol;Acc:MGI:1330848] |
| 5 | protein_coding | 68189178 | 68323741 | ENSMUSG00000068082 | MGI:3577767 | Grxcr1 | glutaredoxin, cysteine rich 1 [Source:MGI Symbol;Acc:MGI:3577767] |
| 5 | protein_coding | 69266628 | 69499022 | ENSMUSG00000037653 | MGI:2443804 | Kctd8 | potassium channel tetramerisation domain containing 8 [Source:MGI Symbol;Acc:MGI:2443804] |
| 5 | protein_coding | 69674014 | 69699991 | ENSMUSG00000029158 | MGI:1922831 | Yipf7 | Yip1 domain family, member 7 [Source:MGI Symbol;Acc:MGI:1922831] |
| 5 | protein_coding | 69714266 | 69733316 | ENSMUSG00000029208 | MGI:2140726 | Guf1 | GUF1 homolog, GTPase [Source:MGI Symbol;Acc:MGI:2140726] |
| 5 | protein_coding | 69730451 | 69749683 | ENSMUSG00000029209 | MGI:1915230 | Gnpda2 | glucosamine-6-phosphate deaminase 2 [Source:MGI Symbol;Acc:MGI:1915230] |
| 5 | protein_coding | 70908390 | 70999960 | ENSMUSG00000001260 | MGI:103156 | Gabrg1 | subunit gamma 1 [Source:MGI Symbol;Acc:MGI:103156] |
| 5 | protein_coding | 71114940 | 71253192 | ENSMUSG00000000560 | MGI:95614 | Gabra2 | subunit alpha 2 [Source:MGI Symbol;Acc:MGI:95614] |
| 5 | protein_coding | 71600167 | 71705548 | ENSMUSG00000049387 | MGI:1925424 | Cox7b2 | cytochrome c oxidase subunit 7B2 [Source:MGI Symbol;Acc:MGI:1925424] |
| 5 | protein_coding | 71727092 | 71815651 | ENSMUSG00000029211 | MGI:95616 | Gabra4 | subunit alpha 4 [Source:MGI Symbol;Acc:MGI:95616] |
| 5 | protein_coding | 71815456 | 72306380 | ENSMUSG00000029212 | MGI:95619 | Gabrb1 | subunit beta 1 [Source:MGI Symbol;Acc:MGI:95619] |
| 5 | protein_coding | 72313918 | 72325532 | ENSMUSG00000029213 | MGI:1343485 | Commd8 | COMM domain containing 8 [Source:MGI Symbol;Acc:MGI:1343485] |
| 5 | protein_coding | 72360672 | 72456118 | ENSMUSG00000046808 | MGI:2450125 | Atp10d | ATPase, class V, type 10D [Source:MGI Symbol;Acc:MGI:2450125] |
| 5 | protein_coding | 72457368 | 72661816 | ENSMUSG00000005220 | MGI:1349451 | Corin | corin, serine peptidase [Source:MGI Symbol;Acc:MGI:1349451] |
| 5 | protein_coding | 72670644 | 72717027 | ENSMUSG00000072889 | MGI:1923646 | Nfxl1 | nuclear transcription factor, X-box binding-like 1 [Source:MGI Symbol;Acc:MGI:1923646] |

| | | | | | | | |
|---|---|---|---|---|---|---|---|
| 5 | protein_coding | 72734149 | 72738597 | ENSMUSG00000063935 | MGI:2180337 | Zar1 | zygote arrest 1 [Source:MGI Symbol;Acc:MGI:2180337] |
| 5 | protein_coding | 72738981 | 72744894 | ENSMUSG00000060204 | MGI:3645333 | Slc10a4-ps | solute carrier family 10 (sodium/bile acid cotransporter family), pseudogene [Source:MGI Symbol;Acc:MGI:3645333] |
| 5 | protein_coding | 72761039 | 72801618 | ENSMUSG00000067220 | MGI:88436 | Cnga1 | cyclic nucleotide gated channel alpha 1 [Source:MGI Symbol;Acc:MGI:88436] |
| 5 | protein_coding | 72805138 | 72828421 | ENSMUSG00000067219 | MGI:1917951 | Nipal1 | NIPA-like domain containing 1 [Source:MGI Symbol;Acc:MGI:1917951] |
| 5 | protein_coding | 72853321 | 72910120 | ENSMUSG00000054892 | MGI:102960 | Txk | TXK tyrosine kinase [Source:MGI Symbol;Acc:MGI:102960] |
| 5 | protein_coding | 72913059 | 73025826 | ENSMUSG00000029217 | MGI:98662 | Tec | tec protein tyrosine kinase [Source:MGI Symbol;Acc:MGI:98662] |
| 5 | protein_coding | 73071647 | 73136172 | ENSMUSG00000036087 | MGI:1923241 | Slain2 | SLAIN motif family, member 2 [Source:MGI Symbol;Acc:MGI:1923241] |
| 5 | protein_coding | 73123151 | 73132045 | ENSMUSG00000063656 | MGI:3710643 | Gm10135 | predicted gene 10135 [Source:MGI Symbol;Acc:MGI:3710643] |
| 5 | protein_coding | 73164226 | 73170298 | ENSMUSG00000029219 | MGI:3606480 | Slc10a4 | solute carrier family 10 (sodium/bile acid cotransporter family), member 4 [Source:MGI Symbol;Acc:MGI:3606480] |
| 5 | protein_coding | 73177330 | 73413962 | ENSMUSG00000070733 | MGI:1919563 | Fryl | FRY like transcription coactivator [Source:MGI Symbol;Acc:MGI:1919563] |
| 5 | protein_coding | 73450127 | 73471412 | ENSMUSG00000029152 | MGI:1915345 | Ociad1 | OCIA domain containing 1 [Source:MGI Symbol;Acc:MGI:1915345] |
| 5 | protein_coding | 73479542 | 73498371 | ENSMUSG00000029153 | MGI:1916377 | Ociad2 | OCIA domain containing 2 [Source:MGI Symbol;Acc:MGI:1916377] |
| 5 | protein_coding | 73563418 | 73610778 | ENSMUSG00000029154 | MGI:2444131 | Cwh43 | cell wall biogenesis 43 C-terminal homolog [Source:MGI Symbol;Acc:MGI:2444131] |
| 5 | protein_coding | 73638343 | 73718137 | ENSMUSG00000051674 | MGI:2140972 | Dcun1d4 | containing 4 [Source:MGI Symbol;Acc:MGI:2140972] |
| 5 | protein_coding | 73763985 | 73789869 | ENSMUSG00000067206 | MGI:2387634 | Lrrc66 | leucine rich repeat containing 66 [Source:MGI Symbol;Acc:MGI:2387634] |
| 5 | protein_coding | 73790092 | 73805133 | ENSMUSG00000029156 | MGI:1346523 | Sgcb | sarcoglycan, beta (dystrophin-associated glycoprotein) [Source:MGI Symbol;Acc:MGI:1346523] |

| | | | | | | | |
|---|---|---|---|---|---|---|---|
| 5 | protein_coding | 73808722 | 73836855 | ENSMUSG00000029155 | MGI:1920722 | Spata18 | spermatogenesis associated 18 [Source:MGI Symbol;Acc:MGI:1920722] |
| 5 | protein_coding | 74159114 | 74229092 | ENSMUSG00000054814 | MGI:1916977 | Usp46 | ubiquitin specific peptidase 46 [Source:MGI Symbol;Acc:MGI:1916977] |
| 5 | protein_coding | 74355947 | 74360142 | ENSMUSG00000049907 | MGI:1916189 | Rasl11b | RAS-like, family 11, member B [Source:MGI Symbol;Acc:MGI:1916189] |
| 5 | protein_coding | 74365477 | 74692420 | ENSMUSG00000062110 | MGI:2443446 | Scfd2 | Sec1 family domain containing 2 [Source:MGI Symbol;Acc:MGI:2443446] |
| 5 | protein_coding | 74696110 | 74759461 | ENSMUSG00000029227 | MGI:1914149 | Fip1l1 | factor interacting with PAPOLA and CPSF1 [Source:MGI Symbol;Acc:MGI:1914149] |
| 5 | protein_coding | 74753108 | 74863573 | ENSMUSG00000029228 | MGI:1278335 | Lnx1 | ligand of numb-protein X 1 [Source:MGI Symbol;Acc:MGI:1278335] |
| 5 | protein_coding | 75158649 | 75205435 | ENSMUSG00000029229 | MGI:1921527 | Chic2 | cysteine-rich hydrophobic domain 2 [Source:MGI Symbol;Acc:MGI:1921527] |
| 5 | protein_coding | 75236262 | 75238554 | ENSMUSG00000035946 | MGI:95843 | Gsx2 | GS homeobox 2 [Source:MGI Symbol;Acc:MGI:95843] |
| 5 | protein_coding | 75312953 | 75358876 | ENSMUSG00000029231 | MGI:97530 | Pdgfra | [illegible] polypeptide [Source:MGI Symbol;Acc:MGI:97530] |
| 5 | protein_coding | 75735576 | 75817382 | ENSMUSG00000005672 | MGI:96677 | Kit | KIT proto-oncogene receptor tyrosine kinase [Source:MGI Symbol;Acc:MGI:96677] |
| 5 | protein_coding | 76093487 | 76139118 | ENSMUSG00000062960 | MGI:96683 | Kdr | kinase insert domain protein receptor [Source:MGI Symbol;Acc:MGI:96683] |
| 5 | protein_coding | 76288118 | 76303351 | ENSMUSG00000029233 | MGI:1930252 | Srd5a3 | steroid 5 alpha-reductase 3 [Source:MGI Symbol;Acc:MGI:1930252] |
| 5 | protein_coding | 76331727 | 76357092 | ENSMUSG00000029234 | MGI:894407 | Tmem165 | transmembrane protein 165 [Source:MGI Symbol;Acc:MGI:894407] |
| 5 | protein_coding | 76357715 | 76452639 | ENSMUSG00000029238 | MGI:99698 | Clock | clock circadian regulator [Source:MGI Symbol;Acc:MGI:99698] |
| 5 | protein_coding | 76459962 | 76479003 | ENSMUSG00000029235 | MGI:1890655 | Pdcl2 | phosducin-like 2 [Source:MGI Symbol;Acc:MGI:1890655] |
| 5 | protein_coding | 76481342 | 76511635 | ENSMUSG00000029236 | MGI:1860476 | Nmu | neuromedin U [Source:MGI Symbol;Acc:MGI:1860476] |
| 5 | protein_coding | 76631887 | 76664496 | ENSMUSG00000091204 | MGI:3647743 | Exoc1l | exocyst complex component 1 like [Source:MGI Symbol;Acc:MGI:3647743] |
| 5 | protein_coding | 76677158 | 76718141 | ENSMUSG00000036435 | MGI:2445020 | Exoc1 | exocyst complex component 1 [Source:MGI Symbol;Acc:MGI:2445020] |

| | | | | | | | |
|---|---|---|---|---|---|---|---|
| 5 | protein_coding | 76736545 | 76794313 | ENSMUSG00000036403 | MGI:2681869 | Cep135 | centrosomal protein 135 [Source:MGI Symbol;Acc:MGI:2681869] |
| 5 | protein_coding | 76804359 | 77021392 | ENSMUSG00000036377 | MGI:2444817 | Cracd | capping protein inhibiting regulator of actin [Source:MGI Symbol;Acc:MGI:2444817] |
| 5 | protein_coding | 77021506 | 77053361 | ENSMUSG00000055923 | MGI:2442517 | Aasdh | aminoadipate-semialdehyde dehydrogenase [Source:MGI Symbol;Acc:MGI:2442517] |
| 5 | protein_coding | 77061096 | 77099425 | ENSMUSG00000029246 | MGI:2387203 | Ppat | [illegible] amidotransferase [Source:MGI Symbol;Acc:MGI:2387203] |
| 5 | protein_coding | 77099154 | 77115356 | ENSMUSG00000029247 | MGI:1914304 | Paics | phosphoribosylaminoimidazole carboxylase, phosphoribosylaminoribosylaminoimidazole, succinocarboxamide synthetase [Source:MGI Symbol;Acc:MGI:1914304] |
| 5 | protein_coding | 77122530 | 77147784 | ENSMUSG00000036323 | MGI:1333795 | Srp72 | signal recognition particle 72 [Source:MGI Symbol;Acc:MGI:1333795] |
| 5 | protein_coding | 77151902 | 77158453 | ENSMUSG00000063820 | MGI:1915496 | Arl9 | ADP-ribosylation factor-like 9 [Source:MGI Symbol;Acc:MGI:1915496] |
| 5 | protein_coding | 77163870 | 77209376 | ENSMUSG00000029248 | MGI:1919118 | Spmap2l | sperm microtubule associated protein 2 like [Source:MGI Symbol;Acc:MGI:1919118] |
| 5 | protein_coding | 77234835 | 77262968 | ENSMUSG00000059325 | MGI:1916782 | Hopx | HOP homeobox [Source:MGI Symbol;Acc:MGI:1916782] |
| 5 | protein_coding | 77352954 | 77359318 | ENSMUSG00000053030 | MGI:1917232 | Spink2 | serine peptidase inhibitor, Kazal type 2 [Source:MGI Symbol;Acc:MGI:1917232] |
| 5 | protein_coding | 77413338 | 77434279 | ENSMUSG00000029249 | MGI:104897 | Rest | RE1-silencing transcription factor [Source:MGI Symbol;Acc:MGI:104897] |
| 5 | protein_coding | 77442029 | 77457931 | ENSMUSG00000036285 | MGI:1914306 | Noa1 | nitric oxide associated 1 [Source:MGI Symbol;Acc:MGI:1914306] |
| 5 | protein_coding | 77457994 | 77497171 | ENSMUSG00000029250 | MGI:2388280 | Polr2b | [illegible] polypeptide B [Source:MGI Symbol;Acc:MGI:2388280] |
| 5 | protein_coding | 77497087 | 77555888 | ENSMUSG00000036256 | MGI:1352480 | Igfbp7 | insulin-like growth factor binding protein 7 [Source:MGI Symbol;Acc:MGI:1352480] |
| 5 | protein_coding | 81167985 | 81972980 | ENSMUSG00000037605 | MGI:2441950 | Adgrl3 | adhesion G protein-coupled receptor L3 [Source:MGI Symbol;Acc:MGI:2441950] |
| 5 | protein_coding | 82270255 | 82272560 | ENSMUSG00000099863 | MGI:1914577 | 1700031L13Rik | RIKEN cDNA 1700031L13 gene [Source:MGI Symbol;Acc:MGI:1914577] |
| 5 | protein_coding | 83425992 | 83503042 | ENSMUSG00000049537 | MGI:2444966 | Tecrl | trans-2,3-enoyl-CoA reductase-like [Source:MGI Symbol;Acc:MGI:2444966] |

| | | | | | | | |
|---|---|---|---|---|---|---|---|
| 5 | protein_coding | 84202620 | 84565241 | ENSMUSG00000029245 | MGI:99654 | Epha5 | Eph receptor A5 [Source:MGI Symbol;Acc:MGI:99654] |
| 5 | protein_coding | 86159883 | 86213442 | ENSMUSG00000029253 | MGI:99700 | Cenpc1 | centromere protein C1 [Source:MGI Symbol;Acc:MGI:99700] |
| 5 | protein_coding | 86219605 | 86253984 | ENSMUSG00000029254 | MGI:1926193 | Stap1 | signal transducing adaptor family member 1 [Source:MGI Symbol;Acc:MGI:1926193] |
| 5 | protein_coding | 86257146 | 86320662 | ENSMUSG00000035898 | MGI:1913894 | Uba6 | ubiquitin-like modifier activating enzyme 6 [Source:MGI Symbol;Acc:MGI:1913894] |
| 5 | protein_coding | 86328613 | 86345760 | ENSMUSG00000029255 | MGI:95790 | Gnrhr | gonadotropin releasing hormone receptor [Source:MGI Symbol;Acc:MGI:95790] |
| 5 | protein_coding | 86379340 | 86437167 | ENSMUSG00000061184 | MGI:3521861 | Tmprss11c | transmembrane protease, serine 11c [Source:MGI Symbol;Acc:MGI:3521861] |
| 5 | protein_coding | 86450076 | 86521279 | ENSMUSG00000061259 | MGI:2385221 | Tmprss11d | transmembrane protease, serine 11d [Source:MGI Symbol;Acc:MGI:2385221] |
| 5 | protein_coding | 86558269 | 86616849 | ENSMUSG00000072845 | MGI:2684853 | Tmprss11a | transmembrane protease, serine 11a [Source:MGI Symbol;Acc:MGI:2684853] |
| 5 | protein_coding | 86633735 | 86666459 | ENSMUSG00000079451 | MGI:2444058 | Tmprss11g | transmembrane protease, serine 11g [Source:MGI Symbol;Acc:MGI:2444058] |
| 5 | protein_coding | 86669757 | 86780283 | ENSMUSG00000048764 | MGI:2442348 | Tmprss11f | transmembrane protease, serine 11f [Source:MGI Symbol;Acc:MGI:2442348] |
| 5 | protein_coding | 86805490 | 86824221 | ENSMUSG00000035861 | MGI:2442893 | Tmprss11b | transmembrane protease, serine 11B [Source:MGI Symbol;Acc:MGI:2442893] |
| 5 | protein_coding | 86853045 | 86893674 | ENSMUSG00000054537 | MGI:3513175 | Tmprss11e | transmembrane protease, serine 11e [Source:MGI Symbol;Acc:MGI:3513175] |
| 5 | protein_coding | 86952080 | 86984518 | ENSMUSG00000035851 | MGI:2443713 | Ythdc1 | YTH domain containing 1 [Source:MGI Symbol;Acc:MGI:2443713] |
| 5 | protein_coding | 87037626 | 87054796 | ENSMUSG00000029260 | MGI:2140962 | Ugt2b34 | [illegible] polypeptide B34 [Source:MGI Symbol;Acc:MGI:2140962] |
| 5 | protein_coding | 87064497 | 87074389 | ENSMUSG00000035836 | MGI:1919023 | Ugt2b1 | [illegible] polypeptide B1 [Source:MGI Symbol;Acc:MGI:1919023] |
| 5 | protein_coding | 87148719 | 87161134 | ENSMUSG00000035811 | MGI:3576100 | Ugt2b35 | [illegible] polypeptide B35 [Source:MGI Symbol;Acc:MGI:3576100] |
| 5 | protein_coding | 87213786 | 87240414 | ENSMUSG00000070704 | MGI:3576103 | Ugt2b36 | [illegible] polypeptide B36 [Source:MGI Symbol;Acc:MGI:3576103] |
| 5 | protein_coding | 87272819 | 87288177 | ENSMUSG00000054630 | MGI:98900 | Ugt2b5 | [illegible] polypeptide B5 [Source:MGI Symbol;Acc:MGI:98900] |

| | | | | | | | |
|---|---|---|---|---|---|---|---|
| 5 | protein_coding | 87388352 | 87402663 | ENSMUSG00000057425 | MGI:2148239 | Ugt2b37 | polypeptide B37 [Source:MGI Symbol;Acc:MGI:2148239] |
| 5 | protein_coding | 87472831 | 87485054 | ENSMUSG00000035780 | MGI:1919344 | Ugt2a3 | polypeptide A3 [Source:MGI Symbol;Acc:MGI:1919344] |
| 5 | protein_coding | 87557801 | 87572062 | ENSMUSG00000061906 | MGI:2140794 | Ugt2b38 | polypeptide B38 [Source:MGI Symbol;Acc:MGI:2140794] |
| 5 | protein_coding | 87607349 | 87630119 | ENSMUSG00000029268 | MGI:3576095 | Ugt2a2 | polypeptide A2 [Source:MGI Symbol;Acc:MGI:3576095] |
| 5 | protein_coding | 87607349 | 87638730 | ENSMUSG00000106677 | MGI:2149905 | Ugt2a1 | polypeptide A1 [Source:MGI Symbol;Acc:MGI:2149905] |
| 5 | protein_coding | 87608099 | 87634665 | ENSMUSG00000107180 | MGI:5663775 | Gm43638 | predicted gene 43638 [Source:MGI Symbol;Acc:MGI:5663775] |
| 5 | protein_coding | 87661198 | 87686054 | ENSMUSG00000029269 | MGI:2136282 | Sult1b1 | sulfotransferase family 1B, member 1 [Source:MGI Symbol;Acc:MGI:2136282] |
| 5 | protein_coding | 87702504 | 87716886 | ENSMUSG00000029273 | MGI:1926341 | Sult1d1 | sulfotransferase family 1D, member 1 [Source:MGI Symbol;Acc:MGI:1926341] |
| 5 | protein_coding | 87723828 | 87739453 | ENSMUSG00000029272 | MGI:98431 | Sult1e1 | sulfotransferase family 1E, member 1 [Source:MGI Symbol;Acc:MGI:98431] |
| 5 | protein_coding | 87814067 | 87830437 | ENSMUSG00000070702 | MGI:88540 | Csn1s1 | casein alpha s1 [Source:MGI Symbol;Acc:MGI:88540] |
| 5 | protein_coding | 87840483 | 87847489 | ENSMUSG00000063157 | MGI:88541 | Csn2 | casein beta [Source:MGI Symbol;Acc:MGI:88541] |
| 5 | protein_coding | 87922426 | 87936656 | ENSMUSG00000061937 | MGI:88542 | Csn1s2a | casein alpha s2-like A [Source:MGI Symbol;Acc:MGI:88542] |
| 5 | protein_coding | 87955941 | 87972285 | ENSMUSG00000061388 | MGI:105312 | Csn1s2b | casein alpha s2-like B [Source:MGI Symbol;Acc:MGI:105312] |
| 5 | protein_coding | 87973545 | 87994246 | ENSMUSG00000002240 | MGI:1921029 | Prr27 | proline rich 27 [Source:MGI Symbol;Acc:MGI:1921029] |
| 5 | protein_coding | 88032888 | 88042033 | ENSMUSG00000009580 | MGI:1916842 | Odam | odontogenic, ameloblast asssociated [Source:MGI Symbol;Acc:MGI:1916842] |
| 5 | protein_coding | 88056638 | 88062030 | ENSMUSG00000105888 | MGI:3512757 | Fdcsp | follicular dendritic cell secreted protein [Source:MGI Symbol;Acc:MGI:3512757] |
| 5 | protein_coding | 88073438 | 88080524 | ENSMUSG00000001622 | MGI:107461 | Csn3 | casein kappa [Source:MGI Symbol;Acc:MGI:107461] |
| 5 | protein_coding | 88117318 | 88120729 | ENSMUSG00000007457 | MGI:1921498 | 2310003L06Rik | RIKEN cDNA 2310003L06 gene [Source:MGI Symbol;Acc:MGI:1921498] |

| | | | | | | | |
|---|---|---|---|---|---|---|---|
| 5 | protein_coding | 88117376 | 88156136 | ENSMUSG00000100704 | MGI:5579140 | Gm28434 | predicted gene 28434 [Source:MGI Symbol;Acc:MGI:5579140] |
| 5 | protein_coding | 88127298 | 88129403 | ENSMUSG00000007907 | MGI:1918227 | Cabs1 | calcium binding protein, spermatid specific 1 [Source:MGI Symbol;Acc:MGI:1918227] |
| 5 | protein_coding | 88150408 | 88156393 | ENSMUSG00000029280 | MGI:102763 | Smr3a | submaxillary gland androgen regulated protein 3A [Source:MGI Symbol;Acc:MGI:102763] |
| 5 | protein_coding | 88234415 | 88256912 | ENSMUSG00000029281 | MGI:102762 | Smr2 | submaxillary gland androgen regulated protein 2 [Source:MGI Symbol;Acc:MGI:102762] |
| 5 | protein_coding | 88416778 | 88430694 | ENSMUSG00000090302 | MGI:3646267 | Smr2l | submaxillary gland androgen regulated protein 2 like [Source:MGI Symbol;Acc:MGI:3646267] |
| 5 | protein_coding | 88465171 | 88476673 | ENSMUSG00000064156 | MGI:107496 | Prol1 | proline rich, lacrimal 1 [Source:MGI Symbol;Acc:MGI:107496] |
| 5 | protein_coding | 88523967 | 88533775 | ENSMUSG00000029282 | MGI:1918671 | Amtn | amelotin [Source:MGI Symbol;Acc:MGI:1918671] |
| 5 | protein_coding | 88603850 | 88616390 | ENSMUSG00000029288 | MGI:104655 | Ambn | ameloblastin [Source:MGI Symbol;Acc:MGI:104655] |
| 5 | protein_coding | 88635834 | 88653908 | ENSMUSG00000029286 | MGI:1333772 | Enam | enamelin [Source:MGI Symbol;Acc:MGI:1333772] |
| 5 | protein_coding | 88667668 | 88675750 | ENSMUSG00000067149 | MGI:96493 | Jchain | immunoglobulin joining chain [Source:MGI Symbol;Acc:MGI:96493] |
| 5 | protein_coding | 88702321 | 88703949 | ENSMUSG00000070697 | MGI:1919230 | Utp3 | UTP3 small subunit processome component [Source:MGI Symbol;Acc:MGI:1919230] |
| 5 | protein_coding | 88712899 | 88799251 | ENSMUSG00000029291 | MGI:106484 | Rufy3 | RUN and FYVE domain containing 3 [Source:MGI Symbol;Acc:MGI:106484] |
| 5 | protein_coding | 88807307 | 88824030 | ENSMUSG00000044221 | MGI:106479 | Grsf1 | G-rich RNA sequence binding factor 1 [Source:MGI Symbol;Acc:MGI:106479] |
| 5 | protein_coding | 88868714 | 88912079 | ENSMUSG00000006262 | MGI:1915723 | Mob1b | MOB kinase activator 1B [Source:MGI Symbol;Acc:MGI:1915723] |
| 5 | protein_coding | 88912855 | 88931140 | ENSMUSG00000029366 | MGI:102726 | Dck | deoxycytidine kinase [Source:MGI Symbol;Acc:MGI:102726] |
| 5 | protein_coding | 89034345 | 89387512 | ENSMUSG00000060961 | MGI:1927555 | Slc4a4 | [illegible] member 4 [Source:MGI Symbol;Acc:MGI:1927555] |
| 5 | protein_coding | 89565381 | 89605757 | ENSMUSG00000035540 | MGI:95669 | Gc | vitamin D binding protein [Source:MGI Symbol;Acc:MGI:95669] |
| 5 | protein_coding | 89675288 | 89731599 | ENSMUSG00000035528 | MGI:1860130 | Npffr2 | neuropeptide FF receptor 2 [Source:MGI Symbol;Acc:MGI:1860130] |

| | | | | | | | |
|---|---|---|---|---|---|---|---|
| 5 | protein_coding | 89824946 | 90031193 | ENSMUSG00000043635 | MGI:3045353 | Adamts3 | type 1 motif 3 [Source:MGI Symbol;Acc:MGI:3045353] |
| 5 | protein_coding | 90362583 | 90371860 | ENSMUSG00000035505 | MGI:2448532 | Cox18 | cytochrome c oxidase assembly protein 18 [Source:MGI Symbol;Acc:MGI:2448532] |
| 5 | protein_coding | 90375025 | 90514436 | ENSMUSG00000055204 | MGI:1932101 | Ankrd17 | ankyrin repeat domain 17 [Source:MGI Symbol;Acc:MGI:1932101] |
| 5 | protein_coding | 90608756 | 90624461 | ENSMUSG00000029368 | MGI:87991 | Alb | albumin [Source:MGI Symbol;Acc:MGI:87991] |
| 5 | protein_coding | 90638596 | 90656766 | ENSMUSG00000054932 | MGI:87951 | Afp | alpha fetoprotein [Source:MGI Symbol;Acc:MGI:87951] |
| 5 | protein_coding | 90666791 | 90701402 | ENSMUSG00000029369 | MGI:2429409 | Afm | afamin [Source:MGI Symbol;Acc:MGI:2429409] |
| 5 | protein_coding | 90708966 | 90745730 | ENSMUSG00000070690 | MGI:1923342 | Albfm1 | albumin superfamily member 1 [Source:MGI Symbol;Acc:MGI:1923342] |
| 5 | protein_coding | 90750935 | 90788516 | ENSMUSG00000029370 | MGI:1920496 | Rassf6 | member 6 [Source:MGI Symbol;Acc:MGI:1920496] |
| 5 | protein_coding | 90907219 | 90909483 | ENSMUSG00000029371 | MGI:1096868 | Cxcl5 | C-X-C motif chemokine ligand 5 [Source:MGI Symbol;Acc:MGI:1096868] |
| 5 | protein_coding | 90916377 | 90917922 | ENSMUSG00000029372 | MGI:1888712 | Ppbp | pro-platelet basic protein [Source:MGI Symbol;Acc:MGI:1888712] |
| 5 | protein_coding | 90920294 | 90921242 | ENSMUSG00000029373 | MGI:1888711 | Pf4 | platelet factor 4 [Source:MGI Symbol;Acc:MGI:1888711] |
| 5 | protein_coding | 90933962 | 90937459 | ENSMUSG00000029379 | MGI:3037818 | Cxcl3 | C-X-C motif chemokine ligand 3 [Source:MGI Symbol;Acc:MGI:3037818] |
| 5 | protein_coding | 90942393 | 90950926 | ENSMUSG00000029375 | MGI:1339941 | Cxcl15 | C-X-C motif chemokine ligand 15 [Source:MGI Symbol;Acc:MGI:1339941] |
| 5 | protein_coding | 91039100 | 91040974 | ENSMUSG00000029380 | MGI:108068 | Cxcl1 | C-X-C motif chemokine ligand 1 [Source:MGI Symbol;Acc:MGI:108068] |
| 5 | protein_coding | 91051730 | 91053797 | ENSMUSG00000058427 | MGI:1340094 | Cxcl2 | C-X-C motif chemokine ligand 2 [Source:MGI Symbol;Acc:MGI:1340094] |
| 5 | protein_coding | 91078976 | 91169227 | ENSMUSG00000029376 | MGI:1915871 | Mthfd2l | methylenetetrahydrofolate dehydrogenase (NADP+ dependent) 2-like [Source:MGI Symbol;Acc:MGI:1915871] |
| 5 | protein_coding | 91175323 | 91183074 | ENSMUSG00000035020 | MGI:1919170 | Epgn | epithelial mitogen [Source:MGI Symbol;Acc:MGI:1919170] |

| | | | | | | | |
|---|---|---|---|---|---|---|---|
| 5 | protein_coding | 91222481 | 91241505 | ENSMUSG00000029377 | MGI:107508 | Ereg | epiregulin [Source:MGI Symbol;Acc:MGI:107508] |
| 5 | protein_coding | 91287458 | 91296291 | ENSMUSG00000029378 | MGI:88068 | Areg | amphiregulin [Source:MGI Symbol;Acc:MGI:88068] |
| 5 | protein_coding | 91505120 | 91550853 | ENSMUSG00000082361 | MGI:99439 | Btc | betacellulin, epidermal growth factor family member [Source:MGI Symbol;Acc:MGI:99439] |
| 5 | protein_coding | 91665474 | 91774753 | ENSMUSG00000034981 | MGI:2443349 | Parm1 | prostate androgen-regulated mucin-like protein 1 [Source:MGI Symbol;Acc:MGI:2443349] |
| 5 | protein_coding | 92096763 | 92110927 | ENSMUSG00000029397 | MGI:1915348 | Rchy1 | [illegible] containing 1 [Source:MGI Symbol;Acc:MGI:1915348] |
| 5 | protein_coding | 92135334 | 92143179 | ENSMUSG00000096035 | MGI:2685891 | Odaph | odontogenesis associated phosphoprotein [Source:MGI Symbol;Acc:MGI:2685891] |
| 5 | protein_coding | 92153933 | 92191742 | ENSMUSG00000029403 | MGI:1858227 | Cdkl2 | cyclin dependent kinase like 2 [Source:MGI Symbol;Acc:MGI:1858227] |
| 5 | protein_coding | 92200005 | 92231578 | ENSMUSG00000029405 | MGI:2442040 | G3bp2 | G3BP stress granule assembly factor 2 [Source:MGI Symbol;Acc:MGI:2442040] |
| 5 | protein_coding | 92285797 | 92350657 | ENSMUSG00000029407 | MGI:1929095 | Uso1 | USO1 vesicle docking factor [Source:MGI Symbol;Acc:MGI:1929095] |
| 5 | protein_coding | 92357752 | 92363267 | ENSMUSG00000029409 | MGI:1930915 | U90926 | cDNA sequence U90926 [Source:MGI Symbol;Acc:MGI:1930915] |
| 5 | protein_coding | 92374538 | 92404137 | ENSMUSG00000029410 | MGI:1342304 | Ppef2 | [illegible] domain 2 [Source:MGI Symbol;Acc:MGI:1342304] |
| 5 | protein_coding | 92405518 | 92426029 | ENSMUSG00000029413 | MGI:1914361 | Naaa | N-acylethanolamine acid amidase [Source:MGI Symbol;Acc:MGI:1914361] |
| 5 | protein_coding | 92431869 | 92458338 | ENSMUSG00000029415 | MGI:2140779 | Sdad1 | SDA1 domain containing 1 [Source:MGI Symbol;Acc:MGI:2140779] |
| 5 | protein_coding | 92469206 | 92475938 | ENSMUSG00000029417 | MGI:1352449 | Cxcl9 | C-X-C motif chemokine ligand 9 [Source:MGI Symbol;Acc:MGI:1352449] |
| 5 | protein_coding | 92479686 | 92562487 | ENSMUSG00000034842 | MGI:1202729 | Art3 | ADP-ribosyltransferase 3 [Source:MGI Symbol;Acc:MGI:1202729] |
| 5 | protein_coding | 92494497 | 92496748 | ENSMUSG00000034855 | MGI:1352450 | Cxcl10 | C-X-C motif chemokine ligand 10 [Source:MGI Symbol;Acc:MGI:1352450] |
| 5 | protein_coding | 92507403 | 92513344 | ENSMUSG00000060183 | MGI:1860203 | Cxcl11 | chemokine (C-X-C motif) ligand 11 [Source:MGI Symbol;Acc:MGI:1860203] |
| 5 | protein_coding | 92563399 | 92583078 | ENSMUSG00000034826 | MGI:1920460 | Nup54 | nucleoporin 54 [Source:MGI Symbol;Acc:MGI:1920460] |

| | | | | | | | |
|---|---|---|---|---|---|---|---|
| 5 | protein_coding | 92589173 | 92654692 | ENSMUSG00000029426 | MGI:1196458 | Scarb2 | scavenger receptor class B, member 2 [Source:MGI Symbol;Acc:MGI:1196458] |
| 5 | protein_coding | 92702928 | 92739138 | ENSMUSG00000057068 | MGI:2686227 | Fam47e | family with sequence similarity 47, member E [Source:MGI Symbol;Acc:MGI:2686227] |
| 5 | protein_coding | 92750900 | 92754438 | ENSMUSG00000047963 | MGI:1261768 | Stbd1 | starch binding domain 1 [Source:MGI Symbol;Acc:MGI:1261768] |
| 5 | protein_coding | 92755813 | 92823130 | ENSMUSG00000050050 | MGI:2444555 | Ccdc158 | coiled-coil domain containing 158 [Source:MGI Symbol;Acc:MGI:2444555] |
| 5 | protein_coding | 92831294 | 93113177 | ENSMUSG00000029381 | MGI:1351655 | Shroom3 | shroom family member 3 [Source:MGI Symbol;Acc:MGI:1351655] |
| 5 | protein_coding | 93188982 | 93192881 | ENSMUSG00000045314 | MGI:1925338 | Sowahb | [illegible] member B [Source:MGI Symbol;Acc:MGI:1925338] |
| 5 | protein_coding | 93241296 | 93324306 | ENSMUSG00000058013 | MGI:1277214 | Septin11 | septin 11 [Source:MGI Symbol;Acc:MGI:1277214] |
| 5 | protein_coding | 93329792 | 93354354 | ENSMUSG00000063015 | MGI:1341077 | Ccni | cyclin I [Source:MGI Symbol;Acc:MGI:1341077] |
| 5 | protein_coding | 93354377 | 93361334 | ENSMUSG00000029384 | MGI:1922860 | 2010109A12Rik | RIKEN cDNA 2010109A12 gene [Source:MGI Symbol;Acc:MGI:1922860] |
| 5 | protein_coding | 93415116 | 93424090 | ENSMUSG00000029385 | MGI:1095734 | Ccng2 | cyclin G2 [Source:MGI Symbol;Acc:MGI:1095734] |
| 5 | protein_coding | 93629688 | 93688059 | ENSMUSG00000096044 | MGI:3704103 | Pramel33 | PRAME like 33 [Source:MGI Symbol;Acc:MGI:3704103] |
| 5 | protein_coding | 93783056 | 93819410 | ENSMUSG00000070686 | MGI:2141341 | Pramel34 | PRAME like 34 [Source:MGI Symbol;Acc:MGI:2141341] |
| 5 | protein_coding | 93991623 | 94003966 | ENSMUSG00000107392 | MGI:3704104 | Pramel35 | PRAME like 35 [Source:MGI Symbol;Acc:MGI:3704104] |
| 5 | protein_coding | 94072636 | 94076911 | ENSMUSG00000096230 | MGI:3704105 | Pramel36 | PRAME like 36 [Source:MGI Symbol;Acc:MGI:3704105] |
| 5 | protein_coding | 94214310 | 94218404 | ENSMUSG00000072822 | MGI:2681870 | Pramel37 | PRAME like 37 [Source:MGI Symbol;Acc:MGI:2681870] |
| 5 | protein_coding | 94224244 | 94228794 | ENSMUSG00000072821 | MGI:3647817 | Pramel53 | PRAME like 53 [Source:MGI Symbol;Acc:MGI:3647817] |
| 5 | protein_coding | 94304436 | 94369394 | ENSMUSG00000096259 | MGI:3781282 | Pramel38 | PRAME like 38 [Source:MGI Symbol;Acc:MGI:3781282] |
| 5 | protein_coding | 94460534 | 94465114 | ENSMUSG00000095718 | MGI:3779601 | Pramel40 | PRAME like 40 [Source:MGI Symbol;Acc:MGI:3779601] |

| | | | | | | | |
|---|---|---|---|---|---|---|---|
| 5 | protein_coding | 94591865 | 94596410 | ENSMUSG00000074011 | MGI:3779756 | Pramel41 | PRAME like 41 [Source:MGI Symbol;Acc:MGI:3779756] |
| 5 | protein_coding | 94674146 | 94694282 | ENSMUSG00000095074 | MGI:3781318 | Pramel42 | PRAME like 42 [Source:MGI Symbol;Acc:MGI:3781318] |
| 5 | protein_coding | 94759928 | 94764201 | ENSMUSG00000095503 | MGI:3781326 | Pramel43 | PRAME like 43 [Source:MGI Symbol;Acc:MGI:3781326] |
| 5 | protein_coding | 94770074 | 94774605 | ENSMUSG00000094195 | MGI:3779602 | Pramel44 | PRAME like 44 [Source:MGI Symbol;Acc:MGI:3779602] |
| 5 | protein_coding | 95013295 | 95033247 | ENSMUSG00000095954 | MGI:3781362 | Pramel56 | PRAME like 56 [Source:MGI Symbol;Acc:MGI:3781362] |
| 5 | protein_coding | 95254563 | 95259017 | ENSMUSG00000092166 | MGI:3646599 | Pramel45 | PRAME like 45 [Source:MGI Symbol;Acc:MGI:3646599] |
| 5 | protein_coding | 95330483 | 95342825 | ENSMUSG00000095996 | MGI:2141041 | Pramel60 | PRAME like 60 [Source:MGI Symbol;Acc:MGI:2141041] |
| 5 | protein_coding | 95416352 | 95420627 | ENSMUSG00000096066 | MGI:3704250 | Pramel46 | PRAME like 46 [Source:MGI Symbol;Acc:MGI:3704250] |
| 5 | protein_coding | 95453816 | 95491776 | ENSMUSG00000079424 | MGI:3781437 | Pramel47 | PRAME like 47 [Source:MGI Symbol;Acc:MGI:3781437] |
| 5 | protein_coding | 95604665 | 95633446 | ENSMUSG00000070677 | MGI:1261918 | Pramel48 | PRAME like 48 [Source:MGI Symbol;Acc:MGI:1261918] |
| 5 | protein_coding | 95658617 | 95670261 | ENSMUSG00000079423 | MGI:3781464 | Pramel57 | PRAME like 57 [Source:MGI Symbol;Acc:MGI:3781464] |
| 5 | protein_coding | 95738458 | 95742734 | ENSMUSG00000094043 | MGI:3779772 | Pramel49 | PRAME like 49 [Source:MGI Symbol;Acc:MGI:3779772] |
| 5 | protein_coding | 95879648 | 95884032 | ENSMUSG00000096139 | MGI:3704107 | Pramel50 | PRAME like 50 [Source:MGI Symbol;Acc:MGI:3704107] |
| 5 | protein_coding | 95889873 | 95894414 | ENSMUSG00000072814 | MGI:3648891 | Pramel54 | PRAME like 54 [Source:MGI Symbol;Acc:MGI:3648891] |
| 5 | protein_coding | 95923729 | 95952089 | ENSMUSG00000072813 | MGI:3615333 | Pramel55 | PRAME like 55 [Source:MGI Symbol;Acc:MGI:3615333] |
| 5 | protein_coding | 96104810 | 96108927 | ENSMUSG00000023078 | MGI:1888499 | Cxcl13 | C-X-C motif chemokine ligand 13 [Source:MGI Symbol;Acc:MGI:1888499] |
| 5 | protein_coding | 96218192 | 96312030 | ENSMUSG00000034724 | MGI:2443154 | Cnot6l | CCR4-NOT transcription complex, subunit 6-like [Source:MGI Symbol;Acc:MGI:2443154] |
| 5 | protein_coding | 96357352 | 96414586 | ENSMUSG00000029486 | MGI:2137202 | Mrpl1 | mitochondrial ribosomal protein L1 [Source:MGI Symbol;Acc:MGI:2137202] |

| | | | | | | | |
|---|---|---|---|---|---|---|---|
| 5 | protein_coding | 96521814 | 96932587 | ENSMUSG00000034687 | MGI:2385368 | Fras1 | Fraser extracellular matrix complex subunit 1 [Source:MGI Symbol;Acc:MGI:2385368] |
| 5 | protein_coding | 96941198 | 96993825 | ENSMUSG00000029484 | MGI:1201378 | Anxa3 | annexin A3 [Source:MGI Symbol;Acc:MGI:1201378] |
| 5 | protein_coding | 97145548 | 97239726 | ENSMUSG00000034663 | MGI:2155456 | Bmp2k | BMP2 inducible kinase [Source:MGI Symbol;Acc:MGI:2155456] |
| 5 | protein_coding | 97230188 | 97259455 | ENSMUSG00000055725 | MGI:2679683 | Paqr3 | progestin and adipoQ receptor family member III [Source:MGI Symbol;Acc:MGI:2679683] |
| 5 | protein_coding | 97530057 | 97540238 | ENSMUSG00000046000 | MGI:2141314 | Naa11 | [illegible] subunit [Source:MGI Symbol;Acc:MGI:2141314] |
| 5 | protein_coding | 97603001 | 97604880 | ENSMUSG00000050553 | MGI:1329027 | Gk2 | glycerol kinase 2 [Source:MGI Symbol;Acc:MGI:1329027] |
| 5 | protein_coding | 98030642 | 98178902 | ENSMUSG00000029338 | MGI:1919164 | Antxr2 | anthrax toxin receptor 2 [Source:MGI Symbol;Acc:MGI:1919164] |
| 5 | protein_coding | 98315057 | 98336850 | ENSMUSG00000035456 | MGI:1924880 | Prdm8 | PR domain containing 8 [Source:MGI Symbol;Acc:MGI:1924880] |
| 5 | protein_coding | 98402043 | 98424889 | ENSMUSG00000029337 | MGI:95519 | Fgf5 | fibroblast growth factor 5 [Source:MGI Symbol;Acc:MGI:95519] |
| 5 | protein_coding | 98477163 | 98949906 | ENSMUSG00000057816 | MGI:1916571 | Cfap299 | cilia and flagella associated protein 299 [Source:MGI Symbol;Acc:MGI:1916571] |
| 5 | protein_coding | 99002274 | 99032255 | ENSMUSG00000029335 | MGI:88179 | Bmp3 | bone morphogenetic protein 3 [Source:MGI Symbol;Acc:MGI:88179] |
| 5 | protein_coding | 99077632 | 99185210 | ENSMUSG00000029334 | MGI:108173 | Prkg2 | protein kinase, cGMP-dependent, type II [Source:MGI Symbol;Acc:MGI:108173] |
| 5 | protein_coding | 99365285 | 99876924 | ENSMUSG00000089809 | MGI:2443755 | Rasgef1b | RasGEF domain family, member 1B [Source:MGI Symbol;Acc:MGI:2443755] |
| 5 | protein_coding | 1.00E+08 | 1.00E+08 | ENSMUSG00000105078 | MGI:5595070 | Vamp9 | vesicle-associated membrane protein 9 [Source:MGI Symbol;Acc:MGI:5595070] |
| 5 | protein_coding | 1.00E+08 | 1.00E+08 | ENSMUSG00000000568 | MGI:101947 | Hnrnpd | heterogeneous nuclear ribonucleoprotein D [Source:MGI Symbol;Acc:MGI:101947] |
| 5 | protein_coding | 1.00E+08 | 1.00E+08 | ENSMUSG00000029328 | MGI:1355299 | Hnrnpdl | heterogeneous nuclear ribonucleoprotein D-like [Source:MGI Symbol;Acc:MGI:1355299] |
| 5 | protein_coding | 1.00E+08 | 1.00E+08 | ENSMUSG00000029326 | MGI:1915120 | Enoph1 | enolase-phosphatase 1 [Source:MGI Symbol;Acc:MGI:1915120] |
| 5 | protein_coding | 1.00E+08 | 1.00E+08 | ENSMUSG00000050640 | MGI:3041258 | Tmem150c | transmembrane protein 150C [Source:MGI Symbol;Acc:MGI:3041258] |

| | | | | | | | |
|---|---|---|---|---|---|---|---|
| 5 | protein_coding | 1.01E+08 | 1.01E+08 | ENSMUSG00000035325 | MGI:1916412 | Sec31a | [illegible] component [Source:MGI Symbol;Acc:MGI:1916412] |
| 5 | protein_coding | 1.01E+08 | 1.01E+08 | ENSMUSG00000118665 | MGI:2140902 | Lin54 | lin-54 DREAM MuvB core complex component [Source:MGI Symbol;Acc:MGI:2140902] |
| 5 | protein_coding | 1.01E+08 | 1.01E+08 | ENSMUSG00000035297 | MGI:1349414 | Cops4 | COP9 signalosome subunit 4 [Source:MGI Symbol;Acc:MGI:1349414] |
| 5 | protein_coding | 1.01E+08 | 1.01E+08 | ENSMUSG00000029322 | MGI:2445289 | Plac8 | placenta-specific 8 [Source:MGI Symbol;Acc:MGI:2445289] |
| 5 | protein_coding | 1.01E+08 | 1.01E+08 | ENSMUSG00000029319 | MGI:1919133 | Coq2 | coenzyme Q2 4-hydroxybenzoate polyprenyltransferase [Source:MGI Symbol;Acc:MGI:1919133] |
| 5 | protein_coding | 1.01E+08 | 1.01E+08 | ENSMUSG00000035273 | MGI:1343124 | Hpse | heparanase [Source:MGI Symbol;Acc:MGI:1343124] |
| 5 | protein_coding | 1.01E+08 | 1.01E+08 | ENSMUSG00000035266 | MGI:2176740 | Helq | helicase, POLQ-like [Source:MGI Symbol;Acc:MGI:2176740] |
| 5 | protein_coding | 1.01E+08 | 1.01E+08 | ENSMUSG00000016833 | MGI:1915985 | Mrps18c | mitochondrial ribosomal protein S18C [Source:MGI Symbol;Acc:MGI:1915985] |
| 5 | protein_coding | 1.01E+08 | 1.01E+08 | ENSMUSG00000035234 | MGI:1917931 | Abraxas1 | BRCA1 A complex subunit [Source:MGI Symbol;Acc:MGI:1917931] |
| 5 | protein_coding | 1.01E+08 | 1.01E+08 | ENSMUSG00000029314 | MGI:3603816 | Gpat3 | glycerol-3-phosphate acyltransferase 3 [Source:MGI Symbol;Acc:MGI:3603816] |
| 5 | protein_coding | 1.02E+08 | 1.02E+08 | ENSMUSG00000035187 | MGI:1206039 | Nkx6-1 | NK6 homeobox 1 [Source:MGI Symbol;Acc:MGI:1206039] |
| 5 | protein_coding | 1.02E+08 | 1.02E+08 | ENSMUSG00000029330 | MGI:1921846 | Cds1 | CDP-diacylglycerol synthase 1 [Source:MGI Symbol;Acc:MGI:1921846] |
| 5 | protein_coding | 1.02E+08 | 1.02E+08 | ENSMUSG00000043940 | MGI:1096875 | Wdfy3 | WD repeat and FYVE domain containing 3 [Source:MGI Symbol;Acc:MGI:1096875] |
| 5 | protein_coding | 1.03E+08 | 1.03E+08 | ENSMUSG00000057315 | MGI:1922647 | Arhgap24 | Rho GTPase activating protein 24 [Source:MGI Symbol;Acc:MGI:1922647] |
| 5 | protein_coding | 1.03E+08 | 1.03E+08 | ENSMUSG00000046709 | MGI:1346863 | Mapk10 | mitogen-activated protein kinase 10 [Source:MGI Symbol;Acc:MGI:1346863] |
| 5 | protein_coding | 1.04E+08 | 1.04E+08 | ENSMUSG00000034573 | MGI:103293 | Ptpn13 | protein tyrosine phosphatase, non-receptor type 13 [Source:MGI Symbol;Acc:MGI:103293] |
| 5 | protein_coding | 1.04E+08 | 1.04E+08 | ENSMUSG00000029321 | MGI:1923000 | Slc10a6 | solute carrier family 10 (sodium/bile acid cotransporter family), member 6 [Source:MGI Symbol;Acc:MGI:1923000] |

| | | | | | | | |
|---|---|---|---|---|---|---|---|
| 5 | protein_coding | 1.04E+08 | 1.04E+08 | ENSMUSG00000029320 | MGI:1921468 | 1700016H13Rik | RIKEN cDNA 1700016H13 gene [Source:MGI Symbol;Acc:MGI:1921468] |
| 5 | protein_coding | 1.04E+08 | 1.04E+08 | ENSMUSG00000029313 | MGI:1100819 | Aff1 | AF4/FMR2 family, member 1 [Source:MGI Symbol;Acc:MGI:1100819] |
| 5 | protein_coding | 1.04E+08 | 1.04E+08 | ENSMUSG00000029312 | MGI:2179430 | Klhl8 | kelch-like 8 [Source:MGI Symbol;Acc:MGI:2179430] |
| 5 | protein_coding | 1.04E+08 | 1.04E+08 | ENSMUSG00000034528 | MGI:2140804 | Hsd17b13 | hydroxysteroid (17-beta) dehydrogenase 13 [Source:MGI Symbol;Acc:MGI:2140804] |
| 5 | protein_coding | 1.04E+08 | 1.04E+08 | ENSMUSG00000029311 | MGI:2149821 | Hsd17b11 | hydroxysteroid (17-beta) dehydrogenase 11 [Source:MGI Symbol;Acc:MGI:2149821] |
| 5 | protein_coding | 1.04E+08 | 1.04E+08 | ENSMUSG00000029310 | MGI:1921417 | Nudt9 | nudix hydrolase 9 [Source:MGI Symbol;Acc:MGI:1921417] |
| 5 | protein_coding | 1.04E+08 | 1.04E+08 | ENSMUSG00000091034 | MGI:4937294 | Scpppq1 | secretory calcium-binding phosphoprotein proline-glutamine rich 1 [Source:MGI Symbol;Acc:MGI:4937294] |
| 5 | protein_coding | 1.04E+08 | 1.04E+08 | ENSMUSG00000029309 | MGI:108110 | Sparcl1 | SPARC-like 1 [Source:MGI Symbol;Acc:MGI:108110] |
| 5 | protein_coding | 1.04E+08 | 1.04E+08 | ENSMUSG00000053268 | MGI:109172 | Dspp | dentin sialophosphoprotein [Source:MGI Symbol;Acc:MGI:109172] |
| 5 | protein_coding | 1.04E+08 | 1.04E+08 | ENSMUSG00000029307 | MGI:94910 | Dmp1 | dentin matrix protein 1 [Source:MGI Symbol;Acc:MGI:94910] |
| 5 | protein_coding | 1.04E+08 | 1.04E+08 | ENSMUSG00000029306 | MGI:96389 | Ibsp | integrin binding sialoprotein [Source:MGI Symbol;Acc:MGI:96389] |
| 5 | protein_coding | 1.04E+08 | 1.04E+08 | ENSMUSG00000053863 | MGI:2137384 | Mepe | matrix extracellular phosphoglycoprotein with ASARM motif (bone) [Source:MGI Symbol;Acc:MGI:2137384] |
| 5 | protein_coding | 1.05E+08 | 1.05E+08 | ENSMUSG00000029304 | MGI:98389 | Spp1 | secreted phosphoprotein 1 [Source:MGI Symbol;Acc:MGI:98389] |
| 5 | protein_coding | 1.05E+08 | 1.05E+08 | ENSMUSG00000034462 | MGI:1099818 | Pkd2 | [illegible] channel [Source:MGI Symbol;Acc:MGI:1099818] |
| 5 | protein_coding | 1.05E+08 | 1.05E+08 | ENSMUSG00000097392 | MGI:3040669 | Thoc2l | THO complex subunit 2-like [Source:MGI Symbol;Acc:MGI:3040669] |
| 5 | protein_coding | 1.05E+08 | 1.05E+08 | ENSMUSG00000120166 | MGI:1916873 | Zfp33b | zinc finger protein 33B [Source:MGI Symbol;Acc:MGI:1916873] |
| 5 | protein_coding | 1.05E+08 | 1.05E+08 | ENSMUSG00000072774 | MGI:2441896 | Zfp951 | zinc finger protein 951 [Source:MGI Symbol;Acc:MGI:2441896] |

| | | | | | | | |
|---|---|---|---|---|---|---|---|
| 5 | protein_coding | 1.05E+08 | 1.05E+08 | ENSMUSG00000029299 | MGI:1351624 | Abcg3 | ATP binding cassette subfamily G member 3 [Source:MGI Symbol;Acc:MGI:1351624] |
| 5 | protein_coding | 1.05E+08 | 1.05E+08 | ENSMUSG00000034438 | MGI:1923324 | Gbp8 | guanylate-binding protein 8 [Source:MGI Symbol;Acc:MGI:1923324] |
| 5 | protein_coding | 1.05E+08 | 1.05E+08 | ENSMUSG00000029298 | MGI:3605620 | Gbp9 | guanylate-binding protein 9 [Source:MGI Symbol;Acc:MGI:3605620] |
| 5 | protein_coding | 1.05E+08 | 1.05E+08 | ENSMUSG00000079363 | MGI:97072 | Gbp4 | guanylate binding protein 4 [Source:MGI Symbol;Acc:MGI:97072] |
| 5 | protein_coding | 1.05E+08 | 1.05E+08 | ENSMUSG00000079362 | MGI:5663439 | Gm43302 | predicted gene 43302 [Source:MGI Symbol;Acc:MGI:5663439] |
| 5 | protein_coding | 1.05E+08 | 1.05E+08 | ENSMUSG00000105096 | MGI:4359647 | Gbp10 | guanylate-binding protein 10 [Source:MGI Symbol;Acc:MGI:4359647] |
| 5 | protein_coding | 1.05E+08 | 1.05E+08 | ENSMUSG00000104713 | MGI:2140937 | Gbp6 | guanylate binding protein 6 [Source:MGI Symbol;Acc:MGI:2140937] |
| 5 | protein_coding | 1.05E+08 | 1.05E+08 | ENSMUSG00000092021 | MGI:3646307 | Gbp11 | guanylate binding protein 11 [Source:MGI Symbol;Acc:MGI:3646307] |
| 5 | protein_coding | 1.06E+08 | 1.06E+08 | ENSMUSG00000070639 | MGI:2141353 | Lrrc8b | [illegible] member B [Source:MGI Symbol;Acc:MGI:2141353] |
| 5 | protein_coding | 1.06E+08 | 1.06E+08 | ENSMUSG00000054720 | MGI:2140839 | Lrrc8c | [illegible] member C [Source:MGI Symbol;Acc:MGI:2140839] |
| 5 | protein_coding | 1.06E+08 | 1.06E+08 | ENSMUSG00000046079 | MGI:1922368 | Lrrc8d | leucine rich repeat containing 8D [Source:MGI Symbol;Acc:MGI:1922368] |
| 5 | protein_coding | 1.06E+08 | 1.06E+08 | ENSMUSG00000029290 | MGI:1927246 | Zfp326 | zinc finger protein 326 [Source:MGI Symbol;Acc:MGI:1927246] |
| 5 | protein_coding | 1.07E+08 | 1.07E+08 | ENSMUSG00000034384 | MGI:1859314 | Barhl2 | BarH like homeobox 2 [Source:MGI Symbol;Acc:MGI:1859314] |
| 5 | protein_coding | 1.07E+08 | 1.07E+08 | ENSMUSG00000049606 | MGI:1277212 | Zfp644 | zinc finger protein 644 [Source:MGI Symbol;Acc:MGI:1277212] |
| 5 | protein_coding | 1.07E+08 | 1.07E+08 | ENSMUSG00000043410 | MGI:3036246 | Hfm1 | HFM1, ATP-dependent DNA helicase homolog [Source:MGI Symbol;Acc:MGI:3036246] |
| 5 | protein_coding | 1.07E+08 | 1.07E+08 | ENSMUSG00000029283 | MGI:1309511 | Cdc7 | cell division cycle 7 [Source:MGI Symbol;Acc:MGI:1309511] |
| 5 | protein_coding | 1.07E+08 | 1.07E+08 | ENSMUSG00000029287 | MGI:104637 | Tgfbr3 | transforming growth factor, beta receptor III [Source:MGI Symbol;Acc:MGI:104637] |
| 5 | protein_coding | 1.07E+08 | 1.08E+08 | ENSMUSG00000029279 | MGI:1891374 | Brdt | bromodomain, testis-specific [Source:MGI Symbol;Acc:MGI:1891374] |

| | | | | | | | |
|---|---|---|---|---|---|---|---|
| 5 | protein_coding | 1.08E+08 | 1.08E+08 | ENSMUSG00000033805 | MGI:2686228 | Ephx4 | epoxide hydrolase 4 [Source:MGI Symbol;Acc:MGI:2686228] |
| 5 | protein_coding | 1.08E+08 | 1.08E+08 | ENSMUSG00000033794 | MGI:1918152 | Lpcat2b | lysophosphatidylcholine acyltransferase 2B [Source:MGI Symbol;Acc:MGI:1918152] |
| 5 | protein_coding | 1.08E+08 | 1.08E+08 | ENSMUSG00000111375 | MGI:3646208 | Btbd8 | BTB domain containing 8 [Source:MGI Symbol;Acc:MGI:3646208] |
| 5 | protein_coding | 1.08E+08 | 1.08E+08 | ENSMUSG00000106631 | MGI:5662806 | Gm42669 | predicted gene 42669 [Source:MGI Symbol;Acc:MGI:5662806] |
| 5 | protein_coding | 1.08E+08 | 1.08E+08 | ENSMUSG00000089798 | MGI:1923671 | 1700028K03Rik | RIKEN cDNA 1700028K03 gene [Source:MGI Symbol;Acc:MGI:1923671] |
| 5 | protein_coding | 1.08E+08 | 1.08E+08 | ENSMUSG00000029276 | MGI:2141180 | Glmn | glomulin, FKBP associated protein [Source:MGI Symbol;Acc:MGI:2141180] |
| 5 | protein_coding | 1.08E+08 | 1.08E+08 | ENSMUSG00000033773 | MGI:2141142 | Rpap2 | RNA polymerase II associated protein 2 [Source:MGI Symbol;Acc:MGI:2141142] |
| 5 | protein_coding | 1.08E+08 | 1.08E+08 | ENSMUSG00000029275 | MGI:103170 | Gfi1 | [illegible] repressor [Source:MGI Symbol;Acc:MGI:103170] |
| 5 | protein_coding | 1.08E+08 | 1.08E+08 | ENSMUSG00000011831 | MGI:104736 | Evi5 | ecotropic viral integration site 5 [Source:MGI Symbol;Acc:MGI:104736] |
| 5 | protein_coding | 1.08E+08 | 1.08E+08 | ENSMUSG00000063447 | MGI:1920568 | Ube2d2b | ubiquitin-conjugating enzyme E2D 2B [Source:MGI Symbol;Acc:MGI:1920568] |
| 5 | protein_coding | 1.08E+08 | 1.08E+08 | ENSMUSG00000058558 | MGI:102854 | Rpl5 | ribosomal protein L5 [Source:MGI Symbol;Acc:MGI:102854] |
| 5 | protein_coding | 1.08E+08 | 1.08E+08 | ENSMUSG00000029270 | MGI:1914516 | Dipk1a | divergent protein kinase domain 1A [Source:MGI Symbol;Acc:MGI:1914516] |
| 5 | protein_coding | 1.08E+08 | 1.08E+08 | ENSMUSG00000029267 | MGI:105050 | Mtf2 | metal response element binding transcription factor 2 [Source:MGI Symbol;Acc:MGI:105050] |
| 5 | protein_coding | 1.08E+08 | 1.08E+08 | ENSMUSG00000063406 | MGI:1921586 | Tmed5 | transmembrane p24 trafficking protein 5 [Source:MGI Symbol;Acc:MGI:1921586] |
| 5 | protein_coding | 1.08E+08 | 1.08E+08 | ENSMUSG00000056531 | MGI:1922974 | Ccdc18 | coiled-coil domain containing 18 [Source:MGI Symbol;Acc:MGI:1922974] |
| 5 | protein_coding | 1.08E+08 | 1.08E+08 | ENSMUSG00000029265 | MGI:1100515 | Dr1 | down-regulator of transcription 1 [Source:MGI Symbol;Acc:MGI:1100515] |
| 5 | protein_coding | 1.08E+08 | 1.08E+08 | ENSMUSG00000029263 | MGI:3576484 | Pigg | [illegible] biosynthesis, class G [Source:MGI Symbol;Acc:MGI:3576484] |
| 5 | protein_coding | 1.09E+08 | 1.09E+08 | ENSMUSG00000105867 | MGI:5662654 | Tmem271rt | transmembrane protein 271, retrotransposed [Source:MGI Symbol;Acc:MGI:5662654] |

| | | | | | | | |
|---|---|---|---|---|---|---|---|
| 5 | protein_coding | 1.09E+08 | 1.09E+08 | ENSMUSG00000029491 | MGI:97525 | Pde6b | beta polypeptide [Source:MGI Symbol;Acc:MGI:97525] |
| 5 | protein_coding | 1.09E+08 | 1.09E+08 | ENSMUSG00000050856 | MGI:106636 | Atp5me | ATP synthase membrane subunit e [Source:MGI Symbol;Acc:MGI:106636] |
| 5 | protein_coding | 1.09E+08 | 1.09E+08 | ENSMUSG00000029490 | MGI:2442629 | Slc49a3 | solute carrier family 49 member 3 [Source:MGI Symbol;Acc:MGI:2442629] |
| 5 | protein_coding | 1.09E+08 | 1.09E+08 | ENSMUSG00000033623 | MGI:1916837 | Pcgf3 | polycomb group ring finger 3 [Source:MGI Symbol;Acc:MGI:1916837] |
| 5 | protein_coding | 1.09E+08 | 1.09E+08 | ENSMUSG00000033615 | MGI:104727 | Cplx1 | complexin 1 [Source:MGI Symbol;Acc:MGI:104727] |
| 5 | protein_coding | 1.09E+08 | 1.09E+08 | ENSMUSG00000062234 | MGI:2442153 | Gak | cyclin G associated kinase [Source:MGI Symbol;Acc:MGI:2442153] |
| 5 | protein_coding | 1.09E+08 | 1.09E+08 | ENSMUSG00000013495 | MGI:1919642 | Tmem175 | transmembrane protein 175 [Source:MGI Symbol;Acc:MGI:1919642] |
| 5 | protein_coding | 1.09E+08 | 1.09E+08 | ENSMUSG00000004815 | MGI:102918 | Dgkq | diacylglycerol kinase, theta [Source:MGI Symbol;Acc:MGI:102918] |
| 5 | protein_coding | 1.09E+08 | 1.09E+08 | ENSMUSG00000033540 | MGI:96418 | Idua | iduronidase, alpha-L [Source:MGI Symbol;Acc:MGI:96418] |
| 5 | protein_coding | 1.09E+08 | 1.09E+08 | ENSMUSG00000046959 | MGI:2385894 | Slc26a1 | member 1 [Source:MGI Symbol;Acc:MGI:2385894] |
| 5 | protein_coding | 1.09E+08 | 1.09E+08 | ENSMUSG00000008090 | MGI:2150920 | Fgfrl1 | fibroblast growth factor receptor-like 1 [Source:MGI Symbol;Acc:MGI:2150920] |
| 5 | protein_coding | 1.09E+08 | 1.09E+08 | ENSMUSG00000055385 | MGI:3645767 | Rnf212 | ring finger protein 212 [Source:MGI Symbol;Acc:MGI:3645767] |
| 5 | protein_coding | 1.09E+08 | 1.09E+08 | ENSMUSG00000004821 | MGI:1914616 | Tmed11 | transmembrane p24 trafficking protein 11 [Source:MGI Symbol;Acc:MGI:1914616] |
| 5 | protein_coding | 1.09E+08 | 1.09E+08 | ENSMUSG00000090961 | MGI:3642986 | Vmn2r8 | vomeronasal 2, receptor 8 [Source:MGI Symbol;Acc:MGI:3642986] |
| 5 | protein_coding | 1.09E+08 | 1.09E+08 | ENSMUSG00000091624 | MGI:3643093 | Vmn2r9 | vomeronasal 2, receptor 9 [Source:MGI Symbol;Acc:MGI:3643093] |
| 5 | protein_coding | 1.09E+08 | 1.09E+08 | ENSMUSG00000067010 | MGI:1316730 | Vmn2r10 | vomeronasal 2, receptor 10 [Source:MGI Symbol;Acc:MGI:1316730] |
| 5 | protein_coding | 1.09E+08 | 1.09E+08 | ENSMUSG00000091450 | MGI:3643806 | Vmn2r11 | vomeronasal 2, receptor 11 [Source:MGI Symbol;Acc:MGI:3643806] |
| 5 | protein_coding | 1.09E+08 | 1.09E+08 | ENSMUSG00000090688 | MGI:3761377 | Vmn2r12 | vomeronasal 2, receptor 12 [Source:MGI Symbol;Acc:MGI:3761377] |

| | | | | | | | |
|---|---|---|---|---|---|---|---|
| 5 | protein_coding | 1.09E+08 | 1.09E+08 | ENSMUSG00000091635 | MGI:3761379 | Vmn2r13 | vomeronasal 2, receptor 13 [Source:MGI Symbol;Acc:MGI:3761379] |
| 5 | protein_coding | 1.09E+08 | 1.09E+08 | ENSMUSG00000091059 | MGI:3649151 | Vmn2r14 | vomeronasal 2, receptor 14 [Source:MGI Symbol;Acc:MGI:3649151] |
| 5 | protein_coding | 1.09E+08 | 1.09E+08 | ENSMUSG00000091375 | MGI:3649165 | Vmn2r15 | vomeronasal 2, receptor 15 [Source:MGI Symbol;Acc:MGI:3649165] |
| 5 | protein_coding | 1.09E+08 | 1.10E+08 | ENSMUSG00000092080 | MGI:3647194 | Vmn2r16 | vomeronasal 2, receptor 16 [Source:MGI Symbol;Acc:MGI:3647194] |
| 5 | protein_coding | 1.10E+08 | 1.10E+08 | ENSMUSG00000091879 | MGI:3647193 | Vmn2r17 | vomeronasal 2, receptor 17 [Source:MGI Symbol;Acc:MGI:3647193] |
| 5 | protein_coding | 1.10E+08 | 1.10E+08 | ENSMUSG00000033467 | MGI:1889506 | Crlf2 | cytokine receptor-like factor 2 [Source:MGI Symbol;Acc:MGI:1889506] |
| 5 | protein_coding | 1.10E+08 | 1.10E+08 | ENSMUSG00000072763 | MGI:1924450 | Zfp1007 | zinc finger protein 1007 [Source:MGI Symbol;Acc:MGI:1924450] |
| 5 | protein_coding | 1.10E+08 | 1.10E+08 | ENSMUSG00000072762 | MGI:1925270 | 4930522L14Rik | RIKEN cDNA 4930522L14 gene [Source:MGI Symbol;Acc:MGI:1925270] |
| 5 | protein_coding | 1.10E+08 | 1.10E+08 | ENSMUSG00000121596 | MGI:3642128 | AB010352 | cDNA sequence AB010352 [Source:MGI Symbol;Acc:MGI:3642128] |
| 5 | protein_coding | 1.10E+08 | 1.10E+08 | ENSMUSG00000090015 | MGI:3709333 | Gm15446 | predicted gene 15446 [Source:MGI Symbol;Acc:MGI:3709333] |
| 5 | protein_coding | 1.10E+08 | 1.10E+08 | ENSMUSG00000121597 | | | novel KRAB containing domain zinc finger protein |
| 5 | protein_coding | 1.10E+08 | 1.10E+08 | ENSMUSG00000066613 | MGI:1916754 | Zfp932 | zinc finger protein 932 [Source:MGI Symbol;Acc:MGI:1916754] |
| 5 | protein_coding | 1.10E+08 | 1.10E+08 | ENSMUSG00000090963 | MGI:4937289 | Gm17655 | predicted gene, 17655 [Source:MGI Symbol;Acc:MGI:4937289] |
| 5 | protein_coding | 1.10E+08 | 1.10E+08 | ENSMUSG00000109771 | MGI:5594474 | Gm35315 | predicted gene, 35315 [Source:MGI Symbol;Acc:MGI:5594474] |
| 5 | protein_coding | 1.10E+08 | 1.10E+08 | ENSMUSG00000064247 | MGI:2685422 | Plcxd1 | phosphatidylinositol-specific phospholipase C, X domain containing 1 [Source:MGI Symbol;Acc:MGI:2685422] |
| 5 | protein_coding | 1.10E+08 | 1.10E+08 | ENSMUSG00000033434 | MGI:1306825 | Gtpbp6 | GTP binding protein 6 (putative) [Source:MGI Symbol;Acc:MGI:1306825] |
| 5 | protein_coding | 1.10E+08 | 1.10E+08 | ENSMUSG00000023284 | MGI:2444933 | Zfp605 | zinc finger protein 605 [Source:MGI Symbol;Acc:MGI:2444933] |

| | | | | | | | |
|---|---|---|---|---|---|---|---|
| 5 | protein_coding | 1.10E+08 | 1.10E+08 | ENSMUSG00000014668 | MGI:2444898 | Chfr | [illegible] domains [Source:MGI Symbol;Acc:MGI:2444898] |
| 5 | protein_coding | 1.10E+08 | 1.10E+08 | ENSMUSG00000029502 | MGI:96958 | Golga3 | golgin A3 [Source:MGI Symbol;Acc:MGI:96958] |
| 5 | protein_coding | 1.10E+08 | 1.10E+08 | ENSMUSG00000029501 | MGI:1261856 | Ankle2 | ankyrin repeat and LEM domain containing 2 [Source:MGI Symbol;Acc:MGI:1261856] |
| 5 | protein_coding | 1.10E+08 | 1.10E+08 | ENSMUSG00000029500 | MGI:1919792 | Pgam5 | phosphoglycerate mutase family member 5 [Source:MGI Symbol;Acc:MGI:1919792] |
| 5 | protein_coding | 1.10E+08 | 1.10E+08 | ENSMUSG00000029499 | MGI:107487 | Pxmp2 | peroxisomal membrane protein 2 [Source:MGI Symbol;Acc:MGI:107487] |
| 5 | protein_coding | 1.10E+08 | 1.10E+08 | ENSMUSG00000007080 | MGI:1196391 | Pole | polymerase (DNA directed), epsilon [Source:MGI Symbol;Acc:MGI:1196391] |
| 5 | protein_coding | 1.10E+08 | 1.10E+08 | ENSMUSG00000029503 | MGI:2665170 | P2rx2 | [illegible] channel, 2 [Source:MGI Symbol;Acc:MGI:2665170] |
| 5 | protein_coding | 1.10E+08 | 1.11E+08 | ENSMUSG00000072754 | MGI:2686525 | Lrcol1 | leucine rich colipase-like 1 [Source:MGI Symbol;Acc:MGI:2686525] |
| 5 | protein_coding | 1.11E+08 | 1.11E+08 | ENSMUSG00000043323 | MGI:1920907 | Fbrsl1 | fibrosin-like 1 [Source:MGI Symbol;Acc:MGI:1920907] |
| 5 | protein_coding | 1.11E+08 | 1.11E+08 | ENSMUSG00000033316 | MGI:2677965 | Galnt9 | polypeptide N-acetylgalactosaminyltransferase 9 [Source:MGI Symbol;Acc:MGI:2677965] |
| 5 | protein_coding | 1.11E+08 | 1.11E+08 | ENSMUSG00000033294 | MGI:2140843 | Noc4l | NOC4 like [Source:MGI Symbol;Acc:MGI:2140843] |
| 5 | protein_coding | 1.11E+08 | 1.11E+08 | ENSMUSG00000029504 | MGI:1916913 | Ddx51 | DEAD box helicase 51 [Source:MGI Symbol;Acc:MGI:1916913] |
| 5 | protein_coding | 1.11E+08 | 1.11E+08 | ENSMUSG00000029505 | MGI:1276124 | Ep400 | E1A binding protein p400 [Source:MGI Symbol;Acc:MGI:1276124] |
| 5 | protein_coding | 1.11E+08 | 1.11E+08 | ENSMUSG00000029507 | MGI:1929237 | Pus1 | pseudouridine synthase 1 [Source:MGI Symbol;Acc:MGI:1929237] |
| 5 | protein_coding | 1.11E+08 | 1.11E+08 | ENSMUSG00000029512 | MGI:1270126 | Ulk1 | unc-51 like kinase 1 [Source:MGI Symbol;Acc:MGI:1270126] |
| 5 | protein_coding | 1.11E+08 | 1.11E+08 | ENSMUSG00000043510 | MGI:2141135 | Hscb | HscB iron-sulfur cluster co-chaperone [Source:MGI Symbol;Acc:MGI:2141135] |
| 5 | protein_coding | 1.11E+08 | 1.11E+08 | ENSMUSG00000029521 | MGI:1355321 | Chek2 | checkpoint kinase 2 [Source:MGI Symbol;Acc:MGI:1355321] |
| 5 | protein_coding | 1.11E+08 | 1.11E+08 | ENSMUSG00000033209 | MGI:2140873 | Ttc28 | tetratricopeptide repeat domain 28 [Source:MGI Symbol;Acc:MGI:2140873] |

| | | | | | | | |
|---|---|---|---|---|---|---|---|
| 5 | protein_coding | 1.11E+08 | 1.12E+08 | ENSMUSG00000050017 | MGI:1927542 | Pitpnb | phosphatidylinositol transfer protein, beta [Source:MGI Symbol;Acc:MGI:1927542] |
| 5 | protein_coding | 1.12E+08 | 1.12E+08 | ENSMUSG00000070576 | MGI:1261813 | Mn1 | meningioma 1 [Source:MGI Symbol;Acc:MGI:1261813] |
| 5 | protein_coding | 1.12E+08 | 1.12E+08 | ENSMUSG00000096949 | MGI:5477391 | Gm26897 | predicted gene, 26897 [Source:MGI Symbol;Acc:MGI:5477391] |
| 5 | protein_coding | 1.12E+08 | 1.12E+08 | ENSMUSG00000066975 | MGI:102716 | Cryba4 | crystallin, beta A4 [Source:MGI Symbol;Acc:MGI:102716] |
| 5 | protein_coding | 1.12E+08 | 1.12E+08 | ENSMUSG00000029343 | MGI:104992 | Crybb1 | crystallin, beta B1 [Source:MGI Symbol;Acc:MGI:104992] |
| 5 | protein_coding | 1.12E+08 | 1.12E+08 | ENSMUSG00000029344 | MGI:1309516 | Tpst2 | protein-tyrosine sulfotransferase 2 [Source:MGI Symbol;Acc:MGI:1309516] |
| 5 | protein_coding | 1.12E+08 | 1.12E+08 | ENSMUSG00000029345 | MGI:1930075 | Tfip11 | tuftelin interacting protein 11 [Source:MGI Symbol;Acc:MGI:1930075] |
| 5 | protein_coding | 1.12E+08 | 1.12E+08 | ENSMUSG00000029346 | MGI:1917368 | Srrd | SRR1 domain containing [Source:MGI Symbol;Acc:MGI:1917368] |
| 5 | protein_coding | 1.12E+08 | 1.13E+08 | ENSMUSG00000042328 | MGI:2177742 | Hps4 | HPS4, biogenesis of lysosomal organelles complex 3 subunit 2 [Source:MGI Symbol;Acc:MGI:2177742] |
| 5 | protein_coding | 1.13E+08 | 1.13E+08 | ENSMUSG00000051503 | MGI:3648511 | Ccdc121rt3 | coiled-coil domain containing 121, retrogene 3 [Source:MGI Symbol;Acc:MGI:3648511] |
| 5 | protein_coding | 1.13E+08 | 1.13E+08 | ENSMUSG00000029348 | MGI:1920148 | Asphd2 | aspartate beta-hydroxylase domain containing 2 [Source:MGI Symbol;Acc:MGI:1920148] |
| 5 | protein_coding | 1.13E+08 | 1.13E+08 | ENSMUSG00000058153 | MGI:1935121 | Sez6l | seizure related 6 homolog like [Source:MGI Symbol;Acc:MGI:1935121] |
| 5 | protein_coding | 1.13E+08 | 1.13E+08 | ENSMUSG00000072722 | MGI:3648937 | Ccdc121rt2 | coiled-coil domain containing 121, retrogene 2 [Source:MGI Symbol;Acc:MGI:3648937] |
| 5 | protein_coding | 1.13E+08 | 1.13E+08 | ENSMUSG00000072720 | MGI:1921626 | Myo18b | myosin XVIIIb [Source:MGI Symbol;Acc:MGI:1921626] |
| 5 | protein_coding | 1.13E+08 | 1.13E+08 | ENSMUSG00000042249 | MGI:87941 | Grk3 | G protein-coupled receptor kinase 3 [Source:MGI Symbol;Acc:MGI:87941] |
| 5 | protein_coding | 1.13E+08 | 1.13E+08 | ENSMUSG00000042240 | MGI:88519 | Crybb2 | crystallin, beta B2 [Source:MGI Symbol;Acc:MGI:88519] |
| 5 | protein_coding | 1.13E+08 | 1.13E+08 | ENSMUSG00000029352 | MGI:102717 | Crybb3 | crystallin, beta B3 [Source:MGI Symbol;Acc:MGI:102717] |

| | | | | | | | |
|---|---|---|---|---|---|---|---|
| 5 | protein_coding | 1.13E+08 | 1.13E+08 | ENSMUSG00000051339 | MGI:1920194 | 2900026A02Rik | RIKEN cDNA 2900026A02 gene [Source:MGI Symbol;Acc:MGI:1920194] |
| 5 | protein_coding | 1.13E+08 | 1.13E+08 | ENSMUSG00000066964 | MGI:2685700 | Lhfpl7 | LHFPL tetraspan subfamily member 7 [Source:MGI Symbol;Acc:MGI:2685700] |
| 5 | protein_coding | 1.13E+08 | 1.13E+08 | ENSMUSG00000042216 | MGI:107320 | Sgsm1 | small G protein signaling modulator 1 [Source:MGI Symbol;Acc:MGI:107320] |
| 5 | protein_coding | 1.14E+08 | 1.14E+08 | ENSMUSG00000063430 | MGI:2445030 | Wscd2 | WSC domain containing 2 [Source:MGI Symbol;Acc:MGI:2445030] |
| 5 | protein_coding | 1.14E+08 | 1.14E+08 | ENSMUSG00000042190 | MGI:109603 | Cmklr1 | chemerin chemokine-like receptor 1 [Source:MGI Symbol;Acc:MGI:109603] |
| 5 | protein_coding | 1.14E+08 | 1.14E+08 | ENSMUSG00000053334 | MGI:1098550 | Ficd | FIC domain containing [Source:MGI Symbol;Acc:MGI:1098550] |
| 5 | protein_coding | 1.14E+08 | 1.14E+08 | ENSMUSG00000018974 | MGI:1858230 | Sart3 | [illegible] by T cells 3 [Source:MGI Symbol;Acc:MGI:1858230] |
| 5 | protein_coding | 1.14E+08 | 1.14E+08 | ENSMUSG00000025825 | MGI:1913633 | Iscu | iron-sulfur cluster assembly enzyme [Source:MGI Symbol;Acc:MGI:1913633] |
| 5 | protein_coding | 1.14E+08 | 1.14E+08 | ENSMUSG00000054675 | MGI:2385228 | Tmem119 | transmembrane protein 119 [Source:MGI Symbol;Acc:MGI:2385228] |
| 5 | protein_coding | 1.14E+08 | 1.14E+08 | ENSMUSG00000048163 | MGI:106689 | Selplg | selectin, platelet (p-selectin) ligand [Source:MGI Symbol;Acc:MGI:106689] |
| 5 | protein_coding | 1.14E+08 | 1.14E+08 | ENSMUSG00000004530 | MGI:1345964 | Coro1c | coronin, actin binding protein 1C [Source:MGI Symbol;Acc:MGI:1345964] |
| 5 | protein_coding | 1.14E+08 | 1.14E+08 | ENSMUSG00000042121 | MGI:2686240 | Ssh1 | slingshot protein phosphatase 1 [Source:MGI Symbol;Acc:MGI:2686240] |
| 5 | protein_coding | 1.14E+08 | 1.14E+08 | ENSMUSG00000042096 | MGI:94859 | Dao | D-amino acid oxidase [Source:MGI Symbol;Acc:MGI:94859] |
| 5 | protein_coding | 1.14E+08 | 1.14E+08 | ENSMUSG00000042078 | MGI:1915916 | Svop | SV2 related protein [Source:MGI Symbol;Acc:MGI:1915916] |
| 5 | protein_coding | 1.14E+08 | 1.14E+08 | ENSMUSG00000029592 | MGI:2140991 | Usp30 | ubiquitin specific peptidase 30 [Source:MGI Symbol;Acc:MGI:2140991] |
| 5 | protein_coding | 1.14E+08 | 1.14E+08 | ENSMUSG00000044339 | MGI:2141032 | Alkbh2 | alkB homolog 2, alpha-ketoglutarate-dependent dioxygenase [Source:MGI Symbol;Acc:MGI:2141032] |
| 5 | protein_coding | 1.14E+08 | 1.14E+08 | ENSMUSG00000029591 | MGI:109352 | Ung | uracil DNA glycosylase [Source:MGI Symbol;Acc:MGI:109352] |

| | | | | | | | |
|---|---|---|---|---|---|---|---|
| 5 | protein_coding | 1.14E+08 | 1.14E+08 | ENSMUSG00000042010 | MGI:2140940 | Acacb | acetyl-Coenzyme A carboxylase beta [Source:MGI Symbol;Acc:MGI:2140940] |
| 5 | protein_coding | 1.14E+08 | 1.14E+08 | ENSMUSG00000042002 | MGI:2151057 | Foxn4 | forkhead box N4 [Source:MGI Symbol;Acc:MGI:2151057] |
| 5 | protein_coding | 1.14E+08 | 1.15E+08 | ENSMUSG00000066952 | MGI:1914674 | Myo1h | myosin 1H [Source:MGI Symbol;Acc:MGI:1914674] |
| 5 | protein_coding | 1.15E+08 | 1.15E+08 | ENSMUSG00000001098 | MGI:2141207 | Kctd10 | potassium channel tetramerisation domain containing 10 [Source:MGI Symbol;Acc:MGI:2141207] |
| 5 | protein_coding | 1.15E+08 | 1.15E+08 | ENSMUSG00000029577 | MGI:1891295 | Ube3b | ubiquitin protein ligase E3B [Source:MGI Symbol;Acc:MGI:1891295] |
| 5 | protein_coding | 1.15E+08 | 1.15E+08 | ENSMUSG00000029575 | MGI:1924947 | Mmab | methylmalonic aciduria (cobalamin deficiency) cblB type homolog (human) [Source:MGI Symbol;Acc:MGI:1924947] |
| 5 | protein_coding | 1.15E+08 | 1.15E+08 | ENSMUSG00000041939 | MGI:107624 | Mvk | mevalonate kinase [Source:MGI Symbol;Acc:MGI:107624] |
| 5 | protein_coding | 1.15E+08 | 1.15E+08 | ENSMUSG00000041930 | MGI:3605543 | Fam222a | family with sequence similarity 222, member A [Source:MGI Symbol;Acc:MGI:3605543] |
| 5 | protein_coding | 1.15E+08 | 1.15E+08 | ENSMUSG00000014158 | MGI:1926945 | Trpv4 | transient receptor potential cation channel, subfamily V, member 4 [Source:MGI Symbol;Acc:MGI:1926945] |
| 5 | protein_coding | 1.15E+08 | 1.15E+08 | ENSMUSG00000011884 | MGI:1929253 | Gltp | glycolipid transfer protein [Source:MGI Symbol;Acc:MGI:1929253] |
| 5 | protein_coding | 1.15E+08 | 1.15E+08 | ENSMUSG00000002486 | MGI:1925082 | Tchp | trichoplein, keratin filament binding [Source:MGI Symbol;Acc:MGI:1925082] |
| 5 | protein_coding | 1.15E+08 | 1.15E+08 | ENSMUSG00000041890 | MGI:1347053 | Git2 | GIT ArfGAP 2 [Source:MGI Symbol;Acc:MGI:1347053] |
| 5 | protein_coding | 1.15E+08 | 1.15E+08 | ENSMUSG00000041870 | MGI:1915670 | Ankrd13a | ankyrin repeat domain 13a [Source:MGI Symbol;Acc:MGI:1915670] |
| 5 | protein_coding | 1.15E+08 | 1.15E+08 | ENSMUSG00000072694 | MGI:7529997 | Gm57857 | predicted gene, 57857 [Source:MGI Symbol;Acc:MGI:7529997] |
| 5 | protein_coding | 1.15E+08 | 1.15E+08 | ENSMUSG00000029564 | MGI:1914843 | 4930519G04Rik | RIKEN cDNA 4930519G04 gene [Source:MGI Symbol;Acc:MGI:1914843] |
| 5 | protein_coding | 1.15E+08 | 1.15E+08 | ENSMUSG00000029561 | MGI:1344390 | Oasl2 | 2'-5' oligoadenylate synthetase-like 2 [Source:MGI Symbol;Acc:MGI:1344390] |

| | | | | | | | |
|---|---|---|---|---|---|---|---|
| 5 | protein_coding | 1.15E+08 | 1.15E+08 | ENSMUSG00000041827 | MGI:2180849 | Oasl1 | 2'-5' oligoadenylate synthetase-like 1 [Source:MGI Symbol;Acc:MGI:2180849] |
| 5 | protein_coding | 1.15E+08 | 1.15E+08 | ENSMUSG00000029559 | MGI:1919607 | 2210016L21Rik | RIKEN cDNA 2210016L21 gene [Source:MGI Symbol;Acc:MGI:1919607] |
| 5 | protein_coding | 1.15E+08 | 1.15E+08 | ENSMUSG00000029556 | MGI:98504 | Hnf1a | HNF1 homeobox A [Source:MGI Symbol;Acc:MGI:98504] |
| 5 | protein_coding | 1.15E+08 | 1.15E+08 | ENSMUSG00000029550 | MGI:1891433 | Sppl3 | signal peptide peptidase 3 [Source:MGI Symbol;Acc:MGI:1891433] |
| 5 | protein_coding | 1.15E+08 | 1.15E+08 | ENSMUSG00000029545 | MGI:87868 | Acads | acyl-Coenzyme A dehydrogenase, short chain [Source:MGI Symbol;Acc:MGI:87868] |
| 5 | protein_coding | 1.15E+08 | 1.15E+08 | ENSMUSG00000046562 | MGI:2147162 | Unc119b | unc-119 lipid binding chaperone B [Source:MGI Symbol;Acc:MGI:2147162] |
| 5 | protein_coding | 1.15E+08 | 1.15E+08 | ENSMUSG00000048578 | MGI:1924015 | Mlec | malectin [Source:MGI Symbol;Acc:MGI:1924015] |
| 5 | protein_coding | 1.15E+08 | 1.15E+08 | ENSMUSG00000029544 | MGI:1352750 | Cabp1 | calcium binding protein 1 [Source:MGI Symbol;Acc:MGI:1352750] |
| 5 | protein_coding | 1.15E+08 | 1.15E+08 | ENSMUSG00000060152 | MGI:2151221 | Pop5 | processing of precursor 5, ribonuclease P/MRP family (S. cerevisiae) [Source:MGI Symbol;Acc:MGI:2151221] |
| 5 | protein_coding | 1.15E+08 | 1.15E+08 | ENSMUSG00000041740 | MGI:1859162 | Rnf10 | ring finger protein 10 [Source:MGI Symbol;Acc:MGI:1859162] |
| 5 | protein_coding | 1.15E+08 | 1.15E+08 | ENSMUSG00000144239 | | | novel protein |
| 5 | protein_coding | 1.15E+08 | 1.15E+08 | ENSMUSG00000041733 | MGI:1098643 | Coq5 | coenzyme Q5 methyltransferase [Source:MGI Symbol;Acc:MGI:1098643] |
| 5 | protein_coding | 1.15E+08 | 1.15E+08 | ENSMUSG00000009013 | MGI:1861457 | Dynll1 | dynein light chain LC8-type 1 [Source:MGI Symbol;Acc:MGI:1861457] |
| 5 | protein_coding | 1.15E+08 | 1.15E+08 | ENSMUSG00000029538 | MGI:104896 | Srsf9 | serine and arginine-rich splicing factor 9 [Source:MGI Symbol;Acc:MGI:104896] |
| 5 | protein_coding | 1.15E+08 | 1.15E+08 | ENSMUSG00000029536 | MGI:1923776 | Gatc | glutamyl-tRNA amidotransferase subunit C [Source:MGI Symbol;Acc:MGI:1923776] |
| 5 | protein_coding | 1.15E+08 | 1.15E+08 | ENSMUSG00000029535 | MGI:1916326 | Triap1 | TP53 regulated inhibitor of apoptosis 1 [Source:MGI Symbol;Acc:MGI:1916326] |
| 5 | protein_coding | 1.15E+08 | 1.15E+08 | ENSMUSG00000041697 | MGI:103099 | Cox6a1 | cytochrome c oxidase subunit 6A1 [Source:MGI Symbol;Acc:MGI:103099] |

| | | | | | | | |
|---|---|---|---|---|---|---|---|
| 5 | protein_coding | 1.16E+08 | 1.16E+08 | ENSMUSG00000054256 | MGI:107376 | Msi1 | musashi RNA-binding protein 1 [Source:MGI Symbol;Acc:MGI:107376] |
| 5 | protein_coding | 1.16E+08 | 1.16E+08 | ENSMUSG00000029522 | MGI:101842 | Pla2g1b | phospholipase A2, group IB, pancreas [Source:MGI Symbol;Acc:MGI:101842] |
| 5 | protein_coding | 1.16E+08 | 1.16E+08 | ENSMUSG00000029524 | MGI:1922637 | Sirt4 | sirtuin 4 [Source:MGI Symbol;Acc:MGI:1922637] |
| 5 | protein_coding | 1.16E+08 | 1.16E+08 | ENSMUSG00000029528 | MGI:108295 | Pxn | paxillin [Source:MGI Symbol;Acc:MGI:108295] |
| 5 | protein_coding | 1.16E+08 | 1.16E+08 | ENSMUSG00000067274 | MGI:1927636 | Rplp0 | ribosomal protein lateral stalk subunit P0 [Source:MGI Symbol;Acc:MGI:1927636] |
| 5 | protein_coding | 1.16E+08 | 1.16E+08 | ENSMUSG00000041638 | MGI:2444248 | Gcn1 | GCN1 activator of EIF2AK4 [Source:MGI Symbol;Acc:MGI:2444248] |
| 5 | protein_coding | 1.16E+08 | 1.16E+08 | ENSMUSG00000029518 | MGI:1924657 | Rab35 | RAB35, member RAS oncogene family [Source:MGI Symbol;Acc:MGI:1924657] |
| 5 | protein_coding | 1.16E+08 | 1.16E+08 | ENSMUSG00000041609 | MGI:1922915 | Bicdl1 | BICD family like cargo adaptor 1 [Source:MGI Symbol;Acc:MGI:1922915] |
| 5 | protein_coding | 1.16E+08 | 1.16E+08 | ENSMUSG00000029516 | MGI:105313 | Cit | citron [Source:MGI Symbol;Acc:MGI:105313] |
| 5 | protein_coding | 1.16E+08 | 1.16E+08 | ENSMUSG00000029513 | MGI:1336167 | Prkab1 | catalytic subunit [Source:MGI Symbol;Acc:MGI:1336167] |
| 5 | protein_coding | 1.16E+08 | 1.16E+08 | ENSMUSG00000079278 | MGI:3651514 | Tmem233 | transmembrane protein 233 [Source:MGI Symbol;Acc:MGI:3651514] |
| 5 | protein_coding | 1.16E+08 | 1.16E+08 | ENSMUSG00000043913 | MGI:2141043 | Ccdc60 | coiled-coil domain containing 60 [Source:MGI Symbol;Acc:MGI:2141043] |
| 5 | protein_coding | 1.17E+08 | 1.17E+08 | ENSMUSG00000041548 | MGI:2135756 | Hspb8 | heat shock protein 8 [Source:MGI Symbol;Acc:MGI:2135756] |
| 5 | protein_coding | 1.17E+08 | 1.17E+08 | ENSMUSG00000063919 | MGI:1916205 | Srrm4 | serine/arginine repetitive matrix 4 [Source:MGI Symbol;Acc:MGI:1916205] |
| 5 | protein_coding | 1.17E+08 | 1.17E+08 | ENSMUSG00000066900 | MGI:1919204 | Suds3 | suppressor of defective silencing 3 homolog (S. cerevisiae) [Source:MGI Symbol;Acc:MGI:1919204] |
| 5 | protein_coding | 1.17E+08 | 1.17E+08 | ENSMUSG00000061288 | MGI:3041177 | Taok3 | TAO kinase 3 [Source:MGI Symbol;Acc:MGI:3041177] |
| 5 | protein_coding | 1.17E+08 | 1.17E+08 | ENSMUSG00000032959 | MGI:1344408 | Pebp1 | phosphatidylethanolamine binding protein 1 [Source:MGI Symbol;Acc:MGI:1344408] |

| | | | | | | | |
|---|---|---|---|---|---|---|---|
| 5 | protein_coding | 1.17E+08 | 1.17E+08 | ENSMUSG00000066894 | MGI:2448533 | Vsig10 | V-set and immunoglobulin domain containing 10 [Source:MGI Symbol;Acc:MGI:2448533] |
| 5 | protein_coding | 1.17E+08 | 1.18E+08 | ENSMUSG00000029364 | MGI:2144041 | Wsb2 | WD repeat and SOCS box-containing 2 [Source:MGI Symbol;Acc:MGI:2144041] |
| 5 | protein_coding | 1.18E+08 | 1.18E+08 | ENSMUSG00000029363 | MGI:1919401 | Rfc5 | replication factor C (activator 1) 5 [Source:MGI Symbol;Acc:MGI:1919401] |
| 5 | protein_coding | 1.18E+08 | 1.18E+08 | ENSMUSG00000061578 | MGI:3610315 | Ksr2 | kinase suppressor of ras 2 [Source:MGI Symbol;Acc:MGI:3610315] |
| 5 | protein_coding | 1.18E+08 | 1.18E+08 | ENSMUSG00000029361 | MGI:97360 | Nos1 | nitric oxide synthase 1, neuronal [Source:MGI Symbol;Acc:MGI:97360] |
| 5 | protein_coding | 1.18E+08 | 1.18E+08 | ENSMUSG00000032898 | MGI:1924223 | Fbxo21 | F-box protein 21 [Source:MGI Symbol;Acc:MGI:1924223] |
| 5 | protein_coding | 1.18E+08 | 1.18E+08 | ENSMUSG00000029359 | MGI:1930803 | Tesc | tescalcin [Source:MGI Symbol;Acc:MGI:1930803] |
| 5 | protein_coding | 1.18E+08 | 1.18E+08 | ENSMUSG00000032867 | MGI:1923041 | Fbxw8 | F-box and WD-40 domain protein 8 [Source:MGI Symbol;Acc:MGI:1923041] |
| 5 | protein_coding | 1.18E+08 | 1.18E+08 | ENSMUSG00000046607 | MGI:1201608 | Hrk | harakiri, BCL2 interacting protein (contains only BH3 domain) [Source:MGI Symbol;Acc:MGI:1201608] |
| 5 | protein_coding | 1.18E+08 | 1.18E+08 | ENSMUSG00000032850 | MGI:2442859 | Rnft2 | ring finger protein, transmembrane 2 [Source:MGI Symbol;Acc:MGI:2442859] |
| 5 | protein_coding | 1.18E+08 | 1.18E+08 | ENSMUSG00000032840 | MGI:1924042 | Spring1 | SREBF pathway regulator in golgi 1 [Source:MGI Symbol;Acc:MGI:1924042] |
| 5 | protein_coding | 1.19E+08 | 1.19E+08 | ENSMUSG00000018076 | MGI:2670178 | Med13l | mediator complex subunit 13-like [Source:MGI Symbol;Acc:MGI:2670178] |
| 5 | protein_coding | 1.20E+08 | 1.20E+08 | ENSMUSG00000018604 | MGI:98495 | Tbx3 | T-box 3 [Source:MGI Symbol;Acc:MGI:98495] |
| 5 | protein_coding | 1.20E+08 | 1.20E+08 | ENSMUSG00000018263 | MGI:102541 | Tbx5 | T-box 5 [Source:MGI Symbol;Acc:MGI:102541] |
| 5 | protein_coding | 1.20E+08 | 1.20E+08 | ENSMUSG00000029594 | MGI:1921361 | Rbm19 | RNA binding motif protein 19 [Source:MGI Symbol;Acc:MGI:1921361] |
| 5 | protein_coding | 1.21E+08 | 1.21E+08 | ENSMUSG00000029595 | MGI:107792 | Lhx5 | LIM homeobox protein 5 [Source:MGI Symbol;Acc:MGI:107792] |
| 5 | protein_coding | 1.21E+08 | 1.21E+08 | ENSMUSG00000029596 | MGI:2182607 | Sdsl | serine dehydratase-like [Source:MGI Symbol;Acc:MGI:2182607] |

| | | | | | | | |
|---|---|---|---|---|---|---|---|
| 5 | protein_coding | 1.21E+08 | 1.21E+08 | ENSMUSG00000029597 | MGI:98270 | Sds | serine dehydratase [Source:MGI Symbol;Acc:MGI:98270] |
| 5 | protein_coding | 1.21E+08 | 1.21E+08 | ENSMUSG00000029598 | MGI:1919022 | Plbd2 | phospholipase B domain containing 2 [Source:MGI Symbol;Acc:MGI:1919022] |
| 5 | protein_coding | 1.21E+08 | 1.21E+08 | ENSMUSG00000032754 | MGI:2180781 | Slc8b1 | solute carrier family 8 (sodium/lithium/calcium exchanger), member B1 [Source:MGI Symbol;Acc:MGI:2180781] |
| 5 | protein_coding | 1.21E+08 | 1.21E+08 | ENSMUSG00000032741 | MGI:2182472 | Tpcn1 | two pore channel 1 [Source:MGI Symbol;Acc:MGI:2182472] |
| 5 | protein_coding | 1.21E+08 | 1.21E+08 | ENSMUSG00000029601 | MGI:1922982 | Iqcd | IQ motif containing D [Source:MGI Symbol;Acc:MGI:1922982] |
| 5 | protein_coding | 1.21E+08 | 1.21E+08 | ENSMUSG00000029600 | MGI:1922021 | Rita1 | RBPJ interacting and tubulin associated 1 [Source:MGI Symbol;Acc:MGI:1922021] |
| 5 | protein_coding | 1.21E+08 | 1.21E+08 | ENSMUSG00000029599 | MGI:1919240 | Ddx54 | DEAD box helicase 54 [Source:MGI Symbol;Acc:MGI:1919240] |
| 5 | protein_coding | 1.21E+08 | 1.21E+08 | ENSMUSG00000094282 | MGI:3779542 | Cfap73 | cilia and flagella associated protein 73 [Source:MGI Symbol;Acc:MGI:3779542] |
| 5 | protein_coding | 1.21E+08 | 1.21E+08 | ENSMUSG00000029602 | MGI:1330842 | Rasal1 | RAS protein activator like 1 (GAP1 like) [Source:MGI Symbol;Acc:MGI:1330842] |
| 5 | protein_coding | 1.21E+08 | 1.21E+08 | ENSMUSG00000029603 | MGI:1352744 | Dtx1 | deltex 1, E3 ubiquitin ligase [Source:MGI Symbol;Acc:MGI:1352744] |
| 5 | protein_coding | 1.21E+08 | 1.21E+08 | ENSMUSG00000032690 | MGI:2180852 | Oas2 | 2'-5' oligoadenylate synthetase 2 [Source:MGI Symbol;Acc:MGI:2180852] |
| 5 | protein_coding | 1.21E+08 | 1.21E+08 | ENSMUSG00000032661 | MGI:2180850 | Oas3 | 2'-5' oligoadenylate synthetase 3 [Source:MGI Symbol;Acc:MGI:2180850] |
| 5 | protein_coding | 1.21E+08 | 1.21E+08 | ENSMUSG00000066867 | MGI:2180856 | Oas1e | 2'-5' oligoadenylate synthetase 1E [Source:MGI Symbol;Acc:MGI:2180856] |
| 5 | protein_coding | 1.21E+08 | 1.21E+08 | ENSMUSG00000001166 | MGI:2149633 | Oas1c | 2'-5' oligoadenylate synthetase 1C [Source:MGI Symbol;Acc:MGI:2149633] |
| 5 | protein_coding | 1.21E+08 | 1.21E+08 | ENSMUSG00000029605 | MGI:97430 | Oas1b | 2'-5' oligoadenylate synthetase 1B [Source:MGI Symbol;Acc:MGI:97430] |
| 5 | protein_coding | 1.21E+08 | 1.21E+08 | ENSMUSG00000053765 | MGI:2180855 | Oas1f | 2'-5' oligoadenylate synthetase 1F [Source:MGI Symbol;Acc:MGI:2180855] |
| 5 | protein_coding | 1.21E+08 | 1.21E+08 | ENSMUSG00000001168 | MGI:2180853 | Oas1h | 2'-5' oligoadenylate synthetase 1H [Source:MGI Symbol;Acc:MGI:2180853] |

| | | | | | | | |
|---|---|---|---|---|---|---|---|
| 5 | protein_coding | 1.21E+08 | 1.21E+08 | ENSMUSG00000066861 | MGI:97429 | Oas1g | 2'-5' oligoadenylate synthetase 1G [Source:MGI Symbol;Acc:MGI:97429] |
| 5 | protein_coding | 1.21E+08 | 1.21E+08 | ENSMUSG00000052776 | MGI:2180860 | Oas1a | 2'-5' oligoadenylate synthetase 1A [Source:MGI Symbol;Acc:MGI:2180860] |
| 5 | protein_coding | 1.21E+08 | 1.21E+08 | ENSMUSG00000032623 | MGI:2140770 | Oas1d | 2'-5' oligoadenylate synthetase 1D [Source:MGI Symbol;Acc:MGI:2140770] |
| 5 | protein_coding | 1.21E+08 | 1.21E+08 | ENSMUSG00000029608 | MGI:102788 | Rph3a | rabphilin 3A [Source:MGI Symbol;Acc:MGI:102788] |
| 5 | protein_coding | 1.21E+08 | 1.21E+08 | ENSMUSG00000043733 | MGI:99511 | Ptpn11 | protein tyrosine phosphatase, non-receptor type 11 [Source:MGI Symbol;Acc:MGI:99511] |
| 5 | protein_coding | 1.21E+08 | 1.21E+08 | ENSMUSG00000029614 | MGI:108057 | Rpl6 | ribosomal protein L6 [Source:MGI Symbol;Acc:MGI:108057] |
| 5 | protein_coding | 1.21E+08 | 1.22E+08 | ENSMUSG00000042744 | MGI:3647820 | Hectd4 | HECT domain E3 ubiquitin protein ligase 4 [Source:MGI Symbol;Acc:MGI:3647820] |
| 5 | protein_coding | 1.22E+08 | 1.22E+08 | ENSMUSG00000042726 | MGI:1923551 | Trafd1 | TRAF type zinc finger domain containing 1 [Source:MGI Symbol;Acc:MGI:1923551] |
| 5 | protein_coding | 1.22E+08 | 1.22E+08 | ENSMUSG00000042719 | MGI:2442563 | Naa25 | [illegible] subunit [Source:MGI Symbol;Acc:MGI:2442563] |
| 5 | protein_coding | 1.22E+08 | 1.22E+08 | ENSMUSG00000029616 | MGI:1914647 | Erp29 | endoplasmic reticulum protein 29 [Source:MGI Symbol;Acc:MGI:1914647] |
| 5 | protein_coding | 1.22E+08 | 1.22E+08 | ENSMUSG00000029452 | MGI:1924712 | Tmem116 | transmembrane protein 116 [Source:MGI Symbol;Acc:MGI:1924712] |
| 5 | protein_coding | 1.22E+08 | 1.22E+08 | ENSMUSG00000062438 | MGI:2429506 | Adam1b | a disintegrin and metallopeptidase domain 1b [Source:MGI Symbol;Acc:MGI:2429506] |
| 5 | protein_coding | 1.22E+08 | 1.22E+08 | ENSMUSG00000072647 | MGI:2429504 | Adam1a | a disintegrin and metallopeptidase domain 1a [Source:MGI Symbol;Acc:MGI:2429504] |
| 5 | protein_coding | 1.22E+08 | 1.22E+08 | ENSMUSG00000105340 | MGI:5663015 | Gm42878 | predicted gene 42878 [Source:MGI Symbol;Acc:MGI:5663015] |
| 5 | protein_coding | 1.22E+08 | 1.22E+08 | ENSMUSG00000029454 | MGI:1333110 | Mapkapk5 | MAP kinase-activated protein kinase 5 [Source:MGI Symbol;Acc:MGI:1333110] |
| 5 | protein_coding | 1.22E+08 | 1.22E+08 | ENSMUSG00000121598 | MGI:7822511 | Gm69851 | predicted gene, 69851 [Source:MGI Symbol;Acc:MGI:7822511] |
| 5 | protein_coding | 1.22E+08 | 1.22E+08 | ENSMUSG00000029455 | MGI:99600 | Aldh2 | aldehyde dehydrogenase 2, mitochondrial [Source:MGI Symbol;Acc:MGI:99600] |
| 5 | protein_coding | 1.22E+08 | 1.22E+08 | ENSMUSG00000042647 | MGI:2443320 | Acad12 | [illegible] member 12 [Source:MGI Symbol;Acc:MGI:2443320] |

| | | | | | | | |
|---|---|---|---|---|---|---|---|
| 5 | protein_coding | 1.22E+08 | 1.22E+08 | ENSMUSG00000029456 | MGI:1919235 | Acad10 | [illegible] member 10 [Source:MGI Symbol;Acc:MGI:1919235] |
| 5 | protein_coding | 1.22E+08 | 1.22E+08 | ENSMUSG00000029458 | MGI:1919649 | Brap | BRCA1 associated protein [Source:MGI Symbol;Acc:MGI:1919649] |
| 5 | protein_coding | 1.22E+08 | 1.22E+08 | ENSMUSG00000042605 | MGI:1277223 | Atxn2 | ataxin 2 [Source:MGI Symbol;Acc:MGI:1277223] |
| 5 | protein_coding | 1.22E+08 | 1.22E+08 | ENSMUSG00000042594 | MGI:893598 | Sh2b3 | SH2B adaptor protein 3 [Source:MGI Symbol;Acc:MGI:893598] |
| 5 | protein_coding | 1.22E+08 | 1.22E+08 | ENSMUSG00000044134 | MGI:2442708 | Pheta1 | [illegible] adaptor 1 [Source:MGI Symbol;Acc:MGI:2442708] |
| 5 | protein_coding | 1.22E+08 | 1.22E+08 | ENSMUSG00000042589 | MGI:107321 | Cux2 | cut-like homeobox 2 [Source:MGI Symbol;Acc:MGI:107321] |
| 5 | protein_coding | 1.22E+08 | 1.22E+08 | ENSMUSG00000013936 | MGI:97272 | Myl2 | myosin, light polypeptide 2, regulatory, cardiac, slow [Source:MGI Symbol;Acc:MGI:97272] |
| 5 | protein_coding | 1.22E+08 | 1.22E+08 | ENSMUSG00000043036 | MGI:3607777 | Ccdc63 | coiled-coil domain containing 63 [Source:MGI Symbol;Acc:MGI:3607777] |
| 5 | protein_coding | 1.22E+08 | 1.22E+08 | ENSMUSG00000004455 | MGI:104872 | Ppp1cc | protein phosphatase 1 catalytic subunit gamma [Source:MGI Symbol;Acc:MGI:104872] |
| 5 | protein_coding | 1.22E+08 | 1.22E+08 | ENSMUSG00000064267 | MGI:1921346 | Hvcn1 | hydrogen voltage-gated channel 1 [Source:MGI Symbol;Acc:MGI:1921346] |
| 5 | protein_coding | 1.22E+08 | 1.22E+08 | ENSMUSG00000038593 | MGI:3603820 | Tctn1 | tectonic family member 1 [Source:MGI Symbol;Acc:MGI:3603820] |
| 5 | protein_coding | 1.22E+08 | 1.22E+08 | ENSMUSG00000038582 | MGI:2444593 | Pptc7 | PTC7 protein phosphatase homolog [Source:MGI Symbol;Acc:MGI:2444593] |
| 5 | protein_coding | 1.22E+08 | 1.22E+08 | ENSMUSG00000038569 | MGI:2385231 | Rad9b | RAD9 checkpoint clamp component B [Source:MGI Symbol;Acc:MGI:2385231] |
| 5 | protein_coding | 1.22E+08 | 1.23E+08 | ENSMUSG00000029462 | MGI:1928344 | Vps29 | VPS29 retromer complex component [Source:MGI Symbol;Acc:MGI:1928344] |
| 5 | protein_coding | 1.23E+08 | 1.23E+08 | ENSMUSG00000029463 | MGI:1916198 | Fam216a | family with sequence similarity 216, member A [Source:MGI Symbol;Acc:MGI:1916198] |
| 5 | protein_coding | 1.23E+08 | 1.23E+08 | ENSMUSG00000029464 | MGI:1289326 | Gpn3 | GPN-loop GTPase 3 [Source:MGI Symbol;Acc:MGI:1289326] |
| 5 | protein_coding | 1.23E+08 | 1.23E+08 | ENSMUSG00000029465 | MGI:1928375 | Arpc3 | actin related protein 2/3 complex, subunit 3 [Source:MGI Symbol;Acc:MGI:1928375] |
| 5 | protein_coding | 1.23E+08 | 1.23E+08 | ENSMUSG00000029466 | MGI:1929711 | Anapc7 | anaphase promoting complex subunit 7 [Source:MGI Symbol;Acc:MGI:1929711] |

| | | | | | | | |
|---|---|---|---|---|---|---|---|
| 5 | protein_coding | 1.23E+08 | 1.23E+08 | ENSMUSG00000029467 | MGI:88110 | Atp2a2 | slow twitch 2 [Source:MGI Symbol;Acc:MGI:88110] |
| 5 | protein_coding | 1.23E+08 | 1.23E+08 | ENSMUSG00000029469 | MGI:1098597 | Ift81 | intraflagellar transport 81 [Source:MGI Symbol;Acc:MGI:1098597] |
| 5 | protein_coding | 1.23E+08 | 1.23E+08 | ENSMUSG00000029468 | MGI:1339957 | P2rx7 | channel, 7 [Source:MGI Symbol;Acc:MGI:1339957] |
| 5 | protein_coding | 1.23E+08 | 1.23E+08 | ENSMUSG00000029470 | MGI:1338859 | P2rx4 | channel 4 [Source:MGI Symbol;Acc:MGI:1338859] |
| 5 | protein_coding | 1.23E+08 | 1.23E+08 | ENSMUSG00000029471 | MGI:2444812 | Camkk2 | calcium/calmodulin-dependent protein kinase kinase 2, beta [Source:MGI Symbol;Acc:MGI:2444812] |
| 5 | protein_coding | 1.23E+08 | 1.23E+08 | ENSMUSG00000029472 | MGI:1929722 | Anapc5 | anaphase-promoting complex subunit 5 [Source:MGI Symbol;Acc:MGI:1929722] |
| 5 | protein_coding | 1.23E+08 | 1.23E+08 | ENSMUSG00000029474 | MGI:2153340 | Rnf34 | ring finger protein 34 [Source:MGI Symbol;Acc:MGI:2153340] |
| 5 | protein_coding | 1.23E+08 | 1.23E+08 | ENSMUSG00000029475 | MGI:1354737 | Kdm2b | lysine (K)-specific demethylase 2B [Source:MGI Symbol;Acc:MGI:1354737] |
| 5 | protein_coding | 1.23E+08 | 1.23E+08 | ENSMUSG00000049686 | MGI:1925542 | Orai1 | modulator 1 [Source:MGI Symbol;Acc:MGI:1925542] |
| 5 | protein_coding | 1.23E+08 | 1.23E+08 | ENSMUSG00000029477 | MGI:1922140 | Morn3 | MORN repeat containing 3 [Source:MGI Symbol;Acc:MGI:1922140] |
| 5 | protein_coding | 1.23E+08 | 1.23E+08 | ENSMUSG00000054434 | MGI:3603158 | Tmem120b | transmembrane protein 120B [Source:MGI Symbol;Acc:MGI:3603158] |
| 5 | protein_coding | 1.23E+08 | 1.23E+08 | ENSMUSG00000029449 | MGI:1345629 | Rhof | ras homolog family member F (in filopodia) [Source:MGI Symbol;Acc:MGI:1345629] |
| 5 | protein_coding | 1.23E+08 | 1.23E+08 | ENSMUSG00000038384 | MGI:2652820 | Setd1b | SET domain containing 1B [Source:MGI Symbol;Acc:MGI:2652820] |
| 5 | protein_coding | 1.23E+08 | 1.23E+08 | ENSMUSG00000029440 | MGI:1914401 | Psmd9 | proteasome (prosome, macropain) 26S subunit, non-ATPase, 9 [Source:MGI Symbol;Acc:MGI:1914401] |
| 5 | protein_coding | 1.23E+08 | 1.23E+08 | ENSMUSG00000029445 | MGI:96213 | Hpd | 4-hydroxyphenylpyruvic acid dioxygenase [Source:MGI Symbol;Acc:MGI:96213] |
| 5 | protein_coding | 1.23E+08 | 1.23E+08 | ENSMUSG00000029442 | MGI:1918495 | Cfap251 | cilia and flagella associated protein 251 [Source:MGI Symbol;Acc:MGI:1918495] |
| 5 | protein_coding | 1.23E+08 | 1.24E+08 | ENSMUSG00000029438 | MGI:1924295 | Bcl7a | B cell CLL/lymphoma 7A [Source:MGI Symbol;Acc:MGI:1924295] |

| | | | | | | | |
|---|---|---|---|---|---|---|---|
| 5 | protein_coding | 1.24E+08 | 1.24E+08 | ENSMUSG00000038342 | MGI:2141183 | Mlxip | MLX interacting protein [Source:MGI Symbol;Acc:MGI:2141183] |
| 5 | protein_coding | 1.24E+08 | 1.24E+08 | ENSMUSG00000029437 | MGI:1923649 | Il31 | interleukin 31 [Source:MGI Symbol;Acc:MGI:1923649] |
| 5 | protein_coding | 1.24E+08 | 1.24E+08 | ENSMUSG00000063409 | MGI:2685907 | Lrrc43 | leucine rich repeat containing 43 [Source:MGI Symbol;Acc:MGI:2685907] |
| 5 | protein_coding | 1.24E+08 | 1.24E+08 | ENSMUSG00000029433 | MGI:1913843 | Diablo | diablo, IAP-binding mitochondrial protein [Source:MGI Symbol;Acc:MGI:1913843] |
| 5 | protein_coding | 1.24E+08 | 1.24E+08 | ENSMUSG00000029431 | MGI:2680208 | B3gnt4 | UDP-GlcNAc:betaGal beta-1,3-N-acetylglucosaminyltransferase 4 [Source:MGI Symbol;Acc:MGI:2680208] |
| 5 | protein_coding | 1.24E+08 | 1.24E+08 | ENSMUSG00000114278 | MGI:6118396 | Gm49027 | predicted gene, 49027 [Source:MGI Symbol;Acc:MGI:6118396] |
| 5 | protein_coding | 1.24E+08 | 1.24E+08 | ENSMUSG00000029434 | MGI:1924823 | Vps33a | VPS33A CORVET/HOPS core subunit [Source:MGI Symbol;Acc:MGI:1924823] |
| 5 | protein_coding | 1.24E+08 | 1.24E+08 | ENSMUSG00000049550 | MGI:1928401 | Clip1 | CAP-GLY domain containing linker protein 1 [Source:MGI Symbol;Acc:MGI:1928401] |
| 5 | protein_coding | 1.24E+08 | 1.24E+08 | ENSMUSG00000029427 | MGI:1917900 | Zcchc8 | zinc finger, CCHC domain containing 8 [Source:MGI Symbol;Acc:MGI:1917900] |
| 5 | protein_coding | 1.24E+08 | 1.24E+08 | ENSMUSG00000029422 | MGI:1913489 | Rsrc2 | arginine/serine-rich coiled-coil 2 [Source:MGI Symbol;Acc:MGI:1913489] |
| 5 | protein_coding | 1.24E+08 | 1.24E+08 | ENSMUSG00000029414 | MGI:2673709 | Kntc1 | kinetochore associated 1 [Source:MGI Symbol;Acc:MGI:2673709] |
| 5 | protein_coding | 1.24E+08 | 1.24E+08 | ENSMUSG00000045502 | MGI:1933383 | Hcar2 | hydroxycarboxylic acid receptor 2 [Source:MGI Symbol;Acc:MGI:1933383] |
| 5 | protein_coding | 1.24E+08 | 1.24E+08 | ENSMUSG00000049241 | MGI:2441671 | Hcar1 | hydrocarboxylic acid receptor 1 [Source:MGI Symbol;Acc:MGI:2441671] |
| 5 | protein_coding | 1.24E+08 | 1.24E+08 | ENSMUSG00000023106 | MGI:1915434 | Denr | density-regulated protein [Source:MGI Symbol;Acc:MGI:1915434] |
| 5 | protein_coding | 1.24E+08 | 1.24E+08 | ENSMUSG00000061882 | MGI:2684996 | Ccdc62 | coiled-coil domain containing 62 [Source:MGI Symbol;Acc:MGI:2684996] |
| 5 | protein_coding | 1.24E+08 | 1.24E+08 | ENSMUSG00000000915 | MGI:1352504 | Hip1r | huntingtin interacting protein 1 related [Source:MGI Symbol;Acc:MGI:1352504] |
| 5 | protein_coding | 1.24E+08 | 1.24E+08 | ENSMUSG00000066278 | MGI:1916724 | Vps37b | vacuolar protein sorting 37B [Source:MGI Symbol;Acc:MGI:1916724] |

| | | | | | | | |
|---|---|---|---|---|---|---|---|
| 5 | protein_coding | 1.24E+08 | 1.24E+08 | ENSMUSG00000029408 | MGI:1861729 | Abcb9 | ATP-binding cassette, sub-family B member 9 [Source:MGI Symbol;Acc:MGI:1861729] |
| 5 | protein_coding | 1.24E+08 | 1.24E+08 | ENSMUSG00000023707 | MGI:1913877 | Ogfod2 | 2-oxoglutarate and iron-dependent oxygenase domain containing 2 [Source:MGI Symbol;Acc:MGI:1913877] |
| 5 | protein_coding | 1.24E+08 | 1.24E+08 | ENSMUSG00000029404 | MGI:1929500 | Arl6ip4 | protein 4 [Source:MGI Symbol;Acc:MGI:1929500] |
| 5 | protein_coding | 1.24E+08 | 1.24E+08 | ENSMUSG00000029406 | MGI:1336192 | Pitpnm2 | phosphatidylinositol transfer protein, membrane-associated 2 [Source:MGI Symbol;Acc:MGI:1336192] |
| 5 | protein_coding | 1.24E+08 | 1.24E+08 | ENSMUSG00000038126 | MGI:2443138 | Mphosph9 | M-phase phosphoprotein 9 [Source:MGI Symbol;Acc:MGI:2443138] |
| 5 | protein_coding | 1.24E+08 | 1.24E+08 | ENSMUSG00000047635 | MGI:1919900 | Mtrfr | rescue [Source:MGI Symbol;Acc:MGI:1919900] |
| 5 | protein_coding | 1.24E+08 | 1.25E+08 | ENSMUSG00000029394 | MGI:1202069 | Cdk2ap1 | cyclin dependent kinase 2 associated protein 1 [Source:MGI Symbol;Acc:MGI:1202069] |
| 5 | protein_coding | 1.25E+08 | 1.25E+08 | ENSMUSG00000038095 | MGI:2384298 | Sbno1 | strawberry notch 1 [Source:MGI Symbol;Acc:MGI:2384298] |
| 5 | protein_coding | 1.25E+08 | 1.25E+08 | ENSMUSG00000049327 | MGI:1915206 | Kmt5a | lysine methyltransferase 5A [Source:MGI Symbol;Acc:MGI:1915206] |
| 5 | protein_coding | 1.25E+08 | 1.25E+08 | ENSMUSG00000029401 | MGI:1933112 | Rilpl2 | Rab interacting lysosomal protein-like 2 [Source:MGI Symbol;Acc:MGI:1933112] |
| 5 | protein_coding | 1.25E+08 | 1.25E+08 | ENSMUSG00000029402 | MGI:1923417 | Snrnp35 | small nuclear ribonucleoprotein 35 (U11/U12) [Source:MGI Symbol;Acc:MGI:1923417] |
| 5 | protein_coding | 1.25E+08 | 1.25E+08 | ENSMUSG00000029392 | MGI:1922945 | Rilpl1 | Rab interacting lysosomal protein-like 1 [Source:MGI Symbol;Acc:MGI:1922945] |
| 5 | protein_coding | 1.25E+08 | 1.25E+08 | ENSMUSG00000029390 | MGI:1929269 | Tmed2 | transmembrane p24 trafficking protein 2 [Source:MGI Symbol;Acc:MGI:1929269] |
| 5 | protein_coding | 1.25E+08 | 1.25E+08 | ENSMUSG00000029389 | MGI:1915098 | Ddx55 | DEAD box helicase 55 [Source:MGI Symbol;Acc:MGI:1915098] |
| 5 | protein_coding | 1.25E+08 | 1.25E+08 | ENSMUSG00000029388 | MGI:2384802 | Eif2b1 | subunit alpha [Source:MGI Symbol;Acc:MGI:2384802] |
| 5 | protein_coding | 1.25E+08 | 1.25E+08 | ENSMUSG00000029387 | MGI:1277143 | Gtf2h3 | general transcription factor IIH, polypeptide 3 [Source:MGI Symbol;Acc:MGI:1277143] |
| 5 | protein_coding | 1.25E+08 | 1.25E+08 | ENSMUSG00000118662 | MGI:1915228 | Tctn2 | tectonic family member 2 [Source:MGI Symbol;Acc:MGI:1915228] |

| | | | | | | | |
|---|---|---|---|---|---|---|---|
| 5 | protein_coding | 1.25E+08 | 1.25E+08 | ENSMUSG00000038023 | MGI:104855 | Atp6v0a2 | [illegible] subunit A2 [Source:MGI Symbol;Acc:MGI:104855] |
| 5 | protein_coding | 1.25E+08 | 1.25E+08 | ENSMUSG00000038011 | MGI:1860299 | Dnah10 | dynein, axonemal, heavy chain 10 [Source:MGI Symbol;Acc:MGI:1860299] |
| 5 | protein_coding | 1.25E+08 | 1.25E+08 | ENSMUSG00000037979 | MGI:106485 | Ccdc92 | coiled-coil domain containing 92 [Source:MGI Symbol;Acc:MGI:106485] |
| 5 | protein_coding | 1.25E+08 | 1.25E+08 | ENSMUSG00000079215 | MGI:2442505 | Zfp664 | zinc finger protein 664 [Source:MGI Symbol;Acc:MGI:2442505] |
| 5 | protein_coding | 1.25E+08 | 1.25E+08 | ENSMUSG00000037962 | MGI:1920371 | Rflna | refilin A [Source:MGI Symbol;Acc:MGI:1920371] |
| 5 | protein_coding | 1.25E+08 | 1.25E+08 | ENSMUSG00000029478 | MGI:1337080 | Ncor2 | nuclear receptor co-repressor 2 [Source:MGI Symbol;Acc:MGI:1337080] |
| 5 | protein_coding | 1.25E+08 | 1.25E+08 | ENSMUSG00000037936 | MGI:893578 | Scarb1 | scavenger receptor class B, member 1 [Source:MGI Symbol;Acc:MGI:893578] |
| 5 | protein_coding | 1.25E+08 | 1.25E+08 | ENSMUSG00000008348 | MGI:98889 | Ubc | ubiquitin C [Source:MGI Symbol;Acc:MGI:98889] |
| 5 | protein_coding | 1.25E+08 | 1.26E+08 | ENSMUSG00000029480 | MGI:3028576 | Dhx37 | DEAH-box helicase 37 [Source:MGI Symbol;Acc:MGI:3028576] |
| 5 | protein_coding | 1.26E+08 | 1.26E+08 | ENSMUSG00000037905 | MGI:1924059 | Bri3bp | Bri3 binding protein [Source:MGI Symbol;Acc:MGI:1924059] |
| 5 | protein_coding | 1.26E+08 | 1.26E+08 | ENSMUSG00000029482 | MGI:1926144 | Aacs | acetoacetyl-CoA synthetase [Source:MGI Symbol;Acc:MGI:1926144] |
| 5 | protein_coding | 1.26E+08 | 1.26E+08 | ENSMUSG00000070498 | MGI:3609245 | Tmem132b | transmembrane protein 132B [Source:MGI Symbol;Acc:MGI:3609245] |
| 5 | rRNA | 53300502 | 53300580 | ENSMUSG00002076332 | MGI:6848481 | Gm56011 | predicted gene, 56011 [Source:MGI Symbol;Acc:MGI:6848481] |
| 5 | rRNA | 53503601 | 53503714 | ENSMUSG00002076314 | MGI:6848483 | Gm56012 | predicted gene, 56012 [Source:MGI Symbol;Acc:MGI:6848483] |
| 5 | rRNA | 53575915 | 53576024 | ENSMUSG00002075450 | MGI:6848485 | Gm56013 | predicted gene, 56013 [Source:MGI Symbol;Acc:MGI:6848485] |
| 5 | rRNA | 65293645 | 65293763 | ENSMUSG00002076210 | MGI:6848657 | Gm56099 | predicted gene, 56099 [Source:MGI Symbol;Acc:MGI:6848657] |
| 5 | rRNA | 76166979 | 76167101 | ENSMUSG00002075417 | MGI:6847279 | Gm55404 | predicted gene, 55404 [Source:MGI Symbol;Acc:MGI:6847279] |
| 5 | rRNA | 91041239 | 91041355 | ENSMUSG00002075252 | MGI:6848414 | Gm55977 | predicted gene, 55977 [Source:MGI Symbol;Acc:MGI:6848414] |

| | | | | | | | |
|---|---|---|---|---|---|---|---|
| 5 | rRNA | 92266036 | 92266142 | ENSMUSG00002075648 | MGI:6848453 | Gm55997 | predicted gene, 55997 [Source:MGI Symbol;Acc:MGI:6848453] |
| 5 | rRNA | 1.08E+08 | 1.08E+08 | ENSMUSG00000064554 | MGI:4422038 | n-R5s173 | nuclear encoded rRNA 5S 173 [Source:MGI Symbol;Acc:MGI:4422038] |
| 5 | rRNA | 1.17E+08 | 1.17E+08 | ENSMUSG00002075044 | MGI:6848505 | Gm56023 | predicted gene, 56023 [Source:MGI Symbol;Acc:MGI:6848505] |
| 5 | rRNA | 1.19E+08 | 1.19E+08 | ENSMUSG00002076829 | MGI:6848615 | Gm56078 | predicted gene, 56078 [Source:MGI Symbol;Acc:MGI:6848615] |
| 5 | rRNA | 1.20E+08 | 1.20E+08 | ENSMUSG00002075687 | MGI:6848651 | Gm56096 | predicted gene, 56096 [Source:MGI Symbol;Acc:MGI:6848651] |
| 5 | scaRNA | 92203923 | 92204051 | ENSMUSG00000088712 | MGI:5452505 | Gm22728 | predicted gene, 22728 [Source:MGI Symbol;Acc:MGI:5452505] |
| 5 | scaRNA | 92577644 | 92577787 | ENSMUSG00002076272 | MGI:6848523 | Gm56032 | predicted gene, 56032 [Source:MGI Symbol;Acc:MGI:6848523] |
| 5 | scaRNA | 1.24E+08 | 1.24E+08 | ENSMUSG00002076894 | MGI:6847105 | Gm55317 | predicted gene, 55317 [Source:MGI Symbol;Acc:MGI:6847105] |
| 5 | snoRNA | 53116693 | 53116804 | ENSMUSG00000077757 | MGI:5453309 | Gm23532 | predicted gene, 23532 [Source:MGI Symbol;Acc:MGI:5453309] |
| 5 | snoRNA | 54984557 | 54984685 | ENSMUSG00000077507 | MGI:5454012 | Gm24235 | predicted gene, 24235 [Source:MGI Symbol;Acc:MGI:5454012] |
| 5 | snoRNA | 56893432 | 56893561 | ENSMUSG00000088734 | MGI:5453673 | Gm23896 | predicted gene, 23896 [Source:MGI Symbol;Acc:MGI:5453673] |
| 5 | snoRNA | 57268166 | 57268288 | ENSMUSG00000088751 | MGI:5454679 | Gm24902 | predicted gene, 24902 [Source:MGI Symbol;Acc:MGI:5454679] |
| 5 | snoRNA | 57285987 | 57286114 | ENSMUSG00000077333 | MGI:5452792 | Gm23015 | predicted gene, 23015 [Source:MGI Symbol;Acc:MGI:5452792] |
| 5 | snoRNA | 58571571 | 58571704 | ENSMUSG00000088796 | MGI:5452313 | Gm22536 | predicted gene, 22536 [Source:MGI Symbol;Acc:MGI:5452313] |
| 5 | snoRNA | 60874660 | 60874782 | ENSMUSG00000077555 | MGI:5453485 | Gm23708 | predicted gene, 23708 [Source:MGI Symbol;Acc:MGI:5453485] |
| 5 | snoRNA | 62289696 | 62289827 | ENSMUSG00000077228 | MGI:5452050 | Gm22273 | predicted gene, 22273 [Source:MGI Symbol;Acc:MGI:5452050] |
| 5 | snoRNA | 62680751 | 62680882 | ENSMUSG00000119858 | MGI:5453856 | Gm24079 | predicted gene, 24079 [Source:MGI Symbol;Acc:MGI:5453856] |
| 5 | snoRNA | 64159696 | 64159827 | ENSMUSG00000077209 | MGI:5455385 | Gm25608 | predicted gene, 25608 [Source:MGI Symbol;Acc:MGI:5455385] |

| | | | | | | | |
|---|---|---|---|---|---|---|---|
| 5 | snoRNA | 64301591 | 64301724 | ENSMUSG00000087954 | MGI:5452336 | Gm22559 | predicted gene, 22559 [Source:MGI Symbol;Acc:MGI:5452336] |
| 5 | snoRNA | 65934357 | 65934550 | ENSMUSG00002076604 | MGI:6848714 | Gm56128 | predicted gene, 56128 [Source:MGI Symbol;Acc:MGI:6848714] |
| 5 | snoRNA | 68723513 | 68723639 | ENSMUSG00000087880 | MGI:5453382 | Gm23605 | predicted gene, 23605 [Source:MGI Symbol;Acc:MGI:5453382] |
| 5 | snoRNA | 74254191 | 74254311 | ENSMUSG00000093355 | MGI:3819495 | Snora26 | small nucleolar RNA, H/ACA box 26 [Source:MGI Symbol;Acc:MGI:3819495] |
| 5 | snoRNA | 77817571 | 77817693 | ENSMUSG00000089415 | MGI:5451788 | Gm22011 | predicted gene, 22011 [Source:MGI Symbol;Acc:MGI:5451788] |
| 5 | snoRNA | 78927706 | 78927841 | ENSMUSG00000077178 | MGI:5455533 | Gm25756 | predicted gene, 25756 [Source:MGI Symbol;Acc:MGI:5455533] |
| 5 | snoRNA | 83796670 | 83796801 | ENSMUSG00000077288 | MGI:5455542 | Gm25765 | predicted gene, 25765 [Source:MGI Symbol;Acc:MGI:5455542] |
| 5 | snoRNA | 83871309 | 83871436 | ENSMUSG00000077474 | MGI:5454569 | Gm24792 | predicted gene, 24792 [Source:MGI Symbol;Acc:MGI:5454569] |
| 5 | snoRNA | 87881387 | 87881509 | ENSMUSG00000119930 | MGI:5454843 | Gm25066 | predicted gene, 25066 [Source:MGI Symbol;Acc:MGI:5454843] |
| 5 | snoRNA | 87912538 | 87912660 | ENSMUSG00000119270 | MGI:5454091 | Gm24314 | predicted gene, 24314 [Source:MGI Symbol;Acc:MGI:5454091] |
| 5 | snoRNA | 89976730 | 89976858 | ENSMUSG00000077175 | MGI:5455535 | Gm25758 | predicted gene, 25758 [Source:MGI Symbol;Acc:MGI:5455535] |
| 5 | snoRNA | 92341181 | 92341324 | ENSMUSG00000084745 | MGI:5455067 | Gm25290 | predicted gene, 25290 [Source:MGI Symbol;Acc:MGI:5455067] |
| 5 | snoRNA | 93221720 | 93221855 | ENSMUSG00000088240 | MGI:5452692 | Gm22915 | predicted gene, 22915 [Source:MGI Symbol;Acc:MGI:5452692] |
| 5 | snoRNA | 98689343 | 98689473 | ENSMUSG00000077424 | MGI:5454056 | Gm24279 | predicted gene, 24279 [Source:MGI Symbol;Acc:MGI:5454056] |
| 5 | snoRNA | 99803560 | 99803691 | ENSMUSG00002076040 | MGI:6848697 | Gm56119 | predicted gene, 56119 [Source:MGI Symbol;Acc:MGI:6848697] |
| 5 | snoRNA | 1.00E+08 | 1.00E+08 | ENSMUSG00002076587 | MGI:6848701 | Gm56121 | predicted gene, 56121 [Source:MGI Symbol;Acc:MGI:6848701] |
| 5 | snoRNA | 1.00E+08 | 1.00E+08 | ENSMUSG00002076905 | MGI:6848730 | Gm56136 | predicted gene, 56136 [Source:MGI Symbol;Acc:MGI:6848730] |
| 5 | snoRNA | 1.01E+08 | 1.01E+08 | ENSMUSG00000077354 | MGI:5456063 | Gm26286 | predicted gene, 26286 [Source:MGI Symbol;Acc:MGI:5456063] |

| | | | | | | | |
|---|---|---|---|---|---|---|---|
| 5 | snoRNA | 1.08E+08 | 1.08E+08 | ENSMUSG00000064453 | MGI:102855 | Snord21 | small nucleolar RNA, C/D box 21 [Source:MGI Symbol;Acc:MGI:102855] |
| 5 | snoRNA | 1.08E+08 | 1.08E+08 | ENSMUSG00000077222 | MGI:5452047 | Gm22270 | predicted gene, 22270 [Source:MGI Symbol;Acc:MGI:5452047] |
| 5 | snoRNA | 1.08E+08 | 1.08E+08 | ENSMUSG00000077426 | MGI:5456164 | Gm26387 | predicted gene, 26387 [Source:MGI Symbol;Acc:MGI:5456164] |
| 5 | snoRNA | 1.10E+08 | 1.10E+08 | ENSMUSG00000087722 | MGI:5456054 | Gm26277 | predicted gene, 26277 [Source:MGI Symbol;Acc:MGI:5456054] |
| 5 | snoRNA | 1.11E+08 | 1.11E+08 | ENSMUSG00000077254 | MGI:5455856 | Gm26079 | predicted gene, 26079 [Source:MGI Symbol;Acc:MGI:5455856] |
| 5 | snoRNA | 1.18E+08 | 1.18E+08 | ENSMUSG00000119733 | MGI:5454853 | Gm25076 | predicted gene, 25076 [Source:MGI Symbol;Acc:MGI:5454853] |
| 5 | snoRNA | 1.21E+08 | 1.21E+08 | ENSMUSG00000064841 | MGI:5455982 | Gm26205 | predicted gene, 26205 [Source:MGI Symbol;Acc:MGI:5455982] |
| 5 | snRNA | 40563093 | 40563272 | ENSMUSG00000089393 | MGI:5452799 | Gm23022 | predicted gene, 23022 [Source:MGI Symbol;Acc:MGI:5452799] |
| 5 | snRNA | 50447722 | 50447828 | ENSMUSG00000119176 | MGI:5452395 | Gm22618 | predicted gene, 22618 [Source:MGI Symbol;Acc:MGI:5452395] |
| 5 | snRNA | 52500460 | 52500564 | ENSMUSG00002075892 | MGI:6848430 | Gm55985 | predicted gene, 55985 [Source:MGI Symbol;Acc:MGI:6848430] |
| 5 | snRNA | 55460762 | 55460923 | ENSMUSG00000065795 | MGI:5455290 | Gm25513 | predicted gene, 25513 [Source:MGI Symbol;Acc:MGI:5455290] |
| 5 | snRNA | 59372289 | 59372395 | ENSMUSG00000119859 | MGI:5452085 | Gm22308 | predicted gene, 22308 [Source:MGI Symbol;Acc:MGI:5452085] |
| 5 | snRNA | 60658176 | 60658283 | ENSMUSG00000088048 | MGI:5453108 | Gm23331 | predicted gene, 23331 [Source:MGI Symbol;Acc:MGI:5453108] |
| 5 | snRNA | 65818149 | 65818222 | ENSMUSG00000088494 | MGI:5454176 | Gm24399 | predicted gene, 24399 [Source:MGI Symbol;Acc:MGI:5454176] |
| 5 | snRNA | 66321899 | 66322005 | ENSMUSG00000093258 | MGI:5452706 | Gm22929 | predicted gene, 22929 [Source:MGI Symbol;Acc:MGI:5452706] |
| 5 | snRNA | 69543409 | 69543515 | ENSMUSG00000119086 | MGI:5453709 | Gm23932 | predicted gene, 23932 [Source:MGI Symbol;Acc:MGI:5453709] |
| 5 | snRNA | 69909746 | 69909871 | ENSMUSG00000080566 | MGI:5455821 | Gm26044 | predicted gene, 26044 [Source:MGI Symbol;Acc:MGI:5455821] |
| 5 | snRNA | 70076438 | 70076524 | ENSMUSG00000084579 | MGI:5453438 | Gm23661 | predicted gene, 23661 [Source:MGI Symbol;Acc:MGI:5453438] |

| | | | | | | | |
|---|---|---|---|---|---|---|---|
| 5 | snRNA | 70344128 | 70344189 | ENSMUSG00000088863 | MGI:5452844 | Gm23067 | predicted gene, 23067 [Source:MGI Symbol;Acc:MGI:5452844] |
| 5 | snRNA | 70708184 | 70708290 | ENSMUSG00000118789 | MGI:5455849 | Gm26072 | predicted gene, 26072 [Source:MGI Symbol;Acc:MGI:5455849] |
| 5 | snRNA | 72462708 | 72462805 | ENSMUSG00000089195 | MGI:5455545 | Gm25768 | predicted gene, 25768 [Source:MGI Symbol;Acc:MGI:5455545] |
| 5 | snRNA | 72576056 | 72576162 | ENSMUSG00000119356 | MGI:5453489 | Gm23712 | predicted gene, 23712 [Source:MGI Symbol;Acc:MGI:5453489] |
| 5 | snRNA | 73128522 | 73128607 | ENSMUSG00002076468 | MGI:6846538 | Gm55032 | predicted gene, 55032 [Source:MGI Symbol;Acc:MGI:6846538] |
| 5 | snRNA | 75532702 | 75532806 | ENSMUSG00002076047 | MGI:6847201 | Gm55365 | predicted gene, 55365 [Source:MGI Symbol;Acc:MGI:6847201] |
| 5 | snRNA | 75721706 | 75721810 | ENSMUSG00000089078 | MGI:5454279 | Gm24502 | predicted gene, 24502 [Source:MGI Symbol;Acc:MGI:5454279] |
| 5 | snRNA | 79898228 | 79898376 | ENSMUSG00000084435 | MGI:5453206 | Gm23429 | predicted gene, 23429 [Source:MGI Symbol;Acc:MGI:5453206] |
| 5 | snRNA | 80246755 | 80246861 | ENSMUSG00000119601 | MGI:5454081 | Gm24304 | predicted gene, 24304 [Source:MGI Symbol;Acc:MGI:5454081] |
| 5 | snRNA | 81041501 | 81041603 | ENSMUSG00000065235 | MGI:5452751 | Gm22974 | predicted gene, 22974 [Source:MGI Symbol;Acc:MGI:5452751] |
| 5 | snRNA | 82951612 | 82951708 | ENSMUSG00000089162 | MGI:5453906 | Gm24129 | predicted gene, 24129 [Source:MGI Symbol;Acc:MGI:5453906] |
| 5 | snRNA | 83837851 | 83837972 | ENSMUSG00000084731 | MGI:5455583 | Gm25806 | predicted gene, 25806 [Source:MGI Symbol;Acc:MGI:5455583] |
| 5 | snRNA | 85977486 | 85977646 | ENSMUSG00000064555 | MGI:5454403 | Gm24626 | predicted gene, 24626 [Source:MGI Symbol;Acc:MGI:5454403] |
| 5 | snRNA | 86227625 | 86227794 | ENSMUSG00000064859 | MGI:5454301 | Gm24524 | predicted gene, 24524 [Source:MGI Symbol;Acc:MGI:5454301] |
| 5 | snRNA | 86384714 | 86384820 | ENSMUSG00000119790 | MGI:5455250 | Gm25473 | predicted gene, 25473 [Source:MGI Symbol;Acc:MGI:5455250] |
| 5 | snRNA | 87064075 | 87064181 | ENSMUSG00000119825 | MGI:5454988 | Gm25211 | predicted gene, 25211 [Source:MGI Symbol;Acc:MGI:5454988] |
| 5 | snRNA | 87076762 | 87076867 | ENSMUSG00000075983 | MGI:5453898 | Gm24121 | predicted gene, 24121 [Source:MGI Symbol;Acc:MGI:5453898] |
| 5 | snRNA | 91365757 | 91365863 | ENSMUSG00000119278 | MGI:5452869 | Gm23092 | predicted gene, 23092 [Source:MGI Symbol;Acc:MGI:5452869] |

| | | | | | | | |
|---|---|---|---|---|---|---|---|
| 5 | snRNA | 92450033 | 92450151 | ENSMUSG00000088623 | MGI:5452808 | Gm23031 | predicted gene, 23031 [Source:MGI Symbol;Acc:MGI:5452808] |
| 5 | snRNA | 93160891 | 93160998 | ENSMUSG00000089018 | MGI:5455298 | Gm25521 | predicted gene, 25521 [Source:MGI Symbol;Acc:MGI:5455298] |
| 5 | snRNA | 96015193 | 96015299 | ENSMUSG00000119041 | MGI:5455714 | Gm25937 | predicted gene, 25937 [Source:MGI Symbol;Acc:MGI:5455714] |
| 5 | snRNA | 97055491 | 97055684 | ENSMUSG00000064550 | MGI:5454400 | Gm24623 | predicted gene, 24623 [Source:MGI Symbol;Acc:MGI:5454400] |
| 5 | snRNA | 1.00E+08 | 1.00E+08 | ENSMUSG00002076497 | MGI:6848703 | Gm56122 | predicted gene, 56122 [Source:MGI Symbol;Acc:MGI:6848703] |
| 5 | snRNA | 1.01E+08 | 1.01E+08 | ENSMUSG00002076815 | MGI:6849022 | Gm56282 | predicted gene, 56282 [Source:MGI Symbol;Acc:MGI:6849022] |
| 5 | snRNA | 1.04E+08 | 1.04E+08 | ENSMUSG00000088121 | MGI:5455498 | Gm25721 | predicted gene, 25721 [Source:MGI Symbol;Acc:MGI:5455498] |
| 5 | snRNA | 1.04E+08 | 1.04E+08 | ENSMUSG00000119515 | MGI:5454674 | Gm24897 | predicted gene, 24897 [Source:MGI Symbol;Acc:MGI:5454674] |
| 5 | snRNA | 1.05E+08 | 1.05E+08 | ENSMUSG00000119653 | MGI:5453874 | Gm24097 | predicted gene, 24097 [Source:MGI Symbol;Acc:MGI:5453874] |
| 5 | snRNA | 1.06E+08 | 1.06E+08 | ENSMUSG00000119467 | MGI:5455449 | Gm25672 | predicted gene, 25672 [Source:MGI Symbol;Acc:MGI:5455449] |
| 5 | snRNA | 1.07E+08 | 1.07E+08 | ENSMUSG00002076219 | MGI:6846582 | Gm55054 | predicted gene, 55054 [Source:MGI Symbol;Acc:MGI:6846582] |
| 5 | snRNA | 1.07E+08 | 1.07E+08 | ENSMUSG00000088429 | MGI:5452334 | Gm22557 | predicted gene, 22557 [Source:MGI Symbol;Acc:MGI:5452334] |
| 5 | snRNA | 1.11E+08 | 1.11E+08 | ENSMUSG00000119880 | MGI:5452493 | Gm22716 | predicted gene, 22716 [Source:MGI Symbol;Acc:MGI:5452493] |
| 5 | snRNA | 1.11E+08 | 1.11E+08 | ENSMUSG00000118827 | MGI:5452360 | Gm22583 | predicted gene, 22583 [Source:MGI Symbol;Acc:MGI:5452360] |
| 5 | snRNA | 1.13E+08 | 1.13E+08 | ENSMUSG00000065373 | MGI:5452517 | Gm22740 | predicted gene, 22740 [Source:MGI Symbol;Acc:MGI:5452517] |
| 5 | snRNA | 1.14E+08 | 1.14E+08 | ENSMUSG00000065320 | MGI:5451833 | Gm22056 | predicted gene, 22056 [Source:MGI Symbol;Acc:MGI:5451833] |
| 5 | snRNA | 1.16E+08 | 1.16E+08 | ENSMUSG00000119520 | MGI:5454042 | Gm24265 | predicted gene, 24265 [Source:MGI Symbol;Acc:MGI:5454042] |
| 5 | snRNA | 1.16E+08 | 1.16E+08 | ENSMUSG00000119132 | MGI:5454184 | Gm24407 | predicted gene, 24407 [Source:MGI Symbol;Acc:MGI:5454184] |

| | | | | | | | |
|---|---|---|---|---|---|---|---|
| 5 | snRNA | 1.18E+08 | 1.18E+08 | ENSMUSG00002075698 | MGI:6848573 | Gm56057 | predicted gene, 56057 [Source:MGI Symbol;Acc:MGI:6848573] |
| 5 | snRNA | 1.18E+08 | 1.18E+08 | ENSMUSG00000088293 | MGI:5456188 | Gm26411 | predicted gene, 26411 [Source:MGI Symbol;Acc:MGI:5456188] |
| 5 | snRNA | 1.21E+08 | 1.21E+08 | ENSMUSG00000077347 | MGI:5454448 | Gm24671 | predicted gene, 24671 [Source:MGI Symbol;Acc:MGI:5454448] |
| 5 | snRNA | 1.23E+08 | 1.23E+08 | ENSMUSG00000088583 | MGI:5452742 | Gm22965 | predicted gene, 22965 [Source:MGI Symbol;Acc:MGI:5452742] |
| 5 | snRNA | 1.23E+08 | 1.23E+08 | ENSMUSG00000065191 | MGI:5453235 | Gm23458 | predicted gene, 23458 [Source:MGI Symbol;Acc:MGI:5453235] |
| 5 | snRNA | 1.24E+08 | 1.24E+08 | ENSMUSG00000084552 | MGI:5454617 | Gm24840 | predicted gene, 24840 [Source:MGI Symbol;Acc:MGI:5454617] |
| 5 | snRNA | 1.25E+08 | 1.25E+08 | ENSMUSG00002074914 | MGI:6847769 | Gm55651 | predicted gene, 55651 [Source:MGI Symbol;Acc:MGI:6847769] |
| 5 | TEC | 41438579 | 41440243 | ENSMUSG00000106865 | MGI:1921963 | 4930519E07Rik | RIKEN cDNA 4930519E07 gene [Source:MGI Symbol;Acc:MGI:1921963] |
| 5 | TEC | 43216598 | 43218226 | ENSMUSG00000105416 | MGI:5662690 | Gm42553 | predicted gene 42553 [Source:MGI Symbol;Acc:MGI:5662690] |
| 5 | TEC | 43301174 | 43305670 | ENSMUSG00000105087 | MGI:5662689 | Gm42552 | predicted gene 42552 [Source:MGI Symbol;Acc:MGI:5662689] |
| 5 | TEC | 43405528 | 43407100 | ENSMUSG00000105374 | MGI:5662688 | Gm42551 | predicted gene 42551 [Source:MGI Symbol;Acc:MGI:5662688] |
| 5 | TEC | 43422500 | 43423711 | ENSMUSG00000105326 | MGI:1924319 | 6030400A10Rik | RIKEN cDNA 6030400A10 gene [Source:MGI Symbol;Acc:MGI:1924319] |
| 5 | TEC | 43685969 | 43690236 | ENSMUSG00000104547 | MGI:5663159 | Gm43022 | predicted gene 43022 [Source:MGI Symbol;Acc:MGI:5663159] |
| 5 | TEC | 43733163 | 43735268 | ENSMUSG00000104912 | MGI:5663160 | Gm43023 | predicted gene 43023 [Source:MGI Symbol;Acc:MGI:5663160] |
| 5 | TEC | 43910251 | 43911613 | ENSMUSG00000104728 | MGI:5662599 | Gm42462 | predicted gene 42462 [Source:MGI Symbol;Acc:MGI:5662599] |
| 5 | TEC | 43939585 | 43942112 | ENSMUSG00000104823 | MGI:5663319 | Gm43182 | predicted gene 43182 [Source:MGI Symbol;Acc:MGI:5663319] |
| 5 | TEC | 43941388 | 43943459 | ENSMUSG00000106609 | MGI:5663318 | Gm43181 | predicted gene 43181 [Source:MGI Symbol;Acc:MGI:5663318] |
| 5 | TEC | 44105458 | 44106004 | ENSMUSG00000106573 | MGI:5663320 | Gm43183 | predicted gene 43183 [Source:MGI Symbol;Acc:MGI:5663320] |

| | | | | | | | |
|---|---|---|---|---|---|---|---|
| 5 | TEC | 44105462 | 44106003 | ENSMUSG00000105831 | MGI:5663321 | Gm43184 | predicted gene 43184 [Source:MGI Symbol;Acc:MGI:5663321] |
| 5 | TEC | 44263191 | 44264881 | ENSMUSG00000105048 | MGI:5663103 | Gm42966 | predicted gene 42966 [Source:MGI Symbol;Acc:MGI:5663103] |
| 5 | TEC | 44291735 | 44292292 | ENSMUSG00000105373 | MGI:5662566 | Gm42429 | predicted gene 42429 [Source:MGI Symbol;Acc:MGI:5662566] |
| 5 | TEC | 44295322 | 44300613 | ENSMUSG00000105263 | MGI:5662564 | Gm42427 | predicted gene 42427 [Source:MGI Symbol;Acc:MGI:5662564] |
| 5 | TEC | 44371143 | 44374601 | ENSMUSG00000105519 | MGI:2444377 | E430021H15Rik | RIKEN cDNA E430021H15 gene [Source:MGI Symbol;Acc:MGI:2444377] |
| 5 | TEC | 44381852 | 44382666 | ENSMUSG00000105149 | MGI:5663120 | Gm42983 | predicted gene 42983 [Source:MGI Symbol;Acc:MGI:5663120] |
| 5 | TEC | 44513047 | 44513389 | ENSMUSG00000104541 | MGI:5663251 | Gm43114 | predicted gene 43114 [Source:MGI Symbol;Acc:MGI:5663251] |
| 5 | TEC | 44649512 | 44652763 | ENSMUSG00000105282 | MGI:5663118 | Gm42981 | predicted gene 42981 [Source:MGI Symbol;Acc:MGI:5663118] |
| 5 | TEC | 44671000 | 44671902 | ENSMUSG00000105136 | MGI:1924798 | 8030487O14Rik | RIKEN cDNA 8030487O14 gene [Source:MGI Symbol;Acc:MGI:1924798] |
| 5 | TEC | 44674303 | 44675055 | ENSMUSG00000106332 | MGI:5663643 | Gm43506 | predicted gene 43506 [Source:MGI Symbol;Acc:MGI:5663643] |
| 5 | TEC | 44742405 | 44742543 | ENSMUSG00000105648 | MGI:5663642 | Gm43505 | predicted gene 43505 [Source:MGI Symbol;Acc:MGI:5663642] |
| 5 | TEC | 44781261 | 44785043 | ENSMUSG00000105861 | MGI:5663645 | Gm43508 | predicted gene 43508 [Source:MGI Symbol;Acc:MGI:5663645] |
| 5 | TEC | 44822036 | 44824073 | ENSMUSG00000104990 | MGI:5663641 | Gm43504 | predicted gene 43504 [Source:MGI Symbol;Acc:MGI:5663641] |
| 5 | TEC | 44837536 | 44840943 | ENSMUSG00000105347 | MGI:5663640 | Gm43503 | predicted gene 43503 [Source:MGI Symbol;Acc:MGI:5663640] |
| 5 | TEC | 44893679 | 44895895 | ENSMUSG00000104851 | MGI:2444788 | E030026E10Rik | RIKEN cDNA E030026E10 gene [Source:MGI Symbol;Acc:MGI:2444788] |
| 5 | TEC | 44908506 | 44911099 | ENSMUSG00000106603 | MGI:5663646 | Gm43509 | predicted gene 43509 [Source:MGI Symbol;Acc:MGI:5663646] |
| 5 | TEC | 45400994 | 45404649 | ENSMUSG00000105247 | MGI:5662656 | Gm42519 | predicted gene 42519 [Source:MGI Symbol;Acc:MGI:5662656] |
| 5 | TEC | 45688414 | 45690453 | ENSMUSG00000105745 | MGI:5662658 | Gm42521 | predicted gene 42521 [Source:MGI Symbol;Acc:MGI:5662658] |

| | | | | | | | |
|---|---|---|---|---|---|---|---|
| 5 | TEC | 45714022 | 45715523 | ENSMUSG00000104837 | MGI:5662657 | Gm42520 | predicted gene 42520 [Source:MGI Symbol;Acc:MGI:5662657] |
| 5 | TEC | 45817409 | 45820063 | ENSMUSG00000104665 | MGI:5663503 | Gm43366 | predicted gene 43366 [Source:MGI Symbol;Acc:MGI:5663503] |
| 5 | TEC | 45826044 | 45827050 | ENSMUSG00000104986 | MGI:1917038 | 1600023N17Rik | RIKEN cDNA 1600023N17 gene [Source:MGI Symbol;Acc:MGI:1917038] |
| 5 | TEC | 45860075 | 45862102 | ENSMUSG00000104342 | MGI:5595560 | Gm36401 | predicted gene, 36401 [Source:MGI Symbol;Acc:MGI:5595560] |
| 5 | TEC | 45915617 | 45919032 | ENSMUSG00000104852 | MGI:5663338 | Gm43201 | predicted gene 43201 [Source:MGI Symbol;Acc:MGI:5663338] |
| 5 | TEC | 45927362 | 45929522 | ENSMUSG00000105457 | MGI:5663337 | Gm43200 | predicted gene 43200 [Source:MGI Symbol;Acc:MGI:5663337] |
| 5 | TEC | 46014984 | 46016106 | ENSMUSG00000105459 | MGI:1921205 | 4930449I04Rik | RIKEN cDNA 4930449I04 gene [Source:MGI Symbol;Acc:MGI:1921205] |
| 5 | TEC | 46659471 | 46660683 | ENSMUSG00000106875 | MGI:5663230 | Gm43093 | predicted gene 43093 [Source:MGI Symbol;Acc:MGI:5663230] |
| 5 | TEC | 46952022 | 46953875 | ENSMUSG00000107113 | MGI:5663229 | Gm43092 | predicted gene 43092 [Source:MGI Symbol;Acc:MGI:5663229] |
| 5 | TEC | 47310993 | 47311937 | ENSMUSG00000104660 | MGI:5663738 | Gm43601 | predicted gene 43601 [Source:MGI Symbol;Acc:MGI:5663738] |
| 5 | TEC | 48215393 | 48217989 | ENSMUSG00000106214 | MGI:5663563 | Gm43426 | predicted gene 43426 [Source:MGI Symbol;Acc:MGI:5663563] |
| 5 | TEC | 48284426 | 48286038 | ENSMUSG00000105502 | MGI:5662673 | Gm42536 | predicted gene 42536 [Source:MGI Symbol;Acc:MGI:5662673] |
| 5 | TEC | 48369014 | 48371587 | ENSMUSG00000105184 | MGI:5662674 | Gm42537 | predicted gene 42537 [Source:MGI Symbol;Acc:MGI:5662674] |
| 5 | TEC | 48437009 | 48438762 | ENSMUSG00000105086 | MGI:5662671 | Gm42534 | predicted gene 42534 [Source:MGI Symbol;Acc:MGI:5662671] |
| 5 | TEC | 48481954 | 48483352 | ENSMUSG00000106604 | MGI:5662672 | Gm42535 | predicted gene 42535 [Source:MGI Symbol;Acc:MGI:5662672] |
| 5 | TEC | 48692686 | 48695283 | ENSMUSG00000105653 | MGI:5663968 | Gm43831 | predicted gene 43831 [Source:MGI Symbol;Acc:MGI:5663968] |
| 5 | TEC | 48782011 | 48784495 | ENSMUSG00000105574 | MGI:5663785 | Gm43648 | predicted gene 43648 [Source:MGI Symbol;Acc:MGI:5663785] |
| 5 | TEC | 48869432 | 48873233 | ENSMUSG00000105530 | MGI:5663966 | Gm43829 | predicted gene 43829 [Source:MGI Symbol;Acc:MGI:5663966] |

| | | | | | | | |
|---|---|---|---|---|---|---|---|
| 5 | TEC | 48948242 | 48949308 | ENSMUSG00000105454 | MGI:5663967 | Gm43830 | predicted gene 43830 [Source:MGI Symbol;Acc:MGI:5663967] |
| 5 | TEC | 48956352 | 48958621 | ENSMUSG00000105958 | MGI:5663456 | Gm43319 | predicted gene 43319 [Source:MGI Symbol;Acc:MGI:5663456] |
| 5 | TEC | 48960070 | 48961353 | ENSMUSG00000106404 | MGI:5663056 | Gm42919 | predicted gene 42919 [Source:MGI Symbol;Acc:MGI:5663056] |
| 5 | TEC | 48976818 | 48979226 | ENSMUSG00000106408 | MGI:5663458 | Gm43321 | predicted gene 43321 [Source:MGI Symbol;Acc:MGI:5663458] |
| 5 | TEC | 49000762 | 49003677 | ENSMUSG00000106414 | MGI:5663457 | Gm43320 | predicted gene 43320 [Source:MGI Symbol;Acc:MGI:5663457] |
| 5 | TEC | 49082296 | 49086082 | ENSMUSG00000103544 | MGI:3644560 | Gm10048 | predicted gene 10048 [Source:MGI Symbol;Acc:MGI:3644560] |
| 5 | TEC | 49105363 | 49107311 | ENSMUSG00000105974 | MGI:5663455 | Gm43318 | predicted gene 43318 [Source:MGI Symbol;Acc:MGI:5663455] |
| 5 | TEC | 49135620 | 49136927 | ENSMUSG00000105791 | MGI:5663478 | Gm43341 | predicted gene 43341 [Source:MGI Symbol;Acc:MGI:5663478] |
| 5 | TEC | 49155054 | 49155881 | ENSMUSG00000106469 | MGI:1924665 | C030015E24Rik | RIKEN cDNA C030015E24 gene [Source:MGI Symbol;Acc:MGI:1924665] |
| 5 | TEC | 49162754 | 49165297 | ENSMUSG00000104655 | MGI:5663454 | Gm43317 | predicted gene 43317 [Source:MGI Symbol;Acc:MGI:5663454] |
| 5 | TEC | 49174855 | 49176149 | ENSMUSG00000105332 | MGI:5663453 | Gm43316 | predicted gene 43316 [Source:MGI Symbol;Acc:MGI:5663453] |
| 5 | TEC | 49178349 | 49179680 | ENSMUSG00000104929 | MGI:5662903 | Gm42766 | predicted gene 42766 [Source:MGI Symbol;Acc:MGI:5662903] |
| 5 | TEC | 49226352 | 49228802 | ENSMUSG00000106244 | MGI:5663183 | Gm43046 | predicted gene 43046 [Source:MGI Symbol;Acc:MGI:5663183] |
| 5 | TEC | 49227345 | 49230804 | ENSMUSG00000105620 | MGI:5663184 | Gm43047 | predicted gene 43047 [Source:MGI Symbol;Acc:MGI:5663184] |
| 5 | TEC | 49231618 | 49232249 | ENSMUSG00000104562 | MGI:5662904 | Gm42767 | predicted gene 42767 [Source:MGI Symbol;Acc:MGI:5662904] |
| 5 | TEC | 49241216 | 49244112 | ENSMUSG00000104661 | MGI:5662905 | Gm42768 | predicted gene 42768 [Source:MGI Symbol;Acc:MGI:5662905] |
| 5 | TEC | 49273101 | 49274759 | ENSMUSG00000105922 | MGI:5662906 | Gm42769 | predicted gene 42769 [Source:MGI Symbol;Acc:MGI:5662906] |
| 5 | TEC | 49281929 | 49286083 | ENSMUSG00000105983 | MGI:5662907 | Gm42770 | predicted gene 42770 [Source:MGI Symbol;Acc:MGI:5662907] |

| | | | | | | | |
|---|---|---|---|---|---|---|---|
| 5 | TEC | 49374643 | 49375030 | ENSMUSG00000105828 | MGI:5663185 | Gm43048 | predicted gene 43048 [Source:MGI Symbol;Acc:MGI:5663185] |
| 5 | TEC | 49377018 | 49379503 | ENSMUSG00000106579 | MGI:5662908 | Gm42771 | predicted gene 42771 [Source:MGI Symbol;Acc:MGI:5662908] |
| 5 | TEC | 49566134 | 49569174 | ENSMUSG00000105700 | MGI:5662909 | Gm42772 | predicted gene 42772 [Source:MGI Symbol;Acc:MGI:5662909] |
| 5 | TEC | 49582359 | 49584125 | ENSMUSG00000106539 | MGI:5663807 | Gm43670 | predicted gene 43670 [Source:MGI Symbol;Acc:MGI:5663807] |
| 5 | TEC | 49673167 | 49676279 | ENSMUSG00000105771 | MGI:1920226 | 2900064K03Rik | RIKEN cDNA 2900064K03 gene [Source:MGI Symbol;Acc:MGI:1920226] |
| 5 | TEC | 50100751 | 50101900 | ENSMUSG00000104634 | MGI:5662598 | Gm42461 | predicted gene 42461 [Source:MGI Symbol;Acc:MGI:5662598] |
| 5 | TEC | 50216438 | 50219004 | ENSMUSG00000106430 | MGI:5662597 | Gm42460 | predicted gene 42460 [Source:MGI Symbol;Acc:MGI:5662597] |
| 5 | TEC | 50995659 | 50998101 | ENSMUSG00000105931 | MGI:5663151 | Gm43014 | predicted gene 43014 [Source:MGI Symbol;Acc:MGI:5663151] |
| 5 | TEC | 51640094 | 51641647 | ENSMUSG00000105442 | MGI:5662751 | Gm42614 | predicted gene 42614 [Source:MGI Symbol;Acc:MGI:5662751] |
| 5 | TEC | 51648439 | 51650258 | ENSMUSG00000105622 | MGI:5662752 | Gm42615 | predicted gene 42615 [Source:MGI Symbol;Acc:MGI:5662752] |
| 5 | TEC | 51656870 | 51658800 | ENSMUSG00000106575 | MGI:5662750 | Gm42613 | predicted gene 42613 [Source:MGI Symbol;Acc:MGI:5662750] |
| 5 | TEC | 51668210 | 51671635 | ENSMUSG00000105762 | MGI:5663742 | Gm43605 | predicted gene 43605 [Source:MGI Symbol;Acc:MGI:5663742] |
| 5 | TEC | 52012562 | 52016117 | ENSMUSG00000105663 | MGI:5663743 | Gm43606 | predicted gene 43606 [Source:MGI Symbol;Acc:MGI:5663743] |
| 5 | TEC | 52032761 | 52036944 | ENSMUSG00000106010 | MGI:5662753 | Gm42616 | predicted gene 42616 [Source:MGI Symbol;Acc:MGI:5662753] |
| 5 | TEC | 52049328 | 52050556 | ENSMUSG00000104919 | MGI:5662754 | Gm42617 | predicted gene 42617 [Source:MGI Symbol;Acc:MGI:5662754] |
| 5 | TEC | 52064464 | 52068273 | ENSMUSG00000105942 | MGI:5663312 | Gm43175 | predicted gene 43175 [Source:MGI Symbol;Acc:MGI:5663312] |
| 5 | TEC | 52237230 | 52239674 | ENSMUSG00000106292 | MGI:5663311 | Gm43174 | predicted gene 43174 [Source:MGI Symbol;Acc:MGI:5663311] |
| 5 | TEC | 52296927 | 52299915 | ENSMUSG00000106648 | MGI:5663313 | Gm43176 | predicted gene 43176 [Source:MGI Symbol;Acc:MGI:5663313] |

| | | | | | | | |
|---|---|---|---|---|---|---|---|
| 5 | TEC | 52331712 | 52333097 | ENSMUSG00000106607 | MGI:5663315 | Gm43178 | predicted gene 43178 [Source:MGI Symbol;Acc:MGI:5663315] |
| 5 | TEC | 52396678 | 52399563 | ENSMUSG00000105328 | MGI:5663317 | Gm43180 | predicted gene 43180 [Source:MGI Symbol;Acc:MGI:5663317] |
| 5 | TEC | 52700248 | 52702686 | ENSMUSG00000104974 | MGI:5663823 | Gm43686 | predicted gene 43686 [Source:MGI Symbol;Acc:MGI:5663823] |
| 5 | TEC | 54062506 | 54066560 | ENSMUSG00000107182 | MGI:5663405 | Gm43268 | predicted gene 43268 [Source:MGI Symbol;Acc:MGI:5663405] |
| 5 | TEC | 54075786 | 54077556 | ENSMUSG00000107173 | MGI:5663403 | Gm43266 | predicted gene 43266 [Source:MGI Symbol;Acc:MGI:5663403] |
| 5 | TEC | 54964676 | 54967356 | ENSMUSG00000106126 | MGI:5663918 | Gm43781 | predicted gene 43781 [Source:MGI Symbol;Acc:MGI:5663918] |
| 5 | TEC | 55660871 | 55664470 | ENSMUSG00000105702 | MGI:5663917 | Gm43780 | predicted gene 43780 [Source:MGI Symbol;Acc:MGI:5663917] |
| 5 | TEC | 55810291 | 55810489 | ENSMUSG00000105186 | MGI:5663915 | Gm43778 | predicted gene 43778 [Source:MGI Symbol;Acc:MGI:5663915] |
| 5 | TEC | 55946513 | 55950229 | ENSMUSG00000105678 | MGI:5663842 | Gm43705 | predicted gene 43705 [Source:MGI Symbol;Acc:MGI:5663842] |
| 5 | TEC | 57608968 | 57611619 | ENSMUSG00000105693 | MGI:5663843 | Gm43706 | predicted gene 43706 [Source:MGI Symbol;Acc:MGI:5663843] |
| 5 | TEC | 57775186 | 57775340 | ENSMUSG00000104924 | MGI:5663139 | Gm43002 | predicted gene 43002 [Source:MGI Symbol;Acc:MGI:5663139] |
| 5 | TEC | 57853339 | 57853719 | ENSMUSG00000106517 | MGI:5662771 | Gm42634 | predicted gene 42634 [Source:MGI Symbol;Acc:MGI:5662771] |
| 5 | TEC | 57881735 | 57884305 | ENSMUSG00000105940 | MGI:5662772 | Gm42635 | predicted gene 42635 [Source:MGI Symbol;Acc:MGI:5662772] |
| 5 | TEC | 57936215 | 57938392 | ENSMUSG00000104821 | MGI:5662618 | Gm42481 | predicted gene 42481 [Source:MGI Symbol;Acc:MGI:5662618] |
| 5 | TEC | 57950501 | 57953095 | ENSMUSG00000105637 | MGI:5662617 | Gm42480 | predicted gene 42480 [Source:MGI Symbol;Acc:MGI:5662617] |
| 5 | TEC | 57955845 | 57957791 | ENSMUSG00000105050 | MGI:5662619 | Gm42482 | predicted gene 42482 [Source:MGI Symbol;Acc:MGI:5662619] |
| 5 | TEC | 58061981 | 58064695 | ENSMUSG00000105079 | MGI:5663852 | Gm43715 | predicted gene 43715 [Source:MGI Symbol;Acc:MGI:5663852] |
| 5 | TEC | 58082962 | 58084317 | ENSMUSG00000105877 | MGI:5662616 | Gm42479 | predicted gene 42479 [Source:MGI Symbol;Acc:MGI:5662616] |

| | | | | | | | |
|---|---|---|---|---|---|---|---|
| 5 | TEC | 58087092 | 58088558 | ENSMUSG00000106337 | MGI:5662615 | Gm42478 | predicted gene 42478 [Source:MGI Symbol;Acc:MGI:5662615] |
| 5 | TEC | 58102384 | 58104254 | ENSMUSG00000104871 | MGI:5662776 | Gm42639 | predicted gene 42639 [Source:MGI Symbol;Acc:MGI:5662776] |
| 5 | TEC | 58118936 | 58120130 | ENSMUSG00000105431 | MGI:5662777 | Gm42640 | predicted gene 42640 [Source:MGI Symbol;Acc:MGI:5662777] |
| 5 | TEC | 58122756 | 58124859 | ENSMUSG00000104709 | MGI:5662621 | Gm42484 | predicted gene 42484 [Source:MGI Symbol;Acc:MGI:5662621] |
| 5 | TEC | 58141554 | 58142794 | ENSMUSG00000103959 | MGI:5610948 | Gm37720 | predicted gene, 37720 [Source:MGI Symbol;Acc:MGI:5610948] |
| 5 | TEC | 58204673 | 58206414 | ENSMUSG00000106646 | MGI:5662620 | Gm42483 | predicted gene 42483 [Source:MGI Symbol;Acc:MGI:5662620] |
| 5 | TEC | 58278781 | 58282266 | ENSMUSG00000105049 | MGI:5662623 | Gm42486 | predicted gene 42486 [Source:MGI Symbol;Acc:MGI:5662623] |
| 5 | TEC | 58330777 | 58332871 | ENSMUSG00000106146 | MGI:5663182 | Gm43045 | predicted gene 43045 [Source:MGI Symbol;Acc:MGI:5663182] |
| 5 | TEC | 59236781 | 59239020 | ENSMUSG00000106467 | MGI:5663532 | Gm43395 | predicted gene 43395 [Source:MGI Symbol;Acc:MGI:5663532] |
| 5 | TEC | 60110854 | 60113483 | ENSMUSG00000104563 | MGI:5663178 | Gm43041 | predicted gene 43041 [Source:MGI Symbol;Acc:MGI:5663178] |
| 5 | TEC | 60154284 | 60156904 | ENSMUSG00000105158 | MGI:5662999 | Gm42862 | predicted gene 42862 [Source:MGI Symbol;Acc:MGI:5662999] |
| 5 | TEC | 60961059 | 60965049 | ENSMUSG00000106044 | MGI:5662997 | Gm42860 | predicted gene 42860 [Source:MGI Symbol;Acc:MGI:5662997] |
| 5 | TEC | 62586062 | 62586420 | ENSMUSG00000105285 | MGI:5663375 | Gm43238 | predicted gene 43238 [Source:MGI Symbol;Acc:MGI:5663375] |
| 5 | TEC | 62769758 | 62770801 | ENSMUSG00000105694 | MGI:5663374 | Gm43237 | predicted gene 43237 [Source:MGI Symbol;Acc:MGI:5663374] |
| 5 | TEC | 62814142 | 62817450 | ENSMUSG00000105707 | MGI:5663890 | Gm43753 | predicted gene 43753 [Source:MGI Symbol;Acc:MGI:5663890] |
| 5 | TEC | 63803513 | 63808237 | ENSMUSG00000104868 | MGI:3642120 | Nwd2os | NACHT and WD repeat domain containing 2, opposite strand [Source:MGI Symbol;Acc:MGI:3642120] |
| 5 | TEC | 64391273 | 64393585 | ENSMUSG00000104947 | MGI:5663858 | Gm43721 | predicted gene 43721 [Source:MGI Symbol;Acc:MGI:5663858] |

| | | | | | | | |
|---|---|---|---|---|---|---|---|
| 5 | TEC | 64625400 | 64626353 | ENSMUSG00000105251 | MGI:1922486 | 4930526M16Rik | RIKEN cDNA 4930526M16 gene [Source:MGI Symbol;Acc:MGI:1922486] |
| 5 | TEC | 64715526 | 64715906 | ENSMUSG00000106566 | MGI:5663973 | Gm43836 | predicted gene 43836 [Source:MGI Symbol;Acc:MGI:5663973] |
| 5 | TEC | 64721114 | 64721262 | ENSMUSG00000104961 | MGI:5663972 | Gm43835 | predicted gene 43835 [Source:MGI Symbol;Acc:MGI:5663972] |
| 5 | TEC | 64721520 | 64722162 | ENSMUSG00000104806 | MGI:5662703 | Gm42566 | predicted gene 42566 [Source:MGI Symbol;Acc:MGI:5662703] |
| 5 | TEC | 64961357 | 64963078 | ENSMUSG00000106205 | MGI:3041194 | C230096K16Rik | RIKEN cDNA C230096K16 gene [Source:MGI Symbol;Acc:MGI:3041194] |
| 5 | TEC | 65129993 | 65132291 | ENSMUSG00000106186 | MGI:5663764 | Gm43627 | predicted gene 43627 [Source:MGI Symbol;Acc:MGI:5663764] |
| 5 | TEC | 65465151 | 65466337 | ENSMUSG00000105935 | MGI:5663765 | Gm43628 | predicted gene 43628 [Source:MGI Symbol;Acc:MGI:5663765] |
| 5 | TEC | 65563094 | 65563860 | ENSMUSG00000106291 | MGI:1915726 | 1110003F10Rik | RIKEN cDNA 1110003F10 gene [Source:MGI Symbol;Acc:MGI:1915726] |
| 5 | TEC | 65567994 | 65570370 | ENSMUSG00000105896 | MGI:5663448 | Gm43311 | predicted gene 43311 [Source:MGI Symbol;Acc:MGI:5663448] |
| 5 | TEC | 65587449 | 65588923 | ENSMUSG00000104917 | MGI:5663426 | Gm43289 | predicted gene 43289 [Source:MGI Symbol;Acc:MGI:5663426] |
| 5 | TEC | 65633848 | 65635404 | ENSMUSG00000104641 | MGI:5663427 | Gm43290 | predicted gene 43290 [Source:MGI Symbol;Acc:MGI:5663427] |
| 5 | TEC | 65650439 | 65651358 | ENSMUSG00000107195 | MGI:1925464 | 4930589O11Rik | RIKEN cDNA 4930589O11 gene [Source:MGI Symbol;Acc:MGI:1925464] |
| 5 | TEC | 65688911 | 65693103 | ENSMUSG00000105601 | MGI:5662862 | Gm42725 | predicted gene 42725 [Source:MGI Symbol;Acc:MGI:5662862] |
| 5 | TEC | 65699272 | 65701023 | ENSMUSG00000107362 | MGI:5623194 | Gm40309 | predicted gene, 40309 [Source:MGI Symbol;Acc:MGI:5623194] |
| 5 | TEC | 65774292 | 65777114 | ENSMUSG00000106767 | MGI:5662864 | Gm42727 | predicted gene 42727 [Source:MGI Symbol;Acc:MGI:5662864] |
| 5 | TEC | 65781277 | 65784206 | ENSMUSG00000106696 | MGI:5662866 | Gm42729 | predicted gene 42729 [Source:MGI Symbol;Acc:MGI:5662866] |
| 5 | TEC | 65829952 | 65831503 | ENSMUSG00000107158 | MGI:5662865 | Gm42728 | predicted gene 42728 [Source:MGI Symbol;Acc:MGI:5662865] |
| 5 | TEC | 65837357 | 65839687 | ENSMUSG00000107050 | MGI:2441875 | C030017G13Rik | RIKEN cDNA C030017G13 gene [Source:MGI Symbol;Acc:MGI:2441875] |

| | | | | | | | |
|---|---|---|---|---|---|---|---|
| 5 | TEC | 65935357 | 65937861 | ENSMUSG00000106682 | MGI:5662785 | Gm42648 | predicted gene 42648 [Source:MGI Symbol;Acc:MGI:5662785] |
| 5 | TEC | 66010675 | 66012301 | ENSMUSG00000106713 | MGI:5662782 | Gm42645 | predicted gene 42645 [Source:MGI Symbol;Acc:MGI:5662782] |
| 5 | TEC | 66014661 | 66015574 | ENSMUSG00000106776 | MGI:5662783 | Gm42646 | predicted gene 42646 [Source:MGI Symbol;Acc:MGI:5662783] |
| 5 | TEC | 66016660 | 66017362 | ENSMUSG00000106681 | MGI:5663141 | Gm43004 | predicted gene 43004 [Source:MGI Symbol;Acc:MGI:5663141] |
| 5 | TEC | 66068967 | 66073289 | ENSMUSG00000107254 | MGI:5663906 | Gm43769 | predicted gene 43769 [Source:MGI Symbol;Acc:MGI:5663906] |
| 5 | TEC | 66108648 | 66109548 | ENSMUSG00000106768 | MGI:1922236 | 4930480C01Rik | RIKEN cDNA 4930480C01 gene [Source:MGI Symbol;Acc:MGI:1922236] |
| 5 | TEC | 66121194 | 66122771 | ENSMUSG00000107049 | MGI:5663908 | Gm43771 | predicted gene 43771 [Source:MGI Symbol;Acc:MGI:5663908] |
| 5 | TEC | 66226398 | 66230238 | ENSMUSG00000107304 | MGI:5663912 | Gm43775 | predicted gene 43775 [Source:MGI Symbol;Acc:MGI:5663912] |
| 5 | TEC | 66237552 | 66241809 | ENSMUSG00000107390 | MGI:5663460 | Gm43323 | predicted gene 43323 [Source:MGI Symbol;Acc:MGI:5663460] |
| 5 | TEC | 66372090 | 66375024 | ENSMUSG00000107125 | MGI:3781994 | Gm3822 | predicted gene 3822 [Source:MGI Symbol;Acc:MGI:3781994] |
| 5 | TEC | 66478348 | 66480360 | ENSMUSG00000106818 | MGI:5663927 | Gm43790 | predicted gene 43790 [Source:MGI Symbol;Acc:MGI:5663927] |
| 5 | TEC | 66503236 | 66504568 | ENSMUSG00000107155 | MGI:5663930 | Gm43793 | predicted gene 43793 [Source:MGI Symbol;Acc:MGI:5663930] |
| 5 | TEC | 66587260 | 66589469 | ENSMUSG00000106732 | MGI:5663929 | Gm43792 | predicted gene 43792 [Source:MGI Symbol;Acc:MGI:5663929] |
| 5 | TEC | 66601075 | 66601766 | ENSMUSG00000106772 | MGI:5663931 | Gm43794 | predicted gene 43794 [Source:MGI Symbol;Acc:MGI:5663931] |
| 5 | TEC | 66681530 | 66683455 | ENSMUSG00000107091 | MGI:5663480 | Gm43343 | predicted gene 43343 [Source:MGI Symbol;Acc:MGI:5663480] |
| 5 | TEC | 66689139 | 66691504 | ENSMUSG00000106720 | MGI:5663932 | Gm43795 | predicted gene 43795 [Source:MGI Symbol;Acc:MGI:5663932] |
| 5 | TEC | 66749805 | 66752913 | ENSMUSG00000106706 | MGI:2444587 | C530043K16Rik | RIKEN cDNA C530043K16 gene [Source:MGI Symbol;Acc:MGI:2444587] |
| 5 | TEC | 66846225 | 66848736 | ENSMUSG00000107300 | MGI:5663416 | Gm43279 | predicted gene 43279 [Source:MGI Symbol;Acc:MGI:5663416] |

| | | | | | | | |
|---|---|---|---|---|---|---|---|
| 5 | TEC | 66875713 | 66877262 | ENSMUSG00000107284 | MGI:5663417 | Gm43280 | predicted gene 43280 [Source:MGI Symbol;Acc:MGI:5663417] |
| 5 | TEC | 67073351 | 67076054 | ENSMUSG00000106749 | MGI:5663418 | Gm43281 | predicted gene 43281 [Source:MGI Symbol;Acc:MGI:5663418] |
| 5 | TEC | 67126772 | 67128928 | ENSMUSG00000107290 | MGI:5663419 | Gm43282 | predicted gene 43282 [Source:MGI Symbol;Acc:MGI:5663419] |
| 5 | TEC | 67195775 | 67196719 | ENSMUSG00000106785 | MGI:5662850 | Gm42713 | predicted gene 42713 [Source:MGI Symbol;Acc:MGI:5662850] |
| 5 | TEC | 67284846 | 67287488 | ENSMUSG00000106995 | MGI:5592326 | Gm33167 | predicted gene, 33167 [Source:MGI Symbol;Acc:MGI:5592326] |
| 5 | TEC | 67585336 | 67587746 | ENSMUSG00000106832 | MGI:5662769 | Gm42632 | predicted gene 42632 [Source:MGI Symbol;Acc:MGI:5662769] |
| 5 | TEC | 67764561 | 67765169 | ENSMUSG00000106947 | MGI:1916650 | 1700025A08Rik | RIKEN cDNA 1700025A08 gene [Source:MGI Symbol;Acc:MGI:1916650] |
| 5 | TEC | 67791006 | 67792651 | ENSMUSG00000107041 | MGI:5662872 | Gm42735 | predicted gene 42735 [Source:MGI Symbol;Acc:MGI:5662872] |
| 5 | TEC | 67914919 | 67916673 | ENSMUSG00000106859 | MGI:5662874 | Gm42737 | predicted gene 42737 [Source:MGI Symbol;Acc:MGI:5662874] |
| 5 | TEC | 67990693 | 67994171 | ENSMUSG00000106837 | MGI:2443017 | D630030B08Rik | RIKEN cDNA D630030B08 gene [Source:MGI Symbol;Acc:MGI:2443017] |
| 5 | TEC | 68000463 | 68001763 | ENSMUSG00000107136 | MGI:5662603 | Gm42466 | predicted gene 42466 [Source:MGI Symbol;Acc:MGI:5662603] |
| 5 | TEC | 68030225 | 68031577 | ENSMUSG00000107206 | MGI:5662873 | Gm42736 | predicted gene 42736 [Source:MGI Symbol;Acc:MGI:5662873] |
| 5 | TEC | 68138014 | 68141704 | ENSMUSG00000106897 | MGI:5663164 | Gm43027 | predicted gene 43027 [Source:MGI Symbol;Acc:MGI:5663164] |
| 5 | TEC | 68239832 | 68241717 | ENSMUSG00000106747 | MGI:5663162 | Gm43025 | predicted gene 43025 [Source:MGI Symbol;Acc:MGI:5663162] |
| 5 | TEC | 69114751 | 69117732 | ENSMUSG00000107117 | MGI:5663979 | Gm43842 | predicted gene 43842 [Source:MGI Symbol;Acc:MGI:5663979] |
| 5 | TEC | 69326015 | 69327591 | ENSMUSG00000107208 | MGI:2140783 | D130004A15Rik | RIKEN cDNA D130004A15 gene [Source:MGI Symbol;Acc:MGI:2140783] |
| 5 | TEC | 69338426 | 69339456 | ENSMUSG00000107234 | MGI:5663508 | Gm43371 | predicted gene 43371 [Source:MGI Symbol;Acc:MGI:5663508] |
| 5 | TEC | 69374764 | 69377633 | ENSMUSG00000107363 | MGI:5663510 | Gm43373 | predicted gene 43373 [Source:MGI Symbol;Acc:MGI:5663510] |

| | | | | | | | |
|---|---|---|---|---|---|---|---|
| 5 | TEC | 69379188 | 69381578 | ENSMUSG00000106673 | MGI:5663511 | Gm43374 | predicted gene 43374 [Source:MGI Symbol;Acc:MGI:5663511] |
| 5 | TEC | 69420928 | 69423056 | ENSMUSG00000106753 | MGI:5663367 | Gm43230 | predicted gene 43230 [Source:MGI Symbol;Acc:MGI:5663367] |
| 5 | TEC | 69433064 | 69434913 | ENSMUSG00000107312 | MGI:5663366 | Gm43229 | predicted gene 43229 [Source:MGI Symbol;Acc:MGI:5663366] |
| 5 | TEC | 69469050 | 69473410 | ENSMUSG00000107148 | MGI:5663025 | Gm42888 | predicted gene 42888 [Source:MGI Symbol;Acc:MGI:5663025] |
| 5 | TEC | 69661185 | 69662303 | ENSMUSG00000106986 | MGI:5663024 | Gm42887 | predicted gene 42887 [Source:MGI Symbol;Acc:MGI:5663024] |
| 5 | TEC | 69713900 | 69714225 | ENSMUSG00000107291 | MGI:1920391 | 3110031N09Rik | RIKEN cDNA 3110031N09 gene [Source:MGI Symbol;Acc:MGI:1920391] |
| 5 | TEC | 70718608 | 70721446 | ENSMUSG00000106549 | MGI:5662790 | Gm42653 | predicted gene 42653 [Source:MGI Symbol;Acc:MGI:5662790] |
| 5 | TEC | 72711583 | 72712572 | ENSMUSG00000105510 | MGI:5663952 | Gm43815 | predicted gene 43815 [Source:MGI Symbol;Acc:MGI:5663952] |
| 5 | TEC | 72967311 | 72969417 | ENSMUSG00000106739 | MGI:5663636 | Gm43499 | predicted gene 43499 [Source:MGI Symbol;Acc:MGI:5663636] |
| 5 | TEC | 73069930 | 73072285 | ENSMUSG00000052154 | MGI:3708720 | Gm9870 | predicted gene 9870 [Source:MGI Symbol;Acc:MGI:3708720] |
| 5 | TEC | 73172542 | 73175028 | ENSMUSG00000107373 | MGI:5662805 | Gm42668 | predicted gene 42668 [Source:MGI Symbol;Acc:MGI:5662805] |
| 5 | TEC | 73199120 | 73202497 | ENSMUSG00000107171 | MGI:5662709 | Gm42572 | predicted gene 42572 [Source:MGI Symbol;Acc:MGI:5662709] |
| 5 | TEC | 73345297 | 73347760 | ENSMUSG00000107109 | MGI:5662708 | Gm42571 | predicted gene 42571 [Source:MGI Symbol;Acc:MGI:5662708] |
| 5 | TEC | 73430215 | 73431092 | ENSMUSG00000106999 | MGI:5662871 | Gm42734 | predicted gene 42734 [Source:MGI Symbol;Acc:MGI:5662871] |
| 5 | TEC | 73465402 | 73465920 | ENSMUSG00000107292 | MGI:5662870 | Gm42733 | predicted gene 42733 [Source:MGI Symbol;Acc:MGI:5662870] |
| 5 | TEC | 73468707 | 73469903 | ENSMUSG00000107331 | MGI:5662869 | Gm42732 | predicted gene 42732 [Source:MGI Symbol;Acc:MGI:5662869] |
| 5 | TEC | 73616672 | 73617065 | ENSMUSG00000106675 | MGI:5663935 | Gm43798 | predicted gene 43798 [Source:MGI Symbol;Acc:MGI:5663935] |
| 5 | TEC | 73804356 | 73804934 | ENSMUSG00000106773 | MGI:5663936 | Gm43799 | predicted gene 43799 [Source:MGI Symbol;Acc:MGI:5663936] |

| | | | | | | | |
|---|---|---|---|---|---|---|---|
| 5 | TEC | 74208960 | 74214166 | ENSMUSG00000106993 | MGI:5663554 | Gm43417 | predicted gene 43417 [Source:MGI Symbol;Acc:MGI:5663554] |
| 5 | TEC | 74220699 | 74221448 | ENSMUSG00000106973 | MGI:5663552 | Gm43415 | predicted gene 43415 [Source:MGI Symbol;Acc:MGI:5663552] |
| 5 | TEC | 74252798 | 74253812 | ENSMUSG00000107255 | MGI:5663553 | Gm43416 | predicted gene 43416 [Source:MGI Symbol;Acc:MGI:5663553] |
| 5 | TEC | 74332822 | 74336711 | ENSMUSG00000106735 | MGI:2442263 | A330058E17Rik | RIKEN cDNA A330058E17 gene [Source:MGI Symbol;Acc:MGI:2442263] |
| 5 | TEC | 74409809 | 74410277 | ENSMUSG00000107209 | MGI:5662736 | Gm42599 | predicted gene 42599 [Source:MGI Symbol;Acc:MGI:5662736] |
| 5 | TEC | 74527824 | 74530576 | ENSMUSG00000107009 | MGI:1925146 | 6720475M21Rik | RIKEN cDNA 6720475M21 gene [Source:MGI Symbol;Acc:MGI:1925146] |
| 5 | TEC | 74590161 | 74591766 | ENSMUSG00000106992 | MGI:5663304 | Gm43167 | predicted gene 43167 [Source:MGI Symbol;Acc:MGI:5663304] |
| 5 | TEC | 74736608 | 74737937 | ENSMUSG00000107165 | MGI:5663884 | Gm43747 | predicted gene 43747 [Source:MGI Symbol;Acc:MGI:5663884] |
| 5 | TEC | 75146517 | 75149727 | ENSMUSG00000107306 | MGI:5662714 | Gm42577 | predicted gene 42577 [Source:MGI Symbol;Acc:MGI:5662714] |
| 5 | TEC | 75180897 | 75183742 | ENSMUSG00000107205 | MGI:5662713 | Gm42576 | predicted gene 42576 [Source:MGI Symbol;Acc:MGI:5662713] |
| 5 | TEC | 75315546 | 75315996 | ENSMUSG00000107245 | MGI:5662940 | Gm42803 | predicted gene 42803 [Source:MGI Symbol;Acc:MGI:5662940] |
| 5 | TEC | 75898542 | 75900865 | ENSMUSG00000107251 | MGI:5663239 | Gm43102 | predicted gene 43102 [Source:MGI Symbol;Acc:MGI:5663239] |
| 5 | TEC | 76006749 | 76008870 | ENSMUSG00000103684 | MGI:5611408 | Gm38180 | predicted gene, 38180 [Source:MGI Symbol;Acc:MGI:5611408] |
| 5 | TEC | 76445447 | 76449622 | ENSMUSG00000106815 | MGI:5663795 | Gm43658 | predicted gene 43658 [Source:MGI Symbol;Acc:MGI:5663795] |
| 5 | TEC | 76803776 | 76806202 | ENSMUSG00000103529 | MGI:2443970 | A730089K16Rik | RIKEN cDNA A730089K16 gene [Source:MGI Symbol;Acc:MGI:2443970] |
| 5 | TEC | 81397096 | 81400537 | ENSMUSG00000107226 | MGI:5663731 | Gm43594 | predicted gene 43594 [Source:MGI Symbol;Acc:MGI:5663731] |
| 5 | TEC | 81683573 | 81684519 | ENSMUSG00000107375 | MGI:5663221 | Gm43084 | predicted gene 43084 [Source:MGI Symbol;Acc:MGI:5663221] |
| 5 | TEC | 81974663 | 81975670 | ENSMUSG00000106746 | MGI:1920293 | 2900064F13Rik | RIKEN cDNA 2900064F13 gene [Source:MGI Symbol;Acc:MGI:1920293] |

| | | | | | | | |
|---|---|---|---|---|---|---|---|
| 5 | TEC | 82267526 | 82269209 | ENSMUSG00000106860 | MGI:1913575 | 1700008H02Rik | RIKEN cDNA 1700008H02 gene [Source:MGI Symbol;Acc:MGI:1913575] |
| 5 | TEC | 87503694 | 87504780 | ENSMUSG00000105192 | MGI:5662933 | Gm42796 | predicted gene 42796 [Source:MGI Symbol;Acc:MGI:5662933] |
| 5 | TEC | 89670719 | 89672859 | ENSMUSG00000104715 | MGI:5663652 | Gm43515 | predicted gene 43515 [Source:MGI Symbol;Acc:MGI:5663652] |
| 5 | TEC | 90545132 | 90547778 | ENSMUSG00000106725 | MGI:5663500 | Gm43363 | predicted gene 43363 [Source:MGI Symbol;Acc:MGI:5663500] |
| 5 | TEC | 90951120 | 90954251 | ENSMUSG00000106957 | MGI:5663222 | Gm43085 | predicted gene 43085 [Source:MGI Symbol;Acc:MGI:5663222] |
| 5 | TEC | 91933920 | 91934741 | ENSMUSG00000106712 | MGI:5011834 | Gm19649 | predicted gene, 19649 [Source:MGI Symbol;Acc:MGI:5011834] |
| 5 | TEC | 92105859 | 92106821 | ENSMUSG00000107200 | MGI:5663176 | Gm43039 | predicted gene 43039 [Source:MGI Symbol;Acc:MGI:5663176] |
| 5 | TEC | 92339219 | 92339776 | ENSMUSG00000107354 | MGI:5663177 | Gm43040 | predicted gene 43040 [Source:MGI Symbol;Acc:MGI:5663177] |
| 5 | TEC | 93238844 | 93241210 | ENSMUSG00000107194 | MGI:5663310 | Gm43173 | predicted gene 43173 [Source:MGI Symbol;Acc:MGI:5663310] |
| 5 | TEC | 93238853 | 93240989 | ENSMUSG00000107374 | MGI:5663309 | Gm43172 | predicted gene 43172 [Source:MGI Symbol;Acc:MGI:5663309] |
| 5 | TEC | 93324829 | 93326918 | ENSMUSG00000107019 | MGI:5663819 | Gm43682 | predicted gene 43682 [Source:MGI Symbol;Acc:MGI:5663819] |
| 5 | TEC | 93331715 | 93333617 | ENSMUSG00000106870 | MGI:5663818 | Gm43681 | predicted gene 43681 [Source:MGI Symbol;Acc:MGI:5663818] |
| 5 | TEC | 96633625 | 96634009 | ENSMUSG00000105446 | MGI:5662742 | Gm42605 | predicted gene 42605 [Source:MGI Symbol;Acc:MGI:5662742] |
| 5 | TEC | 96946066 | 96948211 | ENSMUSG00000105356 | MGI:5662740 | Gm42603 | predicted gene 42603 [Source:MGI Symbol;Acc:MGI:5662740] |
| 5 | TEC | 97027377 | 97030600 | ENSMUSG00000104698 | MGI:5662739 | Gm42602 | predicted gene 42602 [Source:MGI Symbol;Acc:MGI:5662739] |
| 5 | TEC | 97060559 | 97061100 | ENSMUSG00000105871 | MGI:5663284 | Gm43147 | predicted gene 43147 [Source:MGI Symbol;Acc:MGI:5663284] |
| 5 | TEC | 97084672 | 97085191 | ENSMUSG00000105377 | MGI:5663285 | Gm43148 | predicted gene 43148 [Source:MGI Symbol;Acc:MGI:5663285] |
| 5 | TEC | 97088156 | 97088640 | ENSMUSG00000105257 | MGI:5663281 | Gm43144 | predicted gene 43144 [Source:MGI Symbol;Acc:MGI:5663281] |

| | | | | | | | |
|---|---|---|---|---|---|---|---|
| 5 | TEC | 97194040 | 97194667 | ENSMUSG00000104591 | MGI:5663282 | Gm43145 | predicted gene 43145 [Source:MGI Symbol;Acc:MGI:5663282] |
| 5 | TEC | 98147243 | 98148809 | ENSMUSG00000105801 | MGI:3028058 | C430019N01Rik | RIKEN cDNA C430019N01 gene [Source:MGI Symbol;Acc:MGI:3028058] |
| 5 | TEC | 98179426 | 98181864 | ENSMUSG00000106518 | MGI:5662896 | Gm42759 | predicted gene 42759 [Source:MGI Symbol;Acc:MGI:5662896] |
| 5 | TEC | 98494253 | 98497183 | ENSMUSG00000104545 | MGI:2442014 | E030032P16Rik | RIKEN cDNA E030032P16 gene [Source:MGI Symbol;Acc:MGI:2442014] |
| 5 | TEC | 1.01E+08 | 1.01E+08 | ENSMUSG00000106391 | MGI:5662827 | Gm42690 | predicted gene 42690 [Source:MGI Symbol;Acc:MGI:5662827] |
| 5 | TEC | 1.01E+08 | 1.01E+08 | ENSMUSG00000104761 | MGI:5663648 | Gm43511 | predicted gene 43511 [Source:MGI Symbol;Acc:MGI:5663648] |
| 5 | TEC | 1.01E+08 | 1.01E+08 | ENSMUSG00000105607 | MGI:5663650 | Gm43513 | predicted gene 43513 [Source:MGI Symbol;Acc:MGI:5663650] |
| 5 | TEC | 1.01E+08 | 1.01E+08 | ENSMUSG00000106178 | MGI:5663124 | Gm42987 | predicted gene 42987 [Source:MGI Symbol;Acc:MGI:5663124] |
| 5 | TEC | 1.02E+08 | 1.02E+08 | ENSMUSG00000104966 | MGI:5663410 | Gm43273 | predicted gene 43273 [Source:MGI Symbol;Acc:MGI:5663410] |
| 5 | TEC | 1.02E+08 | 1.02E+08 | ENSMUSG00000105632 | MGI:5663409 | Gm43272 | predicted gene 43272 [Source:MGI Symbol;Acc:MGI:5663409] |
| 5 | TEC | 1.02E+08 | 1.02E+08 | ENSMUSG00000106451 | MGI:5663071 | Gm42934 | predicted gene 42934 [Source:MGI Symbol;Acc:MGI:5663071] |
| 5 | TEC | 1.02E+08 | 1.02E+08 | ENSMUSG00000105761 | MGI:5663924 | Gm43787 | predicted gene 43787 [Source:MGI Symbol;Acc:MGI:5663924] |
| 5 | TEC | 1.03E+08 | 1.03E+08 | ENSMUSG00000105947 | MGI:5663926 | Gm43789 | predicted gene 43789 [Source:MGI Symbol;Acc:MGI:5663926] |
| 5 | TEC | 1.03E+08 | 1.03E+08 | ENSMUSG00000104808 | MGI:5662756 | Gm42619 | predicted gene 42619 [Source:MGI Symbol;Acc:MGI:5662756] |
| 5 | TEC | 1.04E+08 | 1.04E+08 | ENSMUSG00000105018 | MGI:5663683 | Gm43546 | predicted gene 43546 [Source:MGI Symbol;Acc:MGI:5663683] |
| 5 | TEC | 1.04E+08 | 1.04E+08 | ENSMUSG00000105019 | MGI:5663684 | Gm43547 | predicted gene 43547 [Source:MGI Symbol;Acc:MGI:5663684] |
| 5 | TEC | 1.04E+08 | 1.04E+08 | ENSMUSG00000106035 | MGI:5663682 | Gm43545 | predicted gene 43545 [Source:MGI Symbol;Acc:MGI:5663682] |
| 5 | TEC | 1.04E+08 | 1.04E+08 | ENSMUSG00000105261 | MGI:5663470 | Gm43333 | predicted gene 43333 [Source:MGI Symbol;Acc:MGI:5663470] |

| | | | | | | | |
|---|---|---|---|---|---|---|---|
| 5 | TEC | 1.05E+08 | 1.05E+08 | ENSMUSG00000105954 | MGI:5662930 | Gm42793 | predicted gene 42793 [Source:MGI Symbol;Acc:MGI:5662930] |
| 5 | TEC | 1.05E+08 | 1.05E+08 | ENSMUSG00000105555 | MGI:5663993 | Gm43856 | predicted gene 43856 [Source:MGI Symbol;Acc:MGI:5663993] |
| 5 | TEC | 1.06E+08 | 1.06E+08 | ENSMUSG00000105826 | MGI:5625026 | Gm42141 | predicted gene, 42141 [Source:MGI Symbol;Acc:MGI:5625026] |
| 5 | TEC | 1.06E+08 | 1.06E+08 | ENSMUSG00000105341 | MGI:5663954 | Gm43817 | predicted gene 43817 [Source:MGI Symbol;Acc:MGI:5663954] |
| 5 | TEC | 1.07E+08 | 1.07E+08 | ENSMUSG00000104706 | MGI:5663558 | Gm43421 | predicted gene 43421 [Source:MGI Symbol;Acc:MGI:5663558] |
| 5 | TEC | 1.07E+08 | 1.07E+08 | ENSMUSG00000106643 | MGI:5663559 | Gm43422 | predicted gene 43422 [Source:MGI Symbol;Acc:MGI:5663559] |
| 5 | TEC | 1.07E+08 | 1.07E+08 | ENSMUSG00000105033 | MGI:5663560 | Gm43423 | predicted gene 43423 [Source:MGI Symbol;Acc:MGI:5663560] |
| 5 | TEC | 1.07E+08 | 1.07E+08 | ENSMUSG00000106671 | MGI:5663037 | Gm42900 | predicted gene 42900 [Source:MGI Symbol;Acc:MGI:5663037] |
| 5 | TEC | 1.08E+08 | 1.08E+08 | ENSMUSG00000105421 | MGI:1923267 | 5830411K02Rik | RIKEN cDNA 5830411K02 gene [Source:MGI Symbol;Acc:MGI:1923267] |
| 5 | TEC | 1.08E+08 | 1.08E+08 | ENSMUSG00000106288 | MGI:5663730 | Gm43593 | predicted gene 43593 [Source:MGI Symbol;Acc:MGI:5663730] |
| 5 | TEC | 1.08E+08 | 1.08E+08 | ENSMUSG00000104748 | MGI:5663729 | Gm43592 | predicted gene 43592 [Source:MGI Symbol;Acc:MGI:5663729] |
| 5 | TEC | 1.08E+08 | 1.08E+08 | ENSMUSG00000106103 | MGI:5663218 | Gm43081 | predicted gene 43081 [Source:MGI Symbol;Acc:MGI:5663218] |
| 5 | TEC | 1.09E+08 | 1.09E+08 | ENSMUSG00000105590 | MGI:5663501 | Gm43364 | predicted gene 43364 [Source:MGI Symbol;Acc:MGI:5663501] |
| 5 | TEC | 1.09E+08 | 1.09E+08 | ENSMUSG00000105084 | MGI:5663502 | Gm43365 | predicted gene 43365 [Source:MGI Symbol;Acc:MGI:5663502] |
| 5 | TEC | 1.09E+08 | 1.09E+08 | ENSMUSG00000103580 | MGI:3642677 | Gm10417 | predicted gene 10417 [Source:MGI Symbol;Acc:MGI:3642677] |
| 5 | TEC | 1.10E+08 | 1.10E+08 | ENSMUSG00000105044 | MGI:3645037 | Gm10416 | predicted pseudogene 10416 [Source:MGI Symbol;Acc:MGI:3645037] |
| 5 | TEC | 1.11E+08 | 1.11E+08 | ENSMUSG00000105112 | MGI:5662915 | Gm42778 | predicted gene 42778 [Source:MGI Symbol;Acc:MGI:5662915] |
| 5 | TEC | 1.11E+08 | 1.11E+08 | ENSMUSG00000105890 | MGI:5662917 | Gm42780 | predicted gene 42780 [Source:MGI Symbol;Acc:MGI:5662917] |

| | | | | | | | |
|---|---|---|---|---|---|---|---|
| 5 | TEC | 1.11E+08 | 1.11E+08 | ENSMUSG00000106153 | MGI:5663814 | Gm43677 | predicted gene 43677 [Source:MGI Symbol;Acc:MGI:5663814] |
| 5 | TEC | 1.11E+08 | 1.11E+08 | ENSMUSG00000105027 | MGI:5663813 | Gm43676 | predicted gene 43676 [Source:MGI Symbol;Acc:MGI:5663813] |
| 5 | TEC | 1.11E+08 | 1.11E+08 | ENSMUSG00000106374 | MGI:5531062 | Gm27680 | predicted gene, 27680 [Source:MGI Symbol;Acc:MGI:5531062] |
| 5 | TEC | 1.12E+08 | 1.12E+08 | ENSMUSG00000106560 | MGI:5663256 | Gm43119 | predicted gene 43119 [Source:MGI Symbol;Acc:MGI:5663256] |
| 5 | TEC | 1.13E+08 | 1.13E+08 | ENSMUSG00000105368 | MGI:5663896 | Gm43759 | predicted gene 43759 [Source:MGI Symbol;Acc:MGI:5663896] |
| 5 | TEC | 1.13E+08 | 1.13E+08 | ENSMUSG00000104885 | MGI:5663895 | Gm43758 | predicted gene 43758 [Source:MGI Symbol;Acc:MGI:5663895] |
| 5 | TEC | 1.13E+08 | 1.13E+08 | ENSMUSG00000105735 | MGI:1918422 | 4933415J04Rik | RIKEN cDNA 4933415J04 gene [Source:MGI Symbol;Acc:MGI:1918422] |
| 5 | TEC | 1.13E+08 | 1.13E+08 | ENSMUSG00000106541 | MGI:5663383 | Gm43246 | predicted gene 43246 [Source:MGI Symbol;Acc:MGI:5663383] |
| 5 | TEC | 1.13E+08 | 1.13E+08 | ENSMUSG00000104576 | MGI:3642056 | F830115B05Rik | RIKEN cDNA F830115B05 gene [Source:MGI Symbol;Acc:MGI:3642056] |
| 5 | TEC | 1.13E+08 | 1.13E+08 | ENSMUSG00000106191 | MGI:1922593 | 4930557B06Rik | RIKEN cDNA 4930557B06 gene [Source:MGI Symbol;Acc:MGI:1922593] |
| 5 | TEC | 1.13E+08 | 1.13E+08 | ENSMUSG00000106206 | MGI:5663231 | Gm43094 | predicted gene 43094 [Source:MGI Symbol;Acc:MGI:5663231] |
| 5 | TEC | 1.14E+08 | 1.14E+08 | ENSMUSG00000106623 | MGI:5663472 | Gm43335 | predicted gene 43335 [Source:MGI Symbol;Acc:MGI:5663472] |
| 5 | TEC | 1.14E+08 | 1.14E+08 | ENSMUSG00000105071 | MGI:5663473 | Gm43336 | predicted gene 43336 [Source:MGI Symbol;Acc:MGI:5663473] |
| 5 | TEC | 1.14E+08 | 1.14E+08 | ENSMUSG00000105514 | MGI:5663050 | Gm42913 | predicted gene 42913 [Source:MGI Symbol;Acc:MGI:5663050] |
| 5 | TEC | 1.15E+08 | 1.15E+08 | ENSMUSG00000107256 | MGI:3652003 | Gm13821 | predicted gene 13821 [Source:MGI Symbol;Acc:MGI:3652003] |
| 5 | TEC | 1.15E+08 | 1.15E+08 | ENSMUSG00000106838 | MGI:1916294 | 1810017P11Rik | RIKEN cDNA 1810017P11 gene [Source:MGI Symbol;Acc:MGI:1916294] |
| 5 | TEC | 1.15E+08 | 1.15E+08 | ENSMUSG00000107054 | MGI:3026961 | C730045M19Rik | RIKEN cDNA C730045M19 gene [Source:MGI Symbol;Acc:MGI:3026961] |
| 5 | TEC | 1.16E+08 | 1.16E+08 | ENSMUSG00000106887 | MGI:3528178 | A930005G22Rik | RIKEN cDNA A930005G22 gene [Source:MGI Symbol;Acc:MGI:3528178] |

| | | | | | | | |
|---|---|---|---|---|---|---|---|
| 5 | TEC | 1.16E+08 | 1.16E+08 | ENSMUSG00000107240 | MGI:5663368 | Gm43231 | predicted gene 43231 [Source:MGI Symbol;Acc:MGI:5663368] |
| 5 | TEC | 1.16E+08 | 1.16E+08 | ENSMUSG00000106869 | MGI:5662610 | Gm42473 | predicted gene 42473 [Source:MGI Symbol;Acc:MGI:5662610] |
| 5 | TEC | 1.16E+08 | 1.16E+08 | ENSMUSG00000107263 | MGI:5662608 | Gm42471 | predicted gene 42471 [Source:MGI Symbol;Acc:MGI:5662608] |
| 5 | TEC | 1.17E+08 | 1.17E+08 | ENSMUSG00000106766 | MGI:1914014 | 4933424N20Rik | RIKEN cDNA 4933424N20 gene [Source:MGI Symbol;Acc:MGI:1914014] |
| 5 | TEC | 1.17E+08 | 1.17E+08 | ENSMUSG00000107021 | MGI:5662990 | Gm42853 | predicted gene 42853 [Source:MGI Symbol;Acc:MGI:5662990] |
| 5 | TEC | 1.17E+08 | 1.17E+08 | ENSMUSG00000107172 | MGI:1918558 | 4933430H06Rik | RIKEN cDNA 4933430H06 gene [Source:MGI Symbol;Acc:MGI:1918558] |
| 5 | TEC | 1.17E+08 | 1.17E+08 | ENSMUSG00000041512 | MGI:1925305 | 4930569F06Rik | RIKEN cDNA 4930569F06 gene [Source:MGI Symbol;Acc:MGI:1925305] |
| 5 | TEC | 1.17E+08 | 1.17E+08 | ENSMUSG00000107225 | MGI:5663774 | Gm43637 | predicted gene 43637 [Source:MGI Symbol;Acc:MGI:5663774] |
| 5 | TEC | 1.17E+08 | 1.17E+08 | ENSMUSG00000106714 | MGI:5662683 | Gm42546 | predicted gene 42546 [Source:MGI Symbol;Acc:MGI:5662683] |
| 5 | TEC | 1.18E+08 | 1.18E+08 | ENSMUSG00000107199 | MGI:5662687 | Gm42550 | predicted gene 42550 [Source:MGI Symbol;Acc:MGI:5662687] |
| 5 | TEC | 1.18E+08 | 1.18E+08 | ENSMUSG00000104071 | MGI:5610256 | Gm37028 | predicted gene, 37028 [Source:MGI Symbol;Acc:MGI:5610256] |
| 5 | TEC | 1.18E+08 | 1.18E+08 | ENSMUSG00000107058 | MGI:5662682 | Gm42545 | predicted gene 42545 [Source:MGI Symbol;Acc:MGI:5662682] |
| 5 | TEC | 1.19E+08 | 1.19E+08 | ENSMUSG00000106912 | MGI:5663922 | Gm43785 | predicted gene 43785 [Source:MGI Symbol;Acc:MGI:5663922] |
| 5 | TEC | 1.19E+08 | 1.19E+08 | ENSMUSG00000107286 | MGI:5663925 | Gm43788 | predicted gene 43788 [Source:MGI Symbol;Acc:MGI:5663925] |
| 5 | TEC | 1.19E+08 | 1.19E+08 | ENSMUSG00000106798 | MGI:5663412 | Gm43275 | predicted gene 43275 [Source:MGI Symbol;Acc:MGI:5663412] |
| 5 | TEC | 1.19E+08 | 1.19E+08 | ENSMUSG00000107116 | MGI:5663411 | Gm43274 | predicted gene 43274 [Source:MGI Symbol;Acc:MGI:5663411] |
| 5 | TEC | 1.19E+08 | 1.19E+08 | ENSMUSG00000107115 | MGI:5663189 | Gm43052 | predicted gene 43052 [Source:MGI Symbol;Acc:MGI:5663189] |
| 5 | TEC | 1.19E+08 | 1.19E+08 | ENSMUSG00000107364 | MGI:5663188 | Gm43051 | predicted gene 43051 [Source:MGI Symbol;Acc:MGI:5663188] |

| | | | | | | | |
|---|---|---|---|---|---|---|---|
| 5 | TEC | 1.19E+08 | 1.19E+08 | ENSMUSG00000107371 | MGI:5663921 | Gm43784 | predicted gene 43784 [Source:MGI Symbol;Acc:MGI:5663921] |
| 5 | TEC | 1.20E+08 | 1.20E+08 | ENSMUSG00000106915 | MGI:5662792 | Gm42655 | predicted gene 42655 [Source:MGI Symbol;Acc:MGI:5662792] |
| 5 | TEC | 1.21E+08 | 1.21E+08 | ENSMUSG00000106825 | MGI:1917539 | 2510016D11Rik | RIKEN cDNA 2510016D11 gene [Source:MGI Symbol;Acc:MGI:1917539] |
| 5 | TEC | 1.21E+08 | 1.21E+08 | ENSMUSG00000107075 | MGI:5663205 | Gm43068 | predicted gene 43068 [Source:MGI Symbol;Acc:MGI:5663205] |
| 5 | TEC | 1.21E+08 | 1.21E+08 | ENSMUSG00000106841 | MGI:5662793 | Gm42656 | predicted gene 42656 [Source:MGI Symbol;Acc:MGI:5662793] |
| 5 | TEC | 1.21E+08 | 1.21E+08 | ENSMUSG00000106940 | MGI:5663067 | Gm42930 | predicted gene 42930 [Source:MGI Symbol;Acc:MGI:5663067] |
| 5 | TEC | 1.22E+08 | 1.22E+08 | ENSMUSG00000106115 | MGI:5663557 | Gm43420 | predicted gene 43420 [Source:MGI Symbol;Acc:MGI:5663557] |
| 5 | TEC | 1.22E+08 | 1.22E+08 | ENSMUSG00000104999 | MGI:1922233 | 4930477O15Rik | RIKEN cDNA 4930477O15 gene [Source:MGI Symbol;Acc:MGI:1922233] |
| 5 | TEC | 1.22E+08 | 1.22E+08 | ENSMUSG00000106590 | MGI:5663016 | Gm42879 | predicted gene 42879 [Source:MGI Symbol;Acc:MGI:5663016] |
| 5 | TEC | 1.22E+08 | 1.22E+08 | ENSMUSG00000106110 | MGI:5663708 | Gm43571 | predicted gene 43571 [Source:MGI Symbol;Acc:MGI:5663708] |
| 5 | TEC | 1.22E+08 | 1.22E+08 | ENSMUSG00000105179 | MGI:5663003 | Gm42866 | predicted gene 42866 [Source:MGI Symbol;Acc:MGI:5663003] |
| 5 | TEC | 1.22E+08 | 1.22E+08 | ENSMUSG00000104719 | MGI:1918553 | 4933437G19Rik | RIKEN cDNA 4933437G19 gene [Source:MGI Symbol;Acc:MGI:1918553] |
| 5 | TEC | 1.22E+08 | 1.22E+08 | ENSMUSG00000105742 | MGI:5662885 | Gm42748 | predicted gene 42748 [Source:MGI Symbol;Acc:MGI:5662885] |
| 5 | TEC | 1.23E+08 | 1.23E+08 | ENSMUSG00000104916 | MGI:5662967 | Gm42830 | predicted gene 42830 [Source:MGI Symbol;Acc:MGI:5662967] |
| 5 | TEC | 1.23E+08 | 1.23E+08 | ENSMUSG00000105434 | MGI:5663496 | Gm43359 | predicted gene 43359 [Source:MGI Symbol;Acc:MGI:5663496] |
| 5 | TEC | 1.23E+08 | 1.23E+08 | ENSMUSG00000105970 | MGI:5663497 | Gm43360 | predicted gene 43360 [Source:MGI Symbol;Acc:MGI:5663497] |
| 5 | TEC | 1.23E+08 | 1.23E+08 | ENSMUSG00000106375 | MGI:5663498 | Gm43361 | predicted gene 43361 [Source:MGI Symbol;Acc:MGI:5663498] |
| 5 | TEC | 1.23E+08 | 1.23E+08 | ENSMUSG00000105815 | MGI:5663678 | Gm43541 | predicted gene 43541 [Source:MGI Symbol;Acc:MGI:5663678] |

| | | | | | | | |
|---|---|---|---|---|---|---|---|
| 5 | TEC | 1.23E+08 | 1.23E+08 | ENSMUSG00000105613 | MGI:5663821 | Gm43684 | predicted gene 43684 [Source:MGI Symbol;Acc:MGI:5663821] |
| 5 | TEC | 1.23E+08 | 1.23E+08 | ENSMUSG00000105979 | MGI:5663548 | Gm43411 | predicted gene 43411 [Source:MGI Symbol;Acc:MGI:5663548] |
| 5 | TEC | 1.23E+08 | 1.23E+08 | ENSMUSG00000105107 | MGI:5663549 | Gm43412 | predicted gene 43412 [Source:MGI Symbol;Acc:MGI:5663549] |
| 5 | TEC | 1.23E+08 | 1.23E+08 | ENSMUSG00000104951 | MGI:5663550 | Gm43413 | predicted gene 43413 [Source:MGI Symbol;Acc:MGI:5663550] |
| 5 | TEC | 1.24E+08 | 1.24E+08 | ENSMUSG00000106254 | MGI:5663044 | Gm42907 | predicted gene 42907 [Source:MGI Symbol;Acc:MGI:5663044] |
| 5 | TEC | 1.24E+08 | 1.24E+08 | ENSMUSG00000104940 | MGI:5663045 | Gm42908 | predicted gene 42908 [Source:MGI Symbol;Acc:MGI:5663045] |
| 5 | TEC | 1.24E+08 | 1.24E+08 | ENSMUSG00000104704 | MGI:5663270 | Gm43133 | predicted gene 43133 [Source:MGI Symbol;Acc:MGI:5663270] |
| 5 | TEC | 1.24E+08 | 1.24E+08 | ENSMUSG00000104877 | MGI:5662788 | Gm42651 | predicted gene 42651 [Source:MGI Symbol;Acc:MGI:5662788] |
| 5 | TEC | 1.24E+08 | 1.24E+08 | ENSMUSG00000104694 | MGI:5663934 | Gm43797 | predicted gene 43797 [Source:MGI Symbol;Acc:MGI:5663934] |
| 5 | TEC | 1.24E+08 | 1.24E+08 | ENSMUSG00000102725 | MGI:5611167 | Gm37939 | predicted gene, 37939 [Source:MGI Symbol;Acc:MGI:5611167] |
| 5 | TEC | 1.24E+08 | 1.24E+08 | ENSMUSG00000105382 | MGI:5663476 | Gm43339 | predicted gene 43339 [Source:MGI Symbol;Acc:MGI:5663476] |
| 5 | TEC | 1.24E+08 | 1.24E+08 | ENSMUSG00000105228 | MGI:5663477 | Gm43340 | predicted gene 43340 [Source:MGI Symbol;Acc:MGI:5663477] |
| 5 | TEC | 1.25E+08 | 1.25E+08 | ENSMUSG00000103864 | MGI:5610643 | Gm37415 | predicted gene, 37415 [Source:MGI Symbol;Acc:MGI:5610643] |
| 5 | TEC | 1.25E+08 | 1.25E+08 | ENSMUSG00000106662 | MGI:5663171 | Gm43034 | predicted gene 43034 [Source:MGI Symbol;Acc:MGI:5663171] |
| 5 | TEC | 1.25E+08 | 1.25E+08 | ENSMUSG00000106458 | MGI:5662770 | Gm42633 | predicted gene 42633 [Source:MGI Symbol;Acc:MGI:5662770] |
| 5 | TEC | 1.26E+08 | 1.26E+08 | ENSMUSG00000104793 | MGI:5663893 | Gm43756 | predicted gene 43756 [Source:MGI Symbol;Acc:MGI:5663893] |
| 5 | transcribed_processed_pseudogene | 44101245 | 44101824 | ENSMUSG00000059565 | MGI:3645663 | Gm5292 | predicted gene 5292 [Source:MGI Symbol;Acc:MGI:3645663] |

| | | | | | | | |
|---|---|---|---|---|---|---|---|
| 5 | transcribed_processed_pseudogene | 45926209 | 45927085 | ENSMUSG00000101587 | MGI:5579742 | Gm29036 | predicted gene 29036 [Source:MGI Symbol;Acc:MGI:5579742] |
| 5 | transcribed_processed_pseudogene | 46571686 | 46571956 | ENSMUSG00000067321 | MGI:3643448 | Gm7931 | predicted pseudogene 7931 [Source:MGI Symbol;Acc:MGI:3643448] |
| 5 | transcribed_processed_pseudogene | 50674815 | 50675556 | ENSMUSG00000106345 | MGI:5662596 | Gm42459 | predicted gene 42459 [Source:MGI Symbol;Acc:MGI:5662596] |
| 5 | transcribed_processed_pseudogene | 50837058 | 50840034 | ENSMUSG00000106038 | MGI:3643111 | Gm4962 | predicted gene 4962 [Source:MGI Symbol;Acc:MGI:3643111] |
| 5 | transcribed_processed_pseudogene | 56571552 | 56647574 | ENSMUSG00000106326 | MGI:5663114 | Gm42977 | predicted gene 42977 [Source:MGI Symbol;Acc:MGI:5663114] |
| 5 | transcribed_processed_pseudogene | 58057076 | 58057649 | ENSMUSG00000104564 | MGI:5010162 | Gm17977 | predicted gene, 17977 [Source:MGI Symbol;Acc:MGI:5010162] |
| 5 | transcribed_processed_pseudogene | 66332308 | 66332686 | ENSMUSG00000107069 | MGI:5663909 | Gm43772 | predicted gene 43772 [Source:MGI Symbol;Acc:MGI:5663909] |
| 5 | transcribed_processed_pseudogene | 67610822 | 67611113 | ENSMUSG00000106925 | MGI:5012257 | Gm20072 | predicted gene, 20072 [Source:MGI Symbol;Acc:MGI:5012257] |
| 5 | transcribed_processed_pseudogene | 72784201 | 72785371 | ENSMUSG00000105124 | MGI:3646414 | Gm5297 | predicted gene 5297 [Source:MGI Symbol;Acc:MGI:3646414] |

| | | | | | | | |
|---|---|---|---|---|---|---|---|
| 5 | transcribed_processed_pseudogene | 73547992 | 73548445 | ENSMUSG00000107272 | MGI:5593742 | Gm34583 | predicted gene, 34583 [Source:MGI Symbol;Acc:MGI:5593742] |
| 5 | transcribed_processed_pseudogene | 73633790 | 73634696 | ENSMUSG00000082039 | MGI:3783097 | Gm15653 | predicted gene 15653 [Source:MGI Symbol;Acc:MGI:3783097] |
| 5 | transcribed_processed_pseudogene | 74477239 | 74477695 | ENSMUSG00000080700 | MGI:3801723 | Gm15981 | predicted gene 15981 [Source:MGI Symbol;Acc:MGI:3801723] |
| 5 | transcribed_processed_pseudogene | 75503104 | 75504493 | ENSMUSG00000106689 | MGI:5010530 | Gm18345 | predicted gene, 18345 [Source:MGI Symbol;Acc:MGI:5010530] |
| 5 | transcribed_processed_pseudogene | 76344472 | 76344941 | ENSMUSG00000083152 | MGI:1930261 | Apc-ps1 | adenomatosis polyposis coli, pseudogene 1 [Source:MGI Symbol;Acc:MGI:1930261] |
| 5 | transcribed_processed_pseudogene | 77659025 | 77659411 | ENSMUSG00000047652 | MGI:3525016 | Pea15b-ps | proliferation and apoptosis adaptor protein 15B, pseudogene [Source:MGI Symbol;Acc:MGI:3525016] |
| 5 | transcribed_processed_pseudogene | 84271172 | 84271450 | ENSMUSG00000079067 | MGI:3783162 | Hmgn2-ps1 | high mobility group nucleosomal binding domain 2, pseudogene 1 [Source:MGI Symbol;Acc:MGI:3783162] |
| 5 | transcribed_processed_pseudogene | 86214616 | 86215615 | ENSMUSG00000106282 | MGI:3780458 | Gm2287 | predicted gene 2287 [Source:MGI Symbol;Acc:MGI:3780458] |
| 5 | transcribed_processed_pseudogene | 87107980 | 87109573 | ENSMUSG00000104798 | MGI:5012306 | Gm20121 | predicted gene, 20121 [Source:MGI Symbol;Acc:MGI:5012306] |

| | | | | | | | |
|---|---|---|---|---|---|---|---|
| 5 | transcribed_processed_pseudogene | 87998377 | 87999501 | ENSMUSG00000067156 | MGI:3646689 | Gm7337 | predicted gene 7337 [Source:MGI Symbol;Acc:MGI:3646689] |
| 5 | transcribed_processed_pseudogene | 88358518 | 88358896 | ENSMUSG00000105576 | MGI:5625049 | Gm42164 | predicted gene, 42164 [Source:MGI Symbol;Acc:MGI:5625049] |
| 5 | transcribed_processed_pseudogene | 91908555 | 91909434 | ENSMUSG00000106917 | MGI:3779767 | Gm7832 | predicted gene 7832 [Source:MGI Symbol;Acc:MGI:3779767] |
| 5 | transcribed_processed_pseudogene | 1.01E+08 | 1.01E+08 | ENSMUSG00000104993 | MGI:5663647 | Gm43510 | predicted gene 43510 [Source:MGI Symbol;Acc:MGI:5663647] |
| 5 | transcribed_processed_pseudogene | 1.08E+08 | 1.08E+08 | ENSMUSG00000084855 | MGI:3642875 | Gm9850 | predicted gene 9850 [Source:MGI Symbol;Acc:MGI:3642875] |
| 5 | transcribed_processed_pseudogene | 1.15E+08 | 1.15E+08 | ENSMUSG00000072692 | MGI:3651519 | Rpl37rt | ribosomal protein L37, retrotransposed [Source:MGI Symbol;Acc:MGI:3651519] |
| 5 | transcribed_processed_pseudogene | 1.16E+08 | 1.16E+08 | ENSMUSG00000081282 | MGI:3650922 | Gm13836 | predicted gene 13836 [Source:MGI Symbol;Acc:MGI:3650922] |
| 5 | transcribed_processed_pseudogene | 1.19E+08 | 1.19E+08 | ENSMUSG00000082365 | MGI:3783197 | Btf3-ps11 | basic transcription factor 3, pseudogene 11 [Source:MGI Symbol;Acc:MGI:3783197] |
| 5 | transcribed_processed_pseudogene | 1.26E+08 | 1.26E+08 | ENSMUSG00000106271 | MGI:5663889 | Gm43752 | predicted gene 43752 [Source:MGI Symbol;Acc:MGI:5663889] |

| | | | | | | | |
|---|---|---|---|---|---|---|---|
| 5 | transcribed_processed_pseudogene | 1.26E+08 | 1.26E+08 | ENSMUSG00000091970 | MGI:3648712 | Gm4868 | predicted gene 4868 [Source:MGI Symbol;Acc:MGI:3648712] |
| 5 | transcribed_unitary_pseudogene | 73170871 | 73174579 | ENSMUSG00000081849 | MGI:3783237 | Zar1-ps | zygote arrest 1, pseudogene [Source:MGI Symbol;Acc:MGI:3783237] |
| 5 | transcribed_unitary_pseudogene | 92111565 | 92117654 | ENSMUSG00000102644 | MGI:1922436 | Thap6 | THAP domain containing 6 [Source:MGI Symbol;Acc:MGI:1922436] |
| 5 | transcribed_unitary_pseudogene | 1.00E+08 | 1.01E+08 | ENSMUSG00000085915 | MGI:3643169 | Gm8091 | predicted gene 8091 [Source:MGI Symbol;Acc:MGI:3643169] |
| 5 | transcribed_unitary_pseudogene | 1.09E+08 | 1.09E+08 | ENSMUSG00000118523 | MGI:5593478 | Gm34319 | predicted gene, 34319 [Source:MGI Symbol;Acc:MGI:5593478] |
| 5 | transcribed_unprocessed_pseudogene | 50374589 | 50548703 | ENSMUSG00000105020 | MGI:3643966 | Gm7205 | predicted gene 7205 [Source:MGI Symbol;Acc:MGI:3643966] |
| 5 | transcribed_unprocessed_pseudogene | 74517244 | 74519080 | ENSMUSG00000084132 | MGI:3801722 | Gm15982 | predicted gene 15982 [Source:MGI Symbol;Acc:MGI:3801722] |
| 5 | transcribed_unprocessed_pseudogene | 77076219 | 77095456 | ENSMUSG00000072852 | MGI:1917534 | 2310040G07Rik | RIKEN cDNA 2310040G07 gene [Source:MGI Symbol;Acc:MGI:1917534] |

| | | | | | | | |
|---|---|---|---|---|---|---|---|
| 5 | transcribed_unprocessed_pseudogene | 87524017 | 87527779 | ENSMUSG00000106397 | MGI:5434404 | Gm21049 | predicted gene, 21049 [Source:MGI Symbol;Acc:MGI:5434404] |
| 5 | transcribed_unprocessed_pseudogene | 88344847 | 88349266 | ENSMUSG00000106065 | MGI:5662758 | Gm42621 | predicted gene 42621 [Source:MGI Symbol;Acc:MGI:5662758] |
| 5 | transcribed_unprocessed_pseudogene | 94450679 | 94453474 | ENSMUSG00000094560 | MGI:2683305 | Pramel39-ps | PRAME like 39, pseudogene [Source:MGI Symbol;Acc:MGI:2683305] |
| 5 | transcribed_unprocessed_pseudogene | 94529232 | 94531954 | ENSMUSG00000073497 | MGI:2140789 | Pramel52-ps | PRAME like 52, pseudogene [Source:MGI Symbol;Acc:MGI:2140789] |
| 5 | transcribed_unprocessed_pseudogene | 95170764 | 95173493 | ENSMUSG00000096742 | MGI:3704106 | Pramel59 | PRAME like 59 [Source:MGI Symbol;Acc:MGI:3704106] |
| 5 | transcribed_unprocessed_pseudogene | 95428321 | 95431051 | ENSMUSG00000107348 | MGI:3646666 | Gm6468 | predicted gene 6468 [Source:MGI Symbol;Acc:MGI:3646666] |
| 5 | transcribed_unprocessed_pseudogene | 95750437 | 95753167 | ENSMUSG00000106690 | MGI:3643967 | Gm6348 | predicted gene 6348 [Source:MGI Symbol;Acc:MGI:3643967] |

| | | | | | | | |
|---|---|---|---|---|---|---|---|
| 5 | transcribed_unprocessed_pseudogene | 1.05E+08 | 1.05E+08 | ENSMUSG00000118625 | MGI:6846487 | Gm55006 | predicted gene, 55006 [Source:MGI Symbol;Acc:MGI:6846487] |
| 5 | transcribed_unprocessed_pseudogene | 1.10E+08 | 1.10E+08 | ENSMUSG00000106028 | MGI:3779800 | Gm8493 | predicted gene 8493 [Source:MGI Symbol;Acc:MGI:3779800] |
| 5 | translated_unprocessed_pseudogene | 1.10E+08 | 1.10E+08 | ENSMUSG00000093663 | MGI:3649028 | Gm7171 | predicted gene 7171 [Source:MGI Symbol;Acc:MGI:3649028] |
| 5 | unitary_pseudogene | 43974000 | 43975191 | ENSMUSG00000144294 | | | family with sequence similarity 200, member B (Fam200B) pseudogene |
| 5 | unitary_pseudogene | 45814290 | 45814914 | ENSMUSG00000105634 | MGI:5663504 | Gm43367 | predicted gene 43367 [Source:MGI Symbol;Acc:MGI:5663504] |
| 5 | unitary_pseudogene | 65054266 | 65056312 | ENSMUSG00000118495 | MGI:5434366 | Gm21011 | predicted gene, 21011 [Source:MGI Symbol;Acc:MGI:5434366] |
| 5 | unitary_pseudogene | 87860712 | 87860762 | ENSMUSG00000144277 | | | Statherin/Histatin pseudogene |
| 5 | unitary_pseudogene | 87901075 | 87901125 | ENSMUSG00000144278 | | | Statherin/Histatin pseudogene |
| 5 | unitary_pseudogene | 1.01E+08 | 1.01E+08 | ENSMUSG00000144266 | | | THAP domain containing 9 (Thap9) pseudogene |
| 5 | unprocessed_pseudogene | 58518976 | 58520854 | ENSMUSG00000105269 | MGI:5663136 | Gm42999 | predicted gene 42999 [Source:MGI Symbol;Acc:MGI:5663136] |

| | | | | | | | |
|---|---|---|---|---|---|---|---|
| 5 | unprocessed_pseudogene | 68726847 | 68752089 | ENSMUSG00000106854 | MGI:5662987 | Gm42850 | predicted gene 42850 [Source:MGI Symbol;Acc:MGI:5662987] |
| 5 | unprocessed_pseudogene | 75302009 | 75302219 | ENSMUSG00000107258 | MGI:3031235 | Olfr1401-ps1 | olfactory receptor 1401, pseudogene 1 [Source:MGI Symbol;Acc:MGI:3031235] |
| 5 | unprocessed_pseudogene | 77835223 | 77837059 | ENSMUSG00000106996 | MGI:5590208 | Gm31049 | predicted gene, 31049 [Source:MGI Symbol;Acc:MGI:5590208] |
| 5 | unprocessed_pseudogene | 84011021 | 84011609 | ENSMUSG00000106576 | MGI:3646530 | Gm4866 | predicted gene 4866 [Source:MGI Symbol;Acc:MGI:3646530] |
| 5 | unprocessed_pseudogene | 87008516 | 87011365 | ENSMUSG00000105309 | MGI:5434816 | Gm21461 | predicted gene, 21461 [Source:MGI Symbol;Acc:MGI:5434816] |
| 5 | unprocessed_pseudogene | 87124679 | 87139636 | ENSMUSG00000106185 | MGI:3644524 | Gm7631 | predicted gene 7631 [Source:MGI Symbol;Acc:MGI:3644524] |
| 5 | unprocessed_pseudogene | 87202462 | 87203524 | ENSMUSG00000106003 | MGI:5662602 | Gm42465 | predicted gene 42465 [Source:MGI Symbol;Acc:MGI:5662602] |
| 5 | unprocessed_pseudogene | 87245041 | 87248254 | ENSMUSG00000105625 | MGI:5590389 | Gm31230 | predicted gene, 31230 [Source:MGI Symbol;Acc:MGI:5590389] |
| 5 | unprocessed_pseudogene | 87310138 | 87323149 | ENSMUSG00000106111 | MGI:3649128 | Gm5717 | predicted gene 5717 [Source:MGI Symbol;Acc:MGI:3649128] |
| 5 | unprocessed_pseudogene | 87354976 | 87365970 | ENSMUSG00000105240 | MGI:5011603 | Gm19418 | predicted gene, 19418 [Source:MGI Symbol;Acc:MGI:5011603] |
| 5 | unprocessed_pseudogene | 87438038 | 87459676 | ENSMUSG00000105897 | MGI:3646880 | Gm7646 | predicted gene 7646 [Source:MGI Symbol;Acc:MGI:3646880] |
| 5 | unprocessed_pseudogene | 87499339 | 87512677 | ENSMUSG00000106303 | MGI:3643731 | Gm7652 | predicted gene 7652 [Source:MGI Symbol;Acc:MGI:3643731] |

| | | | | | | | |
|---|---|---|---|---|---|---|---|
| 5 | unprocessed_pseudogene | 87549178 | 87550837 | ENSMUSG00000106132 | MGI:5663507 | Gm43370 | predicted gene 43370 [Source:MGI Symbol;Acc:MGI:5663507] |
| 5 | unprocessed_pseudogene | 93558621 | 93559719 | ENSMUSG00000107356 | MGI:5010303 | Gm18118 | predicted gene, 18118 [Source:MGI Symbol;Acc:MGI:5010303] |
| 5 | unprocessed_pseudogene | 93569531 | 93571255 | ENSMUSG00000106699 | MGI:5663361 | Gm43224 | predicted gene 43224 [Source:MGI Symbol;Acc:MGI:5663361] |
| 5 | unprocessed_pseudogene | 93842853 | 93845580 | ENSMUSG00000107346 | MGI:5663362 | Gm43225 | predicted gene 43225 [Source:MGI Symbol;Acc:MGI:5663362] |
| 5 | unprocessed_pseudogene | 93885955 | 93888642 | ENSMUSG00000107159 | MGI:3645426 | Gm7919 | predicted gene 7919 [Source:MGI Symbol;Acc:MGI:3645426] |
| 5 | unprocessed_pseudogene | 93918159 | 93919659 | ENSMUSG00000107247 | MGI:5663364 | Gm43227 | predicted gene 43227 [Source:MGI Symbol;Acc:MGI:5663364] |
| 5 | unprocessed_pseudogene | 94084614 | 94087344 | ENSMUSG00000106701 | MGI:3644199 | Gm6346 | predicted gene 6346 [Source:MGI Symbol;Acc:MGI:3644199] |
| 5 | unprocessed_pseudogene | 94144340 | 94147033 | ENSMUSG00000106916 | MGI:3781265 | Gm3089 | predicted gene 3089 [Source:MGI Symbol;Acc:MGI:3781265] |
| 5 | unprocessed_pseudogene | 94582216 | 94584784 | ENSMUSG00000107315 | MGI:3770624 | BC080696 | cDNA sequence BC080696 [Source:MGI Symbol;Acc:MGI:3770624] |
| 5 | unprocessed_pseudogene | 94830504 | 94833233 | ENSMUSG00000092073 | MGI:3779572 | Pramel58 | PRAME like 58 [Source:MGI Symbol;Acc:MGI:3779572] |
| 5 | unprocessed_pseudogene | 94881215 | 94883946 | ENSMUSG00000107064 | MGI:5662698 | Gm42561 | predicted gene 42561 [Source:MGI Symbol;Acc:MGI:5662698] |
| 5 | unprocessed_pseudogene | 94917360 | 94919898 | ENSMUSG00000107187 | MGI:3779755 | Gm7663 | predicted gene 7663 [Source:MGI Symbol;Acc:MGI:3779755] |

| | | | | | | | |
|---|---|---|---|---|---|---|---|
| 5 | unprocessed_pseudogene | 95099396 | 95101993 | ENSMUSG00000107285 | MGI:3781355 | Gm3176 | predicted gene 3176 [Source:MGI Symbol;Acc:MGI:3781355] |
| 5 | unprocessed_pseudogene | 95110877 | 95113553 | ENSMUSG00000073510 | MGI:3779753 | Gm7647 | predicted gene 7647 [Source:MGI Symbol;Acc:MGI:3779753] |
| 5 | unprocessed_pseudogene | 95211435 | 95214166 | ENSMUSG00000107376 | MGI:5662798 | Gm42661 | predicted gene 42661 [Source:MGI Symbol;Acc:MGI:5662798] |
| 5 | unprocessed_pseudogene | 95247883 | 95250141 | ENSMUSG00000106780 | MGI:3647260 | Gm7939 | predicted gene 7939 [Source:MGI Symbol;Acc:MGI:3647260] |
| 5 | unprocessed_pseudogene | 95528913 | 95531643 | ENSMUSG00000107122 | MGI:5591231 | Gm32072 | predicted gene, 32072 [Source:MGI Symbol;Acc:MGI:5591231] |
| 5 | unprocessed_pseudogene | 95564969 | 95567610 | ENSMUSG00000106821 | MGI:3643212 | Gm7961 | predicted gene 7961 [Source:MGI Symbol;Acc:MGI:3643212] |
| 5 | unprocessed_pseudogene | 95573794 | 95576487 | ENSMUSG00000107357 | MGI:5662799 | Gm42662 | predicted gene 42662 [Source:MGI Symbol;Acc:MGI:5662799] |
| 5 | unprocessed_pseudogene | 95810164 | 95812857 | ENSMUSG00000107213 | MGI:3781480 | Gm3302 | predicted gene 3302 [Source:MGI Symbol;Acc:MGI:3781480] |
| 5 | unprocessed_pseudogene | 95982820 | 95985309 | ENSMUSG00000106805 | MGI:5662851 | Gm42714 | predicted gene 42714 [Source:MGI Symbol;Acc:MGI:5662851] |
| 5 | unprocessed_pseudogene | 1.07E+08 | 1.07E+08 | ENSMUSG00000067047 | MGI:3030553 | Olfr719-ps | olfactory receptor 719, pseudogene [Source:MGI Symbol;Acc:MGI:3030553] |
| 5 | unprocessed_pseudogene | 1.09E+08 | 1.09E+08 | ENSMUSG00000093510 | MGI:3761376 | Vmn2r-ps21 | vomeronasal 2, receptor, pseudogene 21 [Source:MGI Symbol;Acc:MGI:3761376] |
| 5 | unprocessed_pseudogene | 1.21E+08 | 1.21E+08 | ENSMUSG00000118628 | MGI:3645862 | Gm7573 | predicted gene 7573 [Source:MGI Symbol;Acc:MGI:3645862] |
